## Supplemental Table 4 for "Inhibition of *O*-GlcNAc transferase activates type I interferon-dependent antitumor immunity by bridging cGAS-STING pathway"

| | | | | | | | | | | KO+GFP_#1 | KO+GFP_#2 | KO+GFP_#3 | KO+WT_#4 | KO+WT_#5 | KO+WT_#6 | KO+GFP_#1 | KO+GFP_#2 | KO+GFP_#3 | KO+WT_#4 | KO+WT_#5 | KO+WT_#6 |
|---|---|---|---|---|---|---|---|---|---|---|---|---|---|---|---|---|---|---|---|---|---|
| Protein FDR | Accession | Description | # PSMs | # Peptides | # Unique Peptide | MW [kDa] | Coverage | # AAs | calc. pI | # PSMs: A2 | # PSMs: B2 | # PSMs: C2 | # PSMs: D2 | # PSMs: E2 | # PSMs: F2 | # Peptides: A2 | # Peptides: B2 | # Peptides: C2 | # Peptides: D2 | # Peptides: E2 | # Peptides: F2 |
| High | P35579 | Myosin-9 OS=Homo sapiens OX=9606 GN=MYH9 PE=1 SV=4 | 6611 | 165 | 135 | 226.4 | 67 | 1960 | 5.6 | 872 | 926 | 1036 | 224 | 345 | 233 | 145 | 143 | 140 | 93 | 113 | 96 |
| High | Q9U6Y5 | SWISS-PROT:Q9U6Y5 Green fluorescent protein (GFP-Cter-HisTag) | 4079 | 24 | 24 | 28.1 | 69 | 249 | 6.52 | 951 | 1033 | 927 | 200 | 186 | 189 | 23 | 23 | 20 | 10 | 9 | 10 |
| High | Q7Z406 | Myosin-14 OS=Homo sapiens OX=9606 GN=MYH14 PE=1 SV=2 | 3222 | 140 | 126 | 227.7 | 65 | 1995 | 5.6 | 482 | 489 | 515 | 118 | 168 | 128 | 124 | 123 | 122 | 75 | 84 | 77 |
| High | O15294 | UDP-N-acetylglucosamine--peptide N-acetylglucosaminyltransferase 110 kDa subunit OS=Homo sapiens OX=9606 GN=OGT PE=1 S | 2542 | 54 | 54 | 116.9 | 61 | 1046 | 6.7 | | | 2 | 293 | 483 | 327 | | | 2 | 37 | 39 | 38 |
| High | P49792 | E3 SUMO-protein ligase RanBP2 OS=Homo sapiens OX=9606 GN=RANBP2 PE=1 SV=2 | 2241 | 182 | 127 | 358 | 67 | 3224 | 6.2 | 8 | 11 | 13 | 40 | 44 | 37 | 8 | 11 | 13 | 39 | 42 | 35 |
| High | P60709 | Actin, cytoplasmic 1 OS=Homo sapiens OX=9606 GN=ACTB PE=1 SV=1 | 2101 | 25 | 10 | 41.7 | 74 | 375 | 5.48 | 246 | 268 | 268 | 120 | 147 | 118 | 21 | 21 | 21 | 16 | 15 | 16 |
| High | P05787 | SWISS-PROT:P05787 Tax_Id=9606 Gene_Symbol=KRT8 Keratin, type II cytoskeletal 8 | 1908 | 49 | 19 | 53.7 | 83 | 483 | 5.59 | 383 | 423 | 460 | 85 | 106 | 84 | 48 | 46 | 46 | 30 | 31 | 31 |
| High | Q15149 | Plectin OS=Homo sapiens OX=9606 GN=PLEC PE=1 SV=3 | 1463 | 228 | 220 | 531.5 | 53 | 4684 | 5.96 | 212 | 220 | 226 | 99 | 113 | 101 | 178 | 182 | 182 | 94 | 104 | 97 |
| High | P68133 | Actin, alpha skeletal muscle OS=Homo sapiens OX=9606 GN=ACTA1 PE=1 SV=1 | 1270 | 19 | 6 | 42 | 54 | 377 | 5.39 | 122 | 124 | 161 | 98 | 91 | 96 | 13 | 15 | 14 | 12 | 10 | 14 |
| High | P51610 | Host cell factor 1 OS=Homo sapiens OX=9606 GN=HCFC1 PE=1 SV=2 | 1227 | 61 | 61 | 208.6 | 44 | 2035 | 7.46 | | 1 | | 161 | 193 | 173 | | 1 | | 49 | 54 | 47 |
| High | P35580 | Myosin-10 OS=Homo sapiens OX=9606 GN=MYH10 PE=1 SV=3 | 1221 | 109 | 82 | 228.9 | 56 | 1976 | 5.54 | 162 | 178 | 176 | 48 | 73 | 53 | 90 | 89 | 90 | 36 | 44 | 37 |
| High | Q13813 | Spectrin alpha chain, non-erythrocytic 1 OS=Homo sapiens OX=9606 GN=SPTAN1 PE=1 SV=3 | 1055 | 141 | 141 | 284.4 | 68 | 2472 | 5.35 | 165 | 176 | 183 | 37 | 46 | 38 | 113 | 119 | 121 | 37 | 45 | 38 |
| High | P78527 | DNA-dependent protein kinase catalytic subunit OS=Homo sapiens OX=9606 GN=PRKDC PE=1 SV=3 | 1009 | 159 | 159 | 468.8 | 42 | 4128 | 7.12 | 143 | 144 | 127 | 67 | 73 | 65 | 127 | 126 | 111 | 63 | 71 | 63 |
| High | Q01082 | Spectrin beta chain, non-erythrocytic 1 OS=Homo sapiens OX=9606 GN=SPTBN1 PE=1 SV=2 | 907 | 127 | 116 | 274.4 | 63 | 2364 | 5.57 | 147 | 151 | 139 | 27 | 33 | 27 | 101 | 107 | 100 | 26 | 33 | 27 |
| High | Q09666 | Neuroblast differentiation-associated protein AHNAK OS=Homo sapiens OX=9606 GN=AHNAK PE=1 SV=2 | 903 | 172 | 172 | 628.7 | 59 | 5890 | 6.15 | 60 | 61 | 90 | 75 | 93 | 76 | 55 | 53 | 77 | 68 | 77 | 68 |
| High | P08727 | Keratin, type I cytoskeletal 19 OS=Homo sapiens OX=9606 GN=KRT19 PE=1 SV=4 | 859 | 40 | 25 | 44.1 | 79 | 400 | 5.14 | 204 | 195 | 217 | 35 | 46 | 37 | 38 | 38 | 38 | 22 | 22 | 22 |
| High | P05783 | Keratin, type I cytoskeletal 18 OS=Homo sapiens OX=9606 GN=KRT18 PE=1 SV=2 | 803 | 30 | 20 | 48 | 61 | 430 | 5.45 | 161 | 160 | 164 | 44 | 52 | 42 | 29 | 26 | 27 | 19 | 19 | 18 |
| High | P60660 | Myosin light polypeptide 6 OS=Homo sapiens OX=9606 GN=MYL6 PE=1 SV=2 | 782 | 13 | 9 | 16.9 | 85 | 151 | 4.65 | 106 | 121 | 122 | 44 | 49 | 47 | 11 | 12 | 13 | 8 | 8 | 9 |
| High | H-INV:HIT | Tax_Id=9606 Gene_Symbol=- Similar to Keratin, type II cytoskeletal 8 | 771 | 24 | 1 | 49.4 | 44 | 443 | 5.2 | 163 | 167 | 176 | 32 | 44 | 34 | 24 | 24 | 22 | 14 | 15 | 15 |
| High | A6NHR9 | Structural maintenance of chromosomes flexible hinge domain-containing protein 1 OS=Homo sapiens OX=9606 GN=SMCHD1 PE= | 769 | 93 | 93 | 226.2 | 49 | 2005 | 7.3 | 5 | 4 | 5 | 91 | 113 | 92 | 5 | 4 | 5 | 62 | 69 | 59 |
| High | Q8WWI1 | LIM domain only protein 7 OS=Homo sapiens OX=9606 GN=LMO7 PE=1 SV=3 | 745 | 73 | 73 | 192.6 | 42 | 1683 | 8.09 | 114 | 107 | 108 | 32 | 37 | 33 | 63 | 63 | 56 | 29 | 30 | 30 |
| High | P35749 | Myosin-11 OS=Homo sapiens OX=9606 GN=MYH11 PE=1 SV=3 | 716 | 26 | 1 | 227.2 | 12 | 1972 | 5.5 | 95 | 94 | 105 | 27 | 52 | 32 | 24 | 23 | 20 | 15 | 20 | 17 |
| High | P07437 | Tubulin beta chain OS=Homo sapiens OX=9606 GN=TUBB PE=1 SV=2 | 676 | 25 | 4 | 49.6 | 73 | 444 | 4.89 | 98 | 97 | 88 | 44 | 54 | 46 | 23 | 25 | 24 | 14 | 14 | 13 |
| High | P02545 | Prelamin-A/C OS=Homo sapiens OX=9606 GN=LMNA PE=1 SV=1 | 675 | 46 | 45 | 74.1 | 69 | 664 | 7.02 | 132 | 130 | 144 | 37 | 43 | 40 | 43 | 44 | 45 | 28 | 31 | 34 |
| High | P68371 | Tubulin beta-4B chain OS=Homo sapiens OX=9606 GN=TUBB4B PE=1 SV=1 | 674 | 25 | 1 | 49.8 | 73 | 445 | 4.89 | 101 | 98 | 88 | 43 | 52 | 43 | 23 | 25 | 24 | 14 | 14 | 13 |
| High | Q562R1 | Beta-actin-like protein 2 OS=Homo sapiens OX=9606 GN=ACTBL2 PE=1 SV=2 | 656 | 12 | 5 | 42 | 36 | 376 | 5.59 | 50 | 52 | 83 | 64 | 50 | 62 | 8 | 6 | 8 | 4 | 5 | 5 |
| High | P52732 | Kinesin-like protein KIF11 OS=Homo sapiens OX=9606 GN=KIF11 PE=1 SV=2 | 638 | 60 | 60 | 119.1 | 53 | 1056 | 5.64 | 69 | 61 | 64 | 70 | 91 | 76 | 49 | 43 | 44 | 44 | 46 | 45 |
| High | Q00610 | Clathrin heavy chain 1 OS=Homo sapiens OX=9606 GN=CLTC PE=1 SV=5 | 631 | 72 | 72 | 191.5 | 52 | 1675 | 5.69 | 81 | 85 | 72 | 36 | 38 | 33 | 66 | 67 | 52 | 31 | 34 | 29 |
| High | P04350 | Tubulin beta-4A chain OS=Homo sapiens OX=9606 GN=TUBB4A PE=1 SV=2 | 607 | 21 | 1 | 49.6 | 62 | 444 | 4.88 | 91 | 88 | 80 | 39 | 44 | 38 | 19 | 20 | 20 | 11 | 10 | 11 |
| High | P49327 | Fatty acid synthase OS=Homo sapiens OX=9606 GN=FASN PE=1 SV=3 | 603 | 85 | 84 | 273.3 | 45 | 2511 | 6.44 | 71 | 78 | 71 | 39 | 48 | 48 | 61 | 65 | 61 | 37 | 44 | 44 |
| High | Q9BTC0 | Death-inducer obliterator 1 OS=Homo sapiens OX=9606 GN=DIDO1 PE=1 SV=5 | 600 | 75 | 74 | 243.7 | 48 | 2240 | 7.88 | | 1 | | 77 | 99 | 85 | | 1 | | 48 | 57 | 56 |
| High | O75369 | Filamin-B OS=Homo sapiens OX=9606 GN=FLNB PE=1 SV=2 | 591 | 100 | 94 | 278 | 53 | 2602 | 5.73 | 62 | 73 | 74 | 47 | 49 | 49 | 58 | 66 | 69 | 43 | 46 | 48 |
| High | Q14974 | Importin subunit beta-1 OS=Homo sapiens OX=9606 GN=KPNB1 PE=1 SV=2 | 584 | 31 | 31 | 97.1 | 44 | 876 | 4.78 | 13 | 14 | 14 | 8 | 11 | 9 | 12 | 13 | 12 | 8 | 11 | 9 |
| High | P11142 | Heat shock cognate 71 kDa protein OS=Homo sapiens OX=9606 GN=HSPA8 PE=1 SV=1 | 571 | 35 | 23 | 70.9 | 69 | 646 | 5.52 | 50 | 47 | 55 | 63 | 77 | 58 | 29 | 26 | 30 | 22 | 27 | 22 |
| High | P15924 | Desmoplakin OS=Homo sapiens OX=9606 GN=DSP PE=1 SV=3 | 543 | 103 | 103 | 331.6 | 40 | 2871 | 6.81 | 76 | 75 | 71 | 45 | 46 | 43 | 70 | 69 | 64 | 44 | 45 | 42 |
| High | Q92614 | Unconventional myosin-XVIIIa OS=Homo sapiens OX=9606 GN=MYO18A PE=1 SV=3 | 537 | 87 | 87 | 233 | 49 | 2054 | 6.3 | 72 | 79 | 73 | 22 | 23 | 22 | 63 | 69 | 62 | 20 | 22 | 20 |
| High | O14950 | Myosin regulatory light chain 12B OS=Homo sapiens OX=9606 GN=MYL12B PE=1 SV=2 | 532 | 12 | 12 | 19.8 | 63 | 172 | 4.84 | 76 | 74 | 74 | 15 | 25 | 16 | 12 | 11 | 11 | 7 | 8 | 7 |
| High | Q13885 | Tubulin beta-2A chain OS=Homo sapiens OX=9606 GN=TUBB2A PE=1 SV=1 | 524 | 22 | 4 | 49.9 | 67 | 445 | 4.89 | 76 | 74 | 71 | 36 | 46 | 38 | 18 | 18 | 18 | 10 | 11 | 9 |
| High | O14715 | RANBP2-like and GRIP domain-containing protein 8 OS=Homo sapiens OX=9606 GN=RGPD8 PE=1 SV=2 | 494 | 66 | 2 | 198.9 | 41 | 1765 | 6.49 | 2 | 2 | 4 | 9 | 12 | 8 | 2 | 2 | 4 | 9 | 11 | 8 |
| High | P38646 | Stress-70 protein, mitochondrial OS=Homo sapiens OX=9606 GN=HSPA9 PE=1 SV=2 | 490 | 35 | 35 | 73.6 | 58 | 679 | 6.16 | 48 | 53 | 56 | 39 | 54 | 41 | 27 | 28 | 31 | 24 | 26 | 25 |
| High | Q99666 | RANBP2-like and GRIP domain-containing protein 5/6 OS=Homo sapiens OX=9606 GN=RGPD5 PE=1 SV=3 | 489 | 64 | 2 | 198.8 | 38 | 1765 | 6.42 | 2 | 2 | 4 | 9 | 12 | 8 | 2 | 2 | 4 | 9 | 11 | 8 |
| High | P21333 | Filamin-A OS=Homo sapiens OX=9606 GN=FLNA PE=1 SV=4 | 474 | 75 | 69 | 280.6 | 45 | 2647 | 6.06 | 58 | 46 | 44 | 48 | 49 | 40 | 39 | 35 | 41 | 37 | 39 | 31 |
| High | P0DJD0 | RANBP2-like and GRIP domain-containing protein 1 OS=Homo sapiens OX=9606 GN=RGPD1 PE=2 SV=1 | 470 | 45 | 2 | 196.5 | 30 | 1748 | 6.14 | | | 3 | 8 | 11 | 9 | | | 3 | 8 | 10 | 9 |
| High | P07355 | Annexin A2 OS=Homo sapiens OX=9606 GN=ANXA2 PE=1 SV=2 | 465 | 27 | 27 | 38.6 | 67 | 339 | 7.75 | 44 | 42 | 49 | 40 | 45 | 40 | 24 | 22 | 22 | 18 | 18 | 20 |
| High | Q06830 | Peroxiredoxin-1 OS=Homo sapiens OX=9606 GN=PRDX1 PE=1 SV=1 | 450 | 17 | 14 | 22.1 | 74 | 199 | 8.13 | 42 | 47 | 46 | 46 | 53 | 49 | 16 | 16 | 15 | 13 | 14 | 14 |
| High | P98088 | Mucin-5AC OS=Homo sapiens OX=9606 GN=MUC5AC PE=1 SV=4 | 443 | 91 | 88 | 585.2 | 34 | 5654 | 7.02 | 16 | 19 | 11 | 92 | 104 | 96 | 15 | 17 | 11 | 69 | 79 | 69 |
| High | Q14204 | Cytoplasmic dynein 1 heavy chain 1 OS=Homo sapiens OX=9606 GN=DYNC1H1 PE=1 SV=5 | 436 | 104 | 104 | 532.1 | 27 | 4646 | 6.4 | 49 | 51 | 53 | 18 | 27 | 24 | 49 | 49 | 50 | 18 | 27 | 24 |
| High | Q9UM54 | Unconventional myosin-VI OS=Homo sapiens OX=9606 GN=MYO6 PE=1 SV=4 | 431 | 66 | 66 | 149.6 | 53 | 1294 | 8.53 | 56 | 61 | 53 | 19 | 21 | 18 | 45 | 48 | 43 | 18 | 21 | 17 |
| High | Q71U36 | Tubulin alpha-1A chain OS=Homo sapiens OX=9606 GN=TUBA1A PE=1 SV=1 | 430 | 24 | 2 | 50.1 | 61 | 451 | 5.06 | 55 | 57 | 52 | 22 | 28 | 22 | 22 | 22 | 22 | 13 | 15 | 13 |
| High | P04264 | Keratin, type II cytoskeletal 1 OS=Homo sapiens OX=9606 GN=KRT1 PE=1 SV=6 | 418 | 34 | 27 | 66 | 57 | 644 | 8.12 | 52 | 42 | 62 | 45 | 45 | 40 | 29 | 26 | 31 | 27 | 24 | 24 |
| High | Q6N021 | Methylcytosine dioxygenase TET2 OS=Homo sapiens OX=9606 GN=TET2 PE=1 SV=3 | 406 | 71 | 68 | 223.7 | 46 | 2002 | 7.99 | | | | 67 | 80 | 63 | | | | 49 | 56 | 48 |
| High | O15047 | Histone-lysine N-methyltransferase SETD1A OS=Homo sapiens OX=9606 GN=SETD1A PE=1 SV=3 | 405 | 43 | 43 | 185.9 | 30 | 1707 | 5.14 | | | | 54 | 70 | 56 | | | | 33 | 37 | 36 |
| High | P68104 | Elongation factor 1-alpha 1 OS=Homo sapiens OX=9606 GN=EEF1A1 PE=1 SV=1 | 403 | 18 | 9 | 50.1 | 63 | 462 | 9.01 | 90 | 84 | 49 | 16 | 29 | 14 | 16 | 17 | 14 | 10 | 12 | 8 |
| High | Q13509 | Tubulin beta-3 chain OS=Homo sapiens OX=9606 GN=TUBB3 PE=1 SV=2 | 394 | 17 | 2 | 50.4 | 43 | 450 | 4.93 | 56 | 50 | 50 | 30 | 32 | 32 | 14 | 15 | 12 | 11 | 10 | 11 |
| High | P58107 | Epiplakin OS=Homo sapiens OX=9606 GN=EPPK1 PE=1 SV=3 | 386 | 90 | 83 | 555.3 | 48 | 5088 | 5.62 | 81 | 75 | 70 | 17 | 23 | 21 | 70 | 62 | 60 | 17 | 18 | 20 |
| High | Q9BQE3 | Tubulin alpha-1C chain OS=Homo sapiens OX=9606 GN=TUBA1C PE=1 SV=1 | 385 | 24 | 3 | 49.9 | 63 | 449 | 5.1 | 51 | 54 | 48 | 22 | 29 | 22 | 22 | 23 | 22 | 13 | 16 | 13 |
| High | P00761 SV | Trypsin - Sus scrofa (Pig). | 381 | 7 | 7 | 24.4 | 38 | 231 | 7.18 | 30 | 35 | 38 | 37 | 51 | 44 | 5 | 4 | 4 | 3 | 4 | 4 |
| High | P19338 | Nucleolin OS=Homo sapiens OX=9606 GN=NCL PE=1 SV=3 | 366 | 26 | 26 | 76.6 | 38 | 710 | 4.7 | 27 | 27 | 27 | 32 | 50 | 28 | 19 | 16 | 17 | 17 | 20 | 16 |
| High | P68366 | Tubulin alpha-4A chain OS=Homo sapiens OX=9606 GN=TUBA4A PE=1 SV=1 | 351 | 23 | 6 | 49.9 | 59 | 448 | 5.06 | 55 | 56 | 44 | 18 | 20 | 17 | 20 | 21 | 20 | 12 | 14 | 11 |
| High | Q9ULV0 | Unconventional myosin-Vb OS=Homo sapiens OX=9606 GN=MYO5B PE=1 SV=3 | 338 | 68 | 64 | 213.5 | 44 | 1848 | 7.2 | 60 | 58 | 53 | 8 | 13 | 11 | 50 | 52 | 47 | 8 | 13 | 11 |
| High | P14618 | Pyruvate kinase PKM OS=Homo sapiens OX=9606 GN=PKM PE=1 SV=4 | 335 | 27 | 27 | 57.9 | 63 | 531 | 7.84 | 40 | 42 | 43 | 20 | 28 | 21 | 24 | 23 | 22 | 14 | 18 | 14 |
| High | P20700 | Lamin-B1 OS=Homo sapiens OX=9606 GN=LMNB1 PE=1 SV=2 | 334 | 36 | 33 | 66.4 | 63 | 586 | 5.16 | 52 | 56 | 63 | 17 | 25 | 20 | 32 | 34 | 33 | 14 | 18 | 18 |
| High | P35900 | SWISS-PROT:P35900 Tax_Id=9606 Gene_Symbol=KRT20 Keratin, type I cytoskeletal 20 | 332 | 30 | 27 | 48.5 | 69 | 424 | 5.69 | 69 | 77 | 63 | 17 | 20 | 16 | 29 | 29 | 28 | 13 | 13 | 14 |
| High | P0DMV9 | Heat shock 70 kDa protein 1B OS=Homo sapiens OX=9606 GN=HSPA1B PE=1 SV=1 | 321 | 27 | 23 | 70 | 54 | 641 | 5.66 | 33 | 32 | 34 | 24 | 30 | 25 | 20 | 20 | 22 | 16 | 17 | 17 |
| High | P35908 | Keratin, type II cytoskeletal 2 epidermal OS=Homo sapiens OX=9606 GN=KRT2 PE=1 SV=2 | 319 | 41 | 1 | 65.4 | 65 | 639 | 8 | 43 | 39 | 91 | 20 | 26 | 24 | 29 | 23 | 40 | 16 | 18 | 17 |
| High | P35908 | SWISS-PROT:P35908 Tax_Id=9606 Gene_Symbol=KRT2 Keratin, type II cytoskeletal 2 epidermal | 319 | 41 | 1 | 65.8 | 66 | 645 | 8 | 43 | 39 | 91 | 20 | 26 | 24 | 29 | 23 | 40 | 16 | 18 | 17 |
| High | P13010 | X-ray repair cross-complementing protein 5 OS=Homo sapiens OX=9606 GN=XRCC5 PE=1 SV=3 | 302 | 37 | 37 | 82.7 | 63 | 732 | 5.81 | 51 | 47 | 52 | 14 | 19 | 17 | 34 | 29 | 28 | 11 | 15 | 12 |
| High | Q86V48 | Leucine zipper protein 1 OS=Homo sapiens OX=9606 GN=LUZP1 PE=1 SV=2 | 302 | 47 | 47 | 120.2 | 50 | 1076 | 8.5 | 42 | 43 | 48 | 13 | 14 | 15 | 34 | 38 | 38 | 13 | 14 | 15 |
| High | O43707 | Alpha-actinin-4 OS=Homo sapiens OX=9606 GN=ACTN4 PE=1 SV=2 | 287 | 42 | 29 | 104.8 | 59 | 911 | 5.44 | 45 | 47 | 49 | 11 | 14 | 13 | 32 | 31 | 33 | 10 | 12 | 11 |
| High | P13639 | Elongation factor 2 OS=Homo sapiens OX=9606 GN=EEF2 PE=1 SV=4 | 285 | 40 | 39 | 95.3 | 56 | 858 | 6.83 | 41 | 35 | 32 | 18 | 33 | 18 | 29 | 24 | 27 | 16 | 20 | 16 |
| High | Q7Z6Z7 | E3 ubiquitin-protein ligase HUWE1 OS=Homo sapiens OX=9606 GN=HUWE1 PE=1 SV=3 | 280 | 73 | 73 | 481.6 | 23 | 4374 | 5.22 | 13 | 12 | 11 | 55 | 63 | 56 | 12 | 11 | 10 | 49 | 54 | 51 |
| High | P11021 | Endoplasmic reticulum chaperone BiP OS=Homo sapiens OX=9606 GN=HSPA5 PE=1 SV=2 | 278 | 28 | 26 | 72.3 | 49 | 654 | 5.16 | 25 | 25 | 27 | 26 | 37 | 27 | 19 | 19 | 21 | 20 | 23 | 22 |
| High | Q8TDZ2 | [F-actin]-monooxygenase MICAL1 OS=Homo sapiens OX=9606 GN=MICAL1 PE=1 SV=2 | 273 | 44 | 44 | 117.8 | 52 | 1067 | 6.4 | 68 | 68 | 80 | 6 | 8 | 6 | 37 | 40 | 38 | 5 | 7 | 5 |
| High | A2NJV5 | Immunoglobulin kappa variable 2-29 OS=Homo sapiens OX=9606 GN=IGKV2-29 PE=3 SV=2 | 272 | 3 | 2 | 13.1 | 17 | 120 | 7.28 | 20 | 23 | 18 | 34 | 51 | 48 | 2 | 2 | 2 | 2 | 2 | 2 |
| High | P49411 | Elongation factor Tu, mitochondrial OS=Homo sapiens OX=9606 GN=TUFM PE=1 SV=2 | 267 | 26 | 26 | 49.5 | 59 | 452 | 7.61 | 38 | 37 | 41 | 15 | 19 | 20 | 26 | 26 | 23 | 14 | 16 | 17 |
| High | P35527 | SWISS-PROT:P35527 Tax_Id=9606 Gene_Symbol=KRT9 Keratin, type I cytoskeletal 9 | 264 | 26 | 25 | 62.1 | 57 | 623 | 5.3 | 41 | 33 | 38 | 22 | 22 | 24 | 22 | 18 | 22 | 15 | 14 | 16 |
| High | P13645 | SWISS-PROT:P13645 Tax_Id=9606 Gene_Symbol=KRT10 Keratin, type I cytoskeletal 10 | 263 | 27 | 23 | 59.5 | 50 | 593 | 5.21 | 38 | 38 | 64 | 18 | 28 | 22 | 24 | 18 | 25 | 14 | 18 | 17 |
| High | P19474 | E3 ubiquitin-protein ligase TRIM21 OS=Homo sapiens OX=9606 GN=TRIM21 PE=1 SV=1 | 262 | 26 | 26 | 54.1 | 52 | 475 | 6.38 | 32 | 30 | 30 | 28 | 32 | 24 | 23 | 22 | 21 | 15 | 17 | 16 |
| High | P46060 | Ran GTPase-activating protein 1 OS=Homo sapiens OX=9606 GN=RANGAP1 PE=1 SV=1 | 260 | 29 | 29 | 63.5 | 53 | 587 | 4.68 | 1 | 1 | 1 | 5 | 6 | 5 | 1 | 1 | 1 | 5 | 6 | 5 |
| High | P12956 | X-ray repair cross-complementing protein 6 OS=Homo sapiens OX=9606 GN=XRCC6 PE=1 SV=2 | 260 | 31 | 31 | 69.8 | 52 | 609 | 6.64 | 47 | 42 | 37 | 15 | 21 | 19 | 28 | 28 | 28 | 12 | 15 | 15 |
| High | O94832 | Unconventional myosin-Id OS=Homo sapiens OX=9606 GN=MYO1D PE=1 SV=2 | 259 | 43 | 43 | 116.1 | 45 | 1006 | 9.39 | 29 | 33 | 29 | 13 | 17 | 11 | 28 | 30 | 26 | 13 | 16 | 11 |
| High | P27708 | CAD protein OS=Homo sapiens OX=9606 GN=CAD PE=1 SV=3 | 258 | 54 | 54 | 242.8 | 33 | 2225 | 6.46 | 38 | 36 | 40 | 13 | 23 | 20 | 36 | 35 | 34 | 12 | 20 | 20 |
| High | Q9P2E9 | Ribosome-binding protein 1 OS=Homo sapiens OX=9606 GN=RRBP1 PE=1 SV=5 | 258 | 43 | 43 | 152.4 | 41 | 1410 | 8.6 | 17 | 19 | 23 | 27 | 30 | 27 | 17 | 18 | 23 | 26 | 28 | 27 |
| High | Q9NYL9 | Tropomodulin-3 OS=Homo sapiens OX=9606 GN=TMOD3 PE=1 SV=1 | 257 | 22 | 22 | 39.6 | 62 | 352 | 5.19 | 38 | 34 | 34 | 10 | 16 | 12 | 22 | 21 | 21 | 10 | 14 | 11 |
| High | P15822 | Zinc finger protein 40 OS=Homo sapiens OX=9606 GN=HIVEP1 PE=1 SV=4 | 253 | 62 | 60 | 296.7 | 31 | 2718 | 7.84 | | | | 48 | 61 | 48 | | | | 43 | 50 | 44 |
| High | Q05639 | Elongation factor 1-alpha 2 OS=Homo sapiens OX=9606 GN=EEF1A2 PE=1 SV=1 | 251 | 14 | 5 | 50.4 | 50 | 463 | 9.03 | 71 | 70 | 29 | 12 | 19 | 12 | 12 | 13 | 10 | 6 | 7 | 6 |
| High | P10809 | 60 kDa heat shock protein, mitochondrial OS=Homo sapiens OX=9606 GN=HSPD1 PE=1 SV=2 | 251 | 28 | 28 | 61 | 63 | 573 | 5.87 | 31 | 30 | 24 | 17 | 19 | 17 | 18 | 19 | 19 | 14 | 17 | 15 |
| High | P05784 | SWISS-PROT:P05784 Tax_Id=10090 Gene_Symbol=Krt18 Keratin, type I cytoskeletal 18 | 248 | 12 | 1 | 47.5 | 24 | 423 | 5.33 | 49 | 55 | 52 | 14 | 17 | 16 | 11 | 11 | 9 | 7 | 7 | 7 |
| High | Q14980 | Nuclear mitotic apparatus protein 1 OS=Homo sapiens OX=9606 GN=NUMA1 PE=1 SV=2 | 248 | 53 | 53 | 238.1 | 32 | 2115 | 5.78 | 28 | 30 | 36 | 20 | 21 | 20 | 25 | 28 | 31 | 19 | 21 | 19 |
| High | Q9UHB6 | LIM domain and actin-binding protein 1 OS=Homo sapiens OX=9606 GN=LIMA1 PE=1 SV=1 | 247 | 36 | 36 | 85.2 | 58 | 759 | 6.84 | 28 | 33 | 33 | 10 | 11 | 10 | 24 | 26 | 24 | 10 | 11 | 10 |
| High | P14649 | Myosin light chain 6B OS=Homo sapiens OX=9606 GN=MYL6B PE=1 SV=1 | 243 | 12 | 10 | 22.8 | 54 | 208 | 5.73 | 24 | 29 | 26 | 25 | 22 | 22 | 9 | 10 | 8 | 7 | 8 | 7 |
| High | P02538 | SWISS-PROT:P02538 Tax_Id=9606 Gene_Symbol=KRT6A Keratin, type II cytoskeletal 6A | 242 | 20 | 5 | 60 | 29 | 564 | 8 | 38 | 40 | 57 | 15 | 15 | 17 | 12 | 10 | 14 | 9 | 8 | 9 |
| High | REFSEQ:XI | Tax_Id=9606 Gene_Symbol=LOC150739 similar to Keratin, type II cytoskeletal 8 | 241 | 4 | 1 | 23.1 | 22 | 213 | 7.59 | 57 | 57 | 73 | 11 | 8 | 8 | 3 | 3 | 4 | 2 | 2 | 2 |
| High | P09493 | Tropomyosin alpha-1 chain OS=Homo sapiens OX=9606 GN=TPM1 PE=1 SV=2 | 240 | 24 | 10 | 32.7 | 52 | 284 | 4.74 | 28 | 25 | 32 | 23 | 30 | 23 | 15 | 13 | 17 | 15 | 21 | 14 |
| High | O43151 | Methylcytosine dioxygenase TET3 OS=Homo sapiens OX=9606 GN=TET3 PE=1 SV=4 | 234 | 54 | 51 | 193.6 | 40 | 1795 | 7.81 | | | | 44 | 53 | 47 | | | | 35 | 41 | 38 |
| High | Q14160 | Protein scribble homolog OS=Homo sapiens OX=9606 GN=SCRIB PE=1 SV=5 | 232 | 49 | 47 | 174.8 | 40 | 1630 | 5.07 | 31 | 32 | 35 | 8 | 8 | 9 | 31 | 32 | 34 | 8 | 8 | 9 |
| High | Q9UPQ0 | LIM and calponin homology domains-containing protein 1 OS=Homo sapiens OX=9606 GN=LIMCH1 PE=1 SV=4 | 232 | 40 | 40 | 121.8 | 46 | 1083 | 6.47 | 38 | 36 | 40 | 9 | 10 | 9 | 30 | 28 | 32 | 9 | 10 | 9 |
| High | P08238 | Heat shock protein HSP 90-beta OS=Homo sapiens OX=9606 GN=HSP90AB1 PE=1 SV=4 | 230 | 30 | 17 | 83.2 | 51 | 724 | 5.03 | 20 | 27 | 33 | 24 | 21 | 20 | 18 | 18 | 24 | 17 | 18 | 15 |
| High | P08730-1 | SWISS-PROT:P08730-1 Tax_Id=10090 Gene_Symbol=Krt13 Isoform 1 of Keratin, type I cytoskeletal 13 | 230 | 9 | 1 | 47.7 | 19 | 437 | 4.86 | 56 | 56 | 53 | 9 | 14 | 10 | 9 | 9 | 9 | 5 | 6 | 5 |
| High | P00352 | Aldehyde dehydrogenase 1A1 OS=Homo sapiens OX=9606 GN=ALDH1A1 PE=1 SV=2 | 229 | 25 | 22 | 54.8 | 56 | 501 | 6.73 | 23 | 23 | 29 | 9 | 16 | 9 | 17 | 18 | 18 | 9 | 14 | 9 |
| High | Q3SX28 | TREMBL:Q3SX28;Q5KR48 (Bos taurus) Tropomyosin 2 | 229 | 21 | 5 | 33 | 46 | 284 | 4.69 | 29 | 26 | 30 | 18 | 25 | 17 | 16 | 15 | 16 | 13 | 17 | 12 |
| High | P61978 | Heterogeneous nuclear ribonucleoprotein K OS=Homo sapiens OX=9606 GN=HNRNPK PE=1 SV=1 | 229 | 19 | 19 | 50.9 | 53 | 463 | 5.54 | 21 | 24 | 23 | 20 | 25 | 19 | 12 | 14 | 15 | 13 | 14 | 13 |
| High | Q08211 | ATP-dependent RNA helicase A OS=Homo sapiens OX=9606 GN=DHX9 PE=1 SV=4 | 228 | 37 | 37 | 140.9 | 36 | 1270 | 6.84 | 24 | 26 | 26 | 20 | 19 | 20 | 22 | 24 | 24 | 18 | 18 | 18 |
| High | P22102 | Trifunctional purine biosynthetic protein adenosine-3 OS=Homo sapiens OX=9606 GN=GART PE=1 SV=1 | 226 | 36 | 36 | 107.7 | 46 | 1010 | 6.7 | 23 | 25 | 29 | 17 | 18 | 16 | 20 | 21 | 22 | 15 | 14 | 16 |
| High | P07384 | Calpain-1 catalytic subunit OS=Homo sapiens OX=9606 GN=CAPN1 PE=1 SV=1 | 226 | 32 | 32 | 81.8 | 45 | 714 | 5.67 | 31 | 36 | 37 | 12 | 14 | 11 | 25 | 28 | 24 | 12 | 14 | 11 |
| High | Q92598 | Heat shock protein 105 kDa OS=Homo sapiens OX=9606 GN=HSPH1 PE=1 SV=1 | 225 | 36 | 33 | 96.8 | 50 | 858 | 5.39 | 20 | 21 | 23 | 16 | 23 | 14 | 18 | 20 | 23 | 15 | 19 | 13 |
| High | P63010 | AP-2 complex subunit beta OS=Homo sapiens OX=9606 GN=AP2B1 PE=1 SV=1 | 224 | 38 | 23 | 104.5 | 47 | 937 | 5.38 | 24 | 32 | 31 | 10 | 9 | 14 | 23 | 30 | 26 | 10 | 7 | 13 |
| High | Q13045 | Protein flightless-1 homolog OS=Homo sapiens OX=9606 GN=FLII PE=1 SV=2 | 223 | 38 | 38 | 144.7 | 38 | 1269 | 6.05 | 34 | 34 | 34 | 7 | 12 | 5 | 26 | 27 | 26 | 6 | 11 | 5 |
| High | P23396 | 40S ribosomal protein S3 OS=Homo sapiens OX=9606 GN=RPS3 PE=1 SV=2 | 223 | 19 | 19 | 26.7 | 79 | 243 | 9.66 | 28 | 29 | 30 | 13 | 19 | 18 | 17 | 16 | 16 | 12 | 11 | 15 |
| High | P23528 | Cofilin-1 OS=Homo sapiens OX=9606 GN=CFL1 PE=1 SV=3 | 222 | 14 | 14 | 18.5 | 67 | 166 | 8.09 | 34 | 32 | 27 | 11 | 17 | 13 | 12 | 12 | 13 | 7 | 9 | 7 |
| High | P07814 | Bifunctional glutamate/proline--tRNA ligase OS=Homo sapiens OX=9606 GN=EPRS1 PE=1 SV=5 | 219 | 43 | 43 | 170.5 | 38 | 1512 | 7.33 | 14 | 19 | 24 | 13 | 19 | 16 | 14 | 18 | 22 | 13 | 18 | 16 |
| High | P02533 | Keratin, type I cytoskeletal 14 OS=Homo sapiens OX=9606 GN=KRT14 PE=1 SV=4 | 218 | 19 | 2 | 51.5 | 36 | 472 | 5.16 | 37 | 40 | 48 | 11 | 17 | 15 | 13 | 12 | 15 | 7 | 9 | 10 |
| High | O00159 | Unconventional myosin-Ic OS=Homo sapiens OX=9606 GN=MYO1C PE=1 SV=4 | 216 | 40 | 40 | 121.6 | 43 | 1063 | 9.41 | 32 | 29 | 31 | 15 | 10 | 12 | 30 | 26 | 28 | 15 | 10 | 12 |
| High | Q2KHR3 | Glutamine and serine-rich protein 1 OS=Homo sapiens OX=9606 GN=QSER1 PE=1 SV=3 | 214 | 41 | 41 | 189.9 | 34 | 1735 | 7.08 | | | | 29 | 46 | 32 | | | | 26 | 33 | 25 |
| High | O75688 | Protein phosphatase 1B OS=Homo sapiens OX=9606 GN=PPM1B PE=1 SV=1 | 214 | 19 | 16 | 52.6 | 48 | 479 | 5.05 | 24 | 29 | 18 | 14 | 30 | 14 | 16 | 18 | 13 | 9 | 14 | 10 |
| High | Q13085 | Acetyl-CoA carboxylase 1 OS=Homo sapiens OX=9606 GN=ACACA PE=1 SV=2 | 213 | 57 | 50 | 265.4 | 32 | 2346 | 6.37 | 28 | 28 | 32 | 14 | 10 | 14 | 27 | 28 | 31 | 14 | 10 | 14 |
| High | O95782 | AP-2 complex subunit alpha-1 OS=Homo sapiens OX=9606 GN=AP2A1 PE=1 SV=3 | 211 | 37 | 27 | 107.5 | 43 | 977 | 7.03 | 25 | 28 | 20 | 16 | 15 | 11 | 24 | 28 | 19 | 14 | 15 | 11 |
| High | P16615 | Sarcoplasmic/endoplasmic reticulum calcium ATPase 2 OS=Homo sapiens OX=9606 GN=ATP2A2 PE=1 SV=1 | 211 | 34 | 29 | 114.7 | 36 | 1042 | 5.34 | 28 | 29 | 34 | 13 | 15 | 16 | 25 | 25 | 27 | 12 | 13 | 14 |
| High | P00390 | Glutathione reductase, mitochondrial OS=Homo sapiens OX=9606 GN=GSR PE=1 SV=2 | 209 | 19 | 19 | 56.2 | 61 | 522 | 8.5 | 22 | 26 | 22 | 14 | 17 | 11 | 16 | 16 | 14 | 10 | 13 | 9 |
| High | Q9P0U4 | CXXC-type zinc finger protein 1 OS=Homo sapiens OX=9606 GN=CXXC1 PE=1 SV=2 | 206 | 29 | 29 | 75.7 | 46 | 656 | 8.24 | | | | 31 | 36 | 23 | | | | 22 | 24 | 19 |
| High | P62937 | Peptidyl-prolyl cis-trans isomerase A OS=Homo sapiens OX=9606 GN=PPIA PE=1 SV=2 | 205 | 9 | 9 | 18 | 53 | 165 | 7.81 | 23 | 18 | 24 | 17 | 29 | 19 | 9 | 8 | 8 | 4 | 5 | 5 |
| High | Q9BUF5 | Tubulin beta-6 chain OS=Homo sapiens OX=9606 GN=TUBB6 PE=1 SV=1 | 204 | 10 | 2 | 49.8 | 29 | 446 | 4.88 | 31 | 25 | 26 | 17 | 18 | 18 | 8 | 8 | 7 | 6 | 5 | 5 |
| High | P13647 | SWISS-PROT:P13647 Tax_Id=9606 Gene_Symbol=KRT5 Keratin, type II cytoskeletal 5 | 204 | 26 | 13 | 62.3 | 38 | 590 | 7.74 | 25 | 25 | 54 | 14 | 14 | 17 | 14 | 8 | 20 | 10 | 9 | 11 |
| High | Q9NR30 | Nucleolar RNA helicase 2 OS=Homo sapiens OX=9606 GN=DDX21 PE=1 SV=5 | 202 | 30 | 28 | 87.3 | 39 | 783 | 9.28 | 11 | 12 | 13 | 23 | 24 | 20 | 11 | 12 | 12 | 21 | 18 | 17 |
| High | P13646-1 | SWISS-PROT:P13646-1 Tax_Id=9606 Gene_Symbol=KRT13 Isoform 1 of Keratin, type I cytoskeletal 13 | 198 | 16 | 8 | 49.6 | 34 | 458 | 4.96 | 49 | 48 | 47 | 6 | 12 | 5 | 13 | 14 | 12 | 5 | 6 | 4 |
| High | P62805 | Histone H4 OS=Homo sapiens OX=9606 GN=H4C1 PE=1 SV=2 | 198 | 7 | 7 | 11.4 | 52 | 103 | 11.36 | 20 | 19 | 28 | 24 | 22 | 23 | 5 | 6 | 7 | 6 | 6 | 6 |
| High | Q92841 | Probable ATP-dependent RNA helicase DDX17 OS=Homo sapiens OX=9606 GN=DDX17 PE=1 SV=2 | 198 | 29 | 21 | 80.2 | 47 | 729 | 8.27 | 20 | 18 | 19 | 20 | 21 | 19 | 19 | 17 | 18 | 18 | 18 | 17 |
| High | Q04695 | SWISS-PROT:Q04695 Tax_Id=9606 Gene_Symbol=KRT17 Keratin, type I cytoskeletal 17 | 198 | 18 | 6 | 48.1 | 33 | 432 | 5.02 | 38 | 42 | 41 | 11 | 20 | 13 | 15 | 16 | 15 | 7 | 12 | 8 |
| High | Q76L83 | Putative Polycomb group protein ASXL2 OS=Homo sapiens OX=9606 GN=ASXL2 PE=1 SV=1 | 196 | 41 | 40 | 153.7 | 40 | 1435 | 8.81 | | | | 31 | 32 | 30 | | | | 31 | 30 | 30 |
| High | P17844 | Probable ATP-dependent RNA helicase DDX5 OS=Homo sapiens OX=9606 GN=DDX5 PE=1 SV=1 | 196 | 28 | 20 | 69.1 | 44 | 614 | 8.92 | 23 | 22 | 22 | 18 | 21 | 19 | 21 | 19 | 20 | 16 | 19 | 16 |
| High | P06733 | Alpha-enolase OS=Homo sapiens OX=9606 GN=ENO1 PE=1 SV=2 | 195 | 20 | 20 | 47.1 | 57 | 434 | 7.39 | 18 | 18 | 24 | 19 | 27 | 19 | 13 | 15 | 16 | 12 | 15 | 12 |
| High | P22626 | Heterogeneous nuclear ribonucleoproteins A2/B1 OS=Homo sapiens OX=9606 GN=HNRNPA2B1 PE=1 SV=2 | 194 | 18 | 16 | 37.4 | 56 | 353 | 8.95 | 31 | 39 | 31 | 14 | 14 | 14 | 15 | 17 | 14 | 10 | 12 | 11 |
| High | P62424 | 60S ribosomal protein L7a OS=Homo sapiens OX=9606 GN=RPL7A PE=1 SV=2 | 193 | 14 | 14 | 30 | 42 | 266 | 10.61 | 14 | 14 | 16 | 17 | 15 | 18 | 10 | 12 | 12 | 11 | 12 | 12 |
| High | Q15233 | Non-POU domain-containing octamer-binding protein OS=Homo sapiens OX=9606 GN=NONO PE=1 SV=4 | 193 | 18 | 17 | 54.2 | 49 | 471 | 8.95 | 17 | 19 | 23 | 19 | 28 | 22 | 13 | 14 | 16 | 15 | 16 | 12 |
| High | Q13126 | S-methyl-5'-thioadenosine phosphorylase OS=Homo sapiens OX=9606 GN=MTAP PE=1 SV=2 | 191 | 16 | 16 | 31.2 | 73 | 283 | 7.18 | 19 | 18 | 22 | 15 | 20 | 14 | 14 | 14 | 14 | 11 | 12 | 12 |
| High | Q6P3W7 | SCY1-like protein 2 OS=Homo sapiens OX=9606 GN=SCYL2 PE=1 SV=1 | 191 | 26 | 26 | 103.6 | 38 | 929 | 8.22 | 19 | 19 | 21 | 13 | 13 | 17 | 18 | 15 | 17 | 12 | 12 | 16 |
| High | Q00839 | Heterogeneous nuclear ribonucleoprotein U OS=Homo sapiens OX=9606 GN=HNRNPU PE=1 SV=6 | 191 | 26 | 26 | 90.5 | 37 | 825 | 6 | 22 | 25 | 33 | 18 | 17 | 18 | 18 | 19 | 21 | 15 | 15 | 14 |
| High | O15027 | Protein transport protein Sec16A OS=Homo sapiens OX=9606 GN=SEC16A PE=1 SV=4 | 190 | 45 | 45 | 251.7 | 31 | 2357 | 5.8 | 20 | 22 | 23 | 11 | 11 | 13 | 18 | 21 | 22 | 11 | 11 | 13 |
| High | P08779 | SWISS-PROT:P08779 Tax_Id=9606 Gene_Symbol=KRT16 Keratin, type I cytoskeletal 16 | 188 | 15 | 3 | 51.2 | 30 | 473 | 5.05 | 30 | 34 | 39 | 11 | 16 | 12 | 9 | 8 | 10 | 7 | 8 | 7 |
| High | P52272 | Heterogeneous nuclear ribonucleoprotein M OS=Homo sapiens OX=9606 GN=HNRNPM PE=1 SV=3 | 184 | 31 | 31 | 77.5 | 48 | 730 | 8.7 | 21 | 20 | 18 | 17 | 13 | 17 | 21 | 20 | 17 | 16 | 13 | 15 |
| High | P67936 | Tropomyosin alpha-4 chain OS=Homo sapiens OX=9606 GN=TPM4 PE=1 SV=3 | 183 | 20 | 8 | 28.5 | 58 | 248 | 4.69 | 24 | 23 | 27 | 14 | 22 | 12 | 12 | 12 | 15 | 11 | 14 | 9 |
| High | Q14764 | Major vault protein OS=Homo sapiens OX=9606 GN=MVP PE=1 SV=4 | 180 | 30 | 30 | 99.3 | 43 | 893 | 5.48 | 13 | 15 | 15 | 17 | 20 | 17 | 13 | 14 | 15 | 16 | 18 | 15 |
| High | P09327 | Villin-1 OS=Homo sapiens OX=9606 GN=VIL1 PE=1 SV=4 | 180 | 29 | 29 | 92.6 | 43 | 827 | 6.39 | 23 | 26 | 24 | 13 | 17 | 11 | 20 | 22 | 21 | 13 | 14 | 11 |
| High | P62979 | Ubiquitin-40S ribosomal protein S27a OS=Homo sapiens OX=9606 GN=RPS27A PE=1 SV=2 | 180 | 7 | 7 | 18 | 51 | 156 | 9.64 | 18 | 16 | 22 | 20 | 23 | 19 | 7 | 7 | 7 | 6 | 7 | 7 |
| High | P51659 | Peroxisomal multifunctional enzyme type 2 OS=Homo sapiens OX=9606 GN=HSD17B4 PE=1 SV=3 | 180 | 24 | 24 | 79.6 | 47 | 736 | 8.84 | 21 | 24 | 26 | 11 | 13 | 12 | 18 | 19 | 20 | 11 | 12 | 11 |
| High | Q15291 | Retinoblastoma-binding protein 5 OS=Homo sapiens OX=9606 GN=RBBP5 PE=1 SV=2 | 180 | 23 | 23 | 59.1 | 51 | 538 | 5.1 | | | 1 | 24 | 25 | 23 | | | 1 | 16 | 17 | 16 |
| High | P07900 | Heat shock protein HSP 90-alpha OS=Homo sapiens OX=9606 GN=HSP90AA1 PE=1 SV=5 | 177 | 26 | 14 | 84.6 | 44 | 732 | 5.02 | 18 | 21 | 21 | 19 | 20 | 18 | 17 | 16 | 16 | 15 | 17 | 14 |
| High | P14625 | Endoplasmin OS=Homo sapiens OX=9606 GN=HSP90B1 PE=1 SV=1 | 176 | 28 | 27 | 92.4 | 39 | 803 | 4.84 | 22 | 25 | 24 | 13 | 22 | 7 | 17 | 15 | 23 | 13 | 16 | 7 |
| High | P46940 | Ras GTPase-activating-like protein IQGAP1 OS=Homo sapiens OX=9606 GN=IQGAP1 PE=1 SV=1 | 176 | 43 | 41 | 189.1 | 33 | 1657 | 6.48 | 22 | 21 | 26 | 5 | 11 | 10 | 22 | 19 | 24 | 5 | 10 | 10 |
| High | Q12906 | Interleukin enhancer-binding factor 3 OS=Homo sapiens OX=9606 GN=ILF3 PE=1 SV=3 | 175 | 22 | 20 | 95.3 | 32 | 894 | 8.76 | 15 | 17 | 16 | 18 | 21 | 17 | 13 | 15 | 14 | 14 | 16 | 14 |
| High | P47756 | F-actin-capping protein subunit beta OS=Homo sapiens OX=9606 GN=CAPZB PE=1 SV=4 | 175 | 13 | 13 | 31.3 | 43 | 277 | 5.59 | 16 | 18 | 23 | 15 | 18 | 12 | 9 | 10 | 12 | 9 | 8 | 7 |
| High | P0CG39 | POTE ankyrin domain family member J OS=Homo sapiens OX=9606 GN=POTEJ PE=3 SV=1 | 174 | 6 | 3 | 117.3 | 8 | 1038 | 5.97 | 32 | 32 | 17 | 8 | 23 | 9 | 5 | 4 | 4 | 3 | 3 | 3 |
| High | P22314 | Ubiquitin-like modifier-activating enzyme 1 OS=Homo sapiens OX=9606 GN=UBA1 PE=1 SV=3 | 174 | 31 | 31 | 117.8 | 37 | 1058 | 5.76 | 16 | 14 | 21 | 16 | 18 | 16 | 13 | 13 | 18 | 15 | 16 | 15 |
| High | P11940 | Polyadenylate-binding protein 1 OS=Homo sapiens OX=9606 GN=PABPC1 PE=1 SV=2 | 174 | 22 | 16 | 70.6 | 44 | 636 | 9.5 | 16 | 17 | 16 | 16 | 20 | 18 | 14 | 14 | 15 | 14 | 17 | 15 |
| High | Q8IZD2 | Inactive histone-lysine N-methyltransferase 2E OS=Homo sapiens OX=9606 GN=KMT2E PE=1 SV=1 | 173 | 39 | 38 | 204.8 | 28 | 1858 | 7.61 | | | | 25 | 29 | 27 | | | | 22 | 26 | 23 |
| High | P53621 | Coatomer subunit alpha OS=Homo sapiens OX=9606 GN=COPA PE=1 SV=2 | 173 | 36 | 36 | 138.3 | 35 | 1224 | 7.66 | 21 | 21 | 20 | 9 | 7 | 10 | 21 | 20 | 20 | 9 | 7 | 10 |
| High | P41250 | Glycine--tRNA ligase OS=Homo sapiens OX=9606 GN=GARS1 PE=1 SV=3 | 171 | 27 | 27 | 83.1 | 45 | 739 | 7.03 | 29 | 28 | 28 | 11 | 13 | 11 | 21 | 19 | 23 | 11 | 12 | 10 |
| High | P25205 | DNA replication licensing factor MCM3 OS=Homo sapiens OX=9606 GN=MCM3 PE=1 SV=3 | 170 | 30 | 30 | 90.9 | 46 | 808 | 5.77 | 15 | 12 | 14 | 20 | 22 | 19 | 14 | 11 | 13 | 16 | 19 | 16 |
| High | P23246 | Splicing factor, proline- and glutamine-rich OS=Homo sapiens OX=9606 GN=SFPQ PE=1 SV=2 | 170 | 22 | 21 | 76.1 | 31 | 707 | 9.44 | 29 | 31 | 30 | 7 | 14 | 6 | 19 | 21 | 20 | 5 | 9 | 6 |
| High | P63244 | Receptor of activated protein C kinase 1 OS=Homo sapiens OX=9606 GN=RACK1 PE=1 SV=3 | 170 | 19 | 19 | 35.1 | 84 | 317 | 7.69 | 21 | 22 | 21 | 18 | 16 | 17 | 15 | 17 | 16 | 13 | 11 | 13 |
| High | Q9Y230 | RuvB-like 2 OS=Homo sapiens OX=9606 GN=RUVBL2 PE=1 SV=3 | 168 | 19 | 19 | 51.1 | 46 | 463 | 5.64 | 19 | 20 | 21 | 17 | 15 | 15 | 16 | 17 | 18 | 16 | 14 | 14 |
| High | O43143 | ATP-dependent RNA helicase DHX15 OS=Homo sapiens OX=9606 GN=DHX15 PE=1 SV=2 | 168 | 22 | 21 | 90.9 | 30 | 795 | 7.46 | 13 | 10 | 15 | 18 | 19 | 20 | 12 | 10 | 13 | 14 | 14 | 16 |
| High | P32119 | Peroxiredoxin-2 OS=Homo sapiens OX=9606 GN=PRDX2 PE=1 SV=5 | 168 | 11 | 10 | 21.9 | 58 | 198 | 5.97 | 21 | 21 | 22 | 12 | 15 | 13 | 11 | 11 | 10 | 7 | 8 | 6 |
| High | Q9UBL3 | Set1/Ash2 histone methyltransferase complex subunit ASH2 OS=Homo sapiens OX=9606 GN=ASH2L PE=1 SV=1 | 167 | 22 | 22 | 68.7 | 42 | 628 | 5.69 | | 1 | | 19 | 21 | 17 | | 1 | | 16 | 16 | 15 |
| High | P11413 | Glucose-6-phosphate 1-dehydrogenase OS=Homo sapiens OX=9606 GN=G6PD PE=1 SV=4 | 166 | 25 | 25 | 59.2 | 55 | 515 | 6.84 | 21 | 15 | 19 | 14 | 14 | 16 | 20 | 15 | 15 | 13 | 13 | 15 |
| High | P06753 | Tropomyosin alpha-3 chain OS=Homo sapiens OX=9606 GN=TPM3 PE=1 SV=2 | 166 | 16 | 5 | 32.9 | 37 | 285 | 4.72 | 19 | 18 | 21 | 17 | 20 | 17 | 9 | 8 | 10 | 10 | 11 | 9 |
| High | Q9UDY2 | Tight junction protein ZO-2 OS=Homo sapiens OX=9606 GN=TJP2 PE=1 SV=2 | 165 | 32 | 32 | 133.9 | 37 | 1190 | 7.4 | 14 | 16 | 18 | 10 | 16 | 11 | 14 | 16 | 17 | 10 | 15 | 11 |

| | | | | | | | | | | | | | | | | | | | | | |
|---|---|---|---|---|---|---|---|---|---|---|---|---|---|---|---|---|---|---|---|---|---|
| High | P54652 | Heat shock-related 70 kDa protein 2 OS=Homo sapiens OX=9606 GN=HSPA2 PE=1 SV=1 | 164 | 12 | 1 | 70 | 18 | 639 | 5.74 | 14 | 13 | 16 | 18 | 26 | 15 | 9 | 8 | 10 | 8 | 10 | 7 |
| High | Q9C0C2 | 182 kDa tankyrase-1-binding protein OS=Homo sapiens OX=9606 GN=TNKS1BP1 PE=1 SV=4 | 164 | 37 | 37 | 181.7 | 33 | 1729 | 4.86 | 10 | 11 | 13 | 15 | 13 | 13 | 10 | 11 | 13 | 15 | 13 | 13 |
| High | P06396 | Gelsolin OS=Homo sapiens OX=9606 GN=GSN PE=1 SV=1 | 162 | 23 | 10 | 85.6 | 39 | 782 | 6.28 | 19 | 21 | 25 | 4 | 12 | 8 | 15 | 16 | 18 | 4 | 9 | 8 |
| High | P14923 | Junction plakoglobin OS=Homo sapiens OX=9606 GN=JUP PE=1 SV=3 | 162 | 23 | 19 | 81.7 | 36 | 745 | 6.14 | 21 | 15 | 21 | 9 | 16 | 9 | 16 | 14 | 17 | 9 | 15 | 9 |
| High | Q9P258 | Protein RCC2 OS=Homo sapiens OX=9606 GN=RCC2 PE=1 SV=2 | 162 | 17 | 17 | 56 | 37 | 522 | 8.78 | 16 | 15 | 18 | 16 | 17 | 16 | 13 | 13 | 12 | 11 | 12 | 13 |
| High | P11216 | Glycogen phosphorylase, brain form OS=Homo sapiens OX=9606 GN=PYGB PE=1 SV=5 | 161 | 32 | 32 | 96.6 | 42 | 843 | 6.86 | 29 | 29 | 25 | 8 | 8 | 9 | 27 | 28 | 23 | 8 | 8 | 9 |
| High | O95678 | SWISS-PROT:O95678 Tax_Id=9606 Gene_Symbol=KRT75 Keratin, type II cytoskeletal 75 | 161 | 10 | 1 | 59.5 | 13 | 551 | 7.74 | 22 | 28 | 45 | 9 | 11 | 10 | 7 | 7 | 10 | 4 | 4 | 4 |
| High | P17987 | T-complex protein 1 subunit alpha OS=Homo sapiens OX=9606 GN=TCP1 PE=1 SV=1 | 161 | 23 | 23 | 60.3 | 51 | 556 | 6.11 | 22 | 22 | 25 | 11 | 11 | 10 | 20 | 19 | 22 | 11 | 11 | 10 |
| High | P23526 | Adenosylhomocysteinase OS=Homo sapiens OX=9606 GN=AHCY PE=1 SV=4 | 161 | 20 | 19 | 47.7 | 49 | 432 | 6.34 | 22 | 26 | 13 | 16 | 11 | 15 | 16 | 18 | 11 | 14 | 10 | 13 |
| High | P05023 | Sodium/potassium-transporting ATPase subunit alpha-1 OS=Homo sapiens OX=9606 GN=ATP1A1 PE=1 SV=1 | 160 | 28 | 28 | 112.8 | 33 | 1023 | 5.49 | 17 | 18 | 18 | 13 | 15 | 14 | 16 | 18 | 15 | 11 | 13 | 13 |
| High | Q7Z3B3 | KAT8 regulatory NSL complex subunit 1 OS=Homo sapiens OX=9606 GN=KANSL1 PE=1 SV=2 | 158 | 31 | 31 | 121 | 42 | 1105 | 8.81 | | | | 23 | 28 | 23 | | | | 19 | 23 | 19 |
| High | P14868 | Aspartate--tRNA ligase, cytoplasmic OS=Homo sapiens OX=9606 GN=DARS1 PE=1 SV=2 | 158 | 27 | 27 | 57.1 | 60 | 501 | 6.55 | 16 | 14 | 18 | 16 | 15 | 13 | 16 | 13 | 17 | 16 | 15 | 13 |
| High | P11387 | DNA topoisomerase 1 OS=Homo sapiens OX=9606 GN=TOP1 PE=1 SV=2 | 158 | 25 | 25 | 90.7 | 32 | 765 | 9.31 | 13 | 14 | 14 | 16 | 18 | 16 | 13 | 13 | 13 | 15 | 16 | 14 |
| High | Q01813 | ATP-dependent 6-phosphofructokinase, platelet type OS=Homo sapiens OX=9606 GN=PFKP PE=1 SV=2 | 156 | 26 | 22 | 85.5 | 41 | 784 | 7.55 | 24 | 24 | 22 | 8 | 11 | 8 | 22 | 21 | 19 | 8 | 10 | 8 |
| High | O00571 | ATP-dependent RNA helicase DDX3X OS=Homo sapiens OX=9606 GN=DDX3X PE=1 SV=3 | 155 | 26 | 25 | 73.2 | 45 | 662 | 7.18 | 20 | 22 | 17 | 13 | 16 | 14 | 17 | 17 | 14 | 12 | 14 | 12 |
| High | Q15365 | Poly(rC)-binding protein 1 OS=Homo sapiens OX=9606 GN=PCBP1 PE=1 SV=2 | 154 | 13 | 9 | 37.5 | 61 | 356 | 7.09 | 13 | 13 | 15 | 17 | 17 | 17 | 11 | 11 | 10 | 9 | 9 | 9 |
| High | P30740 | Leukocyte elastase inhibitor OS=Homo sapiens OX=9606 GN=SERPINB1 PE=1 SV=1 | 154 | 19 | 18 | 42.7 | 53 | 379 | 6.28 | 16 | 15 | 19 | 15 | 14 | 12 | 16 | 15 | 14 | 11 | 7 | 11 |
| High | P06748 | Nucleophosmin OS=Homo sapiens OX=9606 GN=NPM1 PE=1 SV=2 | 153 | 11 | 11 | 32.6 | 52 | 294 | 4.78 | 17 | 13 | 15 | 5 | 14 | 7 | 8 | 7 | 7 | 3 | 7 | 5 |
| High | P62701 | 40S ribosomal protein S4, X isoform OS=Homo sapiens OX=9606 GN=RPS4X PE=1 SV=2 | 152 | 16 | 16 | 29.6 | 51 | 263 | 10.15 | 17 | 15 | 15 | 15 | 17 | 16 | 13 | 12 | 13 | 10 | 13 | 9 |
| High | O14639 | Actin-binding LIM protein 1 OS=Homo sapiens OX=9606 GN=ABLIM1 PE=1 SV=3 | 152 | 25 | 25 | 87.6 | 38 | 778 | 8.59 | 21 | 25 | 23 | 8 | 9 | 6 | 18 | 18 | 17 | 8 | 7 | 6 |
| High | Q03164 | Histone-lysine N-methyltransferase 2A OS=Homo sapiens OX=9606 GN=KMT2A PE=1 SV=5 | 151 | 45 | 41 | 431.5 | 16 | 3969 | 9.09 | | | | 22 | 30 | 24 | | | | 22 | 27 | 24 |
| High | Q5QNW6 | Histone H2B type 2-F OS=Homo sapiens OX=9606 GN=H2BC18 PE=1 SV=3 | 150 | 5 | 2 | 13.9 | 36 | 126 | 10.32 | 22 | 17 | 26 | 12 | 11 | 11 | 4 | 4 | 5 | 5 | 5 | 5 |
| High | Q14247 | Src substrate cortactin OS=Homo sapiens OX=9606 GN=CTTN PE=1 SV=2 | 150 | 20 | 20 | 61.5 | 41 | 550 | 5.4 | 17 | 14 | 15 | 12 | 15 | 15 | 15 | 13 | 12 | 11 | 15 | 14 |
| High | P09874 | Poly [ADP-ribose] polymerase 1 OS=Homo sapiens OX=9606 GN=PARP1 PE=1 SV=4 | 147 | 29 | 29 | 113 | 35 | 1014 | 8.88 | 17 | 16 | 20 | 13 | 14 | 13 | 16 | 16 | 17 | 13 | 14 | 13 |
| High | Q9UJ72 | Annexin A10 OS=Homo sapiens OX=9606 GN=ANXA10 PE=1 SV=3 | 146 | 17 | 17 | 37.3 | 66 | 324 | 5.33 | 17 | 17 | 14 | 8 | 11 | 8 | 14 | 14 | 12 | 7 | 8 | 7 |
| High | Q7L576 | Cytoplasmic FMR1-interacting protein 1 OS=Homo sapiens OX=9606 GN=CYFIP1 PE=1 SV=1 | 146 | 29 | 29 | 145.1 | 27 | 1253 | 6.9 | 20 | 22 | 21 | 9 | 8 | 10 | 17 | 19 | 16 | 9 | 8 | 10 |
| High | P52907 | F-actin-capping protein subunit alpha-1 OS=Homo sapiens OX=9606 GN=CAPZA1 PE=1 SV=3 | 146 | 12 | 10 | 32.9 | 65 | 286 | 5.69 | 15 | 12 | 16 | 5 | 7 | 7 | 9 | 6 | 10 | 4 | 6 | 5 |
| High | P85037 | Forkhead box protein K1 OS=Homo sapiens OX=9606 GN=FOXK1 PE=1 SV=1 | 145 | 21 | 19 | 75.4 | 31 | 733 | 9.32 | 5 | 4 | 3 | 19 | 19 | 19 | 5 | 4 | 3 | 15 | 17 | 14 |
| High | P55265 | Double-stranded RNA-specific adenosine deaminase OS=Homo sapiens OX=9606 GN=ADAR PE=1 SV=4 | 145 | 29 | 29 | 136 | 29 | 1226 | 8.65 | 11 | 12 | 10 | 16 | 17 | 18 | 11 | 12 | 10 | 16 | 15 | 17 |
| High | P41252 | Isoleucine--tRNA ligase, cytoplasmic OS=Homo sapiens OX=9606 GN=IARS1 PE=1 SV=2 | 145 | 30 | 30 | 144.4 | 31 | 1262 | 6.15 | 17 | 14 | 17 | 9 | 7 | 9 | 17 | 14 | 16 | 9 | 7 | 9 |
| High | P10599 | Thioredoxin OS=Homo sapiens OX=9606 GN=TXN PE=1 SV=3 | 144 | 5 | 5 | 11.7 | 50 | 105 | 4.92 | 12 | 16 | 17 | 10 | 23 | 9 | 4 | 5 | 5 | 4 | 4 | 3 |
| High | P37802 | Transgelin-2 OS=Homo sapiens OX=9606 GN=TAGLN2 PE=1 SV=3 | 144 | 13 | 13 | 22.4 | 74 | 199 | 8.25 | 16 | 12 | 12 | 16 | 15 | 12 | 12 | 10 | 9 | 9 | 9 | 8 |
| High | P56470 | Galectin-4 OS=Homo sapiens OX=9606 GN=LGALS4 PE=1 SV=1 | 144 | 11 | 11 | 35.9 | 35 | 323 | 9.16 | 17 | 17 | 20 | 6 | 5 | 8 | 9 | 9 | 10 | 5 | 3 | 5 |
| High | P49915 | GMP synthase [glutamine-hydrolyzing] OS=Homo sapiens OX=9606 GN=GMPS PE=1 SV=1 | 143 | 26 | 26 | 76.7 | 48 | 693 | 6.87 | 20 | 18 | 17 | 11 | 12 | 8 | 17 | 16 | 14 | 10 | 10 | 8 |
| High | P33992 | DNA replication licensing factor MCM5 OS=Homo sapiens OX=9606 GN=MCM5 PE=1 SV=5 | 142 | 28 | 28 | 82.2 | 44 | 734 | 8.37 | 14 | 9 | 15 | 9 | 18 | 10 | 14 | 8 | 15 | 9 | 16 | 10 |
| High | P06576 | ATP synthase subunit beta, mitochondrial OS=Homo sapiens OX=9606 GN=ATP5F1B PE=1 SV=3 | 141 | 22 | 22 | 56.5 | 62 | 529 | 5.4 | 22 | 23 | 25 | 12 | 11 | 13 | 17 | 18 | 20 | 10 | 9 | 11 |
| High | P63104 | 14-3-3 protein zeta/delta OS=Homo sapiens OX=9606 GN=YWHAZ PE=1 SV=1 | 140 | 14 | 11 | 27.7 | 62 | 245 | 4.79 | 15 | 15 | 16 | 7 | 15 | 12 | 11 | 11 | 10 | 7 | 9 | 7 |
| High | Q16778 | Histone H2B type 2-E OS=Homo sapiens OX=9606 GN=H2BC21 PE=1 SV=3 | 140 | 5 | 2 | 13.9 | 36 | 126 | 10.32 | 20 | 16 | 26 | 11 | 10 | 10 | 4 | 4 | 5 | 5 | 5 | 5 |
| High | Q6IFX2 | SWISS-PROT:Q6IFX2 Tax_Id=10090 Gene_Symbol=Krt42 Keratin, type I cytoskeletal 42 | 140 | 11 | 1 | 50.1 | 18 | 452 | 5.16 | 27 | 27 | 32 | 8 | 14 | 7 | 9 | 9 | 11 | 4 | 6 | 4 |
| High | P53618 | Coatomer subunit beta OS=Homo sapiens OX=9606 GN=COPB1 PE=1 SV=3 | 138 | 28 | 28 | 107.1 | 41 | 953 | 6.05 | 26 | 27 | 25 | 6 | 7 | 2 | 22 | 22 | 21 | 5 | 6 | 2 |
| High | P61247 | 40S ribosomal protein S3a OS=Homo sapiens OX=9606 GN=RPS3A PE=1 SV=2 | 137 | 16 | 16 | 29.9 | 53 | 264 | 9.73 | 11 | 9 | 11 | 13 | 15 | 15 | 9 | 7 | 10 | 12 | 12 | 13 |
| High | Q92560 | Ubiquitin carboxyl-terminal hydrolase BAP1 OS=Homo sapiens OX=9606 GN=BAP1 PE=1 SV=2 | 137 | 25 | 25 | 80.3 | 48 | 729 | 6.84 | | | | 15 | 23 | 17 | | | | 13 | 15 | 13 |
| High | P62258 | 14-3-3 protein epsilon OS=Homo sapiens OX=9606 GN=YWHAE PE=1 SV=1 | 137 | 16 | 13 | 29.2 | 59 | 255 | 4.74 | 14 | 16 | 18 | 10 | 12 | 13 | 12 | 12 | 12 | 10 | 10 | 11 |
| High | O14974 | Protein phosphatase 1 regulatory subunit 12A OS=Homo sapiens OX=9606 GN=PPP1R12A PE=1 SV=1 | 136 | 22 | 22 | 115.2 | 26 | 1030 | 5.4 | 14 | 11 | 19 | 12 | 11 | 11 | 12 | 10 | 16 | 10 | 11 | 11 |
| High | P27348 | 14-3-3 protein theta OS=Homo sapiens OX=9606 GN=YWHAQ PE=1 SV=1 | 136 | 12 | 8 | 27.7 | 53 | 245 | 4.78 | 14 | 15 | 16 | 11 | 14 | 14 | 8 | 9 | 8 | 8 | 7 | 8 |
| High | P39023 | 60S ribosomal protein L3 OS=Homo sapiens OX=9606 GN=RPL3 PE=1 SV=2 | 135 | 15 | 15 | 46.1 | 40 | 403 | 10.18 | 13 | 12 | 15 | 7 | 10 | 10 | 10 | 10 | 11 | 7 | 8 | 9 |
| High | P12814 | Alpha-actinin-1 OS=Homo sapiens OX=9606 GN=ACTN1 PE=1 SV=2 | 135 | 23 | 10 | 103 | 32 | 892 | 5.41 | 22 | 29 | 24 | 5 | 7 | 6 | 16 | 20 | 17 | 5 | 6 | 6 |
| High | P04843 | Dolichyl-diphosphooligosaccharide--protein glycosyltransferase subunit 1 OS=Homo sapiens OX=9606 GN=RPN1 PE=1 SV=1 | 135 | 25 | 25 | 68.5 | 52 | 607 | 6.38 | 17 | 22 | 21 | 8 | 9 | 8 | 17 | 21 | 18 | 8 | 8 | 8 |
| High | Q92597 | Protein NDRG1 OS=Homo sapiens OX=9606 GN=NDRG1 PE=1 SV=1 | 135 | 11 | 11 | 42.8 | 54 | 394 | 5.82 | 16 | 20 | 13 | 8 | 11 | 7 | 11 | 11 | 8 | 5 | 6 | 5 |
| High | O94973 | AP-2 complex subunit alpha-2 OS=Homo sapiens OX=9606 GN=AP2A2 PE=1 SV=2 | 134 | 28 | 18 | 103.9 | 40 | 939 | 6.96 | 16 | 17 | 16 | 5 | 7 | 5 | 15 | 16 | 16 | 5 | 7 | 5 |
| High | P25705 | ATP synthase subunit alpha, mitochondrial OS=Homo sapiens OX=9606 GN=ATP5F1A PE=1 SV=1 | 134 | 19 | 19 | 59.7 | 46 | 553 | 9.13 | 22 | 21 | 22 | 7 | 10 | 11 | 17 | 17 | 17 | 7 | 10 | 10 |
| High | P05141 | ADP/ATP translocase 2 OS=Homo sapiens OX=9606 GN=SLC25A5 PE=1 SV=7 | 133 | 16 | 6 | 32.8 | 51 | 298 | 9.69 | 16 | 15 | 15 | 6 | 11 | 8 | 12 | 13 | 10 | 6 | 10 | 6 |
| High | P35080 | Profilin-2 OS=Homo sapiens OX=9606 GN=PFN2 PE=1 SV=3 | 133 | 5 | 5 | 15 | 36 | 140 | 6.99 | 10 | 11 | 15 | 10 | 15 | 10 | 4 | 4 | 4 | 3 | 5 | 4 |
| High | O95789 | Zinc finger MYM-type protein 6 OS=Homo sapiens OX=9606 GN=ZMYM6 PE=1 SV=2 | 133 | 40 | 40 | 148 | 36 | 1325 | 8.22 | | | | | | | | | | | | |
| High | P46013 | Proliferation marker protein Ki-67 OS=Homo sapiens OX=9606 GN=MKI67 PE=1 SV=2 | 132 | 37 | 37 | 358.5 | 17 | 3256 | 9.45 | 8 | 8 | 14 | 12 | 13 | 14 | 8 | 8 | 13 | 12 | 13 | 14 |
| High | O43390 | Heterogeneous nuclear ribonucleoprotein R OS=Homo sapiens OX=9606 GN=HNRNPR PE=1 SV=1 | 131 | 21 | 15 | 70.9 | 39 | 633 | 8.13 | 13 | 14 | 18 | 8 | 12 | 12 | 13 | 14 | 16 | 8 | 12 | 12 |
| High | P38159 | RNA-binding motif protein, X chromosome OS=Homo sapiens OX=9606 GN=RBMX PE=1 SV=3 | 130 | 16 | 4 | 42.3 | 41 | 391 | 10.05 | 13 | 16 | 21 | 10 | 12 | 13 | 11 | 13 | 15 | 10 | 10 | 11 |
| High | P34932 | Heat shock 70 kDa protein 4 OS=Homo sapiens OX=9606 GN=HSPA4 PE=1 SV=4 | 130 | 26 | 25 | 94.3 | 44 | 840 | 5.19 | 10 | 13 | 18 | 8 | 7 | 7 | 10 | 12 | 18 | 7 | 7 | 7 |
| High | P67809 | Y-box-binding protein 1 OS=Homo sapiens OX=9606 GN=YBX1 PE=1 SV=3 | 130 | 9 | 6 | 35.9 | 48 | 324 | 9.88 | 12 | 9 | 14 | 14 | 13 | 14 | 6 | 6 | 7 | 9 | 7 | 9 |
| High | Q92945 | Far upstream element-binding protein 2 OS=Homo sapiens OX=9606 GN=KHSRP PE=1 SV=4 | 130 | 20 | 18 | 73.1 | 40 | 711 | 7.3 | 11 | 11 | 11 | 12 | 16 | 13 | 11 | 10 | 11 | 12 | 14 | 13 |
| High | P43243 | Matrin-3 OS=Homo sapiens OX=9606 GN=MATR3 PE=1 SV=2 | 130 | 25 | 25 | 94.6 | 38 | 847 | 6.25 | 15 | 14 | 16 | 11 | 10 | 9 | 13 | 10 | 14 | 11 | 10 | 9 |
| High | P0DP25 | Calmodulin-3 OS=Homo sapiens OX=9606 GN=CALM3 PE=1 SV=1 | 130 | 9 | 9 | 16.8 | 52 | 149 | 4.22 | 15 | 17 | 15 | 7 | 13 | 6 | 8 | 8 | 7 | 7 | 7 | 6 |
| High | P17812 | CTP synthase 1 OS=Homo sapiens OX=9606 GN=CTPS1 PE=1 SV=2 | 130 | 19 | 18 | 66.6 | 46 | 591 | 6.46 | 18 | 13 | 15 | 12 | 12 | 12 | 17 | 12 | 13 | 11 | 10 | 11 |
| High | P09651 | Heterogeneous nuclear ribonucleoprotein A1 OS=Homo sapiens OX=9606 GN=HNRNPA1 PE=1 SV=5 | 130 | 16 | 14 | 38.7 | 46 | 372 | 9.13 | 21 | 21 | 19 | 9 | 9 | 10 | 15 | 14 | 12 | 6 | 7 | 6 |
| High | O60506 | Heterogeneous nuclear ribonucleoprotein Q OS=Homo sapiens OX=9606 GN=SYNCRIP PE=1 SV=2 | 129 | 20 | 14 | 69.6 | 41 | 623 | 8.59 | 13 | 15 | 16 | 6 | 10 | 11 | 13 | 14 | 14 | 6 | 10 | 10 |
| High | P06703 | Protein S100-A6 OS=Homo sapiens OX=9606 GN=S100A6 PE=1 SV=1 | 129 | 3 | 3 | 10.2 | 19 | 90 | 5.48 | 13 | 9 | 16 | 13 | 13 | 14 | 2 | 2 | 2 | 3 | 2 | 3 |
| High | Q8IXJ9 | Polycomb group protein ASXL1 OS=Homo sapiens OX=9606 GN=ASXL1 PE=1 SV=4 | 128 | 26 | 25 | 165.3 | 23 | 1541 | 6.21 | | | | 21 | 23 | 19 | | | | 20 | 21 | 18 |
| High | P60842 | Eukaryotic initiation factor 4A-I OS=Homo sapiens OX=9606 GN=EIF4A1 PE=1 SV=1 | 128 | 18 | 10 | 46.1 | 54 | 406 | 5.48 | 17 | 16 | 20 | 9 | 13 | 13 | 13 | 14 | 17 | 9 | 11 | 11 |
| High | P35221 | Catenin alpha-1 OS=Homo sapiens OX=9606 GN=CTNNA1 PE=1 SV=1 | 127 | 28 | 28 | 100 | 40 | 906 | 6.29 | 21 | 21 | 17 | 5 | 6 | 7 | 21 | 20 | 17 | 5 | 6 | 7 |
| High | Q03252 | Lamin-B2 OS=Homo sapiens OX=9606 GN=LMNB2 PE=1 SV=4 | 127 | 27 | 24 | 69.9 | 40 | 620 | 5.59 | 23 | 20 | 26 | 7 | 8 | 7 | 16 | 16 | 20 | 6 | 6 | 7 |
| High | P42224 | Signal transducer and activator of transcription 1-alpha/beta OS=Homo sapiens OX=9606 GN=STAT1 PE=1 SV=2 | 126 | 23 | 23 | 87.3 | 36 | 750 | 6.05 | 17 | 14 | 16 | 9 | 17 | 11 | 16 | 13 | 15 | 9 | 14 | 11 |
| High | Q15424 | Scaffold attachment factor B1 OS=Homo sapiens OX=9606 GN=SAFB PE=1 SV=4 | 125 | 19 | 12 | 102.6 | 22 | 915 | 5.47 | 16 | 16 | 19 | 7 | 11 | 10 | 12 | 13 | 15 | 7 | 10 | 9 |
| High | O75694 | Nuclear pore complex protein Nup155 OS=Homo sapiens OX=9606 GN=NUP155 PE=1 SV=1 | 125 | 23 | 23 | 155.1 | 22 | 1391 | 6.16 | 13 | 12 | 17 | 8 | 8 | 6 | 13 | 12 | 14 | 7 | 6 | 6 |
| High | Q15181 | Inorganic pyrophosphatase OS=Homo sapiens OX=9606 GN=PPA1 PE=1 SV=2 | 125 | 16 | 15 | 32.6 | 71 | 289 | 5.86 | 13 | 14 | 16 | 7 | 16 | 10 | 10 | 12 | 13 | 5 | 8 | 7 |
| High | P16144 | Integrin beta-4 OS=Homo sapiens OX=9606 GN=ITGB4 PE=1 SV=5 | 125 | 32 | 32 | 202 | 24 | 1822 | 6.09 | 15 | 13 | 22 | 7 | 6 | 6 | 15 | 13 | 20 | 7 | 6 | 6 |
| High | O60502 | Protein O-GlcNAcase OS=Homo sapiens OX=9606 GN=OGA PE=1 SV=2 | 125 | 32 | 32 | 102.8 | 44 | 916 | 4.91 | 1 | 2 | | 9 | 11 | 10 | 1 | 2 | | 9 | 11 | 10 |
| High | Q16891 | MICOS complex subunit MIC60 OS=Homo sapiens OX=9606 GN=IMMT PE=1 SV=1 | 125 | 25 | 25 | 83.6 | 40 | 758 | 6.48 | 10 | 10 | 14 | 10 | 16 | 11 | 10 | 10 | 13 | 10 | 15 | 11 |
| High | Q6WCQ1 | Myosin phosphatase Rho-interacting protein OS=Homo sapiens OX=9606 GN=MPRIP PE=1 SV=3 | 124 | 37 | 37 | 116.5 | 48 | 1025 | 6.21 | 22 | 25 | 22 | 1 | | 1 | 19 | 22 | 20 | 1 | | 1 |
| High | P36578 | 60S ribosomal protein L4 OS=Homo sapiens OX=9606 GN=RPL4 PE=1 SV=5 | 124 | 16 | 16 | 47.7 | 41 | 427 | 11.06 | 9 | 9 | 11 | 11 | 10 | 13 | 9 | 9 | 11 | 8 | 8 | 9 |
| High | Q9BUJ2 | Heterogeneous nuclear ribonucleoprotein U-like protein 1 OS=Homo sapiens OX=9606 GN=HNRNPUL1 PE=1 SV=2 | 124 | 20 | 20 | 95.7 | 28 | 856 | 6.92 | 18 | 17 | 20 | 7 | 6 | 7 | 17 | 15 | 17 | 7 | 6 | 7 |
| High | Q9UHR4 | Brain-specific angiogenesis inhibitor 1-associated protein 2-like protein 1 OS=Homo sapiens OX=9606 GN=BAIAP2L1 PE=1 SV=2 | 124 | 23 | 23 | 56.8 | 52 | 511 | 8.68 | 12 | 13 | 18 | 9 | 11 | 9 | 12 | 12 | 17 | 9 | 11 | 9 |
| High | P48643 | T-complex protein 1 subunit epsilon OS=Homo sapiens OX=9606 GN=CCT5 PE=1 SV=1 | 124 | 23 | 22 | 59.6 | 54 | 541 | 5.66 | 16 | 15 | 21 | 9 | 11 | 6 | 13 | 13 | 15 | 8 | 10 | 5 |
| High | Q96CW1 | AP-2 complex subunit mu OS=Homo sapiens OX=9606 GN=AP2M1 PE=1 SV=2 | 123 | 18 | 18 | 49.6 | 42 | 435 | 9.54 | 16 | 16 | 15 | 5 | 10 | 7 | 13 | 13 | 12 | 5 | 10 | 7 |
| High | Q9P2N6 | KAT8 regulatory NSL complex subunit 3 OS=Homo sapiens OX=9606 GN=KANSL3 PE=1 SV=2 | 123 | 25 | 25 | 95.9 | 45 | 904 | 9.39 | | | | 19 | 22 | 19 | | | | 17 | 18 | 16 |
| High | Q7Z2W4 | Zinc finger CCCH-type antiviral protein 1 OS=Homo sapiens OX=9606 GN=ZC3HAV1 PE=1 SV=3 | 122 | 23 | 23 | 101.4 | 33 | 902 | 8.4 | 5 | 4 | 5 | 19 | 11 | 19 | 5 | 4 | 5 | 17 | 11 | 17 |
| High | Q9Y2T3 | Guanine deaminase OS=Homo sapiens OX=9606 GN=GDA PE=1 SV=1 | 122 | 19 | 19 | 51 | 52 | 454 | 5.68 | 20 | 15 | 18 | 9 | 7 | 8 | 15 | 12 | 13 | 8 | 6 | 8 |
| High | Q9Y265 | RuvB-like 1 OS=Homo sapiens OX=9606 GN=RUVBL1 PE=1 SV=1 | 122 | 17 | 17 | 50.2 | 49 | 456 | 6.42 | 15 | 14 | 18 | 6 | 8 | 4 | 13 | 13 | 16 | 6 | 7 | 4 |
| High | Q92804 | TATA-binding protein-associated factor 2N OS=Homo sapiens OX=9606 GN=TAF15 PE=1 SV=1 | 122 | 9 | 7 | 61.8 | 22 | 592 | 8.02 | 14 | 18 | 29 | 6 | 16 | 8 | 9 | 9 | 8 | 4 | 7 | 5 |
| High | P11586 | C-1-tetrahydrofolate synthase, cytoplasmic OS=Homo sapiens OX=9606 GN=MTHFD1 PE=1 SV=4 | 122 | 24 | 23 | 101.5 | 30 | 935 | 7.3 | 15 | 14 | 13 | 5 | 16 | 6 | 13 | 13 | 13 | 5 | 15 | 6 |
| High | P27816 | Microtubule-associated protein 4 OS=Homo sapiens OX=9606 GN=MAP4 PE=1 SV=3 | 121 | 28 | 28 | 120.9 | 35 | 1152 | 5.43 | 5 | 9 | 7 | 13 | 13 | 12 | 5 | 9 | 6 | 13 | 13 | 12 |
| High | P54136 | Arginine--tRNA ligase, cytoplasmic OS=Homo sapiens OX=9606 GN=RARS1 PE=1 SV=2 | 121 | 27 | 27 | 75.3 | 46 | 660 | 6.68 | 12 | 13 | 12 | 11 | 10 | 11 | 12 | 11 | 11 | 10 | 10 | 10 |
| High | Q06210 | Glutamine--fructose-6-phosphate aminotransferase [isomerizing] 1 OS=Homo sapiens OX=9606 GN=GFPT1 PE=1 SV=3 | 121 | 22 | 19 | 78.8 | 39 | 699 | 7.11 | 11 | 12 | 13 | 7 | 13 | 11 | 11 | 12 | 12 | 7 | 13 | 10 |
| High | Q9Y5B9 | FACT complex subunit SPT16 OS=Homo sapiens OX=9606 GN=SUPT16H PE=1 SV=1 | 121 | 25 | 25 | 119.8 | 28 | 1047 | 5.66 | 16 | 16 | 13 | 9 | 17 | 12 | 16 | 16 | 13 | 9 | 15 | 12 |
| High | O95994 | Anterior gradient protein 2 homolog OS=Homo sapiens OX=9606 GN=AGR2 PE=1 SV=1 | 120 | 8 | 7 | 20 | 59 | 175 | 9 | 5 | 6 | 5 | 28 | 30 | 30 | 5 | 5 | 4 | 5 | 5 | 4 |
| High | P50990 | T-complex protein 1 subunit theta OS=Homo sapiens OX=9606 GN=CCT8 PE=1 SV=4 | 120 | 21 | 21 | 59.6 | 43 | 548 | 5.6 | 12 | 11 | 15 | 11 | 11 | 11 | 12 | 11 | 15 | 11 | 11 | 11 |
| High | P50995 | Annexin A11 OS=Homo sapiens OX=9606 GN=ANXA11 PE=1 SV=1 | 119 | 17 | 16 | 54.4 | 34 | 505 | 7.65 | 17 | 15 | 17 | 8 | 10 | 7 | 15 | 13 | 14 | 8 | 10 | 7 |
| High | P61964 | WD repeat-containing protein 5 OS=Homo sapiens OX=9606 GN=WDR5 PE=1 SV=1 | 119 | 11 | 11 | 36.6 | 46 | 334 | 8.27 | 3 | 1 | 3 | 14 | 20 | 16 | 3 | 1 | 3 | 8 | 10 | 8 |
| High | Q15366 | Poly(rC)-binding protein 2 OS=Homo sapiens OX=9606 GN=PCBP2 PE=1 SV=1 | 119 | 13 | 8 | 38.6 | 66 | 365 | 6.79 | 15 | 13 | 13 | 11 | 11 | 9 | 12 | 10 | 9 | 6 | 7 | 6 |
| High | O00425 | Insulin-like growth factor 2 mRNA-binding protein 3 OS=Homo sapiens OX=9606 GN=IGF2BP3 PE=1 SV=2 | 119 | 19 | 17 | 63.7 | 42 | 579 | 8.87 | 11 | 9 | 10 | 8 | 11 | 13 | 11 | 9 | 10 | 8 | 10 | 12 |
| High | P49368 | T-complex protein 1 subunit gamma OS=Homo sapiens OX=9606 GN=CCT3 PE=1 SV=4 | 119 | 22 | 22 | 60.5 | 53 | 545 | 6.49 | 12 | 13 | 15 | 10 | 8 | 11 | 10 | 12 | 14 | 10 | 8 | 10 |
| High | P08590 | Myosin light chain 3 OS=Homo sapiens OX=9606 GN=MYL3 PE=1 SV=3 | 118 | 10 | 7 | 21.9 | 51 | 195 | 5.1 | 13 | 13 | 16 | 10 | 10 | 14 | 7 | 7 | 6 | 8 | 5 | 8 |
| High | P35658 | Nuclear pore complex protein Nup214 OS=Homo sapiens OX=9606 GN=NUP214 PE=1 SV=2 | 117 | 29 | 29 | 213.5 | 22 | 2090 | 7.47 | 3 | 3 | 1 | 10 | 10 | 11 | 3 | 3 | 1 | 10 | 10 | 11 |
| High | P31946 | 14-3-3 protein beta/alpha OS=Homo sapiens OX=9606 GN=YWHAB PE=1 SV=3 | 116 | 10 | 5 | 28.1 | 48 | 246 | 4.83 | 13 | 12 | 16 | 11 | 11 | 12 | 8 | 8 | 8 | 8 | 7 | 8 |
| High | Q9NSB2 | SWISS-PROT:Q9NSB2 Keratin, type II cuticular Hb4 (Hair keratin, type II Hb4) - Homo sapiens (Human). | 116 | 4 | 1 | 64.9 | 4 | 600 | 7.74 | 16 | 22 | 32 | 7 | 9 | 8 | 4 | 4 | 3 | 2 | 2 | 2 |
| High | O75533 | Splicing factor 3B subunit 1 OS=Homo sapiens OX=9606 GN=SF3B1 PE=1 SV=3 | 116 | 26 | 26 | 145.7 | 28 | 1304 | 7.09 | 13 | 13 | 10 | 8 | 13 | 9 | 13 | 13 | 10 | 8 | 13 | 9 |
| High | Q9Y2W1 | Thyroid hormone receptor-associated protein 3 OS=Homo sapiens OX=9606 GN=THRAP3 PE=1 SV=2 | 116 | 22 | 21 | 108.6 | 28 | 955 | 10.15 | 11 | 11 | 10 | 10 | 16 | 12 | 11 | 11 | 9 | 10 | 15 | 12 |
| High | P51991 | Heterogeneous nuclear ribonucleoprotein A3 OS=Homo sapiens OX=9606 GN=HNRNPA3 PE=1 SV=2 | 115 | 17 | 16 | 39.6 | 39 | 378 | 9.01 | 17 | 14 | 19 | 5 | 12 | 6 | 13 | 13 | 12 | 4 | 9 | 4 |
| High | P33176 | Kinesin-1 heavy chain OS=Homo sapiens OX=9606 GN=KIF5B PE=1 SV=1 | 115 | 25 | 25 | 109.6 | 30 | 963 | 6.51 | 1 | 4 | 6 | 18 | 16 | 17 | 1 | 4 | 6 | 16 | 16 | 16 |
| High | Q6KB66-1 | SWISS-PROT:Q6KB66-1 Tax_Id=9606 Gene_Symbol=KRT80 Isoform 1 of Keratin, type II cytoskeletal 80 | 115 | 22 | 21 | 50.5 | 49 | 452 | 5.67 | 23 | 15 | 16 | 11 | 10 | 9 | 16 | 14 | 15 | 11 | 10 | 9 |
| High | P31943 | Heterogeneous nuclear ribonucleoprotein H OS=Homo sapiens OX=9606 GN=HNRNPH1 PE=1 SV=4 | 115 | 10 | 4 | 49.2 | 32 | 449 | 6.3 | 15 | 12 | 13 | 7 | 12 | 7 | 8 | 8 | 8 | 6 | 8 | 6 |
| High | O75955 | Flotillin-1 OS=Homo sapiens OX=9606 GN=FLOT1 PE=1 SV=3 | 115 | 23 | 23 | 47.3 | 68 | 427 | 7.49 | 13 | 11 | 12 | 5 | 2 | 3 | 12 | 9 | 12 | 5 | 2 | 3 |
| High | O15020 | Spectrin beta chain, non-erythrocytic 2 OS=Homo sapiens OX=9606 GN=SPTBN2 PE=1 SV=3 | 115 | 29 | 19 | 271.2 | 15 | 2390 | 6.11 | 19 | 20 | 20 | 4 | 2 | 2 | 15 | 16 | 19 | 4 | 2 | 2 |
| High | P27635 | 60S ribosomal protein L10 OS=Homo sapiens OX=9606 GN=RPL10 PE=1 SV=5 | 115 | 10 | 10 | 24.6 | 43 | 214 | 10.08 | 9 | 11 | 11 | 11 | 11 | 8 | 7 | 8 | 8 | 6 | 6 | 5 |
| High | P12236 | ADP/ATP translocase 3 OS=Homo sapiens OX=9606 GN=SLC25A6 PE=1 SV=4 | 114 | 16 | 6 | 32.8 | 57 | 298 | 9.74 | 14 | 13 | 15 | 5 | 9 | 5 | 11 | 12 | 11 | 5 | 8 | 4 |
| High | Q92499 | ATP-dependent RNA helicase DDX1 OS=Homo sapiens OX=9606 GN=DDX1 PE=1 SV=2 | 113 | 22 | 22 | 82.4 | 34 | 740 | 7.23 | 14 | 12 | 15 | 7 | 13 | 7 | 14 | 12 | 14 | 7 | 13 | 7 |
| High | P53992 | Protein transport protein Sec24C OS=Homo sapiens OX=9606 GN=SEC24C PE=1 SV=3 | 113 | 23 | 22 | 118.2 | 27 | 1094 | 7.06 | 8 | 10 | 11 | 10 | 14 | 11 | 8 | 10 | 11 | 9 | 11 | 11 |
| High | Q8N1F7 | Nuclear pore complex protein Nup93 OS=Homo sapiens OX=9606 GN=NUP93 PE=1 SV=2 | 113 | 32 | 32 | 93.4 | 44 | 819 | 5.72 | 23 | 20 | 20 | 2 | 1 | 2 | 23 | 19 | 20 | 2 | 1 | 1 |
| High | Q9NSD9 | Phenylalanine--tRNA ligase beta subunit OS=Homo sapiens OX=9606 GN=FARSB PE=1 SV=3 | 112 | 20 | 20 | 66.1 | 33 | 589 | 6.84 | 13 | 14 | 14 | 9 | 12 | 9 | 13 | 13 | 12 | 9 | 12 | 9 |
| High | P62241 | 40S ribosomal protein S8 OS=Homo sapiens OX=9606 GN=RPS8 PE=1 SV=2 | 112 | 10 | 10 | 24.2 | 52 | 208 | 10.32 | 10 | 9 | 10 | 14 | 11 | 13 | 9 | 9 | 9 | 10 | 9 | 10 |
| High | P26599 | Polypyrimidine tract-binding protein 1 OS=Homo sapiens OX=9606 GN=PTBP1 PE=1 SV=1 | 111 | 14 | 11 | 57.2 | 42 | 531 | 9.17 | 10 | 10 | 11 | 9 | 10 | 12 | 9 | 9 | 10 | 8 | 6 | 8 |
| High | P52597 | Heterogeneous nuclear ribonucleoprotein F OS=Homo sapiens OX=9606 GN=HNRNPF PE=1 SV=3 | 111 | 13 | 11 | 45.6 | 49 | 415 | 5.58 | 13 | 14 | 12 | 5 | 10 | 8 | 9 | 10 | 7 | 5 | 6 | 6 |
| High | P78371 | T-complex protein 1 subunit beta OS=Homo sapiens OX=9606 GN=CCT2 PE=1 SV=4 | 111 | 18 | 18 | 57.5 | 48 | 535 | 6.46 | 13 | 12 | 13 | 6 | 6 | 8 | 13 | 12 | 11 | 6 | 6 | 8 |
| High | P04083 | Annexin A1 OS=Homo sapiens OX=9606 GN=ANXA1 PE=1 SV=2 | 111 | 18 | 18 | 38.7 | 60 | 346 | 7.02 | 10 | 9 | 13 | 9 | 13 | 9 | 8 | 9 | 11 | 9 | 11 | 9 |
| High | P46109 | Crk-like protein OS=Homo sapiens OX=9606 GN=CRKL PE=1 SV=1 | 110 | 13 | 13 | 33.8 | 65 | 303 | 6.74 | 9 | 9 | 10 | 7 | 10 | 6 | 9 | 9 | 9 | 6 | 7 | 5 |
| High | Q99714 | 3-hydroxyacyl-CoA dehydrogenase type-2 OS=Homo sapiens OX=9606 GN=HSD17B10 PE=1 SV=3 | 110 | 14 | 14 | 26.9 | 74 | 261 | 7.78 | 10 | 11 | 7 | 8 | 9 | 10 | 9 | 9 | 6 | 6 | 5 | 8 |
| High | Q07955 | Serine/arginine-rich splicing factor 1 OS=Homo sapiens OX=9606 GN=SRSF1 PE=1 SV=2 | 110 | 11 | 11 | 27.7 | 38 | 248 | 10.36 | 11 | 9 | 11 | 11 | 14 | 10 | 8 | 7 | 8 | 8 | 9 | 8 |
| High | O60701 | UDP-glucose 6-dehydrogenase OS=Homo sapiens OX=9606 GN=UGDH PE=1 SV=1 | 110 | 19 | 19 | 55 | 54 | 494 | 7.12 | 17 | 14 | 18 | 7 | 8 | 9 | 16 | 13 | 16 | 7 | 8 | 8 |
| High | Q99497 | Parkinson disease protein 7 OS=Homo sapiens OX=9606 GN=PARK7 PE=1 SV=2 | 110 | 10 | 10 | 19.9 | 58 | 189 | 6.79 | 12 | 14 | 11 | 7 | 14 | 8 | 8 | 8 | 7 | 5 | 7 | 5 |
| High | Q12929 | Epidermal growth factor receptor kinase substrate 8 OS=Homo sapiens OX=9606 GN=EPS8 PE=1 SV=1 | 109 | 24 | 24 | 91.8 | 42 | 822 | 7.5 | 13 | 14 | 15 | 1 | 2 | 1 | 13 | 13 | 14 | 1 | 2 | 1 |
| High | P47755 | F-actin-capping protein subunit alpha-2 OS=Homo sapiens OX=9606 GN=CAPZA2 PE=1 SV=3 | 109 | 11 | 9 | 32.9 | 65 | 286 | 5.85 | 11 | 12 | 9 | 3 | 6 | 4 | 8 | 7 | 8 | 3 | 5 | 3 |
| High | Q93009 | Ubiquitin carboxyl-terminal hydrolase 7 OS=Homo sapiens OX=9606 GN=USP7 PE=1 SV=2 | 109 | 22 | 22 | 128.2 | 24 | 1102 | 5.55 | 9 | 8 | 5 | 12 | 17 | 12 | 9 | 7 | 5 | 12 | 16 | 12 |
| High | Q9H6S3 | Epidermal growth factor receptor kinase substrate 8-like protein 2 OS=Homo sapiens OX=9606 GN=EPS8L2 PE=1 SV=2 | 108 | 21 | 21 | 80.6 | 37 | 715 | 6.84 | 11 | 11 | 11 | 7 | 13 | 10 | 11 | 11 | 11 | 7 | 13 | 10 |
| High | Q12931 | Heat shock protein 75 kDa, mitochondrial OS=Homo sapiens OX=9606 GN=TRAP1 PE=1 SV=3 | 107 | 19 | 18 | 80.1 | 37 | 704 | 8.21 | 7 | 9 | 14 | 10 | 7 | 7 | 7 | 8 | 10 | 9 | 6 | 6 |
| High | O60437 | Periplakin OS=Homo sapiens OX=9606 GN=PPL PE=1 SV=4 | 107 | 36 | 36 | 204.6 | 24 | 1756 | 5.6 | 17 | 22 | 28 | 6 | 5 | 3 | 17 | 21 | 26 | 6 | 5 | 3 |
| High | O00443 | Phosphatidylinositol 4-phosphate 3-kinase C2 domain-containing subunit alpha OS=Homo sapiens OX=9606 GN=PIK3C2A PE=1 SV | 106 | 35 | 35 | 190.6 | 25 | 1686 | 8.02 | 8 | 12 | 12 | 3 | 2 | 3 | 8 | 12 | 12 | 3 | 2 | 3 |
| High | P55060 | Exportin-2 OS=Homo sapiens OX=9606 GN=CSE1L PE=1 SV=3 | 106 | 17 | 17 | 110.3 | 20 | 971 | 5.77 | 10 | 12 | 15 | 9 | 10 | 10 | 9 | 10 | 14 | 8 | 10 | 10 |
| High | Q96E39 | RNA binding motif protein, X-linked-like-1 OS=Homo sapiens OX=9606 GN=RBMXL1 PE=1 SV=1 | 106 | 13 | 1 | 42.1 | 34 | 390 | 9.89 | 10 | 12 | 19 | 7 | 10 | 11 | 8 | 10 | 13 | 7 | 9 | 9 |
| High | O60716 | Catenin delta-1 OS=Homo sapiens OX=9606 GN=CTNND1 PE=1 SV=1 | 105 | 21 | 21 | 108.1 | 31 | 968 | 6.23 | 13 | 16 | 15 | 7 | 3 | 6 | 13 | 16 | 14 | 7 | 3 | 6 |
| High | P62280 | 40S ribosomal protein S11 OS=Homo sapiens OX=9606 GN=RPS11 PE=1 SV=3 | 105 | 11 | 11 | 18.4 | 52 | 158 | 10.3 | 11 | 12 | 13 | 8 | 11 | 9 | 9 | 10 | 11 | 8 | 7 | 8 |
| High | Q13501 | Sequestosome-1 OS=Homo sapiens OX=9606 GN=SQSTM1 PE=1 SV=1 | 105 | 12 | 12 | 47.7 | 58 | 440 | 5.22 | 4 | 5 | 4 | 18 | 26 | 22 | 4 | 5 | 4 | 10 | 11 | 10 |
| High | Q9UGI8 | Testin OS=Homo sapiens OX=9606 GN=TES PE=1 SV=1 | 104 | 15 | 15 | 48 | 41 | 421 | 7.68 | 11 | 12 | 13 | 8 | 11 | 8 | 10 | 11 | 10 | 8 | 10 | 8 |
| High | P35237 | Serpin B6 OS=Homo sapiens OX=9606 GN=SERPINB6 PE=1 SV=3 | 104 | 13 | 12 | 42.6 | 47 | 376 | 5.27 | 8 | 9 | 16 | 7 | 10 | 6 | 7 | 8 | 13 | 6 | 7 | 6 |
| High | Q9Y608 | Leucine-rich repeat flightless-interacting protein 2 OS=Homo sapiens OX=9606 GN=LRRFIP2 PE=1 SV=1 | 104 | 15 | 14 | 82.1 | 25 | 721 | 6.95 | 13 | 13 | 19 | 6 | 7 | 9 | 10 | 10 | 15 | 6 | 6 | 9 |
| High | P61981 | 14-3-3 protein gamma OS=Homo sapiens OX=9606 GN=YWHAG PE=1 SV=2 | 104 | 9 | 4 | 28.3 | 38 | 247 | 4.89 | 10 | 11 | 15 | 10 | 11 | 13 | 7 | 8 | 7 | 6 | 6 | 7 |
| High | P41240 | Tyrosine-protein kinase CSK OS=Homo sapiens OX=9606 GN=CSK PE=1 SV=1 | 104 | 16 | 15 | 50.7 | 41 | 450 | 7.06 | 9 | 11 | 10 | 10 | 11 | 13 | 9 | 10 | 7 | 10 | 12 | 10 |
| High | P10515 | Dihydrolipoyllysine-residue acetyltransferase component of pyruvate dehydrogenase complex, mitochondrial OS=Homo sapiens ( | 104 | 15 | 15 | 69 | 28 | 647 | 7.84 | 4 | 4 | 7 | 9 | 10 | 9 | 4 | 4 | 7 | 6 | 8 | 7 |
| High | Q92621 | Nuclear pore complex protein Nup205 OS=Homo sapiens OX=9606 GN=NUP205 PE=1 SV=3 | 103 | 34 | 34 | 227.8 | 19 | 2012 | 6.19 | 23 | 25 | 27 | 2 | 2 | 1 | 23 | 25 | 25 | 2 | 2 | 1 |
| High | Q10567 | AP-1 complex subunit beta-1 OS=Homo sapiens OX=9606 GN=AP1B1 PE=1 SV=3 | 103 | 22 | 7 | 104.5 | 25 | 949 | 5.06 | 7 | 10 | 14 | 6 | 5 | 8 | 7 | 10 | 13 | 6 | 6 | 8 |
| High | Q7Z794 | Keratin, type II cytoskeletal 1b OS=Homo sapiens OX=9606 GN=KRT77 PE=1 SV=3 | 102 | 5 | 1 | 61.9 | 6 | 578 | 5.99 | 13 | 18 | 31 | 6 | 8 | 4 | 4 | 4 | 4 | 3 | 2 | 3 |
| High | Q9ULA0 | Aspartyl aminopeptidase OS=Homo sapiens OX=9606 GN=DNPEP PE=1 SV=2 | 102 | 14 | 14 | 53.4 | 33 | 485 | 7.58 | 10 | 10 | 10 | 9 | 11 | 9 | 9 | 8 | 10 | 7 | 8 | 8 |
| High | P30101 | Protein disulfide-isomerase A3 OS=Homo sapiens OX=9606 GN=PDIA3 PE=1 SV=4 | 102 | 20 | 20 | 56.7 | 48 | 505 | 6.35 | 10 | 12 | 16 | 6 | 8 | 9 | 9 | 12 | 14 | 6 | 8 | 9 |
| High | Q02878 | 60S ribosomal protein L6 OS=Homo sapiens OX=9606 GN=RPL6 PE=1 SV=3 | 101 | 12 | 12 | 32.7 | 35 | 288 | 10.58 | 9 | 9 | 8 | 11 | 12 | 11 | 9 | 9 | 7 | 10 | 10 | 10 |
| High | P13797 | Plastin-3 OS=Homo sapiens OX=9606 GN=PLS3 PE=1 SV=4 | 101 | 24 | 16 | 70.8 | 47 | 630 | 5.6 | 8 | 12 | 10 | 11 | 11 | 13 | 7 | 11 | 10 | 11 | 11 | 13 |
| High | Q69YQ0 | Cytospin-A OS=Homo sapiens OX=9606 GN=SPECC1L PE=1 SV=3 | 101 | 27 | 27 | 124.5 | 29 | 1117 | 5.76 | 17 | 22 | 19 | | 1 | | 17 | 21 | 18 | | 1 | |
| High | P61106 | Ras-related protein Rab-14 OS=Homo sapiens OX=9606 GN=RAB14 PE=1 SV=4 | 100 | 14 | 13 | 23.9 | 78 | 215 | 6.21 | 10 | 10 | 9 | 7 | 12 | 7 | 7 | 7 | 7 | 7 | 9 | 7 |
| High | P07744 | SWISS-PROT:P07744 Tax_Id=10090 Gene_Symbol=Krt4 Keratin, type II cytoskeletal 4 | 100 | 5 | 1 | 56.2 | 6 | 525 | 8.15 | 13 | 18 | 30 | 5 | 6 | 7 | 3 | 3 | 4 | 2 | 3 | 2 |
| High | Q13557 | Calcium/calmodulin-dependent protein kinase type II subunit delta OS=Homo sapiens OX=9606 GN=CAMK2D PE=1 SV=3 | 100 | 14 | 11 | 56.3 | 32 | 499 | 7.25 | 10 | 11 | 11 | 8 | 9 | 7 | 10 | 11 | 11 | 8 | 9 | 7 |
| High | Q8WUM4 | Programmed cell death 6-interacting protein OS=Homo sapiens OX=9606 GN=PDCD6IP PE=1 SV=1 | 100 | 25 | 25 | 96 | 32 | 868 | 6.52 | 9 | 11 | 13 | 5 | 3 | 5 | 9 | 11 | 13 | 5 | 3 | 5 |
| High | P18124 | 60S ribosomal protein L7 OS=Homo sapiens OX=9606 GN=RPL7 PE=1 SV=1 | 100 | 17 | 17 | 29.2 | 52 | 248 | 10.65 | 6 | 7 | 11 | 9 | 10 | 10 | 5 | 6 | 9 | 9 | 9 | 8 |
| High | P62249 | 40S ribosomal protein S16 OS=Homo sapiens OX=9606 GN=RPS16 PE=1 SV=2 | 100 | 7 | 7 | 16.4 | 44 | 146 | 10.21 | 9 | 10 | 10 | 6 | 9 | 7 | 5 | 6 | 7 | 6 | 7 | 5 |
| High | Q15393 | Splicing factor 3B subunit 3 OS=Homo sapiens OX=9606 GN=SF3B3 PE=1 SV=4 | 99 | 19 | 19 | 135.5 | 18 | 1217 | 5.26 | 2 | 5 | 9 | 14 | 16 | 14 | 2 | 5 | 8 | 13 | 14 | 13 |
| High | P04899 | Guanine nucleotide-binding protein G(i) subunit alpha-2 OS=Homo sapiens OX=9606 GN=GNAI2 PE=1 SV=3 | 99 | 15 | 9 | 40.4 | 59 | 355 | 5.54 | 11 | 9 | 11 | 7 | 8 | 8 | 9 | 7 | 10 | 6 | 7 | 7 |
| High | Q5SRE5 | Nucleoporin NUP188 OS=Homo sapiens OX=9606 GN=NUP188 PE=1 SV=1 | 99 | 27 | 27 | 195.9 | 19 | 1749 | 6.73 | 5 | 7 | 6 | 5 | 11 | 6 | 5 | 7 | 6 | 5 | 11 | 5 |
| High | P46087 | Probable 28S rRNA (cytosine(4447)-C(5))-methyltransferase OS=Homo sapiens OX=9606 GN=NOP2 PE=1 SV=2 | 99 | 19 | 19 | 89.2 | 29 | 812 | 9.23 | 6 | 3 | 5 | 13 | 16 | 14 | 6 | 3 | 4 | 13 | 16 | 14 |
| High | P62826 | GTP-binding nuclear protein Ran OS=Homo sapiens OX=9606 GN=RAN PE=1 SV=3 | 99 | 9 | 9 | 24.4 | 35 | 216 | 7.49 | 7 | 7 | 6 | 5 | 5 | 5 | 6 | 6 | 5 | 5 | 5 | 5 |
| High | P55795 | Heterogeneous nuclear ribonucleoprotein H2 OS=Homo sapiens OX=9606 GN=HNRNPH2 PE=1 SV=1 | 99 | 11 | 6 | 49.2 | 35 | 449 | 6.3 | 13 | 11 | 10 | 7 | 9 | 6 | 8 | 7 | 7 | 6 | 8 | 6 |
| High | P45974 | Ubiquitin carboxyl-terminal hydrolase 5 OS=Homo sapiens OX=9606 GN=USP5 PE=1 SV=2 | 98 | 16 | 16 | 95.7 | 26 | 858 | 5.03 | 10 | 9 | 13 | 10 | 9 | 10 | 8 | 9 | 10 | 10 | 7 | 10 |
| High | P50991 | T-complex protein 1 subunit delta OS=Homo sapiens OX=9606 GN=CCT4 PE=1 SV=4 | 98 | 18 | 17 | 57.9 | 42 | 539 | 7.83 | 7 | 10 | 13 | 8 | 7 | 10 | 7 | 9 | 11 | 8 | 7 | 8 |
| High | Q13200 | 26S proteasome non-ATPase regulatory subunit 2 OS=Homo sapiens OX=9606 GN=PSMD2 PE=1 SV=3 | 98 | 22 | 22 | 100.1 | 28 | 908 | 5.2 | 3 | 2 | 5 | 15 | 16 | 16 | 3 | 2 | 5 | 15 | 15 | 16 |

| | | | | | | | | | | | | | | | | | | | | | |
|---|---|---|---|---|---|---|---|---|---|---|---|---|---|---|---|---|---|---|---|---|---|
| High | Q9UHX1 | Poly(U)-binding-splicing factor PUF60 OS=Homo sapiens OX=9606 GN=PUF60 PE=1 SV=1 | 98 | 14 | 14 | 59.8 | 35 | 559 | 5.29 | 8 | 7 | 9 | 9 | 11 | 10 | 8 | 7 | 9 | 8 | 10 | 10 |
| High | O43684 | Mitotic checkpoint protein BUB3 OS=Homo sapiens OX=9606 GN=BUB3 PE=1 SV=1 | 97 | 11 | 11 | 37.1 | 35 | 328 | 6.84 | 5 | 5 | 4 | 15 | 18 | 13 | 5 | 5 | 4 | 11 | 11 | 9 |
| High | Q9Y3I0 | RNA-splicing ligase RtcB homolog OS=Homo sapiens OX=9606 GN=RTCB PE=1 SV=1 | 97 | 19 | 19 | 55.2 | 45 | 505 | 7.23 | 12 | 10 | 15 | 6 | 9 | 6 | 12 | 9 | 13 | 6 | 9 | 6 |
| High | P33993 | DNA replication licensing factor MCM7 OS=Homo sapiens OX=9606 GN=MCM7 PE=1 SV=4 | 97 | 26 | 26 | 81.3 | 45 | 719 | 6.46 | 20 | 20 | 15 | 3 | 6 | 4 | 17 | 19 | 14 | 3 | 6 | 4 |
| High | P27694 | Replication protein A 70 kDa DNA-binding subunit OS=Homo sapiens OX=9606 GN=RPA1 PE=1 SV=2 | 97 | 25 | 25 | 68.1 | 44 | 616 | 7.21 | 20 | 18 | 22 | 6 | 6 | 6 | 19 | 18 | 21 | 6 | 6 | 6 |
| High | Q13310 | Polyadenylate-binding protein 4 OS=Homo sapiens OX=9606 GN=PABPC4 PE=1 SV=1 | 96 | 17 | 11 | 70.7 | 29 | 644 | 9.26 | 7 | 7 | 8 | 8 | 12 | 11 | 7 | 7 | 8 | 8 | 11 | 10 |
| High | P04406 | Glyceraldehyde-3-phosphate dehydrogenase OS=Homo sapiens OX=9606 GN=GAPDH PE=1 SV=3 | 96 | 11 | 11 | 36 | 43 | 335 | 8.46 | 13 | 14 | 17 | 5 | 5 | 5 | 9 | 10 | 10 | 5 | 4 | 5 |
| High | P14866 | Heterogeneous nuclear ribonucleoprotein L OS=Homo sapiens OX=9606 GN=HNRNPL PE=1 SV=2 | 96 | 18 | 17 | 64.1 | 48 | 589 | 8.22 | 8 | 9 | 10 | 5 | 15 | 8 | 7 | 9 | 10 | 5 | 14 | 8 |
| High | P84243 | Histone H3.3 OS=Homo sapiens OX=9606 GN=H3-3A PE=1 SV=2 | 95 | 3 | 1 | 15.3 | 17 | 136 | 11.27 | 9 | 9 | 17 | 12 | 7 | 14 | 2 | 2 | 3 | 2 | 3 | 2 |
| High | P50402 | Emerin OS=Homo sapiens OX=9606 GN=EMD PE=1 SV=1 | 95 | 13 | 13 | 29 | 73 | 254 | 5.5 | 16 | 17 | 10 | 5 | 6 | 5 | 12 | 13 | 10 | 4 | 6 | 4 |
| High | P07237 | Protein disulfide-isomerase OS=Homo sapiens OX=9606 GN=P4HB PE=1 SV=3 | 95 | 20 | 20 | 57.1 | 42 | 508 | 4.87 | 10 | 10 | 14 | 3 | 7 | 3 | 8 | 9 | 10 | 3 | 7 | 3 |
| High | P18621 | 60S ribosomal protein L17 OS=Homo sapiens OX=9606 GN=RPL17 PE=1 SV=3 | 95 | 7 | 7 | 21.4 | 42 | 184 | 10.17 | 8 | 9 | 11 | 6 | 6 | 5 | 6 | 6 | 5 | 4 | 4 | 3 |
| High | P00338 | L-lactate dehydrogenase A chain OS=Homo sapiens OX=9606 GN=LDHA PE=1 SV=2 | 95 | 14 | 13 | 36.7 | 52 | 332 | 8.27 | 14 | 16 | 16 | 4 | 8 | 4 | 10 | 12 | 11 | 4 | 8 | 3 |
| High | Q9UJS0 | Calcium-binding mitochondrial carrier protein Aralar2 OS=Homo sapiens OX=9606 GN=SLC25A13 PE=1 SV=2 | 95 | 20 | 15 | 74.1 | 38 | 675 | 8.62 | 13 | 15 | 19 | 3 | 7 | 6 | 13 | 15 | 17 | 3 | 7 | 6 |
| High | Q13428 | Treacle protein OS=Homo sapiens OX=9606 GN=TCOF1 PE=1 SV=3 | 95 | 20 | 20 | 152 | 16 | 1488 | 9.04 | 5 | 6 | 7 | 11 | 10 | 10 | 5 | 5 | 7 | 11 | 9 | 10 |
| High | P05388 | 60S acidic ribosomal protein P0 OS=Homo sapiens OX=9606 GN=RPLP0 PE=1 SV=1 | 94 | 8 | 8 | 34.3 | 28 | 317 | 5.97 | 8 | 8 | 9 | 10 | 7 | 11 | 7 | 7 | 7 | 7 | 6 | 7 |
| High | Q7RTS7 | SWISS-PROT:Q7RTS7 Tax_Id=9606 Gene_Symbol=KRT74 Keratin, type II cytoskeletal 74 | 94 | 5 | 1 | 57.8 | 7 | 529 | 7.71 | 12 | 17 | 30 | 5 | 6 | 7 | 3 | 3 | 5 | 2 | 3 | 2 |
| High | P07947 | Tyrosine-protein kinase Yes OS=Homo sapiens OX=9606 GN=YES1 PE=1 SV=3 | 94 | 18 | 10 | 60.8 | 39 | 543 | 6.74 | 8 | 9 | 10 | 8 | 6 | 8 | 7 | 7 | 7 | 7 | 5 | 8 |
| High | Q9ULV4 | Coronin-1C OS=Homo sapiens OX=9606 GN=CORO1C PE=1 SV=1 | 94 | 18 | 17 | 53.2 | 46 | 474 | 7.08 | 16 | 12 | 15 | 4 | 5 | 4 | 13 | 11 | 13 | 4 | 5 | 4 |
| High | P31629 | Transcription factor HIVEP2 OS=Homo sapiens OX=9606 GN=HIVEP2 PE=1 SV=2 | 94 | 36 | 33 | 268.9 | 18 | 2446 | 6.96 | | | | 23 | 27 | 24 | | | | 23 | 27 | 23 |
| High | Q32MZ4 | Leucine-rich repeat flightless-interacting protein 1 OS=Homo sapiens OX=9606 GN=LRRFIP1 PE=1 SV=2 | 94 | 15 | 14 | 89.2 | 27 | 808 | 4.65 | 8 | 7 | 11 | 7 | 10 | 7 | 8 | 7 | 9 | 6 | 8 | 7 |
| High | P09661 | U2 small nuclear ribonucleoprotein A' OS=Homo sapiens OX=9606 GN=SNRPA1 PE=1 SV=2 | 94 | 12 | 12 | 28.4 | 47 | 255 | 8.62 | 3 | 2 | 1 | 13 | 18 | 13 | 3 | 2 | 1 | 10 | 10 | 8 |
| High | P08865 | 40S ribosomal protein SA OS=Homo sapiens OX=9606 GN=RPSA PE=1 SV=4 | 93 | 10 | 10 | 32.8 | 52 | 295 | 4.87 | 11 | 10 | 11 | 6 | 7 | 7 | 9 | 9 | 7 | 5 | 6 | 6 |
| High | Q15437 | Protein transport protein Sec23B OS=Homo sapiens OX=9606 GN=SEC23B PE=1 SV=2 | 93 | 16 | 13 | 86.4 | 26 | 767 | 6.89 | 11 | 11 | 12 | 7 | 7 | 8 | 10 | 11 | 12 | 7 | 7 | 7 |
| High | Q14151 | Scaffold attachment factor B2 OS=Homo sapiens OX=9606 GN=SAFB2 PE=1 SV=1 | 93 | 18 | 11 | 107.4 | 25 | 953 | 6.16 | 13 | 17 | 15 | 4 | 4 | 6 | 11 | 13 | 12 | 4 | 3 | 5 |
| High | P02769 | SWISS-PROT:P02769 (Bos taurus) Bovine serum albumin precursor | 92 | 20 | 16 | 69.2 | 39 | 607 | 6.18 | 5 | 18 | 3 | 10 | 13 | 9 | 5 | 13 | 3 | 9 | 11 | 8 |
| High | Q04917 | 14-3-3 protein eta OS=Homo sapiens OX=9606 GN=YWHAH PE=1 SV=4 | 92 | 12 | 8 | 28.2 | 49 | 246 | 4.84 | 9 | 9 | 12 | 8 | 9 | 9 | 7 | 6 | 6 | 5 | 6 | 6 |
| High | P31040 | Succinate dehydrogenase [ubiquinone] flavoprotein subunit, mitochondrial OS=Homo sapiens OX=9606 GN=SDHA PE=1 SV=2 | 92 | 18 | 18 | 72.6 | 33 | 664 | 7.39 | 11 | 11 | 12 | 4 | 11 | 8 | 11 | 11 | 10 | 4 | 11 | 8 |
| High | O95372 | Acyl-protein thioesterase 2 OS=Homo sapiens OX=9606 GN=LYPLA2 PE=1 SV=1 | 92 | 9 | 9 | 24.7 | 52 | 231 | 7.23 | 11 | 14 | 12 | 3 | 7 | 4 | 7 | 8 | 8 | 3 | 6 | 4 |
| High | O95793 | Double-stranded RNA-binding protein Staufen homolog 1 OS=Homo sapiens OX=9606 GN=STAU1 PE=1 SV=2 | 91 | 19 | 17 | 63.1 | 41 | 577 | 9.44 | 4 | 7 | 8 | 10 | 11 | 9 | 3 | 6 | 7 | 10 | 11 | 9 |
| High | P31947 | 14-3-3 protein sigma OS=Homo sapiens OX=9606 GN=SFN PE=1 SV=1 | 91 | 12 | 8 | 27.8 | 53 | 248 | 4.74 | 7 | 10 | 13 | 7 | 6 | 7 | 5 | 7 | 7 | 6 | 4 | 4 |
| High | Q07666 | KH domain-containing, RNA-binding, signal transduction-associated protein 1 OS=Homo sapiens OX=9606 GN=KHDRBS1 PE=1 SV= | 91 | 7 | 7 | 48.2 | 24 | 443 | 8.66 | 10 | 6 | 11 | 9 | 5 | 11 | 6 | 4 | 7 | 5 | 5 | 5 |
| High | O43795 | Unconventional myosin-Ib OS=Homo sapiens OX=9606 GN=MYO1B PE=1 SV=3 | 91 | 29 | 29 | 131.9 | 30 | 1136 | 9.38 | 10 | 10 | 12 | 2 | 1 | 2 | 10 | 9 | 12 | 2 | 1 | 2 |
| High | Q08J23 | RNA cytosine C(5)-methyltransferase NSUN2 OS=Homo sapiens OX=9606 GN=NSUN2 PE=1 SV=2 | 90 | 22 | 22 | 86.4 | 40 | 767 | 6.77 | 11 | 10 | 11 | 6 | 11 | 9 | 10 | 9 | 8 | 6 | 10 | 9 |
| High | Q9NQX4 | Unconventional myosin-Vc OS=Homo sapiens OX=9606 GN=MYO5C PE=1 SV=2 | 90 | 27 | 23 | 202.7 | 17 | 1742 | 7.71 | 8 | 11 | 11 | 2 | 3 | 3 | 7 | 11 | 9 | 2 | 3 | 3 |
| High | P63279 | SUMO-conjugating enzyme UBC9 OS=Homo sapiens OX=9606 GN=UBE2I PE=1 SV=1 | 90 | 7 | 7 | 18 | 34 | 158 | 8.66 | 5 | 6 | 5 | 5 | 5 | 3 | 4 | 4 | 4 | 3 | 3 | 2 |
| High | Q14254 | Flotillin-2 OS=Homo sapiens OX=9606 GN=FLOT2 PE=1 SV=2 | 90 | 15 | 15 | 47 | 44 | 428 | 5.25 | 12 | 11 | 11 | 2 | 2 | 2 | 11 | 11 | 10 | 2 | 2 | 2 |
| High | P55196 | Afadin OS=Homo sapiens OX=9606 GN=AFDN PE=1 SV=3 | 90 | 30 | 30 | 206.7 | 23 | 1824 | 6.47 | 11 | 14 | 15 | 4 | 5 | 3 | 11 | 14 | 15 | 4 | 5 | 3 |
| High | O00468 | Agrin OS=Homo sapiens OX=9606 GN=AGRN PE=1 SV=6 | 89 | 27 | 27 | 217.2 | 19 | 2068 | 6.39 | 10 | 8 | 4 | 6 | 8 | 4 | 8 | 6 | 4 | 5 | 8 | 3 |
| High | P31930 | Cytochrome b-c1 complex subunit 1, mitochondrial OS=Homo sapiens OX=9606 GN=UQCRC1 PE=1 SV=3 | 89 | 13 | 12 | 52.6 | 40 | 480 | 6.37 | 12 | 11 | 12 | 9 | 9 | 9 | 10 | 11 | 11 | 8 | 8 | 8 |
| High | P56192 | Methionine--tRNA ligase, cytoplasmic OS=Homo sapiens OX=9606 GN=MARS1 PE=1 SV=2 | 88 | 18 | 18 | 101.1 | 25 | 900 | 6.16 | 10 | 9 | 13 | 1 | 4 | 2 | 9 | 8 | 12 | 1 | 4 | 2 |
| High | Q13057 | Bifunctional coenzyme A synthase OS=Homo sapiens OX=9606 GN=COASY PE=1 SV=4 | 88 | 16 | 16 | 62.3 | 35 | 564 | 6.99 | 7 | 9 | 12 | 6 | 8 | 7 | 7 | 9 | 11 | 6 | 8 | 7 |
| High | Q9HCC0 | Methylcrotonoyl-CoA carboxylase beta chain, mitochondrial OS=Homo sapiens OX=9606 GN=MCCC2 PE=1 SV=1 | 88 | 19 | 19 | 61.3 | 41 | 563 | 7.68 | 8 | 7 | 8 | 8 | 11 | 8 | 7 | 7 | 7 | 7 | 11 | 8 |
| High | P42167 | Lamina-associated polypeptide 2, isoforms beta/gamma OS=Homo sapiens OX=9606 GN=TMPO PE=1 SV=2 | 88 | 12 | 7 | 50.6 | 41 | 454 | 9.38 | 12 | 12 | 10 | 8 | 7 | 6 | 10 | 9 | 8 | 8 | 7 | 6 |
| High | O60547 | GDP-mannose 4,6 dehydratase OS=Homo sapiens OX=9606 GN=GMDS PE=1 SV=1 | 88 | 14 | 14 | 41.9 | 45 | 372 | 7.31 | 9 | 9 | 10 | 6 | 10 | 7 | 9 | 9 | 9 | 6 | 8 | 7 |
| High | Q9P0K7 | Ankycorbin OS=Homo sapiens OX=9606 GN=RAI14 PE=1 SV=2 | 88 | 28 | 28 | 110 | 36 | 980 | 6.21 | 13 | 18 | 17 | | | | 13 | 18 | 17 | | | |
| High | Q6YN16 | Hydroxysteroid dehydrogenase-like protein 2 OS=Homo sapiens OX=9606 GN=HSDL2 PE=1 SV=1 | 88 | 17 | 17 | 45.4 | 50 | 418 | 7.99 | 11 | 10 | 9 | 4 | 5 | 4 | 9 | 9 | 7 | 4 | 5 | 4 |
| High | P12931 | Proto-oncogene tyrosine-protein kinase Src OS=Homo sapiens OX=9606 GN=SRC PE=1 SV=3 | 88 | 18 | 10 | 59.8 | 41 | 536 | 7.42 | 8 | 9 | 10 | 6 | 6 | 8 | 8 | 9 | 9 | 6 | 6 | 7 |
| High | P55786 | Puromycin-sensitive aminopeptidase OS=Homo sapiens OX=9606 GN=NPEPPS PE=1 SV=2 | 88 | 19 | 19 | 103.2 | 23 | 919 | 5.72 | 13 | 16 | 16 | 5 | 6 | 5 | 13 | 14 | 13 | 5 | 6 | 5 |
| High | P62750 | 60S ribosomal protein L23a OS=Homo sapiens OX=9606 GN=RPL23A PE=1 SV=1 | 88 | 8 | 8 | 17.7 | 33 | 156 | 10.45 | 6 | 8 | 8 | 9 | 12 | 9 | 5 | 7 | 6 | 6 | 5 | 5 |
| High | Q9UPU5 | Ubiquitin carboxyl-terminal hydrolase 24 OS=Homo sapiens OX=9606 GN=USP24 PE=1 SV=3 | 88 | 29 | 29 | 294.2 | 14 | 2620 | 6.14 | 14 | 8 | 14 | 2 | 3 | 3 | 14 | 8 | 14 | 2 | 3 | 3 |
| High | Q99832 | T-complex protein 1 subunit eta OS=Homo sapiens OX=9606 GN=CCT7 PE=1 SV=2 | 87 | 17 | 17 | 59.3 | 38 | 543 | 7.65 | 8 | 9 | 7 | 9 | 11 | 6 | 6 | 8 | 6 | 8 | 11 | 6 |
| High | Q96PK6 | RNA-binding protein 14 OS=Homo sapiens OX=9606 GN=RBM14 PE=1 SV=2 | 87 | 17 | 17 | 69.4 | 30 | 669 | 9.67 | 13 | 11 | 16 | 6 | 7 | 7 | 10 | 8 | 14 | 6 | 6 | 7 |
| High | Q14677 | Clathrin interactor 1 OS=Homo sapiens OX=9606 GN=CLINT1 PE=1 SV=1 | 87 | 14 | 14 | 68.2 | 28 | 625 | 6.42 | 5 | 7 | 8 | 3 | 3 | 2 | 5 | 7 | 8 | 3 | 3 | 2 |
| High | P22234 | Multifunctional protein ADE2 OS=Homo sapiens OX=9606 GN=PAICS PE=1 SV=3 | 87 | 13 | 13 | 47 | 39 | 425 | 7.23 | 7 | 9 | 14 | 7 | 9 | 8 | 5 | 6 | 9 | 7 | 8 | 8 |
| High | Q9HBD1 | Roquin-2 OS=Homo sapiens OX=9606 GN=RC3H2 PE=1 SV=2 | 87 | 23 | 19 | 131.6 | 30 | 1191 | 6.89 | | | | 17 | 16 | 16 | | | | 15 | 14 | 15 |
| High | P05091 | Aldehyde dehydrogenase, mitochondrial OS=Homo sapiens OX=9606 GN=ALDH2 PE=1 SV=2 | 87 | 19 | 16 | 56.3 | 44 | 517 | 7.05 | 11 | 9 | 11 | 6 | 6 | 6 | 7 | 7 | 9 | 6 | 6 | 6 |
| High | Q9Y6K5 | 2'-5'-oligoadenylate synthase 3 OS=Homo sapiens OX=9606 GN=OAS3 PE=1 SV=3 | 87 | 21 | 21 | 121.1 | 24 | 1087 | 8.4 | 6 | 5 | 3 | 12 | 14 | 10 | 6 | 5 | 3 | 12 | 14 | 10 |
| High | Q86VP6 | Cullin-associated NEDD8-dissociated protein 1 OS=Homo sapiens OX=9606 GN=CAND1 PE=1 SV=2 | 86 | 22 | 22 | 136.3 | 20 | 1230 | 5.78 | 8 | 12 | 10 | 6 | 7 | 4 | 8 | 11 | 10 | 6 | 7 | 4 |
| High | Q9P2J5 | Leucine--tRNA ligase, cytoplasmic OS=Homo sapiens OX=9606 GN=LARS1 PE=1 SV=2 | 86 | 24 | 24 | 134.4 | 24 | 1176 | 7.3 | 13 | 13 | 14 | | 5 | 2 | 12 | 12 | 14 | | 5 | 2 |
| High | P13489 | Ribonuclease inhibitor OS=Homo sapiens OX=9606 GN=RNH1 PE=1 SV=2 | 86 | 15 | 15 | 49.9 | 48 | 461 | 4.82 | 11 | 10 | 9 | 6 | 7 | 8 | 11 | 10 | 9 | 5 | 7 | 6 |
| High | Q99959 | Plakophilin-2 OS=Homo sapiens OX=9606 GN=PKP2 PE=1 SV=2 | 86 | 20 | 20 | 97.4 | 30 | 881 | 9.33 | 18 | 17 | 14 | 3 | 5 | 4 | 17 | 16 | 12 | 3 | 5 | 4 |
| High | P08754 | Guanine nucleotide-binding protein G(i) subunit alpha-3 OS=Homo sapiens OX=9606 GN=GNAI3 PE=1 SV=3 | 86 | 12 | 6 | 40.5 | 44 | 354 | 5.69 | 8 | 7 | 10 | 5 | 7 | 6 | 6 | 5 | 9 | 4 | 6 | 5 |
| High | P42704 | Leucine-rich PPR motif-containing protein, mitochondrial OS=Homo sapiens OX=9606 GN=LRPPRC PE=1 SV=3 | 85 | 25 | 25 | 157.8 | 21 | 1394 | 6.13 | 10 | 7 | 8 | 6 | 7 | 5 | 10 | 7 | 8 | 6 | 7 | 5 |
| High | P07195 | L-lactate dehydrogenase B chain OS=Homo sapiens OX=9606 GN=LDHB PE=1 SV=2 | 85 | 13 | 12 | 36.6 | 46 | 334 | 6.05 | 13 | 15 | 14 | 4 | 4 | 2 | 10 | 12 | 8 | 4 | 4 | 2 |
| High | O60841 | Eukaryotic translation initiation factor 5B OS=Homo sapiens OX=9606 GN=EIF5B PE=1 SV=4 | 85 | 16 | 16 | 138.7 | 19 | 1220 | 5.49 | 4 | 4 | 7 | 8 | 9 | 9 | 4 | 4 | 7 | 8 | 9 | 9 |
| High | Q16643 | Drebrin OS=Homo sapiens OX=9606 GN=DBN1 PE=1 SV=4 | 85 | 10 | 10 | 71.4 | 24 | 649 | 4.45 | 13 | 9 | 12 | 6 | 5 | 5 | 10 | 7 | 10 | 5 | 4 | 5 |
| High | Q86UE4 | Protein LYRIC OS=Homo sapiens OX=9606 GN=MTDH PE=1 SV=2 | 84 | 17 | 17 | 63.8 | 38 | 582 | 9.32 | 5 | 5 | 6 | 7 | 10 | 9 | 5 | 4 | 6 | 7 | 10 | 9 |
| High | P30153 | Serine/threonine-protein phosphatase 2A 65 kDa regulatory subunit A alpha isoform OS=Homo sapiens OX=9606 GN=PPP2R1A PE | 84 | 15 | 15 | 65.3 | 32 | 589 | 5.11 | 8 | 8 | 6 | 10 | 6 | 9 | 7 | 8 | 5 | 9 | 5 | 9 |
| High | Q6PKG0 | La-related protein 1 OS=Homo sapiens OX=9606 GN=LARP1 PE=1 SV=2 | 84 | 21 | 21 | 123.4 | 27 | 1096 | 8.82 | 5 | 5 | 8 | 7 | 9 | 8 | 5 | 5 | 8 | 7 | 9 | 8 |
| High | Q5T4S7 | E3 ubiquitin-protein ligase UBR4 OS=Homo sapiens OX=9606 GN=UBR4 PE=1 SV=1 | 84 | 35 | 35 | 573.5 | 9 | 5183 | 6.04 | 9 | 9 | 10 | 3 | 3 | 2 | 9 | 9 | 10 | 3 | 3 | 2 |
| High | Q8NC51 | Plasminogen activator inhibitor 1 RNA-binding protein OS=Homo sapiens OX=9606 GN=SERBP1 PE=1 SV=2 | 83 | 10 | 10 | 44.9 | 29 | 408 | 8.65 | 7 | 8 | 10 | 8 | 10 | 8 | 5 | 6 | 7 | 5 | 8 | 6 |
| High | Q9Y2A7 | Nck-associated protein 1 OS=Homo sapiens OX=9606 GN=NCKAP1 PE=1 SV=1 | 83 | 23 | 23 | 128.7 | 23 | 1128 | 6.62 | 6 | 6 | 9 | 7 | 11 | 6 | 6 | 6 | 7 | 7 | 11 | 6 |
| High | Q9H2U2 | Inorganic pyrophosphatase 2, mitochondrial OS=Homo sapiens OX=9606 GN=PPA2 PE=1 SV=2 | 83 | 13 | 12 | 37.9 | 49 | 334 | 7.39 | 11 | 6 | 9 | 6 | 9 | 6 | 9 | 5 | 8 | 5 | 6 | 6 |
| High | O00567 | Nucleolar protein 56 OS=Homo sapiens OX=9606 GN=NOP56 PE=1 SV=4 | 83 | 16 | 16 | 66 | 31 | 594 | 9.19 | 4 | 6 | 4 | 8 | 7 | 10 | 3 | 4 | 4 | 8 | 7 | 9 |
| High | P05976 | Myosin light chain 1/3, skeletal muscle isoform OS=Homo sapiens OX=9606 GN=MYL1 PE=1 SV=3 | 83 | 4 | 2 | 21.1 | 18 | 194 | 5.03 | 9 | 9 | 13 | 6 | 8 | 10 | 3 | 3 | 3 | 3 | 3 | 4 |
| High | P46977 | Dolichyl-diphosphooligosaccharide--protein glycosyltransferase subunit STT3A OS=Homo sapiens OX=9606 GN=STT3A PE=1 SV=2 | 82 | 14 | 13 | 80.5 | 18 | 705 | 8.07 | 9 | 13 | 11 | 6 | 8 | 5 | 8 | 12 | 10 | 6 | 8 | 5 |
| High | Q9Y446 | Plakophilin-3 OS=Homo sapiens OX=9606 GN=PKP3 PE=1 SV=1 | 82 | 16 | 16 | 87 | 22 | 797 | 9.32 | 10 | 10 | 11 | 3 | 9 | 9 | 10 | 10 | 11 | 3 | 8 | 9 |
| High | O00515 | Ladinin-1 OS=Homo sapiens OX=9606 GN=LAD1 PE=1 SV=2 | 82 | 15 | 15 | 57.1 | 32 | 517 | 9.67 | 7 | 4 | 7 | 5 | 7 | 6 | 7 | 4 | 6 | 5 | 7 | 6 |
| High | P33991 | DNA replication licensing factor MCM4 OS=Homo sapiens OX=9606 GN=MCM4 PE=1 SV=5 | 82 | 19 | 19 | 96.5 | 26 | 863 | 6.74 | 12 | 14 | 18 | 6 | 7 | 7 | 11 | 13 | 15 | 6 | 6 | 7 |
| High | Q9BYB4 | Guanine nucleotide-binding protein subunit beta-like protein 1 OS=Homo sapiens OX=9606 GN=GNB1L PE=1 SV=2 | 82 | 13 | 13 | 35.6 | 46 | 327 | 7.97 | 9 | 6 | 7 | 8 | 7 | 9 | 7 | 6 | 7 | 6 | 5 | 7 |
| High | A8MW92 | PHD finger protein 20-like protein 1 OS=Homo sapiens OX=9606 GN=PHF20L1 PE=1 SV=2 | 81 | 19 | 19 | 114.9 | 23 | 1017 | 6.83 | | | | 11 | 17 | 10 | | | | 11 | 14 | 10 |
| High | P62140 | Serine/threonine-protein phosphatase PP1-beta catalytic subunit OS=Homo sapiens OX=9606 GN=PPP1CB PE=1 SV=3 | 81 | 9 | 3 | 37.2 | 36 | 327 | 6.19 | 12 | 9 | 7 | 6 | 5 | 5 | 7 | 7 | 6 | 6 | 5 | 5 |
| High | Q14839 | Chromodomain-helicase-DNA-binding protein 4 OS=Homo sapiens OX=9606 GN=CHD4 PE=1 SV=2 | 81 | 22 | 18 | 217.9 | 15 | 1912 | 5.86 | 4 | 4 | 8 | 3 | 7 | 5 | 4 | 4 | 8 | 3 | 7 | 5 |
| High | Q5T5P2 | Sickle tail protein homolog OS=Homo sapiens OX=9606 GN=KIAA1217 PE=1 SV=2 | 80 | 28 | 28 | 214 | 17 | 1943 | 7.06 | 5 | 8 | 13 | 1 | 4 | 2 | 5 | 8 | 13 | 1 | 4 | 2 |
| High | P29401 | Transketolase OS=Homo sapiens OX=9606 GN=TKT PE=1 SV=3 | 80 | 16 | 16 | 67.8 | 33 | 623 | 7.66 | 14 | 12 | 10 | 7 | 13 | 8 | 14 | 10 | 10 | 6 | 8 | 7 |
| High | Q8TE68 | Epidermal growth factor receptor kinase substrate 8-like protein 1 OS=Homo sapiens OX=9606 GN=EPS8L1 PE=1 SV=3 | 80 | 16 | 16 | 80.2 | 25 | 723 | 6.04 | 8 | 9 | 6 | 4 | 8 | 9 | 8 | 9 | 6 | 4 | 7 | 9 |
| High | Q96TA1 | Protein Niban 2 OS=Homo sapiens OX=9606 GN=NIBAN2 PE=1 SV=3 | 80 | 16 | 16 | 84.1 | 27 | 746 | 6.19 | 11 | 8 | 10 | 7 | 7 | 8 | 11 | 8 | 10 | 7 | 6 | 8 |
| High | Q3SX14 | TREMBL:Q3SX14 (Bos taurus) Similar to Gelsolin | 80 | 14 | 1 | 85.6 | 18 | 781 | 6.21 | 11 | 11 | 13 | 3 | 9 | 5 | 9 | 9 | 9 | 3 | 6 | 5 |
| High | Q99816 | Tumor susceptibility gene 101 protein OS=Homo sapiens OX=9606 GN=TSG101 PE=1 SV=2 | 80 | 13 | 13 | 43.9 | 33 | 390 | 6.46 | 6 | 8 | 9 | 6 | 8 | 8 | 6 | 8 | 8 | 6 | 8 | 8 |
| High | P61353 | 60S ribosomal protein L27 OS=Homo sapiens OX=9606 GN=RPL27 PE=1 SV=2 | 79 | 6 | 6 | 15.8 | 43 | 136 | 10.56 | 6 | 6 | 8 | 7 | 11 | 9 | 4 | 4 | 4 | 5 | 5 | 6 |
| High | P16402 | Histone H1.3 OS=Homo sapiens OX=9606 GN=H1-3 PE=1 SV=2 | 79 | 5 | 5 | 22.3 | 15 | 221 | 11.02 | 8 | 7 | 8 | 11 | 8 | 8 | 3 | 2 | 2 | 5 | 5 | 3 |
| High | P63096 | Guanine nucleotide-binding protein G(i) subunit alpha-1 OS=Homo sapiens OX=9606 GN=GNAI1 PE=1 SV=2 | 79 | 10 | 4 | 40.3 | 32 | 354 | 5.97 | 8 | 7 | 8 | 6 | 7 | 6 | 6 | 5 | 7 | 5 | 6 | 5 |
| High | O75116 | Rho-associated protein kinase 2 OS=Homo sapiens OX=9606 GN=ROCK2 PE=1 SV=4 | 78 | 24 | 21 | 160.8 | 21 | 1388 | 6.02 | 4 | 6 | 4 | 7 | 12 | 8 | 4 | 6 | 4 | 7 | 12 | 8 |
| High | Q5JWF2 | Guanine nucleotide-binding protein G(s) subunit alpha isoforms XLas OS=Homo sapiens OX=9606 GN=GNAS PE=1 SV=2 | 78 | 11 | 10 | 111 | 14 | 1037 | 5.03 | 10 | 8 | 9 | 5 | 7 | 6 | 10 | 8 | 9 | 5 | 7 | 6 |
| High | P62136 | Serine/threonine-protein phosphatase PP1-alpha catalytic subunit OS=Homo sapiens OX=9606 GN=PPP1CA PE=1 SV=1 | 78 | 11 | 4 | 37.5 | 42 | 330 | 6.33 | 12 | 9 | 8 | 6 | 5 | 5 | 9 | 8 | 7 | 6 | 5 | 5 |
| High | Q9NS86 | LanC-like protein 2 OS=Homo sapiens OX=9606 GN=LANCL2 PE=1 SV=1 | 78 | 11 | 11 | 50.8 | 32 | 450 | 7.43 | 8 | 9 | 7 | 8 | 7 | 7 | 6 | 8 | 7 | 7 | 6 | 6 |
| High | Q8WWM7 | Ataxin-2-like protein OS=Homo sapiens OX=9606 GN=ATXN2L PE=1 SV=2 | 78 | 16 | 16 | 113.3 | 22 | 1075 | 8.59 | 3 | 3 | 6 | 9 | 11 | 7 | 3 | 3 | 5 | 9 | 11 | 7 |
| High | P05455 | Lupus La protein OS=Homo sapiens OX=9606 GN=SSB PE=1 SV=2 | 78 | 16 | 16 | 46.8 | 33 | 408 | 7.12 | 9 | 6 | 8 | 7 | 9 | 5 | 9 | 6 | 8 | 7 | 9 | 5 |
| High | Q6ZRV2 | Protein FAM83H OS=Homo sapiens OX=9606 GN=FAM83H PE=1 SV=3 | 78 | 27 | 27 | 127 | 33 | 1179 | 6.98 | 17 | 21 | 16 | 2 | 3 | 1 | 17 | 20 | 16 | 1 | 3 | 1 |
| High | Q08380 | Galectin-3-binding protein OS=Homo sapiens OX=9606 GN=LGALS3BP PE=1 SV=1 | 78 | 15 | 15 | 65.3 | 36 | 585 | 5.27 | 12 | 9 | 7 | 3 | 6 | 5 | 11 | 9 | 6 | 3 | 5 | 5 |
| High | Q9Y383 | Putative RNA-binding protein Luc7-like 2 OS=Homo sapiens OX=9606 GN=LUC7L2 PE=1 SV=2 | 77 | 12 | 8 | 46.5 | 32 | 392 | 10.01 | 6 | 6 | 5 | 10 | 10 | 9 | 5 | 5 | 5 | 9 | 9 | 8 |
| High | P98160 | Basement membrane-specific heparan sulfate proteoglycan core protein OS=Homo sapiens OX=9606 GN=HSPG2 PE=1 SV=4 | 77 | 23 | 23 | 468.5 | 7 | 4391 | 6.51 | 7 | 9 | 10 | 1 | 4 | 3 | 7 | 9 | 10 | 1 | 4 | 3 |
| High | P07910 | Heterogeneous nuclear ribonucleoproteins C1/C2 OS=Homo sapiens OX=9606 GN=HNRNPC PE=1 SV=4 | 77 | 8 | 3 | 33.7 | 25 | 306 | 5.08 | 9 | 9 | 10 | 7 | 7 | 8 | 8 | 7 | 7 | 6 | 6 | 6 |
| High | P14174 | Macrophage migration inhibitory factor OS=Homo sapiens OX=9606 GN=MIF PE=1 SV=4 | 76 | 3 | 3 | 12.5 | 37 | 115 | 7.88 | 12 | 14 | 7 | 2 | 3 | 2 | 3 | 3 | 2 | 2 | 2 | 2 |
| High | Q8NB78 | Lysine-specific histone demethylase 1B OS=Homo sapiens OX=9606 GN=KDM1B PE=1 SV=3 | 76 | 20 | 20 | 92 | 30 | 822 | 7.87 | | | | 14 | 9 | 14 | | | | 14 | 9 | 14 |
| High | Q05397 | Focal adhesion kinase 1 OS=Homo sapiens OX=9606 GN=PTK2 PE=1 SV=2 | 76 | 19 | 19 | 119.2 | 22 | 1052 | 6.62 | 8 | 7 | 7 | 10 | 7 | 7 | 7 | 7 | 7 | 9 | 7 | 7 |
| High | Q96AB3 | Isochorismatase domain-containing protein 2 OS=Homo sapiens OX=9606 GN=ISOC2 PE=1 SV=1 | 76 | 6 | 6 | 22.3 | 52 | 205 | 7.77 | 9 | 9 | 7 | 2 | 1 | 2 | 6 | 5 | 6 | 2 | 1 | 2 |
| High | P35637 | RNA-binding protein FUS OS=Homo sapiens OX=9606 GN=FUS PE=1 SV=1 | 76 | 8 | 6 | 53.4 | 22 | 526 | 9.36 | 11 | 10 | 12 | 6 | 9 | 6 | 6 | 6 | 7 | 5 | 4 | 4 |
| High | Q9NQW6 | Anillin OS=Homo sapiens OX=9606 GN=ANLN PE=1 SV=2 | 76 | 21 | 21 | 124.1 | 23 | 1124 | 8.07 | 2 | 4 | 8 | 6 | 11 | 5 | 2 | 4 | 8 | 6 | 11 | 5 |
| High | P53396 | ATP-citrate synthase OS=Homo sapiens OX=9606 GN=ACLY PE=1 SV=3 | 76 | 24 | 24 | 120.8 | 26 | 1101 | 7.33 | 6 | 5 | 3 | 5 | 8 | 3 | 6 | 5 | 3 | 5 | 8 | 3 |
| High | P78347 | General transcription factor II-I OS=Homo sapiens OX=9606 GN=GTF2I PE=1 SV=2 | 76 | 24 | 24 | 112.3 | 26 | 998 | 6.39 | 5 | 4 | 6 | 3 | 6 | 4 | 5 | 4 | 6 | 3 | 6 | 4 |
| High | Q96EK4 | THAP domain-containing protein 11 OS=Homo sapiens OX=9606 GN=THAP11 PE=1 SV=2 | 76 | 6 | 6 | 34.4 | 19 | 314 | 8.98 | | | | 11 | 13 | 13 | | | | 5 | 6 | 6 |
| High | P60981 | Destrin OS=Homo sapiens OX=9606 GN=DSTN PE=1 SV=3 | 76 | 7 | 7 | 18.5 | 36 | 165 | 7.85 | 11 | 9 | 9 | 5 | 5 | 4 | 6 | 6 | 6 | 3 | 4 | 3 |
| High | P62906 | 60S ribosomal protein L10a OS=Homo sapiens OX=9606 GN=RPL10A PE=1 SV=2 | 75 | 7 | 7 | 24.8 | 33 | 217 | 9.94 | 6 | 6 | 8 | 8 | 10 | 8 | 6 | 6 | 6 | 6 | 6 | 6 |
| High | A0A075B6 | Probable non-functional immunoglobulin kappa variable 2D-24 OS=Homo sapiens OX=9606 GN=IGKV2D-24 PE=1 SV=1 | 75 | 3 | 2 | 13.1 | 13 | 120 | 8.87 | 8 | 10 | 7 | 6 | 9 | 3 | 2 | 2 | 1 | 1 | 2 | 1 |
| High | Q14573 | Inositol 1,4,5-trisphosphate receptor type 3 OS=Homo sapiens OX=9606 GN=ITPR3 PE=1 SV=2 | 75 | 28 | 28 | 303.9 | 12 | 2671 | 6.48 | 6 | 5 | 5 | 2 | 3 | 2 | 6 | 5 | 5 | 2 | 3 | 2 |
| High | P49756 | RNA-binding protein 25 OS=Homo sapiens OX=9606 GN=RBM25 PE=1 SV=3 | 75 | 16 | 16 | 100.1 | 23 | 843 | 6.32 | 2 | 5 | 5 | 8 | 8 | 7 | 2 | 5 | 5 | 7 | 8 | 7 |
| High | Q9NZB2 | Constitutive coactivator of PPAR-gamma-like protein 1 OS=Homo sapiens OX=9606 GN=FAM120A PE=1 SV=2 | 75 | 18 | 18 | 121.8 | 22 | 1118 | 8.88 | 7 | 4 | 7 | 9 | 6 | 7 | 7 | 4 | 6 | 8 | 6 | 7 |
| High | Q5SSJ5 | Heterochromatin protein 1-binding protein 3 OS=Homo sapiens OX=9606 GN=HP1BP3 PE=1 SV=1 | 75 | 15 | 15 | 61.2 | 28 | 553 | 9.67 | 5 | 5 | 5 | 9 | 8 | 9 | 5 | 5 | 5 | 9 | 8 | 9 |
| High | P35555 | Fibrillin-1 OS=Homo sapiens OX=9606 GN=FBN1 PE=1 SV=4 | 74 | 26 | 25 | 312.1 | 11 | 2871 | 4.93 | | | | 22 | 27 | 24 | | | | 20 | 22 | 21 |
| High | P45880 | Voltage-dependent anion-selective channel protein 2 OS=Homo sapiens OX=9606 GN=VDAC2 PE=1 SV=2 | 74 | 11 | 11 | 31.5 | 50 | 294 | 7.56 | 9 | 8 | 10 | 5 | 7 | 5 | 9 | 8 | 9 | 5 | 6 | 5 |
| High | P55011 | Solute carrier family 12 member 2 OS=Homo sapiens OX=9606 GN=SLC12A2 PE=1 SV=1 | 74 | 18 | 18 | 131.4 | 19 | 1212 | 6.4 | 7 | 11 | 9 | 5 | 6 | 4 | 6 | 9 | 8 | 5 | 6 | 4 |
| High | P40227 | T-complex protein 1 subunit zeta OS=Homo sapiens OX=9606 GN=CCT6A PE=1 SV=3 | 74 | 15 | 11 | 58 | 29 | 531 | 6.68 | 10 | 11 | 9 | 2 | 6 | 4 | 9 | 10 | 9 | 2 | 6 | 4 |
| High | Q9ULC5 | Long-chain-fatty-acid--CoA ligase 5 OS=Homo sapiens OX=9606 GN=ACSL5 PE=1 SV=1 | 74 | 19 | 19 | 75.9 | 36 | 683 | 6.92 | 8 | 10 | 10 | 1 | 5 | 4 | 8 | 9 | 10 | 1 | 5 | 4 |
| High | Q9BS26 | Endoplasmic reticulum resident protein 44 OS=Homo sapiens OX=9606 GN=ERP44 PE=1 SV=1 | 74 | 11 | 11 | 46.9 | 28 | 406 | 5.26 | 4 | 4 | 4 | 9 | 12 | 7 | 4 | 4 | 4 | 7 | 9 | 6 |
| High | Q96KR1 | Zinc finger RNA-binding protein OS=Homo sapiens OX=9606 GN=ZFR PE=1 SV=2 | 74 | 20 | 20 | 116.9 | 24 | 1074 | 9.04 | 1 | 1 | 1 | 9 | 10 | 8 | 1 | 1 | 1 | 9 | 10 | 8 |
| High | P61158 | Actin-related protein 3 OS=Homo sapiens OX=9606 GN=ACTR3 PE=1 SV=3 | 74 | 14 | 12 | 47.3 | 46 | 418 | 5.88 | 11 | 10 | 10 | 4 | 6 | 6 | 11 | 10 | 10 | 4 | 6 | 6 |
| High | P15531 | Nucleoside diphosphate kinase A OS=Homo sapiens OX=9606 GN=NME1 PE=1 SV=1 | 73 | 7 | 3 | 17.1 | 53 | 152 | 6.19 | 7 | 7 | 9 | 9 | 7 | 8 | 5 | 5 | 4 | 4 | 4 | 4 |
| High | P28288 | ATP-binding cassette sub-family D member 3 OS=Homo sapiens OX=9606 GN=ABCD3 PE=1 SV=1 | 73 | 18 | 18 | 75.4 | 30 | 659 | 9.36 | 20 | 19 | 18 | 1 | | 1 | 15 | 16 | 15 | 1 | | 1 |
| High | Q6NXG1 | Epithelial splicing regulatory protein 1 OS=Homo sapiens OX=9606 GN=ESRP1 PE=1 SV=2 | 73 | 13 | 11 | 75.5 | 30 | 681 | 6.68 | 6 | 7 | 6 | 5 | 6 | 6 | 6 | 7 | 5 | 5 | 5 | 6 |
| High | P62913 | 60S ribosomal protein L11 OS=Homo sapiens OX=9606 GN=RPL11 PE=1 SV=2 | 73 | 5 | 5 | 20.2 | 34 | 178 | 9.6 | 11 | 11 | 6 | 7 | 9 | 4 | 5 | 5 | 3 | 5 | 3 | 3 |
| High | P62244 | 40S ribosomal protein S15a OS=Homo sapiens OX=9606 GN=RPS15A PE=1 SV=2 | 73 | 6 | 6 | 14.8 | 42 | 130 | 10.13 | 9 | 10 | 7 | 7 | 8 | 6 | 6 | 6 | 5 | 5 | 4 | 4 |
| High | P50914 | 60S ribosomal protein L14 OS=Homo sapiens OX=9606 GN=RPL14 PE=1 SV=4 | 72 | 5 | 5 | 23.4 | 27 | 215 | 10.93 | 3 | 3 | 3 | 8 | 8 | 9 | 3 | 3 | 2 | 5 | 5 | 5 |
| High | Q5TEC6 | Histone H3-7 OS=Homo sapiens OX=9606 GN=H3-7 PE=1 SV=1 | 72 | 3 | 1 | 15.4 | 20 | 136 | 11.27 | 8 | 7 | 13 | 9 | 6 | 10 | 1 | 1 | 3 | 2 | 3 | 2 |
| High | P54886 | Delta-1-pyrroline-5-carboxylate synthase OS=Homo sapiens OX=9606 GN=ALDH18A1 PE=1 SV=2 | 72 | 16 | 16 | 87.2 | 24 | 795 | 7.12 | 11 | 13 | 13 | 2 | 2 | 2 | 11 | 13 | 12 | 2 | 2 | 2 |
| High | Q14257 | Reticulocalbin-2 OS=Homo sapiens OX=9606 GN=RCN2 PE=1 SV=1 | 72 | 13 | 13 | 36.9 | 45 | 317 | 4.4 | 13 | 14 | 17 | 1 | 2 | 1 | 12 | 11 | 12 | 1 | 2 | 1 |
| High | Q9NUQ9 | CYFIP-related Rac1 interactor B OS=Homo sapiens OX=9606 GN=CYRIB PE=1 SV=1 | 72 | 12 | 12 | 36.7 | 52 | 324 | 6.06 | 10 | 12 | 11 | 2 | 4 | 3 | 8 | 10 | 8 | 2 | 4 | 3 |
| High | P15880 | 40S ribosomal protein S2 OS=Homo sapiens OX=9606 GN=RPS2 PE=1 SV=2 | 72 | 9 | 9 | 31.3 | 34 | 293 | 10.24 | 9 | 8 | 7 | 2 | 5 | 3 | 7 | 6 | 6 | 2 | 5 | 3 |
| High | P12004 | Proliferating cell nuclear antigen OS=Homo sapiens OX=9606 GN=PCNA PE=1 SV=1 | 72 | 11 | 11 | 28.8 | 50 | 261 | 4.69 | 10 | 11 | 9 | 6 | 3 | 6 | 8 | 9 | 8 | 6 | 3 | 6 |
| High | P18754 | Regulator of chromosome condensation OS=Homo sapiens OX=9606 GN=RCC1 PE=1 SV=1 | 71 | 14 | 14 | 44.9 | 45 | 421 | 7.52 | 2 | 2 | 4 | 4 | 5 | 3 | 2 | 2 | 3 | 4 | 5 | 3 |
| High | Q63HN8 | E3 ubiquitin-protein ligase RNF213 OS=Homo sapiens OX=9606 GN=RNF213 PE=1 SV=3 | 71 | 23 | 23 | 591 | 5 | 5207 | 6.48 | 14 | 15 | 11 | 2 | 7 | 4 | 14 | 15 | 10 | 2 | 6 | 4 |
| High | P00558 | Phosphoglycerate kinase 1 OS=Homo sapiens OX=9606 GN=PGK1 PE=1 SV=3 | 71 | 13 | 13 | 44.6 | 41 | 417 | 8.1 | 9 | 9 | 10 | 4 | 4 | 4 | 8 | 8 | 9 | 4 | 4 | 4 |
| High | P63241 | Eukaryotic translation initiation factor 5A-1 OS=Homo sapiens OX=9606 GN=EIF5A PE=1 SV=2 | 71 | 10 | 10 | 16.8 | 56 | 154 | 5.24 | 9 | 9 | 6 | 4 | 10 | 4 | 7 | 8 | 6 | 4 | 6 | 4 |
| High | P46781 | 40S ribosomal protein S9 OS=Homo sapiens OX=9606 GN=RPS9 PE=1 SV=3 | 71 | 10 | 10 | 22.6 | 43 | 194 | 10.65 | 8 | 9 | 8 | 5 | 8 | 5 | 8 | 8 | 8 | 5 | 8 | 5 |
| High | Q14240 | Eukaryotic initiation factor 4A-II OS=Homo sapiens OX=9606 GN=EIF4A2 PE=1 SV=2 | 71 | 10 | 2 | 46.4 | 26 | 407 | 5.48 | 8 | 8 | 10 | 7 | 6 | 7 | 7 | 7 | 9 | 7 | 6 | 7 |
| High | P42166 | Lamina-associated polypeptide 2, isoform alpha OS=Homo sapiens OX=9606 GN=TMPO PE=1 SV=2 | 71 | 16 | 11 | 75.4 | 34 | 694 | 7.66 | 8 | 8 | 12 | 4 | 4 | 4 | 7 | 7 | 10 | 4 | 4 | 4 |
| High | Q96HS1 | Serine/threonine-protein phosphatase PGAM5, mitochondrial OS=Homo sapiens OX=9606 GN=PGAM5 PE=1 SV=2 | 71 | 12 | 12 | 32 | 43 | 289 | 8.68 | 10 | 10 | 10 | 6 | 6 | 6 | 9 | 10 | 9 | 6 | 6 | 6 |
| High | P22392 | Nucleoside diphosphate kinase B OS=Homo sapiens OX=9606 GN=NME2 PE=1 SV=1 | 71 | 7 | 3 | 17.3 | 53 | 152 | 8.41 | 6 | 6 | 11 | 9 | 5 | 8 | 5 | 5 | 5 | 5 | 3 | 5 |
| High | P62879 | Guanine nucleotide-binding protein G(I)/G(S)/G(T) subunit beta-2 OS=Homo sapiens OX=9606 GN=GNB2 PE=1 SV=3 | 71 | 11 | 5 | 37.3 | 46 | 340 | 6 | 11 | 11 | 12 | 5 | 5 | 4 | 10 | 10 | 9 | 5 | 5 | 4 |
| High | P18085 | ADP-ribosylation factor 4 OS=Homo sapiens OX=9606 GN=ARF4 PE=1 SV=3 | 70 | 8 | 6 | 20.5 | 53 | 180 | 7.14 | 9 | 12 | 10 | 5 | 5 | 5 | 6 | 8 | 7 | 5 | 5 | 5 |
| High | Q92878 | DNA repair protein RAD50 OS=Homo sapiens OX=9606 GN=RAD50 PE=1 SV=1 | 70 | 24 | 24 | 153.8 | 19 | 1312 | 6.89 | 7 | 5 | 7 | 2 | 6 | 5 | 7 | 5 | 7 | 2 | 6 | 5 |
| High | Q9Y266 | Nuclear migration protein nudC OS=Homo sapiens OX=9606 GN=NUDC PE=1 SV=1 | 70 | 11 | 11 | 38.2 | 36 | 331 | 5.38 | 7 | 6 | 9 | 6 | 7 | 9 | 7 | 4 | 7 | 5 | 6 | 7 |
| High | Q9UQ35 | Serine/arginine repetitive matrix protein 2 OS=Homo sapiens OX=9606 GN=SRRM2 PE=1 SV=2 | 70 | 19 | 19 | 299.4 | 11 | 2752 | 12.06 | 3 | 3 | 5 | 9 | 10 | 8 | 3 | 3 | 5 | 9 | 10 | 8 |
| High | Q12797 | Aspartyl/asparaginyl beta-hydroxylase OS=Homo sapiens OX=9606 GN=ASPH PE=1 SV=3 | 70 | 16 | 16 | 85.8 | 26 | 758 | 5.01 | 4 | 4 | 5 | 7 | 8 | 5 | 4 | 4 | 4 | 7 | 8 | 5 |
| High | P50454 | Serpin H1 OS=Homo sapiens OX=9606 GN=SERPINH1 PE=1 SV=2 | 70 | 14 | 14 | 46.4 | 37 | 418 | 8.69 | 9 | 8 | 11 | 4 | 5 | 4 | 8 | 7 | 9 | 3 | 4 | 3 |
| High | P26641 | Elongation factor 1-gamma OS=Homo sapiens OX=9606 GN=EEF1G PE=1 SV=3 | 69 | 15 | 15 | 50.1 | 37 | 437 | 6.67 | 10 | 10 | 9 | 5 | 7 | 3 | 9 | 9 | 9 | 5 | 7 | 3 |
| High | Q9BYK8 | Helicase with zinc finger domain 2 OS=Homo sapiens OX=9606 GN=HELZ2 PE=1 SV=6 | 69 | 25 | 25 | 294.5 | 11 | 2649 | 7.49 | 3 | 3 | 1 | 13 | 10 | 15 | 3 | 3 | 1 | 12 | 10 | 15 |
| High | O95831 | Apoptosis-inducing factor 1, mitochondrial OS=Homo sapiens OX=9606 GN=AIFM1 PE=1 SV=1 | 69 | 12 | 12 | 66.9 | 25 | 613 | 8.95 | 8 | 9 | 9 | 4 | 7 | 3 | 8 | 9 | 9 | 4 | 7 | 3 |
| High | P23921 | Ribonucleoside-diphosphate reductase large subunit OS=Homo sapiens OX=9606 GN=RRM1 PE=1 SV=1 | 69 | 16 | 16 | 90 | 23 | 792 | 7.15 | 7 | 7 | 5 | 9 | 10 | 6 | 7 | 7 | 5 | 9 | 10 | 6 |
| High | O94905 | Erlin-2 OS=Homo sapiens OX=9606 GN=ERLIN2 PE=1 SV=1 | 69 | 12 | 9 | 37.8 | 44 | 339 | 5.62 | 10 | 11 | 8 | 4 | 3 | 3 | 9 | 10 | 7 | 4 | 3 | 3 |
| High | Q9NW13 | RNA-binding protein 28 OS=Homo sapiens OX=9606 GN=RBM28 PE=1 SV=3 | 69 | 13 | 13 | 85.7 | 18 | 759 | 9.22 | 3 | 3 | 5 | 8 | 8 | 9 | 3 | 3 | 5 | 8 | 8 | 9 |
| High | P61313 | 60S ribosomal protein L15 OS=Homo sapiens OX=9606 GN=RPL15 PE=1 SV=2 | 69 | 8 | 8 | 24.1 | 39 | 204 | 11.62 | 6 | 5 | 5 | 5 | 8 | 6 | 6 | 5 | 4 | 3 | 6 | 4 |
| High | P05387 | 60S acidic ribosomal protein P2 OS=Homo sapiens OX=9606 GN=RPLP2 PE=1 SV=1 | 69 | 9 | 8 | 11.7 | 93 | 115 | 4.54 | 5 | 5 | 5 | 4 | 9 | 6 | 5 | 5 | 5 | 4 | 7 | 4 |
| High | Q99569 | Plakophilin-4 OS=Homo sapiens OX=9606 GN=PKP4 PE=1 SV=2 | 68 | 20 | 20 | 131.8 | 24 | 1192 | 8.94 | 6 | 6 | 11 | 1 | 1 | 2 | 6 | 6 | 10 | 1 | 1 | 2 |
| High | Q9H444 | Charged multivesicular body protein 4b OS=Homo sapiens OX=9606 GN=CHMP4B PE=1 SV=1 | 68 | 9 | 9 | 24.9 | 47 | 224 | 4.82 | 5 | 8 | 8 | 3 | 3 | 3 | 5 | 8 | 8 | 3 | 3 | 3 |
| High | P35914 | Hydroxymethylglutaryl-CoA lyase, mitochondrial OS=Homo sapiens OX=9606 GN=HMGCL PE=1 SV=2 | 68 | 10 | 10 | 34.3 | 36 | 325 | 8.54 | 5 | 6 | 7 | 6 | 6 | 6 | 5 | 6 | 7 | 6 | 6 | 6 |
| High | P35998 | 26S proteasome regulatory subunit 7 OS=Homo sapiens OX=9606 GN=PSMC2 PE=1 SV=3 | 68 | 17 | 17 | 48.6 | 44 | 433 | 5.95 | 2 | 3 | 2 | 11 | 11 | 11 | 2 | 3 | 2 | 10 | 11 | 10 |
| High | P36957 | Dihydrolipoyllysine-residue succinyltransferase component of 2-oxoglutarate dehydrogenase complex, mitochondrial OS=Homo s | 68 | 10 | 10 | 48.7 | 28 | 453 | 8.95 | 6 | 7 | 9 | 6 | 4 | 3 | 6 | 6 | 8 | 5 | 3 | 3 |

| | | | | | | | | | | | | | | | | | | | | | |
|---|---|---|---|---|---|---|---|---|---|---|---|---|---|---|---|---|---|---|---|---|---|
| High | O00148 | ATP-dependent RNA helicase DDX39A OS=Homo sapiens OX=9606 GN=DDX39A PE=1 SV=2 | 68 | 12 | 4 | 49.1 | 34 | 427 | 5.68 | 10 | 8 | 13 | 5 | 6 | 4 | 9 | 8 | 11 | 4 | 6 | 4 |
| High | P09543 | 2',3'-cyclic-nucleotide 3'-phosphodiesterase OS=Homo sapiens OX=9606 GN=CNP PE=1 SV=2 | 68 | 15 | 15 | 47.5 | 34 | 421 | 9.07 | 5 | 5 | 9 | 7 | 6 | 9 | 5 | 5 | 8 | 7 | 6 | 9 |
| High | O15438 | ATP-binding cassette sub-family C member 3 OS=Homo sapiens OX=9606 GN=ABCC3 PE=1 SV=3 | 68 | 22 | 20 | 169.2 | 19 | 1527 | 7.2 | 17 | 20 | 18 | | | 1 | 14 | 17 | 15 | | | 1 |
| High | Q8N1G4 | Leucine-rich repeat-containing protein 47 OS=Homo sapiens OX=9606 GN=LRRC47 PE=1 SV=1 | 68 | 14 | 14 | 63.4 | 30 | 583 | 8.28 | 7 | 5 | 8 | 6 | 6 | 4 | 7 | 5 | 8 | 6 | 6 | 4 |
| High | Q9Y2X3 | Nucleolar protein 58 OS=Homo sapiens OX=9606 GN=NOP58 PE=1 SV=1 | 67 | 13 | 13 | 59.5 | 31 | 529 | 8.92 | 3 | 4 | 7 | 5 | 6 | 6 | 3 | 4 | 7 | 5 | 6 | 6 |
| High | Q15046 | Lysine--tRNA ligase OS=Homo sapiens OX=9606 GN=KARS1 PE=1 SV=3 | 67 | 13 | 13 | 68 | 25 | 597 | 6.35 | 10 | 4 | 8 | 5 | 5 | 5 | 10 | 4 | 7 | 5 | 5 | 5 |
| High | Q13263 | Transcription intermediary factor 1-beta OS=Homo sapiens OX=9606 GN=TRIM28 PE=1 SV=5 | 67 | 13 | 13 | 88.5 | 26 | 835 | 5.77 | 5 | 5 | 8 | 6 | 5 | 6 | 5 | 5 | 8 | 6 | 5 | 6 |
| High | P51114 | Fragile X mental retardation syndrome-related protein 1 OS=Homo sapiens OX=9606 GN=FXR1 PE=1 SV=3 | 67 | 17 | 15 | 69.7 | 36 | 621 | 6.15 | 6 | 7 | 4 | 8 | 8 | 7 | 6 | 7 | 4 | 7 | 7 | 6 |
| High | O43837 | Isocitrate dehydrogenase [NAD] subunit beta, mitochondrial OS=Homo sapiens OX=9606 GN=IDH3B PE=1 SV=2 | 67 | 14 | 14 | 42.2 | 48 | 385 | 8.46 | 7 | 8 | 7 | 5 | 6 | 6 | 7 | 8 | 7 | 5 | 6 | 6 |
| High | P60903 | Protein S100-A10 OS=Homo sapiens OX=9606 GN=S100A10 PE=1 SV=2 | 67 | 3 | 3 | 11.2 | 18 | 97 | 7.37 | 5 | 3 | 5 | 3 | 5 | 4 | 2 | 1 | 2 | 3 | 3 | 3 |
| High | Q6UXN9 | WD repeat-containing protein 82 OS=Homo sapiens OX=9606 GN=WDR82 PE=1 SV=1 | 67 | 11 | 11 | 35.1 | 35 | 313 | 7.69 | 1 | 1 | 1 | 10 | 12 | 11 | 1 | 1 | 1 | 8 | 8 | 9 |
| High | P61204 | ADP-ribosylation factor 3 OS=Homo sapiens OX=9606 GN=ARF3 PE=1 SV=2 | 66 | 7 | 4 | 20.6 | 48 | 181 | 7.43 | 9 | 11 | 11 | 4 | 6 | 4 | 7 | 7 | 7 | 4 | 5 | 4 |
| High | P52789 | Hexokinase-2 OS=Homo sapiens OX=9606 GN=HK2 PE=1 SV=2 | 66 | 20 | 17 | 102.3 | 25 | 917 | 6.05 | 7 | 9 | 10 | 1 | 2 | 1 | 7 | 9 | 10 | 1 | 2 | 1 |
| High | P15170 | Eukaryotic peptide chain release factor GTP-binding subunit ERF3A OS=Homo sapiens OX=9606 GN=GSPT1 PE=1 SV=1 | 66 | 15 | 15 | 55.7 | 37 | 499 | 5.62 | 8 | 7 | 11 | 3 | 2 | 6 | 7 | 7 | 10 | 3 | 2 | 6 |
| High | Q15084 | Protein disulfide-isomerase A6 OS=Homo sapiens OX=9606 GN=PDIA6 PE=1 SV=1 | 66 | 8 | 8 | 48.1 | 26 | 440 | 5.08 | 9 | 9 | 9 | 5 | 5 | 5 | 7 | 7 | 7 | 5 | 5 | 5 |
| High | O15231 | Zinc finger protein 185 OS=Homo sapiens OX=9606 GN=ZNF185 PE=1 SV=3 | 65 | 18 | 18 | 73.5 | 35 | 689 | 7.01 | 10 | 8 | 9 | 2 | 3 | 4 | 10 | 8 | 9 | 2 | 3 | 4 |
| High | Q6XQN6 | Nicotinate phosphoribosyltransferase OS=Homo sapiens OX=9606 GN=NAPRT PE=1 SV=2 | 65 | 11 | 11 | 57.5 | 33 | 538 | 5.68 | 6 | 4 | 9 | 6 | 7 | 7 | 6 | 4 | 9 | 6 | 5 | 7 |
| High | O60264 | SWI/SNF-related matrix-associated actin-dependent regulator of chromatin subfamily A member 5 OS=Homo sapiens OX=9606 GI | 65 | 18 | 18 | 121.8 | 18 | 1052 | 8.09 | 6 | 9 | 8 | 8 | 9 | 5 | 6 | 9 | 8 | 8 | 9 | 5 |
| High | Q9BVI0 | PHD finger protein 20 OS=Homo sapiens OX=9606 GN=PHF20 PE=1 SV=2 | 65 | 18 | 18 | 115.3 | 23 | 1012 | 6.99 | | | | 9 | 10 | 10 | | | | 9 | 10 | 10 |
| High | Q14697 | Neutral alpha-glucosidase AB OS=Homo sapiens OX=9606 GN=GANAB PE=1 SV=3 | 65 | 20 | 20 | 106.8 | 25 | 944 | 6.14 | 8 | 12 | 11 | 2 | 2 | 4 | 8 | 12 | 11 | 2 | 2 | 4 |
| High | P62829 | 60S ribosomal protein L23 OS=Homo sapiens OX=9606 GN=RPL23 PE=1 SV=1 | 65 | 5 | 5 | 14.9 | 44 | 140 | 10.51 | 4 | 7 | 3 | 4 | 7 | 5 | 3 | 4 | 3 | 3 | 5 | 4 |
| High | P36873 | Serine/threonine-protein phosphatase PP1-gamma catalytic subunit OS=Homo sapiens OX=9606 GN=PPP1CC PE=1 SV=1 | 65 | 8 | 1 | 37 | 31 | 323 | 6.54 | 9 | 8 | 7 | 4 | 3 | 3 | 6 | 7 | 6 | 4 | 3 | 3 |
| High | P49589 | Cysteine--tRNA ligase, cytoplasmic OS=Homo sapiens OX=9606 GN=CARS1 PE=1 SV=3 | 65 | 14 | 14 | 85.4 | 21 | 748 | 6.76 | 5 | 5 | 5 | 7 | 6 | 4 | 5 | 5 | 5 | 6 | 6 | 4 |
| High | P35222 | Catenin beta-1 OS=Homo sapiens OX=9606 GN=CTNNB1 PE=1 SV=1 | 65 | 13 | 9 | 85.4 | 18 | 781 | 5.86 | 10 | 9 | 8 | 2 | 3 | 2 | 8 | 9 | 8 | 2 | 3 | 2 |
| High | Q9UQE7 | Structural maintenance of chromosomes protein 3 OS=Homo sapiens OX=9606 GN=SMC3 PE=1 SV=2 | 65 | 22 | 22 | 141.5 | 22 | 1217 | 7.18 | 6 | 4 | 9 | 2 | 6 | 3 | 6 | 4 | 9 | 2 | 6 | 3 |
| High | P30050 | 60S ribosomal protein L12 OS=Homo sapiens OX=9606 GN=RPL12 PE=1 SV=1 | 64 | 5 | 5 | 17.8 | 45 | 165 | 9.42 | 5 | 5 | 5 | 5 | 8 | 5 | 5 | 5 | 5 | 5 | 5 | 5 |
| High | O75534 | Cold shock domain-containing protein E1 OS=Homo sapiens OX=9606 GN=CSDE1 PE=1 SV=2 | 64 | 16 | 16 | 88.8 | 21 | 798 | 6.25 | 3 | 2 | 3 | 8 | 8 | 11 | 3 | 2 | 3 | 8 | 8 | 11 |
| High | P35813 | Protein phosphatase 1A OS=Homo sapiens OX=9606 GN=PPM1A PE=1 SV=1 | 64 | 7 | 4 | 42.4 | 23 | 382 | 5.36 | 6 | 8 | 7 | 6 | 5 | 7 | 4 | 4 | 6 | 3 | 2 | 4 |
| High | P62873 | Guanine nucleotide-binding protein G(I)/G(S)/G(T) subunit beta-1 OS=Homo sapiens OX=9606 GN=GNB1 PE=1 SV=3 | 64 | 11 | 5 | 37.4 | 40 | 340 | 6 | 9 | 8 | 8 | 6 | 4 | 5 | 9 | 8 | 7 | 6 | 4 | 5 |
| High | P62888 | 60S ribosomal protein L30 OS=Homo sapiens OX=9606 GN=RPL30 PE=1 SV=2 | 64 | 5 | 5 | 12.8 | 57 | 115 | 9.63 | 5 | 3 | 6 | 5 | 7 | 4 | 4 | 3 | 4 | 5 | 4 | 4 |
| High | Q13247 | Serine/arginine-rich splicing factor 6 OS=Homo sapiens OX=9606 GN=SRSF6 PE=1 SV=2 | 64 | 9 | 8 | 39.6 | 23 | 344 | 11.43 | 7 | 5 | 4 | 13 | 6 | 7 | 7 | 5 | 4 | 9 | 5 | 6 |
| High | P24752 | Acetyl-CoA acetyltransferase, mitochondrial OS=Homo sapiens OX=9606 GN=ACAT1 PE=1 SV=1 | 64 | 12 | 12 | 45.2 | 30 | 427 | 8.85 | 2 | 4 | 5 | 8 | 7 | 7 | 2 | 4 | 5 | 8 | 7 | 7 |
| High | P11166 | Solute carrier family 2, facilitated glucose transporter member 1 OS=Homo sapiens OX=9606 GN=SLC2A1 PE=1 SV=2 | 64 | 7 | 7 | 54 | 12 | 492 | 8.72 | 10 | 10 | 11 | 2 | 4 | 5 | 6 | 4 | 6 | 2 | 3 | 4 |
| High | P23284 | Peptidyl-prolyl cis-trans isomerase B OS=Homo sapiens OX=9606 GN=PPIB PE=1 SV=2 | 64 | 12 | 12 | 23.7 | 50 | 216 | 9.41 | 8 | 7 | 7 | 9 | 9 | 8 | 8 | 7 | 7 | 8 | 8 | 8 |
| High | Q09028 | Histone-binding protein RBBP4 OS=Homo sapiens OX=9606 GN=RBBP4 PE=1 SV=3 | 64 | 9 | 4 | 47.6 | 22 | 425 | 4.89 | 4 | 3 | 6 | 9 | 8 | 8 | 4 | 3 | 5 | 7 | 7 | 7 |
| High | P04424 | Argininosuccinate lyase OS=Homo sapiens OX=9606 GN=ASL PE=1 SV=4 | 64 | 14 | 14 | 51.6 | 35 | 464 | 6.48 | 6 | 6 | 5 | 4 | 5 | 4 | 6 | 6 | 5 | 4 | 5 | 4 |
| High | Q99543 | DnaJ homolog subfamily C member 2 OS=Homo sapiens OX=9606 GN=DNAJC2 PE=1 SV=4 | 63 | 14 | 14 | 72 | 24 | 621 | 8.7 | 4 | 6 | 5 | 5 | 5 | 8 | 4 | 6 | 5 | 5 | 5 | 8 |
| High | P17655 | Calpain-2 catalytic subunit OS=Homo sapiens OX=9606 GN=CAPN2 PE=1 SV=6 | 63 | 14 | 14 | 79.9 | 28 | 700 | 4.98 | 7 | 6 | 4 | 7 | 3 | 6 | 7 | 6 | 4 | 7 | 3 | 6 |
| High | P04080 | Cystatin-B OS=Homo sapiens OX=9606 GN=CSTB PE=1 SV=2 | 63 | 5 | 5 | 11.1 | 77 | 98 | 7.56 | 5 | 7 | 8 | 4 | 8 | 5 | 4 | 5 | 4 | 3 | 4 | 3 |
| High | P62263 | 40S ribosomal protein S14 OS=Homo sapiens OX=9606 GN=RPS14 PE=1 SV=3 | 63 | 6 | 6 | 16.3 | 38 | 151 | 10.05 | 6 | 5 | 7 | 9 | 7 | 8 | 4 | 4 | 5 | 5 | 5 | 5 |
| High | P62993 | Growth factor receptor-bound protein 2 OS=Homo sapiens OX=9606 GN=GRB2 PE=1 SV=1 | 63 | 12 | 12 | 25.2 | 61 | 217 | 6.32 | 7 | 6 | 10 | 6 | 6 | 6 | 6 | 6 | 8 | 6 | 6 | 6 |
| High | Q96C19 | EF-hand domain-containing protein D2 OS=Homo sapiens OX=9606 GN=EFHD2 PE=1 SV=1 | 63 | 11 | 11 | 26.7 | 38 | 240 | 5.2 | 8 | 10 | 9 | 5 | 4 | 4 | 8 | 9 | 8 | 5 | 4 | 4 |
| High | Q14258 | E3 ubiquitin/ISG15 ligase TRIM25 OS=Homo sapiens OX=9606 GN=TRIM25 PE=1 SV=2 | 63 | 13 | 13 | 70.9 | 29 | 630 | 8.09 | 10 | 12 | 7 | 4 | 4 | 4 | 9 | 11 | 7 | 4 | 3 | 4 |
| High | P09496 | Clathrin light chain A OS=Homo sapiens OX=9606 GN=CLTA PE=1 SV=1 | 63 | 7 | 7 | 27.1 | 20 | 248 | 4.51 | 7 | 7 | 7 | 4 | 6 | 6 | 6 | 6 | 5 | 4 | 6 | 6 |
| High | P11388 | DNA topoisomerase 2-alpha OS=Homo sapiens OX=9606 GN=TOP2A PE=1 SV=3 | 62 | 16 | 8 | 174.3 | 11 | 1531 | 8.72 | 6 | 12 | 12 | 3 | 6 | 3 | 6 | 12 | 10 | 3 | 5 | 3 |
| High | Q04837 | Single-stranded DNA-binding protein, mitochondrial OS=Homo sapiens OX=9606 GN=SSBP1 PE=1 SV=1 | 62 | 7 | 7 | 17.2 | 52 | 148 | 9.6 | 10 | 8 | 8 | 6 | 8 | 6 | 7 | 6 | 7 | 5 | 5 | 5 |
| High | Q9NUQ6 | SPATS2-like protein OS=Homo sapiens OX=9606 GN=SPATS2L PE=1 SV=2 | 62 | 12 | 12 | 61.7 | 28 | 558 | 9.64 | 4 | 4 | 3 | 6 | 9 | 8 | 4 | 4 | 3 | 6 | 8 | 7 |
| High | Q9Y520 | Protein PRRC2C OS=Homo sapiens OX=9606 GN=PRRC2C PE=1 SV=4 | 62 | 19 | 19 | 316.7 | 8 | 2896 | 9.13 | | 1 | 1 | 7 | 9 | 8 | | 1 | 1 | 7 | 9 | 8 |
| High | Q99873 | Protein arginine N-methyltransferase 1 OS=Homo sapiens OX=9606 GN=PRMT1 PE=1 SV=3 | 62 | 10 | 10 | 42.4 | 33 | 371 | 5.35 | 4 | 4 | 8 | 6 | 6 | 7 | 4 | 4 | 8 | 6 | 6 | 7 |
| High | Q13596 | Sorting nexin-1 OS=Homo sapiens OX=9606 GN=SNX1 PE=1 SV=3 | 62 | 12 | 10 | 59 | 27 | 522 | 5.15 | 5 | 3 | 6 | 6 | 9 | 8 | 5 | 3 | 6 | 6 | 9 | 7 |
| High | Q96EZ8 | Microspherule protein 1 OS=Homo sapiens OX=9606 GN=MCRS1 PE=1 SV=1 | 62 | 15 | 15 | 51.8 | 39 | 462 | 9.38 | | | | 9 | 10 | 7 | | | | 9 | 8 | 7 |
| High | Q9H7Z6 | Histone acetyltransferase KAT8 OS=Homo sapiens OX=9606 GN=KAT8 PE=1 SV=2 | 62 | 13 | 12 | 52.4 | 32 | 458 | 8.27 | | | | 6 | 12 | 8 | | | | 6 | 11 | 7 |
| High | Q13435 | Splicing factor 3B subunit 2 OS=Homo sapiens OX=9606 GN=SF3B2 PE=1 SV=2 | 62 | 16 | 16 | 100.2 | 22 | 895 | 5.67 | 2 | 3 | 2 | 5 | 9 | 5 | 2 | 3 | 2 | 5 | 9 | 5 |
| High | Q99439 | Calponin-2 OS=Homo sapiens OX=9606 GN=CNN2 PE=1 SV=4 | 62 | 11 | 11 | 33.7 | 54 | 309 | 7.33 | 6 | 5 | 7 | 5 | 6 | 6 | 5 | 5 | 5 | 5 | 6 | 6 |
| High | Q1KMD3 | Heterogeneous nuclear ribonucleoprotein U-like protein 2 OS=Homo sapiens OX=9606 GN=HNRNPUL2 PE=1 SV=1 | 62 | 13 | 13 | 85.1 | 19 | 747 | 4.91 | 8 | 7 | 9 | 4 | 5 | 5 | 8 | 7 | 9 | 4 | 5 | 5 |
| High | P61081 | NEDD8-conjugating enzyme Ubc12 OS=Homo sapiens OX=9606 GN=UBE2M PE=1 SV=1 | 61 | 7 | 7 | 20.9 | 33 | 183 | 7.69 | 7 | 7 | 8 | 5 | 5 | 7 | 7 | 6 | 7 | 5 | 5 | 7 |
| High | Q7L0Y3 | tRNA methyltransferase 10 homolog C OS=Homo sapiens OX=9606 GN=TRMT10C PE=1 SV=2 | 61 | 16 | 16 | 47.3 | 41 | 403 | 9.36 | 5 | 8 | 5 | 4 | 8 | 3 | 5 | 8 | 5 | 4 | 8 | 3 |
| High | Q8WXF1 | Paraspeckle component 1 OS=Homo sapiens OX=9606 GN=PSPC1 PE=1 SV=1 | 61 | 13 | 13 | 58.7 | 33 | 523 | 6.67 | 1 | 1 | 1 | 9 | 11 | 9 | 1 | 1 | 1 | 9 | 11 | 9 |
| High | Q8TEM1 | Nuclear pore membrane glycoprotein 210 OS=Homo sapiens OX=9606 GN=NUP210 PE=1 SV=3 | 61 | 18 | 18 | 205 | 12 | 1887 | 6.81 | 3 | 2 | 4 | 3 | 6 | 7 | 3 | 2 | 4 | 3 | 6 | 7 |
| High | Q02218 | 2-oxoglutarate dehydrogenase, mitochondrial OS=Homo sapiens OX=9606 GN=OGDH PE=1 SV=3 | 61 | 18 | 18 | 115.9 | 21 | 1023 | 6.86 | 5 | 5 | 5 | 3 | 2 | 4 | 5 | 5 | 5 | 3 | 2 | 4 |
| High | P26368 | Splicing factor U2AF 65 kDa subunit OS=Homo sapiens OX=9606 GN=U2AF2 PE=1 SV=4 | 61 | 8 | 8 | 53.5 | 29 | 475 | 9.09 | 6 | 4 | 5 | 5 | 6 | 7 | 6 | 4 | 5 | 5 | 4 | 5 |
| High | Q14103 | Heterogeneous nuclear ribonucleoprotein D0 OS=Homo sapiens OX=9606 GN=HNRNPD PE=1 SV=1 | 61 | 8 | 6 | 38.4 | 24 | 355 | 7.81 | 8 | 8 | 9 | 4 | 5 | 3 | 7 | 7 | 7 | 3 | 4 | 2 |
| High | Q9BSH5 | Haloacid dehalogenase-like hydrolase domain-containing protein 3 OS=Homo sapiens OX=9606 GN=HDHD3 PE=1 SV=1 | 61 | 8 | 8 | 28 | 45 | 251 | 6.71 | 8 | 8 | 9 | 4 | 3 | 4 | 6 | 6 | 7 | 4 | 3 | 4 |
| High | P38919 | Eukaryotic initiation factor 4A-III OS=Homo sapiens OX=9606 GN=EIF4A3 PE=1 SV=4 | 61 | 13 | 11 | 46.8 | 31 | 411 | 6.73 | 8 | 9 | 5 | 5 | 8 | 7 | 8 | 9 | 5 | 5 | 8 | 7 |
| High | Q01167 | Forkhead box protein K2 OS=Homo sapiens OX=9606 GN=FOXK2 PE=1 SV=3 | 61 | 13 | 11 | 69 | 27 | 660 | 9.54 | | | | 8 | 12 | 8 | | | | 8 | 11 | 8 |
| High | Q9UBU9 | Nuclear RNA export factor 1 OS=Homo sapiens OX=9606 GN=NXF1 PE=1 SV=1 | 60 | 23 | 23 | 70.1 | 44 | 619 | 8.51 | | | | | | | | | | | | |
| High | Q16576 | Histone-binding protein RBBP7 OS=Homo sapiens OX=9606 GN=RBBP7 PE=1 SV=1 | 60 | 10 | 5 | 47.8 | 29 | 425 | 5.05 | 5 | 4 | 5 | 7 | 6 | 6 | 5 | 4 | 5 | 6 | 5 | 6 |
| High | P08670 | Vimentin OS=Homo sapiens OX=9606 GN=VIM PE=1 SV=4 | 60 | 10 | 6 | 53.6 | 20 | 466 | 5.12 | 6 | 7 | 7 | 9 | 10 | 8 | 3 | 3 | 3 | 7 | 6 | 7 |
| High | P52948 | Nuclear pore complex protein Nup98-Nup96 OS=Homo sapiens OX=9606 GN=NUP98 PE=1 SV=4 | 60 | 21 | 21 | 197.5 | 16 | 1817 | 6.4 | 4 | 2 | 2 | 2 | 1 | 3 | 4 | 2 | 2 | 2 | 1 | 3 |
| High | P09622 | Dihydrolipoyl dehydrogenase, mitochondrial OS=Homo sapiens OX=9606 GN=DLD PE=1 SV=2 | 60 | 10 | 10 | 54.1 | 28 | 509 | 7.85 | 10 | 10 | 11 | 3 | 3 | 3 | 8 | 9 | 8 | 3 | 2 | 3 |
| High | P26196 | Probable ATP-dependent RNA helicase DDX6 OS=Homo sapiens OX=9606 GN=DDX6 PE=1 SV=2 | 59 | 11 | 11 | 54.4 | 30 | 483 | 8.66 | 5 | 5 | 4 | 5 | 7 | 7 | 5 | 5 | 4 | 5 | 7 | 7 |
| High | Q12888 | TP53-binding protein 1 OS=Homo sapiens OX=9606 GN=TP53BP1 PE=1 SV=2 | 59 | 20 | 20 | 213.4 | 17 | 1972 | 4.7 | 1 | 4 | 4 | 4 | 7 | 6 | 1 | 4 | 4 | 4 | 7 | 6 |
| High | P21964 | Catechol O-methyltransferase OS=Homo sapiens OX=9606 GN=COMT PE=1 SV=2 | 59 | 11 | 11 | 30 | 61 | 271 | 5.47 | 9 | 8 | 7 | 3 | 4 | 3 | 7 | 5 | 6 | 3 | 4 | 2 |
| High | Q86XN7 | Proline and serine-rich protein 1 OS=Homo sapiens OX=9606 GN=PROSER1 PE=1 SV=2 | 59 | 12 | 12 | 95.6 | 16 | 944 | 8.88 | | | | 11 | 14 | 9 | | | | 8 | 10 | 8 |
| High | P62753 | 40S ribosomal protein S6 OS=Homo sapiens OX=9606 GN=RPS6 PE=1 SV=1 | 59 | 6 | 6 | 28.7 | 20 | 249 | 10.84 | 5 | 4 | 3 | 6 | 8 | 6 | 4 | 4 | 3 | 4 | 4 | 5 |
| High | A5YKK6 | CCR4-NOT transcription complex subunit 1 OS=Homo sapiens OX=9606 GN=CNOT1 PE=1 SV=2 | 59 | 21 | 21 | 266.8 | 11 | 2376 | 7.11 | 3 | 5 | 2 | 4 | 5 | 4 | 3 | 5 | 2 | 4 | 5 | 4 |
| High | Q92900 | Regulator of nonsense transcripts 1 OS=Homo sapiens OX=9606 GN=UPF1 PE=1 SV=2 | 59 | 13 | 13 | 124.3 | 12 | 1129 | 6.61 | 3 | 5 | 4 | 9 | 10 | 9 | 3 | 5 | 4 | 8 | 10 | 9 |
| High | P09497 | Clathrin light chain B OS=Homo sapiens OX=9606 GN=CLTB PE=1 SV=1 | 59 | 10 | 10 | 25.2 | 28 | 229 | 4.64 | 5 | 5 | 5 | 6 | 7 | 7 | 5 | 5 | 5 | 6 | 7 | 7 |
| High | Q9Y5K6 | CD2-associated protein OS=Homo sapiens OX=9606 GN=CD2AP PE=1 SV=1 | 59 | 15 | 15 | 71.4 | 29 | 639 | 6.4 | 5 | 6 | 4 | 2 | 5 | 3 | 5 | 6 | 4 | 2 | 4 | 3 |
| High | Q5JRX3 | Presequence protease, mitochondrial OS=Homo sapiens OX=9606 GN=PITRM1 PE=1 SV=3 | 59 | 17 | 17 | 117.3 | 21 | 1037 | 6.92 | 5 | 5 | 4 | 5 | 6 | 6 | 5 | 5 | 4 | 4 | 5 | 6 |
| High | Q8WVV4 | Protein POF1B OS=Homo sapiens OX=9606 GN=POF1B PE=1 SV=3 | 59 | 15 | 15 | 68 | 33 | 589 | 6.32 | 9 | 7 | 8 | 3 | 4 | 3 | 9 | 7 | 8 | 3 | 4 | 3 |
| High | O43592 | Exportin-T OS=Homo sapiens OX=9606 GN=XPOT PE=1 SV=2 | 59 | 19 | 19 | 109.9 | 25 | 962 | 5.39 | 15 | 14 | 17 | | 2 | | 15 | 14 | 16 | | 2 | |
| High | Q9NS37 | CREB/ATF bZIP transcription factor OS=Homo sapiens OX=9606 GN=CREBZF PE=1 SV=2 | 59 | 14 | 14 | 37.1 | 44 | 354 | 5.24 | | | | 8 | 5 | 8 | | | | 8 | 5 | 8 |
| High | Q9BVP2 | Guanine nucleotide-binding protein-like 3 OS=Homo sapiens OX=9606 GN=GNL3 PE=1 SV=2 | 58 | 14 | 13 | 62 | 27 | 549 | 9.16 | 2 | 6 | 3 | 5 | 6 | 6 | 2 | 4 | 2 | 4 | 6 | 6 |
| High | Q99567 | Nuclear pore complex protein Nup88 OS=Homo sapiens OX=9606 GN=NUP88 PE=1 SV=2 | 58 | 14 | 14 | 83.5 | 27 | 741 | 5.69 | 2 | 1 | 3 | 5 | 7 | 4 | 2 | 1 | 3 | 5 | 7 | 4 |
| High | Q3SX09 | TREMBL:Q3SX09 (Bos taurus) similar to HBG protein | 58 | 9 | 9 | 22 | 45 | 201 | 9.39 | 5 | 6 | 4 | 9 | 10 | 9 | 5 | 5 | 4 | 5 | 7 | 4 |
| High | Q9UE12 | TREMBL:Q9UE12 Type I hair keratin 1 - Homo sapiens (Human). | 58 | 3 | 1 | 47.2 | 6 | 416 | 4.88 | 8 | 9 | 12 | 5 | 8 | 4 | 2 | 2 | 2 | 2 | 2 | 2 |
| High | Q04637 | Eukaryotic translation initiation factor 4 gamma 1 OS=Homo sapiens OX=9606 GN=EIF4G1 PE=1 SV=4 | 58 | 18 | 15 | 175.4 | 13 | 1599 | 5.33 | 2 | 4 | 4 | 4 | 7 | 5 | 2 | 4 | 4 | 4 | 7 | 5 |
| High | O95861 | 3'(2'),5'-bisphosphate nucleotidase 1 OS=Homo sapiens OX=9606 GN=BPNT1 PE=1 SV=1 | 58 | 11 | 11 | 33.4 | 50 | 308 | 5.69 | 3 | 1 | 3 | 6 | 6 | 6 | 3 | 1 | 3 | 6 | 5 | 6 |
| High | P62917 | 60S ribosomal protein L8 OS=Homo sapiens OX=9606 GN=RPL8 PE=1 SV=2 | 58 | 7 | 7 | 28 | 41 | 257 | 11.03 | 5 | 5 | 5 | 5 | 6 | 6 | 4 | 5 | 5 | 5 | 4 | 5 |
| High | P26038 | Moesin OS=Homo sapiens OX=9606 GN=MSN PE=1 SV=3 | 58 | 12 | 8 | 67.8 | 19 | 577 | 6.4 | 4 | 6 | 10 | 4 | 6 | 3 | 4 | 6 | 9 | 4 | 6 | 2 |
| High | P30048 | Thioredoxin-dependent peroxide reductase, mitochondrial OS=Homo sapiens OX=9606 GN=PRDX3 PE=1 SV=3 | 58 | 7 | 7 | 27.7 | 36 | 256 | 7.78 | 6 | 6 | 7 | 6 | 3 | 5 | 6 | 5 | 6 | 5 | 3 | 5 |
| High | Q13242 | Serine/arginine-rich splicing factor 9 OS=Homo sapiens OX=9606 GN=SRSF9 PE=1 SV=1 | 58 | 10 | 10 | 25.5 | 39 | 221 | 8.65 | 5 | 6 | 9 | 6 | 5 | 8 | 5 | 5 | 8 | 5 | 5 | 7 |
| High | Q15436 | Protein transport protein Sec23A OS=Homo sapiens OX=9606 GN=SEC23A PE=1 SV=2 | 58 | 11 | 8 | 86.1 | 18 | 765 | 7.08 | 4 | 7 | 10 | 3 | 3 | 4 | 4 | 7 | 10 | 3 | 3 | 4 |
| High | Q9UG63 | ATP-binding cassette sub-family F member 2 OS=Homo sapiens OX=9606 GN=ABCF2 PE=1 SV=2 | 58 | 13 | 13 | 71.2 | 21 | 623 | 7.37 | 9 | 7 | 6 | 4 | 5 | 3 | 9 | 7 | 6 | 4 | 5 | 3 |
| High | Q09161 | Nuclear cap-binding protein subunit 1 OS=Homo sapiens OX=9606 GN=NCBP1 PE=1 SV=1 | 58 | 13 | 13 | 91.8 | 22 | 790 | 6.43 | 3 | 4 | 5 | 5 | 8 | 8 | 3 | 4 | 5 | 5 | 8 | 8 |
| High | P42330 | Aldo-keto reductase family 1 member C3 OS=Homo sapiens OX=9606 GN=AKR1C3 PE=1 SV=4 | 58 | 10 | 4 | 36.8 | 42 | 323 | 7.94 | 10 | 11 | 9 | 2 | 3 | 2 | 9 | 9 | 8 | 2 | 3 | 2 |
| High | Q9Y3F4 | Serine-threonine kinase receptor-associated protein OS=Homo sapiens OX=9606 GN=STRAP PE=1 SV=1 | 58 | 16 | 16 | 38.4 | 57 | 350 | 5.12 | 2 | 2 | 5 | 7 | 8 | 6 | 2 | 2 | 4 | 7 | 8 | 6 |
| High | P51858 | Hepatoma-derived growth factor OS=Homo sapiens OX=9606 GN=HDGF PE=1 SV=1 | 58 | 13 | 13 | 26.8 | 61 | 240 | 4.73 | 4 | 5 | 6 | 4 | 6 | 7 | 4 | 5 | 6 | 4 | 6 | 7 |
| High | P36551 | Oxygen-dependent coproporphyrinogen-III oxidase, mitochondrial OS=Homo sapiens OX=9606 GN=CPOX PE=1 SV=3 | 58 | 11 | 11 | 50.1 | 29 | 454 | 8.25 | 5 | 4 | 6 | 4 | 4 | 4 | 5 | 4 | 6 | 4 | 4 | 4 |
| High | O75400 | Pre-mRNA-processing factor 40 homolog A OS=Homo sapiens OX=9606 GN=PRPF40A PE=1 SV=2 | 58 | 13 | 13 | 108.7 | 13 | 957 | 7.56 | 2 | 2 | 3 | 9 | 8 | 8 | 2 | 2 | 3 | 9 | 7 | 8 |
| High | Q9NZL9 | Methionine adenosyltransferase 2 subunit beta OS=Homo sapiens OX=9606 GN=MAT2B PE=1 SV=1 | 58 | 10 | 10 | 37.5 | 39 | 334 | 7.36 | 8 | 9 | 7 | 3 | 6 | 4 | 8 | 9 | 7 | 3 | 5 | 4 |
| High | P17980 | 26S proteasome regulatory subunit 6A OS=Homo sapiens OX=9606 GN=PSMC3 PE=1 SV=3 | 57 | 11 | 11 | 49.2 | 33 | 439 | 5.24 | 2 | 2 | 4 | 5 | 8 | 8 | 2 | 2 | 3 | 5 | 7 | 7 |
| High | P04183 | Thymidine kinase, cytosolic OS=Homo sapiens OX=9606 GN=TK1 PE=1 SV=2 | 57 | 8 | 8 | 25.5 | 38 | 234 | 8.51 | 2 | 3 | 3 | 5 | 9 | 4 | 2 | 3 | 3 | 5 | 8 | 4 |
| High | Q8WUM0 | Nuclear pore complex protein Nup133 OS=Homo sapiens OX=9606 GN=NUP133 PE=1 SV=2 | 57 | 16 | 16 | 128.9 | 17 | 1156 | 5.1 | 3 | 5 | 6 | 3 | 3 | 2 | 3 | 5 | 6 | 3 | 3 | 2 |
| High | O60832 | H/ACA ribonucleoprotein complex subunit DKC1 OS=Homo sapiens OX=9606 GN=DKC1 PE=1 SV=3 | 57 | 14 | 14 | 57.6 | 29 | 514 | 9.42 | 3 | 3 | 5 | 3 | 6 | 4 | 3 | 3 | 5 | 3 | 6 | 4 |
| High | Q12905 | Interleukin enhancer-binding factor 2 OS=Homo sapiens OX=9606 GN=ILF2 PE=1 SV=2 | 57 | 10 | 10 | 43 | 37 | 390 | 5.26 | 5 | 4 | 6 | 5 | 4 | 5 | 5 | 4 | 6 | 4 | 3 | 4 |
| High | P17858 | ATP-dependent 6-phosphofructokinase, liver type OS=Homo sapiens OX=9606 GN=PFKL PE=1 SV=6 | 57 | 12 | 7 | 85 | 19 | 780 | 7.5 | 6 | 9 | 10 | 2 | 5 | 3 | 5 | 7 | 7 | 2 | 4 | 3 |
| High | Q99575 | Ribonucleases P/MRP protein subunit POP1 OS=Homo sapiens OX=9606 GN=POP1 PE=1 SV=2 | 57 | 17 | 17 | 114.6 | 24 | 1024 | 9.22 | 3 | 2 | 2 | 3 | 11 | 4 | 3 | 2 | 2 | 3 | 11 | 4 |
| High | Q9H223 | EH domain-containing protein 4 OS=Homo sapiens OX=9606 GN=EHD4 PE=1 SV=1 | 56 | 16 | 15 | 61.1 | 32 | 541 | 6.76 | 9 | 10 | 10 | 1 | | 3 | 9 | 9 | 10 | 1 | | 3 |
| High | O60313 | Dynamin-like 120 kDa protein, mitochondrial OS=Homo sapiens OX=9606 GN=OPA1 PE=1 SV=3 | 56 | 16 | 16 | 111.6 | 20 | 960 | 7.87 | 3 | 4 | 7 | 1 | 3 | 1 | 3 | 4 | 6 | 1 | 3 | 1 |
| High | P18077 | 60S ribosomal protein L35a OS=Homo sapiens OX=9606 GN=RPL35A PE=1 SV=2 | 56 | 6 | 6 | 12.5 | 39 | 110 | 11.06 | 4 | 3 | 7 | 7 | 6 | 8 | 3 | 2 | 5 | 4 | 4 | 4 |
| High | O95425 | Supervillin OS=Homo sapiens OX=9606 GN=SVIL PE=1 SV=2 | 56 | 23 | 23 | 247.6 | 14 | 2214 | 6.98 | 7 | 10 | 10 | | 2 | | 7 | 10 | 10 | | 2 | |
| High | Q13439 | Golgin subfamily A member 4 OS=Homo sapiens OX=9606 GN=GOLGA4 PE=1 SV=1 | 56 | 27 | 27 | 261 | 13 | 2230 | 5.39 | | | | 7 | 11 | 7 | | | | 7 | 11 | 7 |
| High | P08237 | ATP-dependent 6-phosphofructokinase, muscle type OS=Homo sapiens OX=9606 GN=PFKM PE=1 SV=2 | 56 | 14 | 11 | 85.1 | 19 | 780 | 7.99 | 6 | 7 | 9 | 4 | 3 | 3 | 6 | 7 | 9 | 4 | 3 | 3 |
| High | P36542 | ATP synthase subunit gamma, mitochondrial OS=Homo sapiens OX=9606 GN=ATP5F1C PE=1 SV=1 | 56 | 9 | 9 | 33 | 32 | 298 | 9.22 | 8 | 9 | 10 | 2 | 4 | 3 | 7 | 8 | 9 | 2 | 4 | 3 |
| High | Q13838 | Spliceosome RNA helicase DDX39B OS=Homo sapiens OX=9606 GN=DDX39B PE=1 SV=1 | 56 | 12 | 4 | 49 | 36 | 428 | 5.67 | 5 | 5 | 10 | 5 | 5 | 4 | 5 | 5 | 9 | 4 | 5 | 4 |
| High | P46776 | 60S ribosomal protein L27a OS=Homo sapiens OX=9606 GN=RPL27A PE=1 SV=2 | 56 | 5 | 5 | 16.6 | 28 | 148 | 11 | 6 | 5 | 7 | 5 | 7 | 5 | 4 | 4 | 5 | 4 | 5 | 5 |
| High | A6NHL2 | Tubulin alpha chain-like 3 OS=Homo sapiens OX=9606 GN=TUBAL3 PE=1 SV=2 | 55 | 4 | 1 | 49.9 | 9 | 446 | 6.05 | 6 | 8 | 6 | 5 | 6 | 5 | 4 | 4 | 4 | 3 | 3 | 3 |
| High | Q9UPV9 | Trafficking kinesin-binding protein 1 OS=Homo sapiens OX=9606 GN=TRAK1 PE=1 SV=1 | 55 | 15 | 15 | 106 | 21 | 953 | 5.85 | | | | 12 | 14 | 12 | | | | 12 | 13 | 12 |
| High | Q99836 | Myeloid differentiation primary response protein MyD88 OS=Homo sapiens OX=9606 GN=MYD88 PE=1 SV=1 | 55 | 11 | 11 | 33.2 | 43 | 296 | 6.15 | 5 | 6 | 4 | 7 | 8 | 4 | 5 | 6 | 4 | 7 | 8 | 4 |
| High | Q92616 | eIF-2-alpha kinase activator GCN1 OS=Homo sapiens OX=9606 GN=GCN1 PE=1 SV=7 | 55 | 24 | 24 | 292.5 | 11 | 2671 | 7.43 | 5 | 5 | 7 | 1 | 1 | | 5 | 5 | 7 | 1 | 1 | |
| High | Q9HC35 | Echinoderm microtubule-associated protein-like 4 OS=Homo sapiens OX=9606 GN=EML4 PE=1 SV=3 | 55 | 16 | 16 | 108.8 | 18 | 981 | 6.4 | 1 | 4 | 3 | 6 | 4 | 7 | 1 | 4 | 3 | 6 | 4 | 7 |
| High | Q969S8 | Polyamine deacetylase HDAC10 OS=Homo sapiens OX=9606 GN=HDAC10 PE=1 SV=1 | 55 | 9 | 9 | 71.4 | 24 | 669 | 5.74 | 6 | 6 | 7 | 3 | 4 | 4 | 6 | 6 | 5 | 3 | 4 | 4 |
| High | Q13547 | Histone deacetylase 1 OS=Homo sapiens OX=9606 GN=HDAC1 PE=1 SV=1 | 55 | 10 | 6 | 55.1 | 29 | 482 | 5.48 | 5 | 4 | 7 | 3 | 7 | 6 | 5 | 4 | 7 | 3 | 4 | 5 |
| High | P47897 | Glutamine--tRNA ligase OS=Homo sapiens OX=9606 GN=QARS1 PE=1 SV=1 | 55 | 15 | 15 | 87.7 | 22 | 775 | 7.15 | 6 | 6 | 4 | 2 | 6 | 3 | 6 | 6 | 4 | 2 | 5 | 2 |
| High | P00492 | Hypoxanthine-guanine phosphoribosyltransferase OS=Homo sapiens OX=9606 GN=HPRT1 PE=1 SV=2 | 55 | 8 | 8 | 24.6 | 52 | 218 | 6.68 | 5 | 4 | 7 | 6 | 8 | 6 | 5 | 4 | 6 | 5 | 6 | 6 |
| High | Q96DG6 | Carboxymethylenebutenolidase homolog OS=Homo sapiens OX=9606 GN=CMBL PE=1 SV=1 | 55 | 8 | 8 | 28 | 29 | 245 | 7.18 | 6 | 4 | 5 | 7 | 5 | 5 | 5 | 4 | 4 | 6 | 5 | 5 |
| High | Q9UBL6 | Copine-7 OS=Homo sapiens OX=9606 GN=CPNE7 PE=1 SV=1 | 55 | 11 | 10 | 70.2 | 21 | 633 | 6.38 | 3 | 4 | 4 | 2 | 6 | 4 | 3 | 4 | 4 | 2 | 4 | 4 |
| High | Q9NYF8 | Bcl-2-associated transcription factor 1 OS=Homo sapiens OX=9606 GN=BCLAF1 PE=1 SV=2 | 55 | 11 | 10 | 106.1 | 18 | 920 | 9.98 | 5 | 5 | 3 | 6 | 9 | 4 | 4 | 5 | 3 | 5 | 8 | 4 |
| High | O14979 | Heterogeneous nuclear ribonucleoprotein D-like OS=Homo sapiens OX=9606 GN=HNRNPDL PE=1 SV=3 | 55 | 8 | 6 | 46.4 | 21 | 420 | 9.57 | 8 | 9 | 8 | 5 | 4 | 3 | 7 | 8 | 7 | 4 | 2 | 2 |
| High | Q13555 | Calcium/calmodulin-dependent protein kinase type II subunit gamma OS=Homo sapiens OX=9606 GN=CAMK2G PE=1 SV=4 | 55 | 13 | 10 | 62.6 | 30 | 558 | 7.83 | 9 | 7 | 6 | 3 | 4 | 4 | 9 | 7 | 6 | 3 | 4 | 4 |
| High | P69905 | Hemoglobin subunit alpha OS=Homo sapiens OX=9606 GN=HBA1 PE=1 SV=2 | 54 | 4 | 2 | 15.2 | 25 | 142 | 8.68 | 6 | 3 | 4 | 10 | 10 | 8 | 3 | 3 | 3 | 4 | 4 | 4 |
| High | P52943 | Cysteine-rich protein 2 OS=Homo sapiens OX=9606 GN=CRIP2 PE=1 SV=1 | 54 | 6 | 6 | 22.5 | 53 | 208 | 8.72 | 5 | 5 | 6 | 6 | 3 | 5 | 5 | 5 | 5 | 3 | 3 | 3 |
| High | P16989 | Y-box-binding protein 3 OS=Homo sapiens OX=9606 GN=YBX3 PE=1 SV=4 | 54 | 6 | 3 | 40.1 | 22 | 372 | 9.77 | 3 | 3 | 5 | 7 | 5 | 7 | 3 | 3 | 4 | 6 | 4 | 6 |
| High | P60866 | 40S ribosomal protein S20 OS=Homo sapiens OX=9606 GN=RPS20 PE=1 SV=1 | 54 | 5 | 5 | 13.4 | 29 | 119 | 9.94 | 6 | 6 | 5 | 5 | 6 | 5 | 5 | 4 | 4 | 3 | 4 | 3 |
| High | Q9Y490 | Talin-1 OS=Homo sapiens OX=9606 GN=TLN1 PE=1 SV=3 | 54 | 19 | 17 | 269.6 | 10 | 2541 | 6.07 | 3 | 2 | 6 | 3 | 4 | 5 | 3 | 2 | 6 | 3 | 4 | 5 |
| High | P30041 | Peroxiredoxin-6 OS=Homo sapiens OX=9606 GN=PRDX6 PE=1 SV=3 | 54 | 10 | 10 | 25 | 47 | 224 | 6.38 | 7 | 6 | 4 | 6 | 4 | 7 | 7 | 6 | 4 | 6 | 4 | 7 |
| High | Q9UKD2 | mRNA turnover protein 4 homolog OS=Homo sapiens OX=9606 GN=MRTO4 PE=1 SV=2 | 54 | 12 | 12 | 27.5 | 52 | 239 | 8.29 | 2 | 3 | 4 | 6 | 4 | 7 | 2 | 3 | 4 | 6 | 4 | 7 |
| High | Q9P206 | Uncharacterized protein KIAA1522 OS=Homo sapiens OX=9606 GN=KIAA1522 PE=1 SV=2 | 54 | 16 | 16 | 107 | 22 | 1035 | 9.73 | 7 | 2 | 4 | 3 | 4 | 3 | 7 | 2 | 4 | 3 | 4 | 3 |
| High | P49902 | Cytosolic purine 5'-nucleotidase OS=Homo sapiens OX=9606 GN=NT5C2 PE=1 SV=1 | 54 | 11 | 11 | 64.9 | 21 | 561 | 6.14 | 5 | 7 | 5 | 6 | 5 | 6 | 5 | 7 | 5 | 6 | 5 | 6 |
| High | P31153 | S-adenosylmethionine synthase isoform type-2 OS=Homo sapiens OX=9606 GN=MAT2A PE=1 SV=1 | 54 | 9 | 9 | 43.6 | 28 | 395 | 6.48 | 6 | 7 | 5 | 5 | 6 | 5 | 6 | 7 | 5 | 5 | 6 | 5 |
| High | P23588 | Eukaryotic translation initiation factor 4B OS=Homo sapiens OX=9606 GN=EIF4B PE=1 SV=2 | 54 | 7 | 7 | 69.1 | 13 | 611 | 5.73 | 5 | 5 | 7 | 5 | 6 | 4 | 3 | 4 | 5 | 5 | 6 | 4 |
| High | P38117 | Electron transfer flavoprotein subunit beta OS=Homo sapiens OX=9606 GN=ETFB PE=1 SV=3 | 54 | 10 | 10 | 27.8 | 43 | 255 | 8.1 | 7 | 9 | 5 | 5 | 3 | 4 | 7 | 9 | 5 | 5 | 3 | 4 |
| High | P46777 | 60S ribosomal protein L5 OS=Homo sapiens OX=9606 GN=RPL5 PE=1 SV=3 | 54 | 10 | 10 | 34.3 | 38 | 297 | 9.72 | 6 | 7 | 5 | 4 | 3 | 4 | 6 | 7 | 5 | 4 | 3 | 4 |
| High | Q8TD06 | Anterior gradient protein 3 OS=Homo sapiens OX=9606 GN=AGR3 PE=1 SV=1 | 54 | 2 | 1 | 19.2 | 13 | 166 | 7.99 | | | 1 | 13 | 19 | 14 | | 1 | | 1 | 1 | 1 |
| High | Q71UI9 | Histone H2A.V OS=Homo sapiens OX=9606 GN=H2AZ2 PE=1 SV=3 | 53 | 4 | 2 | 13.5 | 31 | 128 | 10.58 | 6 | 7 | 9 | 6 | 5 | 5 | 4 | 4 | 3 | 2 | 3 | 2 |
| High | Q9UKV8 | Protein argonaute-2 OS=Homo sapiens OX=9606 GN=AGO2 PE=1 SV=3 | 53 | 14 | 14 | 97.1 | 20 | 859 | 9.19 | 2 | 3 | 1 | 6 | 8 | 3 | 2 | 2 | 1 | 6 | 8 | 3 |
| High | O15226 | NF-kappa-B-repressing factor OS=Homo sapiens OX=9606 GN=NKRF PE=1 SV=2 | 53 | 13 | 13 | 77.6 | 21 | 690 | 8.79 | 2 | 1 | 3 | 6 | 8 | 7 | 2 | 1 | 3 | 6 | 8 | 7 |
| High | O75477 | Erlin-1 OS=Homo sapiens OX=9606 GN=ERLIN1 PE=1 SV=2 | 53 | 9 | 6 | 39.1 | 27 | 348 | 7.87 | 5 | 7 | 7 | 3 | 2 | 2 | 5 | 6 | 6 | 3 | 2 | 2 |
| High | Q16629 | Serine/arginine-rich splicing factor 7 OS=Homo sapiens OX=9606 GN=SRSF7 PE=1 SV=1 | 53 | 6 | 5 | 27.4 | 24 | 238 | 11.82 | 6 | 8 | 6 | 4 | 3 | 3 | 6 | 6 | 5 | 4 | 3 | 3 |
| High | O76021 | Ribosomal L1 domain-containing protein 1 OS=Homo sapiens OX=9606 GN=RSL1D1 PE=1 SV=3 | 53 | 10 | 10 | 54.9 | 20 | 490 | 10.13 | 3 | 3 | 4 | 5 | 7 | 6 | 3 | 3 | 4 | 5 | 7 | 6 |
| High | Q9H9L4 | KAT8 regulatory NSL complex subunit 2 OS=Homo sapiens OX=9606 GN=KANSL2 PE=1 SV=3 | 53 | 12 | 12 | 55 | 32 | 492 | 6.62 | | | | 7 | 4 | 8 | | | | 6 | 4 | 7 |
| High | P63173 | 60S ribosomal protein L38 OS=Homo sapiens OX=9606 GN=RPL38 PE=1 SV=2 | 53 | 4 | 4 | 8.2 | 41 | 70 | 10.1 | 8 | 10 | 7 | 4 | 5 | 4 | 4 | 4 | 4 | 2 | 3 | 2 |
| High | Q9UMN6 | Histone-lysine N-methyltransferase 2B OS=Homo sapiens OX=9606 GN=KMT2B PE=1 SV=1 | 53 | 17 | 13 | 293.3 | 9 | 2715 | 8.22 | | | | 7 | 8 | 5 | | | | 7 | 8 | 5 |
| High | Q14847 | LIM and SH3 domain protein 1 OS=Homo sapiens OX=9606 GN=LASP1 PE=1 SV=2 | 53 | 8 | 8 | 29.7 | 35 | 261 | 7.05 | 4 | 5 | 5 | 4 | 7 | 4 | 4 | 4 | 5 | 4 | 7 | 4 |
| High | Q9UKM9 | RNA-binding protein Raly OS=Homo sapiens OX=9606 GN=RALY PE=1 SV=1 | 53 | 9 | 9 | 32.4 | 31 | 306 | 9.17 | 4 | 5 | 5 | 6 | 5 | 7 | 4 | 5 | 5 | 6 | 4 | 7 |
| High | P63167 | Dynein light chain 1, cytoplasmic OS=Homo sapiens OX=9606 GN=DYNLL1 PE=1 SV=1 | 53 | 5 | 2 | 10.4 | 58 | 89 | 7.4 | 4 | 5 | 7 | 7 | 6 | 5 | 2 | 2 | 5 | 4 | 3 | 3 |
| High | O75439 | Mitochondrial-processing peptidase subunit beta OS=Homo sapiens OX=9606 GN=PMPCB PE=1 SV=2 | 53 | 9 | 8 | 54.3 | 21 | 489 | 6.83 | 5 | 5 | 3 | 7 | 7 | 6 | 5 | 5 | 3 | 6 | 6 | 5 |
| High | Q06124 | Tyrosine-protein phosphatase non-receptor type 11 OS=Homo sapiens OX=9606 GN=PTPN11 PE=1 SV=3 | 52 | 13 | 13 | 68 | 27 | 593 | 7.3 | 1 | 2 | 5 | 7 | 7 | 7 | 1 | 2 | 5 | 7 | 7 | 7 |
| High | P22061 | Protein-L-isoaspartate(D-aspartate) O-methyltransferase OS=Homo sapiens OX=9606 GN=PCMT1 PE=1 SV=4 | 52 | 10 | 10 | 24.6 | 54 | 227 | 7.21 | 7 | 8 | 6 | 3 | 4 | 2 | 7 | 8 | 4 | 3 | 4 | 2 |
| High | Q8IVT2 | Mitotic interactor and substrate of PLK1 OS=Homo sapiens OX=9606 GN=MISP PE=1 SV=1 | 52 | 17 | 17 | 75.3 | 36 | 679 | 6.83 | 4 | 6 | 10 | 2 | 2 | 4 | 4 | 6 | 10 | 2 | 2 | 4 |
| High | P32969 | 60S ribosomal protein L9 OS=Homo sapiens OX=9606 GN=RPL9 PE=1 SV=1 | 52 | 6 | 6 | 21.9 | 43 | 192 | 9.95 | 7 | 8 | 10 | 3 | 2 | 3 | 4 | 4 | 6 | 3 | 2 | 3 |
| High | P48444 | Coatomer subunit delta OS=Homo sapiens OX=9606 GN=ARCN1 PE=1 SV=1 | 52 | 13 | 13 | 57.2 | 29 | 511 | 6.21 | 6 | 8 | 6 | 4 | 4 | 3 | 6 | 8 | 6 | 4 | 4 | 3 |
| High | P07737 | Profilin-1 OS=Homo sapiens OX=9606 GN=PFN1 PE=1 SV=2 | 52 | 7 | 4 | 15 | 59 | 140 | 8.27 | 6 | 7 | 8 | 3 | 3 | 2 | 6 | 6 | 6 | 3 | 3 | 2 |
| High | P83731 | 60S ribosomal protein L24 OS=Homo sapiens OX=9606 GN=RPL24 PE=1 SV=1 | 52 | 5 | 5 | 17.8 | 31 | 157 | 11.25 | 3 | 4 | 6 | 5 | 7 | 6 | 3 | 4 | 5 | 4 | 5 | 5 |
| High | Q9NXV2 | BTB/POZ domain-containing protein KCTD5 OS=Homo sapiens OX=9606 GN=KCTD5 PE=1 SV=1 | 52 | 8 | 8 | 26.1 | 58 | 234 | 6.24 | 6 | 5 | 5 | 5 | 4 | 5 | 5 | 4 | 4 | 4 | 3 | 4 |
| High | Q9Y2H5 | Pleckstrin homology domain-containing family A member 6 OS=Homo sapiens OX=9606 GN=PLEKHA6 PE=1 SV=4 | 52 | 19 | 19 | 117.1 | 25 | 1048 | 9.1 | 4 | 3 | 8 | 2 | 2 | 1 | 4 | 3 | 8 | 2 | 2 | 1 |
| High | O14818 | Proteasome subunit alpha type-7 OS=Homo sapiens OX=9606 GN=PSMA7 PE=1 SV=1 | 52 | 9 | 9 | 27.9 | 46 | 248 | 8.46 | 4 | 5 | 5 | 8 | 8 | 6 | 4 | 5 | 4 | 7 | 7 | 5 |

| | | | | | | | | | | | | | | | | | | | | | |
|---|---|---|---|---|---|---|---|---|---|---|---|---|---|---|---|---|---|---|---|---|---|
| High | Q93050 | V-type proton ATPase 116 kDa subunit a 1 OS=Homo sapiens OX=9606 GN=ATP6V0A1 PE=1 SV=3 | 52 | 14 | 14 | 96.4 | 20 | 837 | 6.43 | 4 | 4 | 6 | | | | 4 | 4 | 5 | | | |
| High | O15144 | Actin-related protein 2/3 complex subunit 2 OS=Homo sapiens OX=9606 GN=ARPC2 PE=1 SV=1 | 52 | 11 | 11 | 34.3 | 34 | 300 | 7.36 | 7 | 5 | 5 | 4 | 5 | 4 | 7 | 5 | 5 | 4 | 5 | 4 |
| High | P26373 | 60S ribosomal protein L13 OS=Homo sapiens OX=9606 GN=RPL13 PE=1 SV=4 | 51 | 6 | 6 | 24.2 | 28 | 211 | 11.65 | 4 | 5 | 4 | 7 | 6 | 6 | 4 | 5 | 4 | 5 | 5 | 5 |
| High | Q92828 | Coronin-2A OS=Homo sapiens OX=9606 GN=CORO2A PE=1 SV=2 | 51 | 10 | 9 | 59.7 | 21 | 525 | 8.05 | 8 | 7 | 5 | 4 | 3 | 3 | 8 | 6 | 4 | 4 | 3 | 3 |
| High | Q9HC84 | Mucin-5B OS=Homo sapiens OX=9606 GN=MUC5B PE=1 SV=3 | 51 | 11 | 8 | 596 | 4 | 5762 | 6.64 | 2 | 2 | 1 | 10 | 13 | 11 | 2 | 2 | 1 | 7 | 10 | 8 |
| High | O43293 | Death-associated protein kinase 3 OS=Homo sapiens OX=9606 GN=DAPK3 PE=1 SV=1 | 51 | 13 | 13 | 52.5 | 36 | 454 | 6.89 | 9 | 8 | 10 | 1 | 1 | 2 | 9 | 7 | 9 | 1 | 1 | 2 |
| High | Q96FW1 | Ubiquitin thioesterase OTUB1 OS=Homo sapiens OX=9606 GN=OTUB1 PE=1 SV=2 | 51 | 8 | 8 | 31.3 | 42 | 271 | 4.94 | 9 | 5 | 5 | 5 | 4 | 4 | 7 | 5 | 5 | 5 | 4 | 4 |
| High | Q13162 | Peroxiredoxin-4 OS=Homo sapiens OX=9606 GN=PRDX4 PE=1 SV=1 | 51 | 4 | 2 | 30.5 | 13 | 271 | 6.29 | 4 | 4 | 4 | 6 | 10 | 8 | 2 | 2 | 2 | 3 | 4 | 4 |
| High | O95433 | Activator of 90 kDa heat shock protein ATPase homolog 1 OS=Homo sapiens OX=9606 GN=AHSA1 PE=1 SV=1 | 51 | 10 | 10 | 38.3 | 37 | 338 | 5.53 | 7 | 8 | 8 | 3 | 5 | 3 | 5 | 5 | 7 | 3 | 5 | 3 |
| High | Q9H4A4 | Aminopeptidase B OS=Homo sapiens OX=9606 GN=RNPEP PE=1 SV=2 | 51 | 14 | 14 | 72.5 | 27 | 650 | 5.74 | 5 | 4 | 5 | 4 | 6 | 1 | 5 | 4 | 5 | 4 | 6 | 1 |
| High | Q00325 | Phosphate carrier protein, mitochondrial OS=Homo sapiens OX=9606 GN=SLC25A3 PE=1 SV=2 | 51 | 8 | 8 | 40.1 | 25 | 362 | 9.38 | 6 | 8 | 6 | 4 | 5 | 4 | 6 | 7 | 6 | 4 | 5 | 4 |
| High | Q15717 | ELAV-like protein 1 OS=Homo sapiens OX=9606 GN=ELAVL1 PE=1 SV=2 | 51 | 8 | 8 | 36.1 | 30 | 326 | 9.17 | 7 | 5 | 6 | 2 | 4 | 3 | 7 | 5 | 6 | 2 | 4 | 3 |
| High | Q9BZF9 | Uveal autoantigen with coiled-coil domains and ankyrin repeats OS=Homo sapiens OX=9606 GN=UACA PE=1 SV=2 | 51 | 19 | 19 | 162.4 | 16 | 1416 | 7.03 | 8 | 9 | 10 | 1 | 1 | 1 | 8 | 9 | 10 | 1 | 1 | 1 |
| High | P04632 | Calpain small subunit 1 OS=Homo sapiens OX=9606 GN=CAPNS1 PE=1 SV=1 | 50 | 8 | 8 | 28.3 | 39 | 268 | 5.2 | 5 | 5 | 7 | 4 | 5 | 5 | 4 | 4 | 6 | 4 | 5 | 5 |
| High | Q16777 | Histone H2A type 2-C OS=Homo sapiens OX=9606 GN=H2AC20 PE=1 SV=4 | 50 | 3 | 1 | 14 | 27 | 129 | 10.9 | 5 | 6 | 10 | 5 | 4 | 4 | 3 | 3 | 2 | 1 | 2 | 1 |
| High | Q14683 | Structural maintenance of chromosomes protein 1A OS=Homo sapiens OX=9606 GN=SMC1A PE=1 SV=2 | 50 | 16 | 16 | 143.1 | 13 | 1233 | 7.64 | 4 | 3 | 4 | 2 | 2 | 7 | 4 | 3 | 4 | 2 | 2 | 7 |
| High | Q9H3U1 | Protein unc-45 homolog A OS=Homo sapiens OX=9606 GN=UNC45A PE=1 SV=1 | 50 | 14 | 14 | 103 | 18 | 944 | 6.07 | 8 | 6 | 9 | 2 | 2 | 2 | 7 | 6 | 9 | 2 | 2 | 2 |
| High | Q01844 | RNA-binding protein EWS OS=Homo sapiens OX=9606 GN=EWSR1 PE=1 SV=1 | 50 | 7 | 7 | 68.4 | 15 | 656 | 9.33 | 7 | 4 | 6 | 2 | 8 | 4 | 6 | 4 | 5 | 2 | 5 | 4 |
| High | O43583 | Density-regulated protein OS=Homo sapiens OX=9606 GN=DENR PE=1 SV=2 | 50 | 7 | 7 | 22.1 | 37 | 198 | 5.3 | 4 | 3 | 7 | 4 | 4 | 3 | 4 | 3 | 7 | 4 | 4 | 3 |
| High | P57721 | Poly(rC)-binding protein 3 OS=Homo sapiens OX=9606 GN=PCBP3 PE=1 SV=2 | 50 | 5 | 1 | 39.4 | 18 | 371 | 8.07 | 6 | 5 | 5 | 5 | 6 | 4 | 4 | 3 | 2 | 3 | 3 | 3 |
| High | O95630 | STAM-binding protein OS=Homo sapiens OX=9606 GN=STAMBP PE=1 SV=1 | 50 | 10 | 10 | 48 | 31 | 424 | 6.29 | 5 | 6 | 6 | | 1 | | 5 | 6 | 6 | | 1 | |
| High | P84085 | ADP-ribosylation factor 5 OS=Homo sapiens OX=9606 GN=ARF5 PE=1 SV=2 | 50 | 6 | 3 | 20.5 | 37 | 180 | 6.79 | 7 | 9 | 7 | 3 | 3 | 3 | 5 | 6 | 6 | 3 | 3 | 3 |
| High | P62191 | 26S proteasome regulatory subunit 4 OS=Homo sapiens OX=9606 GN=PSMC1 PE=1 SV=1 | 50 | 14 | 12 | 49.2 | 35 | 440 | 6.21 | | 1 | | 8 | 10 | 9 | | 1 | | 7 | 9 | 8 |
| High | P62633 | CCHC-type zinc finger nucleic acid binding protein OS=Homo sapiens OX=9606 GN=CNBP PE=1 SV=1 | 49 | 6 | 6 | 19.5 | 44 | 177 | 7.71 | 6 | 5 | 4 | 5 | 7 | 5 | 5 | 4 | 3 | 4 | 5 | 4 |
| High | Q9UJU6 | Drebrin-like protein OS=Homo sapiens OX=9606 GN=DBNL PE=1 SV=1 | 49 | 11 | 11 | 48.2 | 34 | 430 | 5.05 | 5 | 5 | 6 | 4 | 3 | 4 | 4 | 4 | 6 | 4 | 3 | 4 |
| High | Q9ULC4 | Malignant T-cell-amplified sequence 1 OS=Homo sapiens OX=9606 GN=MCTS1 PE=1 SV=1 | 49 | 7 | 7 | 20.5 | 44 | 181 | 8.82 | 5 | 4 | 3 | 4 | 5 | 5 | 4 | 4 | 3 | 3 | 5 | 4 |
| High | P54868 | Hydroxymethylglutaryl-CoA synthase, mitochondrial OS=Homo sapiens OX=9606 GN=HMGCS2 PE=1 SV=1 | 49 | 14 | 13 | 56.6 | 33 | 508 | 8.16 | | 1 | | 3 | 5 | 4 | | 1 | | 3 | 5 | 4 |
| High | P36952 | Serpin B5 OS=Homo sapiens OX=9606 GN=SERPINB5 PE=1 SV=2 | 49 | 10 | 10 | 42.1 | 31 | 375 | 6.05 | 8 | 7 | 6 | 3 | 5 | 3 | 8 | 7 | 6 | 3 | 5 | 3 |
| High | P61160 | Actin-related protein 2 OS=Homo sapiens OX=9606 GN=ACTR2 PE=1 SV=1 | 49 | 9 | 9 | 44.7 | 27 | 394 | 6.74 | 7 | 6 | 7 | 1 | 5 | 3 | 6 | 5 | 7 | 1 | 5 | 3 |
| High | Q14376 | UDP-glucose 4-epimerase OS=Homo sapiens OX=9606 GN=GALE PE=1 SV=2 | 49 | 10 | 10 | 38.3 | 49 | 348 | 6.73 | 4 | 5 | 9 | 2 | 2 | 2 | 4 | 5 | 9 | 2 | 2 | 2 |
| High | P61254 | 60S ribosomal protein L26 OS=Homo sapiens OX=9606 GN=RPL26 PE=1 SV=1 | 49 | 5 | 5 | 17.2 | 32 | 145 | 10.55 | 5 | 5 | 4 | 5 | 7 | 6 | 4 | 4 | 4 | 3 | 5 | 4 |
| High | P46063 | ATP-dependent DNA helicase Q1 OS=Homo sapiens OX=9606 GN=RECQL PE=1 SV=3 | 49 | 12 | 12 | 73.4 | 22 | 649 | 7.88 | 4 | 6 | 4 | 3 | 6 | 5 | 4 | 6 | 4 | 3 | 6 | 5 |
| High | Q00341 | Vigilin OS=Homo sapiens OX=9606 GN=HDLBP PE=1 SV=2 | 48 | 12 | 12 | 141.4 | 10 | 1268 | 6.87 | 5 | 3 | 4 | 4 | 6 | 3 | 5 | 3 | 4 | 4 | 5 | 3 |
| High | Q8WVV9 | Heterogeneous nuclear ribonucleoprotein L-like OS=Homo sapiens OX=9606 GN=HNRNPLL PE=1 SV=1 | 48 | 13 | 12 | 60 | 36 | 542 | 7.72 | 6 | 3 | 5 | 1 | 6 | 2 | 6 | 3 | 5 | 1 | 6 | 2 |
| High | Q96HC4 | PDZ and LIM domain protein 5 OS=Homo sapiens OX=9606 GN=PDLIM5 PE=1 SV=5 | 48 | 11 | 11 | 63.9 | 26 | 596 | 8.21 | 3 | 6 | 6 | 3 | 3 | 3 | 3 | 5 | 6 | 3 | 3 | 3 |
| High | Q9P2M7 | Cingulin OS=Homo sapiens OX=9606 GN=CGN PE=1 SV=3 | 48 | 16 | 16 | 137 | 16 | 1203 | 5.52 | 6 | 8 | 7 | 2 | 1 | 3 | 6 | 8 | 7 | 2 | 1 | 3 |
| High | O95757 | Heat shock 70 kDa protein 4L OS=Homo sapiens OX=9606 GN=HSPA4L PE=1 SV=3 | 48 | 15 | 12 | 94.5 | 25 | 839 | 5.88 | 2 | 3 | 4 | 1 | 5 | 1 | 2 | 3 | 4 | 1 | 4 | 1 |
| High | Q9HCE1 | Helicase MOV-10 OS=Homo sapiens OX=9606 GN=MOV10 PE=1 SV=2 | 48 | 14 | 14 | 113.6 | 16 | 1003 | 8.82 | 4 | 2 | 2 | 8 | 7 | 7 | 4 | 2 | 2 | 8 | 7 | 6 |
| High | Q96T37 | RNA-binding protein 15 OS=Homo sapiens OX=9606 GN=RBM15 PE=1 SV=2 | 48 | 9 | 9 | 107.1 | 11 | 977 | 10.08 | 5 | 3 | 5 | 5 | 3 | 5 | 5 | 3 | 5 | 5 | 3 | 5 |
| High | Q07020 | 60S ribosomal protein L18 OS=Homo sapiens OX=9606 GN=RPL18 PE=1 SV=2 | 48 | 6 | 6 | 21.6 | 35 | 188 | 11.72 | 2 | 4 | 5 | 5 | 5 | 5 | 2 | 3 | 4 | 5 | 5 | 4 |
| High | Q9Y5X1 | Sorting nexin-9 OS=Homo sapiens OX=9606 GN=SNX9 PE=1 SV=1 | 48 | 10 | 10 | 66.6 | 26 | 595 | 5.58 | 8 | 8 | 6 | 3 | 5 | 3 | 6 | 6 | 4 | 3 | 5 | 3 |
| High | Q15185 | Prostaglandin E synthase 3 OS=Homo sapiens OX=9606 GN=PTGES3 PE=1 SV=1 | 48 | 7 | 7 | 18.7 | 51 | 160 | 4.54 | 7 | 7 | 4 | 3 | 4 | 4 | 6 | 6 | 4 | 3 | 4 | 4 |
| High | Q96B97 | SH3 domain-containing kinase-binding protein 1 OS=Homo sapiens OX=9606 GN=SH3KBP1 PE=1 SV=2 | 48 | 10 | 10 | 73.1 | 17 | 665 | 6.62 | 5 | 5 | 5 | 5 | 3 | 1 | 4 | 4 | 4 | 4 | 3 | 1 |
| High | P60891 | Ribose-phosphate pyrophosphokinase 1 OS=Homo sapiens OX=9606 GN=PRPS1 PE=1 SV=2 | 48 | 8 | 4 | 34.8 | 31 | 318 | 6.98 | 4 | 6 | 7 | 5 | 4 | 5 | 4 | 6 | 7 | 5 | 4 | 5 |
| High | O60884 | DnaJ homolog subfamily A member 2 OS=Homo sapiens OX=9606 GN=DNAJA2 PE=1 SV=1 | 48 | 10 | 10 | 45.7 | 30 | 412 | 6.48 | 6 | 4 | 8 | 5 | 3 | 3 | 6 | 4 | 8 | 5 | 3 | 3 |
| High | Q9H0U4 | Ras-related protein Rab-1B OS=Homo sapiens OX=9606 GN=RAB1B PE=1 SV=1 | 48 | 8 | 6 | 22.2 | 48 | 201 | 5.73 | 7 | 8 | 8 | 2 | 4 | 3 | 5 | 6 | 7 | 2 | 3 | 3 |
| High | P62195 | 26S proteasome regulatory subunit 8 OS=Homo sapiens OX=9606 GN=PSMC5 PE=1 SV=1 | 48 | 11 | 9 | 45.6 | 33 | 406 | 7.55 | 1 | 3 | 2 | 6 | 8 | 6 | 1 | 3 | 2 | 5 | 6 | 5 |
| High | P22087 | rRNA 2'-O-methyltransferase fibrillarin OS=Homo sapiens OX=9606 GN=FBL PE=1 SV=2 | 48 | 10 | 10 | 33.8 | 36 | 321 | 10.18 | 6 | 6 | 8 | 1 | 6 | 3 | 6 | 6 | 8 | 1 | 5 | 3 |
| High | P04075 | Fructose-bisphosphate aldolase A OS=Homo sapiens OX=9606 GN=ALDOA PE=1 SV=2 | 47 | 15 | 14 | 39.4 | 41 | 364 | 8.09 | 7 | 7 | 11 | 2 | 5 | 4 | 7 | 7 | 9 | 2 | 5 | 4 |
| High | Q9Y6J0 | Calcineurin-binding protein cabin-1 OS=Homo sapiens OX=9606 GN=CABIN1 PE=1 SV=1 | 47 | 14 | 14 | 246.2 | 10 | 2220 | 6.02 | 6 | 9 | 6 | 2 | 2 | 2 | 6 | 9 | 6 | 2 | 2 | 2 |
| High | Q9NVH6 | Trimethyllysine dioxygenase, mitochondrial OS=Homo sapiens OX=9606 GN=TMLHE PE=1 SV=1 | 47 | 10 | 10 | 49.5 | 24 | 421 | 7.72 | 6 | 4 | 5 | 3 | 5 | 5 | 6 | 4 | 5 | 3 | 5 | 5 |
| High | P25786 | Proteasome subunit alpha type-1 OS=Homo sapiens OX=9606 GN=PSMA1 PE=1 SV=1 | 47 | 9 | 9 | 29.5 | 36 | 263 | 6.61 | 1 | 3 | 2 | 8 | 8 | 8 | 1 | 3 | 2 | 7 | 7 | 7 |
| High | P62269 | 40S ribosomal protein S18 OS=Homo sapiens OX=9606 GN=RPS18 PE=1 SV=3 | 47 | 8 | 8 | 17.7 | 48 | 152 | 10.99 | 4 | 4 | 9 | 2 | 4 | 4 | 4 | 4 | 7 | 2 | 4 | 3 |
| High | Q9HA64 | Ketosamine-3-kinase OS=Homo sapiens OX=9606 GN=FN3KRP PE=1 SV=2 | 47 | 8 | 8 | 34.4 | 28 | 309 | 7.33 | 6 | 5 | 6 | 3 | 5 | 5 | 5 | 5 | 6 | 3 | 4 | 5 |
| High | Q96QK1 | Vacuolar protein sorting-associated protein 35 OS=Homo sapiens OX=9606 GN=VPS35 PE=1 SV=2 | 47 | 12 | 12 | 91.6 | 19 | 796 | 5.49 | 6 | 8 | 7 | 2 | 6 | 4 | 6 | 8 | 7 | 2 | 6 | 4 |
| High | O60812 | Heterogeneous nuclear ribonucleoprotein C-like 1 OS=Homo sapiens OX=9606 GN=HNRNPCL1 PE=1 SV=1 | 47 | 6 | 1 | 32.1 | 16 | 293 | 5.06 | 5 | 5 | 5 | 4 | 4 | 5 | 5 | 4 | 4 | 4 | 4 | 4 |
| High | Q53H82 | Endoribonuclease LACTB2 OS=Homo sapiens OX=9606 GN=LACTB2 PE=1 SV=2 | 47 | 10 | 10 | 32.8 | 49 | 288 | 6.8 | 3 | 3 | 6 | 3 | 4 | 2 | 3 | 3 | 5 | 3 | 4 | 2 |
| High | P24534 | Elongation factor 1-beta OS=Homo sapiens OX=9606 GN=EEF1B2 PE=1 SV=3 | 47 | 6 | 5 | 24.7 | 44 | 225 | 4.67 | 4 | 6 | 6 | 2 | 2 | 2 | 4 | 5 | 5 | 2 | 2 | 2 |
| High | Q99661 | Kinesin-like protein KIF2C OS=Homo sapiens OX=9606 GN=KIF2C PE=1 SV=2 | 47 | 13 | 12 | 81.3 | 22 | 725 | 7.83 | 1 | 1 | 2 | 5 | 5 | 6 | 1 | 1 | 2 | 5 | 5 | 6 |
| High | P63208 | S-phase kinase-associated protein 1 OS=Homo sapiens OX=9606 GN=SKP1 PE=1 SV=2 | 47 | 7 | 7 | 18.6 | 51 | 163 | 4.54 | 4 | 4 | 4 | 4 | 7 | 4 | 4 | 4 | 4 | 3 | 6 | 4 |
| High | Q9H910 | Jupiter microtubule associated homolog 2 OS=Homo sapiens OX=9606 GN=JPT2 PE=1 SV=1 | 47 | 8 | 8 | 20.1 | 63 | 190 | 9.26 | 4 | 4 | 6 | 5 | 5 | 5 | 4 | 4 | 6 | 5 | 5 | 5 |
| High | Q13492 | Phosphatidylinositol-binding clathrin assembly protein OS=Homo sapiens OX=9606 GN=PICALM PE=1 SV=2 | 47 | 12 | 12 | 70.7 | 25 | 652 | 7.9 | 6 | 4 | 7 | 2 | 4 | 3 | 6 | 4 | 7 | 2 | 4 | 3 |
| High | P84103 | Serine/arginine-rich splicing factor 3 OS=Homo sapiens OX=9606 GN=SRSF3 PE=1 SV=1 | 47 | 6 | 5 | 19.3 | 35 | 164 | 11.65 | 3 | 5 | 4 | 3 | 3 | 4 | 3 | 3 | 3 | 3 | 3 | 4 |
| High | Q9BWF3 | RNA-binding protein 4 OS=Homo sapiens OX=9606 GN=RBM4 PE=1 SV=1 | 47 | 8 | 8 | 40.3 | 27 | 364 | 7.08 | 1 | 3 | 4 | 5 | 6 | 5 | 1 | 3 | 4 | 5 | 4 | 4 |
| High | Q9Y5S2 | Serine/threonine-protein kinase MRCK beta OS=Homo sapiens OX=9606 GN=CDC42BPB PE=1 SV=2 | 47 | 17 | 13 | 194.2 | 13 | 1711 | 6.37 | 3 | 5 | 4 | | | | 3 | 5 | 4 | | | |
| High | A0FGR8 | Extended synaptotagmin-2 OS=Homo sapiens OX=9606 GN=ESYT2 PE=1 SV=1 | 47 | 14 | 14 | 102.3 | 21 | 921 | 9.26 | 9 | 10 | 10 | 1 | 2 | | 9 | 10 | 9 | 1 | 2 | |
| High | Q8N680 | Zinc finger and BTB domain-containing protein 2 OS=Homo sapiens OX=9606 GN=ZBTB2 PE=1 SV=1 | 47 | 10 | 10 | 57.3 | 29 | 514 | 6.1 | | | | 6 | 5 | 5 | | | | 6 | 5 | 5 |
| High | Q13148 | TAR DNA-binding protein 43 OS=Homo sapiens OX=9606 GN=TARDBP PE=1 SV=1 | 47 | 8 | 8 | 44.7 | 28 | 414 | 6.19 | 5 | 5 | 6 | 3 | 4 | 3 | 5 | 5 | 5 | 3 | 4 | 2 |
| High | O75223 | Gamma-glutamylcyclotransferase OS=Homo sapiens OX=9606 GN=GGCT PE=1 SV=1 | 47 | 9 | 9 | 21 | 47 | 188 | 5.14 | 1 | 3 | 3 | 8 | 5 | 5 | 1 | 3 | 3 | 7 | 5 | 5 |
| High | Q9Y6M1 | Insulin-like growth factor 2 mRNA-binding protein 2 OS=Homo sapiens OX=9606 GN=IGF2BP2 PE=1 SV=2 | 46 | 10 | 8 | 66.1 | 22 | 599 | 8.46 | 2 | 2 | 7 | 4 | 4 | 3 | 2 | 2 | 7 | 4 | 4 | 3 |
| High | Q9NVP1 | ATP-dependent RNA helicase DDX18 OS=Homo sapiens OX=9606 GN=DDX18 PE=1 SV=2 | 46 | 11 | 11 | 75.4 | 20 | 670 | 9.5 | 5 | 4 | 4 | 2 | 3 | 2 | 5 | 4 | 4 | 2 | 3 | 2 |
| High | Q14444 | Caprin-1 OS=Homo sapiens OX=9606 GN=CAPRIN1 PE=1 SV=2 | 46 | 6 | 6 | 78.3 | 13 | 709 | 5.25 | 4 | 4 | 4 | 5 | 7 | 6 | 4 | 4 | 4 | 5 | 6 | 6 |
| High | P43246 | DNA mismatch repair protein Msh2 OS=Homo sapiens OX=9606 GN=MSH2 PE=1 SV=1 | 46 | 15 | 15 | 104.7 | 17 | 934 | 5.77 | 3 | 4 | 4 | 3 | 5 | 4 | 3 | 4 | 4 | 3 | 5 | 4 |
| High | Q14690 | Protein RRP5 homolog OS=Homo sapiens OX=9606 GN=PDCD11 PE=1 SV=3 | 46 | 17 | 17 | 208.6 | 10 | 1871 | 8.87 | 1 | | 2 | 3 | 3 | 2 | 1 | | 2 | 3 | 3 | 2 |
| High | P00367 | Glutamate dehydrogenase 1, mitochondrial OS=Homo sapiens OX=9606 GN=GLUD1 PE=1 SV=2 | 46 | 10 | 10 | 61.4 | 23 | 558 | 7.8 | 5 | 6 | 10 | 3 | 4 | 2 | 5 | 6 | 9 | 3 | 4 | 2 |
| High | Q96GQ7 | Probable ATP-dependent RNA helicase DDX27 OS=Homo sapiens OX=9606 GN=DDX27 PE=1 SV=2 | 46 | 15 | 15 | 89.8 | 20 | 796 | 9.28 | | | 1 | 2 | 8 | 4 | | | 1 | 2 | 8 | 4 |
| High | P35232 | Prohibitin 1 OS=Homo sapiens OX=9606 GN=PHB1 PE=1 SV=1 | 46 | 10 | 10 | 29.8 | 42 | 272 | 5.76 | 3 | 3 | 6 | 6 | 1 | 5 | 3 | 3 | 6 | 6 | 1 | 5 |
| High | P06730 | Eukaryotic translation initiation factor 4E OS=Homo sapiens OX=9606 GN=EIF4E PE=1 SV=2 | 46 | 8 | 8 | 25.1 | 34 | 217 | 6.15 | 4 | 4 | 5 | 5 | 4 | 5 | 4 | 3 | 4 | 5 | 4 | 5 |
| High | O43242 | 26S proteasome non-ATPase regulatory subunit 3 OS=Homo sapiens OX=9606 GN=PSMD3 PE=1 SV=2 | 46 | 13 | 13 | 60.9 | 24 | 534 | 8.44 | | | | 9 | 10 | 9 | | | | 9 | 10 | 9 |
| High | Q9BZK7 | F-box-like/WD repeat-containing protein TBL1XR1 OS=Homo sapiens OX=9606 GN=TBL1XR1 PE=1 SV=1 | 46 | 7 | 7 | 55.6 | 24 | 514 | 5.55 | 3 | 2 | 2 | 5 | 4 | 5 | 3 | 2 | 2 | 5 | 4 | 4 |
| High | Q07157 | Tight junction protein ZO-1 OS=Homo sapiens OX=9606 GN=TJP1 PE=1 SV=3 | 46 | 12 | 12 | 195.3 | 10 | 1748 | 6.7 | 6 | 5 | 6 | 4 | 5 | 2 | 6 | 5 | 6 | 4 | 5 | 2 |
| High | Q9BRX2 | Protein pelota homolog OS=Homo sapiens OX=9606 GN=PELO PE=1 SV=2 | 46 | 10 | 10 | 43.3 | 33 | 385 | 6.34 | 4 | 4 | 6 | 3 | 7 | 4 | 4 | 3 | 6 | 3 | 7 | 4 |
| High | Q9Y3B7 | 39S ribosomal protein L11, mitochondrial OS=Homo sapiens OX=9606 GN=MRPL11 PE=1 SV=1 | 46 | 11 | 11 | 20.7 | 54 | 192 | 9.91 | 4 | 4 | 2 | 4 | 4 | 5 | 4 | 4 | 2 | 4 | 4 | 5 |
| High | P08708 | 40S ribosomal protein S17 OS=Homo sapiens OX=9606 GN=RPS17 PE=1 SV=2 | 46 | 5 | 5 | 15.5 | 49 | 135 | 9.85 | 7 | 7 | 8 | 1 | 1 | 1 | 5 | 5 | 5 | 1 | 1 | 1 |
| High | Q9H0D6 | 5'-3' exoribonuclease 2 OS=Homo sapiens OX=9606 GN=XRN2 PE=1 SV=1 | 46 | 12 | 12 | 108.5 | 16 | 950 | 7.47 | 4 | 3 | 4 | 6 | 6 | 4 | 4 | 3 | 4 | 6 | 6 | 4 |
| High | P68036 | Ubiquitin-conjugating enzyme E2 L3 OS=Homo sapiens OX=9606 GN=UBE2L3 PE=1 SV=1 | 46 | 7 | 7 | 17.9 | 65 | 154 | 8.51 | 5 | 8 | 6 | 2 | 4 | 4 | 4 | 6 | 5 | 2 | 4 | 4 |
| High | P58546 | Myotrophin OS=Homo sapiens OX=9606 GN=MTPN PE=1 SV=2 | 45 | 5 | 5 | 12.9 | 55 | 118 | 5.52 | 5 | 4 | 5 | 4 | 5 | 2 | 4 | 3 | 3 | 3 | 3 | 1 |
| High | Q8NE71 | ATP-binding cassette sub-family F member 1 OS=Homo sapiens OX=9606 GN=ABCF1 PE=1 SV=2 | 45 | 13 | 13 | 95.9 | 21 | 845 | 6.8 | 4 | 3 | 1 | 5 | 4 | 4 | 4 | 3 | 1 | 5 | 4 | 4 |
| High | P82650 | 28S ribosomal protein S22, mitochondrial OS=Homo sapiens OX=9606 GN=MRPS22 PE=1 SV=1 | 45 | 9 | 9 | 41.3 | 26 | 360 | 7.9 | 1 | 1 | 2 | 4 | 7 | 5 | 1 | 1 | 2 | 4 | 7 | 5 |
| High | Q0VDF9 | Heat shock 70 kDa protein 14 OS=Homo sapiens OX=9606 GN=HSPA14 PE=1 SV=1 | 45 | 11 | 11 | 54.8 | 28 | 509 | 5.59 | 4 | 2 | 8 | 6 | 6 | 4 | 4 | 2 | 7 | 5 | 6 | 4 |
| High | Q9Y3U8 | 60S ribosomal protein L36 OS=Homo sapiens OX=9606 GN=RPL36 PE=1 SV=3 | 45 | 4 | 4 | 12.2 | 30 | 105 | 11.59 | 4 | 4 | 4 | 5 | 5 | 5 | 3 | 3 | 3 | 4 | 3 | 4 |
| High | Q92974 | Rho guanine nucleotide exchange factor 2 OS=Homo sapiens OX=9606 GN=ARHGEF2 PE=1 SV=4 | 45 | 16 | 16 | 111.5 | 18 | 986 | 7.27 | 10 | 9 | 8 | 2 | | 1 | 10 | 8 | 8 | 2 | | 1 |
| High | Q658Y4 | Protein FAM91A1 OS=Homo sapiens OX=9606 GN=FAM91A1 PE=1 SV=3 | 45 | 11 | 11 | 93.9 | 16 | 838 | 6.39 | 5 | 4 | 5 | 2 | 2 | 2 | 5 | 4 | 5 | 2 | 2 | 2 |
| High | Q9NZT2 | Opioid growth factor receptor OS=Homo sapiens OX=9606 GN=OGFR PE=1 SV=3 | 45 | 10 | 10 | 73.3 | 21 | 677 | 4.84 | 5 | 5 | 4 | 4 | 6 | 4 | 5 | 5 | 4 | 4 | 6 | 4 |
| High | P17096 | High mobility group protein HMG-I/HMG-Y OS=Homo sapiens OX=9606 GN=HMGA1 PE=1 SV=3 | 45 | 8 | 8 | 11.7 | 41 | 107 | 10.32 | 6 | 6 | 10 | 3 | 5 | 2 | 5 | 5 | 7 | 3 | 5 | 2 |
| High | Q14566 | DNA replication licensing factor MCM6 OS=Homo sapiens OX=9606 GN=MCM6 PE=1 SV=1 | 45 | 16 | 16 | 92.8 | 23 | 821 | 5.41 | 7 | 13 | 6 | 3 | 3 | 5 | 7 | 13 | 6 | 3 | 3 | 5 |
| High | Q9H773 | dCTP pyrophosphatase 1 OS=Homo sapiens OX=9606 GN=DCTPP1 PE=1 SV=1 | 45 | 10 | 10 | 18.7 | 59 | 170 | 5.03 | 4 | 3 | 7 | 3 | 3 | 4 | 4 | 3 | 7 | 3 | 3 | 4 |
| High | Q08945 | FACT complex subunit SSRP1 OS=Homo sapiens OX=9606 GN=SSRP1 PE=1 SV=1 | 45 | 13 | 13 | 81 | 25 | 709 | 6.87 | 6 | 6 | 9 | 1 | 4 | 1 | 5 | 5 | 9 | 1 | 4 | 1 |
| High | P31942 | Heterogeneous nuclear ribonucleoprotein H3 OS=Homo sapiens OX=9606 GN=HNRNPH3 PE=1 SV=2 | 45 | 6 | 5 | 36.9 | 25 | 346 | 6.87 | 8 | 6 | 4 | 3 | 5 | 4 | 6 | 5 | 4 | 3 | 4 | 4 |
| High | P62854 | 40S ribosomal protein S26 OS=Homo sapiens OX=9606 GN=RPS26 PE=1 SV=3 | 45 | 4 | 4 | 13 | 44 | 115 | 11 | 2 | 3 | 5 | 3 | 3 | 2 | 2 | 2 | 3 | 2 | 2 | 2 |
| High | P53041 | Serine/threonine-protein phosphatase 5 OS=Homo sapiens OX=9606 GN=PPP5C PE=1 SV=1 | 45 | 10 | 10 | 56.8 | 24 | 499 | 6.28 | 4 | 3 | 3 | 3 | 5 | 4 | 4 | 3 | 3 | 3 | 5 | 4 |
| High | P25789 | Proteasome subunit alpha type-4 OS=Homo sapiens OX=9606 GN=PSMA4 PE=1 SV=1 | 45 | 8 | 8 | 29.5 | 39 | 261 | 7.72 | 5 | 3 | 4 | 4 | 8 | 4 | 3 | 2 | 4 | 4 | 6 | 4 |
| High | Q9ULZ3 | Apoptosis-associated speck-like protein containing a CARD OS=Homo sapiens OX=9606 GN=PYCARD PE=1 SV=2 | 44 | 8 | 8 | 21.6 | 42 | 195 | 6.34 | 5 | 7 | 6 | 3 | 5 | 4 | 5 | 7 | 5 | 3 | 5 | 4 |
| High | Q7L5N7 | Lysophosphatidylcholine acyltransferase 2 OS=Homo sapiens OX=9606 GN=LPCAT2 PE=1 SV=1 | 44 | 10 | 10 | 60.2 | 26 | 544 | 6.55 | 4 | 5 | 4 | 3 | 3 | 4 | 4 | 5 | 4 | 3 | 3 | 4 |
| High | Q92734 | Protein TFG OS=Homo sapiens OX=9606 GN=TFG PE=1 SV=2 | 44 | 8 | 8 | 43.4 | 33 | 400 | 5.1 | 7 | 8 | 8 | 3 | 3 | 2 | 7 | 8 | 7 | 3 | 3 | 2 |
| High | P11277 | Spectrin beta chain, erythrocytic OS=Homo sapiens OX=9606 GN=SPTB PE=1 SV=5 | 44 | 8 | 1 | 246.3 | 5 | 2137 | 5.27 | 8 | 10 | 5 | 3 | 1 | 1 | 5 | 6 | 5 | 3 | 1 | 1 |
| High | Q16881 | Thioredoxin reductase 1, cytoplasmic OS=Homo sapiens OX=9606 GN=TXNRD1 PE=1 SV=3 | 44 | 10 | 10 | 70.9 | 20 | 649 | 7.39 | 2 | 5 | 5 | 5 | 2 | 4 | 2 | 5 | 5 | 5 | 2 | 4 |
| High | Q99460 | 26S proteasome non-ATPase regulatory subunit 1 OS=Homo sapiens OX=9606 GN=PSMD1 PE=1 SV=2 | 44 | 12 | 12 | 105.8 | 18 | 953 | 5.39 | | | | 6 | 9 | 7 | | | | 6 | 8 | 7 |
| High | Q68CZ2 | Tensin-3 OS=Homo sapiens OX=9606 GN=TNS3 PE=1 SV=2 | 44 | 7 | 6 | 155.2 | 8 | 1445 | 6.81 | 4 | 6 | 6 | 3 | 4 | 2 | 3 | 4 | 5 | 3 | 4 | 2 |
| High | P57740 | Nuclear pore complex protein Nup107 OS=Homo sapiens OX=9606 GN=NUP107 PE=1 SV=1 | 44 | 13 | 13 | 106.3 | 19 | 925 | 5.43 | 5 | 2 | 4 | 1 | | 1 | 3 | 2 | 4 | 1 | | 1 |
| High | P60174 | Triosephosphate isomerase OS=Homo sapiens OX=9606 GN=TPI1 PE=1 SV=4 | 44 | 9 | 9 | 26.7 | 49 | 249 | 6.9 | 7 | 6 | 9 | 3 | 3 | 2 | 6 | 5 | 8 | 3 | 3 | 2 |
| High | Q9H6T0 | Epithelial splicing regulatory protein 2 OS=Homo sapiens OX=9606 GN=ESRP2 PE=1 SV=1 | 44 | 10 | 8 | 78.4 | 17 | 727 | 6.71 | 2 | 3 | 5 | 3 | 3 | 3 | 2 | 3 | 4 | 3 | 2 | 3 |
| High | Q15652 | Probable JmjC domain-containing histone demethylation protein 2C OS=Homo sapiens OX=9606 GN=JMJD1C PE=1 SV=2 | 44 | 16 | 16 | 284.3 | 7 | 2540 | 7.87 | | | | 6 | 10 | 7 | | | | 6 | 10 | 7 |
| High | P48047 | ATP synthase subunit O, mitochondrial OS=Homo sapiens OX=9606 GN=ATP5PO PE=1 SV=1 | 44 | 8 | 8 | 23.3 | 45 | 213 | 9.96 | 4 | 4 | 6 | 5 | 5 | 6 | 4 | 4 | 5 | 5 | 4 | 6 |
| High | P20042 | Eukaryotic translation initiation factor 2 subunit 2 OS=Homo sapiens OX=9606 GN=EIF2S2 PE=1 SV=2 | 44 | 10 | 10 | 38.4 | 35 | 333 | 5.8 | 2 | 3 | 3 | 5 | 6 | 4 | 2 | 3 | 3 | 5 | 4 | 4 |
| High | Q99961 | Endophilin-A2 OS=Homo sapiens OX=9606 GN=SH3GL1 PE=1 SV=1 | 43 | 9 | 9 | 41.5 | 26 | 368 | 5.43 | 3 | 2 | 2 | 4 | 7 | 4 | 3 | 2 | 2 | 4 | 7 | 4 |
| High | Q02880 | DNA topoisomerase 2-beta OS=Homo sapiens OX=9606 GN=TOP2B PE=1 SV=3 | 43 | 10 | 2 | 183.2 | 6 | 1626 | 8 | 6 | 7 | 5 | 2 | 5 | 2 | 6 | 7 | 5 | 2 | 4 | 2 |
| High | Q9P0L0 | Vesicle-associated membrane protein-associated protein A OS=Homo sapiens OX=9606 GN=VAPA PE=1 SV=3 | 43 | 6 | 5 | 27.9 | 37 | 249 | 8.62 | 4 | 4 | 6 | 2 | 4 | 2 | 4 | 4 | 4 | 1 | 3 | 2 |
| High | O00291 | Huntingtin-interacting protein 1 OS=Homo sapiens OX=9606 GN=HIP1 PE=1 SV=5 | 43 | 15 | 14 | 116.1 | 17 | 1037 | 5.3 | 2 | 2 | 3 | | 1 | | 2 | 2 | 3 | | 1 | |
| High | Q9UKK3 | Protein mono-ADP-ribosyltransferase PARP4 OS=Homo sapiens OX=9606 GN=PARP4 PE=1 SV=3 | 43 | 16 | 16 | 192.5 | 11 | 1724 | 5.66 | 4 | 3 | 4 | 5 | 7 | 3 | 4 | 3 | 4 | 5 | 7 | 3 |
| High | P46734 | Dual specificity mitogen-activated protein kinase kinase 3 OS=Homo sapiens OX=9606 GN=MAP2K3 PE=1 SV=2 | 43 | 9 | 9 | 39.3 | 31 | 347 | 7.43 | 6 | 5 | 5 | 2 | 5 | 4 | 6 | 5 | 4 | 2 | 5 | 4 |
| High | Q9H9H4 | Vacuolar protein sorting-associated protein 37B OS=Homo sapiens OX=9606 GN=VPS37B PE=1 SV=1 | 43 | 5 | 5 | 31.3 | 25 | 285 | 7.34 | 5 | 4 | 3 | 4 | 5 | 4 | 5 | 4 | 3 | 4 | 4 | 4 |
| High | Q02543 | 60S ribosomal protein L18a OS=Homo sapiens OX=9606 GN=RPL18A PE=1 SV=2 | 43 | 5 | 5 | 20.7 | 30 | 176 | 10.71 | 3 | 3 | 4 | 3 | 3 | 4 | 3 | 3 | 4 | 2 | 2 | 2 |
| High | O75362 | Zinc finger protein 217 OS=Homo sapiens OX=9606 GN=ZNF217 PE=1 SV=1 | 43 | 21 | 21 | 115.2 | 31 | 1048 | 8.48 | | | | | | 1 | | | | | | 1 |
| High | O15355 | Protein phosphatase 1G OS=Homo sapiens OX=9606 GN=PPM1G PE=1 SV=1 | 43 | 10 | 10 | 59.2 | 27 | 546 | 4.36 | | | 2 | 4 | 5 | 6 | | | 2 | 4 | 5 | 6 |
| High | P62910 | 60S ribosomal protein L32 OS=Homo sapiens OX=9606 GN=RPL32 PE=1 SV=2 | 43 | 6 | 6 | 15.9 | 39 | 135 | 11.33 | 5 | 4 | 3 | 3 | 3 | 4 | 4 | 3 | 2 | 2 | 2 | 3 |
| High | Q96H79 | Zinc finger CCCH-type antiviral protein 1-like OS=Homo sapiens OX=9606 GN=ZC3HAV1L PE=1 SV=2 | 43 | 6 | 6 | 32.9 | 29 | 300 | 8.13 | 3 | 3 | 7 | 3 | 4 | 5 | 3 | 3 | 6 | 2 | 4 | 4 |
| High | P25398 | 40S ribosomal protein S12 OS=Homo sapiens OX=9606 GN=RPS12 PE=1 SV=3 | 43 | 6 | 6 | 14.5 | 53 | 132 | 7.21 | 5 | 5 | 6 | 4 | 4 | 3 | 5 | 5 | 6 | 4 | 4 | 3 |
| High | P56545 | C-terminal-binding protein 2 OS=Homo sapiens OX=9606 GN=CTBP2 PE=1 SV=1 | 43 | 8 | 6 | 48.9 | 22 | 445 | 6.95 | 3 | 4 | 4 | 4 | 3 | 3 | 3 | 4 | 4 | 4 | 3 | 3 |
| High | P12268 | Inosine-5'-monophosphate dehydrogenase 2 OS=Homo sapiens OX=9606 GN=IMPDH2 PE=1 SV=2 | 43 | 8 | 7 | 55.8 | 18 | 514 | 6.9 | 3 | 2 | 4 | 2 | 4 | 4 | 3 | 2 | 4 | 2 | 4 | 4 |
| High | P19525 | Interferon-induced, double-stranded RNA-activated protein kinase OS=Homo sapiens OX=9606 GN=EIF2AK2 PE=1 SV=2 | 43 | 9 | 9 | 62.1 | 19 | 551 | 8.4 | 1 | 3 | 1 | 5 | 6 | 7 | 1 | 3 | 1 | 5 | 6 | 7 |
| High | Q12792 | Twinfilin-1 OS=Homo sapiens OX=9606 GN=TWF1 PE=1 SV=3 | 42 | 8 | 7 | 40.3 | 21 | 350 | 6.96 | 5 | 6 | 5 | 3 | 3 | 5 | 4 | 4 | 4 | 3 | 3 | 5 |
| High | O14976 | Cyclin-G-associated kinase OS=Homo sapiens OX=9606 GN=GAK PE=1 SV=2 | 42 | 17 | 17 | 143.1 | 17 | 1311 | 5.73 | 3 | 3 | 2 | | 2 | 1 | 3 | 3 | 2 | | 2 | 1 |
| High | P19367 | Hexokinase-1 OS=Homo sapiens OX=9606 GN=HK1 PE=1 SV=3 | 42 | 14 | 9 | 102.4 | 15 | 917 | 6.8 | 6 | 9 | 5 | 1 | 1 | | 6 | 9 | 5 | 1 | 1 | |
| High | Q9H0A0 | RNA cytidine acetyltransferase OS=Homo sapiens OX=9606 GN=NAT10 PE=1 SV=2 | 42 | 12 | 12 | 115.7 | 14 | 1025 | 8.27 | 1 | 2 | 4 | 4 | 5 | 2 | 1 | 2 | 4 | 4 | 5 | 2 |
| High | P29353 | SHC-transforming protein 1 OS=Homo sapiens OX=9606 GN=SHC1 PE=1 SV=4 | 42 | 10 | 10 | 62.8 | 26 | 583 | 6.44 | 2 | | 3 | 5 | 4 | 6 | 2 | | 3 | 5 | 4 | 6 |
| High | O75179 | Ankyrin repeat domain-containing protein 17 OS=Homo sapiens OX=9606 GN=ANKRD17 PE=1 SV=3 | 42 | 13 | 7 | 274.1 | 7 | 2603 | 6.52 | | | | 7 | 8 | 9 | | | | 7 | 8 | 9 |
| High | Q04828 | Aldo-keto reductase family 1 member C1 OS=Homo sapiens OX=9606 GN=AKR1C1 PE=1 SV=1 | 42 | 7 | 1 | 36.8 | 25 | 323 | 7.88 | 7 | 7 | 6 | 2 | 3 | 2 | 6 | 6 | 5 | 2 | 3 | 2 |
| High | P04844 | Dolichyl-diphosphooligosaccharide--protein glycosyltransferase subunit 2 OS=Homo sapiens OX=9606 GN=RPN2 PE=1 SV=3 | 42 | 12 | 12 | 69.2 | 30 | 631 | 5.69 | 8 | 9 | 3 | 3 | 2 | 1 | 8 | 9 | 3 | 3 | 2 | 1 |
| High | P08240 | Signal recognition particle receptor subunit alpha OS=Homo sapiens OX=9606 GN=SRPRA PE=1 SV=2 | 42 | 11 | 11 | 69.8 | 20 | 638 | 8.95 | 6 | 7 | 7 | 2 | 1 | 1 | 6 | 7 | 7 | 2 | 1 | 1 |
| High | Q5H9F3 | BCL-6 corepressor-like protein 1 OS=Homo sapiens OX=9606 GN=BCORL1 PE=1 SV=2 | 42 | 14 | 13 | 190.4 | 10 | 1785 | 7.85 | | | | 8 | 8 | 7 | | | | 8 | 8 | 7 |
| High | Q9Y4E8 | Ubiquitin carboxyl-terminal hydrolase 15 OS=Homo sapiens OX=9606 GN=USP15 PE=1 SV=3 | 42 | 12 | 12 | 112.3 | 14 | 981 | 5.22 | 5 | 2 | 3 | 4 | 7 | 4 | 5 | 2 | 3 | 4 | 7 | 4 |
| High | Q9BR76 | Coronin-1B OS=Homo sapiens OX=9606 GN=CORO1B PE=1 SV=1 | 42 | 9 | 9 | 54.2 | 20 | 489 | 5.88 | 4 | 5 | 5 | 3 | 4 | 6 | 4 | 5 | 4 | 3 | 4 | 6 |
| High | O75531 | Barrier-to-autointegration factor OS=Homo sapiens OX=9606 GN=BANF1 PE=1 SV=1 | 42 | 4 | 4 | 10.1 | 51 | 89 | 6.09 | 10 | 9 | 11 | 3 | 1 | 2 | 4 | 4 | 4 | 1 | 1 | 1 |
| High | P37108 | Signal recognition particle 14 kDa protein OS=Homo sapiens OX=9606 GN=SRP14 PE=1 SV=2 | 42 | 5 | 5 | 14.6 | 30 | 136 | 10.04 | 4 | 4 | 2 | 3 | 6 | 4 | 3 | 3 | 2 | 3 | 4 | 4 |
| High | O95487 | Protein transport protein Sec24B OS=Homo sapiens OX=9606 GN=SEC24B PE=1 SV=2 | 42 | 11 | 9 | 137.3 | 10 | 1268 | 6.67 | | | 1 | 3 | 5 | 3 | | | 1 | 3 | 5 | 3 |
| High | Q9UNM6 | 26S proteasome non-ATPase regulatory subunit 13 OS=Homo sapiens OX=9606 GN=PSMD13 PE=1 SV=2 | 42 | 11 | 11 | 42.9 | 29 | 376 | 5.81 | 1 | | 1 | 6 | 10 | 6 | 1 | | 1 | 6 | 10 | 6 |
| High | A0AVT1 | Ubiquitin-like modifier-activating enzyme 6 OS=Homo sapiens OX=9606 GN=UBA6 PE=1 SV=1 | 42 | 14 | 14 | 117.9 | 15 | 1052 | 6.14 | 5 | 5 | 3 | 2 | 2 | 4 | 5 | 5 | 3 | 2 | 2 | 4 |
| High | Q6P2Q9 | Pre-mRNA-processing-splicing factor 8 OS=Homo sapiens OX=9606 GN=PRPF8 PE=1 SV=2 | 42 | 18 | 18 | 273.4 | 9 | 2335 | 8.84 | 2 | 3 | 2 | 3 | 5 | 2 | 2 | 3 | 2 | 3 | 5 | 2 |
| High | Q06787 | Synaptic functional regulator FMR1 OS=Homo sapiens OX=9606 GN=FMR1 PE=1 SV=1 | 42 | 8 | 6 | 71.1 | 15 | 632 | 7.42 | 5 | 4 | 2 | 4 | 6 | 4 | 5 | 4 | 2 | 4 | 6 | 4 |
| High | Q8NFJ5 | Retinoic acid-induced protein 3 OS=Homo sapiens OX=9606 GN=GPRC5A PE=1 SV=2 | 42 | 5 | 5 | 40.2 | 14 | 357 | 8.15 | 5 | 5 | 7 | 1 | 3 | 1 | 4 | 4 | 5 | 1 | 3 | 1 |
| High | Q9Y2T2 | AP-3 complex subunit mu-1 OS=Homo sapiens OX=9606 GN=AP3M1 PE=1 SV=1 | 42 | 10 | 8 | 46.9 | 35 | 418 | 6.93 | 6 | 5 | 6 | 3 | 3 | 3 | 6 | 5 | 6 | 3 | 3 | 3 |
| High | P80188 | Neutrophil gelatinase-associated lipocalin OS=Homo sapiens OX=9606 GN=LCN2 PE=1 SV=2 | 41 | 6 | 6 | 22.6 | 40 | 198 | 8.91 | 4 | 3 | 4 | 2 | 4 | 3 | 3 | 2 | 3 | 1 | 3 | 2 |
| High | Q71UM5 | 40S ribosomal protein S27-like OS=Homo sapiens OX=9606 GN=RPS27L PE=1 SV=3 | 41 | 5 | 2 | 9.5 | 40 | 84 | 9.45 | 4 | 5 | 3 | 2 | 3 | 2 | 4 | 4 | 2 | 2 | 3 | 2 |
| High | P61962 | DDB1- and CUL4-associated factor 7 OS=Homo sapiens OX=9606 GN=DCAF7 PE=1 SV=1 | 41 | 9 | 9 | 38.9 | 30 | 342 | 5.52 | 1 | 2 | 3 | 5 | 6 | 4 | 1 | 2 | 3 | 4 | 6 | 4 |
| High | P06493 | Cyclin-dependent kinase 1 OS=Homo sapiens OX=9606 GN=CDK1 PE=1 SV=3 | 41 | 7 | 5 | 34.1 | 26 | 297 | 8.4 | 4 | 5 | 6 | 4 | 3 | 5 | 4 | 4 | 5 | 3 | 2 | 5 |
| High | P34897 | Serine hydroxymethyltransferase, mitochondrial OS=Homo sapiens OX=9606 GN=SHMT2 PE=1 SV=3 | 41 | 12 | 11 | 56 | 27 | 504 | 8.53 | 4 | 7 | 8 | 1 | 2 | 2 | 3 | 7 | 7 | 1 | 2 | 2 |
| High | P51148 | Ras-related protein Rab-5C OS=Homo sapiens OX=9606 GN=RAB5C PE=1 SV=2 | 41 | 7 | 4 | 23.5 | 39 | 216 | 8.41 | 4 | 6 | 6 | 2 | 4 | 2 | 4 | 6 | 6 | 2 | 4 | 2 |
| High | Q7KZI7 | Serine/threonine-protein kinase MARK2 OS=Homo sapiens OX=9606 GN=MARK2 PE=1 SV=2 | 41 | 11 | 8 | 87.9 | 18 | 788 | 9.72 | 5 | 4 | 5 | 1 | 1 | 1 | 5 | 4 | 5 | 1 | 1 | 1 |
| High | O95816 | BAG family molecular chaperone regulator 2 OS=Homo sapiens OX=9606 GN=BAG2 PE=1 SV=1 | 41 | 8 | 8 | 23.8 | 51 | 211 | 6.7 | 5 | 4 | 5 | 4 | 3 | 2 | 4 | 4 | 4 | 4 | 3 | 2 |
| High | Q9H0W9 | Ester hydrolase C11orf54 OS=Homo sapiens OX=9606 GN=C11orf54 PE=1 SV=1 | 41 | 7 | 7 | 35.1 | 30 | 315 | 6.7 | 5 | 6 | 4 | 2 | 3 | 3 | 5 | 6 | 4 | 2 | 3 | 3 |
| High | O14579 | Coatomer subunit epsilon OS=Homo sapiens OX=9606 GN=COPE PE=1 SV=3 | 41 | 7 | 7 | 34.5 | 27 | 308 | 5.12 | 5 | 5 | 4 | 1 | 1 | | 5 | 5 | 4 | 1 | 1 | |
| High | Q9NSE4 | Isoleucine--tRNA ligase, mitochondrial OS=Homo sapiens OX=9606 GN=IARS2 PE=1 SV=2 | 41 | 10 | 10 | 113.7 | 12 | 1012 | 7.2 | 10 | 9 | 8 | 1 | 1 | 1 | 9 | 9 | 8 | 1 | 1 | 1 |
| High | O00299 | Chloride intracellular channel protein 1 OS=Homo sapiens OX=9606 GN=CLIC1 PE=1 SV=4 | 41 | 10 | 10 | 26.9 | 61 | 241 | 5.17 | 4 | 6 | 5 | 2 | 4 | 2 | 3 | 5 | 5 | 2 | 4 | 2 |
| High | P33316 | Deoxyuridine 5'-triphosphate nucleotidohydrolase, mitochondrial OS=Homo sapiens OX=9606 GN=DUT PE=1 SV=4 | 41 | 6 | 6 | 26.5 | 31 | 252 | 9.36 | 5 | 5 | 4 | 3 | 6 | 4 | 4 | 4 | 4 | 3 | 5 | 4 |
| High | P60900 | Proteasome subunit alpha type-6 OS=Homo sapiens OX=9606 GN=PSMA6 PE=1 SV=1 | 41 | 8 | 8 | 27.4 | 36 | 246 | 6.76 | 3 | 1 | 1 | 6 | 11 | 6 | 3 | 1 | 1 | 5 | 8 | 5 |
| High | P22695 | Cytochrome b-c1 complex subunit 2, mitochondrial OS=Homo sapiens OX=9606 GN=UQCRC2 PE=1 SV=3 | 41 | 9 | 9 | 48.4 | 29 | 453 | 8.63 | 6 | 6 | 6 | 2 | 3 | 2 | 5 | 5 | 6 | 2 | 2 | 2 |
| High | P35573 | Glycogen debranching enzyme OS=Homo sapiens OX=9606 GN=AGL PE=1 SV=3 | 41 | 11 | 11 | 174.7 | 9 | 1532 | 6.76 | 2 | 1 | 2 | 3 | 5 | 3 | 2 | 1 | 2 | 3 | 5 | 3 |
| High | P46778 | 60S ribosomal protein L21 OS=Homo sapiens OX=9606 GN=RPL21 PE=1 SV=2 | 41 | 6 | 6 | 18.6 | 34 | 160 | 10.49 | 2 | 5 | 3 | 4 | 4 | 5 | 2 | 4 | 2 | 3 | 3 | 5 |

| | | | | | | | | | | | | | | | | | | | | | |
|---|---|---|---|---|---|---|---|---|---|---|---|---|---|---|---|---|---|---|---|---|---|
| High | Q5VWN6 | Protein TASOR 2 OS=Homo sapiens OX=9606 GN=TASOR2 PE=1 SV=1 | 41 | 16 | 16 | 268.7 | 8 | 2430 | 5.9 | | | | 5 | 6 | 7 | | | | 5 | 6 | 7 |
| High | Q9UEW8 | STE20/SPS1-related proline-alanine-rich protein kinase OS=Homo sapiens OX=9606 GN=STK39 PE=1 SV=3 | 41 | 11 | 7 | 59.4 | 26 | 545 | 6.29 | 6 | 5 | 5 | 2 | 3 | 3 | 6 | 5 | 5 | 2 | 3 | 3 |
| High | O14744 | Protein arginine N-methyltransferase 5 OS=Homo sapiens OX=9606 GN=PRMT5 PE=1 SV=4 | 41 | 12 | 12 | 72.6 | 22 | 637 | 6.29 | 4 | 2 | 2 | 3 | 6 | 5 | 4 | 2 | 2 | 3 | 6 | 4 |
| High | P12694 | 2-oxoisovalerate dehydrogenase subunit alpha, mitochondrial OS=Homo sapiens OX=9606 GN=BCKDHA PE=1 SV=2 | 41 | 9 | 9 | 50.4 | 25 | 445 | 8.27 | 6 | 5 | 6 | 2 | 3 | 4 | 5 | 4 | 6 | 2 | 3 | 4 |
| High | Q9P2R7 | Succinate--CoA ligase [ADP-forming] subunit beta, mitochondrial OS=Homo sapiens OX=9606 GN=SUCLA2 PE=1 SV=3 | 41 | 9 | 9 | 50.3 | 18 | 463 | 7.42 | 5 | 7 | 7 | 2 | 5 | 2 | 5 | 5 | 7 | 2 | 5 | 2 |
| High | P63000 | Ras-related C3 botulinum toxin substrate 1 OS=Homo sapiens OX=9606 GN=RAC1 PE=1 SV=1 | 41 | 6 | 5 | 21.4 | 33 | 192 | 8.5 | 3 | 5 | 4 | 3 | 4 | 3 | 3 | 4 | 3 | 3 | 4 | 3 |
| High | P35659 | Protein DEK OS=Homo sapiens OX=9606 GN=DEK PE=1 SV=1 | 41 | 10 | 10 | 42.6 | 33 | 375 | 8.56 | | | | 5 | 7 | 6 | | | | 5 | 6 | 6 |
| High | P23193 | Transcription elongation factor A protein 1 OS=Homo sapiens OX=9606 GN=TCEA1 PE=1 SV=2 | 41 | 9 | 9 | 33.9 | 35 | 301 | 8.38 | 1 | 1 | 4 | 4 | 4 | 5 | 1 | 1 | 4 | 4 | 4 | 5 |
| High | Q9UHD8 | Septin-9 OS=Homo sapiens OX=9606 GN=SEPTIN9 PE=1 SV=2 | 41 | 10 | 10 | 65.4 | 21 | 586 | 8.97 | 5 | 4 | 7 | 3 | 1 | 2 | 5 | 4 | 7 | 3 | 1 | 2 |
| High | P15927 | Replication protein A 32 kDa subunit OS=Homo sapiens OX=9606 GN=RPA2 PE=1 SV=1 | 41 | 7 | 7 | 29.2 | 46 | 270 | 6.15 | 6 | 6 | 8 | 4 | 4 | 3 | 5 | 5 | 6 | 4 | 4 | 3 |
| High | Q96QV6 | Histone H2A type 1-A OS=Homo sapiens OX=9606 GN=H2AC1 PE=1 SV=3 | 40 | 3 | 1 | 14.2 | 30 | 131 | 10.86 | 5 | 5 | 7 | 5 | 4 | 4 | 3 | 3 | 2 | 1 | 2 | 1 |
| High | P55212 | Caspase-6 OS=Homo sapiens OX=9606 GN=CASP6 PE=1 SV=2 | 40 | 8 | 7 | 33.3 | 29 | 293 | 6.93 | 2 | 4 | 6 | 4 | 4 | 5 | 2 | 4 | 6 | 4 | 4 | 5 |
| High | Q8NFW8 | N-acylneuraminate cytidylyltransferase OS=Homo sapiens OX=9606 GN=CMAS PE=1 SV=2 | 40 | 11 | 11 | 48.3 | 30 | 434 | 7.93 | 2 | 2 | 3 | 3 | 6 | 2 | 2 | 2 | 3 | 3 | 6 | 2 |
| High | Q7Z3B4 | Nucleoporin p54 OS=Homo sapiens OX=9606 GN=NUP54 PE=1 SV=2 | 40 | 10 | 10 | 55.4 | 22 | 507 | 7.02 | | | | 3 | 7 | 5 | | | | 3 | 7 | 4 |
| High | P14621 | Acylphosphatase-2 OS=Homo sapiens OX=9606 GN=ACYP2 PE=1 SV=2 | 40 | 5 | 5 | 11.1 | 46 | 99 | 9.5 | 3 | 4 | 5 | 5 | 4 | 4 | 3 | 4 | 5 | 5 | 4 | 4 |
| High | Q96I24 | Far upstream element-binding protein 3 OS=Homo sapiens OX=9606 GN=FUBP3 PE=1 SV=2 | 40 | 10 | 9 | 61.6 | 27 | 572 | 8.38 | 3 | 4 | 5 | 1 | 1 | 3 | 3 | 4 | 5 | 1 | 1 | 3 |
| High | Q9BZL6 | Serine/threonine-protein kinase D2 OS=Homo sapiens OX=9606 GN=PRKD2 PE=1 SV=3 | 40 | 12 | 12 | 96.7 | 19 | 878 | 6.84 | 6 | 7 | 9 | 1 | 2 | 2 | 6 | 6 | 9 | 1 | 2 | 2 |
| High | P62333 | 26S proteasome regulatory subunit 10B OS=Homo sapiens OX=9606 GN=PSMC6 PE=1 SV=1 | 40 | 10 | 10 | 44.1 | 31 | 389 | 7.49 | 2 | 2 | 2 | 4 | 6 | 6 | 2 | 2 | 2 | 4 | 6 | 6 |
| High | Q9NP81 | Serine--tRNA ligase, mitochondrial OS=Homo sapiens OX=9606 GN=SARS2 PE=1 SV=1 | 40 | 11 | 11 | 58.2 | 31 | 518 | 8.13 | 2 | 2 | 3 | 3 | 2 | 3 | 2 | 2 | 3 | 3 | 2 | 3 |
| High | O75746 | Calcium-binding mitochondrial carrier protein Aralar1 OS=Homo sapiens OX=9606 GN=SLC25A12 PE=1 SV=2 | 40 | 10 | 5 | 74.7 | 16 | 678 | 8.38 | 8 | 8 | 8 | 2 | 3 | 2 | 8 | 8 | 8 | 2 | 3 | 2 |
| High | Q9Y2T7 | Y-box-binding protein 2 OS=Homo sapiens OX=9606 GN=YBX2 PE=1 SV=2 | 40 | 4 | 1 | 38.5 | 14 | 364 | 10.8 | 2 | 2 | 4 | 5 | 3 | 5 | 2 | 2 | 3 | 4 | 2 | 4 |
| High | Q9UBX3 | Mitochondrial dicarboxylate carrier OS=Homo sapiens OX=9606 GN=SLC25A10 PE=1 SV=2 | 40 | 9 | 9 | 31.3 | 34 | 287 | 9.54 | 8 | 5 | 9 | 1 | 3 | 1 | 6 | 4 | 7 | 1 | 3 | 1 |
| High | Q9UQB8 | Brain-specific angiogenesis inhibitor 1-associated protein 2 OS=Homo sapiens OX=9606 GN=BAIAP2 PE=1 SV=1 | 40 | 11 | 11 | 60.8 | 23 | 552 | 8.9 | 3 | 5 | 5 | | 1 | 1 | 3 | 5 | 5 | | 1 | 1 |
| High | P48506 | Glutamate--cysteine ligase catalytic subunit OS=Homo sapiens OX=9606 GN=GCLC PE=1 SV=2 | 40 | 10 | 10 | 72.7 | 19 | 637 | 6.09 | 1 | 3 | 2 | 3 | 5 | 2 | 1 | 3 | 2 | 3 | 5 | 2 |
| High | Q9Y314 | Nitric oxide synthase-interacting protein OS=Homo sapiens OX=9606 GN=NOSIP PE=1 SV=1 | 40 | 8 | 8 | 33.2 | 36 | 301 | 8.82 | 1 | 2 | 2 | 7 | 5 | 6 | 1 | 2 | 2 | 6 | 5 | 6 |
| High | Q9H845 | Complex I assembly factor ACAD9, mitochondrial OS=Homo sapiens OX=9606 GN=ACAD9 PE=1 SV=1 | 40 | 14 | 14 | 68.7 | 23 | 621 | 7.96 | 4 | 5 | 6 | 1 | 2 | | 4 | 5 | 6 | 1 | 2 | |
| High | O75083 | WD repeat-containing protein 1 OS=Homo sapiens OX=9606 GN=WDR1 PE=1 SV=4 | 40 | 9 | 9 | 66.2 | 21 | 606 | 6.65 | 5 | 6 | 6 | 1 | 2 | 2 | 5 | 6 | 6 | 1 | 2 | 2 |
| High | Q7KZF4 | Staphylococcal nuclease domain-containing protein 1 OS=Homo sapiens OX=9606 GN=SND1 PE=1 SV=1 | 40 | 15 | 15 | 101.9 | 20 | 910 | 7.17 | 4 | 3 | 3 | 1 | 3 | 2 | 4 | 3 | 3 | 1 | 3 | 2 |
| High | P36969 | Phospholipid hydroperoxide glutathione peroxidase OS=Homo sapiens OX=9606 GN=GPX4 PE=1 SV=3 | 40 | 9 | 9 | 22.2 | 44 | 197 | 8.37 | 3 | 3 | 2 | 3 | 4 | 3 | 3 | 3 | 2 | 3 | 4 | 3 |
| High | O00763 | Acetyl-CoA carboxylase 2 OS=Homo sapiens OX=9606 GN=ACACB PE=1 SV=3 | 40 | 8 | 1 | 276.4 | 4 | 2458 | 6.49 | 5 | 5 | 4 | 4 | 2 | 4 | 5 | 5 | 4 | 4 | 2 | 4 |
| High | Q86X55 | Histone-arginine methyltransferase CARM1 OS=Homo sapiens OX=9606 GN=CARM1 PE=1 SV=3 | 39 | 9 | 9 | 65.8 | 17 | 608 | 6.73 | 3 | 3 | 3 | 4 | 5 | 4 | 3 | 3 | 3 | 4 | 5 | 4 |
| High | P09012 | U1 small nuclear ribonucleoprotein A OS=Homo sapiens OX=9606 GN=SNRPA PE=1 SV=3 | 39 | 6 | 4 | 31.3 | 27 | 282 | 9.83 | 4 | 4 | 3 | 4 | 4 | 5 | 4 | 4 | 3 | 4 | 3 | 5 |
| High | Q86X29 | Lipolysis-stimulated lipoprotein receptor OS=Homo sapiens OX=9606 GN=LSR PE=1 SV=4 | 39 | 9 | 9 | 71.4 | 18 | 649 | 7.97 | 3 | 3 | 7 | 1 | 1 | 1 | 3 | 3 | 7 | 1 | 1 | 1 |
| High | Q13151 | Heterogeneous nuclear ribonucleoprotein A0 OS=Homo sapiens OX=9606 GN=HNRNPA0 PE=1 SV=1 | 39 | 8 | 7 | 30.8 | 31 | 305 | 9.29 | 6 | 8 | 8 | 2 | 1 | 1 | 5 | 7 | 6 | 2 | 1 | 1 |
| High | Q53H96 | Pyrroline-5-carboxylate reductase 3 OS=Homo sapiens OX=9606 GN=PYCR3 PE=1 SV=3 | 39 | 7 | 7 | 28.6 | 36 | 274 | 7.72 | 5 | 3 | 3 | 3 | 3 | 3 | 4 | 3 | 2 | 3 | 3 | 3 |
| High | Q99829 | Copine-1 OS=Homo sapiens OX=9606 GN=CPNE1 PE=1 SV=1 | 39 | 9 | 9 | 59 | 17 | 537 | 5.83 | 5 | 5 | 4 | 3 | 2 | 4 | 5 | 5 | 4 | 3 | 2 | 4 |
| High | Q9GZP4 | PITH domain-containing protein 1 OS=Homo sapiens OX=9606 GN=PITHD1 PE=1 SV=1 | 39 | 8 | 8 | 24.2 | 52 | 211 | 5.74 | 4 | 5 | 6 | 2 | 3 | 3 | 4 | 5 | 5 | 2 | 3 | 3 |
| High | P06744 | Glucose-6-phosphate isomerase OS=Homo sapiens OX=9606 GN=GPI PE=1 SV=4 | 39 | 10 | 10 | 63.1 | 22 | 558 | 8.32 | 7 | 5 | 9 | | | | 6 | 5 | 8 | | | |
| High | P16401 | Histone H1.5 OS=Homo sapiens OX=9606 GN=H1-5 PE=1 SV=3 | 39 | 4 | 4 | 22.6 | 15 | 226 | 10.92 | 1 | 4 | 4 | 3 | 6 | 5 | 1 | 3 | 3 | 2 | 3 | 4 |
| High | O75608 | Acyl-protein thioesterase 1 OS=Homo sapiens OX=9606 GN=LYPLA1 PE=1 SV=1 | 39 | 5 | 5 | 24.7 | 28 | 230 | 6.77 | 4 | 4 | 5 | 3 | 4 | 4 | 4 | 4 | 5 | 3 | 4 | 4 |
| High | P08195 | 4F2 cell-surface antigen heavy chain OS=Homo sapiens OX=9606 GN=SLC3A2 PE=1 SV=3 | 39 | 10 | 10 | 68 | 22 | 630 | 5.01 | 7 | 3 | 6 | 4 | 5 | 4 | 7 | 3 | 6 | 4 | 4 | 4 |
| High | P41091 | Eukaryotic translation initiation factor 2 subunit 3 OS=Homo sapiens OX=9606 GN=EIF2S3 PE=1 SV=3 | 39 | 8 | 8 | 51.1 | 18 | 472 | 8.4 | 3 | 2 | 2 | 3 | 6 | 5 | 3 | 2 | 2 | 3 | 6 | 4 |
| High | Q56VL3 | OCIA domain-containing protein 2 OS=Homo sapiens OX=9606 GN=OCIAD2 PE=1 SV=1 | 39 | 6 | 6 | 16.9 | 38 | 154 | 9.03 | 6 | 5 | 4 | 3 | 4 | 3 | 5 | 5 | 4 | 3 | 4 | 3 |
| High | Q9UPN3 | Microtubule-actin cross-linking factor 1, isoforms 1/2/3/5 OS=Homo sapiens OX=9606 GN=MACF1 PE=1 SV=4 | 39 | 13 | 12 | 837.8 | 2 | 7388 | 5.39 | 3 | 5 | 4 | | 2 | 1 | 3 | 5 | 4 | | 2 | 1 |
| High | O60296 | Trafficking kinesin-binding protein 2 OS=Homo sapiens OX=9606 GN=TRAK2 PE=1 SV=2 | 39 | 17 | 17 | 101.4 | 26 | 914 | 5.24 | | | | 9 | 14 | 10 | | | | 9 | 13 | 10 |
| High | Q9HD42 | Charged multivesicular body protein 1a OS=Homo sapiens OX=9606 GN=CHMP1A PE=1 SV=1 | 39 | 6 | 6 | 21.7 | 25 | 196 | 8.06 | 5 | 4 | 6 | 3 | 3 | 3 | 5 | 4 | 5 | 3 | 3 | 3 |
| High | O95758 | Polypyrimidine tract-binding protein 3 OS=Homo sapiens OX=9606 GN=PTBP3 PE=1 SV=2 | 39 | 7 | 4 | 59.7 | 16 | 552 | 9.04 | 3 | 4 | 4 | 4 | 5 | 5 | 3 | 4 | 4 | 3 | 3 | 3 |
| High | Q9BXP5 | Serrate RNA effector molecule homolog OS=Homo sapiens OX=9606 GN=SRRT PE=1 SV=1 | 39 | 10 | 10 | 100.6 | 15 | 876 | 5.96 | 3 | 2 | 5 | 3 | 4 | 4 | 3 | 2 | 5 | 3 | 4 | 4 |
| High | Q96IZ0 | PRKC apoptosis WT1 regulator protein OS=Homo sapiens OX=9606 GN=PAWR PE=1 SV=1 | 39 | 8 | 8 | 36.5 | 43 | 340 | 5.41 | 5 | 4 | 7 | 1 | 4 | 1 | 5 | 4 | 5 | 1 | 4 | 1 |
| High | P53814 | Smoothelin OS=Homo sapiens OX=9606 GN=SMTN PE=1 SV=7 | 39 | 9 | 9 | 99 | 11 | 917 | 9.07 | 4 | 3 | 3 | 3 | 4 | 2 | 4 | 3 | 3 | 3 | 4 | 2 |
| High | P01966 | SWISS-PROT:P01966 (Bos taurus) Hemoglobin subunit alpha | 38 | 3 | 1 | 15.2 | 17 | 142 | 8.44 | 1 | 2 | 5 | 5 | 7 | 9 | 1 | 2 | 2 | 3 | 3 | 3 |
| High | Q96BW5 | Phosphotriesterase-related protein OS=Homo sapiens OX=9606 GN=PTER PE=1 SV=1 | 38 | 8 | 8 | 39 | 24 | 349 | 6.52 | 4 | 2 | 4 | 6 | 3 | 6 | 4 | 2 | 4 | 6 | 3 | 6 |
| High | Q9Y3Z3 | Deoxynucleoside triphosphate triphosphohydrolase SAMHD1 OS=Homo sapiens OX=9606 GN=SAMHD1 PE=1 SV=2 | 38 | 13 | 13 | 72.2 | 23 | 626 | 7.14 | 2 | 3 | 1 | 8 | 11 | 7 | 2 | 3 | 1 | 8 | 11 | 7 |
| High | P49207 | 60S ribosomal protein L34 OS=Homo sapiens OX=9606 GN=RPL34 PE=1 SV=3 | 38 | 6 | 6 | 13.3 | 37 | 117 | 11.47 | 3 | 3 | 6 | 3 | 3 | 3 | 3 | 3 | 5 | 3 | 3 | 3 |
| High | O60610 | Protein diaphanous homolog 1 OS=Homo sapiens OX=9606 GN=DIAPH1 PE=1 SV=2 | 38 | 12 | 12 | 141.3 | 11 | 1272 | 5.41 | 1 | 3 | 5 | | 3 | 1 | 1 | 3 | 5 | | 3 | 1 |
| High | O14524 | Nuclear envelope integral membrane protein 1 OS=Homo sapiens OX=9606 GN=NEMP1 PE=1 SV=2 | 38 | 10 | 10 | 50.6 | 19 | 444 | 6.93 | | | | | 1 | 1 | | | | | 1 | 1 |
| High | O95395 | Beta-1,3-galactosyl-O-glycosyl-glycoprotein beta-1,6-N-acetylglucosaminyltransferase 3 OS=Homo sapiens OX=9606 GN=GCNT3 PE | 38 | 11 | 11 | 50.8 | 33 | 438 | 8.25 | 11 | 10 | 10 | | | | 11 | 10 | 10 | | | |
| High | Q4G176 | Malonate--CoA ligase ACSF3, mitochondrial OS=Homo sapiens OX=9606 GN=ACSF3 PE=1 SV=3 | 38 | 10 | 10 | 64.1 | 22 | 576 | 8.37 | 3 | | 2 | 4 | 5 | 3 | 3 | | 2 | 4 | 5 | 3 |
| High | Q14166 | Tubulin--tyrosine ligase-like protein 12 OS=Homo sapiens OX=9606 GN=TTLL12 PE=1 SV=2 | 38 | 9 | 9 | 74.4 | 18 | 644 | 5.53 | 2 | 3 | 5 | 4 | 3 | 4 | 2 | 3 | 5 | 4 | 3 | 4 |
| High | O15014 | Zinc finger protein 609 OS=Homo sapiens OX=9606 GN=ZNF609 PE=1 SV=2 | 38 | 10 | 10 | 151.1 | 11 | 1411 | 8.03 | 2 | 2 | 3 | 3 | 6 | 3 | 2 | 2 | 3 | 3 | 6 | 3 |
| High | Q12933 | TNF receptor-associated factor 2 OS=Homo sapiens OX=9606 GN=TRAF2 PE=1 SV=2 | 38 | 10 | 10 | 55.8 | 23 | 501 | 7.53 | 4 | 3 | 4 | 2 | 3 | 1 | 4 | 3 | 4 | 2 | 3 | 1 |
| High | Q96FJ2 | Dynein light chain 2, cytoplasmic OS=Homo sapiens OX=9606 GN=DYNLL2 PE=1 SV=1 | 38 | 5 | 2 | 10.3 | 58 | 89 | 7.37 | 4 | 5 | 5 | 2 | 3 | 2 | 2 | 2 | 3 | 2 | 2 | 2 |
| High | O15269 | Serine palmitoyltransferase 1 OS=Homo sapiens OX=9606 GN=SPTLC1 PE=1 SV=1 | 38 | 9 | 9 | 52.7 | 25 | 473 | 6.01 | 5 | 5 | 4 | 2 | 1 | 1 | 5 | 5 | 4 | 2 | 1 | 1 |
| High | Q9Y6Y0 | Influenza virus NS1A-binding protein OS=Homo sapiens OX=9606 GN=IVNS1ABP PE=1 SV=3 | 38 | 10 | 10 | 71.7 | 17 | 642 | 5.53 | | | 1 | 6 | 6 | 6 | | | 1 | 6 | 6 | 6 |
| High | P49916 | DNA ligase 3 OS=Homo sapiens OX=9606 GN=LIG3 PE=1 SV=2 | 38 | 13 | 13 | 112.8 | 15 | 1009 | 9.01 | 2 | 1 | 2 | 2 | 3 | | 1 | 1 | 2 | 2 | 3 | |
| High | Q16630 | Cleavage and polyadenylation specificity factor subunit 6 OS=Homo sapiens OX=9606 GN=CPSF6 PE=1 SV=2 | 38 | 8 | 8 | 59.2 | 17 | 551 | 7.15 | 2 | 3 | 3 | 5 | 5 | 4 | 2 | 3 | 2 | 5 | 5 | 4 |
| High | P25189 | Myelin protein P0 OS=Homo sapiens OX=9606 GN=MPZ PE=1 SV=1 | 38 | 6 | 6 | 27.5 | 25 | 248 | 9.54 | | | | 11 | 15 | 12 | | | | 6 | 6 | 6 |
| High | Q15646 | 2'-5'-oligoadenylate synthase-like protein OS=Homo sapiens OX=9606 GN=OASL PE=1 SV=2 | 38 | 9 | 9 | 59.2 | 23 | 514 | 7.87 | 2 | 3 | 3 | 3 | 4 | 3 | 2 | 3 | 3 | 3 | 4 | 3 |
| High | Q01130 | Serine/arginine-rich splicing factor 2 OS=Homo sapiens OX=9606 GN=SRSF2 PE=1 SV=4 | 38 | 5 | 5 | 25.5 | 19 | 221 | 11.85 | 3 | 3 | 4 | 3 | 3 | 4 | 3 | 3 | 3 | 3 | 3 | 4 |
| High | Q13185 | Chromobox protein homolog 3 OS=Homo sapiens OX=9606 GN=CBX3 PE=1 SV=4 | 37 | 5 | 4 | 20.8 | 29 | 183 | 5.33 | 5 | 5 | 4 | 3 | 4 | 3 | 4 | 4 | 4 | 3 | 4 | 3 |
| High | A0A0B4J2 | Putative glutamine amidotransferase-like class 1 domain-containing protein 3B, mitochondrial OS=Homo sapiens OX=9606 GN=G | 37 | 9 | 9 | 28.1 | 56 | 268 | 8.27 | 4 | 4 | 4 | 2 | 6 | 2 | 3 | 3 | 4 | 2 | 4 | 2 |
| High | Q99729 | Heterogeneous nuclear ribonucleoprotein A/B OS=Homo sapiens OX=9606 GN=HNRNPAB PE=1 SV=2 | 37 | 6 | 5 | 36.2 | 21 | 332 | 8.21 | 6 | 5 | 5 | 3 | 3 | 3 | 5 | 4 | 4 | 2 | 2 | 2 |
| High | P51532 | Transcription activator BRG1 OS=Homo sapiens OX=9606 GN=SMARCA4 PE=1 SV=2 | 37 | 12 | 9 | 184.5 | 8 | 1647 | 7.88 | 1 | 2 | 4 | 3 | 3 | 4 | 1 | 2 | 4 | 3 | 3 | 4 |
| High | O00629 | Importin subunit alpha-3 OS=Homo sapiens OX=9606 GN=KPNA4 PE=1 SV=1 | 37 | 9 | 6 | 57.9 | 27 | 521 | 4.96 | | 2 | 3 | 3 | 3 | 2 | | 2 | 3 | 3 | 3 | 2 |
| High | Q16531 | DNA damage-binding protein 1 OS=Homo sapiens OX=9606 GN=DDB1 PE=1 SV=1 | 37 | 15 | 15 | 126.9 | 14 | 1140 | 5.26 | 1 | 3 | 1 | 8 | 5 | 8 | 1 | 3 | 1 | 8 | 5 | 8 |
| High | P04626 | Receptor tyrosine-protein kinase erbB-2 OS=Homo sapiens OX=9606 GN=ERBB2 PE=1 SV=1 | 37 | 14 | 13 | 137.8 | 15 | 1255 | 5.91 | 9 | 10 | 9 | | | | 8 | 9 | 9 | | | |
| High | Q92888 | Rho guanine nucleotide exchange factor 1 OS=Homo sapiens OX=9606 GN=ARHGEF1 PE=1 SV=2 | 37 | 10 | 10 | 102.4 | 14 | 912 | 5.66 | 6 | 5 | 6 | 1 | 1 | 1 | 6 | 5 | 6 | 1 | 1 | 1 |
| High | P42677 | 40S ribosomal protein S27 OS=Homo sapiens OX=9606 GN=RPS27 PE=1 SV=3 | 37 | 4 | 1 | 9.5 | 39 | 84 | 9.45 | 2 | 4 | 1 | 4 | 4 | 4 | 2 | 3 | 1 | 3 | 3 | 3 |
| High | Q14694 | Ubiquitin carboxyl-terminal hydrolase 10 OS=Homo sapiens OX=9606 GN=USP10 PE=1 SV=2 | 37 | 11 | 11 | 87.1 | 20 | 798 | 5.31 | 3 | 1 | 2 | 2 | 7 | 2 | 3 | 1 | 2 | 2 | 7 | 2 |
| High | P46779 | 60S ribosomal protein L28 OS=Homo sapiens OX=9606 GN=RPL28 PE=1 SV=3 | 37 | 4 | 4 | 15.7 | 28 | 137 | 12.02 | 4 | 3 | 4 | 4 | 4 | 3 | 3 | 3 | 4 | 3 | 3 | 2 |
| High | P40939 | Trifunctional enzyme subunit alpha, mitochondrial OS=Homo sapiens OX=9606 GN=HADHA PE=1 SV=2 | 37 | 11 | 11 | 82.9 | 21 | 763 | 9.04 | 4 | 3 | 2 | 4 | 5 | 5 | 4 | 3 | 2 | 4 | 5 | 5 |
| High | Q14684 | Ribosomal RNA processing protein 1 homolog B OS=Homo sapiens OX=9606 GN=RRP1B PE=1 SV=3 | 37 | 11 | 11 | 84.4 | 22 | 758 | 9.76 | | | 1 | 3 | 6 | 5 | | | 1 | 3 | 6 | 5 |
| High | Q96RQ3 | Methylcrotonoyl-CoA carboxylase subunit alpha, mitochondrial OS=Homo sapiens OX=9606 GN=MCCC1 PE=1 SV=3 | 37 | 8 | 8 | 80.4 | 17 | 725 | 7.78 | 5 | 5 | 6 | 2 | 3 | 2 | 5 | 4 | 6 | 2 | 3 | 2 |
| High | P50570 | Dynamin-2 OS=Homo sapiens OX=9606 GN=DNM2 PE=1 SV=2 | 37 | 11 | 6 | 98 | 15 | 870 | 7.44 | 8 | 6 | 6 | | | 2 | 8 | 6 | 6 | | | 2 |
| High | Q9BW92 | Threonine--tRNA ligase, mitochondrial OS=Homo sapiens OX=9606 GN=TARS2 PE=1 SV=1 | 37 | 9 | 9 | 81 | 13 | 718 | 7.3 | 1 | 2 | 3 | 1 | 6 | 4 | 1 | 2 | 3 | 1 | 6 | 4 |
| High | Q01081 | Splicing factor U2AF 35 kDa subunit OS=Homo sapiens OX=9606 GN=U2AF1 PE=1 SV=3 | 37 | 6 | 6 | 27.9 | 29 | 240 | 8.81 | 3 | 4 | 3 | 4 | 4 | 4 | 3 | 4 | 3 | 4 | 4 | 4 |
| High | O95347 | Structural maintenance of chromosomes protein 2 OS=Homo sapiens OX=9606 GN=SMC2 PE=1 SV=2 | 37 | 10 | 10 | 135.6 | 9 | 1197 | 8.43 | 2 | 3 | 6 | 3 | 4 | 3 | 2 | 3 | 6 | 3 | 4 | 3 |
| High | Q15029 | 116 kDa U5 small nuclear ribonucleoprotein component OS=Homo sapiens OX=9606 GN=EFTUD2 PE=1 SV=1 | 37 | 11 | 10 | 109.4 | 17 | 972 | 5 | 6 | 4 | 5 | 3 | 2 | 1 | 5 | 3 | 5 | 3 | 2 | 1 |
| High | O95336 | 6-phosphogluconolactonase OS=Homo sapiens OX=9606 GN=PGLS PE=1 SV=2 | 37 | 8 | 8 | 27.5 | 44 | 258 | 6.05 | 3 | 3 | 4 | 1 | 3 | 2 | 3 | 3 | 3 | 1 | 3 | 2 |
| High | P43487 | Ran-specific GTPase-activating protein OS=Homo sapiens OX=9606 GN=RANBP1 PE=1 SV=1 | 36 | 6 | 6 | 23.3 | 52 | 201 | 5.29 | 3 | 3 | 5 | 3 | 4 | 3 | 3 | 3 | 3 | 3 | 3 | 3 |
| High | Q00535 | Cyclin-dependent-like kinase 5 OS=Homo sapiens OX=9606 GN=CDK5 PE=1 SV=3 | 36 | 6 | 5 | 33.3 | 27 | 292 | 7.66 | 3 | 6 | 6 | 3 | 4 | 2 | 3 | 4 | 5 | 2 | 3 | 2 |
| High | Q16555 | Dihydropyrimidinase-related protein 2 OS=Homo sapiens OX=9606 GN=DPYSL2 PE=1 SV=1 | 36 | 9 | 9 | 62.3 | 22 | 572 | 6.38 | 4 | 3 | 3 | 5 | 3 | 5 | 4 | 3 | 3 | 5 | 3 | 5 |
| High | Q9UN86 | Ras GTPase-activating protein-binding protein 2 OS=Homo sapiens OX=9606 GN=G3BP2 PE=1 SV=2 | 36 | 8 | 8 | 54.1 | 20 | 482 | 5.55 | 2 | 2 | 2 | 5 | 6 | 5 | 2 | 2 | 2 | 5 | 6 | 5 |
| High | P61006 | Ras-related protein Rab-8A OS=Homo sapiens OX=9606 GN=RAB8A PE=1 SV=1 | 36 | 4 | 1 | 23.7 | 21 | 207 | 9.07 | 4 | 6 | 4 | 3 | 4 | 3 | 3 | 4 | 3 | 3 | 3 | 3 |
| High | O00139 | Kinesin-like protein KIF2A OS=Homo sapiens OX=9606 GN=KIF2A PE=1 SV=3 | 36 | 10 | 9 | 79.9 | 14 | 706 | 6.68 | 2 | | 2 | 5 | 3 | 4 | 2 | | 2 | 5 | 3 | 4 |
| High | P13987 | CD59 glycoprotein OS=Homo sapiens OX=9606 GN=CD59 PE=1 SV=1 | 36 | 4 | 4 | 14.2 | 25 | 128 | 6.48 | 5 | 5 | 4 | 2 | 2 | 2 | 4 | 4 | 3 | 2 | 2 | 2 |
| High | P62081 | 40S ribosomal protein S7 OS=Homo sapiens OX=9606 GN=RPS7 PE=1 SV=1 | 36 | 6 | 6 | 22.1 | 36 | 194 | 10.1 | 6 | 6 | 5 | 4 | 1 | 3 | 5 | 5 | 4 | 3 | 1 | 3 |
| High | P26639 | Threonine--tRNA ligase 1, cytoplasmic OS=Homo sapiens OX=9606 GN=TARS1 PE=1 SV=3 | 36 | 11 | 11 | 83.4 | 18 | 723 | 6.67 | 4 | 4 | 4 | | 1 | 3 | 4 | 4 | 4 | | 1 | 3 |
| High | Q3ZBS7 | TREMBL:Q3ZBS7 (Bos taurus) Vitronectin | 36 | 7 | 7 | 54.1 | 15 | 484 | 6.16 | 8 | 6 | 8 | 1 | 2 | 2 | 7 | 6 | 6 | 1 | 2 | 2 |
| High | Q96EY7 | Pentatricopeptide repeat domain-containing protein 3, mitochondrial OS=Homo sapiens OX=9606 GN=PTCD3 PE=1 SV=3 | 36 | 9 | 9 | 78.5 | 18 | 689 | 6.42 | 4 | 2 | 4 | 2 | 4 | 2 | 4 | 2 | 4 | 2 | 3 | 2 |
| High | P10916 | Myosin regulatory light chain 2, ventricular/cardiac muscle isoform OS=Homo sapiens OX=9606 GN=MYL2 PE=1 SV=3 | 36 | 9 | 9 | 18.8 | 52 | 166 | 5 | 5 | 4 | 4 | 6 | 2 | 6 | 4 | 4 | 4 | 6 | 2 | 6 |
| High | Q01518 | Adenylyl cyclase-associated protein 1 OS=Homo sapiens OX=9606 GN=CAP1 PE=1 SV=5 | 36 | 8 | 8 | 51.9 | 23 | 475 | 8.06 | 5 | 3 | 4 | 5 | 3 | 3 | 5 | 3 | 4 | 4 | 3 | 3 |
| High | Q9NVX2 | Notchless protein homolog 1 OS=Homo sapiens OX=9606 GN=NLE1 PE=1 SV=4 | 36 | 6 | 6 | 53.3 | 14 | 485 | 7.34 | 3 | 2 | 3 | 5 | 4 | 4 | 3 | 2 | 3 | 5 | 4 | 4 |
| High | Q9UBN7 | Histone deacetylase 6 OS=Homo sapiens OX=9606 GN=HDAC6 PE=1 SV=2 | 36 | 7 | 7 | 131.3 | 8 | 1215 | 5.3 | 3 | 3 | 3 | 3 | 4 | 4 | 3 | 3 | 3 | 3 | 3 | 4 |
| High | O15042 | U2 snRNP-associated SURP motif-containing protein OS=Homo sapiens OX=9606 GN=U2SURP PE=1 SV=2 | 36 | 10 | 10 | 118.2 | 14 | 1029 | 8.47 | 3 | 3 | 2 | 3 | 5 | 3 | 3 | 3 | 2 | 3 | 5 | 3 |
| High | P29966 | Myristoylated alanine-rich C-kinase substrate OS=Homo sapiens OX=9606 GN=MARCKS PE=1 SV=4 | 36 | 5 | 5 | 31.5 | 23 | 332 | 4.45 | 5 | 5 | 4 | 2 | 2 | 3 | 5 | 5 | 4 | 2 | 2 | 3 |
| High | Q13228 | Methanethiol oxidase OS=Homo sapiens OX=9606 GN=SELENBP1 PE=1 SV=2 | 35 | 11 | 11 | 52.4 | 24 | 472 | 6.37 | 3 | 3 | 3 | 3 | 6 | 5 | 3 | 3 | 2 | 3 | 6 | 5 |
| High | P28074 | Proteasome subunit beta type-5 OS=Homo sapiens OX=9606 GN=PSMB5 PE=1 SV=3 | 35 | 7 | 7 | 28.5 | 31 | 263 | 6.92 | 1 | 2 | 1 | 8 | 5 | 6 | 1 | 2 | 1 | 7 | 5 | 6 |
| High | Q8TAQ2 | SWI/SNF complex subunit SMARCC2 OS=Homo sapiens OX=9606 GN=SMARCC2 PE=1 SV=1 | 35 | 10 | 8 | 132.8 | 13 | 1214 | 5.69 | 2 | 1 | | 1 | 3 | 3 | 2 | 1 | | 1 | 3 | 3 |
| High | Q9Y224 | RNA transcription, translation and transport factor protein OS=Homo sapiens OX=9606 GN=RTRAF PE=1 SV=1 | 35 | 7 | 7 | 28.1 | 30 | 244 | 6.65 | 3 | 3 | 5 | 3 | 5 | 3 | 3 | 3 | 5 | 3 | 4 | 3 |
| High | Q15758 | Neutral amino acid transporter B(0) OS=Homo sapiens OX=9606 GN=SLC1A5 PE=1 SV=2 | 35 | 7 | 7 | 56.6 | 18 | 541 | 5.48 | 5 | 7 | 6 | 2 | 3 | 1 | 5 | 6 | 6 | 2 | 3 | 1 |
| High | Q99848 | Probable rRNA-processing protein EBP2 OS=Homo sapiens OX=9606 GN=EBNA1BP2 PE=1 SV=2 | 35 | 8 | 8 | 34.8 | 29 | 306 | 10.1 | 1 | 1 | 2 | 5 | 3 | 6 | 1 | 1 | 2 | 5 | 3 | 6 |
| High | Q9H3K6 | BolA-like protein 2 OS=Homo sapiens OX=9606 GN=BOLA2 PE=1 SV=1 | 35 | 4 | 4 | 10.1 | 62 | 86 | 6.52 | 3 | 5 | 4 | 4 | 4 | 2 | 3 | 4 | 4 | 3 | 3 | 2 |
| High | A0AV96 | RNA-binding protein 47 OS=Homo sapiens OX=9606 GN=RBM47 PE=1 SV=2 | 35 | 9 | 9 | 64.1 | 19 | 593 | 7.68 | 1 | 1 | | 3 | 3 | 3 | 1 | 1 | | 3 | 3 | 3 |
| High | Q9Y4W6 | AFG3-like protein 2 OS=Homo sapiens OX=9606 GN=AFG3L2 PE=1 SV=2 | 35 | 14 | 14 | 88.5 | 18 | 797 | 8.66 | 5 | 6 | 8 | 1 | 1 | | 5 | 6 | 8 | 1 | 1 | |
| High | Q9Y678 | Coatomer subunit gamma-1 OS=Homo sapiens OX=9606 GN=COPG1 PE=1 SV=1 | 35 | 10 | 10 | 97.7 | 15 | 874 | 5.47 | 3 | 3 | 7 | 2 | 1 | 2 | 3 | 3 | 5 | 2 | 1 | 2 |
| High | Q9UHV9 | Prefoldin subunit 2 OS=Homo sapiens OX=9606 GN=PFDN2 PE=1 SV=1 | 35 | 5 | 5 | 16.6 | 36 | 154 | 6.58 | 2 | 1 | 3 | 4 | 5 | 5 | 2 | 1 | 3 | 3 | 4 | 4 |
| High | Q96RP9 | Elongation factor G, mitochondrial OS=Homo sapiens OX=9606 GN=GFM1 PE=1 SV=2 | 35 | 10 | 10 | 83.4 | 17 | 751 | 7.01 | 2 | 2 | 2 | 2 | 3 | 3 | 2 | 2 | 2 | 2 | 3 | 3 |
| High | Q99426 | Tubulin-folding cofactor B OS=Homo sapiens OX=9606 GN=TBCB PE=1 SV=2 | 35 | 5 | 5 | 27.3 | 23 | 244 | 5.15 | 4 | 4 | 4 | 4 | 5 | 4 | 4 | 4 | 3 | 4 | 5 | 4 |
| High | Q15477 | Helicase SKI2W OS=Homo sapiens OX=9606 GN=SKIV2L PE=1 SV=3 | 35 | 14 | 14 | 137.7 | 16 | 1246 | 6.06 | 9 | 8 | 9 | 1 | | 1 | 9 | 8 | 9 | 1 | | 1 |
| High | P55884 | Eukaryotic translation initiation factor 3 subunit B OS=Homo sapiens OX=9606 GN=EIF3B PE=1 SV=3 | 35 | 8 | 8 | 92.4 | 12 | 814 | 5 | 5 | 4 | 4 | 2 | 2 | 2 | 5 | 4 | 4 | 2 | 2 | 2 |
| High | Q92769 | Histone deacetylase 2 OS=Homo sapiens OX=9606 GN=HDAC2 PE=1 SV=2 | 35 | 7 | 3 | 55.3 | 19 | 488 | 5.91 | 3 | 2 | 3 | 1 | 6 | 4 | 3 | 2 | 3 | 1 | 4 | 3 |
| High | P49790 | Nuclear pore complex protein Nup153 OS=Homo sapiens OX=9606 GN=NUP153 PE=1 SV=2 | 35 | 18 | 18 | 153.8 | 16 | 1475 | 8.73 | | | | 1 | 1 | 2 | | | | 1 | 1 | 2 |
| High | Q96CN7 | Isochorismatase domain-containing protein 1 OS=Homo sapiens OX=9606 GN=ISOC1 PE=1 SV=3 | 35 | 7 | 7 | 32.2 | 32 | 298 | 7.39 | 4 | 4 | 4 | 3 | 5 | 4 | 4 | 4 | 4 | 3 | 5 | 4 |
| High | Q9H4G0 | Band 4.1-like protein 1 OS=Homo sapiens OX=9606 GN=EPB41L1 PE=1 SV=2 | 35 | 13 | 13 | 98.4 | 20 | 881 | 5.62 | 5 | 6 | 6 | | 2 | | 5 | 6 | 6 | | 2 | |
| High | Q2TAA2 | Isoamyl acetate-hydrolyzing esterase 1 homolog OS=Homo sapiens OX=9606 GN=IAH1 PE=1 SV=1 | 34 | 6 | 6 | 27.6 | 31 | 248 | 5.3 | 4 | 4 | 2 | 3 | 3 | 2 | 4 | 4 | 2 | 3 | 3 | 2 |
| High | P63220 | 40S ribosomal protein S21 OS=Homo sapiens OX=9606 GN=RPS21 PE=1 SV=1 | 34 | 5 | 5 | 9.1 | 59 | 83 | 8.5 | 4 | 3 | 2 | 2 | 5 | 3 | 3 | 3 | 2 | 2 | 5 | 3 |
| High | Q8IX04 | Ubiquitin-conjugating enzyme E2 variant 3 OS=Homo sapiens OX=9606 GN=UEVLD PE=1 SV=2 | 34 | 7 | 7 | 52.2 | 19 | 471 | 7.09 | 3 | 4 | 5 | 3 | 3 | 4 | 3 | 4 | 5 | 3 | 3 | 4 |
| High | P07858 | Cathepsin B OS=Homo sapiens OX=9606 GN=CTSB PE=1 SV=3 | 34 | 5 | 5 | 37.8 | 15 | 339 | 6.3 | 5 | 3 | 5 | 4 | 3 | 3 | 4 | 3 | 5 | 4 | 3 | 3 |
| High | Q9Y3D2 | Methionine-R-sulfoxide reductase B2, mitochondrial OS=Homo sapiens OX=9606 GN=MSRB2 PE=1 SV=2 | 34 | 6 | 6 | 19.5 | 51 | 182 | 8.63 | 1 | 1 | 2 | 2 | 6 | 3 | 1 | 1 | 2 | 2 | 4 | 3 |
| High | Q99623 | Prohibitin-2 OS=Homo sapiens OX=9606 GN=PHB2 PE=1 SV=2 | 34 | 8 | 8 | 33.3 | 31 | 299 | 9.83 | 4 | 3 | 4 | 2 | 3 | 2 | 4 | 3 | 4 | 2 | 2 | 2 |
| High | P21980 | Protein-glutamine gamma-glutamyltransferase 2 OS=Homo sapiens OX=9606 GN=TGM2 PE=1 SV=2 | 34 | 11 | 11 | 77.3 | 20 | 687 | 5.22 | 1 | 3 | 1 | 6 | 8 | 7 | 1 | 3 | 1 | 6 | 8 | 7 |
| High | P26447 | Protein S100-A4 OS=Homo sapiens OX=9606 GN=S100A4 PE=1 SV=1 | 34 | 3 | 3 | 11.7 | 28 | 101 | 6.11 | 4 | 5 | 4 | 2 | 2 | 2 | 3 | 3 | 3 | 2 | 2 | 2 |
| High | Q8NEZ5 | F-box only protein 22 OS=Homo sapiens OX=9606 GN=FBXO22 PE=1 SV=1 | 34 | 7 | 7 | 44.5 | 19 | 403 | 7.03 | 5 | 4 | 4 | 4 | 3 | 3 | 4 | 3 | 4 | 4 | 3 | 3 |
| High | P49588 | Alanine--tRNA ligase, cytoplasmic OS=Homo sapiens OX=9606 GN=AARS1 PE=1 SV=2 | 34 | 14 | 14 | 106.7 | 21 | 968 | 5.53 | 2 | 3 | 9 | 1 | 1 | | 2 | 3 | 8 | 1 | 1 | |
| High | Q3ZCQ8 | Mitochondrial import inner membrane translocase subunit TIM50 OS=Homo sapiens OX=9606 GN=TIMM50 PE=1 SV=2 | 34 | 8 | 8 | 39.6 | 27 | 353 | 8.37 | 5 | 5 | 5 | 1 | 1 | 1 | 5 | 5 | 5 | 1 | 1 | 1 |
| High | Q9UK41 | Vacuolar protein sorting-associated protein 28 homolog OS=Homo sapiens OX=9606 GN=VPS28 PE=1 SV=1 | 34 | 8 | 8 | 25.4 | 49 | 221 | 5.54 | 1 | 3 | 5 | 3 | 4 | 3 | 1 | 3 | 5 | 3 | 3 | 3 |
| High | Q01780 | Exosome component 10 OS=Homo sapiens OX=9606 GN=EXOSC10 PE=1 SV=2 | 34 | 9 | 9 | 100.8 | 10 | 885 | 8.46 | | | | 5 | 5 | 4 | | | | 5 | 5 | 4 |
| High | Q8IY21 | Probable ATP-dependent RNA helicase DDX60 OS=Homo sapiens OX=9606 GN=DDX60 PE=1 SV=3 | 34 | 13 | 13 | 197.7 | 9 | 1712 | 7.59 | | 1 | 1 | 6 | 8 | 6 | | 1 | 1 | 6 | 8 | 6 |
| High | P50579 | Methionine aminopeptidase 2 OS=Homo sapiens OX=9606 GN=METAP2 PE=1 SV=1 | 34 | 2 | 2 | 52.9 | 9 | 478 | 5.82 | 7 | 8 | 3 | | 1 | 1 | 2 | 2 | 2 | | 1 | 1 |
| High | O14933 | Ubiquitin/ISG15-conjugating enzyme E2 L6 OS=Homo sapiens OX=9606 GN=UBE2L6 PE=1 SV=4 | 34 | 5 | 5 | 17.8 | 41 | 153 | 7.88 | 4 | 3 | 4 | 3 | 4 | 3 | 3 | 3 | 2 | 1 | 2 | 2 |
| High | Q12769 | Nuclear pore complex protein Nup160 OS=Homo sapiens OX=9606 GN=NUP160 PE=1 SV=3 | 33 | 13 | 13 | 162 | 11 | 1436 | 5.5 | 2 | 4 | 3 | | 1 | | 2 | 4 | 3 | | 1 | |
| High | P05026 | Sodium/potassium-transporting ATPase subunit beta-1 OS=Homo sapiens OX=9606 GN=ATP1B1 PE=1 SV=1 | 33 | 6 | 6 | 35 | 25 | 303 | 8.53 | 3 | 2 | 5 | 1 | 2 | 3 | 3 | 2 | 4 | 1 | 2 | 3 |
| High | P13667 | Protein disulfide-isomerase A4 OS=Homo sapiens OX=9606 GN=PDIA4 PE=1 SV=2 | 33 | 9 | 9 | 72.9 | 20 | 645 | 5.07 | 7 | 8 | 7 | | | | 4 | 5 | 6 | | | |
| High | P18669 | Phosphoglycerate mutase 1 OS=Homo sapiens OX=9606 GN=PGAM1 PE=1 SV=2 | 33 | 7 | 7 | 28.8 | 38 | 254 | 7.18 | 5 | 4 | 5 | 1 | 1 | | 5 | 4 | 5 | 1 | 1 | |
| High | P54819 | Adenylate kinase 2, mitochondrial OS=Homo sapiens OX=9606 GN=AK2 PE=1 SV=2 | 33 | 7 | 7 | 26.5 | 33 | 239 | 7.81 | 2 | 3 | 1 | 3 | 4 | 3 | 2 | 3 | 1 | 3 | 4 | 3 |
| High | Q8IWX8 | Calcium homeostasis endoplasmic reticulum protein OS=Homo sapiens OX=9606 GN=CHERP PE=1 SV=3 | 33 | 8 | 8 | 103.6 | 9 | 916 | 9.04 | 4 | 2 | 1 | 3 | 3 | 2 | 4 | 2 | 1 | 3 | 3 | 2 |
| High | Q14134 | Tripartite motif-containing protein 29 OS=Homo sapiens OX=9606 GN=TRIM29 PE=1 SV=2 | 33 | 14 | 14 | 65.8 | 24 | 588 | 7.15 | 9 | 10 | 10 | 1 | 1 | 1 | 8 | 9 | 10 | 1 | 1 | 1 |
| High | P28066 | Proteasome subunit alpha type-5 OS=Homo sapiens OX=9606 GN=PSMA5 PE=1 SV=3 | 33 | 5 | 5 | 26.4 | 29 | 241 | 4.79 | 3 | 3 | 4 | 3 | 3 | 4 | 3 | 3 | 4 | 3 | 3 | 4 |
| High | Q9UHB9 | Signal recognition particle subunit SRP68 OS=Homo sapiens OX=9606 GN=SRP68 PE=1 SV=2 | 33 | 13 | 13 | 70.7 | 21 | 627 | 8.56 | | | 2 | 2 | 1 | 1 | | | 2 | 2 | 1 | 1 |
| High | O76094 | Signal recognition particle subunit SRP72 OS=Homo sapiens OX=9606 GN=SRP72 PE=1 SV=3 | 33 | 14 | 14 | 74.6 | 24 | 671 | 9.26 | 1 | 1 | 1 | 3 | 3 | 2 | 1 | 1 | 1 | 3 | 3 | 2 |
| High | P49023 | Paxillin OS=Homo sapiens OX=9606 GN=PXN PE=1 SV=3 | 33 | 8 | 8 | 64.5 | 16 | 591 | 6.19 | 4 | 3 | 4 | 3 | 2 | 3 | 4 | 3 | 4 | 3 | 2 | 3 |
| High | Q9HCY8 | Protein S100-A14 OS=Homo sapiens OX=9606 GN=S100A14 PE=1 SV=1 | 33 | 5 | 5 | 11.7 | 63 | 104 | 5.24 | 5 | 3 | 4 | 2 | 1 | 1 | 4 | 3 | 4 | 2 | 1 | 1 |
| High | Q15418 | Ribosomal protein S6 kinase alpha-1 OS=Homo sapiens OX=9606 GN=RPS6KA1 PE=1 SV=2 | 33 | 7 | 3 | 82.7 | 10 | 735 | 7.83 | 2 | 2 | 4 | 4 | 3 | 2 | 2 | 2 | 4 | 3 | 2 | 2 |
| High | O15143 | Actin-related protein 2/3 complex subunit 1B OS=Homo sapiens OX=9606 GN=ARPC1B PE=1 SV=3 | 33 | 6 | 6 | 40.9 | 17 | 372 | 8.35 | 4 | 6 | 4 | 4 | 3 | 3 | 4 | 5 | 4 | 4 | 3 | 3 |
| High | P13533 | Myosin-6 OS=Homo sapiens OX=9606 GN=MYH6 PE=1 SV=5 | 33 | 14 | 9 | 223.6 | 9 | 1939 | 5.73 | 3 | 4 | 4 | 6 | 6 | 5 | 3 | 3 | 2 | 6 | 6 | 5 |
| High | Q15286 | Ras-related protein Rab-35 OS=Homo sapiens OX=9606 GN=RAB35 PE=1 SV=1 | 33 | 4 | 2 | 23 | 22 | 201 | 8.29 | 4 | 5 | 4 | 2 | 3 | 2 | 3 | 3 | 3 | 2 | 2 | 2 |
| High | P06731 | Carcinoembryonic antigen-related cell adhesion molecule 5 OS=Homo sapiens OX=9606 GN=CEACAM5 PE=1 SV=4 | 32 | 6 | 4 | 76.7 | 13 | 702 | 5.69 | 4 | 4 | 4 | 2 | 2 | 1 | 4 | 4 | 4 | 2 | 2 | 1 |
| High | P35251 | Replication factor C subunit 1 OS=Homo sapiens OX=9606 GN=RFC1 PE=1 SV=4 | 32 | 11 | 11 | 128.2 | 11 | 1148 | 9.36 | 3 | 2 | 4 | 3 | 3 | 2 | 3 | 2 | 4 | 3 | 3 | 2 |
| High | Q96EY1 | DnaJ homolog subfamily A member 3, mitochondrial OS=Homo sapiens OX=9606 GN=DNAJA3 PE=1 SV=2 | 32 | 6 | 6 | 52.5 | 19 | 480 | 9.26 | 5 | 3 | 4 | 1 | 1 | 1 | 5 | 3 | 4 | 1 | 1 | 1 |
| High | P55072 | Transitional endoplasmic reticulum ATPase OS=Homo sapiens OX=9606 GN=VCP PE=1 SV=4 | 32 | 12 | 12 | 89.3 | 19 | 806 | 5.26 | 4 | 6 | 4 | 1 | 2 | 2 | 4 | 6 | 4 | 1 | 2 | 2 |
| High | Q92522 | Histone H1.10 OS=Homo sapiens OX=9606 GN=H1-10 PE=1 SV=1 | 32 | 5 | 5 | 22.5 | 23 | 213 | 10.76 | | 2 | 4 | 5 | 3 | 2 | | 2 | 4 | 4 | 3 | 2 |
| High | Q9NQ29 | Putative RNA-binding protein Luc7-like 1 OS=Homo sapiens OX=9606 GN=LUC7L PE=1 SV=1 | 32 | 5 | 1 | 43.7 | 17 | 371 | 9.92 | 3 | 3 | 2 | 3 | 3 | 4 | 2 | 2 | 2 | 3 | 3 | 3 |
| High | Q9Y4P3 | Transducin beta-like protein 2 OS=Homo sapiens OX=9606 GN=TBL2 PE=1 SV=1 | 32 | 9 | 9 | 49.8 | 25 | 447 | 9.44 | 2 | 3 | 3 | 3 | 6 | 4 | 2 | 2 | 3 | 5 | 5 | 4 |
| High | P20618 | Proteasome subunit beta type-1 OS=Homo sapiens OX=9606 GN=PSMB1 PE=1 SV=2 | 32 | 6 | 6 | 26.5 | 32 | 241 | 8.13 | 2 | 2 | 2 | 5 | 5 | 6 | 2 | 2 | 2 | 5 | 3 | 6 |
| High | Q8TDN6 | Ribosome biogenesis protein BRX1 homolog OS=Homo sapiens OX=9606 GN=BRIX1 PE=1 SV=2 | 32 | 12 | 12 | 41.4 | 38 | 353 | 9.92 | 3 | 3 | 3 | 3 | 6 | 2 | 3 | 2 | 2 | 3 | 6 | 2 |
| High | Q8IZW8 | Tensin-4 OS=Homo sapiens OX=9606 GN=TNS4 PE=1 SV=3 | 32 | 9 | 9 | 76.7 | 17 | 715 | 7.34 | 4 | 3 | 2 | 5 | 6 | 5 | 4 | 3 | 2 | 5 | 6 | 5 |
| High | O00232 | 26S proteasome non-ATPase regulatory subunit 12 OS=Homo sapiens OX=9606 GN=PSMD12 PE=1 SV=3 | 32 | 8 | 8 | 52.9 | 22 | 456 | 7.65 | 2 | 1 | 3 | 1 | 7 | 1 | 2 | 1 | 3 | 1 | 6 | 1 |
| High | P28482 | Mitogen-activated protein kinase 1 OS=Homo sapiens OX=9606 GN=MAPK1 PE=1 SV=3 | 32 | 7 | 4 | 41.4 | 23 | 360 | 6.98 | | 1 | 3 | 3 | 2 | 4 | | 1 | 3 | 3 | 2 | 4 |

| | | | | | | | | | | | | | | | | | | | | | |
|---|---|---|---|---|---|---|---|---|---|---|---|---|---|---|---|---|---|---|---|---|---|
| High | P21359 | Neurofibromin OS=Homo sapiens OX=9606 GN=NF1 PE=1 SV=2 | 32 | 16 | 16 | 319.2 | 7 | 2839 | 7.39 | 7 | 7 | 10 | | | | 7 | 7 | 10 | | | |
| High | P33527 | Multidrug resistance-associated protein 1 OS=Homo sapiens OX=9606 GN=ABCC1 PE=1 SV=3 | 32 | 11 | 9 | 171.5 | 8 | 1531 | 7.11 | 5 | 6 | 9 | | | 2 | 5 | 5 | 8 | | | 2 |
| High | P05161 | Ubiquitin-like protein ISG15 OS=Homo sapiens OX=9606 GN=ISG15 PE=1 SV=5 | 32 | 4 | 4 | 17.9 | 30 | 165 | 7.44 | 3 | 3 | 3 | 4 | 5 | 2 | 3 | 3 | 3 | 4 | 4 | 2 |
| High | Q9NVI7 | ATPase family AAA domain-containing protein 3A OS=Homo sapiens OX=9606 GN=ATAD3A PE=1 SV=2 | 32 | 8 | 8 | 71.3 | 12 | 634 | 8.98 | 2 | 2 | 3 | 1 | 3 | 2 | 2 | 2 | 3 | 1 | 3 | 2 |
| High | P63151 | Serine/threonine-protein phosphatase 2A 55 kDa regulatory subunit B alpha isoform OS=Homo sapiens OX=9606 GN=PPP2R2A PE | 32 | 7 | 7 | 51.7 | 21 | 447 | 6.2 | 5 | 4 | 2 | 4 | 3 | 4 | 5 | 4 | 2 | 4 | 3 | 4 |
| High | Q9BUH6 | Protein PAXX OS=Homo sapiens OX=9606 GN=PAXX PE=1 SV=2 | 31 | 6 | 6 | 21.6 | 33 | 204 | 5.48 | 3 | 3 | 3 | 3 | 3 | 3 | 3 | 3 | 3 | 3 | 3 | 3 |
| High | Q14192 | Four and a half LIM domains protein 2 OS=Homo sapiens OX=9606 GN=FHL2 PE=1 SV=3 | 31 | 10 | 10 | 32.2 | 41 | 279 | 7.55 | | | 1 | 2 | 3 | 6 | | | 1 | 2 | 3 | 6 |
| High | Q9BZE4 | GTP-binding protein 4 OS=Homo sapiens OX=9606 GN=GTPBP4 PE=1 SV=3 | 31 | 12 | 12 | 73.9 | 18 | 634 | 9.5 | | 1 | | 3 | 5 | 4 | | 1 | | 3 | 5 | 4 |
| High | P04792 | Heat shock protein beta-1 OS=Homo sapiens OX=9606 GN=HSPB1 PE=1 SV=2 | 31 | 7 | 7 | 22.8 | 39 | 205 | 6.4 | 2 | 3 | 2 | 3 | 4 | 2 | 2 | 3 | 2 | 3 | 4 | 2 |
| High | O00161 | Synaptosomal-associated protein 23 OS=Homo sapiens OX=9606 GN=SNAP23 PE=1 SV=1 | 31 | 10 | 10 | 23.3 | 62 | 211 | 5.01 | 2 | 1 | 3 | | 1 | | 2 | 1 | 2 | | 1 | |
| High | Q8TCJ2 | Dolichyl-diphosphooligosaccharide--protein glycosyltransferase subunit STT3B OS=Homo sapiens OX=9606 GN=STT3B PE=1 SV=1 | 31 | 9 | 8 | 93.6 | 10 | 826 | 8.91 | 3 | 5 | 5 | 1 | 1 | 2 | 2 | 4 | 4 | 1 | 1 | 2 |
| High | P62316 | Small nuclear ribonucleoprotein Sm D2 OS=Homo sapiens OX=9606 GN=SNRPD2 PE=1 SV=1 | 31 | 6 | 6 | 13.5 | 56 | 118 | 9.91 | 4 | 3 | 2 | 3 | 2 | 4 | 4 | 3 | 2 | 3 | 2 | 4 |
| High | Q8TDB6 | E3 ubiquitin-protein ligase DTX3L OS=Homo sapiens OX=9606 GN=DTX3L PE=1 SV=1 | 31 | 9 | 9 | 83.5 | 16 | 740 | 8.06 | 4 | 3 | 4 | 4 | 4 | 3 | 4 | 3 | 4 | 4 | 4 | 3 |
| High | P49959 | Double-strand break repair protein MRE11 OS=Homo sapiens OX=9606 GN=MRE11 PE=1 SV=3 | 31 | 11 | 11 | 80.5 | 20 | 708 | 5.9 | 2 | 4 | 4 | 1 | 1 | 2 | 2 | 4 | 4 | 1 | 1 | 2 |
| High | Q13619 | Cullin-4A OS=Homo sapiens OX=9606 GN=CUL4A PE=1 SV=3 | 31 | 12 | 12 | 87.6 | 16 | 759 | 8.13 | 7 | 8 | 3 | | 3 | 1 | 7 | 8 | 3 | | 3 | 1 |
| High | Q16831 | Uridine phosphorylase 1 OS=Homo sapiens OX=9606 GN=UPP1 PE=1 SV=1 | 31 | 7 | 7 | 33.9 | 22 | 310 | 7.88 | 1 | | 1 | 6 | 5 | 6 | 1 | | 1 | 6 | 5 | 5 |
| High | Q13136 | Liprin-alpha-1 OS=Homo sapiens OX=9606 GN=PPFIA1 PE=1 SV=1 | 31 | 11 | 11 | 135.7 | 13 | 1202 | 6.29 | 5 | 6 | 4 | | 1 | | 5 | 6 | 4 | | 1 | |
| High | Q96CU9 | FAD-dependent oxidoreductase domain-containing protein 1 OS=Homo sapiens OX=9606 GN=FOXRED1 PE=1 SV=2 | 31 | 7 | 7 | 53.8 | 19 | 486 | 7.78 | | 2 | 3 | 3 | 3 | 3 | | 2 | 3 | 3 | 3 | 3 |
| High | P62851 | 40S ribosomal protein S25 OS=Homo sapiens OX=9606 GN=RPS25 PE=1 SV=1 | 31 | 4 | 4 | 13.7 | 28 | 125 | 10.11 | 3 | 4 | 3 | 4 | 4 | 4 | 3 | 4 | 3 | 4 | 4 | 4 |
| High | P11177 | Pyruvate dehydrogenase E1 component subunit beta, mitochondrial OS=Homo sapiens OX=9606 GN=PDHB PE=1 SV=3 | 31 | 9 | 9 | 39.2 | 31 | 359 | 6.65 | 3 | 5 | 1 | | 1 | 1 | 3 | 3 | 1 | | 1 | 1 |
| High | P43490 | Nicotinamide phosphoribosyltransferase OS=Homo sapiens OX=9606 GN=NAMPT PE=1 SV=1 | 31 | 9 | 9 | 55.5 | 26 | 491 | 7.15 | 6 | 5 | 8 | 1 | 3 | 1 | 6 | 5 | 8 | 1 | 3 | 1 |
| High | Q7LBC6 | Lysine-specific demethylase 3B OS=Homo sapiens OX=9606 GN=KDM3B PE=1 SV=2 | 31 | 12 | 12 | 191.5 | 9 | 1761 | 7.18 | 1 | 1 | | 1 | 4 | | 1 | 1 | | 1 | 4 | |
| High | Q9H936 | Mitochondrial glutamate carrier 1 OS=Homo sapiens OX=9606 GN=SLC25A22 PE=1 SV=1 | 31 | 7 | 7 | 34.4 | 21 | 323 | 9.29 | 6 | 5 | 3 | 1 | 2 | 1 | 6 | 5 | 3 | 1 | 2 | 1 |
| High | Q9NZN4 | EH domain-containing protein 2 OS=Homo sapiens OX=9606 GN=EHD2 PE=1 SV=2 | 31 | 9 | 7 | 61.1 | 19 | 543 | 6.46 | 6 | 6 | 4 | 2 | 3 | 3 | 6 | 6 | 4 | 2 | 3 | 3 |
| High | O43852 | Calumenin OS=Homo sapiens OX=9606 GN=CALU PE=1 SV=2 | 31 | 11 | 11 | 37.1 | 47 | 315 | 4.64 | 7 | 7 | 6 | 2 | 4 | 2 | 7 | 7 | 6 | 2 | 3 | 2 |
| High | Q7Z5J4 | Retinoic acid-induced protein 1 OS=Homo sapiens OX=9606 GN=RAI1 PE=1 SV=2 | 31 | 12 | 12 | 203.2 | 11 | 1906 | 8.79 | | 1 | 1 | 5 | 6 | 7 | | 1 | 1 | 5 | 6 | 6 |
| High | P62995 | Transformer-2 protein homolog beta OS=Homo sapiens OX=9606 GN=TRA2B PE=1 SV=1 | 31 | 5 | 5 | 33.6 | 22 | 288 | 11.25 | 3 | 4 | 6 | 3 | 3 | 2 | 3 | 3 | 5 | 3 | 3 | 2 |
| High | O95049 | Tight junction protein ZO-3 OS=Homo sapiens OX=9606 GN=TJP3 PE=1 SV=3 | 31 | 10 | 10 | 101.3 | 13 | 919 | 6.76 | 5 | 4 | 4 | 1 | 2 | 1 | 5 | 4 | 4 | 1 | 2 | 1 |
| High | Q14203 | Dynactin subunit 1 OS=Homo sapiens OX=9606 GN=DCTN1 PE=1 SV=3 | 30 | 10 | 10 | 141.6 | 12 | 1278 | 5.81 | 4 | 3 | 5 | 2 | | 1 | 4 | 3 | 5 | 2 | | 1 |
| High | P24941 | Cyclin-dependent kinase 2 OS=Homo sapiens OX=9606 GN=CDK2 PE=1 SV=2 | 30 | 5 | 3 | 33.9 | 17 | 298 | 8.68 | 2 | 4 | 5 | 2 | 4 | 2 | 2 | 3 | 4 | 1 | 3 | 2 |
| High | Q8IYB3 | Serine/arginine repetitive matrix protein 1 OS=Homo sapiens OX=9606 GN=SRRM1 PE=1 SV=2 | 30 | 8 | 8 | 102.3 | 14 | 904 | 11.84 | 1 | 1 | 2 | 4 | 4 | 4 | 1 | 1 | 2 | 4 | 4 | 4 |
| High | P62277 | 40S ribosomal protein S13 OS=Homo sapiens OX=9606 GN=RPS13 PE=1 SV=2 | 30 | 5 | 5 | 17.2 | 30 | 151 | 10.54 | 3 | 4 | 4 | 3 | 3 | 3 | 3 | 4 | 4 | 3 | 3 | 3 |
| High | Q15370 | Elongin-B OS=Homo sapiens OX=9606 GN=ELOB PE=1 SV=1 | 30 | 5 | 5 | 13.1 | 49 | 118 | 4.88 | 4 | 3 | 3 | 2 | 4 | 2 | 3 | 3 | 3 | 2 | 4 | 2 |
| High | Q9NTJ3 | Structural maintenance of chromosomes protein 4 OS=Homo sapiens OX=9606 GN=SMC4 PE=1 SV=2 | 30 | 10 | 10 | 147.1 | 9 | 1288 | 6.79 | 4 | 4 | 4 | 2 | 1 | 2 | 4 | 4 | 4 | 2 | 1 | 2 |
| High | Q4G0J3 | La-related protein 7 OS=Homo sapiens OX=9606 GN=LARP7 PE=1 SV=1 | 30 | 8 | 8 | 66.9 | 17 | 582 | 9.55 | 1 | 2 | 1 | 2 | 2 | 2 | 1 | 2 | 1 | 2 | 2 | 2 |
| High | Q9UMS4 | Pre-mRNA-processing factor 19 OS=Homo sapiens OX=9606 GN=PRPF19 PE=1 SV=1 | 30 | 7 | 7 | 55.1 | 17 | 504 | 6.61 | 1 | 1 | 1 | 6 | 4 | 5 | 1 | 1 | 1 | 6 | 3 | 5 |
| High | Q9HAN9 | Nicotinamide/nicotinic acid mononucleotide adenylyltransferase 1 OS=Homo sapiens OX=9606 GN=NMNAT1 PE=1 SV=1 | 30 | 6 | 6 | 31.9 | 29 | 279 | 8.87 | 2 | 1 | 2 | 3 | 2 | 4 | 2 | 1 | 2 | 3 | 2 | 3 |
| High | O94901 | SUN domain-containing protein 1 OS=Homo sapiens OX=9606 GN=SUN1 PE=1 SV=4 | 30 | 11 | 11 | 87.1 | 17 | 785 | 6.93 | 6 | 7 | 5 | 1 | 2 | 1 | 5 | 6 | 5 | 1 | 2 | 1 |
| High | O00231 | 26S proteasome non-ATPase regulatory subunit 11 OS=Homo sapiens OX=9606 GN=PSMD11 PE=1 SV=3 | 30 | 12 | 12 | 47.4 | 33 | 422 | 6.48 | | 2 | 1 | 4 | 3 | 5 | | 2 | 1 | 4 | 3 | 5 |
| High | P55769 | NHP2-like protein 1 OS=Homo sapiens OX=9606 GN=SNU13 PE=1 SV=3 | 30 | 5 | 5 | 14.2 | 46 | 128 | 8.46 | 2 | 2 | 3 | 4 | 4 | 4 | 2 | 2 | 3 | 3 | 3 | 3 |
| High | Q8IWS0 | PHD finger protein 6 OS=Homo sapiens OX=9606 GN=PHF6 PE=1 SV=1 | 30 | 4 | 4 | 41.3 | 13 | 365 | 8.68 | 3 | 2 | 3 | 3 | 4 | 3 | 3 | 2 | 3 | 3 | 4 | 3 |
| High | Q9NZ08 | Endoplasmic reticulum aminopeptidase 1 OS=Homo sapiens OX=9606 GN=ERAP1 PE=1 SV=3 | 30 | 11 | 11 | 107.2 | 16 | 941 | 6.46 | 7 | 10 | 9 | 1 | | 1 | 7 | 9 | 9 | 1 | | 1 |
| High | Q9ULX6 | A-kinase anchor protein 8-like OS=Homo sapiens OX=9606 GN=AKAP8L PE=1 SV=4 | 30 | 6 | 6 | 71.6 | 13 | 646 | 5.05 | 7 | 7 | 6 | | | | 6 | 6 | 5 | | | |
| High | P27824 | Calnexin OS=Homo sapiens OX=9606 GN=CANX PE=1 SV=2 | 30 | 12 | 12 | 67.5 | 24 | 592 | 4.6 | 3 | 7 | 3 | | | | 3 | 6 | 3 | | | |
| High | Q86V81 | THO complex subunit 4 OS=Homo sapiens OX=9606 GN=ALYREF PE=1 SV=3 | 30 | 4 | 4 | 26.9 | 23 | 257 | 11.15 | 3 | 3 | 2 | 2 | 4 | 2 | 3 | 3 | 2 | 2 | 3 | 2 |
| High | Q9NX58 | Cell growth-regulating nucleolar protein OS=Homo sapiens OX=9606 GN=LYAR PE=1 SV=3 | 30 | 8 | 8 | 43.6 | 25 | 379 | 9.57 | 1 | 1 | 2 | 2 | 4 | 3 | 1 | 1 | 2 | 2 | 3 | 3 |
| High | P21589 | 5'-nucleotidase OS=Homo sapiens OX=9606 GN=NT5E PE=1 SV=1 | 30 | 10 | 10 | 63.3 | 24 | 574 | 7.03 | 1 | | 3 | | 1 | | 1 | | 3 | | 1 | |
| High | Q92600 | CCR4-NOT transcription complex subunit 9 OS=Homo sapiens OX=9606 GN=CNOT9 PE=1 SV=1 | 30 | 7 | 7 | 33.6 | 24 | 299 | 8.03 | | | 4 | 2 | 3 | 3 | | | 4 | 2 | 3 | 3 |
| High | Q9Y277 | Voltage-dependent anion-selective channel protein 3 OS=Homo sapiens OX=9606 GN=VDAC3 PE=1 SV=1 | 30 | 5 | 4 | 30.6 | 19 | 283 | 8.66 | 1 | 3 | 2 | 1 | 3 | 3 | 1 | 3 | 2 | 1 | 3 | 3 |
| High | P56537 | Eukaryotic translation initiation factor 6 OS=Homo sapiens OX=9606 GN=EIF6 PE=1 SV=1 | 30 | 5 | 5 | 26.6 | 36 | 245 | 4.68 | 3 | 1 | 3 | 3 | 3 | 3 | 3 | 1 | 3 | 3 | 3 | 3 |
| High | P35611 | Alpha-adducin OS=Homo sapiens OX=9606 GN=ADD1 PE=1 SV=2 | 30 | 10 | 10 | 80.9 | 17 | 737 | 5.83 | 3 | 1 | 4 | 2 | 1 | 4 | 3 | 1 | 4 | 2 | 1 | 4 |
| High | O15212 | Prefoldin subunit 6 OS=Homo sapiens OX=9606 GN=PFDN6 PE=1 SV=1 | 30 | 4 | 4 | 14.6 | 36 | 129 | 8.88 | 2 | 2 | 2 | 4 | 4 | 3 | 2 | 2 | 2 | 2 | 2 | 2 |
| High | P61026 | Ras-related protein Rab-10 OS=Homo sapiens OX=9606 GN=RAB10 PE=1 SV=1 | 29 | 5 | 3 | 22.5 | 32 | 200 | 8.38 | 4 | 5 | 3 | 2 | 3 | 2 | 3 | 3 | 2 | 2 | 2 | 2 |
| High | P29692 | Elongation factor 1-delta OS=Homo sapiens OX=9606 GN=EEF1D PE=1 SV=5 | 29 | 9 | 8 | 31.1 | 43 | 281 | 5.01 | 3 | 3 | 2 | 3 | 3 | 2 | 3 | 2 | 2 | 3 | 3 | 2 |
| High | P54578 | Ubiquitin carboxyl-terminal hydrolase 14 OS=Homo sapiens OX=9606 GN=USP14 PE=1 SV=3 | 29 | 9 | 9 | 56 | 23 | 494 | 5.3 | 4 | 1 | 2 | 1 | 2 | 2 | 4 | 1 | 2 | 1 | 2 | 2 |
| High | P07305 | Histone H1.0 OS=Homo sapiens OX=9606 GN=H1-0 PE=1 SV=3 | 29 | 5 | 5 | 20.9 | 21 | 194 | 10.84 | 3 | 1 | 3 | 3 | 1 | 4 | 3 | 1 | 3 | 2 | 1 | 3 |
| High | Q15067 | Peroxisomal acyl-coenzyme A oxidase 1 OS=Homo sapiens OX=9606 GN=ACOX1 PE=1 SV=3 | 29 | 9 | 9 | 74.4 | 21 | 660 | 8.16 | 4 | 3 | 3 | | 3 | 2 | 4 | 3 | 3 | | 3 | 2 |
| High | O95292 | Vesicle-associated membrane protein-associated protein B/C OS=Homo sapiens OX=9606 GN=VAPB PE=1 SV=3 | 29 | 5 | 4 | 27.2 | 29 | 243 | 7.3 | 2 | 2 | 4 | 2 | 1 | 4 | 2 | 2 | 4 | 2 | 1 | 4 |
| High | Q9H0U3 | Magnesium transporter protein 1 OS=Homo sapiens OX=9606 GN=MAGT1 PE=1 SV=1 | 29 | 5 | 5 | 38 | 15 | 335 | 9.63 | 4 | 4 | 5 | 3 | 2 | 3 | 4 | 4 | 4 | 3 | 2 | 3 |
| High | Q86VM9 | Zinc finger CCCH domain-containing protein 18 OS=Homo sapiens OX=9606 GN=ZC3H18 PE=1 SV=2 | 29 | 9 | 9 | 106.3 | 12 | 953 | 8.32 | | | 1 | 4 | 3 | 5 | | | 1 | 4 | 3 | 5 |
| High | P26640 | Valine--tRNA ligase OS=Homo sapiens OX=9606 GN=VARS1 PE=1 SV=4 | 29 | 9 | 9 | 140.4 | 11 | 1264 | 7.59 | 2 | 3 | 3 | 2 | 3 | 2 | 2 | 3 | 3 | 2 | 3 | 2 |
| High | Q5MNZ6 | WD repeat domain phosphoinositide-interacting protein 3 OS=Homo sapiens OX=9606 GN=WDR45B PE=1 SV=2 | 29 | 5 | 5 | 38.1 | 16 | 344 | 7.59 | 2 | 3 | 2 | 3 | 3 | 3 | 2 | 3 | 2 | 3 | 3 | 3 |
| High | P59998 | Actin-related protein 2/3 complex subunit 4 OS=Homo sapiens OX=9606 GN=ARPC4 PE=1 SV=3 | 29 | 4 | 4 | 19.7 | 23 | 168 | 8.43 | 3 | 3 | 3 | 3 | 3 | 3 | 3 | 3 | 3 | 3 | 3 | 3 |
| High | Q12904 | Aminoacyl tRNA synthase complex-interacting multifunctional protein 1 OS=Homo sapiens OX=9606 GN=AIMP1 PE=1 SV=2 | 29 | 6 | 6 | 34.3 | 28 | 312 | 8.43 | 4 | 4 | 4 | 1 | 2 | 2 | 3 | 3 | 3 | 1 | 2 | 2 |
| High | Q96N67 | Dedicator of cytokinesis protein 7 OS=Homo sapiens OX=9606 GN=DOCK7 PE=1 SV=4 | 29 | 10 | 10 | 242.4 | 6 | 2140 | 6.8 | 2 | 4 | 2 | | 2 | 1 | 2 | 4 | 2 | | 2 | 1 |
| High | P49321 | Nuclear autoantigenic sperm protein OS=Homo sapiens OX=9606 GN=NASP PE=1 SV=2 | 29 | 7 | 7 | 85.2 | 9 | 788 | 4.3 | 1 | | 2 | 4 | 4 | 3 | 1 | | 2 | 4 | 4 | 3 |
| High | P15311 | Ezrin OS=Homo sapiens OX=9606 GN=EZR PE=1 SV=4 | 29 | 9 | 5 | 69.4 | 15 | 586 | 6.27 | 2 | 3 | 7 | 2 | 2 | 2 | 2 | 3 | 6 | 2 | 2 | 2 |
| High | O43670 | BUB3-interacting and GLEBS motif-containing protein ZNF207 OS=Homo sapiens OX=9606 GN=ZNF207 PE=1 SV=1 | 29 | 3 | 3 | 50.7 | 9 | 478 | 9.1 | 1 | 2 | 2 | 4 | 4 | 3 | 1 | 2 | 2 | 2 | 2 | 2 |
| High | Q27J81 | Inverted formin-2 OS=Homo sapiens OX=9606 GN=INF2 PE=1 SV=2 | 29 | 12 | 12 | 135.5 | 15 | 1249 | 5.38 | 4 | 5 | 2 | 1 | 2 | 2 | 4 | 5 | 2 | 1 | 2 | 2 |
| High | O43633 | Charged multivesicular body protein 2a OS=Homo sapiens OX=9606 GN=CHMP2A PE=1 SV=1 | 29 | 6 | 6 | 25.1 | 25 | 222 | 5.97 | 3 | 3 | 4 | 1 | 2 | 1 | 3 | 3 | 4 | 1 | 2 | 1 |
| High | Q15637 | Splicing factor 1 OS=Homo sapiens OX=9606 GN=SF1 PE=1 SV=4 | 29 | 6 | 6 | 68.3 | 13 | 639 | 8.98 | 3 | 3 | 4 | 2 | 3 | | 3 | 3 | 4 | 1 | 2 | |
| High | Q9BQL6 | Fermitin family homolog 1 OS=Homo sapiens OX=9606 GN=FERMT1 PE=1 SV=1 | 29 | 11 | 11 | 77.4 | 19 | 677 | 6.28 | 7 | 9 | 8 | | | | 7 | 9 | 8 | | | |
| High | Q8TE67 | Epidermal growth factor receptor kinase substrate 8-like protein 3 OS=Homo sapiens OX=9606 GN=EPS8L3 PE=1 SV=2 | 29 | 10 | 10 | 66.8 | 24 | 593 | 7.99 | 1 | 2 | 4 | 1 | 2 | 1 | 1 | 2 | 3 | 1 | 2 | 1 |
| High | Q6IBS0 | Twinfilin-2 OS=Homo sapiens OX=9606 GN=TWF2 PE=1 SV=2 | 29 | 7 | 6 | 39.5 | 22 | 349 | 6.84 | 3 | 3 | 2 | 2 | 3 | 2 | 3 | 3 | 2 | 2 | 2 | 2 |
| High | O96019 | Actin-like protein 6A OS=Homo sapiens OX=9606 GN=ACTL6A PE=1 SV=1 | 29 | 7 | 7 | 47.4 | 20 | 429 | 5.6 | 2 | | 3 | 3 | 3 | 2 | 2 | | 3 | 3 | 3 | 2 |
| High | Q9HD45 | Transmembrane 9 superfamily member 3 OS=Homo sapiens OX=9606 GN=TM9SF3 PE=1 SV=2 | 29 | 9 | 9 | 67.8 | 16 | 589 | 7.21 | 6 | 4 | 4 | | 1 | | 6 | 4 | 4 | | 1 | |
| High | Q93084 | Sarcoplasmic/endoplasmic reticulum calcium ATPase 3 OS=Homo sapiens OX=9606 GN=ATP2A3 PE=1 SV=3 | 29 | 6 | 1 | 109.2 | 7 | 999 | 5.52 | 3 | 3 | 4 | 2 | 3 | 2 | 3 | 3 | 3 | 2 | 3 | 2 |
| High | Q9UL46 | Proteasome activator complex subunit 2 OS=Homo sapiens OX=9606 GN=PSME2 PE=1 SV=4 | 29 | 5 | 5 | 27.4 | 23 | 239 | 5.73 | 2 | 3 | 1 | 3 | 4 | 4 | 2 | 3 | 1 | 3 | 4 | 4 |
| High | O43809 | Cleavage and polyadenylation specificity factor subunit 5 OS=Homo sapiens OX=9606 GN=NUDT21 PE=1 SV=1 | 29 | 6 | 6 | 26.2 | 27 | 227 | 8.82 | 2 | 2 | 3 | 2 | 4 | 2 | 2 | 2 | 3 | 2 | 3 | 2 |
| High | Q6P1X6 | UPF0598 protein C8orf82 OS=Homo sapiens OX=9606 GN=C8orf82 PE=1 SV=2 | 29 | 5 | 5 | 23.9 | 31 | 216 | 9.14 | 4 | 5 | 3 | 1 | 1 | | 4 | 3 | 3 | 1 | 1 | |
| High | Q9NP79 | Vacuolar protein sorting-associated protein VTA1 homolog OS=Homo sapiens OX=9606 GN=VTA1 PE=1 SV=1 | 29 | 8 | 8 | 33.9 | 30 | 307 | 6.29 | 4 | 4 | 3 | | 2 | | 4 | 4 | 2 | | 2 | |
| High | P34896 | Serine hydroxymethyltransferase, cytosolic OS=Homo sapiens OX=9606 GN=SHMT1 PE=1 SV=1 | 29 | 9 | 8 | 53 | 22 | 483 | 7.71 | 4 | 2 | 4 | 2 | 2 | 2 | 3 | 2 | 3 | 2 | 2 | 2 |
| High | P07203 | Glutathione peroxidase 1 OS=Homo sapiens OX=9606 GN=GPX1 PE=1 SV=4 | 29 | 6 | 6 | 22.1 | 36 | 203 | 6.55 | 3 | 4 | 4 | 2 | 4 | 1 | 3 | 4 | 4 | 2 | 4 | 1 |
| High | Q15208 | Serine/threonine-protein kinase 38 OS=Homo sapiens OX=9606 GN=STK38 PE=1 SV=1 | 29 | 9 | 7 | 54.2 | 23 | 465 | 7.15 | 2 | 4 | 5 | 2 | 2 | 2 | 2 | 4 | 4 | 2 | 2 | 2 |
| High | Q13627 | Dual specificity tyrosine-phosphorylation-regulated kinase 1A OS=Homo sapiens OX=9606 GN=DYRK1A PE=1 SV=2 | 29 | 11 | 11 | 85.5 | 17 | 763 | 8.75 | | 1 | | 2 | 3 | 3 | | 1 | | 2 | 3 | 3 |
| High | P30837 | Aldehyde dehydrogenase X, mitochondrial OS=Homo sapiens OX=9606 GN=ALDH1B1 PE=1 SV=4 | 29 | 4 | 2 | 57.2 | 9 | 517 | 6.99 | 7 | 5 | 4 | 2 | 1 | 2 | 3 | 3 | 3 | 2 | 1 | 2 |
| High | P51116 | Fragile X mental retardation syndrome-related protein 2 OS=Homo sapiens OX=9606 GN=FXR2 PE=1 SV=2 | 29 | 6 | 3 | 74.2 | 11 | 673 | 6.23 | 3 | 4 | 2 | 3 | 3 | 3 | 3 | 4 | 2 | 3 | 3 | 3 |
| High | Q9UBS4 | DnaJ homolog subfamily B member 11 OS=Homo sapiens OX=9606 GN=DNAJB11 PE=1 SV=1 | 28 | 4 | 4 | 40.5 | 19 | 358 | 6.18 | 3 | 5 | 5 | | 1 | | 3 | 3 | 3 | | 1 | |
| High | P29373 | Cellular retinoic acid-binding protein 2 OS=Homo sapiens OX=9606 GN=CRABP2 PE=1 SV=2 | 28 | 5 | 5 | 15.7 | 36 | 138 | 5.4 | 4 | 3 | 3 | 3 | 2 | 3 | 4 | 3 | 3 | 3 | 2 | 3 |
| High | O15427 | Monocarboxylate transporter 4 OS=Homo sapiens OX=9606 GN=SLC16A3 PE=1 SV=1 | 28 | 6 | 6 | 49.4 | 12 | 465 | 7.96 | 6 | 6 | 6 | 1 | 2 | | 6 | 6 | 6 | 1 | 2 | |
| High | P62318 | Small nuclear ribonucleoprotein Sm D3 OS=Homo sapiens OX=9606 GN=SNRPD3 PE=1 SV=1 | 28 | 3 | 3 | 13.9 | 32 | 126 | 10.32 | 4 | 2 | 3 | 3 | 3 | 2 | 3 | 2 | 3 | 2 | 2 | 2 |
| High | P07948 | Tyrosine-protein kinase Lyn OS=Homo sapiens OX=9606 GN=LYN PE=1 SV=3 | 28 | 6 | 3 | 58.5 | 10 | 512 | 7.11 | 3 | 4 | 3 | 3 | 2 | 1 | 3 | 3 | 2 | 2 | 1 | 1 |
| High | P62847 | 40S ribosomal protein S24 OS=Homo sapiens OX=9606 GN=RPS24 PE=1 SV=1 | 28 | 4 | 4 | 15.4 | 35 | 133 | 10.78 | 3 | 2 | 2 | 2 | 3 | 1 | 3 | 2 | 2 | 2 | 2 | 1 |
| High | P51608 | Methyl-CpG-binding protein 2 OS=Homo sapiens OX=9606 GN=MECP2 PE=1 SV=1 | 28 | 9 | 9 | 52.4 | 25 | 486 | 9.95 | 8 | 6 | 8 | 1 | 2 | 1 | 7 | 5 | 6 | 1 | 2 | 1 |
| High | O95573 | Fatty acid CoA ligase Acsl3 OS=Homo sapiens OX=9606 GN=ACSL3 PE=1 SV=3 | 28 | 10 | 9 | 80.4 | 19 | 720 | 8.38 | 3 | 3 | 5 | | | | 3 | 3 | 4 | | | |
| High | P20073 | Annexin A7 OS=Homo sapiens OX=9606 GN=ANXA7 PE=1 SV=3 | 28 | 6 | 6 | 52.7 | 15 | 488 | 5.68 | 4 | 2 | 3 | 3 | 3 | 4 | 4 | 2 | 3 | 3 | 3 | 4 |
| High | P53999 | Activated RNA polymerase II transcriptional coactivator p15 OS=Homo sapiens OX=9606 GN=SUB1 PE=1 SV=3 | 28 | 4 | 4 | 14.4 | 37 | 127 | 9.6 | 2 | 2 | 3 | 2 | 4 | 3 | 2 | 2 | 3 | 2 | 4 | 3 |
| High | Q8IWV7 | E3 ubiquitin-protein ligase UBR1 OS=Homo sapiens OX=9606 GN=UBR1 PE=1 SV=1 | 28 | 13 | 13 | 200.1 | 10 | 1749 | 6.01 | 8 | 8 | 8 | | | | 8 | 8 | 8 | | | |
| High | Q6UB35 | Monofunctional C1-tetrahydrofolate synthase, mitochondrial OS=Homo sapiens OX=9606 GN=MTHFD1L PE=1 SV=1 | 28 | 10 | 9 | 105.7 | 11 | 978 | 8.06 | | | 1 | 5 | 5 | 2 | | | 1 | 5 | 5 | 2 |
| High | Q9UHD1 | Cysteine and histidine-rich domain-containing protein 1 OS=Homo sapiens OX=9606 GN=CHORDC1 PE=1 SV=2 | 28 | 9 | 9 | 37.5 | 33 | 332 | 7.87 | 2 | 3 | 3 | 3 | 3 | 1 | 2 | 3 | 3 | 3 | 3 | 1 |
| High | Q9C005 | Protein dpy-30 homolog OS=Homo sapiens OX=9606 GN=DPY30 PE=1 SV=1 | 28 | 2 | 2 | 11.2 | 38 | 99 | 4.88 | | | 1 | 1 | 3 | 2 | | | 1 | 1 | 2 | 1 |
| High | Q92747 | Actin-related protein 2/3 complex subunit 1A OS=Homo sapiens OX=9606 GN=ARPC1A PE=1 SV=2 | 28 | 8 | 8 | 41.5 | 24 | 370 | 8.18 | 2 | 4 | 3 | 1 | 3 | 2 | 2 | 4 | 3 | 1 | 3 | 2 |
| High | O60341 | Lysine-specific histone demethylase 1A OS=Homo sapiens OX=9606 GN=KDM1A PE=1 SV=2 | 28 | 15 | 15 | 92.8 | 21 | 852 | 6.52 | 1 | 1 | 1 | 1 | 1 | | 1 | 1 | 1 | 1 | 1 | |
| High | O75367 | Core histone macro-H2A.1 OS=Homo sapiens OX=9606 GN=MACROH2A1 PE=1 SV=4 | 28 | 8 | 8 | 39.6 | 26 | 372 | 9.79 | 4 | 3 | 3 | 5 | 3 | 5 | 4 | 3 | 3 | 5 | 3 | 5 |
| High | P31689 | DnaJ homolog subfamily A member 1 OS=Homo sapiens OX=9606 GN=DNAJA1 PE=1 SV=2 | 28 | 9 | 9 | 44.8 | 34 | 397 | 7.08 | 6 | 5 | 3 | 1 | 2 | 2 | 5 | 5 | 3 | 1 | 2 | 2 |
| High | P48729 | Casein kinase I isoform alpha OS=Homo sapiens OX=9606 GN=CSNK1A1 PE=1 SV=2 | 28 | 8 | 8 | 38.9 | 25 | 337 | 9.57 | 3 | 4 | 6 | 1 | 2 | 1 | 3 | 4 | 6 | 1 | 2 | 1 |
| High | Q9BW04 | Specifically androgen-regulated gene protein OS=Homo sapiens OX=9606 GN=SARG PE=1 SV=2 | 28 | 7 | 7 | 63.9 | 17 | 601 | 8.62 | 2 | 2 | 2 | 3 | 4 | 3 | 2 | 2 | 2 | 3 | 4 | 3 |
| High | P84022 | Mothers against decapentaplegic homolog 3 OS=Homo sapiens OX=9606 GN=SMAD3 PE=1 SV=1 | 28 | 5 | 4 | 48.1 | 16 | 425 | 7.15 | 2 | 2 | 2 | 4 | 3 | 3 | 2 | 2 | 2 | 4 | 3 | 3 |
| High | Q9UBM7 | 7-dehydrocholesterol reductase OS=Homo sapiens OX=9606 GN=DHCR7 PE=1 SV=1 | 28 | 6 | 6 | 54.5 | 15 | 475 | 8.7 | 5 | 2 | 4 | 3 | 1 | 1 | 5 | 2 | 4 | 3 | 1 | 1 |
| High | Q52LJ0 | Protein FAM98B OS=Homo sapiens OX=9606 GN=FAM98B PE=1 SV=2 | 28 | 6 | 6 | 45.5 | 21 | 433 | 8.69 | | | 1 | 3 | 3 | 3 | | | 1 | 3 | 3 | 2 |
| High | Q06323 | Proteasome activator complex subunit 1 OS=Homo sapiens OX=9606 GN=PSME1 PE=1 SV=1 | 28 | 8 | 8 | 28.7 | 37 | 249 | 6.02 | 2 | 3 | 1 | 3 | 6 | 2 | 2 | 3 | 1 | 3 | 4 | 2 |
| High | Q9Y285 | Phenylalanine--tRNA ligase alpha subunit OS=Homo sapiens OX=9606 GN=FARSA PE=1 SV=3 | 28 | 11 | 11 | 57.5 | 29 | 508 | 7.8 | 2 | 4 | 4 | 2 | 1 | 2 | 2 | 4 | 4 | 2 | 1 | 2 |
| High | Q13283 | Ras GTPase-activating protein-binding protein 1 OS=Homo sapiens OX=9606 GN=G3BP1 PE=1 SV=1 | 27 | 7 | 7 | 52.1 | 23 | 466 | 5.52 | 3 | 3 | 5 | 1 | 4 | 1 | 3 | 3 | 5 | 1 | 2 | 1 |
| High | Q9C0A6 | Histone-lysine N-methyltransferase SETD5 OS=Homo sapiens OX=9606 GN=SETD5 PE=1 SV=2 | 27 | 6 | 5 | 157.4 | 6 | 1442 | 8.46 | | | | 4 | 4 | 4 | | | | 4 | 4 | 4 |
| High | Q9UNF1 | Melanoma-associated antigen D2 OS=Homo sapiens OX=9606 GN=MAGED2 PE=1 SV=2 | 27 | 8 | 7 | 64.9 | 20 | 606 | 9.32 | 3 | 3 | 3 | 1 | | | 3 | 3 | 3 | 1 | | |
| High | Q9GZR7 | ATP-dependent RNA helicase DDX24 OS=Homo sapiens OX=9606 GN=DDX24 PE=1 SV=1 | 27 | 9 | 9 | 96.3 | 17 | 859 | 9.06 | 1 | 1 | 1 | 2 | 2 | 2 | 1 | 1 | 1 | 2 | 2 | 2 |
| High | P21291 | Cysteine and glycine-rich protein 1 OS=Homo sapiens OX=9606 GN=CSRP1 PE=1 SV=3 | 27 | 5 | 5 | 20.6 | 38 | 193 | 8.57 | 3 | 3 | 2 | 3 | 2 | 3 | 3 | 3 | 2 | 3 | 2 | 3 |
| High | P57764 | Gasdermin-D OS=Homo sapiens OX=9606 GN=GSDMD PE=1 SV=1 | 27 | 7 | 7 | 52.8 | 20 | 484 | 5.08 | 3 | 3 | 6 | 2 | 4 | 1 | 2 | 3 | 6 | 2 | 4 | 1 |
| High | Q5VZK9 | F-actin-uncapping protein LRRC16A OS=Homo sapiens OX=9606 GN=CARMIL1 PE=1 SV=1 | 27 | 13 | 13 | 151.5 | 11 | 1371 | 7.85 | 6 | 7 | 6 | | | 1 | 6 | 7 | 6 | | | 1 |
| High | Q9Y6W5 | Wiskott-Aldrich syndrome protein family member 2 OS=Homo sapiens OX=9606 GN=WASF2 PE=1 SV=3 | 27 | 6 | 6 | 54.3 | 14 | 498 | 5.53 | 3 | 3 | 3 | 1 | 4 | 2 | 3 | 3 | 3 | 1 | 3 | 2 |
| High | P51812 | Ribosomal protein S6 kinase alpha-3 OS=Homo sapiens OX=9606 GN=RPS6KA3 PE=1 SV=1 | 27 | 8 | 4 | 83.7 | 13 | 740 | 6.89 | 1 | 2 | 1 | 3 | 4 | 1 | 1 | 2 | 1 | 2 | 3 | 1 |
| High | P30044 | Peroxiredoxin-5, mitochondrial OS=Homo sapiens OX=9606 GN=PRDX5 PE=1 SV=4 | 27 | 6 | 6 | 22.1 | 39 | 214 | 8.7 | 4 | 1 | 4 | 1 | 1 | 3 | 4 | 1 | 4 | 1 | 1 | 3 |
| High | Q9NSY1 | BMP-2-inducible protein kinase OS=Homo sapiens OX=9606 GN=BMP2K PE=1 SV=2 | 27 | 9 | 9 | 129.1 | 12 | 1161 | 6.51 | 4 | 3 | 4 | | | | 4 | 3 | 4 | | | |
| High | P21796 | Voltage-dependent anion-selective channel protein 1 OS=Homo sapiens OX=9606 GN=VDAC1 PE=1 SV=2 | 27 | 6 | 5 | 30.8 | 26 | 283 | 8.54 | 2 | 2 | 3 | | 2 | 1 | 2 | 2 | 3 | | 2 | 1 |
| High | P51149 | Ras-related protein Rab-7a OS=Homo sapiens OX=9606 GN=RAB7A PE=1 SV=1 | 27 | 6 | 6 | 23.5 | 33 | 207 | 6.7 | 5 | 4 | 4 | | 2 | | 5 | 4 | 4 | | 2 | |
| High | O94906 | Pre-mRNA-processing factor 6 OS=Homo sapiens OX=9606 GN=PRPF6 PE=1 SV=1 | 27 | 11 | 11 | 106.9 | 14 | 941 | 8.25 | 1 | 2 | 1 | | 2 | 1 | 1 | 2 | 1 | | 2 | 1 |
| High | P57735 | Ras-related protein Rab-25 OS=Homo sapiens OX=9606 GN=RAB25 PE=1 SV=2 | 27 | 4 | 3 | 23.5 | 19 | 213 | 5.96 | 2 | 3 | 4 | 2 | 3 | 3 | 2 | 3 | 3 | 2 | 3 | 3 |
| High | Q9H074 | Polyadenylate-binding protein-interacting protein 1 OS=Homo sapiens OX=9606 GN=PAIP1 PE=1 SV=1 | 27 | 6 | 6 | 53.5 | 14 | 479 | 4.81 | 2 | 2 | 3 | 2 | 2 | 3 | 2 | 2 | 3 | 2 | 2 | 2 |
| High | Q3YEC7 | Rab-like protein 6 OS=Homo sapiens OX=9606 GN=RABL6 PE=1 SV=2 | 27 | 7 | 7 | 79.5 | 13 | 729 | 5.22 | 4 | 2 | 3 | 2 | 2 | 1 | 4 | 2 | 3 | 2 | 2 | 1 |
| High | Q2TB90 | Hexokinase HKDC1 OS=Homo sapiens OX=9606 GN=HKDC1 PE=1 SV=3 | 27 | 11 | 8 | 102.5 | 11 | 917 | 7.12 | 3 | 3 | 3 | 1 | 1 | | 3 | 3 | 3 | 1 | 1 | |
| High | Q15645 | Pachytene checkpoint protein 2 homolog OS=Homo sapiens OX=9606 GN=TRIP13 PE=1 SV=2 | 27 | 11 | 11 | 48.5 | 26 | 432 | 6.09 | 4 | 5 | 7 | | 1 | | 4 | 5 | 7 | | 1 | |
| High | Q15008 | 26S proteasome non-ATPase regulatory subunit 6 OS=Homo sapiens OX=9606 GN=PSMD6 PE=1 SV=1 | 27 | 8 | 8 | 45.5 | 21 | 389 | 5.62 | 1 | 2 | 1 | 3 | 4 | 4 | 1 | 2 | 1 | 3 | 4 | 4 |
| High | Q14126 | Desmoglein-2 OS=Homo sapiens OX=9606 GN=DSG2 PE=1 SV=2 | 27 | 7 | 7 | 122.2 | 11 | 1118 | 5.24 | 4 | 5 | 3 | 2 | 1 | 1 | 4 | 5 | 3 | 2 | 1 | 1 |
| High | P50416 | Carnitine O-palmitoyltransferase 1, liver isoform OS=Homo sapiens OX=9606 GN=CPT1A PE=1 SV=2 | 27 | 10 | 10 | 88.3 | 14 | 773 | 8.65 | 7 | 4 | 5 | | | | 7 | 4 | 5 | | | |
| High | Q96I25 | Splicing factor 45 OS=Homo sapiens OX=9606 GN=RBM17 PE=1 SV=1 | 27 | 6 | 6 | 44.9 | 16 | 401 | 5.97 | 2 | 1 | | 3 | 3 | 3 | 2 | 1 | | 3 | 3 | 3 |
| High | P27797 | Calreticulin OS=Homo sapiens OX=9606 GN=CALR PE=1 SV=1 | 27 | 7 | 7 | 48.1 | 23 | 417 | 4.44 | 7 | 5 | 5 | 1 | 1 | 1 | 6 | 5 | 5 | 1 | 1 | 1 |
| High | Q9BQ67 | Glutamate-rich WD repeat-containing protein 1 OS=Homo sapiens OX=9606 GN=GRWD1 PE=1 SV=1 | 26 | 5 | 5 | 49.4 | 17 | 446 | 4.92 | 2 | 2 | 1 | 3 | 3 | 3 | 2 | 2 | 1 | 3 | 3 | 3 |
| High | P61758 | Prefoldin subunit 3 OS=Homo sapiens OX=9606 GN=VBP1 PE=1 SV=4 | 26 | 6 | 6 | 22.6 | 26 | 197 | 7.11 | 1 | | 2 | 4 | 5 | 5 | 1 | | 2 | 4 | 5 | 5 |
| High | Q6PGP7 | Tetratricopeptide repeat protein 37 OS=Homo sapiens OX=9606 GN=TTC37 PE=1 SV=1 | 26 | 12 | 12 | 175.4 | 11 | 1564 | 7.53 | 9 | 6 | 7 | | | | 9 | 6 | 7 | | | |
| High | P21912 | Succinate dehydrogenase [ubiquinone] iron-sulfur subunit, mitochondrial OS=Homo sapiens OX=9606 GN=SDHB PE=1 SV=3 | 26 | 6 | 6 | 31.6 | 22 | 280 | 8.76 | 3 | 3 | 1 | 1 | 5 | 3 | 3 | 3 | 1 | 1 | 4 | 3 |
| High | Q5T440 | Putative transferase CAF17, mitochondrial OS=Homo sapiens OX=9606 GN=IBA57 PE=1 SV=1 | 26 | 4 | 4 | 38.1 | 11 | 356 | 9.83 | | 1 | 3 | 3 | 3 | 4 | | 1 | 3 | 3 | 3 | 4 |
| High | P02768-1 | SWISS-PROT:P02768-1 Tax_Id=9606 Gene_Symbol=ALB Isoform 1 of Serum albumin precursor | 26 | 7 | 3 | 69.3 | 10 | 609 | 6.28 | 2 | 3 | 1 | 3 | 6 | 2 | 2 | 3 | 1 | 3 | 6 | 2 |
| High | Q8N4T8 | 3-oxoacyl-[acyl-carrier-protein] reductase OS=Homo sapiens OX=9606 GN=CBR4 PE=1 SV=3 | 26 | 6 | 6 | 25.3 | 35 | 237 | 9.33 | 4 | 2 | 2 | 2 | 3 | 2 | 4 | 2 | 2 | 2 | 3 | 2 |
| High | P61619 | Protein transport protein Sec61 subunit alpha isoform 1 OS=Homo sapiens OX=9606 GN=SEC61A1 PE=1 SV=2 | 26 | 5 | 5 | 52.2 | 11 | 476 | 8.06 | 2 | 2 | 2 | 3 | 2 | 3 | 2 | 2 | 2 | 3 | 2 | 3 |
| High | Q12800 | Alpha-globin transcription factor CP2 OS=Homo sapiens OX=9606 GN=TFCP2 PE=1 SV=2 | 26 | 7 | 6 | 57.2 | 21 | 502 | 5.8 | 6 | 7 | 5 | | 1 | | 6 | 7 | 5 | | 1 | |
| High | Q9Y6E2 | eIF5-mimic protein 1 OS=Homo sapiens OX=9606 GN=BZW2 PE=1 SV=1 | 26 | 10 | 9 | 48.1 | 24 | 419 | 6.68 | 2 | 1 | 3 | 2 | 4 | 2 | 2 | 1 | 3 | 2 | 4 | 2 |
| High | Q8N684 | Cleavage and polyadenylation specificity factor subunit 7 OS=Homo sapiens OX=9606 GN=CPSF7 PE=1 SV=1 | 26 | 7 | 7 | 52 | 16 | 471 | 8 | | | | 3 | 6 | 4 | | | | 3 | 6 | 4 |
| High | P13796 | Plastin-2 OS=Homo sapiens OX=9606 GN=LCP1 PE=1 SV=6 | 26 | 7 | 1 | 70.2 | 11 | 627 | 5.43 | 1 | 2 | 2 | 3 | 5 | 5 | 1 | 2 | 2 | 3 | 5 | 5 |
| High | O14618 | Copper chaperone for superoxide dismutase OS=Homo sapiens OX=9606 GN=CCS PE=1 SV=1 | 26 | 5 | 5 | 29 | 16 | 274 | 5.58 | 1 | 1 | | 2 | 5 | 3 | 1 | 1 | | 2 | 5 | 3 |
| High | P52292 | Importin subunit alpha-1 OS=Homo sapiens OX=9606 GN=KPNA2 PE=1 SV=1 | 26 | 7 | 7 | 57.8 | 16 | 529 | 5.4 | 1 | | 3 | 3 | 4 | 4 | 1 | | 3 | 3 | 4 | 4 |
| High | Q13765 | Nascent polypeptide-associated complex subunit alpha OS=Homo sapiens OX=9606 GN=NACA PE=1 SV=1 | 26 | 3 | 3 | 23.4 | 20 | 215 | 4.56 | 3 | 2 | 3 | 3 | 2 | 2 | 2 | 1 | 2 | 2 | 1 | 1 |
| High | O94903 | Pyridoxal phosphate homeostasis protein OS=Homo sapiens OX=9606 GN=PLPBP PE=1 SV=1 | 26 | 7 | 7 | 30.3 | 29 | 275 | 7.5 | 1 | 3 | 3 | 4 | | 4 | 1 | 3 | 2 | 4 | | 4 |
| High | O00505 | Importin subunit alpha-4 OS=Homo sapiens OX=9606 GN=KPNA3 PE=1 SV=2 | 26 | 6 | 3 | 57.8 | 13 | 521 | 4.94 | | 1 | 2 | 2 | 3 | 2 | | 1 | 2 | 2 | 3 | 2 |
| High | Q9NQ88 | Fructose-2,6-bisphosphatase TIGAR OS=Homo sapiens OX=9606 GN=TIGAR PE=1 SV=1 | 26 | 5 | 5 | 30 | 26 | 270 | 7.69 | 1 | 2 | 2 | 2 | 3 | 2 | 1 | 2 | 2 | 2 | 3 | 2 |
| High | Q8NAP3 | Zinc finger and BTB domain-containing protein 38 OS=Homo sapiens OX=9606 GN=ZBTB38 PE=1 SV=2 | 26 | 8 | 8 | 134.2 | 7 | 1195 | 8.03 | | | | 3 | 4 | 3 | | | | 3 | 4 | 3 |
| High | O15305 | Phosphomannomutase 2 OS=Homo sapiens OX=9606 GN=PMM2 PE=1 SV=1 | 26 | 6 | 6 | 28.1 | 23 | 246 | 6.77 | 1 | 1 | 2 | 3 | 3 | 2 | 1 | 1 | 2 | 3 | 3 | 2 |
| High | Q12849 | G-rich sequence factor 1 OS=Homo sapiens OX=9606 GN=GRSF1 PE=1 SV=3 | 26 | 7 | 7 | 53.1 | 19 | 480 | 6.19 | | 2 | 1 | 3 | 3 | 3 | | 2 | 1 | 3 | 3 | 3 |
| High | Q02750 | Dual specificity mitogen-activated protein kinase kinase 1 OS=Homo sapiens OX=9606 GN=MAP2K1 PE=1 SV=2 | 26 | 7 | 5 | 43.4 | 25 | 393 | 6.62 | 4 | 4 | 4 | 1 | 2 | 1 | 4 | 4 | 3 | 1 | 2 | 1 |
| High | P40926 | Malate dehydrogenase, mitochondrial OS=Homo sapiens OX=9606 GN=MDH2 PE=1 SV=3 | 26 | 8 | 8 | 35.5 | 29 | 338 | 8.68 | 3 | 1 | 6 | 5 | 3 | 5 | 3 | 1 | 6 | 5 | 2 | 5 |
| High | Q8N2K0 | Lysophosphatidylserine lipase ABHD12 OS=Homo sapiens OX=9606 GN=ABHD12 PE=1 SV=2 | 26 | 7 | 7 | 45.1 | 21 | 398 | 8.65 | 4 | 4 | 4 | 1 | 2 | 1 | 4 | 4 | 2 | 1 | 2 | 1 |
| High | P62899 | 60S ribosomal protein L31 OS=Homo sapiens OX=9606 GN=RPL31 PE=1 SV=1 | 26 | 3 | 3 | 14.5 | 25 | 125 | 10.54 | 3 | 3 | 3 | 2 | 2 | 2 | 3 | 3 | 2 | 2 | 2 | 2 |
| High | P53007 | Tricarboxylate transport protein, mitochondrial OS=Homo sapiens OX=9606 GN=SLC25A1 PE=1 SV=2 | 26 | 5 | 5 | 34 | 17 | 311 | 9.89 | 3 | 4 | 2 | 2 | 1 | 1 | 3 | 3 | 1 | 2 | 1 | 1 |
| High | Q13573 | SNW domain-containing protein 1 OS=Homo sapiens OX=9606 GN=SNW1 PE=1 SV=1 | 26 | 7 | 7 | 61.5 | 15 | 536 | 9.52 | 3 | 3 | 2 | 1 | 2 | 3 | 3 | 3 | 2 | 1 | 2 | 3 |
| High | O43823 | A-kinase anchor protein 8 OS=Homo sapiens OX=9606 GN=AKAP8 PE=1 SV=1 | 26 | 10 | 10 | 76.1 | 22 | 692 | 5.15 | 1 | 3 | 3 | | 1 | | 1 | 3 | 3 | | 1 | |
| High | Q12789 | General transcription factor 3C polypeptide 1 OS=Homo sapiens OX=9606 GN=GTF3C1 PE=1 SV=4 | 26 | 13 | 13 | 238.7 | 8 | 2109 | 7.3 | 1 | | 2 | | 5 | 2 | 1 | | 2 | | 5 | 2 |
| High | O60381 | HMG box-containing protein 1 OS=Homo sapiens OX=9606 GN=HBP1 PE=1 SV=2 | 26 | 9 | 9 | 57.6 | 25 | 514 | 6.19 | | | | 2 | 4 | 2 | | | | 2 | 4 | 2 |
| High | Q9NX24 | H/ACA ribonucleoprotein complex subunit 2 OS=Homo sapiens OX=9606 GN=NHP2 PE=1 SV=1 | 26 | 3 | 3 | 17.2 | 30 | 153 | 8.22 | 2 | 2 | 3 | 1 | 2 | 1 | 2 | 2 | 3 | 1 | 2 | 1 |
| High | Q9P287 | BRCA2 and CDKN1A-interacting protein OS=Homo sapiens OX=9606 GN=BCCIP PE=1 SV=1 | 26 | 6 | 6 | 36 | 25 | 314 | 4.61 | 4 | 4 | 4 | 2 | 1 | 1 | 4 | 4 | 4 | 2 | 1 | 1 |
| High | Q96IU4 | Protein ABHD14B OS=Homo sapiens OX=9606 GN=ABHD14B PE=1 SV=1 | 26 | 8 | 8 | 22.3 | 48 | 210 | 6.4 | 5 | 5 | 5 | 1 | 1 | | 5 | 5 | 5 | 1 | 1 | |

| | | | | | | | | | | | | | | | | | | | | | |
|---|---|---|---|---|---|---|---|---|---|---|---|---|---|---|---|---|---|---|---|---|---|
| High | P50851 | Lipopolysaccharide-responsive and beige-like anchor protein OS=Homo sapiens OX=9606 GN=LRBA PE=1 SV=4 | 26 | 12 | 12 | 318.9 | 5 | 2863 | 5.6 | 2 | 2 | 7 | 1 | | 1 | 2 | 2 | 7 | 1 | | 1 |
| High | P16455 | Methylated-DNA--protein-cysteine methyltransferase OS=Homo sapiens OX=9606 GN=MGMT PE=1 SV=1 | 26 | 6 | 6 | 21.6 | 33 | 207 | 8.1 | 3 | 3 | 2 | 2 | 3 | 2 | 3 | 3 | 2 | 2 | 3 | 2 |
| High | Q92552 | 28S ribosomal protein S27, mitochondrial OS=Homo sapiens OX=9606 GN=MRPS27 PE=1 SV=3 | 26 | 9 | 9 | 47.6 | 25 | 414 | 6.18 | 2 | 3 | 1 | 2 | 2 | 3 | 2 | 3 | 1 | 2 | 2 | 3 |
| High | Q9BRK5 | 45 kDa calcium-binding protein OS=Homo sapiens OX=9606 GN=SDF4 PE=1 SV=1 | 26 | 6 | 6 | 41.8 | 20 | 362 | 4.86 | 5 | 5 | 6 | 2 | 1 | 2 | 4 | 4 | 6 | 2 | 1 | 2 |
| High | Q96SB4 | SRSF protein kinase 1 OS=Homo sapiens OX=9606 GN=SRPK1 PE=1 SV=2 | 26 | 9 | 7 | 74.3 | 17 | 655 | 6.16 | | | 2 | 2 | 4 | 3 | | | 2 | 2 | 4 | 3 |
| High | P62857 | 40S ribosomal protein S28 OS=Homo sapiens OX=9606 GN=RPS28 PE=1 SV=1 | 26 | 2 | 2 | 7.8 | 30 | 69 | 10.7 | 3 | 2 | 3 | 2 | 3 | 3 | 2 | 2 | 2 | 1 | 2 | 2 |
| High | O00410 | Importin-5 OS=Homo sapiens OX=9606 GN=IPO5 PE=1 SV=4 | 26 | 7 | 7 | 123.6 | 9 | 1097 | 4.94 | 3 | 2 | | 1 | | 2 | 3 | 2 | | 1 | | 2 |
| High | Q9NY12 | H/ACA ribonucleoprotein complex subunit 1 OS=Homo sapiens OX=9606 GN=GAR1 PE=1 SV=1 | 25 | 4 | 4 | 22.3 | 25 | 217 | 10.92 | 3 | 3 | 4 | 2 | 1 | 1 | 3 | 3 | 4 | 2 | 1 | 1 |
| High | P43304 | Glycerol-3-phosphate dehydrogenase, mitochondrial OS=Homo sapiens OX=9606 GN=GPD2 PE=1 SV=3 | 25 | 10 | 10 | 80.8 | 15 | 727 | 7.69 | | | | 2 | 2 | 4 | | | | 2 | 2 | 4 |
| High | Q9UKV3 | Apoptotic chromatin condensation inducer in the nucleus OS=Homo sapiens OX=9606 GN=ACIN1 PE=1 SV=2 | 25 | 8 | 8 | 151.8 | 8 | 1341 | 6.43 | 2 | 1 | 2 | 4 | 5 | 2 | 2 | 1 | 2 | 4 | 4 | 2 |
| High | P63165 | Small ubiquitin-related modifier 1 OS=Homo sapiens OX=9606 GN=SUMO1 PE=1 SV=1 | 25 | 3 | 3 | 11.6 | 28 | 101 | 5.52 | | | | 1 | 2 | 1 | | | | 1 | 2 | 1 |
| High | O60784 | Target of Myb protein 1 OS=Homo sapiens OX=9606 GN=TOM1 PE=1 SV=2 | 25 | 7 | 7 | 53.8 | 25 | 492 | 4.7 | 3 | 3 | 3 | | | | 3 | 3 | 3 | | | |
| High | Q14244 | Ensconsin OS=Homo sapiens OX=9606 GN=MAP7 PE=1 SV=1 | 25 | 5 | 5 | 84 | 7 | 749 | 9.61 | 2 | 3 | 3 | 2 | 3 | 2 | 2 | 3 | 3 | 2 | 3 | 2 |
| High | O15145 | Actin-related protein 2/3 complex subunit 3 OS=Homo sapiens OX=9606 GN=ARPC3 PE=1 SV=3 | 25 | 4 | 4 | 20.5 | 21 | 178 | 8.59 | 4 | 3 | 3 | 1 | 2 | 1 | 4 | 3 | 3 | 1 | 2 | 1 |
| High | Q9Y5M8 | Signal recognition particle receptor subunit beta OS=Homo sapiens OX=9606 GN=SRPRB PE=1 SV=3 | 25 | 6 | 6 | 29.7 | 27 | 271 | 9.04 | 3 | 2 | 4 | 2 | 2 | 2 | 3 | 2 | 4 | 2 | 2 | 2 |
| High | P62070 | Ras-related protein R-Ras2 OS=Homo sapiens OX=9606 GN=RRAS2 PE=1 SV=1 | 25 | 5 | 3 | 23.4 | 31 | 204 | 6.01 | 3 | 4 | 3 | 1 | 1 | 1 | 3 | 4 | 2 | 1 | 1 | 1 |
| High | P08559 | Pyruvate dehydrogenase E1 component subunit alpha, somatic form, mitochondrial OS=Homo sapiens OX=9606 GN=PDHA1 PE=1 | 25 | 9 | 9 | 43.3 | 22 | 390 | 8.06 | 1 | | | 2 | 3 | 1 | 1 | | | 2 | 3 | 1 |
| High | Q9NR19 | Acetyl-coenzyme A synthetase, cytoplasmic OS=Homo sapiens OX=9606 GN=ACSS2 PE=1 SV=1 | 25 | 5 | 5 | 78.5 | 6 | 701 | 6.46 | 3 | 3 | 4 | 2 | 2 | 1 | 3 | 3 | 4 | 2 | 2 | 1 |
| High | Q99471 | Prefoldin subunit 5 OS=Homo sapiens OX=9606 GN=PFDN5 PE=1 SV=2 | 25 | 5 | 5 | 17.3 | 38 | 154 | 6.33 | 2 | 2 | 2 | 2 | 2 | 2 | 2 | 2 | 2 | 2 | 2 | 2 |
| High | Q92925 | SWI/SNF-related matrix-associated actin-dependent regulator of chromatin subfamily D member 2 OS=Homo sapiens OX=9606 G | 25 | 8 | 7 | 58.9 | 19 | 531 | 9.64 | 1 | 1 | 1 | 2 | 2 | 1 | 1 | 1 | 1 | 2 | 2 | 1 |
| High | O95299 | NADH dehydrogenase [ubiquinone] 1 alpha subcomplex subunit 10, mitochondrial OS=Homo sapiens OX=9606 GN=NDUFA10 PE= | 25 | 4 | 4 | 40.7 | 11 | 355 | 8.48 | 2 | 3 | 3 | 2 | 2 | 3 | 2 | 3 | 3 | 2 | 2 | 3 |
| High | O75663 | TIP41-like protein OS=Homo sapiens OX=9606 GN=TIPRL PE=1 SV=2 | 25 | 7 | 7 | 31.4 | 38 | 272 | 5.91 | 4 | 3 | 3 | 1 | 2 | | 4 | 3 | 3 | 1 | 2 | |
| High | Q05193 | Dynamin-1 OS=Homo sapiens OX=9606 GN=DNM1 PE=1 SV=2 | 25 | 6 | 1 | 97.3 | 7 | 864 | 7.17 | 4 | 3 | 5 | | | 2 | 4 | 3 | 5 | | | 2 |
| High | P60953 | Cell division control protein 42 homolog OS=Homo sapiens OX=9606 GN=CDC42 PE=1 SV=2 | 25 | 4 | 3 | 21.2 | 26 | 191 | 6.55 | 3 | 5 | 2 | 2 | 2 | 2 | 3 | 4 | 2 | 2 | 2 | 2 |
| High | O00193 | Small acidic protein OS=Homo sapiens OX=9606 GN=SMAP PE=1 SV=1 | 25 | 4 | 4 | 20.3 | 22 | 183 | 4.72 | 1 | 1 | 3 | 2 | 4 | 3 | 1 | 1 | 3 | 2 | 4 | 2 |
| High | O94776 | Metastasis-associated protein MTA2 OS=Homo sapiens OX=9606 GN=MTA2 PE=1 SV=1 | 25 | 8 | 7 | 75 | 14 | 668 | 9.66 | 1 | 2 | 4 | 2 | 3 | 1 | 1 | 2 | 3 | 2 | 3 | 1 |
| High | Q8TD30 | Alanine aminotransferase 2 OS=Homo sapiens OX=9606 GN=GPT2 PE=1 SV=1 | 25 | 8 | 8 | 57.9 | 25 | 523 | 7.71 | 1 | 4 | 4 | | 2 | | 1 | 4 | 4 | | 2 | |
| High | Q93052 | Lipoma-preferred partner OS=Homo sapiens OX=9606 GN=LPP PE=1 SV=1 | 25 | 6 | 6 | 65.7 | 17 | 612 | 7.37 | 2 | 2 | 2 | 3 | 3 | 2 | 2 | 2 | 2 | 3 | 3 | 2 |
| High | Q96FJ0 | AMSH-like protease OS=Homo sapiens OX=9606 GN=STAMBPL1 PE=1 SV=2 | 25 | 6 | 6 | 49.8 | 20 | 436 | 7.23 | 3 | 3 | 4 | 1 | | 1 | 3 | 3 | 4 | 1 | | 1 |
| High | P98082 | Disabled homolog 2 OS=Homo sapiens OX=9606 GN=DAB2 PE=1 SV=3 | 25 | 7 | 7 | 82.4 | 17 | 770 | 5.53 | 2 | 2 | 4 | | | | 2 | 2 | 4 | | | |
| High | O00469 | Procollagen-lysine,2-oxoglutarate 5-dioxygenase 2 OS=Homo sapiens OX=9606 GN=PLOD2 PE=1 SV=2 | 25 | 10 | 10 | 84.6 | 16 | 737 | 6.71 | 2 | | 3 | | 2 | | 2 | | 3 | | 2 | |
| High | Q14498 | RNA-binding protein 39 OS=Homo sapiens OX=9606 GN=RBM39 PE=1 SV=2 | 25 | 5 | 5 | 59.3 | 12 | 530 | 10.1 | 2 | 2 | 2 | 2 | 2 | 2 | 2 | 2 | 2 | 2 | 2 | 2 |
| High | P31948 | Stress-induced-phosphoprotein 1 OS=Homo sapiens OX=9606 GN=STIP1 PE=1 SV=1 | 25 | 7 | 7 | 62.6 | 16 | 543 | 6.8 | 4 | 3 | 5 | 1 | 2 | 1 | 4 | 3 | 5 | 1 | 2 | 1 |
| High | Q92506 | (3R)-3-hydroxyacyl-CoA dehydrogenase OS=Homo sapiens OX=9606 GN=HSD17B8 PE=1 SV=2 | 25 | 5 | 5 | 27 | 25 | 261 | 6.54 | 1 | 1 | 1 | 3 | 3 | 3 | 1 | 1 | 1 | 3 | 3 | 3 |
| High | P09211 | Glutathione S-transferase P OS=Homo sapiens OX=9606 GN=GSTP1 PE=1 SV=2 | 25 | 5 | 5 | 23.3 | 36 | 210 | 5.64 | 3 | 3 | 3 | 1 | 2 | 1 | 3 | 3 | 3 | 1 | 2 | 1 |
| High | O75436 | Vacuolar protein sorting-associated protein 26A OS=Homo sapiens OX=9606 GN=VPS26A PE=1 SV=2 | 25 | 5 | 4 | 38.1 | 22 | 327 | 6.57 | 2 | 3 | 3 | 3 | 4 | 2 | 2 | 3 | 3 | 3 | 4 | 2 |
| High | Q9NQT8 | Kinesin-like protein KIF13B OS=Homo sapiens OX=9606 GN=KIF13B PE=1 SV=2 | 25 | 7 | 6 | 202.7 | 5 | 1826 | 5.88 | 4 | 4 | 2 | | 2 | | 4 | 4 | 2 | | 2 | |
| High | Q16543 | Hsp90 co-chaperone Cdc37 OS=Homo sapiens OX=9606 GN=CDC37 PE=1 SV=1 | 25 | 7 | 7 | 44.4 | 21 | 378 | 5.25 | 2 | 2 | 5 | 4 | 2 | 2 | 2 | 2 | 5 | 4 | 2 | 2 |
| High | Q9Y6R7 | IgGFc-binding protein OS=Homo sapiens OX=9606 GN=FCGBP PE=1 SV=3 | 25 | 11 | 11 | 571.6 | 6 | 5405 | 5.34 | | | | | | | | | | | | |
| High | Q9Y5A9 | YTH domain-containing family protein 2 OS=Homo sapiens OX=9606 GN=YTHDF2 PE=1 SV=2 | 25 | 4 | 3 | 62.3 | 7 | 579 | 8.79 | 3 | 3 | 3 | 3 | 3 | 2 | 3 | 3 | 3 | 3 | 3 | 2 |
| High | Q9UHD2 | Serine/threonine-protein kinase TBK1 OS=Homo sapiens OX=9606 GN=TBK1 PE=1 SV=1 | 25 | 9 | 9 | 83.6 | 14 | 729 | 6.79 | 2 | 1 | 2 | 1 | 2 | | 2 | 1 | 2 | 1 | 2 | |
| High | Q8TCT9 | Minor histocompatibility antigen H13 OS=Homo sapiens OX=9606 GN=HM13 PE=1 SV=1 | 25 | 5 | 5 | 41.5 | 16 | 377 | 6.43 | 2 | 3 | 3 | | 3 | 1 | 2 | 3 | 3 | | 3 | 1 |
| High | O75643 | U5 small nuclear ribonucleoprotein 200 kDa helicase OS=Homo sapiens OX=9606 GN=SNRNP200 PE=1 SV=2 | 25 | 15 | 15 | 244.4 | 10 | 2136 | 6.06 | 3 | | 2 | 1 | 3 | | 3 | | 2 | 1 | 3 | |
| High | Q92905 | COP9 signalosome complex subunit 5 OS=Homo sapiens OX=9606 GN=COPS5 PE=1 SV=4 | 25 | 6 | 6 | 37.6 | 23 | 334 | 6.54 | 2 | 2 | 4 | | 3 | | 2 | 2 | 4 | | 3 | |
| High | P08758 | Annexin A5 OS=Homo sapiens OX=9606 GN=ANXA5 PE=1 SV=2 | 25 | 8 | 8 | 35.9 | 26 | 320 | 5.05 | 3 | 2 | 3 | 4 | 5 | 4 | 3 | 2 | 3 | 2 | 5 | 3 |
| High | Q9NZ01 | Very-long-chain enoyl-CoA reductase OS=Homo sapiens OX=9606 GN=TECR PE=1 SV=1 | 25 | 8 | 8 | 36 | 25 | 308 | 9.45 | 5 | 6 | 3 | | | 1 | 5 | 5 | 3 | | | 1 |
| High | Q9H7C9 | Mth938 domain-containing protein OS=Homo sapiens OX=9606 GN=AAMDC PE=1 SV=1 | 25 | 5 | 5 | 13.3 | 43 | 122 | 8.46 | 2 | | 3 | 3 | | 4 | 2 | | 3 | 3 | | 3 |
| High | Q9P035 | Very-long-chain (3R)-3-hydroxyacyl-CoA dehydratase 3 OS=Homo sapiens OX=9606 GN=HACD3 PE=1 SV=2 | 24 | 5 | 5 | 43.1 | 14 | 362 | 8.94 | 5 | 4 | 2 | 1 | 2 | | 5 | 4 | 2 | 1 | 2 | |
| High | P28331 | NADH-ubiquinone oxidoreductase 75 kDa subunit, mitochondrial OS=Homo sapiens OX=9606 GN=NDUFS1 PE=1 SV=3 | 24 | 11 | 11 | 79.4 | 19 | 727 | 6.23 | 1 | 2 | 3 | 1 | 2 | | 1 | 2 | 3 | 1 | 2 | |
| High | A3KMH1 | von Willebrand factor A domain-containing protein 8 OS=Homo sapiens OX=9606 GN=VWA8 PE=1 SV=2 | 24 | 7 | 7 | 214.7 | 4 | 1905 | 7.4 | 7 | 6 | 3 | | 1 | | 7 | 6 | 3 | | 1 | |
| High | P11908 | Ribose-phosphate pyrophosphokinase 2 OS=Homo sapiens OX=9606 GN=PRPS2 PE=1 SV=2 | 24 | 5 | 1 | 34.7 | 20 | 318 | 6.61 | 3 | 3 | 3 | 3 | 1 | 3 | 3 | 3 | 3 | 3 | 1 | 3 |
| High | Q9UBV8 | Peflin OS=Homo sapiens OX=9606 GN=PEF1 PE=1 SV=1 | 24 | 4 | 4 | 30.4 | 13 | 284 | 6.54 | 3 | 3 | 3 | 3 | 2 | 1 | 3 | 3 | 3 | 3 | 2 | 1 |
| High | Q9H2P0 | Activity-dependent neuroprotector homeobox protein OS=Homo sapiens OX=9606 GN=ADNP PE=1 SV=1 | 24 | 7 | 7 | 123.5 | 8 | 1102 | 7.34 | 3 | 2 | 3 | 2 | 2 | 3 | 3 | 2 | 3 | 2 | 2 | 3 |
| High | Q8IYS1 | Xaa-Arg dipeptidase OS=Homo sapiens OX=9606 GN=PM20D2 PE=1 SV=2 | 24 | 3 | 3 | 47.7 | 11 | 436 | 5.85 | 2 | 3 | 2 | 2 | 2 | 2 | 2 | 3 | 2 | 2 | 2 | 2 |
| High | P62837 | Ubiquitin-conjugating enzyme E2 D2 OS=Homo sapiens OX=9606 GN=UBE2D2 PE=1 SV=1 | 24 | 4 | 3 | 16.7 | 41 | 147 | 7.83 | 2 | 4 | 4 | 1 | 3 | 2 | 2 | 4 | 3 | 1 | 2 | 2 |
| High | P40429 | 60S ribosomal protein L13a OS=Homo sapiens OX=9606 GN=RPL13A PE=1 SV=2 | 24 | 4 | 4 | 23.6 | 21 | 203 | 10.93 | 2 | 1 | 2 | 2 | 2 | 2 | 2 | 1 | 1 | 2 | 2 | 2 |
| High | O14980 | Exportin-1 OS=Homo sapiens OX=9606 GN=XPO1 PE=1 SV=1 | 24 | 9 | 9 | 123.3 | 9 | 1071 | 6.06 | 3 | 4 | 2 | | 2 | | 2 | 4 | 2 | | 2 | |
| High | P12882 | Myosin-1 OS=Homo sapiens OX=9606 GN=MYH1 PE=1 SV=3 | 24 | 9 | 4 | 223 | 6 | 1939 | 5.74 | 2 | 2 | 4 | 3 | 6 | 3 | 2 | 1 | 2 | 3 | 6 | 3 |
| High | P62266 | 40S ribosomal protein S23 OS=Homo sapiens OX=9606 GN=RPS23 PE=1 SV=3 | 24 | 3 | 3 | 15.8 | 21 | 143 | 10.49 | 2 | 2 | 3 | 2 | 2 | 3 | 2 | 2 | 3 | 2 | 2 | 2 |
| High | P02452 | Collagen alpha-1(I) chain OS=Homo sapiens OX=9606 GN=COL1A1 PE=1 SV=6 | 24 | 4 | 4 | 138.8 | 3 | 1464 | 5.8 | | | | 5 | 14 | 5 | | | | 3 | 4 | 3 |
| High | Q7L2E3 | ATP-dependent RNA helicase DHX30 OS=Homo sapiens OX=9606 GN=DHX30 PE=1 SV=1 | 24 | 8 | 8 | 133.9 | 9 | 1194 | 8.78 | 1 | 1 | 1 | 2 | 2 | 3 | 1 | 1 | 1 | 2 | 2 | 3 |
| High | Q9Y3A5 | Ribosome maturation protein SBDS OS=Homo sapiens OX=9606 GN=SBDS PE=1 SV=4 | 24 | 6 | 6 | 28.7 | 24 | 250 | 8.75 | 4 | 3 | 3 | 1 | 1 | 1 | 4 | 3 | 3 | 1 | 1 | 1 |
| High | Q96HE7 | ERO1-like protein alpha OS=Homo sapiens OX=9606 GN=ERO1A PE=1 SV=2 | 24 | 9 | 9 | 54.4 | 27 | 468 | 5.68 | 2 | 3 | 2 | 2 | 1 | | 2 | 3 | 2 | 2 | 1 | |
| High | P02584 | SWISS-PROT:P02584 (Bos taurus) Profilin-1 | 24 | 4 | 1 | 15 | 31 | 140 | 8.28 | 3 | 3 | 3 | 3 | 2 | 2 | 3 | 3 | 3 | 3 | 2 | 2 |
| High | Q99700 | Ataxin-2 OS=Homo sapiens OX=9606 GN=ATXN2 PE=1 SV=2 | 24 | 5 | 5 | 140.2 | 5 | 1313 | 9.57 | 1 | 1 | 1 | 4 | 2 | 4 | 1 | 1 | 1 | 4 | 2 | 4 |
| High | Q02978 | Mitochondrial 2-oxoglutarate/malate carrier protein OS=Homo sapiens OX=9606 GN=SLC25A11 PE=1 SV=3 | 24 | 7 | 7 | 34 | 23 | 314 | 9.91 | 5 | 4 | 7 | | | | 5 | 4 | 7 | | | |
| High | Q15382 | GTP-binding protein Rheb OS=Homo sapiens OX=9606 GN=RHEB PE=1 SV=1 | 24 | 5 | 5 | 20.5 | 28 | 184 | 5.92 | 2 | 3 | 2 | 1 | 2 | 1 | 2 | 3 | 2 | 1 | 2 | 1 |
| High | Q9HC78 | Zinc finger and BTB domain-containing protein 20 OS=Homo sapiens OX=9606 GN=ZBTB20 PE=1 SV=3 | 24 | 10 | 10 | 81 | 20 | 741 | 6.49 | | | | | | | | | | | | |
| High | P20340 | Ras-related protein Rab-6A OS=Homo sapiens OX=9606 GN=RAB6A PE=1 SV=3 | 24 | 5 | 4 | 23.6 | 25 | 208 | 5.54 | 3 | 2 | 3 | 2 | 2 | 1 | 2 | 1 | 2 | 2 | 1 | 1 |
| High | Q9UEY8 | Gamma-adducin OS=Homo sapiens OX=9606 GN=ADD3 PE=1 SV=1 | 24 | 9 | 9 | 79.1 | 16 | 706 | 6.32 | | 1 | 2 | | 4 | | | 1 | 2 | | 4 | |
| High | Q9BY42 | Replication termination factor 2 OS=Homo sapiens OX=9606 GN=RTF2 PE=1 SV=3 | 24 | 6 | 6 | 33.9 | 25 | 306 | 8.59 | 1 | 3 | 2 | 4 | 4 | 3 | 1 | 3 | 2 | 4 | 4 | 3 |
| High | P38606 | V-type proton ATPase catalytic subunit A OS=Homo sapiens OX=9606 GN=ATP6V1A PE=1 SV=2 | 23 | 8 | 8 | 68.3 | 16 | 617 | 5.52 | 4 | 4 | 2 | | | 1 | 4 | 4 | 2 | | | 1 |
| High | Q9BQA1 | Methylosome protein 50 OS=Homo sapiens OX=9606 GN=WDR77 PE=1 SV=1 | 23 | 6 | 6 | 36.7 | 25 | 342 | 5.17 | | 1 | 2 | 4 | 4 | 2 | | 1 | 2 | 3 | 3 | 2 |
| High | Q9BV57 | 1,2-dihydroxy-3-keto-5-methylthiopentene dioxygenase OS=Homo sapiens OX=9606 GN=ADI1 PE=1 SV=1 | 23 | 4 | 4 | 21.5 | 24 | 179 | 5.68 | 1 | 3 | 2 | 2 | 2 | 2 | 1 | 2 | 2 | 2 | 2 | 2 |
| High | Q8IZP0 | Abl interactor 1 OS=Homo sapiens OX=9606 GN=ABI1 PE=1 SV=4 | 23 | 7 | 4 | 55 | 18 | 508 | 7.06 | 2 | 2 | | 1 | 1 | 1 | 2 | 2 | | 1 | 1 | 1 |
| High | Q9BW27 | Nuclear pore complex protein Nup85 OS=Homo sapiens OX=9606 GN=NUP85 PE=1 SV=1 | 23 | 10 | 10 | 75 | 21 | 656 | 5.55 | 1 | 2 | 2 | | | 1 | 1 | 2 | 2 | | | 1 |
| High | Q8WXI9 | Transcriptional repressor p66-beta OS=Homo sapiens OX=9606 GN=GATAD2B PE=1 SV=1 | 23 | 5 | 4 | 65.2 | 13 | 593 | 9.7 | | 2 | 2 | 2 | 3 | 3 | | 2 | 2 | 2 | 3 | 3 |
| High | Q9UBE0 | SUMO-activating enzyme subunit 1 OS=Homo sapiens OX=9606 GN=SAE1 PE=1 SV=1 | 23 | 8 | 8 | 38.4 | 30 | 346 | 5.3 | 3 | 3 | 6 | | 1 | | 3 | 3 | 6 | | 1 | |
| High | Q9Y263 | Phospholipase A-2-activating protein OS=Homo sapiens OX=9606 GN=PLAA PE=1 SV=2 | 23 | 8 | 8 | 87.1 | 16 | 795 | 6.37 | 1 | | 3 | 2 | 2 | 2 | 1 | | 3 | 2 | 2 | 2 |
| High | Q5TC82 | Roquin-1 OS=Homo sapiens OX=9606 GN=RC3H1 PE=1 SV=1 | 23 | 9 | 5 | 125.7 | 12 | 1133 | 7.27 | | | | 5 | 2 | 4 | | | | 4 | 2 | 4 |
| High | O43615 | Mitochondrial import inner membrane translocase subunit TIM44 OS=Homo sapiens OX=9606 GN=TIMM44 PE=1 SV=2 | 23 | 10 | 10 | 51.3 | 25 | 452 | 8.32 | 4 | 4 | 6 | | 1 | | 4 | 4 | 6 | | 1 | |
| High | O43175 | D-3-phosphoglycerate dehydrogenase OS=Homo sapiens OX=9606 GN=PHGDH PE=1 SV=4 | 23 | 10 | 10 | 56.6 | 21 | 533 | 6.71 | 1 | 2 | 1 | | | 1 | 1 | 2 | 1 | | | 1 |
| High | Q14653 | Interferon regulatory factor 3 OS=Homo sapiens OX=9606 GN=IRF3 PE=1 SV=1 | 23 | 6 | 6 | 47.2 | 20 | 427 | 5.34 | 3 | 4 | 6 | 1 | | 1 | 3 | 3 | 5 | 1 | | 1 |
| High | P42566 | Epidermal growth factor receptor substrate 15 OS=Homo sapiens OX=9606 GN=EPS15 PE=1 SV=2 | 23 | 7 | 7 | 98.6 | 8 | 896 | 4.64 | 2 | 1 | 1 | | 1 | | 2 | 1 | 1 | | 1 | |
| High | Q9UBI6 | Guanine nucleotide-binding protein G(I)/G(S)/G(O) subunit gamma-12 OS=Homo sapiens OX=9606 GN=GNG12 PE=1 SV=3 | 23 | 4 | 4 | 8 | 69 | 72 | 8.97 | 2 | 3 | 3 | 1 | 1 | 2 | 2 | 3 | 3 | 1 | 1 | 2 |
| High | Q96C86 | m7GpppX diphosphatase OS=Homo sapiens OX=9606 GN=DCPS PE=1 SV=2 | 23 | 6 | 6 | 38.6 | 21 | 337 | 6.38 | 1 | 1 | 1 | 3 | 5 | 1 | 1 | 1 | 1 | 3 | 5 | 1 |
| High | P49006 | MARCKS-related protein OS=Homo sapiens OX=9606 GN=MARCKSL1 PE=1 SV=2 | 23 | 5 | 5 | 19.5 | 55 | 195 | 4.67 | 2 | 2 | 2 | 1 | 2 | 1 | 2 | 2 | 2 | 1 | 2 | 1 |
| High | Q9H6Q4 | Cytosolic iron-sulfur assembly component 3 OS=Homo sapiens OX=9606 GN=CIAO3 PE=1 SV=1 | 23 | 8 | 8 | 53 | 26 | 476 | 7.23 | 2 | 2 | 3 | 3 | 1 | 3 | 2 | 2 | 3 | 3 | 1 | 3 |
| High | P40121 | Macrophage-capping protein OS=Homo sapiens OX=9606 GN=CAPG PE=1 SV=2 | 23 | 6 | 6 | 38.5 | 23 | 348 | 6.19 | 3 | 3 | 4 | 1 | 1 | 1 | 3 | 3 | 4 | 1 | 1 | 1 |
| High | O75947 | ATP synthase subunit d, mitochondrial OS=Homo sapiens OX=9606 GN=ATP5PD PE=1 SV=3 | 23 | 6 | 6 | 18.5 | 43 | 161 | 5.3 | 1 | 1 | 1 | 2 | 3 | 3 | 1 | 1 | 1 | 2 | 3 | 3 |
| High | Q86V15 | Zinc finger protein castor homolog 1 OS=Homo sapiens OX=9606 GN=CASZ1 PE=1 SV=4 | 23 | 12 | 12 | 189.9 | 10 | 1759 | 7.03 | 7 | 9 | 7 | | | | 7 | 9 | 7 | | | |
| High | P28799 | Progranulin OS=Homo sapiens OX=9606 GN=GRN PE=1 SV=2 | 23 | 5 | 5 | 63.5 | 16 | 593 | 6.83 | 2 | 2 | | 1 | 2 | 1 | 2 | 2 | | 1 | 2 | 1 |
| High | Q8NI27 | THO complex subunit 2 OS=Homo sapiens OX=9606 GN=THOC2 PE=1 SV=2 | 23 | 10 | 10 | 182.7 | 7 | 1593 | 8.44 | 2 | | 1 | 1 | 3 | 3 | 2 | | 1 | 1 | 3 | 3 |
| High | Q14966 | Zinc finger protein 638 OS=Homo sapiens OX=9606 GN=ZNF638 PE=1 SV=2 | 23 | 10 | 10 | 220.5 | 7 | 1978 | 6.38 | 2 | 3 | 2 | 2 | 1 | 2 | 2 | 3 | 2 | 2 | 1 | 2 |
| High | Q9Y3E7 | Charged multivesicular body protein 3 OS=Homo sapiens OX=9606 GN=CHMP3 PE=1 SV=3 | 23 | 6 | 6 | 25.1 | 34 | 222 | 5.12 | 2 | 3 | 2 | | | | 2 | 3 | 2 | | | |
| High | P40938 | Replication factor C subunit 3 OS=Homo sapiens OX=9606 GN=RFC3 PE=1 SV=2 | 23 | 5 | 5 | 40.5 | 22 | 356 | 8.34 | 4 | 4 | 4 | | | 1 | 4 | 4 | 3 | | | 1 |
| High | Q9HB71 | Calcyclin-binding protein OS=Homo sapiens OX=9606 GN=CACYBP PE=1 SV=2 | 23 | 4 | 4 | 26.2 | 22 | 228 | 8.25 | 1 | 1 | 2 | 3 | 2 | 3 | 1 | 1 | 2 | 3 | 2 | 3 |
| High | P50452 | Serpin B8 OS=Homo sapiens OX=9606 GN=SERPINB8 PE=1 SV=2 | 23 | 6 | 4 | 42.7 | 15 | 374 | 5.57 | 2 | 2 | 2 | 1 | 5 | 4 | 2 | 2 | 2 | 1 | 4 | 4 |
| High | Q9Y2Z0 | Protein SGT1 homolog OS=Homo sapiens OX=9606 GN=SUGT1 PE=1 SV=3 | 23 | 8 | 8 | 41 | 27 | 365 | 5.16 | 1 | | 3 | 2 | 3 | 2 | 1 | | 3 | 2 | 3 | 2 |
| High | O95232 | Luc7-like protein 3 OS=Homo sapiens OX=9606 GN=LUC7L3 PE=1 SV=2 | 23 | 5 | 5 | 51.4 | 13 | 432 | 9.79 | | 1 | 1 | 5 | 5 | 4 | | 1 | 1 | 5 | 4 | 4 |
| High | Q9NX63 | MICOS complex subunit MIC19 OS=Homo sapiens OX=9606 GN=CHCHD3 PE=1 SV=1 | 23 | 5 | 5 | 26.1 | 25 | 227 | 8.28 | 1 | 1 | 1 | 2 | 3 | 3 | 1 | 1 | 1 | 2 | 3 | 3 |
| High | P04439 | HLA class I histocompatibility antigen, A alpha chain OS=Homo sapiens OX=9606 GN=HLA-A PE=1 SV=2 | 23 | 5 | 4 | 40.8 | 22 | 365 | 6 | 3 | 3 | 1 | 2 | 3 | 2 | 2 | 2 | 1 | 2 | 2 | 2 |
| High | P55084 | Trifunctional enzyme subunit beta, mitochondrial OS=Homo sapiens OX=9606 GN=HADHB PE=1 SV=3 | 23 | 8 | 8 | 51.3 | 23 | 474 | 9.41 | 2 | 3 | 2 | 3 | 6 | 3 | 2 | 3 | 2 | 3 | 6 | 3 |
| High | P41212 | Transcription factor ETV6 OS=Homo sapiens OX=9606 GN=ETV6 PE=1 SV=1 | 23 | 6 | 6 | 53 | 11 | 452 | 7.43 | 1 | 1 | 1 | 2 | 2 | 2 | 1 | 1 | 1 | 2 | 2 | 2 |
| High | P24666 | Low molecular weight phosphotyrosine protein phosphatase OS=Homo sapiens OX=9606 GN=ACP1 PE=1 SV=3 | 23 | 5 | 5 | 18 | 47 | 158 | 6.74 | 2 | 2 | 3 | 1 | 2 | 1 | 2 | 2 | 3 | 1 | 2 | 1 |
| High | O43290 | U4/U6.U5 tri-snRNP-associated protein 1 OS=Homo sapiens OX=9606 GN=SART1 PE=1 SV=1 | 23 | 9 | 9 | 90.2 | 14 | 800 | 6.13 | 2 | 1 | | 4 | 2 | 3 | 2 | 1 | | 4 | 2 | 3 |
| High | O43813 | Glutathione S-transferase LANCL1 OS=Homo sapiens OX=9606 GN=LANCL1 PE=1 SV=1 | 23 | 6 | 6 | 45.3 | 16 | 399 | 7.75 | 2 | | 3 | 1 | 3 | 3 | 2 | | 2 | 1 | 3 | 2 |
| High | O75153 | Clustered mitochondria protein homolog OS=Homo sapiens OX=9606 GN=CLUH PE=1 SV=2 | 23 | 9 | 9 | 146.6 | 8 | 1309 | 6.13 | 2 | 2 | 3 | | 5 | | 2 | 2 | 3 | | 5 | |
| High | O00170 | AH receptor-interacting protein OS=Homo sapiens OX=9606 GN=AIP PE=1 SV=3 | 23 | 6 | 6 | 37.6 | 24 | 330 | 6.42 | 1 | 1 | 1 | 2 | 3 | 2 | 1 | 1 | 1 | 2 | 3 | 2 |
| High | Q9Y2R4 | Probable ATP-dependent RNA helicase DDX52 OS=Homo sapiens OX=9606 GN=DDX52 PE=1 SV=4 | 23 | 6 | 5 | 67.4 | 12 | 599 | 9.67 | 1 | 1 | 1 | 2 | 2 | 2 | 1 | 1 | 1 | 2 | 2 | 2 |
| High | P50750 | Cyclin-dependent kinase 9 OS=Homo sapiens OX=9606 GN=CDK9 PE=1 SV=3 | 23 | 5 | 4 | 42.8 | 17 | 372 | 8.79 | 2 | 4 | 3 | 3 | 3 | 2 | 2 | 3 | 2 | 2 | 2 | 2 |
| High | P18887 | DNA repair protein XRCC1 OS=Homo sapiens OX=9606 GN=XRCC1 PE=1 SV=3 | 23 | 5 | 5 | 69.5 | 8 | 633 | 6.28 | 2 | 2 | 1 | 3 | 3 | 3 | 2 | 2 | 1 | 3 | 3 | 3 |
| High | Q8IZ83 | Aldehyde dehydrogenase family 16 member A1 OS=Homo sapiens OX=9606 GN=ALDH16A1 PE=1 SV=2 | 22 | 6 | 6 | 85.1 | 8 | 802 | 6.79 | | 1 | 3 | 1 | 3 | 4 | | 1 | 3 | 1 | 3 | 4 |
| High | O95747 | Serine/threonine-protein kinase OSR1 OS=Homo sapiens OX=9606 GN=OXSR1 PE=1 SV=1 | 22 | 6 | 2 | 58 | 11 | 527 | 6.43 | 2 | 3 | 2 | | 2 | 1 | 2 | 3 | 2 | | 2 | 1 |
| High | O43865 | S-adenosylhomocysteine hydrolase-like protein 1 OS=Homo sapiens OX=9606 GN=AHCYL1 PE=1 SV=2 | 22 | 6 | 5 | 58.9 | 11 | 530 | 6.89 | 1 | 1 | 1 | 1 | 1 | 1 | 1 | 1 | 1 | 1 | 1 | 1 |
| High | Q01581 | Hydroxymethylglutaryl-CoA synthase, cytoplasmic OS=Homo sapiens OX=9606 GN=HMGCS1 PE=1 SV=2 | 22 | 6 | 5 | 57.3 | 13 | 520 | 5.41 | 1 | 2 | 3 | 3 | | 4 | 1 | 2 | 3 | 3 | | 4 |
| High | P61421 | V-type proton ATPase subunit d 1 OS=Homo sapiens OX=9606 GN=ATP6V0D1 PE=1 SV=1 | 22 | 7 | 7 | 40.3 | 28 | 351 | 5 | 4 | 3 | 2 | | | | 4 | 3 | 2 | | | |
| High | P09972 | Fructose-bisphosphate aldolase C OS=Homo sapiens OX=9606 GN=ALDOC PE=1 SV=2 | 22 | 3 | 2 | 39.4 | 9 | 364 | 6.87 | 2 | 3 | 3 | 1 | 3 | 2 | 2 | 3 | 3 | 1 | 3 | 2 |
| High | O14929 | Histone acetyltransferase type B catalytic subunit OS=Homo sapiens OX=9606 GN=HAT1 PE=1 SV=2 | 22 | 4 | 4 | 49.5 | 14 | 419 | 5.69 | 2 | 2 | 1 | 3 | 2 | 3 | 2 | 2 | 1 | 3 | 2 | 3 |
| High | Q86U86 | Protein polybromo-1 OS=Homo sapiens OX=9606 GN=PBRM1 PE=1 SV=1 | 22 | 8 | 8 | 192.8 | 6 | 1689 | 6.89 | | | | 3 | 5 | 4 | | | | 3 | 5 | 4 |
| High | P05386 | 60S acidic ribosomal protein P1 OS=Homo sapiens OX=9606 GN=RPLP1 PE=1 SV=1 | 22 | 3 | 2 | 11.5 | 67 | 114 | 4.32 | 3 | 3 | 4 | | | | 1 | 1 | 2 | | | |
| High | Q9BY32 | Inosine triphosphate pyrophosphatase OS=Homo sapiens OX=9606 GN=ITPA PE=1 SV=2 | 22 | 5 | 5 | 21.4 | 45 | 194 | 5.66 | 4 | 4 | 4 | 1 | 1 | 1 | 4 | 4 | 3 | 1 | 1 | 1 |
| High | Q53GQ0 | Very-long-chain 3-oxoacyl-CoA reductase OS=Homo sapiens OX=9606 GN=HSD17B12 PE=1 SV=2 | 22 | 8 | 8 | 34.3 | 30 | 312 | 9.32 | 5 | 5 | 3 | | | | 5 | 5 | 3 | | | |
| High | Q8N6T3 | ADP-ribosylation factor GTPase-activating protein 1 OS=Homo sapiens OX=9606 GN=ARFGAP1 PE=1 SV=2 | 22 | 5 | 5 | 44.6 | 16 | 406 | 5.66 | 1 | 1 | 2 | 3 | 2 | 3 | 1 | 1 | 2 | 3 | 2 | 3 |
| High | P35244 | Replication protein A 14 kDa subunit OS=Homo sapiens OX=9606 GN=RPA3 PE=1 SV=1 | 22 | 4 | 4 | 13.6 | 51 | 121 | 5.08 | 4 | 5 | 7 | 1 | 1 | 1 | 3 | 4 | 4 | 1 | 1 | 1 |
| High | Q96EE3 | Nucleoporin SEH1 OS=Homo sapiens OX=9606 GN=SEH1L PE=1 SV=3 | 22 | 6 | 6 | 39.6 | 23 | 360 | 8.09 | 1 | 2 | 2 | 2 | 1 | 2 | 1 | 2 | 2 | 2 | 1 | 2 |
| High | Q12873 | Chromodomain-helicase-DNA-binding protein 3 OS=Homo sapiens OX=9606 GN=CHD3 PE=1 SV=3 | 22 | 5 | 1 | 226.5 | 4 | 2000 | 7.3 | 1 | 2 | 4 | 1 | 3 | 1 | 1 | 2 | 4 | 1 | 3 | 1 |
| High | Q9BYG3 | MKI67 FHA domain-interacting nucleolar phosphoprotein OS=Homo sapiens OX=9606 GN=NIFK PE=1 SV=1 | 22 | 5 | 5 | 34.2 | 23 | 293 | 9.88 | 1 | 1 | 1 | 2 | 3 | 2 | 1 | 1 | 1 | 2 | 3 | 2 |
| High | Q04323 | UBX domain-containing protein 1 OS=Homo sapiens OX=9606 GN=UBXN1 PE=1 SV=2 | 22 | 4 | 4 | 33.3 | 23 | 297 | 5.25 | 3 | 1 | 2 | 1 | 3 | 2 | 3 | 1 | 2 | 1 | 3 | 2 |
| High | O76003 | Glutaredoxin-3 OS=Homo sapiens OX=9606 GN=GLRX3 PE=1 SV=2 | 22 | 4 | 4 | 37.4 | 13 | 335 | 5.39 | 1 | 4 | 2 | 2 | 2 | 2 | 1 | 3 | 2 | 2 | 2 | 2 |
| High | P02686 | Myelin basic protein OS=Homo sapiens OX=9606 GN=MBP PE=1 SV=3 | 22 | 3 | 3 | 33.1 | 11 | 304 | 9.79 | | | | 6 | 8 | 8 | | | | 3 | 3 | 3 |
| High | P17931 | Galectin-3 OS=Homo sapiens OX=9606 GN=LGALS3 PE=1 SV=5 | 22 | 5 | 5 | 26.1 | 22 | 250 | 8.56 | 4 | 3 | 4 | | | | 4 | 3 | 4 | | | |
| High | P43897 | Elongation factor Ts, mitochondrial OS=Homo sapiens OX=9606 GN=TSFM PE=1 SV=2 | 22 | 6 | 6 | 35.4 | 24 | 325 | 8.38 | 2 | 1 | 3 | 2 | | 2 | 2 | 1 | 3 | 2 | | 2 |
| High | Q9BTT6 | Leucine-rich repeat-containing protein 1 OS=Homo sapiens OX=9606 GN=LRRC1 PE=1 SV=1 | 22 | 4 | 2 | 59.2 | 11 | 524 | 5.02 | 4 | 4 | 4 | | 1 | | 4 | 4 | 4 | | 1 | |
| High | Q96RS6 | NudC domain-containing protein 1 OS=Homo sapiens OX=9606 GN=NUDCD1 PE=1 SV=2 | 22 | 8 | 8 | 66.7 | 16 | 583 | 5.11 | 1 | 1 | | 2 | 2 | 1 | 1 | 1 | | 2 | 2 | 1 |
| High | Q99798 | Aconitate hydratase, mitochondrial OS=Homo sapiens OX=9606 GN=ACO2 PE=1 SV=2 | 22 | 7 | 7 | 85.4 | 13 | 780 | 7.61 | 2 | 2 | 2 | 3 | 2 | 2 | 2 | 2 | 2 | 3 | 2 | 2 |
| High | Q8N1N4 | Keratin, type II cytoskeletal 78 OS=Homo sapiens OX=9606 GN=KRT78 PE=1 SV=2 | 22 | 4 | 1 | 56.8 | 7 | 520 | 6.02 | 3 | 3 | 5 | 1 | 1 | 1 | 2 | 2 | 4 | 1 | 1 | 1 |
| High | Q86UP2 | Kinectin OS=Homo sapiens OX=9606 GN=KTN1 PE=1 SV=1 | 22 | 7 | 7 | 156.2 | 6 | 1357 | 5.64 | 2 | 1 | 2 | 1 | 2 | 1 | 2 | 1 | 2 | 1 | 2 | 1 |
| High | Q9UJZ1 | Stomatin-like protein 2, mitochondrial OS=Homo sapiens OX=9606 GN=STOML2 PE=1 SV=1 | 22 | 7 | 7 | 38.5 | 29 | 356 | 7.39 | 2 | 1 | 4 | | | | 2 | 1 | 4 | | | |
| High | Q06203 | Amidophosphoribosyltransferase OS=Homo sapiens OX=9606 GN=PPAT PE=1 SV=1 | 22 | 6 | 6 | 57.4 | 13 | 517 | 6.76 | 1 | 2 | 4 | 2 | 2 | 1 | 1 | 2 | 4 | 2 | 2 | 1 |
| High | Q8IWZ3 | Ankyrin repeat and KH domain-containing protein 1 OS=Homo sapiens OX=9606 GN=ANKHD1 PE=1 SV=1 | 21 | 7 | 1 | 269.3 | 3 | 2542 | 5.73 | | | 1 | 2 | 2 | 3 | | | 1 | 2 | 2 | 3 |
| High | P11172 | Uridine 5'-monophosphate synthase OS=Homo sapiens OX=9606 GN=UMPS PE=1 SV=1 | 21 | 6 | 6 | 52.2 | 10 | 480 | 7.24 | 1 | | 3 | | 2 | | 1 | | 3 | | 2 | |
| High | Q96T51 | RUN and FYVE domain-containing protein 1 OS=Homo sapiens OX=9606 GN=RUFY1 PE=1 SV=2 | 21 | 7 | 7 | 79.8 | 10 | 708 | 5.74 | | 2 | 2 | 1 | 2 | | | 2 | 2 | 1 | 2 | |
| High | P09382 | Galectin-1 OS=Homo sapiens OX=9606 GN=LGALS1 PE=1 SV=2 | 21 | 4 | 4 | 14.7 | 28 | 135 | 5.5 | | 1 | 2 | 2 | 3 | 4 | | 1 | 2 | 2 | 3 | 4 |
| High | O00483 | Cytochrome c oxidase subunit NDUFA4 OS=Homo sapiens OX=9606 GN=NDUFA4 PE=1 SV=1 | 21 | 3 | 3 | 9.4 | 32 | 81 | 9.38 | 2 | 2 | 2 | 3 | 3 | 3 | 2 | 2 | 2 | 3 | 3 | 3 |
| High | Q15050 | Ribosome biogenesis regulatory protein homolog OS=Homo sapiens OX=9606 GN=RRS1 PE=1 SV=2 | 21 | 6 | 6 | 41.2 | 20 | 365 | 10.7 | 1 | 1 | 1 | 1 | 2 | 1 | 1 | 1 | 1 | 1 | 2 | 1 |
| High | O60763 | General vesicular transport factor p115 OS=Homo sapiens OX=9606 GN=USO1 PE=1 SV=2 | 21 | 8 | 8 | 107.8 | 10 | 962 | 4.91 | 2 | 1 | 3 | 1 | | 1 | 2 | 1 | 3 | 1 | | 1 |
| High | P20592 | Interferon-induced GTP-binding protein Mx2 OS=Homo sapiens OX=9606 GN=MX2 PE=1 SV=1 | 21 | 10 | 10 | 82 | 15 | 715 | 8.76 | | | | 6 | 8 | 6 | | | | 6 | 8 | 6 |
| High | P54198 | Protein HIRA OS=Homo sapiens OX=9606 GN=HIRA PE=1 SV=2 | 21 | 8 | 8 | 111.8 | 10 | 1017 | 8.07 | 4 | 6 | 3 | | 1 | 1 | 4 | 6 | 3 | | 1 | 1 |
| High | P82673 | 28S ribosomal protein S35, mitochondrial OS=Homo sapiens OX=9606 GN=MRPS35 PE=1 SV=1 | 21 | 6 | 6 | 36.8 | 27 | 323 | 8.24 | 1 | 1 | 3 | 1 | 3 | 2 | 1 | 1 | 3 | 1 | 3 | 2 |
| High | Q15599 | Na(+)/H(+) exchange regulatory cofactor NHE-RF2 OS=Homo sapiens OX=9606 GN=SLC9A3R2 PE=1 SV=2 | 21 | 6 | 6 | 37.4 | 23 | 337 | 7.93 | | 2 | 2 | 1 | 3 | 1 | | 2 | 2 | 1 | 3 | 1 |
| High | Q96A33 | PAT complex subunit CCDC47 OS=Homo sapiens OX=9606 GN=CCDC47 PE=1 SV=1 | 21 | 5 | 5 | 55.8 | 14 | 483 | 4.87 | 3 | 5 | 2 | | | | 3 | 3 | 2 | | | |
| High | Q9Y221 | 60S ribosome subunit biogenesis protein NIP7 homolog OS=Homo sapiens OX=9606 GN=NIP7 PE=1 SV=1 | 21 | 6 | 6 | 20.4 | 38 | 180 | 8.51 | 1 | 1 | 1 | 4 | 3 | 2 | 1 | 1 | 1 | 4 | 3 | 2 |
| High | Q00534 | Cyclin-dependent kinase 6 OS=Homo sapiens OX=9606 GN=CDK6 PE=1 SV=1 | 21 | 4 | 3 | 36.9 | 11 | 326 | 6.46 | 2 | 3 | 3 | 3 | 2 | 2 | 2 | 2 | 2 | 2 | 1 | 2 |
| High | O14908 | PDZ domain-containing protein GIPC1 OS=Homo sapiens OX=9606 GN=GIPC1 PE=1 SV=2 | 21 | 7 | 7 | 36 | 26 | 333 | 6.28 | | 1 | 3 | | 1 | 1 | | 1 | 3 | | 1 | 1 |
| High | P25788 | Proteasome subunit alpha type-3 OS=Homo sapiens OX=9606 GN=PSMA3 PE=1 SV=2 | 21 | 4 | 4 | 28.4 | 16 | 255 | 5.33 | | | 2 | 4 | 3 | 3 | | | 2 | 4 | 3 | 3 |
| High | P19623 | Spermidine synthase OS=Homo sapiens OX=9606 GN=SRM PE=1 SV=1 | 21 | 8 | 8 | 33.8 | 38 | 302 | 5.49 | 1 | 1 | 2 | 1 | | 2 | 1 | 1 | 2 | 1 | | 2 |
| High | P51153 | Ras-related protein Rab-13 OS=Homo sapiens OX=9606 GN=RAB13 PE=1 SV=1 | 21 | 4 | 2 | 22.8 | 21 | 203 | 9.19 | 2 | 3 | 2 | 2 | 2 | 2 | 2 | 2 | 2 | 2 | 2 | 2 |
| High | O43776 | Asparagine--tRNA ligase, cytoplasmic OS=Homo sapiens OX=9606 GN=NARS1 PE=1 SV=1 | 21 | 8 | 8 | 62.9 | 19 | 548 | 6.25 | | 2 | 4 | 1 | 1 | 1 | | 2 | 4 | 1 | 1 | 1 |
| High | Q9UJA5 | tRNA (adenine(58)-N(1))-methyltransferase non-catalytic subunit TRM6 OS=Homo sapiens OX=9606 GN=TRMT6 PE=1 SV=1 | 21 | 6 | 6 | 55.8 | 15 | 497 | 7.55 | 2 | 2 | 1 | 2 | 1 | 3 | 2 | 2 | 1 | 2 | 1 | 3 |
| High | Q96QR8 | Transcriptional activator protein Pur-beta OS=Homo sapiens OX=9606 GN=PURB PE=1 SV=3 | 21 | 5 | 5 | 33.2 | 27 | 312 | 5.43 | 1 | 1 | 1 | 1 | 2 | 1 | 1 | 1 | 1 | 1 | 2 | 1 |
| High | P04637 | Cellular tumor antigen p53 OS=Homo sapiens OX=9606 GN=TP53 PE=1 SV=4 | 21 | 7 | 7 | 43.6 | 27 | 393 | 6.79 | 1 | | 1 | 2 | 3 | 2 | 1 | | 1 | 2 | 3 | 2 |
| High | Q9UKF6 | Cleavage and polyadenylation specificity factor subunit 3 OS=Homo sapiens OX=9606 GN=CPSF3 PE=1 SV=1 | 21 | 5 | 5 | 77.4 | 9 | 684 | 5.6 | 2 | 1 | 1 | 2 | 3 | 2 | 2 | 1 | 1 | 2 | 3 | 2 |
| High | P13861 | cAMP-dependent protein kinase type II-alpha regulatory subunit OS=Homo sapiens OX=9606 GN=PRKAR2A PE=1 SV=2 | 21 | 5 | 5 | 45.5 | 15 | 404 | 5.07 | 1 | 1 | 2 | 1 | 2 | 2 | 1 | 1 | 2 | 1 | 2 | 2 |
| High | P17029 | Zinc finger protein with KRAB and SCAN domains 1 OS=Homo sapiens OX=9606 GN=ZKSCAN1 PE=1 SV=3 | 21 | 7 | 7 | 63.6 | 17 | 563 | 7.05 | 4 | 5 | 4 | | | | 4 | 5 | 4 | | | |
| High | P48735 | Isocitrate dehydrogenase [NADP], mitochondrial OS=Homo sapiens OX=9606 GN=IDH2 PE=1 SV=2 | 21 | 7 | 7 | 50.9 | 19 | 452 | 8.69 | 2 | 2 | 5 | 1 | 1 | 1 | 2 | 2 | 5 | 1 | 1 | 1 |
| High | P78362 | SRSF protein kinase 2 OS=Homo sapiens OX=9606 GN=SRPK2 PE=1 SV=3 | 21 | 8 | 6 | 77.5 | 16 | 688 | 4.97 | 1 | 1 | 1 | 4 | 3 | 3 | 1 | 1 | 1 | 4 | 3 | 3 |
| High | P46782 | 40S ribosomal protein S5 OS=Homo sapiens OX=9606 GN=RPS5 PE=1 SV=4 | 21 | 4 | 4 | 22.9 | 23 | 204 | 9.72 | 1 | 2 | 2 | 1 | 1 | 1 | 1 | 2 | 2 | 1 | 1 | 1 |
| High | Q5H9R7 | Serine/threonine-protein phosphatase 6 regulatory subunit 3 OS=Homo sapiens OX=9606 GN=PPP6R3 PE=1 SV=2 | 21 | 7 | 7 | 97.6 | 10 | 873 | 4.6 | 2 | 2 | 1 | 1 | 2 | 2 | 2 | 2 | 1 | 1 | 2 | 2 |

| | | | | | | | | | | | | | | | | | | | | | |
|---|---|---|---|---|---|---|---|---|---|---|---|---|---|---|---|---|---|---|---|---|---|
| High | P61586 | Transforming protein RhoA OS=Homo sapiens OX=9606 GN=RHOA PE=1 SV=1 | 21 | 4 | 3 | 21.8 | 21 | 193 | 6.1 | 3 | 3 | 2 | 2 | 2 | 2 | 2 | 2 | 1 | 2 | 2 | 2 |
| High | P31146 | Coronin-1A OS=Homo sapiens OX=9606 GN=CORO1A PE=1 SV=4 | 21 | 5 | 4 | 51 | 11 | 461 | 6.68 | 2 | 2 | 2 | 3 | 4 | 3 | 2 | 2 | 2 | 3 | 4 | 3 |
| High | P35613 | Basigin OS=Homo sapiens OX=9606 GN=BSG PE=1 SV=2 | 21 | 5 | 5 | 42.2 | 18 | 385 | 5.66 | 3 | 3 | 4 | | 1 | | 3 | 3 | 4 | | 1 | |
| High | Q7L014 | Probable ATP-dependent RNA helicase DDX46 OS=Homo sapiens OX=9606 GN=DDX46 PE=1 SV=2 | 21 | 10 | 10 | 117.3 | 10 | 1031 | 9.29 | | | | 2 | 5 | 4 | | | | 2 | 5 | 4 |
| High | P40616 | ADP-ribosylation factor-like protein 1 OS=Homo sapiens OX=9606 GN=ARL1 PE=1 SV=1 | 21 | 6 | 6 | 20.4 | 53 | 181 | 5.72 | 5 | 2 | 5 | 1 | 1 | 1 | 5 | 2 | 5 | 1 | 1 | 1 |
| High | P47914 | 60S ribosomal protein L29 OS=Homo sapiens OX=9606 GN=RPL29 PE=1 SV=2 | 21 | 2 | 2 | 17.7 | 14 | 159 | 11.66 | 1 | 1 | 2 | 2 | 3 | 3 | 1 | 1 | 1 | 2 | 2 | 2 |
| High | Q14165 | Malectin OS=Homo sapiens OX=9606 GN=MLEC PE=1 SV=1 | 20 | 6 | 6 | 32.2 | 24 | 292 | 5.41 | | | 3 | | | 1 | | | 3 | | | 1 |
| High | Q9P2K3 | REST corepressor 3 OS=Homo sapiens OX=9606 GN=RCOR3 PE=1 SV=2 | 20 | 5 | 1 | 55.5 | 11 | 495 | 8.27 | 1 | | 1 | 1 | 2 | 1 | 1 | | 1 | 1 | 2 | 1 |
| High | P36507 | Dual specificity mitogen-activated protein kinase kinase 2 OS=Homo sapiens OX=9606 GN=MAP2K2 PE=1 SV=1 | 20 | 5 | 3 | 44.4 | 15 | 400 | 6.55 | 2 | 2 | 2 | 1 | 2 | 1 | 2 | 2 | 2 | 1 | 2 | 1 |
| High | P61513 | 60S ribosomal protein L37a OS=Homo sapiens OX=9606 GN=RPL37A PE=1 SV=2 | 20 | 3 | 3 | 10.3 | 41 | 92 | 10.43 | 1 | 1 | 2 | 2 | 3 | 2 | 1 | 1 | 2 | 2 | 2 | 2 |
| High | Q9BRJ6 | Uncharacterized protein C7orf50 OS=Homo sapiens OX=9606 GN=C7orf50 PE=1 SV=1 | 20 | 5 | 5 | 22.1 | 42 | 194 | 9.64 | | | 1 | 5 | 4 | 3 | | | 1 | 4 | 4 | 3 |
| High | O95486 | Protein transport protein Sec24A OS=Homo sapiens OX=9606 GN=SEC24A PE=1 SV=2 | 20 | 7 | 5 | 119.7 | 6 | 1093 | 7.66 | | | | 2 | 3 | 2 | | | | 2 | 3 | 2 |
| High | Q9NWH9 | SAFB-like transcription modulator OS=Homo sapiens OX=9606 GN=SLTM PE=1 SV=2 | 20 | 6 | 6 | 117.1 | 8 | 1034 | 7.87 | 2 | 2 | 3 | 1 | 3 | 1 | 2 | 2 | 3 | 1 | 3 | 1 |
| High | Q9H4I2 | Zinc fingers and homeoboxes protein 3 OS=Homo sapiens OX=9606 GN=ZHX3 PE=1 SV=3 | 20 | 7 | 6 | 104.6 | 12 | 956 | 6.07 | | | | 3 | | 2 | | | | 3 | | 2 |
| High | Q9UKL0 | REST corepressor 1 OS=Homo sapiens OX=9606 GN=RCOR1 PE=1 SV=2 | 20 | 7 | 3 | 53.3 | 18 | 485 | 7.03 | 1 | | 1 | 1 | 1 | 1 | 1 | | 1 | 1 | 1 | 1 |
| High | Q13576 | Ras GTPase-activating-like protein IQGAP2 OS=Homo sapiens OX=9606 GN=IQGAP2 PE=1 SV=4 | 20 | 8 | 6 | 180.5 | 6 | 1575 | 5.64 | 1 | 2 | 2 | 1 | 1 | 1 | 1 | 1 | 2 | 1 | 1 | 1 |
| High | P41743 | Protein kinase C iota type OS=Homo sapiens OX=9606 GN=PRKCI PE=1 SV=2 | 20 | 6 | 6 | 68.2 | 13 | 596 | 5.85 | 2 | 1 | 2 | 2 | 1 | 2 | 2 | 1 | 2 | 2 | 1 | 2 |
| High | P55036 | 26S proteasome non-ATPase regulatory subunit 4 OS=Homo sapiens OX=9606 GN=PSMD4 PE=1 SV=1 | 20 | 7 | 7 | 40.7 | 31 | 377 | 4.79 | 1 | | | 4 | 4 | 4 | 1 | | | 4 | 4 | 4 |
| High | O00233 | 26S proteasome non-ATPase regulatory subunit 9 OS=Homo sapiens OX=9606 GN=PSMD9 PE=1 SV=3 | 20 | 6 | 6 | 24.7 | 27 | 223 | 6.95 | 2 | 1 | 1 | 2 | 1 | 2 | 2 | 1 | 1 | 2 | 1 | 2 |
| High | Q15020 | Squamous cell carcinoma antigen recognized by T-cells 3 OS=Homo sapiens OX=9606 GN=SART3 PE=1 SV=1 | 20 | 5 | 5 | 109.9 | 6 | 963 | 5.57 | | | 1 | 3 | 1 | 2 | | | 1 | 3 | 1 | 2 |
| High | P42285 | Exosome RNA helicase MTR4 OS=Homo sapiens OX=9606 GN=MTREX PE=1 SV=3 | 20 | 9 | 9 | 117.7 | 9 | 1042 | 6.52 | 1 | | | 3 | 2 | 2 | 1 | | | 3 | 2 | 2 |
| High | Q9BT17 | Mitochondrial ribosome-associated GTPase 1 OS=Homo sapiens OX=9606 GN=MTG1 PE=1 SV=2 | 20 | 3 | 3 | 37.2 | 11 | 334 | 9.47 | 3 | 3 | 2 | 2 | 2 | 1 | 3 | 3 | 2 | 2 | 2 | 1 |
| High | O75351 | Vacuolar protein sorting-associated protein 4B OS=Homo sapiens OX=9606 GN=VPS4B PE=1 SV=2 | 20 | 6 | 4 | 49.3 | 18 | 444 | 7.23 | 1 | 1 | 3 | 1 | 2 | 2 | 1 | 1 | 3 | 1 | 2 | 2 |
| High | P23258 | Tubulin gamma-1 chain OS=Homo sapiens OX=9606 GN=TUBG1 PE=1 SV=2 | 20 | 6 | 6 | 51.1 | 22 | 451 | 6.14 | 5 | 4 | 5 | | | | 4 | 3 | 4 | | | |
| High | P10301 | Ras-related protein R-Ras OS=Homo sapiens OX=9606 GN=RRAS PE=1 SV=1 | 20 | 4 | 2 | 23.5 | 21 | 218 | 6.93 | 3 | 3 | 3 | 1 | 1 | 1 | 3 | 3 | 2 | 1 | 1 | 1 |
| High | P61604 | 10 kDa heat shock protein, mitochondrial OS=Homo sapiens OX=9606 GN=HSPE1 PE=1 SV=2 | 20 | 4 | 4 | 10.9 | 38 | 102 | 8.92 | 3 | 2 | 4 | 1 | 2 | 3 | 3 | 2 | 4 | 1 | 2 | 3 |
| High | P13807 | Glycogen [starch] synthase, muscle OS=Homo sapiens OX=9606 GN=GYS1 PE=1 SV=2 | 20 | 5 | 5 | 83.7 | 8 | 737 | 6.18 | 1 | | 1 | 2 | 3 | 2 | 1 | | 1 | 2 | 3 | 2 |
| High | O94808 | Glutamine--fructose-6-phosphate aminotransferase [isomerizing] 2 OS=Homo sapiens OX=9606 GN=GFPT2 PE=1 SV=3 | 20 | 4 | 1 | 76.9 | 6 | 682 | 7.37 | 3 | 3 | 3 | 2 | 1 | 2 | 3 | 3 | 3 | 2 | 1 | 2 |
| High | P43307 | Translocon-associated protein subunit alpha OS=Homo sapiens OX=9606 GN=SSR1 PE=1 SV=3 | 20 | 4 | 4 | 32.2 | 21 | 286 | 4.49 | 2 | 2 | 4 | 2 | | 2 | 2 | 2 | 4 | 2 | | 2 |
| High | P61221 | ATP-binding cassette sub-family E member 1 OS=Homo sapiens OX=9606 GN=ABCE1 PE=1 SV=1 | 20 | 9 | 9 | 67.3 | 15 | 599 | 8.34 | | | | | 3 | 1 | | | | | 3 | 1 |
| High | P61626 | Lysozyme C OS=Homo sapiens OX=9606 GN=LYZ PE=1 SV=1 | 20 | 3 | 3 | 16.5 | 18 | 148 | 9.16 | 3 | 1 | 2 | 2 | 3 | 3 | 3 | 1 | 2 | 2 | 3 | 2 |
| High | Q9BW19 | Kinesin-like protein KIFC1 OS=Homo sapiens OX=9606 GN=KIFC1 PE=1 SV=2 | 20 | 6 | 6 | 73.7 | 11 | 673 | 8.98 | 1 | | | 3 | 2 | 3 | 1 | | | 3 | 2 | 3 |
| High | Q96SI9 | Spermatid perinuclear RNA-binding protein OS=Homo sapiens OX=9606 GN=STRBP PE=1 SV=1 | 20 | 3 | 1 | 73.6 | 4 | 672 | 8.72 | 2 | 2 | 1 | 2 | 2 | 2 | 2 | 2 | 1 | 2 | 2 | 2 |
| High | P31949 | Protein S100-A11 OS=Homo sapiens OX=9606 GN=S100A11 PE=1 SV=2 | 20 | 4 | 4 | 11.7 | 43 | 105 | 7.12 | 3 | 3 | 3 | 2 | 1 | 2 | 2 | 2 | 2 | 2 | 1 | 2 |
| High | P43686 | 26S proteasome regulatory subunit 6B OS=Homo sapiens OX=9606 GN=PSMC4 PE=1 SV=2 | 20 | 9 | 8 | 47.3 | 24 | 418 | 5.21 | 1 | | | 2 | 6 | 3 | 1 | | | 2 | 6 | 3 |
| High | Q9BQG0 | Myb-binding protein 1A OS=Homo sapiens OX=9606 GN=MYBBP1A PE=1 SV=2 | 20 | 8 | 8 | 148.8 | 7 | 1328 | 9.28 | | | | 1 | | 2 | | | | 1 | | 2 |
| High | O95163 | Elongator complex protein 1 OS=Homo sapiens OX=9606 GN=ELP1 PE=1 SV=3 | 20 | 6 | 6 | 150.2 | 6 | 1332 | 5.94 | 2 | 2 | 1 | 1 | 2 | 2 | 2 | 2 | 1 | 1 | 2 | 2 |
| High | Q9HAV7 | GrpE protein homolog 1, mitochondrial OS=Homo sapiens OX=9606 GN=GRPEL1 PE=1 SV=2 | 20 | 4 | 4 | 24.3 | 23 | 217 | 8.12 | 2 | 3 | 2 | 1 | 1 | 1 | 2 | 3 | 2 | 1 | 1 | 1 |
| High | P52294 | Importin subunit alpha-5 OS=Homo sapiens OX=9606 GN=KPNA1 PE=1 SV=3 | 20 | 5 | 2 | 60.2 | 13 | 538 | 5.01 | 2 | 3 | 1 | 2 | 3 | 1 | 2 | 3 | 1 | 2 | 3 | 1 |
| High | P35606 | Coatomer subunit beta' OS=Homo sapiens OX=9606 GN=COPB2 PE=1 SV=2 | 20 | 6 | 6 | 102.4 | 9 | 906 | 5.27 | 3 | 5 | 3 | | 1 | 1 | 3 | 4 | 3 | | 1 | 1 |
| High | Q14157 | Ubiquitin-associated protein 2-like OS=Homo sapiens OX=9606 GN=UBAP2L PE=1 SV=2 | 20 | 7 | 7 | 114.5 | 10 | 1087 | 7.11 | 4 | 5 | 4 | | | | 4 | 5 | 4 | | | |
| High | Q9NUL3 | Double-stranded RNA-binding protein Staufen homolog 2 OS=Homo sapiens OX=9606 GN=STAU2 PE=1 SV=2 | 20 | 7 | 5 | 62.6 | 15 | 570 | 9.61 | | 1 | 2 | 3 | 2 | 1 | | 1 | 2 | 3 | 2 | 1 |
| High | Q8IXI2 | Mitochondrial Rho GTPase 1 OS=Homo sapiens OX=9606 GN=RHOT1 PE=1 SV=2 | 20 | 5 | 4 | 70.7 | 9 | 618 | 6.27 | | 1 | 1 | 3 | 4 | 3 | | 1 | 1 | 3 | 4 | 3 |
| High | Q9Y6Y8 | SEC23-interacting protein OS=Homo sapiens OX=9606 GN=SEC23IP PE=1 SV=1 | 20 | 6 | 6 | 111 | 9 | 1000 | 5.54 | 2 | 1 | 2 | | 1 | 2 | 2 | 1 | 2 | | 1 | 2 |
| High | P82930 | 28S ribosomal protein S34, mitochondrial OS=Homo sapiens OX=9606 GN=MRPS34 PE=1 SV=2 | 20 | 5 | 5 | 25.6 | 26 | 218 | 9.98 | 1 | 1 | 2 | 1 | 2 | 1 | 1 | 1 | 2 | 1 | 2 | 1 |
| High | P61289 | Proteasome activator complex subunit 3 OS=Homo sapiens OX=9606 GN=PSME3 PE=1 SV=1 | 20 | 4 | 4 | 29.5 | 17 | 254 | 5.95 | 3 | 2 | 2 | 1 | 2 | 1 | 3 | 2 | 2 | 1 | 2 | 1 |
| High | Q5T6F2 | Ubiquitin-associated protein 2 OS=Homo sapiens OX=9606 GN=UBAP2 PE=1 SV=1 | 20 | 7 | 7 | 117 | 9 | 1119 | 7.34 | | | 1 | 1 | 3 | 1 | | | 1 | 1 | 3 | 1 |
| High | P30084 | Enoyl-CoA hydratase, mitochondrial OS=Homo sapiens OX=9606 GN=ECHS1 PE=1 SV=4 | 20 | 7 | 7 | 31.4 | 33 | 290 | 8.07 | 4 | 3 | 3 | | 2 | 1 | 4 | 3 | 3 | | 2 | 1 |
| High | Q9NYB9 | Abl interactor 2 OS=Homo sapiens OX=9606 GN=ABI2 PE=1 SV=1 | 19 | 5 | 2 | 55.6 | 13 | 513 | 6.16 | 1 | 1 | 1 | 1 | 1 | 1 | 1 | 1 | 1 | 1 | 1 | 1 |
| High | P78417 | Glutathione S-transferase omega-1 OS=Homo sapiens OX=9606 GN=GSTO1 PE=1 SV=2 | 19 | 4 | 4 | 27.5 | 16 | 241 | 6.6 | 2 | 2 | 1 | 1 | 2 | 2 | 2 | 2 | 1 | 1 | 2 | 2 |
| High | Q9BXJ9 | N-alpha-acetyltransferase 15, NatA auxiliary subunit OS=Homo sapiens OX=9606 GN=NAA15 PE=1 SV=1 | 19 | 6 | 6 | 101.2 | 8 | 866 | 7.42 | 2 | 2 | 1 | 2 | 2 | 2 | 2 | 2 | 1 | 2 | 2 | 2 |
| High | Q08495 | Dematin OS=Homo sapiens OX=9606 GN=DMTN PE=1 SV=3 | 19 | 6 | 6 | 45.5 | 18 | 405 | 8.88 | 2 | 2 | 3 | | | | 2 | 2 | 3 | | | |
| High | Q13257 | Mitotic spindle assembly checkpoint protein MAD2A OS=Homo sapiens OX=9606 GN=MAD2L1 PE=1 SV=1 | 19 | 4 | 4 | 23.5 | 17 | 205 | 5.08 | 1 | 2 | 3 | 1 | 3 | 2 | 1 | 2 | 3 | 1 | 3 | 2 |
| High | O43432 | Eukaryotic translation initiation factor 4 gamma 3 OS=Homo sapiens OX=9606 GN=EIF4G3 PE=1 SV=2 | 19 | 9 | 6 | 176.5 | 7 | 1585 | 5.38 | | 2 | 2 | 1 | 1 | 1 | | 2 | 2 | 1 | 1 | 1 |
| High | Q99459 | Cell division cycle 5-like protein OS=Homo sapiens OX=9606 GN=CDC5L PE=1 SV=2 | 19 | 8 | 8 | 92.2 | 15 | 802 | 8.18 | 3 | 2 | 3 | 1 | 2 | 1 | 3 | 2 | 3 | 1 | 2 | 1 |
| High | Q08209 | Serine/threonine-protein phosphatase 2B catalytic subunit alpha isoform OS=Homo sapiens OX=9606 GN=PPP3CA PE=1 SV=1 | 19 | 5 | 5 | 58.7 | 11 | 521 | 5.86 | 1 | 1 | 1 | 1 | 2 | 2 | 1 | 1 | 1 | 1 | 2 | 2 |
| High | Q96T58 | Msx2-interacting protein OS=Homo sapiens OX=9606 GN=SPEN PE=1 SV=1 | 19 | 3 | 3 | 402 | 1 | 3664 | 7.64 | 3 | 1 | 3 | 2 | | | 2 | 1 | 2 | 1 | | |
| High | Q7Z417 | Nuclear fragile X mental retardation-interacting protein 2 OS=Homo sapiens OX=9606 GN=NUFIP2 PE=1 SV=1 | 19 | 7 | 7 | 76.1 | 17 | 695 | 8.7 | 1 | 1 | 2 | | 3 | 2 | 1 | 1 | 2 | | 3 | 2 |
| High | P28072 | Proteasome subunit beta type-6 OS=Homo sapiens OX=9606 GN=PSMB6 PE=1 SV=4 | 19 | 4 | 4 | 25.3 | 22 | 239 | 4.92 | 2 | | | 3 | 3 | 2 | 2 | | | 3 | 3 | 2 |
| High | Q5GLZ8 | Probable E3 ubiquitin-protein ligase HERC4 OS=Homo sapiens OX=9606 GN=HERC4 PE=1 SV=1 | 19 | 5 | 5 | 118.5 | 7 | 1057 | 6.19 | 1 | 2 | 2 | | 2 | 2 | 1 | 2 | 2 | | 2 | 2 |
| High | Q9Y4K1 | Beta/gamma crystallin domain-containing protein 1 OS=Homo sapiens OX=9606 GN=CRYBG1 PE=1 SV=3 | 19 | 8 | 8 | 188.6 | 7 | 1723 | 5.86 | 5 | 7 | 3 | | | | 5 | 7 | 3 | | | |
| High | O15371 | Eukaryotic translation initiation factor 3 subunit D OS=Homo sapiens OX=9606 GN=EIF3D PE=1 SV=1 | 19 | 5 | 5 | 63.9 | 16 | 548 | 6.05 | 3 | 3 | 4 | | 1 | | 3 | 3 | 4 | | 1 | |
| High | Q5BKZ1 | DBIRD complex subunit ZNF326 OS=Homo sapiens OX=9606 GN=ZNF326 PE=1 SV=2 | 19 | 5 | 5 | 65.6 | 11 | 582 | 5.15 | 2 | 2 | 3 | 1 | 2 | | 2 | 2 | 2 | 1 | 2 | |
| High | Q7LBR1 | Charged multivesicular body protein 1b OS=Homo sapiens OX=9606 GN=CHMP1B PE=1 SV=1 | 19 | 3 | 3 | 22.1 | 12 | 199 | 8.1 | 2 | 3 | 3 | 2 | 2 | 2 | 2 | 2 | 2 | 2 | 2 | 2 |
| High | P52888 | Thimet oligopeptidase OS=Homo sapiens OX=9606 GN=THOP1 PE=1 SV=2 | 19 | 6 | 6 | 78.8 | 10 | 689 | 6.05 | 1 | 1 | | 2 | 3 | 2 | 1 | 1 | | 2 | 3 | 2 |
| High | O75475 | PC4 and SFRS1-interacting protein OS=Homo sapiens OX=9606 GN=PSIP1 PE=1 SV=1 | 19 | 9 | 9 | 60.1 | 19 | 530 | 9.13 | 4 | 2 | 5 | | 2 | | 3 | 2 | 5 | | 2 | |
| High | P00973 | 2'-5'-oligoadenylate synthase 1 OS=Homo sapiens OX=9606 GN=OAS1 PE=1 SV=4 | 19 | 5 | 5 | 46 | 13 | 400 | 8.22 | | 1 | 1 | 3 | 1 | 3 | | 1 | 1 | 3 | 1 | 3 |
| High | Q8N1W1 | Rho guanine nucleotide exchange factor 28 OS=Homo sapiens OX=9606 GN=ARHGEF28 PE=1 SV=3 | 19 | 8 | 8 | 191.8 | 6 | 1705 | 6.04 | 3 | 3 | 8 | | | | 3 | 3 | 8 | | | |
| High | Q0JRZ9 | F-BAR domain only protein 2 OS=Homo sapiens OX=9606 GN=FCHO2 PE=1 SV=2 | 19 | 6 | 6 | 88.9 | 8 | 810 | 6.86 | 1 | 2 | 1 | 1 | | 1 | 1 | 2 | 1 | 1 | | 1 |
| High | Q14789 | Golgin subfamily B member 1 OS=Homo sapiens OX=9606 GN=GOLGB1 PE=1 SV=2 | 19 | 9 | 8 | 375.8 | 3 | 3259 | 5 | | 1 | | | | 1 | | 1 | | | | 1 |
| High | Q9BZH6 | WD repeat-containing protein 11 OS=Homo sapiens OX=9606 GN=WDR11 PE=1 SV=1 | 19 | 9 | 9 | 136.6 | 10 | 1224 | 6.92 | | 1 | 1 | | 2 | | | 1 | 1 | | 2 | |
| High | O60333 | Kinesin-like protein KIF1B OS=Homo sapiens OX=9606 GN=KIF1B PE=1 SV=5 | 19 | 6 | 5 | 204.3 | 5 | 1816 | 5.6 | 4 | 4 | 4 | | 2 | | 4 | 4 | 4 | | 2 | |
| High | Q9BYJ9 | YTH domain-containing family protein 1 OS=Homo sapiens OX=9606 GN=YTHDF1 PE=1 SV=1 | 19 | 5 | 4 | 60.8 | 9 | 559 | 8.79 | 4 | 4 | 2 | 1 | 2 | | 4 | 4 | 2 | 1 | 2 | |
| High | P62834 | Ras-related protein Rap-1A OS=Homo sapiens OX=9606 GN=RAP1A PE=1 SV=1 | 19 | 4 | 2 | 21 | 28 | 184 | 6.67 | 2 | | 1 | 1 | 3 | 1 | 2 | | 1 | 1 | 3 | 1 |
| High | P61020 | Ras-related protein Rab-5B OS=Homo sapiens OX=9606 GN=RAB5B PE=1 SV=1 | 19 | 5 | 2 | 23.7 | 27 | 215 | 8.13 | 2 | 3 | 2 | 1 | 2 | 1 | 2 | 3 | 2 | 1 | 2 | 1 |
| High | O43889 | Cyclic AMP-responsive element-binding protein 3 OS=Homo sapiens OX=9606 GN=CREB3 PE=1 SV=2 | 19 | 4 | 4 | 41.4 | 14 | 371 | 4.91 | | | | 3 | 2 | 3 | | | | 3 | 2 | 3 |
| High | Q9NVE7 | 4'-phosphopantetheine phosphatase OS=Homo sapiens OX=9606 GN=PANK4 PE=1 SV=1 | 19 | 8 | 8 | 85.9 | 12 | 773 | 6.28 | 5 | 4 | 6 | | | | 5 | 4 | 6 | | | |
| High | Q8IXB1 | DnaJ homolog subfamily C member 10 OS=Homo sapiens OX=9606 GN=DNAJC10 PE=1 SV=2 | 19 | 4 | 4 | 91 | 7 | 793 | 7.18 | 1 | 2 | 1 | 2 | 1 | 3 | 1 | 2 | 1 | 2 | 1 | 3 |
| High | Q13363 | C-terminal-binding protein 1 OS=Homo sapiens OX=9606 GN=CTBP1 PE=1 SV=2 | 19 | 4 | 2 | 47.5 | 10 | 440 | 6.77 | 1 | 1 | 2 | 2 | 1 | 2 | 1 | 1 | 2 | 2 | 1 | 2 |
| High | Q6DD88 | Atlastin-3 OS=Homo sapiens OX=9606 GN=ATL3 PE=1 SV=1 | 19 | 7 | 7 | 60.5 | 21 | 541 | 5.66 | 5 | 3 | 5 | | | | 5 | 3 | 4 | | | |
| High | Q96PY5 | Formin-like protein 2 OS=Homo sapiens OX=9606 GN=FMNL2 PE=1 SV=3 | 19 | 10 | 10 | 123.2 | 10 | 1086 | 7.4 | 2 | 3 | 2 | 1 | | 1 | 2 | 3 | 2 | 1 | | 1 |
| High | P49841 | Glycogen synthase kinase-3 beta OS=Homo sapiens OX=9606 GN=GSK3B PE=1 SV=2 | 19 | 4 | 4 | 46.7 | 15 | 420 | 8.78 | 2 | 2 | 2 | 3 | 2 | 2 | 2 | 2 | 2 | 3 | 2 | 2 |
| High | P28340 | DNA polymerase delta catalytic subunit OS=Homo sapiens OX=9606 GN=POLD1 PE=1 SV=2 | 19 | 7 | 6 | 123.6 | 7 | 1107 | 7.03 | 2 | 2 | 2 | | | 1 | 2 | 2 | 2 | | | 1 |
| High | P40763 | Signal transducer and activator of transcription 3 OS=Homo sapiens OX=9606 GN=STAT3 PE=1 SV=2 | 18 | 6 | 6 | 88 | 13 | 770 | 6.3 | 3 | 2 | 3 | | 3 | | 3 | 2 | 3 | | 3 | |
| High | P12763 | SWISS-PROT:P12763 (Bos taurus) Alpha-2-HS-glycoprotein precursor | 18 | 4 | 4 | 38.4 | 13 | 359 | 5.5 | 1 | 1 | 2 | 1 | | 1 | 1 | 1 | 2 | 1 | | 1 |
| High | P52298 | Nuclear cap-binding protein subunit 2 OS=Homo sapiens OX=9606 GN=NCBP2 PE=1 SV=1 | 18 | 3 | 3 | 18 | 17 | 156 | 8.21 | 1 | 1 | 3 | 2 | 2 | 2 | 1 | 1 | 2 | 2 | 2 | 2 |
| High | P82675 | 28S ribosomal protein S5, mitochondrial OS=Homo sapiens OX=9606 GN=MRPS5 PE=1 SV=2 | 18 | 5 | 5 | 48 | 11 | 430 | 9.92 | 1 | 2 | | 3 | 1 | 2 | 1 | 2 | | 3 | 1 | 2 |
| High | P42685 | Tyrosine-protein kinase FRK OS=Homo sapiens OX=9606 GN=FRK PE=1 SV=1 | 18 | 3 | 1 | 58.2 | 5 | 505 | 6.67 | 2 | 3 | 1 | 1 | 2 | 2 | 2 | 2 | 1 | 1 | 2 | 2 |
| High | Q9Y508 | E3 ubiquitin-protein ligase RNF114 OS=Homo sapiens OX=9606 GN=RNF114 PE=1 SV=1 | 18 | 5 | 5 | 25.7 | 26 | 228 | 7.25 | | 1 | 1 | 2 | 3 | 2 | | 1 | 1 | 2 | 3 | 2 |
| High | Q9H0C8 | Integrin-linked kinase-associated serine/threonine phosphatase 2C OS=Homo sapiens OX=9606 GN=ILKAP PE=1 SV=1 | 18 | 6 | 6 | 42.9 | 15 | 392 | 7.09 | 1 | 1 | 2 | 3 | | 2 | 1 | 1 | 2 | 3 | | 2 |
| High | O60218 | Aldo-keto reductase family 1 member B10 OS=Homo sapiens OX=9606 GN=AKR1B10 PE=1 SV=2 | 18 | 5 | 5 | 36 | 16 | 316 | 7.84 | 2 | 3 | 2 | 1 | 2 | 2 | 2 | 3 | 2 | 1 | 2 | 2 |
| High | Q15369 | Elongin-C OS=Homo sapiens OX=9606 GN=ELOC PE=1 SV=1 | 18 | 3 | 3 | 12.5 | 41 | 112 | 4.78 | 2 | 2 | 1 | 1 | 2 | 1 | 2 | 2 | 1 | 1 | 2 | 1 |
| High | P40199 | Carcinoembryonic antigen-related cell adhesion molecule 6 OS=Homo sapiens OX=9606 GN=CEACAM6 PE=1 SV=4 | 18 | 5 | 4 | 37.2 | 24 | 344 | 5.82 | | | 2 | | | 1 | | | 2 | | | 1 |
| High | P51571 | Translocon-associated protein subunit delta OS=Homo sapiens OX=9606 GN=SSR4 PE=1 SV=1 | 18 | 4 | 4 | 19 | 31 | 173 | 6.15 | 1 | 3 | 4 | 1 | 1 | 1 | 1 | 3 | 3 | 1 | 1 | 1 |
| High | Q9Y295 | Developmentally-regulated GTP-binding protein 1 OS=Homo sapiens OX=9606 GN=DRG1 PE=1 SV=1 | 18 | 4 | 4 | 40.5 | 13 | 367 | 8.9 | 1 | | 2 | 1 | 4 | 1 | 1 | | 2 | 1 | 4 | 1 |
| High | Q96SZ5 | 2-aminoethanethiol dioxygenase OS=Homo sapiens OX=9606 GN=ADO PE=1 SV=2 | 18 | 5 | 5 | 29.7 | 19 | 270 | 6.04 | 1 | 1 | 2 | 2 | 2 | 2 | 1 | 1 | 2 | 2 | 2 | 2 |
| High | Q9Y570 | Protein phosphatase methylesterase 1 OS=Homo sapiens OX=9606 GN=PPME1 PE=1 SV=3 | 18 | 7 | 7 | 42.3 | 21 | 386 | 5.97 | 1 | 2 | 2 | 1 | 1 | | 1 | 2 | 2 | 1 | 1 | |
| High | Q9NWB6 | Arginine and glutamate-rich protein 1 OS=Homo sapiens OX=9606 GN=ARGLU1 PE=1 SV=1 | 18 | 3 | 3 | 33.2 | 8 | 273 | 10.35 | | | 3 | 2 | 3 | 2 | | | 3 | 2 | 3 | 2 |
| High | Q7L2J0 | 7SK snRNA methylphosphate capping enzyme OS=Homo sapiens OX=9606 GN=MEPCE PE=1 SV=1 | 18 | 6 | 6 | 74.3 | 13 | 689 | 9.57 | 1 | 1 | 2 | 1 | 2 | | 1 | 1 | 2 | 1 | 2 | |
| High | P49748 | Very long-chain specific acyl-CoA dehydrogenase, mitochondrial OS=Homo sapiens OX=9606 GN=ACADVL PE=1 SV=1 | 18 | 8 | 8 | 70.3 | 14 | 655 | 8.75 | 1 | 2 | 3 | | | 1 | 1 | 2 | 3 | | | 1 |
| High | P06868 | SWISS-PROT:P06868 (Bos taurus) Plasminogen precursor | 18 | 8 | 8 | 91.2 | 18 | 812 | 7.5 | 1 | 3 | 1 | 1 | 3 | 3 | 1 | 3 | 1 | 1 | 3 | 3 |
| High | P15559 | NAD(P)H dehydrogenase [quinone] 1 OS=Homo sapiens OX=9606 GN=NQO1 PE=1 SV=1 | 18 | 5 | 5 | 30.8 | 25 | 274 | 8.88 | 5 | 4 | 2 | 1 | | | 4 | 3 | 2 | 1 | | |
| High | Q07954 | Prolow-density lipoprotein receptor-related protein 1 OS=Homo sapiens OX=9606 GN=LRP1 PE=1 SV=2 | 18 | 7 | 7 | 504.3 | 2 | 4544 | 5.39 | 1 | 1 | 1 | 1 | 1 | | 1 | 1 | 1 | 1 | 1 | |
| High | Q92485 | Acid sphingomyelinase-like phosphodiesterase 3b OS=Homo sapiens OX=9606 GN=SMPDL3B PE=1 SV=2 | 18 | 4 | 4 | 50.8 | 12 | 455 | 5.64 | 3 | 4 | 2 | 1 | 1 | 1 | 3 | 4 | 2 | 1 | 1 | 1 |
| High | Q53EL6 | Programmed cell death protein 4 OS=Homo sapiens OX=9606 GN=PDCD4 PE=1 SV=2 | 18 | 10 | 10 | 51.7 | 31 | 469 | 5.21 | 1 | 1 | 2 | 1 | | 1 | 1 | 1 | 2 | 1 | | 1 |
| High | P51531 | Probable global transcription activator SNF2L2 OS=Homo sapiens OX=9606 GN=SMARCA2 PE=1 SV=2 | 18 | 4 | 1 | 181.2 | 3 | 1590 | 7.2 | 2 | 1 | 2 | 2 | 1 | 2 | 2 | 1 | 2 | 2 | 1 | 2 |
| High | Q969V3 | Nicalin OS=Homo sapiens OX=9606 GN=NCLN PE=1 SV=2 | 18 | 3 | 3 | 62.9 | 6 | 563 | 6.89 | 2 | 2 | 3 | | 2 | 3 | 2 | 2 | 3 | | 2 | 2 |
| High | Q6UB28 | Methionine aminopeptidase 1D, mitochondrial OS=Homo sapiens OX=9606 GN=METAP1D PE=1 SV=1 | 18 | 2 | 2 | 37.1 | 9 | 335 | 6.84 | 2 | 2 | 2 | 1 | | 2 | 2 | 2 | 2 | 1 | | 2 |
| High | Q9GZR2 | RNA exonuclease 4 OS=Homo sapiens OX=9606 GN=REXO4 PE=1 SV=2 | 18 | 5 | 5 | 46.6 | 14 | 422 | 9.77 | | | | 2 | 2 | 2 | | | | 2 | 2 | 2 |
| High | P81605 | Dermcidin OS=Homo sapiens OX=9606 GN=DCD PE=1 SV=2 | 18 | 2 | 2 | 11.3 | 20 | 110 | 6.54 | 1 | 1 | 2 | 4 | 3 | 3 | 1 | 1 | 2 | 2 | 2 | 2 |
| High | P23229 | Integrin alpha-6 OS=Homo sapiens OX=9606 GN=ITGA6 PE=1 SV=5 | 18 | 10 | 10 | 126.5 | 9 | 1130 | 6.61 | 3 | 1 | 4 | 1 | 1 | 1 | 3 | 1 | 4 | 1 | 1 | 1 |
| High | O75340 | Programmed cell death protein 6 OS=Homo sapiens OX=9606 GN=PDCD6 PE=1 SV=1 | 18 | 6 | 6 | 21.9 | 44 | 191 | 5.4 | 6 | 3 | 3 | | 1 | | 6 | 3 | 3 | | 1 | |
| High | Q969Q0 | 60S ribosomal protein L36a-like OS=Homo sapiens OX=9606 GN=RPL36AL PE=1 SV=3 | 18 | 3 | 1 | 12.5 | 24 | 106 | 10.65 | 1 | 1 | 1 | 2 | 3 | 2 | 1 | 1 | 1 | 2 | 3 | 2 |
| High | Q13418 | Integrin-linked protein kinase OS=Homo sapiens OX=9606 GN=ILK PE=1 SV=2 | 18 | 4 | 4 | 51.4 | 8 | 452 | 8.07 | 3 | 2 | 3 | | 1 | 1 | 3 | 2 | 3 | | 1 | 1 |
| High | Q9UGV2 | Protein NDRG3 OS=Homo sapiens OX=9606 GN=NDRG3 PE=1 SV=2 | 18 | 4 | 4 | 41.4 | 14 | 375 | 5.31 | 3 | 2 | 1 | 1 | 2 | 1 | 3 | 2 | 1 | 1 | 2 | 1 |
| High | Q6P1Q9 | tRNA N(3)-methylcytidine methyltransferase METTL2B OS=Homo sapiens OX=9606 GN=METTL2B PE=1 SV=3 | 18 | 5 | 5 | 43.4 | 13 | 378 | 5.86 | 1 | 1 | 1 | 3 | 1 | 2 | 1 | 1 | 1 | 3 | 1 | 2 |
| High | P61088 | Ubiquitin-conjugating enzyme E2 N OS=Homo sapiens OX=9606 GN=UBE2N PE=1 SV=1 | 18 | 6 | 6 | 17.1 | 47 | 152 | 6.57 | 3 | 3 | 2 | 1 | 3 | 2 | 3 | 3 | 2 | 1 | 3 | 2 |
| High | Q9Y676 | 28S ribosomal protein S18b, mitochondrial OS=Homo sapiens OX=9606 GN=MRPS18B PE=1 SV=1 | 18 | 3 | 3 | 29.4 | 15 | 258 | 9.38 | 1 | | 2 | 2 | 2 | 3 | 1 | | 2 | 2 | 2 | 3 |
| High | Q92526 | T-complex protein 1 subunit zeta-2 OS=Homo sapiens OX=9606 GN=CCT6B PE=1 SV=5 | 18 | 5 | 1 | 57.8 | 9 | 530 | 7.24 | 3 | 3 | 2 | 1 | 3 | 1 | 3 | 3 | 2 | 1 | 3 | 1 |
| High | P82933 | 28S ribosomal protein S9, mitochondrial OS=Homo sapiens OX=9606 GN=MRPS9 PE=1 SV=2 | 18 | 6 | 6 | 45.8 | 19 | 396 | 9.51 | 2 | | 2 | 1 | 1 | | 2 | | 2 | 1 | 1 | |
| High | Q8WVY7 | Ubiquitin-like domain-containing CTD phosphatase 1 OS=Homo sapiens OX=9606 GN=UBLCP1 PE=1 SV=2 | 18 | 6 | 6 | 36.8 | 17 | 318 | 6.46 | 1 | 1 | 2 | 2 | 6 | 2 | 1 | 1 | 1 | 2 | 6 | 2 |
| High | Q14562 | ATP-dependent RNA helicase DHX8 OS=Homo sapiens OX=9606 GN=DHX8 PE=1 SV=1 | 18 | 3 | 2 | 139.2 | 2 | 1220 | 8.32 | 1 | 1 | 1 | 3 | 3 | 3 | 1 | 1 | 1 | 2 | 2 | 2 |
| High | Q16401 | 26S proteasome non-ATPase regulatory subunit 5 OS=Homo sapiens OX=9606 GN=PSMD5 PE=1 SV=3 | 18 | 6 | 6 | 56.2 | 15 | 504 | 5.48 | 5 | 3 | 5 | | | | 5 | 3 | 5 | | | |
| High | Q6P1M3 | LLGL scribble cell polarity complex component 2 OS=Homo sapiens OX=9606 GN=LLGL2 PE=1 SV=2 | 18 | 5 | 5 | 113.4 | 6 | 1020 | 7.52 | 1 | 2 | 1 | 2 | 1 | 1 | 1 | 2 | 1 | 2 | 1 | 1 |
| High | Q8NBF2 | NHL repeat-containing protein 2 OS=Homo sapiens OX=9606 GN=NHLRC2 PE=1 SV=1 | 18 | 7 | 7 | 79.4 | 10 | 726 | 5.55 | | 2 | 1 | 3 | 1 | 1 | | 2 | 1 | 3 | 1 | 1 |
| High | O75381 | Peroxisomal membrane protein PEX14 OS=Homo sapiens OX=9606 GN=PEX14 PE=1 SV=1 | 17 | 4 | 4 | 41.2 | 16 | 377 | 4.94 | 2 | 3 | 4 | | | | 2 | 3 | 4 | | | |
| High | Q86YP4 | Transcriptional repressor p66-alpha OS=Homo sapiens OX=9606 GN=GATAD2A PE=1 SV=1 | 17 | 4 | 3 | 68 | 9 | 633 | 9.94 | 1 | 1 | 2 | | 4 | 1 | 1 | 1 | 2 | | 4 | 1 |
| High | Q9Y6N5 | Sulfide:quinone oxidoreductase, mitochondrial OS=Homo sapiens OX=9606 GN=SQOR PE=1 SV=1 | 17 | 6 | 6 | 49.9 | 18 | 450 | 9.11 | 3 | 5 | 3 | | | | 3 | 5 | 3 | | | |
| High | Q8IYQ7 | Threonine synthase-like 1 OS=Homo sapiens OX=9606 GN=THNSL1 PE=1 SV=2 | 17 | 6 | 6 | 83 | 11 | 743 | 7.12 | 1 | 1 | 1 | 2 | 2 | | 1 | 1 | 1 | 2 | 2 | |
| High | Q15262 | Receptor-type tyrosine-protein phosphatase kappa OS=Homo sapiens OX=9606 GN=PTPRK PE=1 SV=2 | 17 | 6 | 6 | 162 | 6 | 1439 | 5.9 | 1 | 1 | 1 | | 1 | 1 | 1 | 1 | 1 | | 1 | 1 |
| High | P27449 | V-type proton ATPase 16 kDa proteolipid subunit c OS=Homo sapiens OX=9606 GN=ATP6V0C PE=1 SV=1 | 17 | 1 | 1 | 15.7 | 12 | 155 | 8.44 | 1 | 1 | | | 1 | | 1 | 1 | 1 | | 1 | |
| High | Q9UJV9 | Probable ATP-dependent RNA helicase DDX41 OS=Homo sapiens OX=9606 GN=DDX41 PE=1 SV=2 | 17 | 7 | 7 | 69.8 | 11 | 622 | 6.84 | 2 | 1 | | 4 | 5 | 3 | 2 | 1 | | 4 | 5 | 3 |
| High | P62714 | Serine/threonine-protein phosphatase 2A catalytic subunit beta isoform OS=Homo sapiens OX=9606 GN=PPP2CB PE=1 SV=1 | 17 | 5 | 5 | 35.6 | 22 | 309 | 5.43 | 1 | 1 | 1 | | 2 | 2 | 1 | 1 | 1 | | 2 | 2 |
| High | P49720 | Proteasome subunit beta type-3 OS=Homo sapiens OX=9606 GN=PSMB3 PE=1 SV=2 | 17 | 5 | 5 | 22.9 | 25 | 205 | 6.55 | 2 | 1 | 1 | 3 | 4 | 2 | 2 | 1 | 1 | 3 | 4 | 2 |
| High | O75131 | Copine-3 OS=Homo sapiens OX=9606 GN=CPNE3 PE=1 SV=1 | 17 | 4 | 3 | 60.1 | 9 | 537 | 5.85 | 1 | 1 | 2 | 1 | 1 | 3 | 1 | 1 | 2 | 1 | 1 | 3 |
| High | P39656 | Dolichyl-diphosphooligosaccharide--protein glycosyltransferase 48 kDa subunit OS=Homo sapiens OX=9606 GN=DDOST PE=1 SV=4 | 17 | 7 | 7 | 50.8 | 23 | 456 | 6.55 | 4 | 2 | 4 | 1 | 2 | 1 | 4 | 2 | 4 | 1 | 2 | 1 |
| High | P82932 | 28S ribosomal protein S6, mitochondrial OS=Homo sapiens OX=9606 GN=MRPS6 PE=1 SV=3 | 17 | 4 | 4 | 14.2 | 30 | 125 | 9.26 | 1 | 1 | | 2 | 2 | 2 | 1 | 1 | | 2 | 2 | 2 |
| High | Q9BSJ8 | Extended synaptotagmin-1 OS=Homo sapiens OX=9606 GN=ESYT1 PE=1 SV=1 | 17 | 4 | 4 | 122.8 | 5 | 1104 | 5.83 | 1 | 2 | 3 | 1 | 1 | 2 | 1 | 2 | 3 | 1 | 1 | 2 |
| High | P00403 | Cytochrome c oxidase subunit 2 OS=Homo sapiens OX=9606 GN=MT-CO2 PE=1 SV=1 | 17 | 4 | 4 | 25.5 | 23 | 227 | 4.82 | 3 | 3 | 3 | 1 | 1 | | 3 | 3 | 3 | 1 | 1 | |
| High | Q9Y2F5 | Little elongation complex subunit 1 OS=Homo sapiens OX=9606 GN=ICE1 PE=1 SV=5 | 17 | 8 | 8 | 247.7 | 5 | 2266 | 5.48 | | | | 2 | 2 | 5 | | | | 2 | 2 | 5 |
| High | Q9NRF8 | CTP synthase 2 OS=Homo sapiens OX=9606 GN=CTPS2 PE=1 SV=1 | 17 | 7 | 6 | 65.6 | 15 | 586 | 6.9 | 2 | 2 | 1 | 1 | | 1 | 2 | 2 | 1 | 1 | | 1 |
| High | P53634 | Dipeptidyl peptidase 1 OS=Homo sapiens OX=9606 GN=CTSC PE=1 SV=2 | 17 | 5 | 5 | 51.8 | 16 | 463 | 6.99 | 2 | 1 | 3 | 1 | 1 | 2 | 2 | 1 | 3 | 1 | 1 | 2 |
| High | Q8NBQ5 | Estradiol 17-beta-dehydrogenase 11 OS=Homo sapiens OX=9606 GN=HSD17B11 PE=1 SV=3 | 17 | 5 | 5 | 32.9 | 19 | 300 | 9.07 | 1 | | 3 | | 1 | | 1 | | 3 | | 1 | |
| High | Q9H2D6 | TRIO and F-actin-binding protein OS=Homo sapiens OX=9606 GN=TRIOBP PE=1 SV=3 | 17 | 5 | 5 | 261.2 | 3 | 2365 | 8.48 | 3 | 3 | 4 | | 1 | | 3 | 3 | 4 | | 1 | |
| High | P84098 | 60S ribosomal protein L19 OS=Homo sapiens OX=9606 GN=RPL19 PE=1 SV=1 | 17 | 3 | 3 | 23.5 | 18 | 196 | 11.47 | 3 | 2 | 1 | 1 | 2 | 2 | 3 | 1 | 1 | 1 | 2 | 2 |
| High | P49721 | Proteasome subunit beta type-2 OS=Homo sapiens OX=9606 GN=PSMB2 PE=1 SV=1 | 17 | 3 | 3 | 22.8 | 12 | 201 | 7.02 | | | | 3 | 3 | 3 | | | | 3 | 3 | 3 |
| High | Q9BY43 | Charged multivesicular body protein 4a OS=Homo sapiens OX=9606 GN=CHMP4A PE=1 SV=3 | 17 | 4 | 4 | 25.1 | 22 | 222 | 4.7 | 1 | 1 | 1 | | | | 1 | 1 | 1 | | | |
| High | Q96PV6 | Leukocyte receptor cluster member 8 OS=Homo sapiens OX=9606 GN=LENG8 PE=1 SV=3 | 17 | 7 | 7 | 88.1 | 15 | 800 | 9.11 | 3 | 2 | 2 | | 1 | | 3 | 2 | 2 | | 1 | |
| High | Q92572 | AP-3 complex subunit sigma-1 OS=Homo sapiens OX=9606 GN=AP3S1 PE=1 SV=1 | 17 | 4 | 4 | 21.7 | 23 | 193 | 5.39 | 1 | 1 | 3 | 1 | 2 | 1 | 1 | 1 | 3 | 1 | 2 | 1 |
| High | Q9Y2Q3 | Glutathione S-transferase kappa 1 OS=Homo sapiens OX=9606 GN=GSTK1 PE=1 SV=3 | 17 | 3 | 3 | 25.5 | 18 | 226 | 8.41 | 2 | 2 | 3 | 1 | 1 | 1 | 2 | 2 | 3 | 1 | 1 | 1 |
| High | Q92785 | Zinc finger protein ubi-d4 OS=Homo sapiens OX=9606 GN=DPF2 PE=1 SV=2 | 17 | 5 | 5 | 44.1 | 17 | 391 | 6.33 | 3 | 1 | 1 | | 1 | | 3 | 1 | 1 | | 1 | |
| High | Q8NBS9 | Thioredoxin domain-containing protein 5 OS=Homo sapiens OX=9606 GN=TXNDC5 PE=1 SV=2 | 17 | 5 | 5 | 47.6 | 15 | 432 | 5.97 | 1 | 1 | 2 | 2 | | 1 | 1 | 1 | 2 | 2 | | 1 |
| High | P12036 | Neurofilament heavy polypeptide OS=Homo sapiens OX=9606 GN=NEFH PE=1 SV=4 | 17 | 4 | 1 | 112.4 | 3 | 1026 | 6.18 | 1 | 1 | 2 | 3 | 4 | 3 | 1 | 1 | 2 | 3 | 3 | 3 |
| High | P14678 | Small nuclear ribonucleoprotein-associated proteins B and B' OS=Homo sapiens OX=9606 GN=SNRPB PE=1 SV=2 | 17 | 4 | 4 | 24.6 | 15 | 240 | 11.19 | 3 | 2 | 2 | | 3 | 1 | 3 | 2 | 2 | | 2 | 1 |
| High | Q8WUJ3 | Cell migration-inducing and hyaluronan-binding protein OS=Homo sapiens OX=9606 GN=CEMIP PE=1 SV=2 | 17 | 7 | 7 | 152.9 | 7 | 1361 | 7.85 | 4 | 4 | 7 | | | | 4 | 4 | 6 | | | |
| High | Q9Y4F1 | FERM, ARHGEF and pleckstrin domain-containing protein 1 OS=Homo sapiens OX=9606 GN=FARP1 PE=1 SV=1 | 17 | 6 | 6 | 118.6 | 8 | 1045 | 8.15 | 3 | 3 | 3 | 2 | | 1 | 3 | 3 | 3 | 2 | | 1 |
| High | Q7Z5K2 | Wings apart-like protein homolog OS=Homo sapiens OX=9606 GN=WAPL PE=1 SV=1 | 17 | 7 | 7 | 132.9 | 7 | 1190 | 5.44 | 1 | 1 | | 3 | 2 | 1 | 1 | 1 | | 3 | 2 | 1 |
| High | P82663 | 28S ribosomal protein S25, mitochondrial OS=Homo sapiens OX=9606 GN=MRPS25 PE=1 SV=1 | 17 | 4 | 4 | 20.1 | 29 | 173 | 8.82 | 1 | 1 | 1 | 1 | 2 | | 1 | 1 | 1 | 1 | 2 | |
| High | Q8IXT5 | RNA-binding protein 12B OS=Homo sapiens OX=9606 GN=RBM12B PE=1 SV=2 | 17 | 6 | 6 | 118 | 6 | 1001 | 6.81 | 5 | 6 | 2 | | | | 5 | 6 | 2 | | | |
| High | Q5T1R4 | Transcription factor HIVEP3 OS=Homo sapiens OX=9606 GN=HIVEP3 PE=2 SV=1 | 17 | 6 | 3 | 259.3 | 3 | 2406 | 7.81 | | | | 4 | 4 | 5 | | | | 4 | 4 | 5 |
| High | O95671 | Probable bifunctional dTTP/UTP pyrophosphatase/methyltransferase protein OS=Homo sapiens OX=9606 GN=ASMTL PE=1 SV=3 | 17 | 5 | 5 | 68.8 | 12 | 621 | 6.07 | 1 | 1 | 2 | 1 | 1 | 1 | 1 | 1 | 2 | 1 | 1 | 1 |
| High | P61011 | Signal recognition particle 54 kDa protein OS=Homo sapiens OX=9606 GN=SRP54 PE=1 SV=1 | 17 | 5 | 5 | 55.7 | 10 | 504 | 8.75 | 1 | | 1 | 2 | 2 | 3 | 1 | | 1 | 2 | 2 | 3 |
| High | Q96FZ2 | Abasic site processing protein HMCES OS=Homo sapiens OX=9606 GN=HMCES PE=1 SV=1 | 17 | 5 | 5 | 40.5 | 15 | 354 | 8.15 | 3 | 2 | 1 | 2 | 1 | 2 | 3 | 2 | 1 | 2 | 1 | 2 |
| High | Q9P2N5 | RNA-binding protein 27 OS=Homo sapiens OX=9606 GN=RBM27 PE=1 SV=2 | 17 | 6 | 5 | 118.6 | 6 | 1060 | 9.19 | | | | 1 | 2 | 1 | | | | 1 | 2 | 1 |
| High | Q13464 | Rho-associated protein kinase 1 OS=Homo sapiens OX=9606 GN=ROCK1 PE=1 SV=1 | 17 | 4 | 1 | 158.1 | 3 | 1354 | 5.9 | | 1 | 2 | 1 | 1 | 2 | | 1 | 2 | 1 | 1 | 2 |
| High | O95571 | Persulfide dioxygenase ETHE1, mitochondrial OS=Homo sapiens OX=9606 GN=ETHE1 PE=1 SV=2 | 17 | 6 | 6 | 27.9 | 36 | 254 | 6.83 | 2 | 2 | 1 | | 1 | | 2 | 2 | 1 | | 1 | |
| High | Q8IYB5 | Stromal membrane-associated protein 1 OS=Homo sapiens OX=9606 GN=SMAP1 PE=1 SV=2 | 17 | 5 | 5 | 50.4 | 15 | 467 | 8.75 | 1 | 1 | 2 | 1 | 2 | 2 | 1 | 1 | 2 | 1 | 2 | 2 |
| High | P20839 | Inosine-5'-monophosphate dehydrogenase 1 OS=Homo sapiens OX=9606 GN=IMPDH1 PE=1 SV=2 | 17 | 4 | 3 | 55.4 | 11 | 514 | 6.9 | | 1 | | 2 | 3 | 3 | | 1 | | 2 | 3 | 3 |
| High | P02786 | Transferrin receptor protein 1 OS=Homo sapiens OX=9606 GN=TFRC PE=1 SV=2 | 17 | 8 | 8 | 84.8 | 13 | 760 | 6.61 | | 2 | 2 | 1 | | 1 | | 2 | 2 | 1 | | 1 |

| | | | | | | | | | | | | | | | | | | | | | |
|---|---|---|---|---|---|---|---|---|---|---|---|---|---|---|---|---|---|---|---|---|---|
| High | P08579 | U2 small nuclear ribonucleoprotein B'' OS=Homo sapiens OX=9606 GN=SNRPB2 PE=1 SV=1 | 17 | 3 | 1 | 25.5 | 12 | 225 | 9.72 | 2 | 1 | 1 | 1 | 3 | 2 | 2 | 1 | 1 | 1 | 2 | 2 |
| High | P46379 | Large proline-rich protein BAG6 OS=Homo sapiens OX=9606 GN=BAG6 PE=1 SV=2 | 17 | 5 | 5 | 119.3 | 7 | 1132 | 5.6 | | | | 3 | 2 | 4 | | | | 3 | 2 | 4 |
| High | Q13243 | Serine/arginine-rich splicing factor 5 OS=Homo sapiens OX=9606 GN=SRSF5 PE=1 SV=1 | 17 | 5 | 4 | 31.2 | 25 | 272 | 11.59 | 2 | 2 | 1 | 2 | | 2 | 2 | 2 | 1 | 2 | | 2 |
| High | O00330 | Pyruvate dehydrogenase protein X component, mitochondrial OS=Homo sapiens OX=9606 GN=PDHX PE=1 SV=3 | 16 | 5 | 5 | 54.1 | 12 | 501 | 8.66 | | 1 | | 1 | 1 | | | 1 | | 1 | 1 | |
| High | Q14651 | Plastin-1 OS=Homo sapiens OX=9606 GN=PLS1 PE=1 SV=2 | 16 | 5 | 1 | 70.2 | 11 | 629 | 5.41 | 2 | 2 | 2 | 1 | 2 | 2 | 2 | 2 | 2 | 1 | 2 | 2 |
| High | Q96ST3 | Paired amphipathic helix protein Sin3a OS=Homo sapiens OX=9606 GN=SIN3A PE=1 SV=2 | 16 | 8 | 8 | 145.1 | 6 | 1273 | 7.25 | | | | 1 | 3 | 1 | | | | 1 | 3 | 1 |
| High | Q9NXH8 | Torsin-4A OS=Homo sapiens OX=9606 GN=TOR4A PE=1 SV=2 | 16 | 6 | 6 | 46.9 | 15 | 423 | 9.94 | 1 | | 4 | | 2 | 1 | 1 | | 4 | | 2 | 1 |
| High | Q9BPX5 | Actin-related protein 2/3 complex subunit 5-like protein OS=Homo sapiens OX=9606 GN=ARPC5L PE=1 SV=1 | 16 | 4 | 3 | 16.9 | 34 | 153 | 6.6 | 2 | 2 | 1 | 1 | 2 | 1 | 2 | 2 | 1 | 1 | 1 | 1 |
| High | Q5JPE7 | Nodal modulator 2 OS=Homo sapiens OX=9606 GN=NOMO2 PE=1 SV=1 | 16 | 4 | 4 | 139.4 | 3 | 1267 | 5.76 | 3 | 3 | 2 | | | | 3 | 3 | 2 | | | |
| High | Q06330 | Recombining binding protein suppressor of hairless OS=Homo sapiens OX=9606 GN=RBPJ PE=1 SV=3 | 16 | 5 | 5 | 55.6 | 12 | 500 | 7.18 | | | | 2 | 3 | 3 | | | | 2 | 3 | 3 |
| High | Q96GM8 | Target of EGR1 protein 1 OS=Homo sapiens OX=9606 GN=TOE1 PE=1 SV=1 | 16 | 4 | 4 | 56.5 | 13 | 510 | 7.18 | 2 | 1 | 2 | 1 | 1 | 1 | 2 | 1 | 2 | 1 | 1 | 1 |
| High | Q92922 | SWI/SNF complex subunit SMARCC1 OS=Homo sapiens OX=9606 GN=SMARCC1 PE=1 SV=3 | 16 | 5 | 3 | 122.8 | 7 | 1105 | 5.76 | | | 1 | | 1 | 1 | | | 1 | | 1 | 1 |
| High | P09525 | Annexin A4 OS=Homo sapiens OX=9606 GN=ANXA4 PE=1 SV=4 | 16 | 5 | 5 | 35.9 | 16 | 319 | 6.13 | 2 | 2 | 1 | 1 | 1 | 2 | 2 | 2 | 1 | 1 | 1 | 2 |
| High | P10620 | Microsomal glutathione S-transferase 1 OS=Homo sapiens OX=9606 GN=MGST1 PE=1 SV=1 | 16 | 3 | 3 | 17.6 | 26 | 155 | 9.39 | 3 | 3 | 3 | | 1 | | 3 | 3 | 3 | | 1 | |
| High | Q96EP5 | DAZ-associated protein 1 OS=Homo sapiens OX=9606 GN=DAZAP1 PE=1 SV=1 | 16 | 3 | 3 | 43.4 | 10 | 407 | 8.56 | 2 | 2 | 2 | 1 | 2 | 1 | 2 | 2 | 2 | 1 | 2 | 1 |
| High | P09234 | U1 small nuclear ribonucleoprotein C OS=Homo sapiens OX=9606 GN=SNRPC PE=1 SV=1 | 16 | 2 | 2 | 17.4 | 19 | 159 | 9.67 | 1 | 2 | 2 | 2 | 2 | 2 | 1 | 2 | 2 | 2 | 2 | 2 |
| High | A8CG34 | Nuclear envelope pore membrane protein POM 121C OS=Homo sapiens OX=9606 GN=POM121C PE=1 SV=3 | 16 | 5 | 5 | 125 | 6 | 1229 | 10.37 | | | 1 | | 1 | 1 | | | 1 | | 1 | 1 |
| High | Q96P47 | Arf-GAP with GTPase, ANK repeat and PH domain-containing protein 3 OS=Homo sapiens OX=9606 GN=AGAP3 PE=1 SV=2 | 16 | 3 | 3 | 95 | 5 | 875 | 7.97 | 2 | 3 | 3 | 1 | 1 | 1 | 2 | 3 | 2 | 1 | 1 | 1 |
| High | O00182 | Galectin-9 OS=Homo sapiens OX=9606 GN=LGALS9 PE=1 SV=2 | 16 | 5 | 2 | 39.5 | 15 | 355 | 9.17 | 1 | | 1 | 2 | 3 | 2 | 1 | | 1 | 2 | 3 | 2 |
| High | Q9GZS3 | WD repeat-containing protein 61 OS=Homo sapiens OX=9606 GN=WDR61 PE=1 SV=1 | 16 | 4 | 4 | 33.6 | 14 | 305 | 5.47 | 1 | 2 | 3 | | 2 | 1 | 1 | 2 | 3 | | 2 | 1 |
| High | Q96A65 | Exocyst complex component 4 OS=Homo sapiens OX=9606 GN=EXOC4 PE=1 SV=1 | 16 | 8 | 8 | 110.4 | 11 | 974 | 6.49 | 5 | 4 | 5 | | | | 5 | 4 | 5 | | | |
| High | O60925 | Prefoldin subunit 1 OS=Homo sapiens OX=9606 GN=PFDN1 PE=1 SV=2 | 16 | 3 | 3 | 14.2 | 21 | 122 | 6.81 | | 1 | 1 | 3 | 1 | 3 | | 1 | 1 | 3 | 1 | 3 |
| High | Q6P996 | Pyridoxal-dependent decarboxylase domain-containing protein 1 OS=Homo sapiens OX=9606 GN=PDXDC1 PE=1 SV=2 | 16 | 7 | 7 | 86.7 | 12 | 788 | 5.38 | 2 | 1 | 3 | | 1 | | 2 | 1 | 3 | | 1 | |
| High | Q96DN6 | Methyl-CpG-binding domain protein 6 OS=Homo sapiens OX=9606 GN=MBD6 PE=1 SV=2 | 16 | 6 | 6 | 101.1 | 10 | 1003 | 9.63 | | | | 3 | 2 | 2 | | | | 3 | 2 | 2 |
| High | Q13404 | Ubiquitin-conjugating enzyme E2 variant 1 OS=Homo sapiens OX=9606 GN=UBE2V1 PE=1 SV=2 | 16 | 3 | 3 | 16.5 | 18 | 147 | 7.93 | 1 | 1 | | 2 | 3 | 2 | 1 | 1 | | 2 | 3 | 2 |
| High | P12277 | Creatine kinase B-type OS=Homo sapiens OX=9606 GN=CKB PE=1 SV=1 | 16 | 4 | 4 | 42.6 | 20 | 381 | 5.59 | 2 | 3 | 1 | | | | 2 | 3 | 1 | | | |
| High | P27361 | Mitogen-activated protein kinase 3 OS=Homo sapiens OX=9606 GN=MAPK3 PE=1 SV=4 | 16 | 4 | 1 | 43.1 | 15 | 379 | 6.74 | | 1 | 2 | 1 | 1 | 2 | | 1 | 2 | 1 | 1 | 2 |
| High | O60762 | Dolichol-phosphate mannosyltransferase subunit 1 OS=Homo sapiens OX=9606 GN=DPM1 PE=1 SV=1 | 16 | 6 | 6 | 29.6 | 28 | 260 | 9.57 | 3 | 2 | 5 | | | | 3 | 2 | 5 | | | |
| High | Q96SQ9 | Cytochrome P450 2S1 OS=Homo sapiens OX=9606 GN=CYP2S1 PE=1 SV=2 | 16 | 4 | 4 | 55.8 | 8 | 504 | 8.62 | 4 | 4 | 3 | | | | 4 | 4 | 3 | | | |
| High | Q99828 | Calcium and integrin-binding protein 1 OS=Homo sapiens OX=9606 GN=CIB1 PE=1 SV=4 | 16 | 3 | 3 | 21.7 | 20 | 191 | 4.78 | 2 | 2 | 2 | | 1 | | 2 | 2 | 2 | | 1 | |
| High | Q9NZJ9 | Diphosphoinositol polyphosphate phosphohydrolase 2 OS=Homo sapiens OX=9606 GN=NUDT4 PE=1 SV=2 | 16 | 4 | 4 | 20.3 | 31 | 180 | 6.35 | 2 | 5 | 2 | | 1 | | 2 | 4 | 2 | | 1 | |
| High | Q9UN37 | Vacuolar protein sorting-associated protein 4A OS=Homo sapiens OX=9606 GN=VPS4A PE=1 SV=1 | 16 | 4 | 2 | 48.9 | 11 | 437 | 7.8 | | 1 | | 1 | 2 | 2 | | 1 | | 1 | 2 | 2 |
| High | P42338 | Phosphatidylinositol 4,5-bisphosphate 3-kinase catalytic subunit beta isoform OS=Homo sapiens OX=9606 GN=PIK3CB PE=1 SV=1 | 16 | 5 | 5 | 122.7 | 5 | 1070 | 7.09 | 4 | 3 | 3 | | 1 | 1 | 4 | 3 | 3 | | 1 | 1 |
| High | P52701 | DNA mismatch repair protein Msh6 OS=Homo sapiens OX=9606 GN=MSH6 PE=1 SV=2 | 16 | 8 | 8 | 152.7 | 7 | 1360 | 6.9 | | 1 | | 1 | | 1 | | 1 | | 1 | | 1 |
| High | P31939 | Bifunctional purine biosynthesis protein ATIC OS=Homo sapiens OX=9606 GN=ATIC PE=1 SV=3 | 16 | 8 | 8 | 64.6 | 18 | 592 | 6.71 | 2 | 2 | 4 | | | | 2 | 2 | 4 | | | |
| High | O95373 | Importin-7 OS=Homo sapiens OX=9606 GN=IPO7 PE=1 SV=1 | 16 | 10 | 10 | 119.4 | 13 | 1038 | 4.82 | 5 | 2 | 3 | | 1 | | 5 | 2 | 3 | | 1 | |
| High | P35250 | Replication factor C subunit 2 OS=Homo sapiens OX=9606 GN=RFC2 PE=1 SV=3 | 16 | 7 | 7 | 39.1 | 23 | 354 | 6.44 | 1 | 1 | | 1 | 3 | 1 | 1 | 1 | | 1 | 3 | 1 |
| High | Q9UI30 | Multifunctional methyltransferase subunit TRM112-like protein OS=Homo sapiens OX=9606 GN=TRMT112 PE=1 SV=1 | 16 | 2 | 2 | 14.2 | 22 | 125 | 5.26 | 2 | 1 | 1 | 2 | 2 | 2 | 2 | 1 | 1 | 2 | 2 | 2 |
| High | O43148 | mRNA cap guanine-N7 methyltransferase OS=Homo sapiens OX=9606 GN=RNMT PE=1 SV=1 | 16 | 4 | 4 | 54.8 | 10 | 476 | 6.61 | | 1 | 3 | 2 | 1 | 1 | | 1 | 3 | 2 | 1 | 1 |
| High | Q14919 | Dr1-associated corepressor OS=Homo sapiens OX=9606 GN=DRAP1 PE=1 SV=3 | 16 | 3 | 3 | 22.3 | 21 | 205 | 5.17 | 1 | 2 | 2 | 1 | 1 | 2 | 1 | 2 | 2 | 1 | 1 | 2 |
| High | P83881 | 60S ribosomal protein L36a OS=Homo sapiens OX=9606 GN=RPL36A PE=1 SV=2 | 16 | 3 | 1 | 12.4 | 24 | 106 | 10.58 | 1 | 1 | 1 | 2 | 3 | 2 | 1 | 1 | 1 | 2 | 3 | 2 |
| High | O15446 | DNA-directed RNA polymerase I subunit RPA34 OS=Homo sapiens OX=9606 GN=POLR1G PE=1 SV=1 | 16 | 5 | 5 | 55 | 15 | 510 | 8.51 | | | 1 | 1 | 2 | 1 | | | 1 | 1 | 2 | 1 |
| High | Q14451 | Growth factor receptor-bound protein 7 OS=Homo sapiens OX=9606 GN=GRB7 PE=1 SV=2 | 16 | 7 | 7 | 59.6 | 18 | 532 | 8.5 | 4 | 2 | 1 | 1 | | | 4 | 2 | 1 | 1 | | |
| High | P08174 | Complement decay-accelerating factor OS=Homo sapiens OX=9606 GN=CD55 PE=1 SV=4 | 16 | 4 | 4 | 41.4 | 9 | 381 | 7.59 | 2 | 2 | 1 | | | | 2 | 2 | 1 | | | |
| High | Q8TBC3 | SH3KBP1-binding protein 1 OS=Homo sapiens OX=9606 GN=SHKBP1 PE=1 SV=2 | 16 | 4 | 3 | 76.3 | 8 | 707 | 8.28 | | | | 3 | 2 | 3 | | | | 3 | 2 | 3 |
| High | Q9NQC3 | Reticulon-4 OS=Homo sapiens OX=9606 GN=RTN4 PE=1 SV=2 | 16 | 5 | 5 | 129.9 | 8 | 1192 | 4.5 | 1 | 2 | 2 | | 1 | | 1 | 2 | 1 | | 1 | |
| High | P50238 | Cysteine-rich protein 1 OS=Homo sapiens OX=9606 GN=CRIP1 PE=1 SV=3 | 16 | 3 | 3 | 8.5 | 56 | 77 | 8.75 | 3 | 2 | 2 | | 2 | 1 | 1 | 2 | 2 | | 2 | 1 |
| High | O43447 | Peptidyl-prolyl cis-trans isomerase H OS=Homo sapiens OX=9606 GN=PPIH PE=1 SV=1 | 16 | 4 | 4 | 19.2 | 23 | 177 | 8.07 | 2 | 1 | 2 | 1 | 1 | 1 | 2 | 1 | 2 | 1 | 1 | 1 |
| High | Q9NRX3 | NADH dehydrogenase [ubiquinone] 1 alpha subcomplex subunit 4-like 2 OS=Homo sapiens OX=9606 GN=NDUFA4L2 PE=3 SV=1 | 16 | 3 | 3 | 10 | 29 | 87 | 9.92 | 1 | 3 | 3 | | | | 1 | 3 | 3 | | | |
| High | Q9P0J7 | E3 ubiquitin-protein ligase KCMF1 OS=Homo sapiens OX=9606 GN=KCMF1 PE=1 SV=2 | 16 | 4 | 4 | 41.9 | 19 | 381 | 5.66 | 1 | 1 | 1 | 1 | 1 | 1 | 1 | 1 | 1 | 1 | 1 | 1 |
| High | Q9C0J8 | pre-mRNA 3' end processing protein WDR33 OS=Homo sapiens OX=9606 GN=WDR33 PE=1 SV=2 | 15 | 5 | 5 | 145.8 | 6 | 1336 | 9.17 | | | | 2 | 3 | 2 | | | | 2 | 3 | 2 |
| High | Q86SX6 | Glutaredoxin-related protein 5, mitochondrial OS=Homo sapiens OX=9606 GN=GLRX5 PE=1 SV=2 | 15 | 3 | 3 | 16.6 | 28 | 157 | 6.79 | 2 | 1 | 1 | 1 | 2 | 1 | 2 | 1 | 1 | 1 | 2 | 1 |
| High | P08621 | U1 small nuclear ribonucleoprotein 70 kDa OS=Homo sapiens OX=9606 GN=SNRNP70 PE=1 SV=2 | 15 | 4 | 4 | 51.5 | 12 | 437 | 9.94 | 1 | 1 | 2 | 1 | 1 | 1 | 1 | 1 | 2 | 1 | 1 | 1 |
| High | O43760 | Synaptogyrin-2 OS=Homo sapiens OX=9606 GN=SYNGR2 PE=1 SV=1 | 15 | 4 | 4 | 24.8 | 17 | 224 | 4.94 | 2 | 3 | 1 | | 1 | | 2 | 3 | 1 | | 1 | |
| High | Q9NTK5 | Obg-like ATPase 1 OS=Homo sapiens OX=9606 GN=OLA1 PE=1 SV=2 | 15 | 5 | 5 | 44.7 | 11 | 396 | 7.81 | | | 1 | 2 | 1 | 1 | | | 1 | 2 | 1 | 1 |
| High | Q9UBC2 | Epidermal growth factor receptor substrate 15-like 1 OS=Homo sapiens OX=9606 GN=EPS15L1 PE=1 SV=1 | 15 | 5 | 5 | 94.2 | 5 | 864 | 5.11 | | | 3 | | 1 | 1 | | | 3 | | 1 | 1 |
| High | Q99615 | DnaJ homolog subfamily C member 7 OS=Homo sapiens OX=9606 GN=DNAJC7 PE=1 SV=2 | 15 | 6 | 6 | 56.4 | 15 | 494 | 6.96 | 1 | 1 | 1 | 2 | 1 | 3 | 1 | 1 | 1 | 2 | 1 | 3 |
| High | P25787 | Proteasome subunit alpha type-2 OS=Homo sapiens OX=9606 GN=PSMA2 PE=1 SV=2 | 15 | 4 | 4 | 25.9 | 23 | 234 | 7.43 | 1 | 1 | 1 | 2 | 3 | 2 | 1 | 1 | 1 | 2 | 2 | 2 |
| High | Q13769 | THO complex subunit 5 homolog OS=Homo sapiens OX=9606 GN=THOC5 PE=1 SV=2 | 15 | 5 | 5 | 78.5 | 9 | 683 | 6.87 | 1 | 1 | | 1 | | 1 | 1 | 1 | | 1 | | 1 |
| High | Q7Z434 | Mitochondrial antiviral-signaling protein OS=Homo sapiens OX=9606 GN=MAVS PE=1 SV=2 | 15 | 4 | 4 | 56.5 | 12 | 540 | 5.52 | 1 | 2 | | 1 | 2 | 2 | 1 | 2 | | 1 | 2 | 2 |
| High | Q13098 | COP9 signalosome complex subunit 1 OS=Homo sapiens OX=9606 GN=GPS1 PE=1 SV=4 | 15 | 6 | 6 | 55.5 | 16 | 491 | 6.74 | 1 | 1 | | 1 | 1 | 1 | 1 | 1 | | 1 | 1 | 1 |
| High | O94916 | Nuclear factor of activated T-cells 5 OS=Homo sapiens OX=9606 GN=NFAT5 PE=1 SV=1 | 15 | 7 | 7 | 165.7 | 7 | 1531 | 5.24 | 2 | 2 | 8 | | | | 2 | 2 | 7 | | | |
| High | Q8TF72 | Protein Shroom3 OS=Homo sapiens OX=9606 GN=SHROOM3 PE=1 SV=2 | 15 | 8 | 8 | 216.7 | 7 | 1996 | 7.8 | | | 1 | | 1 | | | | 1 | | 1 | |
| High | O15551 | Claudin-3 OS=Homo sapiens OX=9606 GN=CLDN3 PE=1 SV=1 | 15 | 3 | 3 | 23.3 | 15 | 220 | 8.05 | | 1 | 2 | 2 | 2 | 2 | | 1 | 2 | 2 | 2 | 2 |
| High | Q9UKY1 | Zinc fingers and homeoboxes protein 1 OS=Homo sapiens OX=9606 GN=ZHX1 PE=1 SV=1 | 15 | 6 | 6 | 98 | 15 | 873 | 6.05 | 1 | | 1 | | | | 1 | | 1 | | | |
| High | Q8IY37 | Probable ATP-dependent RNA helicase DHX37 OS=Homo sapiens OX=9606 GN=DHX37 PE=1 SV=1 | 15 | 8 | 8 | 129.5 | 8 | 1157 | 8.1 | 2 | 1 | 1 | 1 | 2 | 2 | 2 | 1 | 1 | 1 | 2 | 2 |
| High | Q96FQ6 | Protein S100-A16 OS=Homo sapiens OX=9606 GN=S100A16 PE=1 SV=1 | 15 | 3 | 3 | 11.8 | 34 | 103 | 6.79 | 2 | 2 | 3 | 1 | 1 | 1 | 2 | 2 | 1 | 1 | 1 | 1 |
| High | P47985 | Cytochrome b-c1 complex subunit Rieske, mitochondrial OS=Homo sapiens OX=9606 GN=UQCRFS1 PE=1 SV=2 | 15 | 5 | 1 | 29.6 | 20 | 274 | 8.32 | 3 | 1 | 4 | 2 | 1 | 1 | 3 | 1 | 4 | 2 | 1 | 1 |
| High | Q9ULW0 | Targeting protein for Xklp2 OS=Homo sapiens OX=9606 GN=TPX2 PE=1 SV=2 | 15 | 7 | 7 | 85.6 | 15 | 747 | 9.23 | 1 | 1 | 1 | | | | 1 | 1 | 1 | | | |
| High | Q9UKB1 | F-box/WD repeat-containing protein 11 OS=Homo sapiens OX=9606 GN=FBXW11 PE=1 SV=1 | 15 | 6 | 6 | 62.1 | 11 | 542 | 7.12 | | | | 3 | 5 | 3 | | | | 3 | 5 | 3 |
| High | P53677 | AP-3 complex subunit mu-2 OS=Homo sapiens OX=9606 GN=AP3M2 PE=1 SV=1 | 15 | 3 | 1 | 46.9 | 9 | 418 | 7.56 | 2 | 2 | 2 | 1 | 2 | 1 | 2 | 2 | 2 | 1 | 2 | 1 |
| High | P83916 | Chromobox protein homolog 1 OS=Homo sapiens OX=9606 GN=CBX1 PE=1 SV=1 | 15 | 2 | 1 | 21.4 | 15 | 185 | 4.93 | 2 | 2 | 1 | | 1 | | 1 | 1 | 1 | | 1 | |
| High | P48634 | Protein PRRC2A OS=Homo sapiens OX=9606 GN=PRRC2A PE=1 SV=3 | 15 | 6 | 6 | 228.7 | 6 | 2157 | 9.45 | | | 1 | 1 | 1 | 2 | | | 1 | 1 | 1 | 2 |
| High | Q9UNS2 | COP9 signalosome complex subunit 3 OS=Homo sapiens OX=9606 GN=COPS3 PE=1 SV=3 | 15 | 5 | 5 | 47.8 | 17 | 423 | 6.65 | 1 | 1 | 2 | 1 | 2 | 1 | 1 | 1 | 2 | 1 | 2 | 1 |
| High | Q6UN15 | Pre-mRNA 3'-end-processing factor FIP1 OS=Homo sapiens OX=9606 GN=FIP1L1 PE=1 SV=1 | 15 | 5 | 5 | 66.5 | 13 | 594 | 5.59 | 1 | 2 | | 1 | 2 | 3 | 1 | 2 | | 1 | 2 | 3 |
| High | O60216 | Double-strand-break repair protein rad21 homolog OS=Homo sapiens OX=9606 GN=RAD21 PE=1 SV=2 | 15 | 5 | 5 | 71.6 | 9 | 631 | 4.65 | 2 | 2 | 1 | 2 | 1 | | 2 | 2 | 1 | 2 | 1 | |
| High | Q8N0W3 | L-fucose kinase OS=Homo sapiens OX=9606 GN=FCSK PE=1 SV=2 | 15 | 6 | 6 | 117.5 | 8 | 1084 | 6.25 | 5 | 6 | 3 | | | | 5 | 6 | 3 | | | |
| High | P28070 | Proteasome subunit beta type-4 OS=Homo sapiens OX=9606 GN=PSMB4 PE=1 SV=4 | 15 | 3 | 3 | 29.2 | 13 | 264 | 5.97 | 1 | 1 | 1 | 3 | 3 | 3 | 1 | 1 | 1 | 3 | 3 | 3 |
| High | Q07021 | Complement component 1 Q subcomponent-binding protein, mitochondrial OS=Homo sapiens OX=9606 GN=C1QBP PE=1 SV=1 | 15 | 4 | 4 | 31.3 | 26 | 282 | 4.84 | 1 | 2 | 3 | | 2 | | 1 | 2 | 3 | | 2 | |
| High | Q9NZZ3 | Charged multivesicular body protein 5 OS=Homo sapiens OX=9606 GN=CHMP5 PE=1 SV=1 | 15 | 3 | 3 | 24.6 | 20 | 219 | 4.83 | 2 | 2 | 1 | | 1 | 1 | 2 | 2 | 1 | | 1 | 1 |
| High | Q9NZN5 | Rho guanine nucleotide exchange factor 12 OS=Homo sapiens OX=9606 GN=ARHGEF12 PE=1 SV=1 | 15 | 8 | 8 | 173.1 | 7 | 1544 | 5.74 | | | | | | | | | | | | |
| High | P37198 | Nuclear pore glycoprotein p62 OS=Homo sapiens OX=9606 GN=NUP62 PE=1 SV=3 | 15 | 6 | 6 | 53.2 | 19 | 522 | 5.31 | 1 | 1 | | 1 | 1 | 1 | 1 | 1 | | 1 | 1 | 1 |
| High | Q15942 | Zyxin OS=Homo sapiens OX=9606 GN=ZYX PE=1 SV=1 | 15 | 6 | 6 | 61.2 | 16 | 572 | 6.67 | 3 | 3 | 2 | 3 | | | 3 | 3 | 2 | 3 | | |
| High | Q96PU5 | E3 ubiquitin-protein ligase NEDD4-like OS=Homo sapiens OX=9606 GN=NEDD4L PE=1 SV=2 | 15 | 4 | 3 | 111.9 | 4 | 975 | 5.82 | 1 | 1 | 1 | 3 | 1 | 2 | 1 | 1 | 1 | 3 | 1 | 2 |
| High | P23458 | Tyrosine-protein kinase JAK1 OS=Homo sapiens OX=9606 GN=JAK1 PE=1 SV=2 | 15 | 5 | 5 | 133.2 | 6 | 1154 | 7.55 | 3 | 3 | 2 | | | | 3 | 3 | 2 | | | |
| High | P68400 | Casein kinase II subunit alpha OS=Homo sapiens OX=9606 GN=CSNK2A1 PE=1 SV=1 | 15 | 5 | 5 | 45.1 | 21 | 391 | 7.74 | 3 | 3 | 3 | 1 | | 1 | 3 | 3 | 3 | 1 | | 1 |
| High | O95239 | Chromosome-associated kinesin KIF4A OS=Homo sapiens OX=9606 GN=KIF4A PE=1 SV=3 | 15 | 7 | 7 | 139.8 | 8 | 1232 | 6.27 | 1 | | 1 | 1 | 2 | 1 | 1 | | 1 | 1 | 2 | 1 |
| High | P25490 | Transcriptional repressor protein YY1 OS=Homo sapiens OX=9606 GN=YY1 PE=1 SV=2 | 15 | 3 | 3 | 44.7 | 7 | 414 | 6.25 | | 1 | 2 | 2 | 1 | 2 | | 1 | 2 | 2 | 1 | 2 |
| High | Q9Y399 | 28S ribosomal protein S2, mitochondrial OS=Homo sapiens OX=9606 GN=MRPS2 PE=1 SV=1 | 15 | 5 | 5 | 33.2 | 14 | 296 | 9.26 | | | 1 | 1 | 3 | 1 | | | 1 | 1 | 3 | 1 |
| High | Q8WVB6 | Chromosome transmission fidelity protein 18 homolog OS=Homo sapiens OX=9606 GN=CHTF18 PE=1 SV=1 | 15 | 3 | 3 | 107.3 | 4 | 975 | 7.21 | 1 | 2 | 1 | 1 | 1 | 1 | 1 | 2 | 1 | 1 | 1 | 1 |
| High | Q9Y618 | Nuclear receptor corepressor 2 OS=Homo sapiens OX=9606 GN=NCOR2 PE=1 SV=3 | 15 | 8 | 8 | 273.5 | 4 | 2514 | 7.59 | 1 | | 1 | 1 | 2 | 1 | 1 | | 1 | 1 | 2 | 1 |
| High | Q14558 | Phosphoribosyl pyrophosphate synthase-associated protein 1 OS=Homo sapiens OX=9606 GN=PRPSAP1 PE=1 SV=2 | 15 | 4 | 3 | 39.4 | 15 | 356 | 7.2 | 3 | 3 | 2 | | 1 | 1 | 3 | 3 | 2 | | 1 | 1 |
| High | P40222 | Alpha-taxilin OS=Homo sapiens OX=9606 GN=TXLNA PE=1 SV=3 | 15 | 7 | 7 | 61.9 | 19 | 546 | 6.52 | | 2 | 2 | 1 | 3 | 2 | | 2 | 2 | 1 | 3 | 2 |
| High | Q9BRU9 | rRNA-processing protein UTP23 homolog OS=Homo sapiens OX=9606 GN=UTP23 PE=1 SV=2 | 15 | 3 | 3 | 28.4 | 14 | 249 | 10.07 | | | 1 | 1 | 2 | 2 | | | 1 | 1 | 2 | 2 |
| High | P29034 | Protein S100-A2 OS=Homo sapiens OX=9606 GN=S100A2 PE=1 SV=3 | 15 | 2 | 2 | 11.1 | 19 | 98 | 4.78 | 2 | 2 | 2 | 1 | 1 | | 2 | 2 | 2 | 1 | 1 | |
| High | P50552 | Vasodilator-stimulated phosphoprotein OS=Homo sapiens OX=9606 GN=VASP PE=1 SV=3 | 15 | 5 | 5 | 39.8 | 16 | 380 | 8.94 | 3 | 2 | 2 | | | | 3 | 2 | 2 | | | |
| High | O00154 | Cytosolic acyl coenzyme A thioester hydrolase OS=Homo sapiens OX=9606 GN=ACOT7 PE=1 SV=3 | 15 | 4 | 4 | 41.8 | 11 | 380 | 8.54 | 2 | 3 | 2 | | 2 | | 2 | 3 | 2 | | 2 | |
| High | Q9P1Y5 | Calmodulin-regulated spectrin-associated protein 3 OS=Homo sapiens OX=9606 GN=CAMSAP3 PE=1 SV=2 | 15 | 5 | 5 | 134.7 | 5 | 1249 | 8.35 | 1 | 1 | 2 | 1 | 1 | 1 | 1 | 1 | 2 | 1 | 1 | 1 |
| High | Q9H4M9 | EH domain-containing protein 1 OS=Homo sapiens OX=9606 GN=EHD1 PE=1 SV=2 | 15 | 5 | 4 | 60.6 | 13 | 534 | 6.83 | 2 | 3 | 3 | 1 | 1 | 1 | 2 | 3 | 2 | 1 | 1 | 1 |
| High | Q9NVS9 | Pyridoxine-5'-phosphate oxidase OS=Homo sapiens OX=9606 GN=PNPO PE=1 SV=1 | 15 | 6 | 6 | 30 | 26 | 261 | 7.06 | | | 1 | 1 | 2 | 2 | | | 1 | 1 | 2 | 2 |
| High | Q9Y6G9 | Cytoplasmic dynein 1 light intermediate chain 1 OS=Homo sapiens OX=9606 GN=DYNC1LI1 PE=1 SV=3 | 15 | 4 | 4 | 56.5 | 8 | 523 | 6.42 | 2 | 2 | 2 | | 1 | | 2 | 2 | 2 | | 1 | |
| High | P53985 | Monocarboxylate transporter 1 OS=Homo sapiens OX=9606 GN=SLC16A1 PE=1 SV=3 | 15 | 2 | 2 | 53.9 | 6 | 500 | 8.66 | 2 | 2 | 2 | 1 | 1 | 1 | 2 | 2 | 1 | 1 | 1 | 1 |
| High | Q9H3P7 | Golgi resident protein GCP60 OS=Homo sapiens OX=9606 GN=ACBD3 PE=1 SV=4 | 15 | 6 | 6 | 60.6 | 14 | 528 | 5.06 | | 1 | 1 | | 2 | | | 1 | 1 | | 2 | |
| High | P78385 | SWISS-PROT:P78385 Keratin, type II cuticular Hb3 (Hair keratin, type II Hb3) - Homo sapiens (Human). | 14 | 2 | 1 | 54.2 | 4 | 493 | 5.4 | 2 | 2 | 2 | 1 | 1 | 1 | 1 | 1 | 1 | 1 | 1 | 1 |
| High | Q12815 | Tastin OS=Homo sapiens OX=9606 GN=TROAP PE=1 SV=3 | 14 | 8 | 8 | 83.8 | 13 | 778 | 7.36 | | | | | 1 | | | | | | 1 | |
| High | Q9HAU0 | Pleckstrin homology domain-containing family A member 5 OS=Homo sapiens OX=9606 GN=PLEKHA5 PE=1 SV=1 | 14 | 5 | 5 | 127.4 | 5 | 1116 | 7.53 | 1 | 1 | 2 | | 2 | | 1 | 1 | 2 | | 2 | |
| High | Q96AE4 | Far upstream element-binding protein 1 OS=Homo sapiens OX=9606 GN=FUBP1 PE=1 SV=3 | 14 | 4 | 2 | 67.5 | 6 | 644 | 7.61 | 1 | 2 | 1 | | 1 | 2 | 1 | 2 | 1 | | 1 | 2 |
| High | Q13131 | 5'-AMP-activated protein kinase catalytic subunit alpha-1 OS=Homo sapiens OX=9606 GN=PRKAA1 PE=1 SV=4 | 14 | 5 | 5 | 64 | 11 | 559 | 8.12 | 1 | 1 | 1 | | 3 | 1 | 1 | 1 | 1 | | 3 | 1 |
| High | Q14517 | Protocadherin Fat 1 OS=Homo sapiens OX=9606 GN=FAT1 PE=1 SV=2 | 14 | 6 | 6 | 506 | 2 | 4588 | 5 | 2 | 4 | 4 | | 2 | | 2 | 4 | 4 | | 2 | |
| High | P55199 | RNA polymerase II elongation factor ELL OS=Homo sapiens OX=9606 GN=ELL PE=1 SV=1 | 14 | 5 | 5 | 68.2 | 10 | 621 | 9.33 | | | | 2 | 3 | 3 | | | | 2 | 3 | 3 |
| High | Q8N6M0 | Deubiquitinase OTUD6B OS=Homo sapiens OX=9606 GN=OTUD6B PE=1 SV=1 | 14 | 3 | 3 | 33.8 | 11 | 293 | 6.05 | 1 | 2 | 2 | 2 | 2 | 1 | 1 | 2 | 2 | 2 | 2 | 1 |
| High | Q05086 | Ubiquitin-protein ligase E3A OS=Homo sapiens OX=9606 GN=UBE3A PE=1 SV=4 | 14 | 5 | 5 | 100.6 | 7 | 875 | 5.22 | | | 3 | 2 | 1 | 2 | | | 3 | 2 | 1 | 2 |
| High | Q9NUI1 | Peroxisomal 2,4-dienoyl-CoA reductase [(3E)-enoyl-CoA-producing] OS=Homo sapiens OX=9606 GN=DECR2 PE=1 SV=1 | 14 | 6 | 6 | 30.8 | 21 | 292 | 9.22 | 1 | | | 1 | 2 | | 1 | | | 1 | 2 | |
| High | Q01105 | Protein SET OS=Homo sapiens OX=9606 GN=SET PE=1 SV=3 | 14 | 3 | 3 | 33.5 | 12 | 290 | 4.32 | | | 2 | 3 | 1 | 2 | | | 2 | 3 | 1 | 2 |
| High | Q4G0N4 | NAD kinase 2, mitochondrial OS=Homo sapiens OX=9606 GN=NADK2 PE=1 SV=2 | 14 | 6 | 6 | 49.4 | 13 | 442 | 8.18 | | | | 2 | 1 | 2 | | | | 2 | 1 | 2 |
| High | Q8NFF5 | FAD synthase OS=Homo sapiens OX=9606 GN=FLAD1 PE=1 SV=1 | 14 | 8 | 8 | 65.2 | 19 | 587 | 6.93 | 1 | | | 2 | 3 | 2 | 1 | | | 2 | 3 | 2 |
| High | Q8IUC4 | Rhophilin-2 OS=Homo sapiens OX=9606 GN=RHPN2 PE=1 SV=1 | 14 | 5 | 5 | 76.9 | 11 | 686 | 6.8 | 2 | 1 | 2 | | | | 2 | 1 | 2 | | | |
| High | Q86UE8 | Serine/threonine-protein kinase tousled-like 2 OS=Homo sapiens OX=9606 GN=TLK2 PE=1 SV=2 | 14 | 3 | 2 | 87.6 | 3 | 772 | 8.41 | | | 1 | 2 | 2 | 1 | | | 1 | 1 | 1 | 1 |
| High | P49184 | Deoxyribonuclease-1-like 1 OS=Homo sapiens OX=9606 GN=DNASE1L1 PE=1 SV=1 | 14 | 3 | 3 | 33.9 | 15 | 302 | 5.74 | 2 | 2 | 3 | | | | 2 | 2 | 3 | | | |
| High | P13688 | Carcinoembryonic antigen-related cell adhesion molecule 1 OS=Homo sapiens OX=9606 GN=CEACAM1 PE=1 SV=2 | 14 | 3 | 2 | 57.5 | 8 | 526 | 5.97 | 2 | 1 | 1 | 1 | 1 | 1 | 2 | 1 | 1 | 1 | 1 | 1 |
| High | Q02338 | D-beta-hydroxybutyrate dehydrogenase, mitochondrial OS=Homo sapiens OX=9606 GN=BDH1 PE=1 SV=3 | 14 | 6 | 6 | 38.1 | 22 | 343 | 8.95 | 1 | 1 | | | | | 1 | 1 | | | | |
| High | Q6P3X3 | Tetratricopeptide repeat protein 27 OS=Homo sapiens OX=9606 GN=TTC27 PE=1 SV=1 | 14 | 6 | 6 | 96.6 | 9 | 843 | 5.59 | 4 | 4 | 5 | | | | 4 | 4 | 5 | | | |
| High | Q9NS69 | Mitochondrial import receptor subunit TOM22 homolog OS=Homo sapiens OX=9606 GN=TOMM22 PE=1 SV=3 | 14 | 4 | 4 | 15.5 | 51 | 142 | 4.34 | 3 | 3 | 4 | 1 | | 1 | 2 | 2 | 3 | 1 | | 1 |
| High | P62312 | U6 snRNA-associated Sm-like protein LSm6 OS=Homo sapiens OX=9606 GN=LSM6 PE=1 SV=1 | 14 | 2 | 2 | 9.1 | 19 | 80 | 9.58 | 1 | | 1 | 2 | 2 | 2 | 1 | | 1 | 2 | 2 | 2 |
| High | P35249 | Replication factor C subunit 4 OS=Homo sapiens OX=9606 GN=RFC4 PE=1 SV=2 | 14 | 7 | 7 | 39.7 | 19 | 363 | 8.02 | | | 1 | | 2 | | | | 1 | | 1 | |
| High | P49757 | Protein numb homolog OS=Homo sapiens OX=9606 GN=NUMB PE=1 SV=2 | 14 | 5 | 5 | 70.8 | 9 | 651 | 8.51 | 2 | 2 | | 1 | 2 | 1 | 2 | 2 | | 1 | 2 | 1 |
| High | Q9BUA3 | Spindlin interactor and repressor of chromatin-binding protein OS=Homo sapiens OX=9606 GN=SPINDOC PE=1 SV=3 | 14 | 6 | 6 | 41 | 20 | 381 | 5.01 | 4 | 4 | 5 | | 1 | | 4 | 4 | 5 | | 1 | |
| High | P0C7P4 | Putative cytochrome b-c1 complex subunit Rieske-like protein 1 OS=Homo sapiens OX=9606 GN=UQCRFS1P1 PE=5 SV=1 | 14 | 5 | 1 | 30.8 | 17 | 283 | 8.87 | 2 | 1 | 4 | 2 | 1 | 1 | 2 | 1 | 4 | 2 | 1 | 1 |
| High | P23919 | Thymidylate kinase OS=Homo sapiens OX=9606 GN=DTYMK PE=1 SV=4 | 14 | 4 | 4 | 23.8 | 17 | 212 | 8.27 | | 1 | 2 | 1 | | 1 | | 1 | 2 | 1 | | 1 |
| High | Q8IXI1 | Mitochondrial Rho GTPase 2 OS=Homo sapiens OX=9606 GN=RHOT2 PE=1 SV=2 | 14 | 6 | 5 | 68.1 | 17 | 618 | 5.86 | 1 | 1 | 1 | 2 | 3 | 2 | 1 | 1 | 1 | 2 | 3 | 2 |
| High | Q8IWU2 | Serine/threonine-protein kinase LMTK2 OS=Homo sapiens OX=9606 GN=LMTK2 PE=1 SV=2 | 14 | 5 | 5 | 164.8 | 5 | 1503 | 4.48 | 1 | 1 | 4 | | | | 1 | 1 | 3 | | | |
| High | Q8TDD1 | ATP-dependent RNA helicase DDX54 OS=Homo sapiens OX=9606 GN=DDX54 PE=1 SV=2 | 14 | 7 | 6 | 98.5 | 10 | 881 | 10.02 | | | | | | | | | | | | |
| High | P22059 | Oxysterol-binding protein 1 OS=Homo sapiens OX=9606 GN=OSBP PE=1 SV=1 | 14 | 4 | 4 | 89.4 | 6 | 807 | 7.3 | 1 | 1 | 2 | | 1 | | 1 | 1 | 2 | | 1 | |
| High | Q92643 | GPI-anchor transamidase OS=Homo sapiens OX=9606 GN=PIGK PE=1 SV=2 | 14 | 5 | 5 | 45.2 | 16 | 395 | 6.16 | 2 | 3 | 3 | 1 | 1 | 1 | 2 | 3 | 3 | 1 | 1 | 1 |
| High | Q8TEQ6 | Gem-associated protein 5 OS=Homo sapiens OX=9606 GN=GEMIN5 PE=1 SV=3 | 14 | 5 | 5 | 168.5 | 4 | 1508 | 6.62 | | 1 | 1 | | 2 | | | 1 | 1 | | 2 | |
| High | Q9NX20 | 39S ribosomal protein L16, mitochondrial OS=Homo sapiens OX=9606 GN=MRPL16 PE=1 SV=1 | 14 | 5 | 5 | 28.4 | 20 | 251 | 10.13 | 3 | 1 | 2 | | 1 | | 3 | 1 | 2 | | 1 | |
| High | Q9BXS6 | Nucleolar and spindle-associated protein 1 OS=Homo sapiens OX=9606 GN=NUSAP1 PE=1 SV=1 | 14 | 4 | 4 | 49.4 | 13 | 441 | 9.91 | 1 | 1 | | 1 | 2 | 1 | 1 | 1 | | 1 | 2 | 1 |
| High | P60468 | Protein transport protein Sec61 subunit beta OS=Homo sapiens OX=9606 GN=SEC61B PE=1 SV=2 | 14 | 2 | 2 | 10 | 22 | 96 | 11.56 | 2 | 2 | 1 | 1 | 1 | 1 | 2 | 2 | 1 | 1 | 1 | 1 |
| High | Q8WVJ2 | NudC domain-containing protein 2 OS=Homo sapiens OX=9606 GN=NUDCD2 PE=1 SV=1 | 14 | 2 | 2 | 17.7 | 13 | 157 | 5.07 | 3 | 2 | 2 | | 1 | 1 | 2 | 1 | 2 | | 1 | 1 |
| High | Q8NB16 | Mixed lineage kinase domain-like protein OS=Homo sapiens OX=9606 GN=MLKL PE=1 SV=1 | 14 | 5 | 5 | 54.4 | 14 | 471 | 8.82 | 1 | 2 | 1 | | 1 | | 1 | 2 | 1 | | 1 | |
| High | Q9Y316 | Protein MEMO1 OS=Homo sapiens OX=9606 GN=MEMO1 PE=1 SV=1 | 14 | 5 | 5 | 33.7 | 18 | 297 | 7.14 | 2 | 1 | 2 | 1 | | 1 | 2 | 1 | 2 | 1 | | 1 |
| High | Q92530 | Proteasome inhibitor PI31 subunit OS=Homo sapiens OX=9606 GN=PSMF1 PE=1 SV=2 | 14 | 3 | 3 | 29.8 | 14 | 271 | 5.74 | 1 | 2 | 1 | 1 | 2 | 1 | 1 | 2 | 1 | 1 | 2 | 1 |
| High | Q13155 | Aminoacyl tRNA synthase complex-interacting multifunctional protein 2 OS=Homo sapiens OX=9606 GN=AIMP2 PE=1 SV=2 | 14 | 5 | 5 | 35.3 | 23 | 320 | 8.22 | 1 | | 2 | 1 | 1 | | 1 | | 2 | 1 | 1 | |
| High | Q9BT78 | COP9 signalosome complex subunit 4 OS=Homo sapiens OX=9606 GN=COPS4 PE=1 SV=1 | 14 | 6 | 6 | 46.2 | 18 | 406 | 5.83 | 2 | | 2 | 1 | | 2 | 2 | | 2 | 1 | | 2 |
| High | P62861 | FAU ubiquitin-like and ribosomal protein S30 OS=Homo sapiens OX=9606 GN=FAU PE=1 SV=2 | 14 | 2 | 2 | 14.4 | 8 | 133 | 10.17 | 1 | 1 | 2 | 1 | 1 | 1 | 1 | 1 | 1 | 1 | 1 | 1 |
| High | Q02241 | Kinesin-like protein KIF23 OS=Homo sapiens OX=9606 GN=KIF23 PE=1 SV=3 | 14 | 5 | 5 | 110 | 7 | 960 | 8.51 | 2 | 3 | 2 | | | | 2 | 3 | 2 | | | |
| High | Q14004 | Cyclin-dependent kinase 13 OS=Homo sapiens OX=9606 GN=CDK13 PE=1 SV=2 | 14 | 2 | 1 | 164.8 | 1 | 1512 | 9.69 | 1 | 2 | 2 | 2 | 2 | 1 | 1 | 1 | 1 | 1 | 1 | 1 |
| High | Q9BQ39 | ATP-dependent RNA helicase DDX50 OS=Homo sapiens OX=9606 GN=DDX50 PE=1 SV=1 | 14 | 4 | 2 | 82.5 | 6 | 737 | 9.17 | | | | 1 | 4 | 1 | | | | 1 | 3 | 1 |
| High | Q9NZI7 | Upstream-binding protein 1 OS=Homo sapiens OX=9606 GN=UBP1 PE=1 SV=1 | 14 | 5 | 4 | 60.5 | 13 | 540 | 6.35 | 4 | 3 | 4 | | | | 4 | 3 | 4 | | | |
| High | Q9BWE0 | Replication initiator 1 OS=Homo sapiens OX=9606 GN=REPIN1 PE=1 SV=1 | 14 | 3 | 3 | 63.5 | 8 | 567 | 9.98 | | | | 1 | | 2 | | | | 1 | | 2 |
| High | Q8WV41 | Sorting nexin-33 OS=Homo sapiens OX=9606 GN=SNX33 PE=1 SV=1 | 14 | 4 | 4 | 65.2 | 9 | 574 | 6.79 | | | 2 | 2 | 1 | 2 | | | 2 | 2 | 1 | 2 |
| High | Q9UID6 | Zinc finger protein 639 OS=Homo sapiens OX=9606 GN=ZNF639 PE=1 SV=1 | 14 | 7 | 7 | 56 | 17 | 485 | 5.57 | | | | 2 | 2 | 2 | | | | 2 | 2 | 2 |
| High | Q12830 | Nucleosome-remodeling factor subunit BPTF OS=Homo sapiens OX=9606 GN=BPTF PE=1 SV=3 | 14 | 7 | 7 | 338.1 | 2 | 3046 | 6.54 | | | | 2 | 4 | 3 | | | | 2 | 4 | 3 |
| High | Q02790 | Peptidyl-prolyl cis-trans isomerase FKBP4 OS=Homo sapiens OX=9606 GN=FKBP4 PE=1 SV=3 | 14 | 5 | 5 | 51.8 | 9 | 459 | 5.43 | 2 | 3 | 3 | 1 | | | 2 | 3 | 3 | 1 | | |
| High | Q5TFE4 | 5'-nucleotidase domain-containing protein 1 OS=Homo sapiens OX=9606 GN=NT5DC1 PE=1 SV=1 | 14 | 3 | 3 | 51.8 | 8 | 455 | 6.35 | 1 | 1 | 1 | 1 | 3 | 1 | 1 | 1 | 1 | 1 | 3 | 1 |
| High | Q16875 | 6-phosphofructo-2-kinase/fructose-2,6-bisphosphatase 3 OS=Homo sapiens OX=9606 GN=PFKFB3 PE=1 SV=1 | 14 | 7 | 5 | 59.6 | 16 | 520 | 8.21 | | | 1 | 1 | 1 | | | | 1 | 1 | 1 | |
| High | Q9NYV4 | Cyclin-dependent kinase 12 OS=Homo sapiens OX=9606 GN=CDK12 PE=1 SV=2 | 14 | 2 | 1 | 164.1 | 1 | 1490 | 9.44 | 1 | 2 | 2 | 2 | 2 | 1 | 1 | 1 | 1 | 1 | 1 | 1 |
| High | P06400 | Retinoblastoma-associated protein OS=Homo sapiens OX=9606 GN=RB1 PE=1 SV=2 | 14 | 5 | 5 | 106.1 | 6 | 928 | 7.94 | 2 | 3 | | 1 | 2 | | 2 | 3 | | 1 | 2 | |
| High | Q9P016 | Thymocyte nuclear protein 1 OS=Homo sapiens OX=9606 GN=THYN1 PE=1 SV=1 | 14 | 4 | 4 | 25.7 | 16 | 225 | 9.25 | 1 | 1 | 2 | 1 | 2 | 1 | 1 | 1 | 2 | 1 | 2 | 1 |
| High | Q14152 | Eukaryotic translation initiation factor 3 subunit A OS=Homo sapiens OX=9606 GN=EIF3A PE=1 SV=1 | 14 | 5 | 5 | 166.5 | 4 | 1382 | 6.79 | 1 | | 2 | 2 | 2 | 1 | 1 | | 2 | 2 | 2 | 1 |
| High | Q9HB21 | Pleckstrin homology domain-containing family A member 1 OS=Homo sapiens OX=9606 GN=PLEKHA1 PE=1 SV=2 | 14 | 4 | 4 | 45.5 | 13 | 404 | 8.56 | 2 | 2 | 1 | | | | 2 | 2 | 1 | | | |
| High | Q9BWM7 | Sideroflexin-3 OS=Homo sapiens OX=9606 GN=SFXN3 PE=1 SV=3 | 14 | 3 | 3 | 35.5 | 10 | 321 | 9.1 | 2 | 2 | 3 | 1 | | 1 | 2 | 2 | 3 | 1 | | |
| High | Q9Y3B4 | Splicing factor 3B subunit 6 OS=Homo sapiens OX=9606 GN=SF3B6 PE=1 SV=1 | 14 | 4 | 4 | 14.6 | 38 | 125 | 9.38 | 1 | 1 | 1 | 1 | 2 | 1 | 1 | 1 | 1 | 1 | 2 | 1 |
| High | P49750 | YLP motif-containing protein 1 OS=Homo sapiens OX=9606 GN=YLPM1 PE=1 SV=4 | 14 | 5 | 5 | 241.5 | 4 | 2146 | 6.55 | 3 | 3 | 2 | 1 | | | 3 | 3 | 2 | 1 | | |
| High | P52434 | DNA-directed RNA polymerases I, II, and III subunit RPABC3 OS=Homo sapiens OX=9606 GN=POLR2H PE=1 SV=4 | 14 | 3 | 3 | 17.1 | 27 | 150 | 4.68 | | 1 | 1 | 1 | 2 | 2 | | 1 | 1 | 1 | 2 | 2 |
| High | Q96JP5 | E3 ubiquitin-protein ligase ZFP91 OS=Homo sapiens OX=9606 GN=ZFP91 PE=1 SV=1 | 14 | 4 | 4 | 63.4 | 10 | 570 | 7.36 | | | | 1 | | 1 | | | | 1 | | 1 |
| High | Q9P1Z2 | Calcium-binding and coiled-coil domain-containing protein 1 OS=Homo sapiens OX=9606 GN=CALCOCO1 PE=1 SV=2 | 14 | 4 | 4 | 77.3 | 6 | 691 | 4.82 | 1 | | 2 | 1 | 1 | 1 | 1 | | 2 | 1 | 1 | 1 |
| High | O60307 | Microtubule-associated serine/threonine-protein kinase 3 OS=Homo sapiens OX=9606 GN=MAST3 PE=1 SV=2 | 14 | 2 | 1 | 143 | 1 | 1309 | 8.06 | | | 1 | 2 | 3 | 1 | | | 1 | 1 | 2 | 1 |
| High | Q9HCD5 | Nuclear receptor coactivator 5 OS=Homo sapiens OX=9606 GN=NCOA5 PE=1 SV=2 | 14 | 6 | 6 | 65.5 | 11 | 579 | 9.6 | 4 | 4 | 2 | 1 | 2 | 1 | 4 | 4 | 2 | 1 | 2 | 1 |

| | | | | | | | | | | | | | | | | | | | | | |
|---|---|---|---|---|---|---|---|---|---|---|---|---|---|---|---|---|---|---|---|---|---|
| High | Q6Y7W6 | GRB10-interacting GYF protein 2 OS=Homo sapiens OX=9606 GN=GIGYF2 PE=1 SV=1 | 14 | 5 | 5 | 150 | 6 | 1299 | 5.54 | 1 | 1 | 2 | | | | 1 | 1 | 2 | | | |
| High | Q53EP0 | Fibronectin type III domain-containing protein 3B OS=Homo sapiens OX=9606 GN=FNDC3B PE=1 SV=2 | 13 | 4 | 4 | 132.8 | 4 | 1204 | 5.95 | 1 | 1 | 2 | | 2 | | 1 | 1 | 2 | | 2 | |
| High | Q96FV9 | THO complex subunit 1 OS=Homo sapiens OX=9606 GN=THOC1 PE=1 SV=1 | 13 | 5 | 5 | 75.6 | 9 | 657 | 4.98 | | | | 2 | 1 | 2 | | | | 2 | 1 | 2 |
| High | Q9Y6D5 | Brefeldin A-inhibited guanine nucleotide-exchange protein 2 OS=Homo sapiens OX=9606 GN=ARFGEF2 PE=1 SV=3 | 13 | 5 | 4 | 201.9 | 3 | 1785 | 6.33 | 1 | 1 | 3 | | | | 1 | 1 | 3 | | | |
| High | P31944 | Caspase-14 OS=Homo sapiens OX=9606 GN=CASP14 PE=1 SV=2 | 13 | 6 | 6 | 27.7 | 28 | 242 | 5.58 | | | | 4 | 5 | 4 | | | | 4 | 5 | 4 |
| High | O43824 | Putative GTP-binding protein 6 OS=Homo sapiens OX=9606 GN=GTPBP6 PE=1 SV=4 | 13 | 4 | 4 | 56.9 | 7 | 516 | 9.42 | 1 | 2 | | 1 | 1 | 1 | 1 | 2 | | 1 | 1 | 1 |
| High | P28838 | Cytosol aminopeptidase OS=Homo sapiens OX=9606 GN=LAP3 PE=1 SV=3 | 13 | 4 | 4 | 56.1 | 11 | 519 | 7.93 | 2 | 3 | | 1 | 1 | | 2 | 3 | | 1 | 1 | |
| High | P23497 | Nuclear autoantigen Sp-100 OS=Homo sapiens OX=9606 GN=SP100 PE=1 SV=3 | 13 | 5 | 5 | 100.4 | 8 | 879 | 8.22 | | | | 1 | 2 | 3 | | | | 1 | 2 | 3 |
| High | Q96I15 | Selenocysteine lyase OS=Homo sapiens OX=9606 GN=SCLY PE=1 SV=4 | 13 | 5 | 5 | 48.1 | 18 | 445 | 7.12 | 1 | | 2 | | | 1 | 1 | | 2 | | | 1 |
| High | P30085 | UMP-CMP kinase OS=Homo sapiens OX=9606 GN=CMPK1 PE=1 SV=3 | 13 | 5 | 5 | 22.2 | 22 | 196 | 5.57 | | | 1 | | | 1 | | | 1 | | | 1 |
| High | Q9H0H5 | Rac GTPase-activating protein 1 OS=Homo sapiens OX=9606 GN=RACGAP1 PE=1 SV=1 | 13 | 3 | 3 | 71 | 5 | 632 | 8.88 | 2 | 2 | 3 | | 1 | 1 | 2 | 2 | 3 | | 1 | 1 |
| High | Q9Y2L1 | Exosome complex exonuclease RRP44 OS=Homo sapiens OX=9606 GN=DIS3 PE=1 SV=2 | 13 | 5 | 5 | 108.9 | 9 | 958 | 7.14 | 1 | 1 | 1 | | | | 1 | 1 | 1 | | | |
| High | Q10589 | Bone marrow stromal antigen 2 OS=Homo sapiens OX=9606 GN=BST2 PE=1 SV=1 | 13 | 2 | 2 | 19.8 | 13 | 180 | 5.6 | 2 | 1 | 1 | 1 | 2 | 1 | 1 | 1 | 1 | 1 | 1 | 1 |
| High | Q5VT25 | Serine/threonine-protein kinase MRCK alpha OS=Homo sapiens OX=9606 GN=CDC42BPA PE=1 SV=1 | 13 | 7 | 3 | 197.2 | 5 | 1732 | 6.58 | | | 1 | | | | | | 1 | | | |
| High | Q15007 | Pre-mRNA-splicing regulator WTAP OS=Homo sapiens OX=9606 GN=WTAP PE=1 SV=2 | 13 | 5 | 5 | 44.2 | 17 | 396 | 5.19 | | 1 | 1 | 3 | 2 | 2 | | 1 | 1 | 3 | 2 | 2 |
| High | O95394 | Phosphoacetylglucosamine mutase OS=Homo sapiens OX=9606 GN=PGM3 PE=1 SV=1 | 13 | 5 | 5 | 59.8 | 8 | 542 | 6.25 | 1 | 2 | | 2 | 1 | | 1 | 2 | | 2 | 1 | |
| High | P12829 | Myosin light chain 4 OS=Homo sapiens OX=9606 GN=MYL4 PE=1 SV=3 | 13 | 5 | 4 | 21.6 | 24 | 197 | 5.03 | 1 | 1 | 1 | 5 | | 4 | 1 | 1 | 1 | 5 | | 4 |
| High | Q9NP72 | Ras-related protein Rab-18 OS=Homo sapiens OX=9606 GN=RAB18 PE=1 SV=1 | 13 | 5 | 5 | 23 | 30 | 206 | 5.24 | 1 | 1 | 1 | 1 | | 2 | 1 | 1 | 1 | 1 | | 2 |
| High | Q96RT1 | Erbin OS=Homo sapiens OX=9606 GN=ERBIN PE=1 SV=2 | 13 | 6 | 6 | 158.2 | 6 | 1412 | 5.5 | 2 | 3 | 4 | | 1 | | 2 | 3 | 4 | | 1 | |
| High | O75164 | Lysine-specific demethylase 4A OS=Homo sapiens OX=9606 GN=KDM4A PE=1 SV=2 | 13 | 8 | 8 | 120.6 | 8 | 1064 | 5.85 | | | | | 3 | 2 | | | | | 3 | 2 |
| High | P51665 | 26S proteasome non-ATPase regulatory subunit 7 OS=Homo sapiens OX=9606 GN=PSMD7 PE=1 SV=2 | 13 | 3 | 3 | 37 | 10 | 324 | 6.77 | | | 1 | 2 | 2 | 2 | | | 1 | 2 | 2 | 2 |
| High | Q96EY5 | Multivesicular body subunit 12A OS=Homo sapiens OX=9606 GN=MVB12A PE=1 SV=1 | 13 | 4 | 4 | 28.8 | 26 | 273 | 8.91 | 1 | 1 | 2 | 1 | | | 1 | 1 | 2 | 1 | | |
| High | P63218 | Guanine nucleotide-binding protein G(I)/G(S)/G(O) subunit gamma-5 OS=Homo sapiens OX=9606 GN=GNG5 PE=1 SV=3 | 13 | 2 | 2 | 7.3 | 24 | 68 | 9.85 | 1 | 1 | 2 | 2 | 1 | 1 | 1 | 1 | 2 | 2 | 1 | 1 |
| High | P00533 | Epidermal growth factor receptor OS=Homo sapiens OX=9606 GN=EGFR PE=1 SV=2 | 13 | 4 | 3 | 134.2 | 6 | 1210 | 6.68 | 2 | 3 | 2 | | | | 2 | 3 | 2 | | | |
| High | P14550 | Aldo-keto reductase family 1 member A1 OS=Homo sapiens OX=9606 GN=AKR1A1 PE=1 SV=3 | 13 | 2 | 2 | 36.6 | 5 | 325 | 6.79 | 1 | 1 | 2 | 2 | 2 | 2 | 1 | 1 | 2 | 2 | 2 | 2 |
| High | O15067 | Phosphoribosylformylglycinamidine synthase OS=Homo sapiens OX=9606 GN=PFAS PE=1 SV=4 | 13 | 8 | 8 | 144.6 | 7 | 1338 | 5.76 | | 1 | 1 | | 3 | 1 | | 1 | 1 | | 3 | 1 |
| High | Q5RKV6 | Exosome complex component MTR3 OS=Homo sapiens OX=9606 GN=EXOSC6 PE=1 SV=1 | 13 | 4 | 4 | 28.2 | 16 | 272 | 6.28 | | 1 | 1 | 2 | 2 | 1 | | 1 | 1 | 2 | 2 | 1 |
| High | Q9BUQ8 | Probable ATP-dependent RNA helicase DDX23 OS=Homo sapiens OX=9606 GN=DDX23 PE=1 SV=3 | 13 | 4 | 4 | 95.5 | 4 | 820 | 9.55 | | 1 | | 1 | 1 | 2 | | 1 | | 1 | 1 | 1 |
| High | Q6DKI2 | Galectin-9C OS=Homo sapiens OX=9606 GN=LGALS9C PE=1 SV=2 | 13 | 4 | 1 | 39.6 | 13 | 356 | 9.19 | 1 | | 1 | 1 | 2 | 2 | 1 | | 1 | 1 | 2 | 2 |
| High | Q9BVL2 | Nucleoporin p58/p45 OS=Homo sapiens OX=9606 GN=NUP58 PE=1 SV=1 | 13 | 5 | 5 | 60.9 | 9 | 599 | 9.33 | | 1 | | 1 | | 1 | | 1 | | 1 | | 1 |
| High | O15511 | Actin-related protein 2/3 complex subunit 5 OS=Homo sapiens OX=9606 GN=ARPC5 PE=1 SV=3 | 13 | 4 | 3 | 16.3 | 34 | 151 | 5.67 | 2 | 1 | 2 | | 1 | | 2 | 1 | 2 | | 1 | |
| High | P61224 | Ras-related protein Rap-1b OS=Homo sapiens OX=9606 GN=RAP1B PE=1 SV=1 | 13 | 3 | 1 | 20.8 | 21 | 184 | 5.78 | 2 | | 1 | 1 | 2 | 1 | 2 | | 1 | 1 | 2 | 1 |
| High | Q9HBH1 | Peptide deformylase, mitochondrial OS=Homo sapiens OX=9606 GN=PDF PE=1 SV=1 | 13 | 4 | 4 | 27 | 20 | 243 | 9.16 | 1 | 1 | 1 | | | | 1 | 1 | 1 | | | |
| High | Q8WYP5 | Protein ELYS OS=Homo sapiens OX=9606 GN=AHCTF1 PE=1 SV=3 | 13 | 5 | 5 | 252.3 | 3 | 2266 | 6.6 | | | 1 | 1 | 1 | | | | 1 | 1 | 1 | |
| High | P02675 | Fibrinogen beta chain OS=Homo sapiens OX=9606 GN=FGB PE=1 SV=2 | 13 | 6 | 3 | 55.9 | 12 | 491 | 8.27 | | | | 4 | 4 | 5 | | | | 4 | 4 | 5 |
| High | P78312 | Protein FAM193A OS=Homo sapiens OX=9606 GN=FAM193A PE=1 SV=2 | 13 | 6 | 6 | 139.9 | 6 | 1265 | 6.48 | | | | 4 | 4 | 4 | | | | 3 | 4 | 4 |
| High | O96017 | Serine/threonine-protein kinase Chk2 OS=Homo sapiens OX=9606 GN=CHEK2 PE=1 SV=1 | 13 | 5 | 4 | 60.9 | 9 | 543 | 5.91 | 1 | 1 | 1 | 1 | 3 | | 1 | 1 | 1 | 1 | 3 | |
| High | Q86YT6 | E3 ubiquitin-protein ligase MIB1 OS=Homo sapiens OX=9606 GN=MIB1 PE=1 SV=1 | 13 | 4 | 4 | 110.1 | 4 | 1006 | 6.92 | 2 | 1 | 2 | | 1 | | 2 | 1 | 2 | | 1 | |
| High | P17676 | CCAAT/enhancer-binding protein beta OS=Homo sapiens OX=9606 GN=CEBPB PE=1 SV=2 | 13 | 4 | 4 | 36.1 | 17 | 345 | 8.31 | 2 | 2 | 2 | | 1 | 2 | 2 | 2 | 2 | | 1 | 2 |
| High | O43663 | Protein regulator of cytokinesis 1 OS=Homo sapiens OX=9606 GN=PRC1 PE=1 SV=2 | 13 | 3 | 3 | 71.6 | 6 | 620 | 6.68 | | | 1 | 1 | 2 | 1 | | | 1 | 1 | 2 | 1 |
| High | O14735 | CDP-diacylglycerol--inositol 3-phosphatidyltransferase OS=Homo sapiens OX=9606 GN=CDIPT PE=1 SV=1 | 13 | 2 | 2 | 23.5 | 10 | 213 | 8.03 | 2 | 1 | 1 | 2 | 1 | | 2 | 1 | 1 | 2 | 1 | |
| High | Q16762 | Thiosulfate sulfurtransferase OS=Homo sapiens OX=9606 GN=TST PE=1 SV=4 | 13 | 6 | 6 | 33.4 | 33 | 297 | 7.25 | 2 | 3 | 1 | | | | 2 | 3 | 1 | | | |
| High | P05198 | Eukaryotic translation initiation factor 2 subunit 1 OS=Homo sapiens OX=9606 GN=EIF2S1 PE=1 SV=3 | 13 | 6 | 6 | 36.1 | 21 | 315 | 5.08 | 1 | 2 | 1 | 2 | 1 | 1 | 1 | 2 | 1 | 2 | 1 | 1 |
| High | O75909 | Cyclin-K OS=Homo sapiens OX=9606 GN=CCNK PE=1 SV=2 | 13 | 4 | 4 | 64.2 | 11 | 580 | 8.41 | | | | 2 | 2 | 1 | | | | 2 | 2 | 1 |
| High | Q6IAA8 | Ragulator complex protein LAMTOR1 OS=Homo sapiens OX=9606 GN=LAMTOR1 PE=1 SV=2 | 13 | 2 | 2 | 17.7 | 15 | 161 | 5.15 | 2 | 1 | 2 | | 1 | | 2 | 1 | 2 | | 1 | |
| High | P55735 | Protein SEC13 homolog OS=Homo sapiens OX=9606 GN=SEC13 PE=1 SV=3 | 13 | 3 | 3 | 35.5 | 15 | 322 | 5.48 | 1 | | | 1 | 1 | | 1 | | | 1 | 1 | |
| High | Q16769 | Glutaminyl-peptide cyclotransferase OS=Homo sapiens OX=9606 GN=QPCT PE=1 SV=1 | 13 | 3 | 3 | 40.9 | 12 | 361 | 6.61 | 1 | 1 | 1 | 1 | 2 | 2 | 1 | 1 | 1 | 1 | 2 | 2 |
| High | O95218 | Zinc finger Ran-binding domain-containing protein 2 OS=Homo sapiens OX=9606 GN=ZRANB2 PE=1 SV=2 | 13 | 6 | 6 | 37.4 | 24 | 330 | 10.01 | | 1 | 1 | | 2 | | | 1 | 1 | | 2 | |
| High | P13804 | Electron transfer flavoprotein subunit alpha, mitochondrial OS=Homo sapiens OX=9606 GN=ETFA PE=1 SV=1 | 13 | 6 | 6 | 35.1 | 26 | 333 | 8.38 | 3 | 4 | 3 | | | | 3 | 4 | 3 | | | |
| High | O43772 | Mitochondrial carnitine/acylcarnitine carrier protein OS=Homo sapiens OX=9606 GN=SLC25A20 PE=1 SV=1 | 13 | 5 | 5 | 32.9 | 19 | 301 | 9.41 | 2 | 1 | 4 | | | | 2 | 1 | 4 | | | |
| High | P41229 | Lysine-specific demethylase 5C OS=Homo sapiens OX=9606 GN=KDM5C PE=1 SV=2 | 13 | 7 | 6 | 175.6 | 6 | 1560 | 5.58 | 1 | 1 | 2 | | 1 | | 1 | 1 | 2 | | 1 | |
| High | Q9UQ80 | Proliferation-associated protein 2G4 OS=Homo sapiens OX=9606 GN=PA2G4 PE=1 SV=3 | 13 | 4 | 4 | 43.8 | 18 | 394 | 6.55 | 1 | 1 | 1 | | 3 | | 1 | 1 | 1 | | 2 | |
| High | Q9P1U1 | Actin-related protein 3B OS=Homo sapiens OX=9606 GN=ACTR3B PE=2 SV=1 | 13 | 3 | 1 | 47.6 | 10 | 418 | 5.88 | 2 | 2 | 1 | 1 | 1 | 2 | 2 | 2 | 1 | 1 | 1 | 2 |
| High | P18283 | Glutathione peroxidase 2 OS=Homo sapiens OX=9606 GN=GPX2 PE=1 SV=3 | 13 | 5 | 5 | 21.9 | 33 | 190 | 7.78 | | | 3 | 1 | 2 | | | | 3 | 1 | 2 | |
| High | Q14344 | Guanine nucleotide-binding protein subunit alpha-13 OS=Homo sapiens OX=9606 GN=GNA13 PE=1 SV=2 | 13 | 3 | 1 | 44 | 8 | 377 | 8 | 1 | 2 | 1 | 2 | 1 | 2 | 1 | 2 | 1 | 2 | 1 | 2 |
| High | Q86X76 | Deaminated glutathione amidase OS=Homo sapiens OX=9606 GN=NIT1 PE=1 SV=2 | 13 | 4 | 4 | 35.9 | 15 | 327 | 7.74 | 1 | | 2 | | 1 | | 1 | | 2 | | 1 | |
| High | Q9UJC5 | SH3 domain-binding glutamic acid-rich-like protein 2 OS=Homo sapiens OX=9606 GN=SH3BGRL2 PE=1 SV=2 | 13 | 3 | 3 | 12.3 | 36 | 107 | 6.71 | 1 | 2 | 3 | 1 | 1 | 1 | 1 | 2 | 3 | 1 | 1 | 1 |
| High | Q14692 | Ribosome biogenesis protein BMS1 homolog OS=Homo sapiens OX=9606 GN=BMS1 PE=1 SV=1 | 13 | 6 | 6 | 145.7 | 6 | 1282 | 6.44 | | | | | 2 | | | | | | 2 | |
| High | P09110 | 3-ketoacyl-CoA thiolase, peroxisomal OS=Homo sapiens OX=9606 GN=ACAA1 PE=1 SV=2 | 13 | 7 | 7 | 44.3 | 30 | 424 | 8.44 | | 2 | 1 | | 2 | | | 2 | 1 | | 2 | |
| High | Q9Y3A4 | Ribosomal RNA-processing protein 7 homolog A OS=Homo sapiens OX=9606 GN=RRP7A PE=1 SV=2 | 13 | 4 | 4 | 32.3 | 23 | 280 | 9.58 | 1 | 1 | | | 1 | 2 | 1 | 1 | | | 1 | 2 |
| High | P39748 | Flap endonuclease 1 OS=Homo sapiens OX=9606 GN=FEN1 PE=1 SV=1 | 13 | 2 | 2 | 42.6 | 8 | 380 | 8.62 | 2 | 2 | 2 | 1 | 1 | 1 | 2 | 2 | 2 | 1 | 1 | 1 |
| High | Q96BS2 | Calcineurin B homologous protein 3 OS=Homo sapiens OX=9606 GN=TESC PE=1 SV=3 | 13 | 4 | 4 | 24.7 | 21 | 214 | 4.93 | 3 | 3 | 2 | 1 | | 1 | 3 | 3 | 2 | 1 | | 1 |
| High | O15031 | Plexin-B2 OS=Homo sapiens OX=9606 GN=PLXNB2 PE=1 SV=3 | 13 | 5 | 5 | 205 | 4 | 1838 | 6.24 | 2 | 2 | 2 | | | | 2 | 2 | 2 | | | |
| High | Q96J01 | THO complex subunit 3 OS=Homo sapiens OX=9606 GN=THOC3 PE=1 SV=1 | 13 | 3 | 3 | 38.7 | 9 | 351 | 6.09 | | 1 | 2 | 1 | 1 | 1 | | 1 | 2 | 1 | 1 | 1 |
| High | Q6P179 | Endoplasmic reticulum aminopeptidase 2 OS=Homo sapiens OX=9606 GN=ERAP2 PE=1 SV=2 | 13 | 7 | 7 | 110.4 | 8 | 960 | 6.71 | 4 | 1 | 4 | | | 1 | 4 | 1 | 4 | | | 1 |
| High | Q9Y3D3 | 28S ribosomal protein S16, mitochondrial OS=Homo sapiens OX=9606 GN=MRPS16 PE=1 SV=1 | 13 | 3 | 3 | 15.3 | 25 | 137 | 9.5 | | | | 2 | 3 | 2 | | | | 2 | 3 | 2 |
| High | Q6ISB3 | Grainyhead-like protein 2 homolog OS=Homo sapiens OX=9606 GN=GRHL2 PE=1 SV=1 | 13 | 5 | 5 | 71.1 | 12 | 625 | 6.44 | 1 | 1 | 3 | 1 | 1 | 1 | 1 | 1 | 3 | 1 | 1 | 1 |
| High | Q9BSD7 | Cancer-related nucleoside-triphosphatase OS=Homo sapiens OX=9606 GN=NTPCR PE=1 SV=1 | 13 | 2 | 2 | 20.7 | 13 | 190 | 9.54 | 1 | | 2 | 2 | 1 | 1 | 1 | | 2 | 1 | 1 | 1 |
| High | Q8TE77 | Protein phosphatase Slingshot homolog 3 OS=Homo sapiens OX=9606 GN=SSH3 PE=1 SV=2 | 13 | 4 | 4 | 73 | 6 | 659 | 5.3 | | | 3 | 1 | | 1 | | | 2 | 1 | | 1 |
| High | Q9C0F0 | Putative Polycomb group protein ASXL3 OS=Homo sapiens OX=9606 GN=ASXL3 PE=2 SV=3 | 13 | 3 | 1 | 241.8 | 1 | 2248 | 6.14 | | | | 2 | 2 | 2 | | | | 2 | 2 | 2 |
| High | Q8N983 | 39S ribosomal protein L43, mitochondrial OS=Homo sapiens OX=9606 GN=MRPL43 PE=1 SV=1 | 13 | 4 | 4 | 23.4 | 17 | 215 | 8.65 | | 1 | | | 2 | 1 | | 1 | | | 2 | 1 |
| High | O14647 | Chromodomain-helicase-DNA-binding protein 2 OS=Homo sapiens OX=9606 GN=CHD2 PE=1 SV=2 | 12 | 5 | 4 | 211.2 | 3 | 1828 | 8.1 | 4 | 3 | 4 | | 1 | | 4 | 3 | 4 | | 1 | |
| High | P20020 | Plasma membrane calcium-transporting ATPase 1 OS=Homo sapiens OX=9606 GN=ATP2B1 PE=1 SV=4 | 12 | 6 | 6 | 134.6 | 7 | 1220 | 5.91 | 2 | 2 | 2 | | 1 | | 2 | 2 | 2 | | 1 | |
| High | Q13232 | Nucleoside diphosphate kinase 3 OS=Homo sapiens OX=9606 GN=NME3 PE=1 SV=2 | 12 | 4 | 4 | 19 | 28 | 169 | 7.84 | 2 | 2 | | 1 | 1 | 1 | 2 | 2 | | 1 | 1 | 1 |
| High | O75940 | Survival of motor neuron-related-splicing factor 30 OS=Homo sapiens OX=9606 GN=SMNDC1 PE=1 SV=1 | 12 | 3 | 3 | 26.7 | 18 | 238 | 7.24 | | | | 1 | 2 | 2 | | | | 1 | 2 | 2 |
| High | Q9H6R4 | Nucleolar protein 6 OS=Homo sapiens OX=9606 GN=NOL6 PE=1 SV=2 | 12 | 4 | 4 | 127.5 | 5 | 1146 | 7.64 | | | 1 | | 1 | 2 | | | 1 | | 1 | 2 |
| High | P10155 | RNA-binding protein RO60 OS=Homo sapiens OX=9606 GN=RO60 PE=1 SV=2 | 12 | 3 | 3 | 60.6 | 5 | 538 | 8.03 | 1 | 1 | 2 | 1 | 1 | 1 | 1 | 1 | 2 | 1 | 1 | 1 |
| High | P21281 | V-type proton ATPase subunit B, brain isoform OS=Homo sapiens OX=9606 GN=ATP6V1B2 PE=1 SV=3 | 12 | 2 | 2 | 56.5 | 6 | 511 | 5.81 | 2 | 2 | 1 | 1 | 1 | 1 | 2 | 2 | 1 | 1 | 1 | 1 |
| High | P02754 | SWISS-PROT:P02754 Beta-lactoglobulin - Bos taurus (Bovine). | 12 | 5 | 5 | 18.3 | 31 | 162 | 4.92 | | | | 1 | 1 | 1 | | | | 1 | 1 | 1 |
| High | P11498 | Pyruvate carboxylase, mitochondrial OS=Homo sapiens OX=9606 GN=PC PE=1 SV=2 | 12 | 5 | 5 | 129.6 | 6 | 1178 | 6.84 | 1 | 2 | | 1 | | 1 | 1 | 2 | | 1 | | 1 |
| High | Q8N1B4 | Vacuolar protein sorting-associated protein 52 homolog OS=Homo sapiens OX=9606 GN=VPS52 PE=1 SV=1 | 12 | 5 | 5 | 82.2 | 7 | 723 | 5.99 | 4 | 2 | 5 | | | | 3 | 2 | 5 | | | |
| High | O95819 | Mitogen-activated protein kinase kinase kinase kinase 4 OS=Homo sapiens OX=9606 GN=MAP4K4 PE=1 SV=2 | 12 | 5 | 5 | 142 | 5 | 1239 | 7.46 | 1 | | 3 | 1 | 2 | 1 | 1 | | 3 | 1 | 2 | 1 |
| High | Q99436 | Proteasome subunit beta type-7 OS=Homo sapiens OX=9606 GN=PSMB7 PE=1 SV=1 | 12 | 3 | 3 | 29.9 | 10 | 277 | 7.68 | | | | 3 | 3 | 2 | | | | 3 | 3 | 2 |
| High | Q16698 | 2,4-dienoyl-CoA reductase [(3E)-enoyl-CoA-producing], mitochondrial OS=Homo sapiens OX=9606 GN=DECR1 PE=1 SV=1 | 12 | 3 | 3 | 36 | 14 | 335 | 9.28 | 1 | 2 | 2 | 1 | 1 | 1 | 1 | 2 | 2 | 1 | 1 | 1 |
| High | P62491 | Ras-related protein Rab-11A OS=Homo sapiens OX=9606 GN=RAB11A PE=1 SV=3 | 12 | 3 | 2 | 24.4 | 14 | 216 | 6.57 | 1 | 1 | 1 | 1 | 1 | 2 | 1 | 1 | 1 | 1 | 1 | 2 |
| High | Q14739 | Delta(14)-sterol reductase LBR OS=Homo sapiens OX=9606 GN=LBR PE=1 SV=2 | 12 | 4 | 4 | 70.7 | 7 | 615 | 9.36 | 3 | 3 | 2 | 1 | 1 | 1 | 3 | 3 | 2 | 1 | 1 | 1 |
| High | Q92673 | Sortilin-related receptor OS=Homo sapiens OX=9606 GN=SORL1 PE=1 SV=2 | 12 | 6 | 6 | 248.3 | 3 | 2214 | 5.55 | | | 1 | | | | | | 1 | | | |
| High | Q53GT1 | Kelch-like protein 22 OS=Homo sapiens OX=9606 GN=KLHL22 PE=1 SV=2 | 12 | 8 | 8 | 71.6 | 12 | 634 | 5.49 | | | | 2 | 7 | 3 | | | | 2 | 7 | 3 |
| High | O43488 | Aflatoxin B1 aldehyde reductase member 2 OS=Homo sapiens OX=9606 GN=AKR7A2 PE=1 SV=3 | 12 | 4 | 3 | 39.6 | 14 | 359 | 7.17 | | 1 | 2 | 1 | | 2 | | 1 | 2 | 1 | | 2 |
| High | P55209 | Nucleosome assembly protein 1-like 1 OS=Homo sapiens OX=9606 GN=NAP1L1 PE=1 SV=1 | 12 | 4 | 3 | 45.3 | 14 | 391 | 4.46 | 1 | 3 | 2 | 1 | | | 1 | 3 | 2 | 1 | | |
| High | O75175 | CCR4-NOT transcription complex subunit 3 OS=Homo sapiens OX=9606 GN=CNOT3 PE=1 SV=1 | 12 | 3 | 3 | 81.8 | 9 | 753 | 6.2 | 1 | 2 | 1 | 1 | 1 | 1 | 1 | 2 | 1 | 1 | 1 | 1 |
| High | Q969G3 | SWI/SNF-related matrix-associated actin-dependent regulator of chromatin subfamily E member 1 OS=Homo sapiens OX=9606 GI | 12 | 5 | 5 | 46.6 | 15 | 411 | 4.88 | 1 | | | 1 | | | 1 | | | 1 | | |
| High | Q8N122 | Regulatory-associated protein of mTOR OS=Homo sapiens OX=9606 GN=RPTOR PE=1 SV=1 | 12 | 5 | 5 | 148.9 | 5 | 1335 | 6.89 | 2 | 3 | 4 | | | | 2 | 3 | 4 | | | |
| High | P51398 | 28S ribosomal protein S29, mitochondrial OS=Homo sapiens OX=9606 GN=DAP3 PE=1 SV=1 | 12 | 7 | 7 | 45.5 | 22 | 398 | 8.88 | | | | | 1 | 1 | | | | | 1 | 1 |
| High | O15484 | Calpain-5 OS=Homo sapiens OX=9606 GN=CAPN5 PE=1 SV=2 | 12 | 3 | 3 | 73.1 | 7 | 640 | 7.64 | 1 | 1 | 3 | | | | 1 | 1 | 2 | | | |
| High | Q9HAW8 | UDP-glucuronosyltransferase 1A10 OS=Homo sapiens OX=9606 GN=UGT1A10 PE=1 SV=1 | 12 | 4 | 4 | 59.8 | 8 | 530 | 7.3 | 3 | 3 | 2 | | | | 3 | 3 | 2 | | | |
| High | Q9Y657 | Spindlin-1 OS=Homo sapiens OX=9606 GN=SPIN1 PE=1 SV=3 | 12 | 4 | 4 | 29.6 | 26 | 262 | 6.96 | 3 | 4 | 2 | 1 | | 1 | 3 | 4 | 2 | 1 | | 1 |
| High | O00560 | Syntenin-1 OS=Homo sapiens OX=9606 GN=SDCBP PE=1 SV=1 | 12 | 3 | 3 | 32.4 | 22 | 298 | 7.53 | 2 | 2 | 2 | | 1 | | 2 | 2 | 2 | | 1 | |
| High | Q9BRP1 | Programmed cell death protein 2-like OS=Homo sapiens OX=9606 GN=PDCD2L PE=1 SV=1 | 12 | 4 | 4 | 39.4 | 19 | 358 | 4.86 | | | 1 | | 1 | 1 | | | 1 | | 1 | 1 |
| High | Q9BYM8 | RanBP-type and C3HC4-type zinc finger-containing protein 1 OS=Homo sapiens OX=9606 GN=RBCK1 PE=1 SV=2 | 12 | 4 | 4 | 57.5 | 7 | 510 | 5.67 | | | 2 | | 2 | 1 | | | 2 | | 2 | 1 |
| High | Q16563 | Synaptophysin-like protein 1 OS=Homo sapiens OX=9606 GN=SYPL1 PE=1 SV=1 | 12 | 3 | 3 | 28.5 | 20 | 259 | 8.43 | 2 | 2 | 4 | 1 | | 1 | 2 | 2 | 3 | 1 | | 1 |
| High | Q96LR5 | Ubiquitin-conjugating enzyme E2 E2 OS=Homo sapiens OX=9606 GN=UBE2E2 PE=1 SV=1 | 12 | 3 | 3 | 22.2 | 17 | 201 | 7.71 | 2 | 1 | 1 | 2 | 2 | 2 | 2 | 1 | 1 | 2 | 2 | 2 |
| High | Q9ULD2 | Microtubule-associated tumor suppressor 1 OS=Homo sapiens OX=9606 GN=MTUS1 PE=1 SV=2 | 12 | 3 | 3 | 141.3 | 3 | 1270 | 7.5 | 1 | 2 | 1 | | 1 | | 1 | 2 | 1 | | 1 | |
| High | O75396 | Vesicle-trafficking protein SEC22b OS=Homo sapiens OX=9606 GN=SEC22B PE=1 SV=5 | 12 | 3 | 3 | 24.7 | 16 | 215 | 8.51 | 1 | | | 1 | 1 | 1 | 1 | | | 1 | 1 | 1 |
| High | P78346 | Ribonuclease P protein subunit p30 OS=Homo sapiens OX=9606 GN=RPP30 PE=1 SV=1 | 12 | 3 | 3 | 29.3 | 15 | 268 | 8.91 | 2 | 1 | | 2 | 1 | 2 | 2 | 1 | | 2 | 1 | 2 |
| High | Q8WWC4 | m-AAA protease-interacting protein 1, mitochondrial OS=Homo sapiens OX=9606 GN=MAIP1 PE=1 SV=1 | 12 | 5 | 5 | 32.5 | 19 | 291 | 9.17 | 2 | 1 | 5 | | | | 2 | 1 | 5 | | | |
| High | Q9P013 | Spliceosome-associated protein CWC15 homolog OS=Homo sapiens OX=9606 GN=CWC15 PE=1 SV=2 | 12 | 3 | 3 | 26.6 | 19 | 229 | 5.71 | 1 | 2 | 2 | | 2 | 1 | 1 | 2 | 2 | | 2 | 1 |
| High | P84095 | Rho-related GTP-binding protein RhoG OS=Homo sapiens OX=9606 GN=RHOG PE=1 SV=1 | 12 | 3 | 2 | 21.3 | 23 | 191 | 8.12 | 1 | 2 | 1 | 1 | 1 | 1 | 1 | 1 | 1 | 1 | 1 | 1 |
| High | Q8N9U0 | Tandem C2 domains nuclear protein OS=Homo sapiens OX=9606 GN=TC2N PE=1 SV=2 | 12 | 3 | 3 | 55.2 | 7 | 490 | 9.19 | 1 | 1 | | | 2 | 1 | 1 | 1 | | | 2 | 1 |
| High | P84090 | Enhancer of rudimentary homolog OS=Homo sapiens OX=9606 GN=ERH PE=1 SV=1 | 12 | 2 | 2 | 12.3 | 27 | 104 | 5.92 | 1 | 1 | 1 | 1 | 2 | 1 | 1 | 1 | 1 | 1 | 1 | 1 |
| High | Q9ULX9 | Transcription factor MafF OS=Homo sapiens OX=9606 GN=MAFF PE=1 SV=2 | 12 | 3 | 3 | 17.7 | 29 | 164 | 9.8 | 1 | 1 | 1 | 2 | 1 | 1 | 1 | 1 | 1 | 2 | 1 | 1 |
| High | Q9NYK5 | 39S ribosomal protein L39, mitochondrial OS=Homo sapiens OX=9606 GN=MRPL39 PE=1 SV=3 | 12 | 4 | 4 | 38.7 | 12 | 338 | 7.65 | | | | | 2 | 1 | | | | | 2 | 1 |
| High | Q8WUA4 | General transcription factor 3C polypeptide 2 OS=Homo sapiens OX=9606 GN=GTF3C2 PE=1 SV=2 | 12 | 6 | 6 | 100.6 | 8 | 911 | 7.31 | | | | | 2 | 1 | | | | | 2 | 1 |
| High | Q86U90 | YrdC domain-containing protein, mitochondrial OS=Homo sapiens OX=9606 GN=YRDC PE=1 SV=1 | 12 | 4 | 4 | 29.3 | 23 | 279 | 8.57 | | | | 1 | 1 | 1 | | | | 1 | 1 | 1 |
| High | P08123 | Collagen alpha-2(I) chain OS=Homo sapiens OX=9606 GN=COL1A2 PE=1 SV=7 | 12 | 2 | 2 | 129.2 | 1 | 1366 | 8.95 | | | | 5 | 3 | 4 | | | | 2 | 2 | 1 |
| High | P02679 | Fibrinogen gamma chain OS=Homo sapiens OX=9606 GN=FGG PE=1 SV=3 | 12 | 3 | 3 | 51.5 | 8 | 453 | 5.62 | | | | 4 | 4 | 4 | | | | 3 | 3 | 3 |
| High | P45984 | Mitogen-activated protein kinase 9 OS=Homo sapiens OX=9606 GN=MAPK9 PE=1 SV=2 | 12 | 2 | 1 | 48.1 | 4 | 424 | 5.63 | | 1 | 1 | 2 | | 2 | | 1 | 1 | 2 | | 2 |
| High | Q9Y2L9 | Leucine-rich repeat and calponin homology domain-containing protein 1 OS=Homo sapiens OX=9606 GN=LRCH1 PE=1 SV=3 | 12 | 4 | 4 | 80.8 | 10 | 728 | 5.94 | 2 | 2 | 3 | | | | 2 | 2 | 3 | | | |
| High | Q16181 | Septin-7 OS=Homo sapiens OX=9606 GN=SEPTIN7 PE=1 SV=2 | 12 | 4 | 3 | 50.6 | 13 | 437 | 8.63 | 2 | 2 | 2 | 2 | | 1 | 2 | 2 | 2 | 2 | | 1 |
| High | Q9UQN3 | Charged multivesicular body protein 2b OS=Homo sapiens OX=9606 GN=CHMP2B PE=1 SV=1 | 12 | 3 | 2 | 23.9 | 13 | 213 | 8.76 | 1 | 1 | 1 | | | | 1 | 1 | 1 | | | |
| High | P62306 | Small nuclear ribonucleoprotein F OS=Homo sapiens OX=9606 GN=SNRPF PE=1 SV=1 | 12 | 2 | 2 | 9.7 | 24 | 86 | 4.67 | 1 | 1 | 1 | 2 | 2 | 1 | 1 | 1 | 1 | 2 | 2 | 1 |
| High | Q9Y6C9 | Mitochondrial carrier homolog 2 OS=Homo sapiens OX=9606 GN=MTCH2 PE=1 SV=1 | 12 | 5 | 5 | 33.3 | 16 | 303 | 7.97 | 1 | | 1 | | | | 1 | | 1 | | | |
| High | Q8WY36 | HMG box transcription factor BBX OS=Homo sapiens OX=9606 GN=BBX PE=1 SV=1 | 12 | 6 | 6 | 105.1 | 9 | 941 | 8.79 | | 1 | 1 | | 2 | 1 | | 1 | 1 | | 2 | 1 |
| High | Q9NUP9 | Protein lin-7 homolog C OS=Homo sapiens OX=9606 GN=LIN7C PE=1 SV=1 | 12 | 5 | 5 | 21.8 | 25 | 197 | 8.43 | 1 | 1 | 1 | 2 | 1 | 1 | 1 | 1 | 1 | 2 | 1 | 1 |
| High | Q08379 | Golgin subfamily A member 2 OS=Homo sapiens OX=9606 GN=GOLGA2 PE=1 SV=3 | 12 | 4 | 4 | 113 | 6 | 1002 | 5.02 | 1 | | | 1 | 2 | | 1 | | | 1 | 2 | |
| High | Q13616 | Cullin-1 OS=Homo sapiens OX=9606 GN=CUL1 PE=1 SV=2 | 12 | 4 | 4 | 89.6 | 6 | 776 | 8 | 1 | | 1 | 1 | 1 | 1 | 1 | | 1 | 1 | 1 | 1 |
| High | Q15397 | Pumilio homolog 3 OS=Homo sapiens OX=9606 GN=PUM3 PE=1 SV=3 | 12 | 4 | 4 | 73.5 | 5 | 648 | 9.64 | | | | 2 | 3 | | | | | 2 | 3 | |
| High | P10321 | HLA class I histocompatibility antigen, C alpha chain OS=Homo sapiens OX=9606 GN=HLA-C PE=1 SV=3 | 12 | 3 | 2 | 40.6 | 12 | 366 | 6.04 | | | | 1 | 1 | 2 | | | | 1 | 1 | 2 |
| High | Q96P70 | Importin-9 OS=Homo sapiens OX=9606 GN=IPO9 PE=1 SV=3 | 12 | 5 | 5 | 115.9 | 6 | 1041 | 4.81 | 4 | 1 | 4 | | | | 4 | 1 | 4 | | | |
| High | Q9UPN7 | Serine/threonine-protein phosphatase 6 regulatory subunit 1 OS=Homo sapiens OX=9606 GN=PPP6R1 PE=1 SV=5 | 12 | 4 | 4 | 96.7 | 8 | 881 | 4.55 | | | 3 | 1 | | 1 | | | 3 | 1 | | 1 |
| High | O75607 | Nucleoplasmin-3 OS=Homo sapiens OX=9606 GN=NPM3 PE=1 SV=3 | 12 | 3 | 3 | 19.3 | 26 | 178 | 4.63 | 1 | 1 | 1 | | | | 1 | 1 | 1 | | | |
| High | Q9Y3B9 | RRP15-like protein OS=Homo sapiens OX=9606 GN=RRP15 PE=1 SV=2 | 12 | 4 | 4 | 31.5 | 15 | 282 | 5.52 | | 1 | 1 | | | | | 1 | 1 | | | |
| High | P29992 | Guanine nucleotide-binding protein subunit alpha-11 OS=Homo sapiens OX=9606 GN=GNA11 PE=1 SV=2 | 12 | 4 | 1 | 42.1 | 9 | 359 | 5.69 | | 1 | 1 | 3 | 2 | 2 | | 1 | 1 | 3 | 2 | 2 |
| High | Q8TB45 | DEP domain-containing mTOR-interacting protein OS=Homo sapiens OX=9606 GN=DEPTOR PE=1 SV=2 | 12 | 6 | 6 | 46.3 | 19 | 409 | 8.07 | 1 | | | | | | 1 | | | | | |
| High | O15084 | Serine/threonine-protein phosphatase 6 regulatory ankyrin repeat subunit A OS=Homo sapiens OX=9606 GN=ANKRD28 PE=1 SV= | 12 | 2 | 2 | 112.9 | 2 | 1053 | 6.25 | 1 | 1 | 2 | | 3 | | 1 | 1 | 2 | | 2 | |
| High | Q15124 | Phosphoglucomutase-like protein 5 OS=Homo sapiens OX=9606 GN=PGM5 PE=1 SV=2 | 12 | 2 | 2 | 62.2 | 4 | 567 | 7.21 | 1 | 1 | 2 | 1 | 1 | 1 | 1 | 1 | 1 | 1 | 1 | 1 |
| High | Q15293 | Reticulocalbin-1 OS=Homo sapiens OX=9606 GN=RCN1 PE=1 SV=1 | 12 | 5 | 5 | 38.9 | 18 | 331 | 5 | 3 | 4 | 2 | | | | 3 | 4 | 2 | | | |
| High | O95881 | Thioredoxin domain-containing protein 12 OS=Homo sapiens OX=9606 GN=TXNDC12 PE=1 SV=1 | 11 | 3 | 3 | 19.2 | 18 | 172 | 5.4 | | | 1 | 2 | 3 | 1 | | | 1 | 2 | 3 | 1 |
| High | Q9BYG5 | Partitioning defective 6 homolog beta OS=Homo sapiens OX=9606 GN=PARD6B PE=1 SV=1 | 11 | 2 | 2 | 41.2 | 6 | 372 | 5.58 | 1 | 1 | 2 | 1 | 1 | 1 | 1 | 1 | 1 | 1 | 1 | 1 |
| High | P53597 | Succinate--CoA ligase [ADP/GDP-forming] subunit alpha, mitochondrial OS=Homo sapiens OX=9606 GN=SUCLG1 PE=1 SV=4 | 11 | 2 | 2 | 36.2 | 7 | 346 | 8.79 | 1 | 1 | 1 | 2 | 1 | 2 | 1 | 1 | 1 | 2 | 1 | 2 |
| High | Q8ND24 | RING finger protein 214 OS=Homo sapiens OX=9606 GN=RNF214 PE=1 SV=2 | 11 | 4 | 4 | 77.6 | 5 | 703 | 6.95 | 1 | 1 | | | | | 1 | 1 | | | | |
| High | Q15631 | Translin OS=Homo sapiens OX=9606 GN=TSN PE=1 SV=1 | 11 | 2 | 2 | 26.2 | 11 | 228 | 6.44 | 1 | | 2 | 1 | 1 | | 1 | | 2 | 1 | 1 | |
| High | P48556 | 26S proteasome non-ATPase regulatory subunit 8 OS=Homo sapiens OX=9606 GN=PSMD8 PE=1 SV=2 | 11 | 3 | 3 | 39.6 | 7 | 350 | 9.7 | | | | 2 | 2 | 1 | | | | 2 | 2 | 1 |
| High | Q9C0B1 | Alpha-ketoglutarate-dependent dioxygenase FTO OS=Homo sapiens OX=9606 GN=FTO PE=1 SV=3 | 11 | 5 | 5 | 58.2 | 11 | 505 | 5.22 | | 1 | 1 | | | | | 1 | 1 | | | |
| High | O00217 | NADH dehydrogenase [ubiquinone] iron-sulfur protein 8, mitochondrial OS=Homo sapiens OX=9606 GN=NDUFS8 PE=1 SV=1 | 11 | 2 | 2 | 23.7 | 8 | 210 | 6.34 | 2 | 2 | 1 | 1 | 1 | 1 | 2 | 2 | 1 | 1 | 1 | 1 |
| High | Q96T76 | MMS19 nucleotide excision repair protein homolog OS=Homo sapiens OX=9606 GN=MMS19 PE=1 SV=2 | 11 | 3 | 3 | 113.2 | 3 | 1030 | 6.35 | 1 | | 3 | | | | 1 | | 2 | | | |
| High | Q96ME7 | Zinc finger protein 512 OS=Homo sapiens OX=9606 GN=ZNF512 PE=1 SV=2 | 11 | 3 | 3 | 64.6 | 7 | 567 | 9.76 | | | 1 | | 1 | 1 | | | 1 | | 1 | 1 |
| High | P18031 | Tyrosine-protein phosphatase non-receptor type 1 OS=Homo sapiens OX=9606 GN=PTPN1 PE=1 SV=1 | 11 | 4 | 4 | 49.9 | 14 | 435 | 6.27 | 3 | 2 | 1 | | 1 | | 3 | 2 | 1 | | 1 | |
| High | P41227 | N-alpha-acetyltransferase 10 OS=Homo sapiens OX=9606 GN=NAA10 PE=1 SV=1 | 11 | 4 | 4 | 26.4 | 20 | 235 | 5.64 | 1 | 1 | 1 | 1 | 1 | 2 | 1 | 1 | 1 | 1 | 1 | 2 |
| High | Q9NR50 | Translation initiation factor eIF-2B subunit gamma OS=Homo sapiens OX=9606 GN=EIF2B3 PE=1 SV=1 | 11 | 2 | 2 | 50.2 | 9 | 452 | 6.47 | 1 | 1 | 2 | 1 | 1 | | 1 | 1 | 2 | 1 | 1 | |
| High | Q8WTV1 | THAP domain-containing protein 3 OS=Homo sapiens OX=9606 GN=THAP3 PE=1 SV=1 | 11 | 4 | 4 | 27 | 28 | 239 | 10.24 | | | | 1 | 2 | 1 | | | | 1 | 2 | 1 |
| High | Q9Y5Y2 | Cytosolic Fe-S cluster assembly factor NUBP2 OS=Homo sapiens OX=9606 GN=NUBP2 PE=1 SV=1 | 11 | 3 | 3 | 28.8 | 23 | 271 | 5.83 | 2 | 2 | 2 | | 1 | | 2 | 2 | 2 | | 1 | |
| High | Q9NZW5 | Protein PALS2 OS=Homo sapiens OX=9606 GN=PALS2 PE=1 SV=2 | 11 | 3 | 2 | 61.1 | 6 | 540 | 6.18 | 2 | 2 | 1 | | 1 | 1 | 2 | 2 | 1 | | 1 | 1 |
| High | Q5T8P6 | RNA-binding protein 26 OS=Homo sapiens OX=9606 GN=RBM26 PE=1 SV=3 | 11 | 3 | 2 | 113.5 | 3 | 1007 | 9.16 | | | | 1 | 2 | 1 | | | | 1 | 2 | 1 |
| High | Q7RTV0 | PHD finger-like domain-containing protein 5A OS=Homo sapiens OX=9606 GN=PHF5A PE=1 SV=1 | 11 | 3 | 3 | 12.4 | 25 | 110 | 8.41 | | | | 2 | 2 | 1 | | | | 2 | 2 | 1 |
| High | Q96JQ2 | Calmin OS=Homo sapiens OX=9606 GN=CLMN PE=1 SV=1 | 11 | 3 | 3 | 111.6 | 3 | 1002 | 4.94 | 2 | 2 | 3 | | | 1 | 2 | 2 | 3 | | | 1 |
| High | Q8WXE9 | Stonin-2 OS=Homo sapiens OX=9606 GN=STON2 PE=1 SV=1 | 11 | 1 | 1 | 101.1 | 1 | 905 | 5.39 | 2 | 4 | 2 | | | | 1 | 1 | 1 | | | |
| High | P48507 | Glutamate--cysteine ligase regulatory subunit OS=Homo sapiens OX=9606 GN=GCLM PE=1 SV=1 | 11 | 4 | 4 | 30.7 | 19 | 274 | 6.02 | 1 | 1 | 3 | | 1 | 1 | 1 | 1 | 3 | | 1 | 1 |
| High | P31151 | Protein S100-A7 OS=Homo sapiens OX=9606 GN=S100A7 PE=1 SV=4 | 11 | 3 | 3 | 11.5 | 24 | 101 | 6.77 | | | | 3 | 4 | 4 | | | | 3 | 3 | 3 |
| High | P67870 | Casein kinase II subunit beta OS=Homo sapiens OX=9606 GN=CSNK2B PE=1 SV=1 | 11 | 3 | 3 | 24.9 | 20 | 215 | 5.55 | | | 1 | 2 | 2 | 1 | | | 1 | 2 | 2 | 1 |
| High | P29317 | Ephrin type-A receptor 2 OS=Homo sapiens OX=9606 GN=EPHA2 PE=1 SV=2 | 11 | 6 | 4 | 108.2 | 7 | 976 | 6.23 | | 2 | 1 | 1 | 1 | 1 | | 2 | 1 | 1 | 1 | 1 |
| High | Q86VP1 | Tax1-binding protein 1 OS=Homo sapiens OX=9606 GN=TAX1BP1 PE=1 SV=2 | 11 | 5 | 5 | 90.8 | 6 | 789 | 5.43 | 1 | 1 | 1 | | 1 | | 1 | 1 | 1 | | 1 | |
| High | Q9UK59 | Lariat debranching enzyme OS=Homo sapiens OX=9606 GN=DBR1 PE=1 SV=2 | 11 | 3 | 3 | 61.5 | 6 | 544 | 5.47 | 1 | 1 | 1 | 1 | 2 | 1 | 1 | 1 | 1 | 1 | 2 | 1 |
| High | O14745 | Na(+)/H(+) exchange regulatory cofactor NHE-RF1 OS=Homo sapiens OX=9606 GN=SLC9A3R1 PE=1 SV=4 | 11 | 3 | 3 | 38.8 | 11 | 358 | 5.77 | | 1 | 2 | 1 | | 1 | | 1 | 2 | 1 | | 1 |
| High | P49458 | Signal recognition particle 9 kDa protein OS=Homo sapiens OX=9606 GN=SRP9 PE=1 SV=2 | 11 | 2 | 2 | 10.1 | 22 | 86 | 7.97 | 1 | 2 | | 1 | 1 | 2 | 1 | 2 | | 1 | 1 | 2 |
| High | Q96DV4 | 39S ribosomal protein L38, mitochondrial OS=Homo sapiens OX=9606 GN=MRPL38 PE=1 SV=2 | 11 | 3 | 3 | 44.6 | 10 | 380 | 7.53 | 1 | 1 | 2 | 1 | 1 | 1 | 1 | 1 | 2 | 1 | 1 | 1 |
| High | Q9P270 | SLAIN motif-containing protein 2 OS=Homo sapiens OX=9606 GN=SLAIN2 PE=1 SV=2 | 11 | 3 | 3 | 62.5 | 8 | 581 | 9.45 | | 2 | 2 | | 1 | 1 | | 2 | 2 | | 1 | 1 |
| High | P61163 | Alpha-centractin OS=Homo sapiens OX=9606 GN=ACTR1A PE=1 SV=1 | 11 | 3 | 3 | 42.6 | 11 | 376 | 6.64 | 2 | 1 | | | | | 2 | 1 | | | | |
| High | Q01650 | Large neutral amino acids transporter small subunit 1 OS=Homo sapiens OX=9606 GN=SLC7A5 PE=1 SV=2 | 11 | 3 | 3 | 55 | 8 | 507 | 7.72 | 1 | 1 | 1 | 2 | 1 | 1 | 1 | 1 | 1 | 2 | 1 | 1 |

| | | | | | | | | | | | | | | | | | | | | | |
|---|---|---|---|---|---|---|---|---|---|---|---|---|---|---|---|---|---|---|---|---|---|
| High | P54920 | Alpha-soluble NSF attachment protein OS=Homo sapiens OX=9606 GN=NAPA PE=1 SV=3 | 11 | 4 | 4 | 33.2 | 16 | 295 | 5.36 | 1 | 1 | | | 1 | 1 | 1 | 1 | | | 1 | 1 |
| High | Q10570 | Cleavage and polyadenylation specificity factor subunit 1 OS=Homo sapiens OX=9606 GN=CPSF1 PE=1 SV=2 | 11 | 5 | 5 | 160.8 | 4 | 1443 | 6.4 | | 1 | | | 2 | | | 1 | | | 2 | |
| High | Q3ZAQ7 | Vacuolar ATPase assembly integral membrane protein VMA21 OS=Homo sapiens OX=9606 GN=VMA21 PE=1 SV=1 | 11 | 2 | 2 | 11.3 | 22 | 101 | 7.24 | 1 | 1 | 1 | 1 | | 1 | 1 | 1 | 1 | 1 | | 1 |
| High | P47813 | Eukaryotic translation initiation factor 1A, X-chromosomal OS=Homo sapiens OX=9606 GN=EIF1AX PE=1 SV=2 | 11 | 2 | 2 | 16.5 | 15 | 144 | 5.24 | 1 | | 1 | 2 | 2 | 2 | 1 | | 1 | 2 | 2 | 2 |
| High | Q9H2G2 | STE20-like serine/threonine-protein kinase OS=Homo sapiens OX=9606 GN=SLK PE=1 SV=1 | 11 | 4 | 4 | 142.6 | 5 | 1235 | 5.15 | 1 | 1 | 2 | 1 | | 1 | 1 | 1 | 2 | 1 | | 1 |
| High | Q93008 | Probable ubiquitin carboxyl-terminal hydrolase FAF-X OS=Homo sapiens OX=9606 GN=USP9X PE=1 SV=4 | 11 | 7 | 4 | 290.3 | 3 | 2554 | 5.8 | 1 | 2 | 2 | | | | 1 | 2 | 2 | | | |
| High | O60749 | Sorting nexin-2 OS=Homo sapiens OX=9606 GN=SNX2 PE=1 SV=2 | 11 | 4 | 2 | 58.4 | 8 | 519 | 5.12 | 1 | | | | 3 | | 1 | | | | 3 | |
| High | P04732 | Metallothionein-1E OS=Homo sapiens OX=9606 GN=MT1E PE=1 SV=1 | 11 | 2 | 1 | 6 | 52 | 61 | 7.96 | 1 | 1 | 1 | 2 | 2 | 1 | 1 | 1 | 1 | 2 | 2 | 1 |
| High | O14578 | Citron Rho-interacting kinase OS=Homo sapiens OX=9606 GN=CIT PE=1 SV=2 | 11 | 4 | 4 | 231.3 | 2 | 2027 | 6.57 | 1 | 1 | | 1 | 4 | 2 | 1 | 1 | | 1 | 4 | 2 |
| High | Q14596 | Next to BRCA1 gene 1 protein OS=Homo sapiens OX=9606 GN=NBR1 PE=1 SV=3 | 11 | 2 | 2 | 107.3 | 3 | 966 | 5.12 | 1 | 1 | 2 | | | | 1 | 1 | 1 | | | |
| High | Q14008 | Cytoskeleton-associated protein 5 OS=Homo sapiens OX=9606 GN=CKAP5 PE=1 SV=3 | 11 | 3 | 3 | 225.4 | 1 | 2032 | 7.8 | | | 1 | | 2 | 2 | | | 1 | | 2 | 2 |
| High | Q8NFQ8 | Torsin-1A-interacting protein 2 OS=Homo sapiens OX=9606 GN=TOR1AIP2 PE=1 SV=1 | 11 | 3 | 3 | 51.2 | 9 | 470 | 4.96 | 1 | 1 | 1 | 1 | 1 | 1 | 1 | 1 | 1 | 1 | 1 | 1 |
| High | Q99733 | Nucleosome assembly protein 1-like 4 OS=Homo sapiens OX=9606 GN=NAP1L4 PE=1 SV=1 | 11 | 5 | 4 | 42.8 | 17 | 375 | 4.69 | | 2 | 3 | 1 | 1 | | | 2 | 3 | 1 | 1 | |
| High | Q9H9A6 | Leucine-rich repeat-containing protein 40 OS=Homo sapiens OX=9606 GN=LRRC40 PE=1 SV=1 | 11 | 3 | 3 | 68.2 | 5 | 602 | 6.43 | 1 | 1 | 1 | 3 | 2 | 2 | 1 | 1 | 1 | 3 | 2 | 2 |
| High | O00264 | Membrane-associated progesterone receptor component 1 OS=Homo sapiens OX=9606 GN=PGRMC1 PE=1 SV=3 | 11 | 3 | 3 | 21.7 | 16 | 195 | 4.7 | 1 | 1 | 1 | | 1 | 1 | 1 | 1 | 1 | | 1 | 1 |
| High | Q96S19 | Methyltransferase-like 26 OS=Homo sapiens OX=9606 GN=METTL26 PE=1 SV=2 | 11 | 4 | 4 | 22.6 | 24 | 204 | 7.83 | | | 2 | 1 | 2 | 1 | | | 2 | 1 | 2 | 1 |
| High | P98175 | RNA-binding protein 10 OS=Homo sapiens OX=9606 GN=RBM10 PE=1 SV=3 | 11 | 6 | 6 | 103.5 | 7 | 930 | 5.97 | | | | | | | | | | | | |
| High | Q9H0E2 | Toll-interacting protein OS=Homo sapiens OX=9606 GN=TOLLIP PE=1 SV=1 | 11 | 3 | 3 | 30.3 | 15 | 274 | 5.97 | 3 | 3 | 2 | | | | 3 | 3 | 2 | | | |
| High | Q00059 | Transcription factor A, mitochondrial OS=Homo sapiens OX=9606 GN=TFAM PE=1 SV=1 | 11 | 2 | 2 | 29.1 | 7 | 246 | 9.72 | | | 1 | 1 | 1 | 2 | | | 1 | 1 | 1 | 2 |
| High | Q2TAL8 | Transcriptional regulator QRICH1 OS=Homo sapiens OX=9606 GN=QRICH1 PE=1 SV=1 | 11 | 3 | 3 | 86.4 | 3 | 776 | 5.87 | | | | 2 | 1 | 2 | | | | 2 | 1 | 2 |
| High | Q13643 | Four and a half LIM domains protein 3 OS=Homo sapiens OX=9606 GN=FHL3 PE=1 SV=4 | 11 | 5 | 5 | 31.2 | 20 | 280 | 6.2 | | | | 1 | 3 | 1 | | | | 1 | 3 | 1 |
| High | P01100 | Protein c-Fos OS=Homo sapiens OX=9606 GN=FOS PE=1 SV=1 | 11 | 3 | 3 | 40.7 | 11 | 380 | 4.87 | | 1 | | | 2 | | | 1 | | | 2 | |
| High | Q00796 | Sorbitol dehydrogenase OS=Homo sapiens OX=9606 GN=SORD PE=1 SV=4 | 11 | 4 | 4 | 38.3 | 15 | 357 | 7.97 | 1 | 1 | 2 | 1 | | 1 | 1 | 1 | 2 | 1 | | 1 |
| High | Q92667 | A-kinase anchor protein 1, mitochondrial OS=Homo sapiens OX=9606 GN=AKAP1 PE=1 SV=1 | 11 | 3 | 3 | 97.3 | 4 | 903 | 4.94 | 1 | | 1 | 1 | 1 | 1 | 1 | | 1 | 1 | 1 | 1 |
| High | P29144 | Tripeptidyl-peptidase 2 OS=Homo sapiens OX=9606 GN=TPP2 PE=1 SV=4 | 11 | 6 | 6 | 138.3 | 7 | 1249 | 6.32 | | 1 | | | 1 | | | 1 | | | 1 | |
| High | Q8N3C0 | Activating signal cointegrator 1 complex subunit 3 OS=Homo sapiens OX=9606 GN=ASCC3 PE=1 SV=3 | 11 | 5 | 5 | 251.3 | 3 | 2202 | 7.09 | 2 | 3 | 1 | | | | 2 | 3 | 1 | | | |
| High | P19388 | DNA-directed RNA polymerases I, II, and III subunit RPABC1 OS=Homo sapiens OX=9606 GN=POLR2E PE=1 SV=4 | 11 | 4 | 4 | 24.5 | 23 | 210 | 5.95 | | | 1 | 1 | | 2 | | | 1 | 1 | | 2 |
| High | O95837 | Guanine nucleotide-binding protein subunit alpha-14 OS=Homo sapiens OX=9606 GN=GNA14 PE=1 SV=1 | 11 | 4 | 1 | 41.5 | 9 | 355 | 6.07 | | 1 | 1 | 3 | 1 | 2 | | 1 | 1 | 3 | 1 | 2 |
| High | P78406 | mRNA export factor RAE1 OS=Homo sapiens OX=9606 GN=RAE1 PE=1 SV=1 | 11 | 6 | 6 | 40.9 | 24 | 368 | 7.83 | 1 | | | | 1 | | 1 | | | | 1 | |
| High | Q9HBL0 | Tensin-1 OS=Homo sapiens OX=9606 GN=TNS1 PE=1 SV=2 | 11 | 3 | 2 | 185.6 | 2 | 1735 | 7.75 | 1 | 2 | 1 | 1 | 2 | 1 | 1 | 1 | 1 | 1 | 2 | 1 |
| High | Q659A1 | Little elongation complex subunit 2 OS=Homo sapiens OX=9606 GN=ICE2 PE=1 SV=2 | 11 | 5 | 5 | 109.9 | 5 | 982 | 7.09 | | | | 3 | 3 | 3 | | | | 3 | 3 | 3 |
| High | Q9H4A3 | Serine/threonine-protein kinase WNK1 OS=Homo sapiens OX=9606 GN=WNK1 PE=1 SV=2 | 11 | 5 | 5 | 250.6 | 2 | 2382 | 6.34 | | | 1 | | 1 | 1 | | | 1 | | 1 | 1 |
| High | Q9NPG3 | Ubinuclein-1 OS=Homo sapiens OX=9606 GN=UBN1 PE=1 SV=2 | 11 | 3 | 3 | 121.4 | 3 | 1134 | 9.33 | 2 | 2 | 1 | 1 | 1 | 1 | 2 | 2 | 1 | 1 | 1 | 1 |
| High | Q9H6Z4 | Ran-binding protein 3 OS=Homo sapiens OX=9606 GN=RANBP3 PE=1 SV=1 | 11 | 3 | 3 | 60.2 | 7 | 567 | 4.78 | 1 | 1 | 1 | 1 | 1 | 1 | 1 | 1 | 1 | 1 | 1 | 1 |
| High | Q9NYL2 | Mitogen-activated protein kinase kinase kinase 20 OS=Homo sapiens OX=9606 GN=MAP3K20 PE=1 SV=3 | 11 | 4 | 4 | 91.1 | 5 | 800 | 7.87 | 2 | 1 | 1 | 1 | 1 | 1 | 2 | 1 | 1 | 1 | 1 | 1 |
| High | Q16186 | Proteasomal ubiquitin receptor ADRM1 OS=Homo sapiens OX=9606 GN=ADRM1 PE=1 SV=2 | 11 | 3 | 3 | 42.1 | 11 | 407 | 5.07 | | | 1 | 2 | 1 | 1 | | | 1 | 2 | 1 | 1 |
| High | Q9UKN8 | General transcription factor 3C polypeptide 4 OS=Homo sapiens OX=9606 GN=GTF3C4 PE=1 SV=2 | 11 | 4 | 4 | 91.9 | 6 | 822 | 6.65 | 1 | 1 | 1 | | 1 | | 1 | 1 | 1 | | 1 | |
| High | O75494 | Serine/arginine-rich splicing factor 10 OS=Homo sapiens OX=9606 GN=SRSF10 PE=1 SV=1 | 11 | 4 | 4 | 31.3 | 16 | 262 | 11.27 | 2 | 1 | 1 | 1 | | 1 | 2 | 1 | 1 | 1 | | 1 |
| High | Q8TBB5 | Kelch domain-containing protein 4 OS=Homo sapiens OX=9606 GN=KLHDC4 PE=1 SV=1 | 11 | 2 | 2 | 57.9 | 4 | 520 | 5.72 | | 1 | 1 | 1 | 1 | 1 | | 1 | 1 | 1 | 1 | 1 |
| High | P01111 | GTPase NRas OS=Homo sapiens OX=9606 GN=NRAS PE=1 SV=1 | 11 | 3 | 3 | 21.2 | 17 | 189 | 5.17 | 1 | 1 | 3 | | 1 | | 1 | 1 | 2 | | 1 | |
| High | P30566 | Adenylosuccinate lyase OS=Homo sapiens OX=9606 GN=ADSL PE=1 SV=2 | 11 | 4 | 4 | 54.9 | 13 | 484 | 7.11 | | | 2 | 1 | 1 | 1 | | | 2 | 1 | 1 | 1 |
| High | Q9H307 | Pinin OS=Homo sapiens OX=9606 GN=PNN PE=1 SV=5 | 11 | 3 | 3 | 81.6 | 4 | 717 | 7.14 | 1 | | 1 | 1 | 3 | 2 | 1 | | 1 | 1 | 3 | 2 |
| High | P02676 | SWISS-PROT:P02676 (Bos taurus) similar to Fibrinogen beta chain precursor | 11 | 4 | 1 | 56.4 | 8 | 495 | 8.22 | | | | 4 | 3 | 4 | | | | 4 | 3 | 4 |
| High | Q99613 | Eukaryotic translation initiation factor 3 subunit C OS=Homo sapiens OX=9606 GN=EIF3C PE=1 SV=1 | 11 | 3 | 3 | 105.3 | 4 | 913 | 5.68 | 2 | 2 | 1 | | | | 2 | 2 | 1 | | | |
| High | Q9BPW8 | Protein NipSnap homolog 1 OS=Homo sapiens OX=9606 GN=NIPSNAP1 PE=1 SV=1 | 11 | 2 | 2 | 33.3 | 7 | 284 | 9.31 | 2 | 1 | 1 | 1 | 1 | 1 | 2 | 1 | 1 | 1 | 1 | 1 |
| High | P31350 | Ribonucleoside-diphosphate reductase subunit M2 OS=Homo sapiens OX=9606 GN=RRM2 PE=1 SV=1 | 11 | 6 | 6 | 44.8 | 22 | 389 | 5.38 | 1 | 2 | 1 | | 1 | 1 | 1 | 2 | 1 | | 1 | 1 |
| High | O94929 | Actin-binding LIM protein 3 OS=Homo sapiens OX=9606 GN=ABLIM3 PE=1 SV=3 | 11 | 5 | 5 | 77.8 | 10 | 683 | 8.54 | 3 | 2 | 3 | | | | 3 | 2 | 3 | | | |
| High | Q9BZL4 | Protein phosphatase 1 regulatory subunit 12C OS=Homo sapiens OX=9606 GN=PPP1R12C PE=1 SV=1 | 11 | 4 | 4 | 84.8 | 10 | 782 | 5.57 | 1 | 2 | 3 | | | | 1 | 2 | 3 | | | |
| High | Q9NVI1 | Fanconi anemia group I protein OS=Homo sapiens OX=9606 GN=FANCI PE=1 SV=4 | 10 | 3 | 3 | 149.2 | 3 | 1328 | 6.74 | 1 | 1 | 1 | | 1 | | 1 | 1 | 1 | | 1 | |
| High | O95251 | Histone acetyltransferase KAT7 OS=Homo sapiens OX=9606 GN=KAT7 PE=1 SV=1 | 10 | 3 | 2 | 70.6 | 6 | 611 | 8.85 | | 1 | | 1 | 1 | 1 | | 1 | | 1 | 1 | 1 |
| High | Q9UQ88 | Cyclin-dependent kinase 11A OS=Homo sapiens OX=9606 GN=CDK11A PE=1 SV=4 | 10 | 5 | 4 | 91.3 | 7 | 783 | 5.36 | | | | | 4 | 1 | | | | | 4 | 1 |
| High | Q9NPE3 | H/ACA ribonucleoprotein complex subunit 3 OS=Homo sapiens OX=9606 GN=NOP10 PE=1 SV=1 | 10 | 2 | 2 | 7.7 | 44 | 64 | 9.99 | | | 2 | 1 | 1 | 1 | | | 2 | 1 | 1 | 1 |
| High | O75165 | DnaJ homolog subfamily C member 13 OS=Homo sapiens OX=9606 GN=DNAJC13 PE=1 SV=5 | 10 | 6 | 6 | 254.3 | 4 | 2243 | 6.74 | 4 | 2 | 2 | | | | 2 | 2 | 2 | | | |
| High | O60343 | TBC1 domain family member 4 OS=Homo sapiens OX=9606 GN=TBC1D4 PE=1 SV=2 | 10 | 6 | 6 | 146.5 | 6 | 1298 | 7.01 | 1 | | | | | | 1 | | | | | |
| High | P98172 | Ephrin-B1 OS=Homo sapiens OX=9606 GN=EFNB1 PE=1 SV=1 | 10 | 2 | 2 | 38 | 9 | 346 | 8.94 | 1 | 1 | 1 | 2 | | 2 | 1 | 1 | 1 | 2 | | 2 |
| High | Q9H8H2 | Probable ATP-dependent RNA helicase DDX31 OS=Homo sapiens OX=9606 GN=DDX31 PE=1 SV=2 | 10 | 5 | 5 | 94 | 6 | 851 | 9.99 | | | | | | | | | | | | |
| High | Q9UNH7 | Sorting nexin-6 OS=Homo sapiens OX=9606 GN=SNX6 PE=1 SV=1 | 10 | 3 | 3 | 46.6 | 9 | 406 | 6.16 | | | 2 | 2 | 3 | 1 | | | 2 | 1 | 2 | 1 |
| High | P16333 | Cytoplasmic protein NCK1 OS=Homo sapiens OX=9606 GN=NCK1 PE=1 SV=1 | 10 | 3 | 3 | 42.8 | 13 | 377 | 6.47 | 3 | 3 | 3 | | | | 3 | 3 | 3 | | | |
| High | P62273 | 40S ribosomal protein S29 OS=Homo sapiens OX=9606 GN=RPS29 PE=1 SV=2 | 10 | 1 | 1 | 6.7 | 13 | 56 | 10.13 | 1 | 1 | 1 | 2 | 1 | 1 | 1 | 1 | 1 | 1 | 1 | 1 |
| High | Q06265 | Exosome complex component RRP45 OS=Homo sapiens OX=9606 GN=EXOSC9 PE=1 SV=3 | 10 | 2 | 2 | 48.9 | 5 | 439 | 5.29 | | | | 2 | 2 | 1 | | | | 2 | 2 | 1 |
| High | P01859 | Immunoglobulin heavy constant gamma 2 OS=Homo sapiens OX=9606 GN=IGHG2 PE=1 SV=2 | 10 | 2 | 1 | 35.9 | 6 | 326 | 7.59 | | 1 | | 2 | 2 | 2 | | 1 | | 2 | 2 | 2 |
| High | Q04760 | Lactoylglutathione lyase OS=Homo sapiens OX=9606 GN=GLO1 PE=1 SV=4 | 10 | 4 | 4 | 20.8 | 23 | 184 | 5.31 | 2 | 3 | 1 | | | | 2 | 3 | 1 | | | |
| High | O43818 | U3 small nucleolar RNA-interacting protein 2 OS=Homo sapiens OX=9606 GN=RRP9 PE=1 SV=1 | 10 | 2 | 2 | 51.8 | 4 | 475 | 7.85 | | | | 1 | 1 | 2 | | | | 1 | 1 | 2 |
| High | O75376 | Nuclear receptor corepressor 1 OS=Homo sapiens OX=9606 GN=NCOR1 PE=1 SV=2 | 10 | 6 | 6 | 270 | 4 | 2440 | 7.11 | | | | | 1 | | | | | | 1 | |
| High | Q9Y6Q5 | AP-1 complex subunit mu-2 OS=Homo sapiens OX=9606 GN=AP1M2 PE=1 SV=4 | 10 | 2 | 2 | 48.1 | 4 | 423 | 8.22 | 2 | 2 | 2 | | | | 2 | 2 | 2 | | | |
| High | Q9Y6K9 | NF-kappa-B essential modulator OS=Homo sapiens OX=9606 GN=IKBKG PE=1 SV=2 | 10 | 2 | 2 | 48.2 | 7 | 419 | 5.71 | 2 | 1 | 2 | | 1 | | 2 | 1 | 2 | | 1 | |
| High | P53582 | Methionine aminopeptidase 1 OS=Homo sapiens OX=9606 GN=METAP1 PE=1 SV=2 | 10 | 2 | 2 | 43.2 | 5 | 386 | 7.17 | 1 | 1 | 1 | | 1 | 1 | 1 | 1 | 1 | | 1 | 1 |
| High | Q13356 | RING-type E3 ubiquitin-protein ligase PPIL2 OS=Homo sapiens OX=9606 GN=PPIL2 PE=1 SV=1 | 10 | 5 | 5 | 58.8 | 12 | 520 | 8.78 | 1 | 1 | 1 | 1 | 1 | 1 | 1 | 1 | 1 | 1 | 1 | 1 |
| High | Q9H2U1 | ATP-dependent DNA/RNA helicase DHX36 OS=Homo sapiens OX=9606 GN=DHX36 PE=1 SV=2 | 10 | 4 | 4 | 114.7 | 5 | 1008 | 7.68 | | 1 | 2 | | 1 | 1 | | 1 | 2 | | 1 | 1 |
| High | Q9Y6M4 | Casein kinase I isoform gamma-3 OS=Homo sapiens OX=9606 GN=CSNK1G3 PE=1 SV=2 | 10 | 3 | 1 | 51.4 | 6 | 447 | 9.22 | 2 | 2 | 3 | | | | 1 | 1 | 2 | | | |
| High | Q96S97 | Myeloid-associated differentiation marker OS=Homo sapiens OX=9606 GN=MYADM PE=1 SV=2 | 10 | 2 | 2 | 35.3 | 10 | 322 | 8.15 | 1 | 1 | 1 | | | 1 | 1 | 1 | 1 | | | 1 |
| High | Q86W42 | THO complex subunit 6 homolog OS=Homo sapiens OX=9606 GN=THOC6 PE=1 SV=1 | 10 | 5 | 5 | 37.5 | 18 | 341 | 7.43 | | | | 1 | 1 | | | | | 1 | 1 | |
| High | Q9UKX7 | Nuclear pore complex protein Nup50 OS=Homo sapiens OX=9606 GN=NUP50 PE=1 SV=2 | 10 | 4 | 4 | 50.1 | 15 | 468 | 7.06 | | | 1 | 1 | | | | | 1 | 1 | | |
| High | P60228 | Eukaryotic translation initiation factor 3 subunit E OS=Homo sapiens OX=9606 GN=EIF3E PE=1 SV=1 | 10 | 5 | 5 | 52.2 | 15 | 445 | 6.04 | 2 | 1 | 2 | | | | 2 | 1 | 2 | | | |
| High | Q9NPH0 | Lysophosphatidic acid phosphatase type 6 OS=Homo sapiens OX=9606 GN=ACP6 PE=1 SV=3 | 10 | 4 | 4 | 48.9 | 12 | 428 | 6.47 | | | | | 1 | | | | | | 1 | |
| High | P49736 | DNA replication licensing factor MCM2 OS=Homo sapiens OX=9606 GN=MCM2 PE=1 SV=4 | 10 | 2 | 2 | 101.8 | 3 | 904 | 5.52 | 1 | 2 | 1 | 1 | 1 | | 1 | 2 | 1 | 1 | 1 | |
| High | Q6DD87 | Zinc finger protein 787 OS=Homo sapiens OX=9606 GN=ZNF787 PE=1 SV=4 | 10 | 4 | 4 | 40.4 | 15 | 382 | 7.96 | | 1 | 1 | | 1 | 1 | | 1 | 1 | | 1 | 1 |
| High | Q9BQ61 | Telomerase RNA component interacting RNase OS=Homo sapiens OX=9606 GN=TRIR PE=1 SV=1 | 10 | 2 | 2 | 18.4 | 12 | 176 | 9.44 | 1 | 1 | 1 | 1 | 2 | 1 | 1 | 1 | 1 | 1 | 2 | 1 |
| High | P27216 | Annexin A13 OS=Homo sapiens OX=9606 GN=ANXA13 PE=1 SV=3 | 10 | 4 | 3 | 35.4 | 15 | 316 | 5.6 | 1 | 1 | 1 | 1 | | | 1 | 1 | 1 | 1 | | |
| High | P27448 | MAP/microtubule affinity-regulating kinase 3 OS=Homo sapiens OX=9606 GN=MARK3 PE=1 SV=5 | 10 | 4 | 1 | 84.4 | 6 | 753 | 9.57 | 2 | 1 | 1 | | | | 2 | 1 | 1 | | | |
| High | Q7Z3C6 | Autophagy-related protein 9A OS=Homo sapiens OX=9606 GN=ATG9A PE=1 SV=3 | 10 | 3 | 3 | 94.4 | 5 | 839 | 6.67 | 3 | 3 | 1 | | | | 3 | 3 | 1 | | | |
| High | Q9NTI5 | Sister chromatid cohesion protein PDS5 homolog B OS=Homo sapiens OX=9606 GN=PDS5B PE=1 SV=1 | 10 | 4 | 4 | 164.6 | 3 | 1447 | 8.47 | 2 | 2 | 2 | | | | 2 | 2 | 1 | | | |
| High | Q9Y512 | Sorting and assembly machinery component 50 homolog OS=Homo sapiens OX=9606 GN=SAMM50 PE=1 SV=3 | 10 | 5 | 5 | 51.9 | 18 | 469 | 6.9 | 1 | 1 | | | 1 | | 1 | 1 | | | 1 | |
| High | A1L0T0 | 2-hydroxyacyl-CoA lyase 2 OS=Homo sapiens OX=9606 GN=ILVBL PE=1 SV=2 | 10 | 6 | 6 | 67.8 | 19 | 632 | 8.15 | 1 | 1 | 2 | | | | 1 | 1 | 2 | | | |
| High | O75489 | NADH dehydrogenase [ubiquinone] iron-sulfur protein 3, mitochondrial OS=Homo sapiens OX=9606 GN=NDUFS3 PE=1 SV=1 | 10 | 4 | 4 | 30.2 | 20 | 264 | 7.5 | 2 | 2 | 2 | | 1 | 1 | 2 | 2 | 2 | | 1 | 1 |
| High | P52815 | 39S ribosomal protein L12, mitochondrial OS=Homo sapiens OX=9606 GN=MRPL12 PE=1 SV=2 | 10 | 2 | 2 | 21.3 | 11 | 198 | 8.87 | | | | 1 | 2 | 1 | | | | 1 | 2 | 1 |
| High | P61966 | AP-1 complex subunit sigma-1A OS=Homo sapiens OX=9606 GN=AP1S1 PE=1 SV=1 | 10 | 3 | 3 | 18.7 | 20 | 158 | 5.73 | 1 | 1 | 1 | 1 | 1 | 1 | 1 | 1 | 1 | 1 | 1 | 1 |
| High | Q15025 | TNFAIP3-interacting protein 1 OS=Homo sapiens OX=9606 GN=TNIP1 PE=1 SV=2 | 10 | 4 | 4 | 71.8 | 9 | 636 | 6.62 | 1 | 1 | 2 | | | | 1 | 1 | 2 | | | |
| High | Q8IX12 | Cell division cycle and apoptosis regulator protein 1 OS=Homo sapiens OX=9606 GN=CCAR1 PE=1 SV=2 | 10 | 2 | 2 | 132.7 | 1 | 1150 | 5.76 | 1 | 1 | 1 | | 2 | | 1 | 1 | 1 | | 2 | |
| High | Q9BYN8 | 28S ribosomal protein S26, mitochondrial OS=Homo sapiens OX=9606 GN=MRPS26 PE=1 SV=1 | 10 | 5 | 5 | 24.2 | 22 | 205 | 10.39 | | | | | | 1 | | | | | | 1 |
| High | Q96CP2 | FLYWCH family member 2 OS=Homo sapiens OX=9606 GN=FLYWCH2 PE=1 SV=1 | 10 | 2 | 2 | 14.6 | 16 | 140 | 8.46 | 2 | 1 | 2 | 1 | 1 | | 2 | 1 | 2 | 1 | 1 | |
| High | Q6IA17 | Single Ig IL-1-related receptor OS=Homo sapiens OX=9606 GN=SIGIRR PE=1 SV=3 | 10 | 4 | 4 | 45.7 | 14 | 410 | 6.46 | | 1 | 1 | | | | | 1 | 1 | | | |
| High | Q13011 | Delta(3,5)-Delta(2,4)-dienoyl-CoA isomerase, mitochondrial OS=Homo sapiens OX=9606 GN=ECH1 PE=1 SV=2 | 10 | 3 | 3 | 35.8 | 11 | 328 | 8 | 1 | 1 | 3 | 1 | | 1 | 1 | 1 | 3 | 1 | | 1 |
| High | Q9UL25 | Ras-related protein Rab-21 OS=Homo sapiens OX=9606 GN=RAB21 PE=1 SV=3 | 10 | 2 | 2 | 24.3 | 9 | 225 | 7.94 | | 1 | 2 | | 1 | 1 | | 1 | 2 | | 1 | 1 |
| High | Q15019 | Septin-2 OS=Homo sapiens OX=9606 GN=SEPTIN2 PE=1 SV=1 | 10 | 4 | 4 | 41.5 | 11 | 361 | 6.6 | 3 | 2 | 2 | | | | 3 | 2 | 2 | | | |
| High | Q9BVJ7 | Dual specificity protein phosphatase 23 OS=Homo sapiens OX=9606 GN=DUSP23 PE=1 SV=1 | 10 | 3 | 3 | 16.6 | 26 | 150 | 8.21 | | | | 1 | 1 | 1 | | | | 1 | 1 | 1 |
| High | P49674 | Casein kinase I isoform epsilon OS=Homo sapiens OX=9606 GN=CSNK1E PE=1 SV=1 | 10 | 3 | 3 | 47.3 | 9 | 416 | 9.66 | | | 1 | | 1 | 1 | | | 1 | | 1 | 1 |
| High | Q13206 | Probable ATP-dependent RNA helicase DDX10 OS=Homo sapiens OX=9606 GN=DDX10 PE=1 SV=2 | 10 | 1 | 1 | 100.8 | 2 | 875 | 8.63 | 1 | 1 | 1 | 1 | | 1 | 1 | 1 | 1 | 1 | | 1 |
| High | Q9Y3P9 | Rab GTPase-activating protein 1 OS=Homo sapiens OX=9606 GN=RABGAP1 PE=1 SV=3 | 10 | 3 | 3 | 121.7 | 4 | 1069 | 5.25 | 1 | 1 | 1 | | 1 | | 1 | 1 | 1 | | 1 | |
| High | A0MZ66 | Shootin-1 OS=Homo sapiens OX=9606 GN=SHTN1 PE=1 SV=4 | 10 | 3 | 3 | 71.6 | 6 | 631 | 5.33 | 1 | 1 | 2 | | 1 | 2 | 1 | 1 | 2 | | 1 | 2 |
| High | Q9Y6D6 | Brefeldin A-inhibited guanine nucleotide-exchange protein 1 OS=Homo sapiens OX=9606 GN=ARFGEF1 PE=1 SV=2 | 10 | 3 | 2 | 208.6 | 1 | 1849 | 5.85 | 1 | 1 | 2 | | | | 1 | 1 | 2 | | | |
| High | O76031 | ATP-dependent Clp protease ATP-binding subunit clpX-like, mitochondrial OS=Homo sapiens OX=9606 GN=CLPX PE=1 SV=2 | 10 | 7 | 7 | 69.2 | 13 | 633 | 7.58 | 1 | 4 | 3 | | | | 1 | 4 | 3 | | | |
| High | Q9H0U6 | 39S ribosomal protein L18, mitochondrial OS=Homo sapiens OX=9606 GN=MRPL18 PE=1 SV=1 | 10 | 2 | 2 | 20.6 | 9 | 180 | 9.54 | 2 | 1 | 2 | | 1 | | 2 | 1 | 2 | | 1 | |
| High | O60684 | Importin subunit alpha-7 OS=Homo sapiens OX=9606 GN=KPNA6 PE=1 SV=1 | 10 | 4 | 1 | 60 | 10 | 536 | 4.98 | 1 | 3 | | | 1 | | 1 | 3 | | | 1 | |
| High | Q86Y56 | Dynein axonemal assembly factor 5 OS=Homo sapiens OX=9606 GN=DNAAF5 PE=1 SV=4 | 10 | 5 | 5 | 93.5 | 9 | 855 | 6.42 | 1 | 2 | 1 | | | | 1 | 2 | 1 | | | |
| High | Q6P1X5 | Transcription initiation factor TFIID subunit 2 OS=Homo sapiens OX=9606 GN=TAF2 PE=1 SV=3 | 10 | 3 | 3 | 136.9 | 4 | 1199 | 8.19 | 1 | 1 | 2 | | | | 1 | 1 | 2 | | | |
| High | Q9Y3C6 | Peptidyl-prolyl cis-trans isomerase-like 1 OS=Homo sapiens OX=9606 GN=PPIL1 PE=1 SV=1 | 10 | 2 | 2 | 18.2 | 19 | 166 | 7.99 | 1 | 1 | 1 | 1 | 1 | 1 | 1 | 1 | 1 | 1 | 1 | 1 |
| High | Q9P267 | Methyl-CpG-binding domain protein 5 OS=Homo sapiens OX=9606 GN=MBD5 PE=1 SV=3 | 10 | 3 | 3 | 159.8 | 4 | 1494 | 8.98 | | | | 1 | 1 | | | | | 1 | 1 | |
| High | Q6PD62 | RNA polymerase-associated protein CTR9 homolog OS=Homo sapiens OX=9606 GN=CTR9 PE=1 SV=1 | 10 | 3 | 3 | 133.4 | 3 | 1173 | 6.77 | | | | | 1 | 1 | | | | | 1 | 1 |
| High | Q92665 | 28S ribosomal protein S31, mitochondrial OS=Homo sapiens OX=9606 GN=MRPS31 PE=1 SV=3 | 10 | 3 | 3 | 45.3 | 7 | 395 | 9.29 | | | 1 | 2 | 1 | 2 | | | 1 | 2 | 1 | 2 |
| High | Q16539 | Mitogen-activated protein kinase 14 OS=Homo sapiens OX=9606 GN=MAPK14 PE=1 SV=3 | 10 | 2 | 1 | 41.3 | 4 | 360 | 5.78 | | 1 | 1 | 1 | | 2 | | 1 | 1 | 1 | | 2 |
| High | Q6P161 | 39S ribosomal protein L54, mitochondrial OS=Homo sapiens OX=9606 GN=MRPL54 PE=1 SV=1 | 10 | 1 | 1 | 15.8 | 7 | 138 | 9.6 | | 1 | 1 | 1 | 1 | 1 | | 1 | 1 | 1 | 1 | 1 |
| High | P17480 | Nucleolar transcription factor 1 OS=Homo sapiens OX=9606 GN=UBTF PE=1 SV=1 | 10 | 6 | 6 | 89.4 | 10 | 764 | 5.81 | 2 | 2 | 1 | | | 1 | 2 | 2 | 1 | | | 1 |
| High | P51798 | H(+)/Cl(-) exchange transporter 7 OS=Homo sapiens OX=9606 GN=CLCN7 PE=1 SV=2 | 10 | 3 | 3 | 88.6 | 3 | 805 | 8.43 | 3 | 3 | 1 | 1 | | | 3 | 3 | 1 | 1 | | |
| High | Q92615 | La-related protein 4B OS=Homo sapiens OX=9606 GN=LARP4B PE=1 SV=3 | 10 | 6 | 6 | 80.5 | 9 | 738 | 6.92 | 1 | 1 | 1 | | 1 | | 1 | 1 | 1 | | 1 | |
| High | Q5T9L3 | Protein wntless homolog OS=Homo sapiens OX=9606 GN=WLS PE=1 SV=2 | 10 | 3 | 3 | 62.2 | 7 | 541 | 7.36 | 2 | 2 | 2 | | | | 2 | 2 | 2 | | | |
| High | Q8NI36 | WD repeat-containing protein 36 OS=Homo sapiens OX=9606 GN=WDR36 PE=1 SV=1 | 10 | 6 | 6 | 105.3 | 8 | 951 | 7.53 | | | 1 | | 2 | | | | 1 | | 2 | |
| High | Q9Y5X3 | Sorting nexin-5 OS=Homo sapiens OX=9606 GN=SNX5 PE=1 SV=1 | 10 | 2 | 2 | 46.8 | 4 | 404 | 6.76 | 2 | 1 | 1 | 1 | 1 | 1 | 2 | 1 | 1 | 1 | 1 | 1 |
| High | Q9HCP0 | Casein kinase I isoform gamma-1 OS=Homo sapiens OX=9606 GN=CSNK1G1 PE=1 SV=1 | 10 | 3 | 1 | 48.5 | 9 | 422 | 9.01 | 2 | 2 | 2 | | | | 1 | 1 | 1 | | | |
| High | P02795 | Metallothionein-2 OS=Homo sapiens OX=9606 GN=MT2A PE=1 SV=1 | 10 | 2 | 1 | 6 | 52 | 61 | 7.83 | 1 | 1 | 1 | 1 | 2 | 1 | 1 | 1 | 1 | 1 | 2 | 1 |
| High | P12270 | Nucleoprotein TPR OS=Homo sapiens OX=9606 GN=TPR PE=1 SV=3 | 10 | 3 | 3 | 267.1 | 2 | 2363 | 5.02 | 1 | 1 | 1 | | | | 1 | 1 | 1 | | | |
| High | P53680 | AP-2 complex subunit sigma OS=Homo sapiens OX=9606 GN=AP2S1 PE=1 SV=2 | 10 | 2 | 2 | 17 | 11 | 142 | 6.18 | 2 | 1 | 1 | 1 | 1 | 1 | 2 | 1 | 1 | 1 | 1 | 1 |
| High | Q13907 | Isopentenyl-diphosphate Delta-isomerase 1 OS=Homo sapiens OX=9606 GN=IDI1 PE=1 SV=2 | 10 | 3 | 3 | 26.3 | 18 | 227 | 6.34 | 2 | 3 | 3 | | | | 2 | 3 | 3 | | | |
| High | P04040 | Catalase OS=Homo sapiens OX=9606 GN=CAT PE=1 SV=3 | 10 | 4 | 4 | 59.7 | 10 | 527 | 7.39 | 2 | 2 | 2 | | 1 | | 2 | 2 | 2 | | 1 | |
| High | O00192 | Armadillo repeat protein deleted in velo-cardio-facial syndrome OS=Homo sapiens OX=9606 GN=ARVCF PE=1 SV=1 | 10 | 3 | 3 | 104.6 | 6 | 962 | 6.81 | 2 | 2 | 2 | | | | 2 | 2 | 2 | | | |
| High | Q96CF2 | Charged multivesicular body protein 4c OS=Homo sapiens OX=9606 GN=CHMP4C PE=1 SV=1 | 10 | 2 | 2 | 26.4 | 11 | 233 | 6.07 | 1 | 2 | 1 | | | | 1 | 1 | 1 | | | |
| High | Q05209 | Tyrosine-protein phosphatase non-receptor type 12 OS=Homo sapiens OX=9606 GN=PTPN12 PE=1 SV=3 | 10 | 7 | 7 | 88.1 | 12 | 780 | 5.62 | 1 | 1 | 1 | | 2 | 2 | 1 | 1 | 1 | | 2 | 2 |
| High | Q9Y4G6 | Talin-2 OS=Homo sapiens OX=9606 GN=TLN2 PE=1 SV=4 | 9 | 3 | 1 | 271.4 | 1 | 2542 | 5.57 | 1 | | 1 | | 1 | | 1 | | 1 | | 1 | |
| High | Q6ZMG9 | Ceramide synthase 6 OS=Homo sapiens OX=9606 GN=CERS6 PE=1 SV=1 | 9 | 1 | 1 | 44.9 | 2 | 384 | 7.65 | 1 | 1 | 1 | 1 | 1 | 1 | 1 | 1 | 1 | 1 | 1 | 1 |
| High | Q92567 | Protein FAM168A OS=Homo sapiens OX=9606 GN=FAM168A PE=1 SV=2 | 9 | 2 | 2 | 26.2 | 7 | 244 | 8.98 | 1 | 1 | 1 | 1 | 1 | 2 | 1 | 1 | 1 | 1 | 1 | 2 |
| High | P80297 | Metallothionein-1X OS=Homo sapiens OX=9606 GN=MT1X PE=1 SV=1 | 9 | 2 | 1 | 6.1 | 52 | 61 | 7.96 | 1 | 1 | 1 | 2 | 1 | | 1 | 1 | 1 | 2 | 1 | |
| High | Q9BVM4 | Gamma-glutamylaminecyclotransferase OS=Homo sapiens OX=9606 GN=GGACT PE=1 SV=2 | 9 | 2 | 1 | 17.3 | 14 | 153 | 6.87 | | | | 1 | 1 | 1 | | | | 1 | 1 | 1 |
| High | Q9NXE4 | Sphingomyelin phosphodiesterase 4 OS=Homo sapiens OX=9606 GN=SMPD4 PE=1 SV=3 | 9 | 4 | 4 | 97.7 | 7 | 866 | 8.27 | 2 | 1 | 2 | | | | 2 | 1 | 2 | | | |
| High | P62891 | 60S ribosomal protein L39 OS=Homo sapiens OX=9606 GN=RPL39 PE=1 SV=2 | 9 | 1 | 1 | 6.4 | 20 | 51 | 12.56 | 1 | 1 | 1 | 1 | 1 | 1 | 1 | 1 | 1 | 1 | 1 | 1 |
| High | Q03393 | 6-pyruvoyl tetrahydrobiopterin synthase OS=Homo sapiens OX=9606 GN=PTS PE=1 SV=1 | 9 | 3 | 3 | 16.4 | 21 | 145 | 6.68 | 1 | 1 | 1 | 1 | 1 | 1 | 1 | 1 | 1 | 1 | 1 | 1 |
| High | Q16658 | Fascin OS=Homo sapiens OX=9606 GN=FSCN1 PE=1 SV=3 | 9 | 2 | 2 | 54.5 | 5 | 493 | 7.24 | 1 | 1 | 1 | 2 | 2 | 2 | 1 | 1 | 1 | 2 | 2 | 2 |
| High | Q96N66 | Lysophospholipid acyltransferase 7 OS=Homo sapiens OX=9606 GN=MBOAT7 PE=1 SV=2 | 9 | 3 | 3 | 52.7 | 8 | 472 | 8.97 | 1 | 1 | 1 | | | | 1 | 1 | 1 | | | |
| High | Q9Y2S7 | Polymerase delta-interacting protein 2 OS=Homo sapiens OX=9606 GN=POLDIP2 PE=1 SV=1 | 9 | 4 | 4 | 42 | 14 | 368 | 8.63 | 1 | 1 | | | 1 | 1 | 1 | 1 | | | 1 | 1 |
| High | Q9BXW7 | Haloacid dehalogenase-like hydrolase domain-containing 5 OS=Homo sapiens OX=9606 GN=HDHD5 PE=1 SV=1 | 9 | 3 | 3 | 46.3 | 9 | 423 | 8.13 | | 1 | | 1 | | 1 | | 1 | | 1 | | 1 |
| High | P36405 | ADP-ribosylation factor-like protein 3 OS=Homo sapiens OX=9606 GN=ARL3 PE=1 SV=2 | 9 | 4 | 4 | 20.4 | 35 | 182 | 7.24 | 1 | 2 | | | | | 1 | 2 | | | | |
| High | Q6P6C2 | RNA demethylase ALKBH5 OS=Homo sapiens OX=9606 GN=ALKBH5 PE=1 SV=2 | 9 | 2 | 2 | 44.2 | 5 | 394 | 9.09 | | | 1 | 2 | 2 | 1 | | | 1 | 2 | 2 | 1 |
| High | O15160 | DNA-directed RNA polymerases I and III subunit RPAC1 OS=Homo sapiens OX=9606 GN=POLR1C PE=1 SV=1 | 9 | 3 | 3 | 39.2 | 10 | 346 | 5.5 | | | 1 | | | 1 | | | 1 | | | 1 |
| High | Q15334 | Lethal(2) giant larvae protein homolog 1 OS=Homo sapiens OX=9606 GN=LLGL1 PE=1 SV=3 | 9 | 4 | 4 | 115.3 | 5 | 1064 | 6.29 | | | 1 | | 1 | 1 | | | 1 | | 1 | 1 |
| High | P42766 | 60S ribosomal protein L35 OS=Homo sapiens OX=9606 GN=RPL35 PE=1 SV=2 | 9 | 1 | 1 | 14.5 | 8 | 123 | 11.05 | 1 | 1 | 1 | 1 | 1 | 1 | 1 | 1 | 1 | 1 | 1 | 1 |
| High | Q9Y4L1 | Hypoxia up-regulated protein 1 OS=Homo sapiens OX=9606 GN=HYOU1 PE=1 SV=1 | 9 | 4 | 4 | 111.3 | 8 | 999 | 5.22 | 2 | 1 | 3 | | | | 2 | 1 | 3 | | | |
| High | Q12834 | Cell division cycle protein 20 homolog OS=Homo sapiens OX=9606 GN=CDC20 PE=1 SV=2 | 9 | 1 | 1 | 54.7 | 3 | 499 | 9.23 | 1 | 1 | 1 | 1 | 1 | 1 | 1 | 1 | 1 | 1 | 1 | 1 |
| High | Q96JH7 | Deubiquitinating protein VCPIP1 OS=Homo sapiens OX=9606 GN=VCPIP1 PE=1 SV=2 | 9 | 3 | 3 | 134.2 | 2 | 1222 | 7.2 | | | 1 | | 1 | 2 | | | 1 | | 1 | 2 |
| High | Q13595 | Transformer-2 protein homolog alpha OS=Homo sapiens OX=9606 GN=TRA2A PE=1 SV=1 | 9 | 5 | 5 | 32.7 | 20 | 282 | 11.27 | 1 | 1 | 2 | | 1 | 2 | 1 | 1 | 2 | | 1 | 2 |
| High | Q9UMZ2 | Synergin gamma OS=Homo sapiens OX=9606 GN=SYNRG PE=1 SV=2 | 9 | 2 | 2 | 140.6 | 2 | 1314 | 5.03 | 2 | 2 | 1 | | 1 | | 2 | 2 | 1 | | 1 | |
| High | Q8WXG6 | MAP kinase-activating death domain protein OS=Homo sapiens OX=9606 GN=MADD PE=1 SV=2 | 9 | 3 | 3 | 183.2 | 3 | 1647 | 6.04 | 1 | 1 | 2 | | | | 1 | 1 | 2 | | | |
| High | P62304 | Small nuclear ribonucleoprotein E OS=Homo sapiens OX=9606 GN=SNRPE PE=1 SV=1 | 9 | 2 | 2 | 10.8 | 25 | 92 | 9.44 | 1 | 1 | 2 | 2 | 1 | | 1 | 1 | 2 | 1 | 1 | |
| High | Q9Y6E0 | Serine/threonine-protein kinase 24 OS=Homo sapiens OX=9606 GN=STK24 PE=1 SV=1 | 9 | 4 | 4 | 49.3 | 9 | 443 | 5.69 | | | | | | | | | | | | |
| High | Q96AG4 | Leucine-rich repeat-containing protein 59 OS=Homo sapiens OX=9606 GN=LRRC59 PE=1 SV=1 | 9 | 4 | 4 | 34.9 | 15 | 307 | 9.57 | | | 2 | 1 | 1 | 1 | | | 2 | 1 | 1 | 1 |
| High | Q9BVJ6 | U3 small nucleolar RNA-associated protein 14 homolog A OS=Homo sapiens OX=9606 GN=UTP14A PE=1 SV=1 | 9 | 3 | 3 | 87.9 | 6 | 771 | 7.87 | | 1 | 1 | 1 | 3 | 1 | | 1 | 1 | 1 | 3 | 1 |
| High | Q9Y2D5 | A-kinase anchor protein 2 OS=Homo sapiens OX=9606 GN=AKAP2 PE=1 SV=3 | 9 | 4 | 4 | 94.6 | 4 | 859 | 5.11 | 1 | 3 | 2 | | 1 | | 1 | 3 | 2 | | 1 | |
| High | Q8N9T8 | Protein KRI1 homolog OS=Homo sapiens OX=9606 GN=KRI1 PE=1 SV=3 | 9 | 3 | 3 | 82.5 | 7 | 703 | 5.14 | 2 | | | | | | 1 | | | | | |
| High | Q8NEY8 | Periphilin-1 OS=Homo sapiens OX=9606 GN=PPHLN1 PE=1 SV=2 | 9 | 2 | 2 | 52.7 | 4 | 458 | 9.11 | 1 | | | 1 | 2 | 1 | 1 | | | 1 | 2 | 1 |
| High | Q9P2X0 | Dolichol-phosphate mannosyltransferase subunit 3 OS=Homo sapiens OX=9606 GN=DPM3 PE=1 SV=2 | 9 | 2 | 2 | 10.1 | 24 | 92 | 5.94 | 1 | 1 | 1 | 1 | | 1 | 1 | 1 | 1 | 1 | | 1 |
| High | Q15014 | Mortality factor 4-like protein 2 OS=Homo sapiens OX=9606 GN=MORF4L2 PE=1 SV=1 | 9 | 3 | 2 | 32.3 | 11 | 288 | 9.72 | | | | 3 | 1 | 1 | | | | 3 | 1 | 1 |
| High | O14561 | Acyl carrier protein, mitochondrial OS=Homo sapiens OX=9606 GN=NDUFAB1 PE=1 SV=3 | 9 | 1 | 1 | 17.4 | 9 | 156 | 4.93 | 1 | 1 | 2 | | | | 1 | 1 | 1 | | | |
| High | O75146 | Huntingtin-interacting protein 1-related protein OS=Homo sapiens OX=9606 GN=HIP1R PE=1 SV=2 | 9 | 4 | 3 | 119.3 | 4 | 1068 | 6.67 | | 1 | 1 | | | | | 1 | 1 | | | |
| High | P18510 | Interleukin-1 receptor antagonist protein OS=Homo sapiens OX=9606 GN=IL1RN PE=1 SV=1 | 9 | 3 | 3 | 20 | 19 | 177 | 6.19 | | | | 2 | 1 | 1 | | | | 2 | 1 | 1 |
| High | Q9Y3B2 | Exosome complex component CSL4 OS=Homo sapiens OX=9606 GN=EXOSC1 PE=1 SV=1 | 9 | 3 | 3 | 21.4 | 17 | 195 | 8.24 | 1 | | | | 2 | 1 | 1 | | | | 2 | 1 |
| High | Q9NVJ2 | ADP-ribosylation factor-like protein 8B OS=Homo sapiens OX=9606 GN=ARL8B PE=1 SV=1 | 9 | 3 | 3 | 21.5 | 18 | 186 | 8.43 | 1 | 2 | 2 | | | | 1 | 2 | 2 | | | |
| High | Q96BR5 | Cytochrome c oxidase assembly factor 7 OS=Homo sapiens OX=9606 GN=COA7 PE=1 SV=2 | 9 | 3 | 3 | 25.7 | 14 | 231 | 6.02 | | | 1 | 1 | 1 | 1 | | | 1 | 1 | 1 | 1 |
| High | Q07866 | Kinesin light chain 1 OS=Homo sapiens OX=9606 GN=KLC1 PE=1 SV=2 | 9 | 3 | 1 | 65.3 | 4 | 573 | 6.2 | 1 | 1 | 1 | 1 | 1 | | 1 | 1 | 1 | 1 | 1 | |
| High | P49711 | Transcriptional repressor CTCF OS=Homo sapiens OX=9606 GN=CTCF PE=1 SV=1 | 9 | 4 | 4 | 82.7 | 8 | 727 | 6.96 | | | | 1 | | | | | | 1 | | |
| High | Q8TBB0 | THAP domain-containing protein 6 OS=Homo sapiens OX=9606 GN=THAP6 PE=1 SV=1 | 9 | 3 | 3 | 25.7 | 20 | 222 | 8.7 | | | | 1 | 1 | 1 | | | | 1 | 1 | 1 |
| High | Q10713 | Mitochondrial-processing peptidase subunit alpha OS=Homo sapiens OX=9606 GN=PMPCA PE=1 SV=2 | 9 | 4 | 4 | 58.2 | 7 | 525 | 6.92 | 3 | 3 | 3 | | | | 3 | 3 | 3 | | | |

| | | | | | | | | | | | | | | | | | | | | | |
|---|---|---|---|---|---|---|---|---|---|---|---|---|---|---|---|---|---|---|---|---|---|
| High | Q9Y2R9 | 28S ribosomal protein S7, mitochondrial OS=Homo sapiens OX=9606 GN=MRPS7 PE=1 SV=2 | 9 | 5 | 5 | 28.1 | 26 | 242 | 9.99 | | | | 1 | 1 | 1 | | | | 1 | 1 | 1 |
| High | Q7Z6K1 | THAP domain-containing protein 5 OS=Homo sapiens OX=9606 GN=THAP5 PE=1 SV=2 | 9 | 1 | 1 | 45.4 | 2 | 395 | 6.71 | 1 | 1 | 1 | 1 | 1 | 1 | 1 | 1 | 1 | 1 | 1 | 1 |
| High | Q16854 | Deoxyguanosine kinase, mitochondrial OS=Homo sapiens OX=9606 GN=DGUOK PE=1 SV=2 | 9 | 3 | 3 | 32 | 15 | 277 | 8.66 | 1 | | 2 | 1 | 1 | 2 | 1 | | 2 | 1 | 1 | 2 |
| High | Q14669 | E3 ubiquitin-protein ligase TRIP12 OS=Homo sapiens OX=9606 GN=TRIP12 PE=1 SV=1 | 9 | 5 | 5 | 220.3 | 3 | 1992 | 8.48 | | | | 1 | | 1 | | | | 1 | | 1 |
| High | P35268 | 60S ribosomal protein L22 OS=Homo sapiens OX=9606 GN=RPL22 PE=1 SV=2 | 9 | 1 | 1 | 14.8 | 9 | 128 | 9.19 | 1 | 1 | 1 | 1 | 1 | 1 | 1 | 1 | 1 | 1 | 1 | 1 |
| High | Q8IVF2 | Protein AHNAK2 OS=Homo sapiens OX=9606 GN=AHNAK2 PE=1 SV=2 | 9 | 2 | 2 | 616.2 | 3 | 5795 | 5.36 | 1 | 1 | 2 | | 1 | 1 | 1 | 1 | 2 | | 1 | 1 |
| High | O00541 | Pescadillo homolog OS=Homo sapiens OX=9606 GN=PES1 PE=1 SV=1 | 9 | 3 | 3 | 68 | 6 | 588 | 7.33 | 1 | 1 | 1 | 1 | | 1 | 1 | 1 | 1 | 1 | | 1 |
| High | Q14493 | Histone RNA hairpin-binding protein OS=Homo sapiens OX=9606 GN=SLBP PE=1 SV=1 | 9 | 3 | 3 | 31.3 | 14 | 270 | 7.47 | | | 1 | 1 | | | | | 1 | 1 | | |
| High | Q15154 | Pericentriolar material 1 protein OS=Homo sapiens OX=9606 GN=PCM1 PE=1 SV=5 | 9 | 3 | 3 | 228.4 | 2 | 2024 | 5.02 | 2 | 1 | 1 | | | | 2 | 1 | 1 | | | |
| High | Q96RQ1 | Endoplasmic reticulum-Golgi intermediate compartment protein 2 OS=Homo sapiens OX=9606 GN=ERGIC2 PE=1 SV=2 | 9 | 4 | 4 | 42.5 | 15 | 377 | 6.77 | 1 | 3 | 3 | | | 1 | 1 | 3 | 3 | | | 1 |
| High | P51553 | Isocitrate dehydrogenase [NAD] subunit gamma, mitochondrial OS=Homo sapiens OX=9606 GN=IDH3G PE=1 SV=1 | 9 | 2 | 2 | 42.8 | 4 | 393 | 8.5 | 2 | 2 | | 1 | 1 | 1 | 2 | 2 | | 1 | 1 | 1 |
| High | Q9H8V3 | Protein ECT2 OS=Homo sapiens OX=9606 GN=ECT2 PE=1 SV=4 | 9 | 4 | 4 | 103.4 | 7 | 914 | 7.71 | 1 | 1 | 1 | | | | 1 | 1 | 1 | | | |
| High | Q9Y6X8 | Zinc fingers and homeoboxes protein 2 OS=Homo sapiens OX=9606 GN=ZHX2 PE=1 SV=1 | 9 | 4 | 3 | 92.3 | 5 | 837 | 6.86 | | | | 2 | | 1 | | | | 2 | | 1 |
| High | Q01968 | Inositol polyphosphate 5-phosphatase OCRL OS=Homo sapiens OX=9606 GN=OCRL PE=1 SV=3 | 9 | 2 | 2 | 104.1 | 2 | 901 | 6.55 | 1 | 1 | 1 | 1 | 1 | | 1 | 1 | 1 | 1 | 1 | |
| High | Q9BU76 | Multiple myeloma tumor-associated protein 2 OS=Homo sapiens OX=9606 GN=MMTAG2 PE=1 SV=1 | 9 | 5 | 5 | 29.4 | 20 | 263 | 10.02 | | | | | 2 | 2 | | | | | 2 | 2 |
| High | Q969S3 | Zinc finger protein 622 OS=Homo sapiens OX=9606 GN=ZNF622 PE=1 SV=1 | 9 | 3 | 3 | 54.2 | 8 | 477 | 6.15 | 1 | 1 | 1 | 1 | | | 1 | 1 | 1 | 1 | | |
| High | Q14671 | Pumilio homolog 1 OS=Homo sapiens OX=9606 GN=PUM1 PE=1 SV=3 | 9 | 5 | 5 | 126.4 | 6 | 1186 | 6.84 | | 2 | 4 | | | | | 2 | 4 | | | |
| High | P54259 | Atrophin-1 OS=Homo sapiens OX=9606 GN=ATN1 PE=1 SV=3 | 9 | 2 | 2 | 125.3 | 3 | 1190 | 8.98 | 1 | 1 | 2 | 1 | 1 | | 1 | 1 | 2 | 1 | 1 | |
| High | Q9C0B0 | RING finger protein unkempt homolog OS=Homo sapiens OX=9606 GN=UNK PE=1 SV=2 | 9 | 3 | 3 | 88 | 5 | 810 | 6.86 | 2 | 3 | 1 | | | | 2 | 2 | 1 | | | |
| High | O94855 | Protein transport protein Sec24D OS=Homo sapiens OX=9606 GN=SEC24D PE=1 SV=2 | 9 | 5 | 4 | 112.9 | 6 | 1032 | 7.25 | | 1 | | 1 | | 1 | | 1 | | 1 | | 1 |
| High | Q9H7E9 | UPF0488 protein C8orf33 OS=Homo sapiens OX=9606 GN=C8orf33 PE=1 SV=1 | 9 | 3 | 3 | 25 | 19 | 229 | 9.95 | | | | 1 | 2 | 1 | | | | 1 | 2 | 1 |
| High | Q6R327 | Rapamycin-insensitive companion of mTOR OS=Homo sapiens OX=9606 GN=RICTOR PE=1 SV=1 | 9 | 4 | 4 | 192.1 | 4 | 1708 | 7.47 | 3 | 2 | 3 | | | | 3 | 2 | 3 | | | |
| High | P61019 | Ras-related protein Rab-2A OS=Homo sapiens OX=9606 GN=RAB2A PE=1 SV=1 | 9 | 4 | 4 | 23.5 | 22 | 212 | 6.54 | 1 | | 1 | 2 | | 2 | 1 | | 1 | 2 | | 2 |
| High | Q9NXV6 | CDKN2A-interacting protein OS=Homo sapiens OX=9606 GN=CDKN2AIP PE=1 SV=3 | 9 | 3 | 3 | 61.1 | 6 | 580 | 9.01 | 1 | | | | 1 | | 1 | | | | 1 | |
| High | Q99081 | Transcription factor 12 OS=Homo sapiens OX=9606 GN=TCF12 PE=1 SV=1 | 9 | 2 | 2 | 72.9 | 6 | 682 | 7.02 | 1 | 1 | 1 | 1 | 1 | 1 | 1 | 1 | 1 | 1 | 1 | 1 |
| High | P42574 | Caspase-3 OS=Homo sapiens OX=9606 GN=CASP3 PE=1 SV=2 | 9 | 2 | 1 | 31.6 | 7 | 277 | 6.54 | | | | 1 | 1 | 2 | | | | 1 | 1 | 2 |
| High | Q6P1L8 | 39S ribosomal protein L14, mitochondrial OS=Homo sapiens OX=9606 GN=MRPL14 PE=1 SV=1 | 9 | 2 | 2 | 15.9 | 17 | 145 | 10.24 | 1 | 1 | 1 | | 1 | | 1 | 1 | 1 | | 1 | |
| High | Q8IXQ6 | Protein mono-ADP-ribosyltransferase PARP9 OS=Homo sapiens OX=9606 GN=PARP9 PE=1 SV=2 | 9 | 2 | 2 | 96.3 | 2 | 854 | 7.91 | 1 | 1 | 1 | 2 | 1 | | 1 | 1 | 1 | 2 | 1 | |
| High | Q969N2 | GPI transamidase component PIG-T OS=Homo sapiens OX=9606 GN=PIGT PE=1 SV=1 | 9 | 6 | 6 | 65.7 | 12 | 578 | 8.38 | 2 | 2 | 3 | | | | 2 | 2 | 3 | | | |
| High | P16930 | Fumarylacetoacetase OS=Homo sapiens OX=9606 GN=FAH PE=1 SV=2 | 9 | 3 | 3 | 46.3 | 10 | 419 | 6.95 | 2 | 2 | | | 1 | | 2 | 2 | | | 1 | |
| High | Q9Y580 | RNA-binding protein 7 OS=Homo sapiens OX=9606 GN=RBM7 PE=1 SV=1 | 9 | 2 | 2 | 30.5 | 6 | 266 | 9.57 | 1 | 1 | 1 | | 1 | | 1 | 1 | 1 | | 1 | |
| High | P08581 | Hepatocyte growth factor receptor OS=Homo sapiens OX=9606 GN=MET PE=1 SV=4 | 9 | 2 | 1 | 155.4 | 2 | 1390 | 7.33 | 1 | 1 | 1 | | 1 | 1 | 1 | 1 | 1 | | 1 | 1 |
| High | P07339 | Cathepsin D OS=Homo sapiens OX=9606 GN=CTSD PE=1 SV=1 | 9 | 6 | 6 | 44.5 | 21 | 412 | 6.54 | | | 2 | 1 | | 1 | | | 2 | 1 | | 1 |
| High | P48200 | Iron-responsive element-binding protein 2 OS=Homo sapiens OX=9606 GN=IREB2 PE=1 SV=4 | 9 | 4 | 4 | 105 | 5 | 963 | 7.05 | 1 | 1 | | | | | 1 | 1 | | | | |
| High | P78344 | Eukaryotic translation initiation factor 4 gamma 2 OS=Homo sapiens OX=9606 GN=EIF4G2 PE=1 SV=1 | 9 | 5 | 5 | 102.3 | 5 | 907 | 7.14 | 2 | 2 | 3 | | | | 2 | 2 | 3 | | | |
| High | O15379 | Histone deacetylase 3 OS=Homo sapiens OX=9606 GN=HDAC3 PE=1 SV=2 | 9 | 3 | 2 | 48.8 | 9 | 428 | 5.16 | 1 | 1 | 2 | | | 1 | 1 | 1 | 2 | | | 1 |
| High | P10586 | Receptor-type tyrosine-protein phosphatase F OS=Homo sapiens OX=9606 GN=PTPRF PE=1 SV=2 | 9 | 4 | 4 | 212.7 | 3 | 1907 | 6.3 | 1 | 1 | 1 | | 1 | | 1 | 1 | 1 | | 1 | |
| High | Q96JJ7 | Protein disulfide-isomerase TMX3 OS=Homo sapiens OX=9606 GN=TMX3 PE=1 SV=2 | 9 | 3 | 3 | 51.8 | 6 | 454 | 4.91 | | | 1 | | 1 | | | | 1 | | 1 | |
| High | Q6NZY4 | Zinc finger CCHC domain-containing protein 8 OS=Homo sapiens OX=9606 GN=ZCCHC8 PE=1 SV=2 | 9 | 3 | 3 | 78.5 | 7 | 707 | 4.87 | 1 | 1 | 1 | | 1 | 1 | 1 | 1 | 1 | | 1 | 1 |
| High | Q9BZE1 | 39S ribosomal protein L37, mitochondrial OS=Homo sapiens OX=9606 GN=MRPL37 PE=1 SV=2 | 9 | 4 | 4 | 48.1 | 11 | 423 | 8.59 | | | | 1 | 3 | | | | | 1 | 3 | |
| High | P29728 | 2'-5'-oligoadenylate synthase 2 OS=Homo sapiens OX=9606 GN=OAS2 PE=1 SV=3 | 8 | 2 | 2 | 82.4 | 4 | 719 | 8.25 | | | | 1 | 2 | 1 | | | | 1 | 2 | 1 |
| High | Q9BWJ5 | Splicing factor 3B subunit 5 OS=Homo sapiens OX=9606 GN=SF3B5 PE=1 SV=1 | 8 | 2 | 2 | 10.1 | 33 | 86 | 6.35 | 1 | 1 | | | 1 | 1 | 1 | 1 | | | 1 | 1 |
| High | Q9P003 | Protein cornichon homolog 4 OS=Homo sapiens OX=9606 GN=CNIH4 PE=1 SV=1 | 8 | 1 | 1 | 16.1 | 14 | 139 | 6.65 | 3 | 2 | 1 | | | | 1 | 1 | 1 | | | |
| High | P21741 | Midkine OS=Homo sapiens OX=9606 GN=MDK PE=1 SV=1 | 8 | 3 | 3 | 15.6 | 26 | 143 | 9.79 | | | | 1 | 1 | 1 | | | | 1 | 1 | 1 |
| High | P98194 | Calcium-transporting ATPase type 2C member 1 OS=Homo sapiens OX=9606 GN=ATP2C1 PE=1 SV=3 | 8 | 4 | 4 | 100.5 | 5 | 919 | 6.74 | 1 | 2 | 2 | | | | 1 | 2 | 2 | | | |
| High | Q8TAA9 | Vang-like protein 1 OS=Homo sapiens OX=9606 GN=VANGL1 PE=1 SV=1 | 8 | 3 | 3 | 59.9 | 8 | 524 | 8.81 | | | | 1 | 2 | 2 | | | | 1 | 2 | 2 |
| High | O96013 | Serine/threonine-protein kinase PAK 4 OS=Homo sapiens OX=9606 GN=PAK4 PE=1 SV=1 | 8 | 3 | 3 | 64 | 7 | 591 | 9.73 | 2 | 1 | 1 | | | | 2 | 1 | 1 | | | |
| High | P51668 | Ubiquitin-conjugating enzyme E2 D1 OS=Homo sapiens OX=9606 GN=UBE2D1 PE=1 SV=1 | 8 | 2 | 1 | 16.6 | 20 | 147 | 7.42 | 2 | 2 | 1 | 1 | 1 | 1 | 2 | 2 | 1 | 1 | 1 | 1 |
| High | Q9BRZ2 | E3 ubiquitin-protein ligase TRIM56 OS=Homo sapiens OX=9606 GN=TRIM56 PE=1 SV=3 | 8 | 6 | 6 | 81.4 | 10 | 755 | 7.74 | | | 1 | | 1 | | | | 1 | | 1 | |
| High | Q92575 | UBX domain-containing protein 4 OS=Homo sapiens OX=9606 GN=UBXN4 PE=1 SV=2 | 8 | 2 | 2 | 56.7 | 7 | 508 | 6.38 | 1 | | 1 | | | | 1 | | 1 | | | |
| High | Q1ED39 | Lysine-rich nucleolar protein 1 OS=Homo sapiens OX=9606 GN=KNOP1 PE=1 SV=1 | 8 | 5 | 5 | 51.6 | 16 | 458 | 9.86 | | | | | 1 | | | | | | 1 | |
| High | Q9NW64 | Pre-mRNA-splicing factor RBM22 OS=Homo sapiens OX=9606 GN=RBM22 PE=1 SV=1 | 8 | 2 | 2 | 46.9 | 5 | 420 | 8.54 | | 1 | 1 | | 1 | 1 | | 1 | 1 | | 1 | 1 |
| High | Q9P265 | Disco-interacting protein 2 homolog B OS=Homo sapiens OX=9606 GN=DIP2B PE=1 SV=3 | 8 | 4 | 4 | 171.4 | 3 | 1576 | 8.09 | 1 | 1 | 2 | | | | 1 | 1 | 2 | | | |
| High | Q9NY93 | Probable ATP-dependent RNA helicase DDX56 OS=Homo sapiens OX=9606 GN=DDX56 PE=1 SV=1 | 8 | 5 | 5 | 61.6 | 12 | 547 | 9.26 | | 1 | | 1 | 1 | 2 | | 1 | | 1 | 1 | 2 |
| High | Q13451 | Peptidyl-prolyl cis-trans isomerase FKBP5 OS=Homo sapiens OX=9606 GN=FKBP5 PE=1 SV=2 | 8 | 4 | 4 | 51.2 | 11 | 457 | 5.9 | 1 | 1 | 1 | | | | 1 | 1 | 1 | | | |
| High | Q96CB8 | Integrator complex subunit 12 OS=Homo sapiens OX=9606 GN=INTS12 PE=1 SV=1 | 8 | 2 | 2 | 48.8 | 7 | 462 | 9.69 | 2 | 2 | 1 | | | | 2 | 2 | 1 | | | |
| High | Q8TEA1 | tRNA (cytosine(72)-C(5))-methyltransferase NSUN6 OS=Homo sapiens OX=9606 GN=NSUN6 PE=1 SV=1 | 8 | 1 | 1 | 51.7 | 4 | 469 | 7.96 | 1 | 1 | 1 | | | | 1 | 1 | 1 | | | |
| High | Q9UHR5 | SAP30-binding protein OS=Homo sapiens OX=9606 GN=SAP30BP PE=1 SV=1 | 8 | 1 | 1 | 33.9 | 3 | 308 | 4.84 | 1 | 1 | | 1 | 1 | 1 | 1 | 1 | | 1 | 1 | 1 |
| High | P28290 | Protein ITPRID2 OS=Homo sapiens OX=9606 GN=ITPRID2 PE=1 SV=3 | 8 | 4 | 4 | 138.3 | 4 | 1259 | 5.19 | | | | | | | | | | | | |
| High | Q92879 | CUGBP Elav-like family member 1 OS=Homo sapiens OX=9606 GN=CELF1 PE=1 SV=2 | 8 | 3 | 3 | 52 | 6 | 486 | 8.46 | | 1 | 1 | | | | | 1 | 1 | | | |
| High | P49591 | Serine--tRNA ligase, cytoplasmic OS=Homo sapiens OX=9606 GN=SARS1 PE=1 SV=3 | 8 | 2 | 2 | 58.7 | 9 | 514 | 6.43 | 2 | 1 | 2 | | | | 2 | 1 | 2 | | | |
| High | O15230 | Laminin subunit alpha-5 OS=Homo sapiens OX=9606 GN=LAMA5 PE=1 SV=8 | 8 | 2 | 2 | 399.5 | 1 | 3695 | 7.02 | | | 1 | 1 | 2 | 1 | | | 1 | 1 | 2 | 1 |
| High | Q96CW5 | Gamma-tubulin complex component 3 OS=Homo sapiens OX=9606 GN=TUBGCP3 PE=1 SV=2 | 8 | 4 | 4 | 103.5 | 8 | 907 | 8.12 | 1 | 2 | 2 | | | | 1 | 2 | 2 | | | |
| High | Q6PJG2 | Mitotic deacetylase-associated SANT domain protein OS=Homo sapiens OX=9606 GN=MIDEAS PE=1 SV=2 | 8 | 5 | 5 | 114.9 | 9 | 1045 | 9.19 | | | | | | | | | | | | |
| High | Q9NRN7 | L-aminoadipate-semialdehyde dehydrogenase-phosphopantetheinyl transferase OS=Homo sapiens OX=9606 GN=AASDHPPT PE=1 | 8 | 2 | 2 | 35.8 | 10 | 309 | 6.8 | 2 | | 1 | | | | 2 | | 1 | | | |
| High | Q5VV67 | Peroxisome proliferator-activated receptor gamma coactivator-related protein 1 OS=Homo sapiens OX=9606 GN=PPRC1 PE=1 SV= | 8 | 5 | 5 | 177.4 | 5 | 1664 | 6.51 | | | | 3 | 4 | 1 | | | | 3 | 4 | 1 |
| High | P36639 | Oxidized purine nucleoside triphosphate hydrolase OS=Homo sapiens OX=9606 GN=NUDT1 PE=1 SV=4 | 8 | 2 | 2 | 17.9 | 16 | 156 | 5.08 | 1 | 1 | 2 | | | 1 | 1 | 1 | 2 | | | 1 |
| High | Q9NPD3 | Exosome complex component RRP41 OS=Homo sapiens OX=9606 GN=EXOSC4 PE=1 SV=3 | 8 | 2 | 2 | 26.4 | 9 | 245 | 6.52 | 1 | | | 1 | 1 | 1 | 1 | | | 1 | 1 | 1 |
| High | Q96GM5 | SWI/SNF-related matrix-associated actin-dependent regulator of chromatin subfamily D member 1 OS=Homo sapiens OX=9606 GI | 8 | 3 | 2 | 58.2 | 8 | 515 | 9.25 | | | 1 | 1 | | | | | 1 | 1 | | |
| High | Q9P2I0 | Cleavage and polyadenylation specificity factor subunit 2 OS=Homo sapiens OX=9606 GN=CPSF2 PE=1 SV=2 | 8 | 4 | 4 | 88.4 | 5 | 782 | 5.11 | 1 | 1 | 1 | | | | 1 | 1 | 1 | | | |
| High | Q6IA86 | Elongator complex protein 2 OS=Homo sapiens OX=9606 GN=ELP2 PE=1 SV=2 | 8 | 4 | 4 | 92.4 | 6 | 826 | 5.96 | 1 | 1 | 1 | | | | 1 | 1 | 1 | | | |
| High | Q9BRT6 | Protein LLP homolog OS=Homo sapiens OX=9606 GN=LLPH PE=1 SV=1 | 8 | 2 | 2 | 15.2 | 19 | 129 | 10.37 | | | 2 | 1 | 1 | 1 | | | 2 | 1 | 1 | 1 |
| High | Q13541 | Eukaryotic translation initiation factor 4E-binding protein 1 OS=Homo sapiens OX=9606 GN=EIF4EBP1 PE=1 SV=3 | 8 | 3 | 3 | 12.6 | 59 | 118 | 5.48 | | | 2 | | | | | | 2 | | | |
| High | Q15796 | Mothers against decapentaplegic homolog 2 OS=Homo sapiens OX=9606 GN=SMAD2 PE=1 SV=1 | 8 | 2 | 1 | 52.3 | 5 | 467 | 6.58 | | | | 1 | 2 | 1 | | | | 1 | 2 | 1 |
| High | Q9NP66 | High mobility group protein 20A OS=Homo sapiens OX=9606 GN=HMG20A PE=1 SV=1 | 8 | 4 | 4 | 40.1 | 24 | 347 | 6.49 | 1 | 2 | 5 | | | | 1 | 2 | 4 | | | |
| High | Q00587 | Cdc42 effector protein 1 OS=Homo sapiens OX=9606 GN=CDC42EP1 PE=1 SV=1 | 8 | 3 | 3 | 40.3 | 8 | 391 | 7.15 | 1 | 1 | 1 | 1 | | 1 | 1 | 1 | 1 | 1 | | 1 |
| High | Q8N3E9 | 1-phosphatidylinositol 4,5-bisphosphate phosphodiesterase delta-3 OS=Homo sapiens OX=9606 GN=PLCD3 PE=1 SV=3 | 8 | 3 | 3 | 89.2 | 7 | 789 | 6.98 | | 1 | 2 | | | | | 1 | 2 | | | |
| High | Q96SK2 | Transmembrane protein 209 OS=Homo sapiens OX=9606 GN=TMEM209 PE=1 SV=2 | 8 | 5 | 5 | 62.9 | 14 | 561 | 8.63 | | | 3 | | | | | | 3 | | | |
| High | Q12774 | Rho guanine nucleotide exchange factor 5 OS=Homo sapiens OX=9606 GN=ARHGEF5 PE=1 SV=3 | 8 | 2 | 2 | 176.7 | 2 | 1597 | 5.53 | | | 1 | 1 | 1 | 2 | | | 1 | 1 | 1 | 2 |
| High | Q9H8W4 | Pleckstrin homology domain-containing family F member 2 OS=Homo sapiens OX=9606 GN=PLEKHF2 PE=1 SV=1 | 8 | 2 | 2 | 27.8 | 12 | 249 | 8.21 | | 1 | | 1 | 1 | 1 | | 1 | | 1 | 1 | 1 |
| High | P19256 | Lymphocyte function-associated antigen 3 OS=Homo sapiens OX=9606 GN=CD58 PE=1 SV=1 | 8 | 2 | 2 | 28.1 | 7 | 250 | 6.76 | 1 | 1 | 1 | | 1 | | 1 | 1 | 1 | | 1 | |
| High | P62330 | ADP-ribosylation factor 6 OS=Homo sapiens OX=9606 GN=ARF6 PE=1 SV=2 | 8 | 2 | 2 | 20.1 | 15 | 175 | 8.95 | 2 | 1 | 2 | | | | 2 | 1 | 2 | | | |
| High | Q9ULU4 | Protein kinase C-binding protein 1 OS=Homo sapiens OX=9606 GN=ZMYND8 PE=1 SV=2 | 8 | 4 | 4 | 131.6 | 5 | 1186 | 7.2 | 1 | | 1 | 1 | | 1 | 1 | | 1 | 1 | | 1 |
| High | P55263 | Adenosine kinase OS=Homo sapiens OX=9606 GN=ADK PE=1 SV=2 | 8 | 2 | 2 | 40.5 | 8 | 362 | 6.7 | | 1 | 1 | | 1 | 1 | | 1 | 1 | | 1 | 1 |
| High | P20674 | Cytochrome c oxidase subunit 5A, mitochondrial OS=Homo sapiens OX=9606 GN=COX5A PE=1 SV=2 | 8 | 3 | 3 | 16.8 | 21 | 150 | 6.79 | 2 | 2 | 1 | | 1 | | 2 | 2 | 1 | | 1 | |
| High | O75363 | Breast carcinoma-amplified sequence 1 OS=Homo sapiens OX=9606 GN=BCAS1 PE=1 SV=2 | 8 | 2 | 2 | 61.7 | 5 | 584 | 5.1 | 1 | 1 | 1 | | | | 1 | 1 | 1 | | | |
| High | O75506 | Heat shock factor-binding protein 1 OS=Homo sapiens OX=9606 GN=HSBP1 PE=1 SV=1 | 8 | 2 | 2 | 8.5 | 58 | 76 | 4.36 | 2 | 1 | 3 | | | | 2 | 1 | 2 | | | |
| High | Q9NZQ3 | NCK-interacting protein with SH3 domain OS=Homo sapiens OX=9606 GN=NCKIPSD PE=1 SV=1 | 8 | 4 | 4 | 78.9 | 7 | 722 | 6.38 | 3 | 3 | 2 | | | | 3 | 3 | 2 | | | |
| High | Q5VYY1 | Ankyrin repeat domain-containing protein 22 OS=Homo sapiens OX=9606 GN=ANKRD22 PE=2 SV=1 | 8 | 2 | 2 | 21.8 | 9 | 191 | 8.84 | 2 | 2 | | | | | 2 | 2 | | | | |
| High | P06702 | Protein S100-A9 OS=Homo sapiens OX=9606 GN=S100A9 PE=1 SV=1 | 8 | 3 | 3 | 13.2 | 31 | 114 | 6.13 | | | | 2 | 3 | 2 | | | | 2 | 3 | 2 |
| High | Q13887 | Krueppel-like factor 5 OS=Homo sapiens OX=9606 GN=KLF5 PE=1 SV=2 | 8 | 3 | 3 | 50.8 | 15 | 457 | 8.6 | 1 | 1 | 1 | | | | 1 | 1 | 1 | | | |
| High | Q92896 | Golgi apparatus protein 1 OS=Homo sapiens OX=9606 GN=GLG1 PE=1 SV=2 | 8 | 4 | 4 | 134.5 | 3 | 1179 | 6.9 | | | | | 2 | | | | | | 2 | |
| High | Q12857 | Nuclear factor 1 A-type OS=Homo sapiens OX=9606 GN=NFIA PE=1 SV=2 | 8 | 2 | 1 | 55.9 | 4 | 509 | 8.44 | 1 | | 1 | 1 | 1 | 1 | 1 | | 1 | 1 | 1 | 1 |
| High | Q9UI10 | Translation initiation factor eIF-2B subunit delta OS=Homo sapiens OX=9606 GN=EIF2B4 PE=1 SV=2 | 8 | 3 | 3 | 57.5 | 12 | 523 | 9.38 | 1 | 2 | 2 | | | | 1 | 2 | 2 | | | |
| High | Q9H3M7 | Thioredoxin-interacting protein OS=Homo sapiens OX=9606 GN=TXNIP PE=1 SV=1 | 8 | 5 | 5 | 43.6 | 22 | 391 | 7.5 | | 2 | 3 | | | | | 2 | 2 | | | |
| High | Q9BRD0 | BUD13 homolog OS=Homo sapiens OX=9606 GN=BUD13 PE=1 SV=1 | 8 | 2 | 2 | 70.5 | 3 | 619 | 9.86 | 1 | 1 | | | 1 | 1 | 1 | 1 | | | 1 | 1 |
| High | Q9NZL4 | Hsp70-binding protein 1 OS=Homo sapiens OX=9606 GN=HSPBP1 PE=1 SV=2 | 8 | 2 | 2 | 39.3 | 9 | 359 | 5.21 | | | | 1 | | 1 | | | | 1 | | 1 |
| High | Q9NV88 | Integrator complex subunit 9 OS=Homo sapiens OX=9606 GN=INTS9 PE=1 SV=2 | 8 | 5 | 5 | 73.8 | 12 | 658 | 6.33 | 3 | 2 | 3 | | | | 3 | 2 | 3 | | | |
| High | P04258 | SWISS-PROT:P04258 (Bos taurus) Similar to Collagen alpha 1(III) chain | 8 | 3 | 3 | 138.4 | 3 | 1466 | 6.43 | | | | 3 | 2 | 3 | | | | 2 | 2 | 2 |
| High | Q6IA69 | Glutamine-dependent NAD(+) synthetase OS=Homo sapiens OX=9606 GN=NADSYN1 PE=1 SV=3 | 8 | 2 | 2 | 79.2 | 4 | 706 | 6.44 | 1 | 1 | 1 | 1 | 2 | 1 | 1 | 1 | 1 | 1 | 2 | 1 |
| High | Q9BTE6 | Alanyl-tRNA editing protein Aarsd1 OS=Homo sapiens OX=9606 GN=AARSD1 PE=1 SV=2 | 8 | 2 | 2 | 45.5 | 8 | 412 | 6.42 | 1 | 1 | | 1 | | 1 | 1 | 1 | | 1 | | 1 |
| High | Q5VTU8 | ATP synthase subunit epsilon-like protein, mitochondrial OS=Homo sapiens OX=9606 GN=ATP5F1EP2 PE=1 SV=1 | 8 | 1 | 1 | 5.8 | 16 | 51 | 10.14 | 1 | 1 | | 1 | 1 | 1 | 1 | 1 | | 1 | 1 | 1 |
| High | P13073 | Cytochrome c oxidase subunit 4 isoform 1, mitochondrial OS=Homo sapiens OX=9606 GN=COX4I1 PE=1 SV=1 | 8 | 3 | 3 | 19.6 | 19 | 169 | 9.51 | 1 | 1 | 1 | | | | 1 | 1 | 1 | | | |
| High | Q9P0J0 | NADH dehydrogenase [ubiquinone] 1 alpha subcomplex subunit 13 OS=Homo sapiens OX=9606 GN=NDUFA13 PE=1 SV=3 | 8 | 3 | 3 | 16.7 | 21 | 144 | 8.43 | 2 | 3 | 2 | | | | 2 | 3 | 2 | | | |
| High | P32320 | Cytidine deaminase OS=Homo sapiens OX=9606 GN=CDA PE=1 SV=2 | 8 | 4 | 4 | 16.2 | 55 | 146 | 6.92 | | | | 1 | | | | | | 1 | | |
| High | P14635 | G2/mitotic-specific cyclin-B1 OS=Homo sapiens OX=9606 GN=CCNB1 PE=1 SV=1 | 8 | 3 | 3 | 48.3 | 8 | 433 | 7.47 | | | | 2 | 2 | 2 | | | | 2 | 2 | 2 |
| High | O75569 | Interferon-inducible double-stranded RNA-dependent protein kinase activator A OS=Homo sapiens OX=9606 GN=PRKRA PE=1 SV= | 8 | 3 | 3 | 34.4 | 13 | 313 | 8.41 | | | 1 | | 1 | | | | 1 | | 1 | |
| High | Q5JTH9 | RRP12-like protein OS=Homo sapiens OX=9606 GN=RRP12 PE=1 SV=2 | 8 | 4 | 4 | 143.6 | 4 | 1297 | 8.75 | | | 1 | 1 | | 1 | | | 1 | 1 | | 1 |
| High | P20936 | Ras GTPase-activating protein 1 OS=Homo sapiens OX=9606 GN=RASA1 PE=1 SV=1 | 8 | 3 | 3 | 116.3 | 3 | 1047 | 6.54 | | | 1 | 1 | 1 | | | | 1 | 1 | 1 | |
| High | Q96EY8 | Corrinoid adenosyltransferase OS=Homo sapiens OX=9606 GN=MMAB PE=1 SV=1 | 8 | 3 | 3 | 27.4 | 18 | 250 | 8.6 | 1 | 1 | 1 | | 1 | | 1 | 1 | 1 | | 1 | |
| High | O43291 | Kunitz-type protease inhibitor 2 OS=Homo sapiens OX=9606 GN=SPINT2 PE=1 SV=2 | 8 | 3 | 3 | 28.2 | 13 | 252 | 8.29 | | 1 | 1 | | | | | 1 | 1 | | | |
| High | O75419 | Cell division control protein 45 homolog OS=Homo sapiens OX=9606 GN=CDC45 PE=1 SV=1 | 8 | 2 | 2 | 65.5 | 5 | 566 | 5.53 | 1 | 2 | 1 | | 1 | | 1 | 2 | 1 | | 1 | |
| High | Q9UDW1 | Cytochrome b-c1 complex subunit 9 OS=Homo sapiens OX=9606 GN=UQCR10 PE=1 SV=3 | 8 | 2 | 2 | 7.3 | 38 | 63 | 9.47 | 2 | 2 | 2 | | 1 | | 2 | 2 | 2 | | 1 | |
| High | P54577 | Tyrosine--tRNA ligase, cytoplasmic OS=Homo sapiens OX=9606 GN=YARS1 PE=1 SV=4 | 8 | 5 | 5 | 59.1 | 9 | 528 | 7.05 | 2 | 1 | 3 | | | | 2 | 1 | 3 | | | |
| High | Q96MF7 | E3 SUMO-protein ligase NSE2 OS=Homo sapiens OX=9606 GN=NSMCE2 PE=1 SV=2 | 8 | 3 | 3 | 27.9 | 15 | 247 | 7.74 | | | | | 1 | | | | | | 1 | |
| High | Q6P2E9 | Enhancer of mRNA-decapping protein 4 OS=Homo sapiens OX=9606 GN=EDC4 PE=1 SV=1 | 8 | 3 | 3 | 151.6 | 2 | 1401 | 5.86 | | | 2 | 1 | | 1 | | | 2 | 1 | | 1 |
| High | O94915 | Protein furry homolog-like OS=Homo sapiens OX=9606 GN=FRYL PE=1 SV=2 | 8 | 2 | 2 | 339.4 | 1 | 3013 | 5.58 | 1 | | | | 1 | | 1 | | | | 1 | |
| High | Q96KP4 | Cytosolic non-specific dipeptidase OS=Homo sapiens OX=9606 GN=CNDP2 PE=1 SV=2 | 8 | 3 | 3 | 52.8 | 8 | 475 | 5.97 | 1 | | | | | | 1 | | | | | |
| High | Q9NVN8 | Guanine nucleotide-binding protein-like 3-like protein OS=Homo sapiens OX=9606 GN=GNL3L PE=1 SV=1 | 8 | 2 | 1 | 65.5 | 5 | 582 | 8.44 | | 1 | 1 | | 1 | | | 1 | 1 | | 1 | |
| High | Q9UNL2 | Translocon-associated protein subunit gamma OS=Homo sapiens OX=9606 GN=SSR3 PE=1 SV=1 | 8 | 1 | 1 | 21.1 | 8 | 185 | 9.61 | 1 | 1 | 1 | | 2 | | 1 | 1 | 1 | | 1 | |
| High | Q9BTC8 | Metastasis-associated protein MTA3 OS=Homo sapiens OX=9606 GN=MTA3 PE=1 SV=2 | 8 | 4 | 3 | 67.5 | 7 | 594 | 8.57 | | | | 1 | | 1 | | | | 1 | | 1 |
| High | P50453 | Serpin B9 OS=Homo sapiens OX=9606 GN=SERPINB9 PE=1 SV=1 | 8 | 4 | 3 | 42.4 | 13 | 376 | 5.86 | 1 | 1 | 2 | | 1 | 2 | 1 | 1 | 2 | | 1 | 2 |
| High | Q9UKK6 | NTF2-related export protein 1 OS=Homo sapiens OX=9606 GN=NXT1 PE=1 SV=1 | 8 | 3 | 3 | 15.8 | 25 | 140 | 5.03 | | | | | 1 | | | | | | 1 | |
| High | Q68CR1 | Protein sel-1 homolog 3 OS=Homo sapiens OX=9606 GN=SEL1L3 PE=1 SV=2 | 8 | 3 | 3 | 128.5 | 5 | 1132 | 7.28 | 1 | 2 | 2 | | 1 | | 1 | 2 | 2 | | 1 | |
| High | Q9BY89 | Uncharacterized protein KIAA1671 OS=Homo sapiens OX=9606 GN=KIAA1671 PE=1 SV=2 | 8 | 2 | 2 | 196.6 | 2 | 1806 | 8.47 | 2 | | 2 | | | | 2 | | 2 | | | |
| High | Q6KC79 | Nipped-B-like protein OS=Homo sapiens OX=9606 GN=NIPBL PE=1 SV=2 | 8 | 5 | 5 | 315.9 | 2 | 2804 | 7.91 | 2 | 1 | 1 | | | | 2 | 1 | 1 | | | |
| High | Q8NFH4 | Nucleoporin Nup37 OS=Homo sapiens OX=9606 GN=NUP37 PE=1 SV=1 | 8 | 3 | 3 | 36.7 | 11 | 326 | 5.92 | | 1 | | | | | | 1 | | | | |
| High | P30876 | DNA-directed RNA polymerase II subunit RPB2 OS=Homo sapiens OX=9606 GN=POLR2B PE=1 SV=1 | 8 | 3 | 3 | 133.8 | 4 | 1174 | 6.87 | | | 1 | | 1 | 1 | | | 1 | | 1 | 1 |
| High | Q9NVX7 | Kelch repeat and BTB domain-containing protein 4 OS=Homo sapiens OX=9606 GN=KBTBD4 PE=1 SV=3 | 8 | 3 | 3 | 58.1 | 5 | 518 | 5.67 | 1 | 2 | 1 | | | | 1 | 2 | 1 | | | |
| High | Q9NYU2 | UDP-glucose:glycoprotein glucosyltransferase 1 OS=Homo sapiens OX=9606 GN=UGGT1 PE=1 SV=3 | 8 | 4 | 4 | 177.1 | 3 | 1555 | 5.63 | | | 1 | | 1 | | | | 1 | | 1 | |
| High | Q15003 | Condensin complex subunit 2 OS=Homo sapiens OX=9606 GN=NCAPH PE=1 SV=3 | 8 | 3 | 3 | 82.5 | 5 | 741 | 5.06 | | | | 1 | 2 | 2 | | | | 1 | 2 | 2 |
| High | O00442 | RNA 3'-terminal phosphate cyclase OS=Homo sapiens OX=9606 GN=RTCA PE=1 SV=1 | 8 | 2 | 2 | 39.3 | 5 | 366 | 7.85 | | | | 1 | 1 | 1 | | | | 1 | 1 | 1 |
| High | Q7L1Q6 | eIF5-mimic protein 2 OS=Homo sapiens OX=9606 GN=BZW1 PE=1 SV=1 | 8 | 2 | 1 | 48 | 4 | 419 | 5.92 | 2 | 2 | 2 | | | 2 | 2 | 2 | 2 | | | 2 |
| High | Q9UD71 | Protein phosphatase 1 regulatory subunit 1B OS=Homo sapiens OX=9606 GN=PPP1R1B PE=1 SV=2 | 8 | 2 | 2 | 22.9 | 23 | 204 | 4.53 | 1 | 1 | 1 | | | | 1 | 1 | 1 | | | |
| High | Q9BST9 | Rhotekin OS=Homo sapiens OX=9606 GN=RTKN PE=1 SV=2 | 8 | 2 | 2 | 62.6 | 4 | 563 | 7.44 | 2 | 1 | 1 | 1 | | 1 | 2 | 1 | 1 | 1 | | 1 |
| High | P14324 | Farnesyl pyrophosphate synthase OS=Homo sapiens OX=9606 GN=FDPS PE=1 SV=4 | 8 | 3 | 3 | 48.2 | 9 | 419 | 6.15 | | | 1 | 1 | | | | | 1 | 1 | | |
| High | Q7Z5L9 | Interferon regulatory factor 2-binding protein 2 OS=Homo sapiens OX=9606 GN=IRF2BP2 PE=1 SV=2 | 8 | 3 | 3 | 61 | 8 | 587 | 8.69 | | 1 | 2 | | 1 | | | 1 | 2 | | 1 | |
| High | P30049 | ATP synthase subunit delta, mitochondrial OS=Homo sapiens OX=9606 GN=ATP5F1D PE=1 SV=2 | 8 | 1 | 1 | 17.5 | 5 | 168 | 5.49 | | 1 | 1 | 1 | 1 | 1 | | 1 | 1 | 1 | 1 | 1 |
| High | Q9UBR2 | Cathepsin Z OS=Homo sapiens OX=9606 GN=CTSZ PE=1 SV=1 | 8 | 3 | 3 | 33.8 | 13 | 303 | 7.11 | 1 | 1 | | | 1 | 1 | 1 | 1 | | | 1 | 1 |
| High | Q9BQ75 | Protein CMSS1 OS=Homo sapiens OX=9606 GN=CMSS1 PE=1 SV=2 | 8 | 6 | 6 | 31.9 | 21 | 279 | 9.19 | | | | | | | | | | | | |
| High | P50213 | Isocitrate dehydrogenase [NAD] subunit alpha, mitochondrial OS=Homo sapiens OX=9606 GN=IDH3A PE=1 SV=1 | 8 | 3 | 3 | 39.6 | 13 | 366 | 6.92 | 1 | 1 | 3 | 1 | | 2 | 1 | 1 | 3 | 1 | | 2 |
| High | Q15738 | Sterol-4-alpha-carboxylate 3-dehydrogenase, decarboxylating OS=Homo sapiens OX=9606 GN=NSDHL PE=1 SV=2 | 8 | 5 | 5 | 41.9 | 19 | 373 | 8.06 | 3 | 3 | 1 | | | | 3 | 3 | 1 | | | |
| High | P42226 | Signal transducer and activator of transcription 6 OS=Homo sapiens OX=9606 GN=STAT6 PE=1 SV=1 | 8 | 2 | 2 | 94.1 | 3 | 847 | 6.23 | 1 | 1 | 1 | | | 1 | 1 | 1 | 1 | | | 1 |
| High | P60983 | Glia maturation factor beta OS=Homo sapiens OX=9606 GN=GMFB PE=1 SV=2 | 8 | 2 | 2 | 16.7 | 11 | 142 | 5.29 | | | 1 | 2 | 2 | 2 | | | 1 | 2 | 2 | 2 |
| High | Q9Y5Q9 | General transcription factor 3C polypeptide 3 OS=Homo sapiens OX=9606 GN=GTF3C3 PE=1 SV=1 | 8 | 4 | 4 | 101.2 | 8 | 886 | 5.07 | | | | | | 1 | | | | | | 1 |
| High | Q9NSK0 | Kinesin light chain 4 OS=Homo sapiens OX=9606 GN=KLC4 PE=1 SV=3 | 7 | 3 | 1 | 68.6 | 4 | 619 | 6.18 | 1 | 1 | 1 | 1 | 1 | | 1 | 1 | 1 | 1 | 1 | |
| High | Q9UID3 | Vacuolar protein sorting-associated protein 51 homolog OS=Homo sapiens OX=9606 GN=VPS51 PE=1 SV=2 | 7 | 2 | 2 | 86 | 3 | 782 | 6.47 | 1 | 1 | 2 | | | | 1 | 1 | 2 | | | |
| High | P35270 | Sepiapterin reductase OS=Homo sapiens OX=9606 GN=SPR PE=1 SV=1 | 7 | 4 | 4 | 28 | 21 | 261 | 8.05 | | 1 | | 1 | | 1 | | 1 | | 1 | | 1 |
| High | O15260 | Surfeit locus protein 4 OS=Homo sapiens OX=9606 GN=SURF4 PE=1 SV=3 | 7 | 2 | 2 | 30.4 | 9 | 269 | 7.78 | 1 | | 2 | | 1 | | 1 | | 1 | | 1 | |
| High | O75348 | V-type proton ATPase subunit G 1 OS=Homo sapiens OX=9606 GN=ATP6V1G1 PE=1 SV=3 | 7 | 2 | 2 | 13.7 | 21 | 118 | 8.79 | 1 | | 1 | | 2 | | 1 | | 1 | | 1 | |
| High | Q9UBP9 | PTB domain-containing engulfment adapter protein 1 OS=Homo sapiens OX=9606 GN=GULP1 PE=1 SV=1 | 7 | 5 | 5 | 34.5 | 17 | 304 | 7.9 | 1 | 3 | 1 | | | | 1 | 3 | 1 | | | |
| High | P18206 | Vinculin OS=Homo sapiens OX=9606 GN=VCL PE=1 SV=4 | 7 | 2 | 2 | 123.7 | 3 | 1134 | 5.66 | | 1 | 1 | | 2 | 2 | | 1 | 1 | | 2 | 2 |
| High | Q8N465 | D-2-hydroxyglutarate dehydrogenase, mitochondrial OS=Homo sapiens OX=9606 GN=D2HGDH PE=1 SV=3 | 7 | 3 | 3 | 56.4 | 8 | 521 | 7.99 | | | | | | | | | | | | |
| High | Q8NFC6 | Biorientation of chromosomes in cell division protein 1-like 1 OS=Homo sapiens OX=9606 GN=BOD1L1 PE=1 SV=2 | 7 | 2 | 2 | 330.3 | 1 | 3051 | 5.08 | | | | 1 | 2 | 1 | | | | 1 | 2 | 1 |
| High | Q92576 | PHD finger protein 3 OS=Homo sapiens OX=9606 GN=PHF3 PE=1 SV=3 | 7 | 2 | 1 | 229.3 | 1 | 2039 | 6.96 | | | | 1 | 1 | 1 | | | | 1 | 1 | 1 |
| High | Q9Y2H1 | Serine/threonine-protein kinase 38-like OS=Homo sapiens OX=9606 GN=STK38L PE=1 SV=3 | 7 | 3 | 1 | 54 | 8 | 464 | 6.81 | | 1 | 1 | 1 | | | | 1 | 1 | 1 | | |
| High | P25815 | Protein S100-P OS=Homo sapiens OX=9606 GN=S100P PE=1 SV=2 | 7 | 1 | 1 | 10.4 | 14 | 95 | 4.88 | | | 1 | 1 | 1 | 1 | | | 1 | 1 | 1 | 1 |
| High | Q9P107 | GEM-interacting protein OS=Homo sapiens OX=9606 GN=GMIP PE=1 SV=2 | 7 | 4 | 4 | 106.6 | 6 | 970 | 5.72 | | | | | | | | | | | | |
| High | Q93034 | Cullin-5 OS=Homo sapiens OX=9606 GN=CUL5 PE=1 SV=4 | 7 | 3 | 3 | 90.9 | 4 | 780 | 7.94 | 1 | | | 1 | 1 | 1 | 1 | | | 1 | 1 | 1 |
| High | Q9NQX3 | Gephyrin OS=Homo sapiens OX=9606 GN=GPHN PE=1 SV=1 | 7 | 3 | 3 | 79.7 | 4 | 736 | 5.43 | | | | | | 1 | | | | | | 1 |
| High | Q12824 | SWI/SNF-related matrix-associated actin-dependent regulator of chromatin subfamily B member 1 OS=Homo sapiens OX=9606 GI | 7 | 4 | 4 | 44.1 | 18 | 385 | 6.23 | | | | | 1 | | | | | | 1 | |
| High | Q9NXA8 | NAD-dependent protein deacylase sirtuin-5, mitochondrial OS=Homo sapiens OX=9606 GN=SIRT5 PE=1 SV=2 | 7 | 2 | 2 | 33.9 | 9 | 310 | 8.47 | 1 | 1 | 1 | 1 | | | 1 | 1 | 1 | 1 | | |
| High | O43399 | Tumor protein D54 OS=Homo sapiens OX=9606 GN=TPD52L2 PE=1 SV=2 | 7 | 1 | 1 | 22.2 | 4 | 206 | 5.36 | 1 | 1 | 1 | 1 | 1 | | 1 | 1 | 1 | 1 | 1 | |
| High | Q92947 | Glutaryl-CoA dehydrogenase, mitochondrial OS=Homo sapiens OX=9606 GN=GCDH PE=1 SV=1 | 7 | 3 | 3 | 48.1 | 13 | 438 | 8.06 | | 2 | 3 | | | | | 2 | 3 | | | |
| High | Q9BQ70 | Transcription factor 25 OS=Homo sapiens OX=9606 GN=TCF25 PE=1 SV=1 | 7 | 2 | 2 | 76.6 | 4 | 676 | 6.35 | | | | | | 1 | | | | | | 1 |
| High | Q9Y6I3 | Epsin-1 OS=Homo sapiens OX=9606 GN=EPN1 PE=1 SV=2 | 7 | 5 | 5 | 60.3 | 17 | 576 | 4.83 | | | | | | | | | | | | |
| High | Q9BQ69 | ADP-ribose glycohydrolase MACROD1 OS=Homo sapiens OX=9606 GN=MACROD1 PE=1 SV=2 | 7 | 2 | 2 | 35.5 | 10 | 325 | 9.51 | | | 1 | | 1 | | | | 1 | | 1 | |
| High | P57088 | Transmembrane protein 33 OS=Homo sapiens OX=9606 GN=TMEM33 PE=1 SV=1 | 7 | 2 | 2 | 28 | 9 | 247 | 9.7 | 2 | 2 | 1 | | 1 | | 2 | 2 | 1 | | 1 | |
| High | Q8TBC4 | NEDD8-activating enzyme E1 catalytic subunit OS=Homo sapiens OX=9606 GN=UBA3 PE=1 SV=2 | 7 | 4 | 4 | 51.8 | 14 | 463 | 5.45 | | 1 | 1 | 1 | | 2 | | 1 | 1 | 1 | | 2 |

| | | | | | | | | | | | | | | | | | | | | | |
|---|---|---|---|---|---|---|---|---|---|---|---|---|---|---|---|---|---|---|---|---|---|
| High | Q9H857 | 5'-nucleotidase domain-containing protein 2 OS=Homo sapiens OX=9606 GN=NT5DC2 PE=1 SV=1 | 7 | 4 | 4 | 60.7 | 8 | 520 | 6.77 | 1 | 1 | 1 | | | | 1 | 1 | 1 | | | |
| High | Q9Y6M9 | NADH dehydrogenase [ubiquinone] 1 beta subcomplex subunit 9 OS=Homo sapiens OX=9606 GN=NDUFB9 PE=1 SV=3 | 7 | 2 | 2 | 21.8 | 18 | 179 | 8.38 | 2 | 2 | 2 | | | | 2 | 2 | 2 | | | |
| High | Q9Y4F5 | Centrosomal protein of 170 kDa protein B OS=Homo sapiens OX=9606 GN=CEP170B PE=1 SV=4 | 7 | 3 | 3 | 171.6 | 3 | 1589 | 6.84 | | | | | | | | | | | | |
| High | Q14145 | Kelch-like ECH-associated protein 1 OS=Homo sapiens OX=9606 GN=KEAP1 PE=1 SV=2 | 7 | 3 | 3 | 69.6 | 5 | 624 | 6.44 | | | | 2 | 3 | 2 | | | | 2 | 3 | 2 |
| High | P49419 | Alpha-aminoadipic semialdehyde dehydrogenase OS=Homo sapiens OX=9606 GN=ALDH7A1 PE=1 SV=5 | 7 | 2 | 2 | 58.5 | 7 | 539 | 7.99 | | 1 | | | 1 | 1 | | 1 | | | 1 | 1 |
| High | Q9H788 | SH2 domain-containing protein 4A OS=Homo sapiens OX=9606 GN=SH2D4A PE=1 SV=1 | 7 | 1 | 1 | 52.7 | 3 | 454 | 8.06 | | | 1 | 1 | 1 | 1 | | | 1 | 1 | 1 | 1 |
| High | Q96D46 | 60S ribosomal export protein NMD3 OS=Homo sapiens OX=9606 GN=NMD3 PE=1 SV=1 | 7 | 3 | 3 | 57.6 | 8 | 503 | 7.14 | 1 | | | 1 | 1 | 1 | 1 | | | 1 | 1 | 1 |
| High | Q9NV56 | MRG/MORF4L-binding protein OS=Homo sapiens OX=9606 GN=MRGBP PE=1 SV=1 | 7 | 1 | 1 | 22.4 | 7 | 204 | 5.83 | | 1 | 1 | 1 | | 1 | | 1 | 1 | 1 | | 1 |
| High | Q8N1G0 | Zinc finger protein 687 OS=Homo sapiens OX=9606 GN=ZNF687 PE=1 SV=1 | 7 | 3 | 3 | 129.4 | 5 | 1237 | 8.19 | | | | 1 | | 1 | | | | 1 | | 1 |
| High | Q9UKU7 | Isobutyryl-CoA dehydrogenase, mitochondrial OS=Homo sapiens OX=9606 GN=ACAD8 PE=1 SV=1 | 7 | 2 | 2 | 45 | 4 | 415 | 7.85 | 1 | 1 | 1 | | | | 1 | 1 | 1 | | | |
| High | O43414 | ERI1 exoribonuclease 3 OS=Homo sapiens OX=9606 GN=ERI3 PE=1 SV=2 | 7 | 4 | 4 | 37.2 | 14 | 337 | 8.07 | 1 | 1 | | 1 | 2 | | 1 | 1 | | 1 | 1 | |
| High | P62308 | Small nuclear ribonucleoprotein G OS=Homo sapiens OX=9606 GN=SNRPG PE=1 SV=1 | 7 | 2 | 2 | 8.5 | 25 | 76 | 8.88 | | 2 | 1 | 1 | 1 | | | 2 | 1 | 1 | 1 | |
| High | P26440 | Isovaleryl-CoA dehydrogenase, mitochondrial OS=Homo sapiens OX=9606 GN=IVD PE=1 SV=2 | 7 | 1 | 1 | 46.6 | 2 | 426 | 8.05 | 1 | 1 | 1 | | 1 | | 1 | 1 | 1 | | 1 | |
| High | P11234 | Ras-related protein Ral-B OS=Homo sapiens OX=9606 GN=RALB PE=1 SV=1 | 7 | 3 | 3 | 23.4 | 16 | 206 | 6.62 | | 1 | | | 1 | | | 1 | | | 1 | |
| High | P62841 | 40S ribosomal protein S15 OS=Homo sapiens OX=9606 GN=RPS15 PE=1 SV=2 | 7 | 1 | 1 | 17 | 9 | 145 | 10.39 | 1 | 1 | 1 | | 1 | | 1 | 1 | 1 | | 1 | |
| High | O75190 | DnaJ homolog subfamily B member 6 OS=Homo sapiens OX=9606 GN=DNAJB6 PE=1 SV=2 | 7 | 4 | 3 | 36.1 | 17 | 326 | 9.16 | 2 | 1 | 2 | | | | 2 | 1 | 2 | | | |
| High | P47736 | Rap1 GTPase-activating protein 1 OS=Homo sapiens OX=9606 GN=RAP1GAP PE=1 SV=2 | 7 | 3 | 3 | 73.3 | 6 | 663 | 5.82 | 2 | 2 | 1 | | | | 2 | 2 | 1 | | | |
| High | Q9Y3B8 | Oligoribonuclease, mitochondrial OS=Homo sapiens OX=9606 GN=REXO2 PE=1 SV=3 | 7 | 1 | 1 | 26.8 | 3 | 237 | 6.87 | 1 | 1 | | 1 | 1 | 1 | 1 | 1 | | 1 | 1 | 1 |
| High | Q9Y5J1 | U3 small nucleolar RNA-associated protein 18 homolog OS=Homo sapiens OX=9606 GN=UTP18 PE=1 SV=3 | 7 | 2 | 2 | 62 | 6 | 556 | 8.76 | 1 | 1 | 1 | | | | 1 | 1 | 1 | | | |
| High | Q06587 | E3 ubiquitin-protein ligase RING1 OS=Homo sapiens OX=9606 GN=RING1 PE=1 SV=2 | 7 | 3 | 2 | 42.4 | 11 | 406 | 5.62 | 1 | | 1 | | 1 | 2 | 1 | | 1 | | 1 | 2 |
| High | O00507 | Probable ubiquitin carboxyl-terminal hydrolase FAF-Y OS=Homo sapiens OX=9606 GN=USP9Y PE=2 SV=2 | 7 | 4 | 1 | 290.9 | 2 | 2555 | 5.86 | 1 | | 1 | | | | 1 | | 1 | | | |
| High | Q02413 | Desmoglein-1 OS=Homo sapiens OX=9606 GN=DSG1 PE=1 SV=2 | 7 | 6 | 6 | 113.7 | 8 | 1049 | 5.03 | 1 | | 5 | | 1 | | 1 | | 5 | | 1 | |
| High | Q92973 | Transportin-1 OS=Homo sapiens OX=9606 GN=TNPO1 PE=1 SV=2 | 7 | 2 | 2 | 102.3 | 2 | 898 | 4.98 | | | 1 | | | | | | 1 | | | |
| High | P57678 | Gem-associated protein 4 OS=Homo sapiens OX=9606 GN=GEMIN4 PE=1 SV=2 | 7 | 4 | 4 | 120 | 5 | 1058 | 6.04 | 2 | 1 | | | | | 2 | 1 | | | | |
| High | Q5T653 | 39S ribosomal protein L2, mitochondrial OS=Homo sapiens OX=9606 GN=MRPL2 PE=1 SV=2 | 7 | 3 | 3 | 33.3 | 14 | 305 | 11.3 | | | | 1 | 1 | 1 | | | | 1 | 1 | 1 |
| High | Q5T6V5 | Queuosine salvage protein OS=Homo sapiens OX=9606 GN=C9orf64 PE=1 SV=1 | 7 | 1 | 1 | 39 | 2 | 341 | 5.88 | 1 | 1 | 1 | | 1 | | 1 | 1 | 1 | | 1 | |
| High | P02666 | SWISS-PROT:P02666 Beta-casein - Bos taurus (Bovine). | 7 | 2 | 2 | 23.6 | 12 | 209 | 5.19 | | 1 | | | | 1 | | 1 | | | | 1 |
| High | Q9BRP8 | Partner of Y14 and mago OS=Homo sapiens OX=9606 GN=PYM1 PE=1 SV=1 | 7 | 2 | 2 | 22.6 | 14 | 204 | 9.45 | | | 1 | | | | | | 1 | | | |
| High | Q92817 | Envoplakin OS=Homo sapiens OX=9606 GN=EVPL PE=1 SV=3 | 7 | 4 | 4 | 231.5 | 3 | 2033 | 6.96 | | | | | | | | | | | | |
| High | P05114 | Non-histone chromosomal protein HMG-14 OS=Homo sapiens OX=9606 GN=HMGN1 PE=1 SV=3 | 7 | 2 | 2 | 10.7 | 16 | 100 | 9.6 | 1 | 2 | | 1 | 1 | 2 | 1 | 2 | | 1 | 1 | 2 |
| High | P25311 | Zinc-alpha-2-glycoprotein OS=Homo sapiens OX=9606 GN=AZGP1 PE=1 SV=2 | 7 | 3 | 3 | 34.2 | 12 | 298 | 6.05 | | | | 3 | 2 | 2 | | | | 3 | 2 | 2 |
| High | Q9BYD2 | 39S ribosomal protein L9, mitochondrial OS=Homo sapiens OX=9606 GN=MRPL9 PE=1 SV=2 | 7 | 2 | 2 | 30.2 | 7 | 267 | 10.08 | | | | 1 | 1 | 1 | | | | 1 | 1 | 1 |
| High | P20810 | Calpastatin OS=Homo sapiens OX=9606 GN=CAST PE=1 SV=4 | 7 | 3 | 3 | 76.5 | 9 | 708 | 5.07 | 1 | 1 | 1 | | 1 | | 1 | 1 | 1 | | 1 | |
| High | Q13188 | Serine/threonine-protein kinase 3 OS=Homo sapiens OX=9606 GN=STK3 PE=1 SV=2 | 7 | 4 | 3 | 56.3 | 12 | 491 | 5.24 | 2 | 1 | 2 | | 1 | | 2 | 1 | 2 | | 1 | |
| High | P55064 | Aquaporin-5 OS=Homo sapiens OX=9606 GN=AQP5 PE=1 SV=1 | 7 | 2 | 2 | 28.3 | 10 | 265 | 8.62 | 1 | 1 | 1 | | | | 1 | 1 | 1 | | | |
| High | Q9NVV4 | Poly(A) RNA polymerase, mitochondrial OS=Homo sapiens OX=9606 GN=MTPAP PE=1 SV=1 | 7 | 2 | 2 | 66.1 | 4 | 582 | 9.04 | 1 | 1 | 1 | | 2 | | 1 | 1 | 1 | | 2 | |
| High | Q9BXB5 | Oxysterol-binding protein-related protein 10 OS=Homo sapiens OX=9606 GN=OSBPL10 PE=1 SV=2 | 7 | 1 | 1 | 83.9 | 2 | 764 | 8.31 | | 1 | 1 | 1 | 1 | | | 1 | 1 | 1 | 1 | |
| High | Q8TB61 | Adenosine 3'-phospho 5'-phosphosulfate transporter 1 OS=Homo sapiens OX=9606 GN=SLC35B2 PE=1 SV=1 | 7 | 3 | 3 | 47.5 | 7 | 432 | 9.16 | 1 | 1 | 2 | | | 1 | 1 | 1 | 2 | | | 1 |
| High | O00273 | DNA fragmentation factor subunit alpha OS=Homo sapiens OX=9606 GN=DFFA PE=1 SV=1 | 7 | 2 | 2 | 36.5 | 9 | 331 | 4.79 | 1 | 1 | 1 | | | | 1 | 1 | 1 | | | |
| High | Q9Y5V3 | Melanoma-associated antigen D1 OS=Homo sapiens OX=9606 GN=MAGED1 PE=1 SV=3 | 7 | 3 | 2 | 86.1 | 5 | 778 | 5.83 | 1 | 3 | 2 | 1 | | | 1 | 3 | 2 | 1 | | |
| High | Q9HC52 | Chromobox protein homolog 8 OS=Homo sapiens OX=9606 GN=CBX8 PE=1 SV=3 | 7 | 2 | 2 | 43.4 | 8 | 389 | 9.91 | | | | 1 | 2 | | | | | 1 | 2 | |
| High | Q15691 | Microtubule-associated protein RP/EB family member 1 OS=Homo sapiens OX=9606 GN=MAPRE1 PE=1 SV=3 | 7 | 2 | 2 | 30 | 9 | 268 | 5.14 | 2 | 1 | | 1 | 1 | 1 | 2 | 1 | | 1 | 1 | 1 |
| High | Q9BRJ7 | Tudor-interacting repair regulator protein OS=Homo sapiens OX=9606 GN=NUDT16L1 PE=1 SV=1 | 7 | 2 | 2 | 23.3 | 13 | 211 | 8.91 | 1 | 1 | 1 | | | 1 | 1 | 1 | 1 | | | 1 |
| High | P62877 | E3 ubiquitin-protein ligase RBX1 OS=Homo sapiens OX=9606 GN=RBX1 PE=1 SV=1 | 7 | 1 | 1 | 12.3 | 7 | 108 | 6.96 | 1 | | | 1 | 1 | 1 | 1 | | | 1 | 1 | 1 |
| High | Q96LD4 | E3 ubiquitin-protein ligase TRIM47 OS=Homo sapiens OX=9606 GN=TRIM47 PE=1 SV=2 | 7 | 4 | 4 | 69.5 | 8 | 638 | 6.44 | | | | | | | | | | | | |
| High | Q68CP9 | AT-rich interactive domain-containing protein 2 OS=Homo sapiens OX=9606 GN=ARID2 PE=1 SV=2 | 7 | 2 | 2 | 197.3 | 2 | 1835 | 7.42 | 1 | | 1 | | | | 1 | | 1 | | | |
| High | P54619 | 5'-AMP-activated protein kinase subunit gamma-1 OS=Homo sapiens OX=9606 GN=PRKAG1 PE=1 SV=1 | 7 | 2 | 2 | 37.6 | 6 | 331 | 6.92 | | | 1 | 1 | 1 | 1 | | | 1 | 1 | 1 | 1 |
| High | Q9HAS0 | Protein Njmu-R1 OS=Homo sapiens OX=9606 GN=C17orf75 PE=1 SV=2 | 7 | 1 | 1 | 44.6 | 3 | 396 | 5.03 | | | 1 | 1 | 1 | 1 | | | 1 | 1 | 1 | 1 |
| High | Q07507 | Dermatopontin OS=Homo sapiens OX=9606 GN=DPT PE=1 SV=2 | 7 | 3 | 3 | 24 | 13 | 201 | 4.82 | | | | 3 | 2 | 2 | | | | 3 | 2 | 2 |
| High | Q69YN2 | CWF19-like protein 1 OS=Homo sapiens OX=9606 GN=CWF19L1 PE=1 SV=2 | 7 | 2 | 2 | 60.6 | 4 | 538 | 7.24 | 1 | | | 1 | | | 1 | | | 1 | | |
| High | O00743 | Serine/threonine-protein phosphatase 6 catalytic subunit OS=Homo sapiens OX=9606 GN=PPP6C PE=1 SV=1 | 7 | 3 | 3 | 35.1 | 12 | 305 | 5.69 | | | | 1 | 1 | | | | | 1 | 1 | |
| High | Q5VZF2 | Muscleblind-like protein 2 OS=Homo sapiens OX=9606 GN=MBNL2 PE=1 SV=2 | 7 | 2 | 2 | 40.5 | 5 | 373 | 8.38 | | | | 1 | 1 | 1 | | | | 1 | 1 | 1 |
| High | O95363 | Phenylalanine--tRNA ligase, mitochondrial OS=Homo sapiens OX=9606 GN=FARS2 PE=1 SV=1 | 7 | 2 | 2 | 52.3 | 4 | 451 | 7.46 | | | | 1 | | 2 | | | | 1 | | 2 |
| High | Q9H9J2 | 39S ribosomal protein L44, mitochondrial OS=Homo sapiens OX=9606 GN=MRPL44 PE=1 SV=1 | 7 | 3 | 3 | 37.5 | 7 | 332 | 8.4 | | | | 1 | 1 | | | | | 1 | 1 | |
| High | Q96EK6 | Glucosamine 6-phosphate N-acetyltransferase OS=Homo sapiens OX=9606 GN=GNPNAT1 PE=1 SV=1 | 7 | 4 | 4 | 20.7 | 23 | 184 | 7.99 | 1 | | | 1 | | 1 | 1 | | | 1 | | 1 |
| High | P30536 | Translocator protein OS=Homo sapiens OX=9606 GN=TSPO PE=1 SV=3 | 7 | 1 | 1 | 18.8 | 5 | 169 | 9.36 | 1 | 1 | 1 | 1 | | | 1 | 1 | 1 | 1 | | |
| High | O94842 | TOX high mobility group box family member 4 OS=Homo sapiens OX=9606 GN=TOX4 PE=1 SV=1 | 7 | 2 | 2 | 66.2 | 4 | 621 | 5.06 | | | | 2 | 2 | 1 | | | | 2 | 2 | 1 |
| High | Q9C098 | Serine/threonine-protein kinase DCLK3 OS=Homo sapiens OX=9606 GN=DCLK3 PE=2 SV=2 | 7 | 2 | 1 | 73.8 | 2 | 648 | 8.84 | 1 | 1 | 1 | | | 1 | 1 | 1 | 1 | | | 1 |
| High | Q9GZS1 | DNA-directed RNA polymerase I subunit RPA49 OS=Homo sapiens OX=9606 GN=POLR1E PE=1 SV=3 | 7 | 4 | 4 | 47.2 | 12 | 419 | 8.94 | | | | | 4 | | | | | | 3 | |
| High | P16152 | Carbonyl reductase [NADPH] 1 OS=Homo sapiens OX=9606 GN=CBR1 PE=1 SV=3 | 7 | 2 | 2 | 30.4 | 8 | 277 | 8.32 | 2 | 2 | 1 | | | | 2 | 2 | 1 | | | |
| High | Q9UPN9 | E3 ubiquitin-protein ligase TRIM33 OS=Homo sapiens OX=9606 GN=TRIM33 PE=1 SV=3 | 7 | 3 | 3 | 122.5 | 3 | 1127 | 6.67 | | | | | | 1 | | | | | | 1 |
| High | P16422 | Epithelial cell adhesion molecule OS=Homo sapiens OX=9606 GN=EPCAM PE=1 SV=2 | 7 | 2 | 2 | 34.9 | 12 | 314 | 7.46 | 1 | 1 | 1 | | | | 1 | 1 | 1 | | | |
| High | Q9Y3D0 | Cytosolic iron-sulfur assembly component 2B OS=Homo sapiens OX=9606 GN=CIAO2B PE=1 SV=1 | 7 | 1 | 1 | 17.7 | 13 | 163 | 5.19 | | | 1 | 1 | 1 | 1 | | | 1 | 1 | 1 | 1 |
| High | P82921 | 28S ribosomal protein S21, mitochondrial OS=Homo sapiens OX=9606 GN=MRPS21 PE=1 SV=3 | 7 | 3 | 3 | 10.7 | 39 | 87 | 9.92 | | 1 | | 1 | | 1 | | 1 | | 1 | | 1 |
| High | Q9UHF7 | Zinc finger transcription factor Trps1 OS=Homo sapiens OX=9606 GN=TRPS1 PE=1 SV=2 | 7 | 3 | 3 | 141.4 | 4 | 1281 | 7.58 | | | 1 | | | | | | 1 | | | |
| High | Q9P2R3 | Rabankyrin-5 OS=Homo sapiens OX=9606 GN=ANKFY1 PE=1 SV=2 | 7 | 2 | 2 | 128.3 | 1 | 1169 | 6.1 | 1 | 1 | 2 | | | | 1 | 1 | 2 | | | |
| High | Q15773 | Myeloid leukemia factor 2 OS=Homo sapiens OX=9606 GN=MLF2 PE=1 SV=1 | 7 | 1 | 1 | 28.1 | 4 | 248 | 6.9 | 1 | 1 | 1 | | 1 | 1 | 1 | 1 | 1 | | 1 | 1 |
| High | Q9Y2Q5 | Ragulator complex protein LAMTOR2 OS=Homo sapiens OX=9606 GN=LAMTOR2 PE=1 SV=1 | 7 | 2 | 2 | 13.5 | 17 | 125 | 5.4 | 1 | 1 | 1 | | | | 1 | 1 | 1 | | | |
| High | P49406 | 39S ribosomal protein L19, mitochondrial OS=Homo sapiens OX=9606 GN=MRPL19 PE=1 SV=2 | 7 | 3 | 3 | 33.5 | 14 | 292 | 9.5 | 1 | | | | 1 | | 1 | | | | 1 | |
| High | Q12802 | A-kinase anchor protein 13 OS=Homo sapiens OX=9606 GN=AKAP13 PE=1 SV=2 | 7 | 4 | 4 | 307.4 | 3 | 2813 | 5.24 | | 1 | 2 | 1 | | | | 1 | 2 | 1 | | |
| High | Q13325 | Interferon-induced protein with tetratricopeptide repeats 5 OS=Homo sapiens OX=9606 GN=IFIT5 PE=1 SV=1 | 7 | 3 | 3 | 55.8 | 10 | 482 | 7.4 | 2 | | | 1 | 1 | | 2 | | | 1 | 1 | |
| High | P20290 | Transcription factor BTF3 OS=Homo sapiens OX=9606 GN=BTF3 PE=1 SV=1 | 7 | 2 | 2 | 22.2 | 20 | 206 | 9.38 | 1 | 1 | 1 | | 2 | | 1 | 1 | 1 | | 2 | |
| High | O43396 | Thioredoxin-like protein 1 OS=Homo sapiens OX=9606 GN=TXNL1 PE=1 SV=3 | 7 | 1 | 1 | 32.2 | 6 | 289 | 4.96 | 1 | 1 | 1 | | 1 | | 1 | 1 | 1 | | 1 | |
| High | P00387 | NADH-cytochrome b5 reductase 3 OS=Homo sapiens OX=9606 GN=CYB5R3 PE=1 SV=3 | 7 | 3 | 3 | 34.2 | 14 | 301 | 7.59 | 1 | 2 | 2 | | | | 1 | 2 | 2 | | | |
| High | P10398 | Serine/threonine-protein kinase A-Raf OS=Homo sapiens OX=9606 GN=ARAF PE=1 SV=2 | 7 | 3 | 3 | 67.5 | 6 | 606 | 9.01 | 1 | 1 | | | 1 | | 1 | 1 | | | 1 | |
| High | Q9UPY8 | Microtubule-associated protein RP/EB family member 3 OS=Homo sapiens OX=9606 GN=MAPRE3 PE=1 SV=1 | 7 | 2 | 2 | 32 | 5 | 281 | 5.54 | | | | 1 | 1 | 1 | | | | 1 | 1 | 1 |
| High | Q7Z6E9 | E3 ubiquitin-protein ligase RBBP6 OS=Homo sapiens OX=9606 GN=RBBP6 PE=1 SV=1 | 7 | 4 | 4 | 201.4 | 3 | 1792 | 9.64 | | | | | 2 | | | | | | 2 | |
| High | Q9BTE3 | Mini-chromosome maintenance complex-binding protein OS=Homo sapiens OX=9606 GN=MCMBP PE=1 SV=2 | 7 | 3 | 3 | 72.9 | 5 | 642 | 5.87 | | | | 1 | 2 | 2 | | | | 1 | 2 | 2 |
| High | Q9UNA1 | Rho GTPase-activating protein 26 OS=Homo sapiens OX=9606 GN=ARHGAP26 PE=1 SV=1 | 7 | 2 | 2 | 92.2 | 3 | 814 | 6.64 | 2 | 2 | 2 | | | | 2 | 2 | 2 | | | |
| High | Q7Z3K3 | Pogo transposable element with ZNF domain OS=Homo sapiens OX=9606 GN=POGZ PE=1 SV=2 | 7 | 3 | 3 | 155.2 | 3 | 1410 | 7.4 | | 1 | | | | | | 1 | | | | |
| High | O60256 | Phosphoribosyl pyrophosphate synthase-associated protein 2 OS=Homo sapiens OX=9606 GN=PRPSAP2 PE=1 SV=1 | 7 | 2 | 1 | 40.9 | 6 | 369 | 7.44 | 1 | 1 | | | 2 | 1 | 1 | 1 | | | 2 | 1 |
| High | Q9NVS2 | 39S ribosomal protein S18a, mitochondrial OS=Homo sapiens OX=9606 GN=MRPS18A PE=1 SV=1 | 7 | 1 | 1 | 22.2 | 5 | 196 | 10.33 | 1 | | 1 | 1 | | 1 | 1 | | 1 | 1 | | 1 |
| High | Q8NEC7 | Glutathione S-transferase C-terminal domain-containing protein OS=Homo sapiens OX=9606 GN=GSTCD PE=1 SV=2 | 7 | 1 | 1 | 71 | 2 | 633 | 7.78 | | 1 | 1 | 1 | 1 | | | 1 | 1 | 1 | 1 | |
| High | O43660 | Pleiotropic regulator 1 OS=Homo sapiens OX=9606 GN=PLRG1 PE=1 SV=1 | 7 | 2 | 2 | 57.2 | 4 | 514 | 9.17 | | | 1 | | 1 | | | | 1 | | 1 | |
| High | Q9Y4E1 | WASH complex subunit 2C OS=Homo sapiens OX=9606 GN=WASHC2C PE=1 SV=4 | 7 | 3 | 3 | 144.8 | 4 | 1320 | 4.79 | | | 1 | | 1 | | | | 1 | | 1 | |
| High | Q8NEU8 | DCC-interacting protein 13-beta OS=Homo sapiens OX=9606 GN=APPL2 PE=1 SV=3 | 7 | 2 | 2 | 74.4 | 5 | 664 | 4.94 | 1 | 1 | 1 | | | | 1 | 1 | 1 | | | |
| High | P62942 | Peptidyl-prolyl cis-trans isomerase FKBP1A OS=Homo sapiens OX=9606 GN=FKBP1A PE=1 SV=2 | 7 | 3 | 3 | 11.9 | 30 | 108 | 8.16 | 3 | 2 | | | | | 3 | 2 | | | | |
| High | O00303 | Eukaryotic translation initiation factor 3 subunit F OS=Homo sapiens OX=9606 GN=EIF3F PE=1 SV=1 | 7 | 2 | 2 | 37.5 | 8 | 357 | 5.45 | | | | 1 | | 1 | | | | 1 | | 1 |
| High | Q969L2 | Protein MAL2 OS=Homo sapiens OX=9606 GN=MAL2 PE=1 SV=1 | 7 | 1 | 1 | 19.1 | 6 | 176 | 6.24 | 1 | 1 | 1 | | 1 | 1 | 1 | 1 | 1 | | 1 | 1 |
| High | Q8IUF8 | Ribosomal oxygenase 2 OS=Homo sapiens OX=9606 GN=RIOX2 PE=1 SV=1 | 7 | 2 | 2 | 52.8 | 5 | 465 | 6.7 | | 1 | | 1 | 1 | 1 | | 1 | | 1 | 1 | 1 |
| High | Q9Y3D9 | 28S ribosomal protein S23, mitochondrial OS=Homo sapiens OX=9606 GN=MRPS23 PE=1 SV=2 | 7 | 2 | 2 | 21.8 | 11 | 190 | 8.9 | | 1 | 1 | | 1 | | | 1 | 1 | | 1 | |
| High | Q66K74 | Microtubule-associated protein 1S OS=Homo sapiens OX=9606 GN=MAP1S PE=1 SV=2 | 7 | 2 | 2 | 112.1 | 4 | 1059 | 7.3 | 1 | 1 | 1 | 1 | | 1 | 1 | 1 | 1 | 1 | | 1 |
| High | Q13490 | Baculoviral IAP repeat-containing protein 2 OS=Homo sapiens OX=9606 GN=BIRC2 PE=1 SV=2 | 7 | 2 | 2 | 69.9 | 4 | 618 | 6.7 | 1 | 1 | 1 | | | | 1 | 1 | 1 | | | |
| High | Q8NC96 | Adaptin ear-binding coat-associated protein 1 OS=Homo sapiens OX=9606 GN=NECAP1 PE=1 SV=2 | 7 | 2 | 2 | 29.7 | 7 | 275 | 6.8 | | | 1 | | | | | | 1 | | | |
| High | O15235 | 28S ribosomal protein S12, mitochondrial OS=Homo sapiens OX=9606 GN=MRPS12 PE=1 SV=1 | 7 | 2 | 2 | 15.2 | 14 | 138 | 10.29 | | | 1 | 1 | 1 | 1 | | | 1 | 1 | 1 | 1 |
| High | P30519 | Heme oxygenase 2 OS=Homo sapiens OX=9606 GN=HMOX2 PE=1 SV=2 | 7 | 2 | 2 | 36 | 7 | 316 | 5.41 | 1 | 1 | 1 | | | | 1 | 1 | 1 | | | |
| High | Q14814 | Myocyte-specific enhancer factor 2D OS=Homo sapiens OX=9606 GN=MEF2D PE=1 SV=1 | 7 | 2 | 2 | 55.9 | 4 | 521 | 7.88 | 1 | | 1 | | | 1 | 1 | | 1 | | | 1 |
| High | P36776 | Lon protease homolog, mitochondrial OS=Homo sapiens OX=9606 GN=LONP1 PE=1 SV=2 | 7 | 4 | 4 | 106.4 | 5 | 959 | 6.39 | 1 | | 2 | | | | 1 | | 2 | | | |
| High | Q8IUQ4 | E3 ubiquitin-protein ligase SIAH1 OS=Homo sapiens OX=9606 GN=SIAH1 PE=1 SV=2 | 7 | 2 | 2 | 31.1 | 8 | 282 | 6.8 | 1 | | | 1 | 1 | 1 | 1 | | | 1 | 1 | 1 |
| High | O75844 | CAAX prenyl protease 1 homolog OS=Homo sapiens OX=9606 GN=ZMPSTE24 PE=1 SV=2 | 7 | 2 | 2 | 54.8 | 6 | 475 | 7.49 | 1 | 1 | 2 | | | | 1 | 1 | 2 | | | |
| High | O00764 | Pyridoxal kinase OS=Homo sapiens OX=9606 GN=PDXK PE=1 SV=1 | 7 | 1 | 1 | 35.1 | 3 | 312 | 6.13 | 1 | 1 | 1 | | 1 | | 1 | 1 | 1 | | 1 | |
| High | Q16790 | Carbonic anhydrase 9 OS=Homo sapiens OX=9606 GN=CA9 PE=1 SV=2 | 7 | 2 | 2 | 49.7 | 7 | 459 | 4.72 | 1 | 1 | 2 | | | | 1 | 1 | 2 | | | |
| High | P99999 | Cytochrome c OS=Homo sapiens OX=9606 GN=CYCS PE=1 SV=2 | 7 | 2 | 1 | 11.7 | 18 | 105 | 9.57 | | | 2 | 2 | 1 | 2 | | | 2 | 2 | 1 | 2 |
| High | Q16644 | MAP kinase-activated protein kinase 3 OS=Homo sapiens OX=9606 GN=MAPKAPK3 PE=1 SV=1 | 7 | 3 | 3 | 43 | 7 | 382 | 7.28 | | | 1 | | 2 | 1 | | | 1 | | 2 | 1 |
| High | Q9BSJ2 | Gamma-tubulin complex component 2 OS=Homo sapiens OX=9606 GN=TUBGCP2 PE=1 SV=2 | 6 | 2 | 2 | 102.5 | 3 | 902 | 6.84 | 2 | 1 | 1 | | | | 2 | 1 | 1 | | | |
| High | Q92538 | Golgi-specific brefeldin A-resistance guanine nucleotide exchange factor 1 OS=Homo sapiens OX=9606 GN=GBF1 PE=1 SV=2 | 6 | 4 | 4 | 206.3 | 2 | 1859 | 5.73 | 2 | 2 | 2 | | | | 2 | 2 | 2 | | | |
| High | O14686 | Histone-lysine N-methyltransferase 2D OS=Homo sapiens OX=9606 GN=KMT2D PE=1 SV=2 | 6 | 3 | 3 | 593 | 1 | 5537 | 5.58 | | | | | 2 | | | | | | 2 | |
| High | Q14C86 | GTPase-activating protein and VPS9 domain-containing protein 1 OS=Homo sapiens OX=9606 GN=GAPVD1 PE=1 SV=2 | 6 | 3 | 3 | 164.9 | 3 | 1478 | 5.22 | | 2 | 1 | | | | | 2 | 1 | | | |
| High | O14965 | Aurora kinase A OS=Homo sapiens OX=9606 GN=AURKA PE=1 SV=3 | 6 | 3 | 3 | 45.8 | 10 | 403 | 9.39 | | | | | 2 | 1 | | | | | 2 | 1 |
| High | Q16718 | NADH dehydrogenase [ubiquinone] 1 alpha subcomplex subunit 5 OS=Homo sapiens OX=9606 GN=NDUFA5 PE=1 SV=3 | 6 | 1 | 1 | 13.5 | 8 | 116 | 5.99 | | | 1 | 1 | 1 | 1 | | | 1 | 1 | 1 | 1 |
| High | P18583 | Protein SON OS=Homo sapiens OX=9606 GN=SON PE=1 SV=4 | 6 | 3 | 3 | 263.7 | 3 | 2426 | 5.64 | | | | 1 | 3 | 1 | | | | 1 | 2 | 1 |
| High | Q9NP77 | RNA polymerase II subunit A C-terminal domain phosphatase SSU72 OS=Homo sapiens OX=9606 GN=SSU72 PE=1 SV=1 | 6 | 1 | 1 | 22.6 | 6 | 194 | 5.33 | | | | 1 | 1 | 1 | | | | 1 | 1 | 1 |
| High | Q6ZW49 | PAX-interacting protein 1 OS=Homo sapiens OX=9606 GN=PAXIP1 PE=1 SV=2 | 6 | 2 | 2 | 121.3 | 2 | 1069 | 6.7 | 1 | 1 | 1 | | 1 | 1 | 1 | 1 | 1 | | 1 | 1 |
| High | O00116 | Alkyldihydroxyacetonephosphate synthase, peroxisomal OS=Homo sapiens OX=9606 GN=AGPS PE=1 SV=1 | 6 | 4 | 4 | 72.9 | 9 | 658 | 7.34 | | | 1 | | | 1 | | | 1 | | | 1 |
| High | Q12959 | Disks large homolog 1 OS=Homo sapiens OX=9606 GN=DLG1 PE=1 SV=2 | 6 | 1 | 1 | 100.4 | 1 | 904 | 5.76 | | | 1 | 1 | | 1 | | | 1 | 1 | | 1 |
| High | P0CG13 | Chromosome transmission fidelity protein 8 homolog OS=Homo sapiens OX=9606 GN=CHTF8 PE=1 SV=1 | 6 | 2 | 2 | 13.3 | 17 | 121 | 8.18 | 1 | | 1 | | | | 1 | | 1 | | | |
| High | Q9Y606 | tRNA pseudouridine synthase A OS=Homo sapiens OX=9606 GN=PUS1 PE=1 SV=3 | 6 | 3 | 3 | 47.4 | 9 | 427 | 8.41 | | | | 1 | 1 | 1 | | | | 1 | 1 | 1 |
| High | Q9H4Z3 | mRNA (2'-O-methyladenosine-N(6)-)-methyltransferase OS=Homo sapiens OX=9606 GN=PCIF1 PE=1 SV=1 | 6 | 1 | 1 | 80.6 | 1 | 704 | 7.42 | 1 | | | 1 | 1 | 1 | 1 | | | 1 | 1 | 1 |
| High | Q01085 | Nucleolysin TIAR OS=Homo sapiens OX=9606 GN=TIAL1 PE=1 SV=1 | 6 | 2 | 2 | 41.6 | 8 | 375 | 7.74 | 2 | 1 | 1 | | 1 | | 2 | 1 | 1 | | 1 | |
| High | Q96EL2 | 28S ribosomal protein S24, mitochondrial OS=Homo sapiens OX=9606 GN=MRPS24 PE=1 SV=1 | 6 | 1 | 1 | 19 | 5 | 167 | 9.38 | | | | 1 | 1 | 1 | | | | 1 | 1 | 1 |
| High | P28347 | Transcriptional enhancer factor TEF-1 OS=Homo sapiens OX=9606 GN=TEAD1 PE=1 SV=2 | 6 | 1 | 1 | 47.9 | 2 | 426 | 8.15 | 1 | 1 | 1 | 1 | 1 | 1 | 1 | 1 | 1 | 1 | 1 | 1 |
| High | Q8WUF5 | RelA-associated inhibitor OS=Homo sapiens OX=9606 GN=PPP1R13L PE=1 SV=4 | 6 | 2 | 2 | 89 | 4 | 828 | 6.81 | | | | | 1 | 2 | | | | | 1 | 2 |
| High | Q14676 | Mediator of DNA damage checkpoint protein 1 OS=Homo sapiens OX=9606 GN=MDC1 PE=1 SV=3 | 6 | 3 | 3 | 226.5 | 3 | 2089 | 5.47 | 1 | 1 | 2 | | | | 1 | 1 | 2 | | | |
| High | Q8IX01 | SURP and G-patch domain-containing protein 2 OS=Homo sapiens OX=9606 GN=SUGP2 PE=1 SV=2 | 6 | 2 | 2 | 120.1 | 2 | 1082 | 7.28 | | | | 1 | 1 | | | | | 1 | 1 | |
| High | O15156 | Zinc finger and BTB domain-containing protein 7B OS=Homo sapiens OX=9606 GN=ZBTB7B PE=1 SV=2 | 6 | 1 | 1 | 58 | 2 | 539 | 5.86 | | | 1 | 1 | 1 | 1 | | | 1 | 1 | 1 | 1 |
| High | Q9UNX4 | WD repeat-containing protein 3 OS=Homo sapiens OX=9606 GN=WDR3 PE=1 SV=1 | 6 | 1 | 1 | 106 | 1 | 943 | 6.64 | 1 | | 1 | | 1 | | 1 | | 1 | | 1 | |
| High | O75676 | Ribosomal protein S6 kinase alpha-4 OS=Homo sapiens OX=9606 GN=RPS6KA4 PE=1 SV=1 | 6 | 4 | 4 | 85.6 | 5 | 772 | 8.28 | 2 | 1 | 2 | | | | 2 | 1 | 2 | | | |
| High | Q9UMD9 | Collagen alpha-1(XVII) chain OS=Homo sapiens OX=9606 GN=COL17A1 PE=1 SV=3 | 6 | 2 | 2 | 150.3 | 1 | 1497 | 8.79 | | | | | | 1 | | | | | | 1 |
| High | Q14728 | Major facilitator superfamily domain-containing protein 10 OS=Homo sapiens OX=9606 GN=MFSD10 PE=1 SV=1 | 6 | 1 | 1 | 48.3 | 3 | 455 | 9.6 | 1 | 1 | 1 | | | | 1 | 1 | 1 | | | |
| High | O14617 | AP-3 complex subunit delta-1 OS=Homo sapiens OX=9606 GN=AP3D1 PE=1 SV=1 | 6 | 4 | 4 | 130.1 | 5 | 1153 | 8.48 | | | | | | | | | | | | |
| High | O43251 | RNA binding protein fox-1 homolog 2 OS=Homo sapiens OX=9606 GN=RBFOX2 PE=1 SV=3 | 6 | 2 | 2 | 41.3 | 6 | 390 | 7.27 | | 1 | | 1 | | 1 | | 1 | | 1 | | 1 |
| High | O75390 | Citrate synthase, mitochondrial OS=Homo sapiens OX=9606 GN=CS PE=1 SV=2 | 6 | 2 | 2 | 51.7 | 5 | 466 | 8.32 | | | | 1 | 2 | 1 | | | | 1 | 2 | 1 |
| High | O95456 | Proteasome assembly chaperone 1 OS=Homo sapiens OX=9606 GN=PSMG1 PE=1 SV=1 | 6 | 2 | 2 | 32.8 | 10 | 288 | 7.17 | | 2 | 1 | | | 1 | | 2 | 1 | | | 1 |
| High | P31150 | Rab GDP dissociation inhibitor alpha OS=Homo sapiens OX=9606 GN=GDI1 PE=1 SV=2 | 6 | 2 | 1 | 50.6 | 6 | 447 | 5.14 | 1 | 1 | 2 | | | | 1 | 1 | 2 | | | |
| High | Q99496 | E3 ubiquitin-protein ligase RING2 OS=Homo sapiens OX=9606 GN=RNF2 PE=1 SV=1 | 6 | 3 | 2 | 37.6 | 14 | 336 | 6.84 | 1 | | 1 | | 1 | | 1 | | 1 | | 1 | |
| High | Q9HCN8 | Stromal cell-derived factor 2-like protein 1 OS=Homo sapiens OX=9606 GN=SDF2L1 PE=1 SV=2 | 6 | 1 | 1 | 23.6 | 9 | 221 | 7.03 | 1 | 1 | 1 | | | | 1 | 1 | 1 | | | |
| High | Q92686 | Neurogranin OS=Homo sapiens OX=9606 GN=NRGN PE=1 SV=1 | 6 | 2 | 2 | 7.6 | 60 | 78 | 7.87 | | | 1 | 1 | 2 | 1 | | | 1 | 1 | 2 | 1 |
| High | P62314 | Small nuclear ribonucleoprotein Sm D1 OS=Homo sapiens OX=9606 GN=SNRPD1 PE=1 SV=1 | 6 | 1 | 1 | 13.3 | 11 | 119 | 11.56 | 1 | | | 1 | 1 | | 1 | | | 1 | 1 | |
| High | Q9NT62 | Ubiquitin-like-conjugating enzyme ATG3 OS=Homo sapiens OX=9606 GN=ATG3 PE=1 SV=1 | 6 | 2 | 2 | 35.8 | 7 | 314 | 4.74 | | | | | 1 | | | | | | 1 | |
| High | Q86V21 | Acetoacetyl-CoA synthetase OS=Homo sapiens OX=9606 GN=AACS PE=1 SV=1 | 6 | 2 | 2 | 75.1 | 3 | 672 | 6.24 | 1 | | 1 | | | | 1 | | 1 | | | |
| High | O14545 | TRAF-type zinc finger domain-containing protein 1 OS=Homo sapiens OX=9606 GN=TRAFD1 PE=1 SV=1 | 6 | 1 | 1 | 64.8 | 2 | 582 | 5.29 | 1 | 1 | | | | 1 | 1 | 1 | | | | 1 |
| High | Q14318 | Peptidyl-prolyl cis-trans isomerase FKBP8 OS=Homo sapiens OX=9606 GN=FKBP8 PE=1 SV=2 | 6 | 1 | 1 | 44.5 | 2 | 412 | 4.84 | | | | 1 | 1 | 1 | | | | 1 | 1 | 1 |
| High | Q15386 | Ubiquitin-protein ligase E3C OS=Homo sapiens OX=9606 GN=UBE3C PE=1 SV=3 | 6 | 4 | 4 | 123.8 | 4 | 1083 | 6.71 | 1 | 1 | 3 | | | | 1 | 1 | 3 | | | |
| High | P32321 | Deoxycytidylate deaminase OS=Homo sapiens OX=9606 GN=DCTD PE=1 SV=2 | 6 | 2 | 2 | 20 | 15 | 178 | 7.56 | 1 | | | | 2 | | 1 | | | | 2 | |
| High | Q5TC63 | Growth hormone-regulated TBC protein 1 OS=Homo sapiens OX=9606 GN=GRTP1 PE=1 SV=4 | 6 | 3 | 3 | 38.5 | 13 | 336 | 8.78 | 1 | 1 | 1 | | | | 1 | 1 | 1 | | | |
| High | Q9UMR2 | ATP-dependent RNA helicase DDX19B OS=Homo sapiens OX=9606 GN=DDX19B PE=1 SV=1 | 6 | 4 | 4 | 53.9 | 14 | 479 | 6.3 | | 1 | 1 | | 1 | | | 1 | 1 | | 1 | |
| High | Q9BTV4 | Transmembrane protein 43 OS=Homo sapiens OX=9606 GN=TMEM43 PE=1 SV=1 | 6 | 4 | 4 | 44.8 | 16 | 400 | 8.13 | 1 | | 1 | | | | 1 | | 1 | | | |
| High | P84074 | Neuron-specific calcium-binding protein hippocalcin OS=Homo sapiens OX=9606 GN=HPCA PE=1 SV=2 | 6 | 3 | 3 | 22.4 | 16 | 193 | 4.97 | | 1 | 1 | | | | | 1 | 1 | | | |
| High | Q9UI12 | V-type proton ATPase subunit H OS=Homo sapiens OX=9606 GN=ATP6V1H PE=1 SV=1 | 6 | 3 | 3 | 55.8 | 10 | 483 | 6.48 | 2 | 2 | 2 | | | | 2 | 2 | 2 | | | |
| High | Q9Y2R5 | 28S ribosomal protein S17, mitochondrial OS=Homo sapiens OX=9606 GN=MRPS17 PE=1 SV=1 | 6 | 2 | 2 | 14.5 | 29 | 130 | 9.85 | | | | | 1 | | | | | | 1 | |
| High | Q9BSG0 | Protease-associated domain-containing protein 1 OS=Homo sapiens OX=9606 GN=PRADC1 PE=1 SV=1 | 6 | 2 | 2 | 21 | 11 | 188 | 5.47 | 2 | 2 | 2 | | | | 2 | 2 | 2 | | | |
| High | O43318 | Mitogen-activated protein kinase kinase kinase 7 OS=Homo sapiens OX=9606 GN=MAP3K7 PE=1 SV=1 | 6 | 2 | 2 | 67.2 | 5 | 606 | 7.11 | 1 | 1 | 1 | | 1 | | 1 | 1 | 1 | | 1 | |
| High | O43324 | Eukaryotic translation elongation factor 1 epsilon-1 OS=Homo sapiens OX=9606 GN=EEF1E1 PE=1 SV=1 | 6 | 3 | 3 | 19.8 | 18 | 174 | 8.54 | | | 1 | | 1 | | | | 1 | | 1 | |
| High | P12111 | Collagen alpha-3(VI) chain OS=Homo sapiens OX=9606 GN=COL6A3 PE=1 SV=5 | 6 | 3 | 3 | 343.5 | 1 | 3177 | 6.68 | | | | 1 | 3 | 2 | | | | 1 | 3 | 2 |
| High | Q01970 | 1-phosphatidylinositol 4,5-bisphosphate phosphodiesterase beta-3 OS=Homo sapiens OX=9606 GN=PLCB3 PE=1 SV=2 | 6 | 4 | 4 | 138.7 | 4 | 1234 | 5.9 | 1 | | 1 | | | | 1 | | 1 | | | |
| High | Q9Y696 | Chloride intracellular channel protein 4 OS=Homo sapiens OX=9606 GN=CLIC4 PE=1 SV=4 | 6 | 1 | 1 | 28.8 | 5 | 253 | 5.59 | 1 | 1 | 3 | 1 | | | 1 | 1 | 1 | 1 | | |
| High | Q9BYD6 | 39S ribosomal protein L1, mitochondrial OS=Homo sapiens OX=9606 GN=MRPL1 PE=1 SV=2 | 6 | 3 | 3 | 36.9 | 14 | 325 | 8.78 | | | 1 | | | 1 | | | 1 | | | 1 |
| High | Q99570 | Phosphoinositide 3-kinase regulatory subunit 4 OS=Homo sapiens OX=9606 GN=PIK3R4 PE=1 SV=3 | 6 | 5 | 5 | 153 | 5 | 1358 | 7.17 | 1 | 2 | 2 | | | | 1 | 2 | 2 | | | |
| High | Q8N1G2 | Cap-specific mRNA (nucleoside-2'-O-)-methyltransferase 1 OS=Homo sapiens OX=9606 GN=CMTR1 PE=1 SV=1 | 6 | 4 | 4 | 95.3 | 6 | 835 | 7.05 | | 1 | 1 | 1 | 1 | | | 1 | 1 | 1 | 1 | |
| High | Q6W2J9 | BCL-6 corepressor OS=Homo sapiens OX=9606 GN=BCOR PE=1 SV=1 | 6 | 2 | 1 | 192.1 | 1 | 1755 | 6.48 | | | | 2 | 1 | 1 | | | | 2 | 1 | 1 |
| High | O95202 | Mitochondrial proton/calcium exchanger protein OS=Homo sapiens OX=9606 GN=LETM1 PE=1 SV=1 | 6 | 2 | 2 | 83.3 | 3 | 739 | 6.7 | 1 | | 1 | | | 1 | 1 | | 1 | | | 1 |
| High | P09001 | 39S ribosomal protein L3, mitochondrial OS=Homo sapiens OX=9606 GN=MRPL3 PE=1 SV=1 | 6 | 1 | 1 | 38.6 | 3 | 348 | 9.48 | | | 1 | 1 | | 1 | | | 1 | 1 | | 1 |
| High | Q13042 | Cell division cycle protein 16 homolog OS=Homo sapiens OX=9606 GN=CDC16 PE=1 SV=2 | 6 | 3 | 3 | 71.6 | 6 | 620 | 5.85 | 3 | 2 | 1 | | | | 3 | 2 | 1 | | | |
| High | Q96QC0 | Serine/threonine-protein phosphatase 1 regulatory subunit 10 OS=Homo sapiens OX=9606 GN=PPP1R10 PE=1 SV=1 | 6 | 2 | 2 | 99 | 4 | 940 | 9.17 | | | | | 1 | 1 | | | | | 1 | 1 |
| High | P56182 | Ribosomal RNA processing protein 1 homolog A OS=Homo sapiens OX=9606 GN=RRP1 PE=1 SV=1 | 6 | 1 | 1 | 52.8 | 2 | 461 | 9.33 | 1 | 1 | | 1 | | | 1 | 1 | | 1 | | |
| High | P52926 | High mobility group protein HMGI-C OS=Homo sapiens OX=9606 GN=HMGA2 PE=1 SV=1 | 6 | 2 | 2 | 11.8 | 31 | 109 | 10.62 | 2 | 1 | 1 | | 1 | | 2 | 1 | 1 | | 1 | |
| High | Q9P253 | Vacuolar protein sorting-associated protein 18 homolog OS=Homo sapiens OX=9606 GN=VPS18 PE=1 SV=2 | 6 | 3 | 3 | 110.1 | 3 | 973 | 6.07 | 1 | 2 | 1 | | | | 1 | 2 | 1 | | | |
| High | Q16851 | UTP--glucose-1-phosphate uridylyltransferase OS=Homo sapiens OX=9606 GN=UGP2 PE=1 SV=5 | 6 | 1 | 1 | 56.9 | 3 | 508 | 8.15 | 1 | 1 | 1 | | | | 1 | 1 | 1 | | | |
| High | Q04206 | Transcription factor p65 OS=Homo sapiens OX=9606 GN=RELA PE=1 SV=2 | 6 | 3 | 3 | 60.2 | 7 | 551 | 5.68 | 1 | 1 | | | 1 | 1 | 1 | 1 | | | 1 | 1 |

| | | | | | | | | | | | | | | | | | | | | | |
|---|---|---|---|---|---|---|---|---|---|---|---|---|---|---|---|---|---|---|---|---|---|
| High | O60725 | Protein-S-isoprenylcysteine O-methyltransferase OS=Homo sapiens OX=9606 GN=ICMT PE=1 SV=1 | 6 | 1 | 1 | 31.9 | 6 | 284 | 7.96 | 1 | 1 | 1 | | | | 1 | 1 | 1 | | | |
| High | Q12965 | Unconventional myosin-Ie OS=Homo sapiens OX=9606 GN=MYO1E PE=1 SV=2 | 6 | 2 | 2 | 127 | 2 | 1108 | 8.92 | 1 | 1 | 1 | | | 1 | 1 | 1 | 1 | | | 1 |
| High | Q8WUY1 | Protein THEM6 OS=Homo sapiens OX=9606 GN=THEM6 PE=1 SV=2 | 6 | 3 | 3 | 23.9 | 11 | 208 | 9.55 | | | 1 | | | | | | 1 | | | |
| High | Q96BI1 | Solute carrier family 22 member 18 OS=Homo sapiens OX=9606 GN=SLC22A18 PE=1 SV=3 | 6 | 2 | 2 | 44.8 | 4 | 424 | 9.57 | 1 | 1 | 1 | | | | 1 | 1 | 1 | | | |
| High | Q13823 | Nucleolar GTP-binding protein 2 OS=Homo sapiens OX=9606 GN=GNL2 PE=1 SV=1 | 6 | 3 | 3 | 83.6 | 6 | 731 | 9.25 | | | | | | | | | | | | |
| High | O94992 | Protein HEXIM1 OS=Homo sapiens OX=9606 GN=HEXIM1 PE=1 SV=1 | 6 | 3 | 3 | 40.6 | 11 | 359 | 4.89 | | | | 1 | 2 | 1 | | | | 1 | 2 | 1 |
| High | P49247 | Ribose-5-phosphate isomerase OS=Homo sapiens OX=9606 GN=RPIA PE=1 SV=3 | 6 | 2 | 2 | 33.2 | 8 | 311 | 8.54 | | | | 1 | | 1 | | | | 1 | | 1 |
| High | O95639 | Cleavage and polyadenylation specificity factor subunit 4 OS=Homo sapiens OX=9606 GN=CPSF4 PE=1 SV=1 | 6 | 2 | 2 | 30.2 | 15 | 269 | 8.31 | | | | 2 | 1 | 1 | | | | 2 | 1 | 1 |
| High | O75683 | Surfeit locus protein 6 OS=Homo sapiens OX=9606 GN=SURF6 PE=1 SV=3 | 6 | 2 | 2 | 41.4 | 6 | 361 | 10.64 | | | | 1 | | 1 | | | | 1 | | 1 |
| High | O14683 | Tumor protein p53-inducible protein 11 OS=Homo sapiens OX=9606 GN=TP53I11 PE=1 SV=2 | 6 | 2 | 2 | 21 | 13 | 189 | 9.55 | | | | | 1 | | | | | | 1 | |
| High | Q96CS3 | FAS-associated factor 2 OS=Homo sapiens OX=9606 GN=FAF2 PE=1 SV=2 | 6 | 3 | 3 | 52.6 | 9 | 445 | 5.62 | 2 | 2 | 2 | | | | 2 | 2 | 2 | | | |
| High | Q8WVX9 | Fatty acyl-CoA reductase 1 OS=Homo sapiens OX=9606 GN=FAR1 PE=1 SV=1 | 6 | 4 | 4 | 59.3 | 10 | 515 | 9.17 | 2 | 1 | 3 | | | | 2 | 1 | 3 | | | |
| High | P45973 | Chromobox protein homolog 5 OS=Homo sapiens OX=9606 GN=CBX5 PE=1 SV=1 | 6 | 2 | 2 | 22.2 | 13 | 191 | 5.86 | 1 | | | | 1 | 1 | 1 | | | | 1 | 1 |
| High | Q9Y6A5 | Transforming acidic coiled-coil-containing protein 3 OS=Homo sapiens OX=9606 GN=TACC3 PE=1 SV=1 | 6 | 2 | 2 | 90.3 | 2 | 838 | 5.05 | | | | 1 | 1 | | | | | 1 | 1 | |
| High | O14772 | Fucose-1-phosphate guanylyltransferase OS=Homo sapiens OX=9606 GN=FPGT PE=1 SV=3 | 6 | 2 | 2 | 68 | 3 | 607 | 6.87 | | 1 | | | | | | 1 | | | | |
| High | O00151 | PDZ and LIM domain protein 1 OS=Homo sapiens OX=9606 GN=PDLIM1 PE=1 SV=4 | 6 | 2 | 2 | 36 | 6 | 329 | 7.02 | | 1 | | | 1 | | | 1 | | | 1 | |
| High | Q01664 | Transcription factor AP-4 OS=Homo sapiens OX=9606 GN=TFAP4 PE=1 SV=2 | 6 | 3 | 3 | 38.7 | 13 | 338 | 5.87 | 2 | 2 | 2 | | | | 2 | 2 | 2 | | | |
| High | Q9UBW7 | Zinc finger MYM-type protein 2 OS=Homo sapiens OX=9606 GN=ZMYM2 PE=1 SV=1 | 6 | 4 | 4 | 154.8 | 5 | 1377 | 6.34 | | | | | 1 | | | | | | 1 | |
| High | Q9BTW9 | Tubulin-specific chaperone D OS=Homo sapiens OX=9606 GN=TBCD PE=1 SV=2 | 6 | 1 | 1 | 132.5 | 1 | 1192 | 6.19 | | 1 | 1 | | | 1 | | 1 | 1 | | | 1 |
| High | Q13488 | V-type proton ATPase 116 kDa subunit a 3 OS=Homo sapiens OX=9606 GN=TCIRG1 PE=1 SV=3 | 6 | 4 | 4 | 92.9 | 5 | 830 | 7.12 | 1 | | | | | | 1 | | | | | |
| High | P13693 | Translationally-controlled tumor protein OS=Homo sapiens OX=9606 GN=TPT1 PE=1 SV=1 | 6 | 1 | 1 | 19.6 | 8 | 172 | 4.93 | 1 | 1 | 1 | | | | 1 | 1 | 1 | | | |
| High | O75787 | Renin receptor OS=Homo sapiens OX=9606 GN=ATP6AP2 PE=1 SV=2 | 6 | 2 | 2 | 39 | 5 | 350 | 6.1 | 1 | 1 | 1 | | | | 1 | 1 | 1 | | | |
| High | Q9UBU8 | Mortality factor 4-like protein 1 OS=Homo sapiens OX=9606 GN=MORF4L1 PE=1 SV=2 | 6 | 2 | 1 | 41.4 | 6 | 362 | 9.28 | | | | 1 | 1 | 2 | | | | 1 | 1 | 2 |
| High | Q8NEV8 | Exophilin-5 OS=Homo sapiens OX=9606 GN=EXPH5 PE=1 SV=4 | 6 | 1 | 1 | 222.4 | 1 | 1989 | 7.81 | 1 | 1 | 1 | | | | 1 | 1 | 1 | | | |
| High | P35556 | Fibrillin-2 OS=Homo sapiens OX=9606 GN=FBN2 PE=1 SV=3 | 6 | 2 | 1 | 314.6 | 1 | 2912 | 4.86 | | | | 1 | 3 | 2 | | | | 1 | 2 | 2 |
| High | Q8WXE1 | ATR-interacting protein OS=Homo sapiens OX=9606 GN=ATRIP PE=1 SV=1 | 6 | 1 | 1 | 85.8 | 1 | 791 | 6.32 | | | | 1 | 1 | 1 | | | | 1 | 1 | 1 |
| High | P19838 | Nuclear factor NF-kappa-B p105 subunit OS=Homo sapiens OX=9606 GN=NFKB1 PE=1 SV=2 | 6 | 4 | 4 | 105.3 | 5 | 968 | 5.4 | | | | | | | | | | | | |
| High | Q9P2D0 | Inhibitor of Bruton tyrosine kinase OS=Homo sapiens OX=9606 GN=IBTK PE=1 SV=3 | 6 | 4 | 4 | 150.4 | 5 | 1353 | 7.71 | | | | | | | | | | | | |
| High | Q8WXH0 | Nesprin-2 OS=Homo sapiens OX=9606 GN=SYNE2 PE=1 SV=3 | 6 | 4 | 4 | 795.9 | 1 | 6885 | 5.36 | | 1 | 2 | | | | | 1 | 2 | | | |
| High | Q04656 | Copper-transporting ATPase 1 OS=Homo sapiens OX=9606 GN=ATP7A PE=1 SV=4 | 6 | 3 | 3 | 163.3 | 3 | 1500 | 6.33 | 1 | 1 | 1 | | | | 1 | 1 | 1 | | | |
| High | Q9Y6K0 | Choline/ethanolaminephosphotransferase 1 OS=Homo sapiens OX=9606 GN=CEPT1 PE=1 SV=1 | 6 | 1 | 1 | 46.5 | 2 | 416 | 8.21 | 1 | 1 | 1 | | 1 | | 1 | 1 | 1 | | 1 | |
| High | P13051 | Uracil-DNA glycosylase OS=Homo sapiens OX=9606 GN=UNG PE=1 SV=2 | 6 | 3 | 3 | 34.6 | 14 | 313 | 9.32 | | | | | 2 | | | | | | 2 | |
| High | P51570 | Galactokinase OS=Homo sapiens OX=9606 GN=GALK1 PE=1 SV=1 | 6 | 3 | 3 | 42.2 | 8 | 392 | 6.46 | 1 | 1 | 1 | | | | 1 | 1 | 1 | | | |
| High | P00167 | Cytochrome b5 OS=Homo sapiens OX=9606 GN=CYB5A PE=1 SV=2 | 6 | 2 | 2 | 15.3 | 25 | 134 | 4.96 | | | | 1 | 1 | 2 | | | | 1 | 1 | 2 |
| High | Q08378 | Golgin subfamily A member 3 OS=Homo sapiens OX=9606 GN=GOLGA3 PE=1 SV=2 | 6 | 3 | 3 | 167.3 | 3 | 1498 | 5.44 | | 1 | | | | | | 1 | | | | |
| High | Q9NQ55 | Suppressor of SWI4 1 homolog OS=Homo sapiens OX=9606 GN=PPAN PE=2 SV=1 | 6 | 2 | 2 | 53.2 | 4 | 473 | 10.13 | | | | 1 | | 1 | | | | 1 | | 1 |
| High | O00487 | 26S proteasome non-ATPase regulatory subunit 14 OS=Homo sapiens OX=9606 GN=PSMD14 PE=1 SV=1 | 6 | 1 | 1 | 34.6 | 4 | 310 | 6.52 | | | | 1 | 1 | 1 | | | | 1 | 1 | 1 |
| High | Q15427 | Splicing factor 3B subunit 4 OS=Homo sapiens OX=9606 GN=SF3B4 PE=1 SV=1 | 6 | 1 | 1 | 44.4 | 3 | 424 | 8.56 | | | | 1 | 1 | 1 | | | | 1 | 1 | 1 |
| High | Q8TBX8 | Phosphatidylinositol 5-phosphate 4-kinase type-2 gamma OS=Homo sapiens OX=9606 GN=PIP4K2C PE=1 SV=3 | 6 | 2 | 2 | 47.3 | 6 | 421 | 6.84 | 2 | 1 | 1 | | | | 2 | 1 | 1 | | | |
| High | Q7Z4G4 | tRNA (guanine(10)-N2)-methyltransferase homolog OS=Homo sapiens OX=9606 GN=TRMT11 PE=1 SV=1 | 6 | 2 | 2 | 53.4 | 3 | 463 | 7.78 | | | 1 | | | 1 | | | 1 | | | 1 |
| High | P46439 | Glutathione S-transferase Mu 5 OS=Homo sapiens OX=9606 GN=GSTM5 PE=1 SV=3 | 6 | 2 | 2 | 25.7 | 8 | 218 | 7.39 | | | | 2 | 2 | 2 | | | | 2 | 2 | 2 |
| High | P29590 | Protein PML OS=Homo sapiens OX=9606 GN=PML PE=1 SV=3 | 6 | 1 | 1 | 97.5 | 1 | 882 | 6.21 | | | 1 | 1 | | 1 | | | 1 | 1 | | 1 |
| High | Q9BSH4 | Translational activator of cytochrome c oxidase 1 OS=Homo sapiens OX=9606 GN=TACO1 PE=1 SV=1 | 6 | 2 | 2 | 32.5 | 9 | 297 | 8.13 | | 1 | | | | | | 1 | | | | |
| High | Q9ULX3 | RNA-binding protein NOB1 OS=Homo sapiens OX=9606 GN=NOB1 PE=1 SV=1 | 6 | 2 | 2 | 46.6 | 3 | 412 | 7.18 | | 1 | | 1 | | 1 | | 1 | | 1 | | 1 |
| High | P02662 | SWISS-PROT:P02662 Alpha-S1-casein - Bos taurus (Bovine). | 6 | 2 | 2 | 23 | 14 | 199 | 4.94 | | 1 | | | | | | 1 | | | | |
| High | Q96IX5 | ATP synthase membrane subunit K, mitochondrial OS=Homo sapiens OX=9606 GN=ATP5MK PE=1 SV=1 | 6 | 1 | 1 | 6.5 | 17 | 58 | 9.76 | 1 | | 1 | 1 | 1 | 1 | 1 | | 1 | 1 | 1 | 1 |
| High | Q8IXK0 | Polyhomeotic-like protein 2 OS=Homo sapiens OX=9606 GN=PHC2 PE=1 SV=1 | 6 | 3 | 3 | 90.7 | 3 | 858 | 8.69 | | | | | | | | | | | | |
| High | Q96EV2 | RNA-binding protein 33 OS=Homo sapiens OX=9606 GN=RBM33 PE=1 SV=3 | 6 | 4 | 4 | 129.9 | 6 | 1170 | 6.93 | | | | | 1 | | | | | | 1 | |
| High | O95071 | E3 ubiquitin-protein ligase UBR5 OS=Homo sapiens OX=9606 GN=UBR5 PE=1 SV=2 | 6 | 4 | 4 | 309.2 | 2 | 2799 | 5.85 | | | | | | | | | | | | |
| High | O94888 | UBX domain-containing protein 7 OS=Homo sapiens OX=9606 GN=UBXN7 PE=1 SV=2 | 6 | 1 | 1 | 54.8 | 3 | 489 | 5.16 | | | | | 2 | | | | | | 1 | |
| High | Q9NWT1 | p21-activated protein kinase-interacting protein 1 OS=Homo sapiens OX=9606 GN=PAK1IP1 PE=1 SV=2 | 6 | 2 | 2 | 43.9 | 6 | 392 | 8.91 | | | | 1 | 1 | 1 | | | | 1 | 1 | 1 |
| High | Q9H583 | HEAT repeat-containing protein 1 OS=Homo sapiens OX=9606 GN=HEATR1 PE=1 SV=3 | 6 | 4 | 4 | 242.2 | 2 | 2144 | 6.54 | 2 | 3 | 1 | | | | 2 | 3 | 1 | | | |
| High | P40692 | DNA mismatch repair protein Mlh1 OS=Homo sapiens OX=9606 GN=MLH1 PE=1 SV=1 | 6 | 2 | 2 | 84.5 | 3 | 756 | 5.72 | 1 | | 1 | | | 1 | 1 | | 1 | | | 1 |
| High | Q7Z3U7 | Protein MON2 homolog OS=Homo sapiens OX=9606 GN=MON2 PE=1 SV=3 | 6 | 3 | 3 | 190.2 | 2 | 1717 | 6.06 | 2 | 3 | 1 | | | | 2 | 3 | 1 | | | |
| High | Q05048 | Cleavage stimulation factor subunit 1 OS=Homo sapiens OX=9606 GN=CSTF1 PE=1 SV=1 | 6 | 2 | 2 | 48.3 | 6 | 431 | 6.58 | | | | 1 | 1 | 1 | | | | 1 | 1 | 1 |
| High | P40925 | Malate dehydrogenase, cytoplasmic OS=Homo sapiens OX=9606 GN=MDH1 PE=1 SV=4 | 6 | 2 | 2 | 36.4 | 9 | 334 | 7.36 | | | | 2 | | 2 | | | | 2 | | 2 |
| High | P53602 | Diphosphomevalonate decarboxylase OS=Homo sapiens OX=9606 GN=MVD PE=1 SV=1 | 6 | 3 | 3 | 43.4 | 8 | 400 | 7.23 | 1 | 1 | | 1 | 1 | | 1 | 1 | | 1 | 1 | |
| High | Q9HCM4 | Band 4.1-like protein 5 OS=Homo sapiens OX=9606 GN=EPB41L5 PE=1 SV=3 | 6 | 2 | 2 | 81.8 | 4 | 733 | 6.58 | 1 | 1 | 1 | | | | 1 | 1 | 1 | | | |
| High | Q15287 | RNA-binding protein with serine-rich domain 1 OS=Homo sapiens OX=9606 GN=RNPS1 PE=1 SV=1 | 6 | 1 | 1 | 34.2 | 3 | 305 | 11.84 | | | | 1 | 1 | 1 | | | | 1 | 1 | 1 |
| High | Q9NRX2 | 39S ribosomal protein L17, mitochondrial OS=Homo sapiens OX=9606 GN=MRPL17 PE=1 SV=1 | 6 | 1 | 1 | 20 | 4 | 175 | 10.11 | | | 1 | 1 | | 1 | | | 1 | 1 | | 1 |
| High | Q6FI81 | Anamorsin OS=Homo sapiens OX=9606 GN=CIAPIN1 PE=1 SV=2 | 6 | 3 | 3 | 33.6 | 12 | 312 | 5.62 | 1 | | | | 1 | | 1 | | | | 1 | |
| High | Q92610 | Zinc finger protein 592 OS=Homo sapiens OX=9606 GN=ZNF592 PE=1 SV=2 | 6 | 3 | 3 | 137.4 | 5 | 1267 | 7.84 | | | | | | | | | | | | |
| High | Q9UEG4 | Zinc finger protein 629 OS=Homo sapiens OX=9606 GN=ZNF629 PE=1 SV=2 | 6 | 1 | 1 | 96.6 | 2 | 869 | 7.93 | 1 | 1 | 1 | 1 | 1 | 1 | 1 | 1 | 1 | 1 | 1 | 1 |
| High | Q9NRP0 | Oligosaccharyltransferase complex subunit OSTC OS=Homo sapiens OX=9606 GN=OSTC PE=1 SV=1 | 6 | 1 | 1 | 16.8 | 8 | 149 | 9.13 | 1 | 1 | 1 | | 1 | | 1 | 1 | 1 | | 1 | |
| High | Q13330 | Metastasis-associated protein MTA1 OS=Homo sapiens OX=9606 GN=MTA1 PE=1 SV=2 | 6 | 3 | 2 | 80.7 | 6 | 715 | 9.26 | | | | 1 | | | | | | 1 | | |
| High | Q5XKP0 | MICOS complex subunit MIC13 OS=Homo sapiens OX=9606 GN=MICOS13 PE=1 SV=1 | 6 | 2 | 2 | 13.1 | 23 | 118 | 9.42 | | 1 | 1 | | | | | 1 | 1 | | | |
| High | Q9BUT9 | MAPK regulated corepressor interacting protein 2 OS=Homo sapiens OX=9606 GN=MCRIP2 PE=1 SV=2 | 6 | 1 | 1 | 17.8 | 7 | 160 | 9.41 | | | 1 | 1 | 1 | | | | 1 | 1 | 1 | |
| High | O00767 | Stearoyl-CoA desaturase OS=Homo sapiens OX=9606 GN=SCD PE=1 SV=2 | 6 | 2 | 2 | 41.5 | 11 | 359 | 9 | 2 | 2 | 2 | | | | 2 | 2 | 2 | | | |
| High | Q9GZT9 | Egl nine homolog 1 OS=Homo sapiens OX=9606 GN=EGLN1 PE=1 SV=1 | 6 | 4 | 4 | 46 | 13 | 426 | 8.53 | | | | 1 | | 1 | | | | 1 | | 1 |
| High | Q7L8W6 | Diphthine--ammonia ligase OS=Homo sapiens OX=9606 GN=DPH6 PE=1 SV=3 | 6 | 3 | 3 | 30.3 | 12 | 267 | 5.4 | | 1 | | 1 | 1 | 1 | | 1 | | 1 | 1 | 1 |
| High | Q96GW9 | Methionine--tRNA ligase, mitochondrial OS=Homo sapiens OX=9606 GN=MARS2 PE=1 SV=2 | 6 | 2 | 2 | 66.5 | 3 | 593 | 8.09 | 1 | 1 | | 1 | | 1 | 1 | 1 | | 1 | | 1 |
| High | P05556 | Integrin beta-1 OS=Homo sapiens OX=9606 GN=ITGB1 PE=1 SV=2 | 6 | 2 | 2 | 88.4 | 3 | 798 | 5.39 | 1 | 1 | 2 | | | | 1 | 1 | 2 | | | |
| High | O94880 | PHD finger protein 14 OS=Homo sapiens OX=9606 GN=PHF14 PE=1 SV=2 | 6 | 3 | 3 | 100 | 5 | 888 | 5.33 | 1 | 1 | 3 | | | | 1 | 1 | 2 | | | |
| High | Q29RF7 | Sister chromatid cohesion protein PDS5 homolog A OS=Homo sapiens OX=9606 GN=PDS5A PE=1 SV=1 | 6 | 2 | 2 | 150.7 | 2 | 1337 | 7.91 | | | 1 | | 1 | | | | 1 | | 1 | |
| High | Q6PCB5 | Lysine-specific demethylase RSBN1L OS=Homo sapiens OX=9606 GN=RSBN1L PE=1 SV=2 | 6 | 3 | 3 | 94.8 | 3 | 846 | 8.78 | | | | | | 1 | | | | | | 1 |
| High | O14776 | Transcription elongation regulator 1 OS=Homo sapiens OX=9606 GN=TCERG1 PE=1 SV=2 | 6 | 2 | 2 | 123.8 | 2 | 1098 | 8.65 | 1 | 1 | 1 | | | | 1 | 1 | 1 | | | |
| High | Q9BYC9 | 39S ribosomal protein L20, mitochondrial OS=Homo sapiens OX=9606 GN=MRPL20 PE=1 SV=1 | 6 | 1 | 1 | 17.4 | 5 | 149 | 10.86 | | | | 1 | 1 | 1 | | | | 1 | 1 | 1 |
| High | Q96RE7 | Nucleus accumbens-associated protein 1 OS=Homo sapiens OX=9606 GN=NACC1 PE=1 SV=1 | 6 | 2 | 2 | 57.2 | 4 | 527 | 5.74 | | | | | | | | | | | | |
| High | Q9NZN8 | CCR4-NOT transcription complex subunit 2 OS=Homo sapiens OX=9606 GN=CNOT2 PE=1 SV=1 | 6 | 4 | 4 | 59.7 | 11 | 540 | 7.66 | | | | | | | | | | | | |
| High | Q9HD33 | 39S ribosomal protein L47, mitochondrial OS=Homo sapiens OX=9606 GN=MRPL47 PE=1 SV=2 | 6 | 2 | 2 | 29.4 | 8 | 250 | 10.37 | | | | | 1 | | | | | | 1 | |
| High | Q96BA8 | Cyclic AMP-responsive element-binding protein 3-like protein 1 OS=Homo sapiens OX=9606 GN=CREB3L1 PE=1 SV=1 | 6 | 3 | 3 | 57 | 8 | 519 | 5.17 | | | | 1 | 2 | | | | | 1 | 2 | |
| High | Q00577 | Transcriptional activator protein Pur-alpha OS=Homo sapiens OX=9606 GN=PURA PE=1 SV=2 | 5 | 3 | 3 | 34.9 | 17 | 322 | 6.44 | | | | | 1 | | | | | | 1 | |
| High | O43617 | Trafficking protein particle complex subunit 3 OS=Homo sapiens OX=9606 GN=TRAPPC3 PE=1 SV=1 | 5 | 2 | 2 | 20.3 | 10 | 180 | 4.96 | | | | | 1 | | | | | | 1 | |
| High | O14879 | Interferon-induced protein with tetratricopeptide repeats 3 OS=Homo sapiens OX=9606 GN=IFIT3 PE=1 SV=1 | 5 | 1 | 1 | 56 | 2 | 490 | 5.2 | 1 | | 1 | 1 | 1 | 1 | 1 | | 1 | 1 | 1 | 1 |
| High | Q86XN8 | RNA-binding protein MEX3D OS=Homo sapiens OX=9606 GN=MEX3D PE=1 SV=3 | 5 | 1 | 1 | 64.8 | 1 | 651 | 5.35 | 1 | 1 | | 1 | 1 | 1 | 1 | 1 | | 1 | 1 | 1 |
| High | Q8TED0 | U3 small nucleolar RNA-associated protein 15 homolog OS=Homo sapiens OX=9606 GN=UTP15 PE=1 SV=3 | 5 | 4 | 4 | 58.4 | 8 | 518 | 9.11 | | | | | 2 | | | | | | 2 | |
| High | Q96B21 | Transmembrane protein 45B OS=Homo sapiens OX=9606 GN=TMEM45B PE=1 SV=1 | 5 | 1 | 1 | 31.8 | 3 | 275 | 6.95 | | | | 1 | 1 | | | | | 1 | 1 | |
| High | P08133 | Annexin A6 OS=Homo sapiens OX=9606 GN=ANXA6 PE=1 SV=3 | 5 | 4 | 4 | 75.8 | 5 | 673 | 5.6 | | | | 3 | 1 | 1 | | | | 3 | 1 | 1 |
| High | O43688 | Phospholipid phosphatase 2 OS=Homo sapiens OX=9606 GN=PLPP2 PE=1 SV=1 | 5 | 2 | 2 | 32.6 | 8 | 288 | 8.35 | | | 1 | | 1 | | | | 1 | | 1 | |
| High | Q8WZ82 | Esterase OVCA2 OS=Homo sapiens OX=9606 GN=OVCA2 PE=1 SV=1 | 5 | 2 | 2 | 24.4 | 10 | 227 | 6.89 | | 1 | 1 | 1 | | | | 1 | 1 | 1 | | |
| High | Q13882 | Protein-tyrosine kinase 6 OS=Homo sapiens OX=9606 GN=PTK6 PE=1 SV=1 | 5 | 2 | 1 | 51.8 | 4 | 451 | 7.03 | | | | | 1 | | | | | | 1 | |
| High | O43570 | Carbonic anhydrase 12 OS=Homo sapiens OX=9606 GN=CA12 PE=1 SV=1 | 5 | 1 | 1 | 39.4 | 4 | 354 | 7.23 | | 1 | 1 | | | | | 1 | 1 | | | |
| High | Q86T03 | Type 1 phosphatidylinositol 4,5-bisphosphate 4-phosphatase OS=Homo sapiens OX=9606 GN=PIP4P1 PE=1 SV=1 | 5 | 1 | 1 | 29.5 | 8 | 277 | 8.91 | 1 | 1 | 1 | | | | 1 | 1 | 1 | | | |
| High | Q03519 | Antigen peptide transporter 2 OS=Homo sapiens OX=9606 GN=TAP2 PE=1 SV=1 | 5 | 1 | 1 | 75.6 | 1 | 686 | 8.02 | 1 | 1 | 1 | | | 1 | 1 | 1 | 1 | | | 1 |
| High | Q7L8L6 | FAST kinase domain-containing protein 5, mitochondrial OS=Homo sapiens OX=9606 GN=FASTKD5 PE=1 SV=1 | 5 | 2 | 2 | 86.5 | 2 | 764 | 8.13 | 2 | 2 | 1 | | | | 2 | 2 | 1 | | | |
| High | O14682 | Ectoderm-neural cortex protein 1 OS=Homo sapiens OX=9606 GN=ENC1 PE=1 SV=2 | 5 | 2 | 2 | 66.1 | 3 | 589 | 6.84 | 1 | | 1 | | 2 | | 1 | | 1 | | 2 | |
| High | Q9H147 | Deoxynucleotidyltransferase terminal-interacting protein 1 OS=Homo sapiens OX=9606 GN=DNTTIP1 PE=1 SV=2 | 5 | 3 | 3 | 37 | 14 | 329 | 8.97 | | | | | | | | | | | | |
| High | Q8IWR0 | Zinc finger CCCH domain-containing protein 7A OS=Homo sapiens OX=9606 GN=ZC3H7A PE=1 SV=1 | 5 | 1 | 1 | 110.5 | 2 | 971 | 7.3 | 1 | 1 | 1 | | | | 1 | 1 | 1 | | | |
| High | Q6UX53 | Thiol S-methyltransferase METTL7B OS=Homo sapiens OX=9606 GN=METTL7B PE=1 SV=2 | 5 | 2 | 2 | 27.8 | 9 | 244 | 8.38 | | | 1 | | 1 | | | | 1 | | 1 | |
| High | Q16514 | Transcription initiation factor TFIID subunit 12 OS=Homo sapiens OX=9606 GN=TAF12 PE=1 SV=1 | 5 | 1 | 1 | 17.9 | 6 | 161 | 7.99 | | | | 1 | 1 | | | | | 1 | 1 | |
| High | Q15542 | Transcription initiation factor TFIID subunit 5 OS=Homo sapiens OX=9606 GN=TAF5 PE=1 SV=3 | 5 | 2 | 2 | 86.8 | 4 | 800 | 5.64 | | | | 1 | | | | | | 1 | | |
| High | Q8NB15 | Zinc finger protein 511 OS=Homo sapiens OX=9606 GN=ZNF511 PE=1 SV=2 | 5 | 1 | 1 | 28.2 | 6 | 252 | 7.66 | | 1 | 1 | | | 1 | | 1 | 1 | | | 1 |
| High | Q9BYD3 | 39S ribosomal protein L4, mitochondrial OS=Homo sapiens OX=9606 GN=MRPL4 PE=1 SV=1 | 5 | 2 | 2 | 34.9 | 9 | 311 | 9.72 | 1 | 1 | 1 | | | | 1 | 1 | 1 | | | |
| High | O43278 | Kunitz-type protease inhibitor 1 OS=Homo sapiens OX=9606 GN=SPINT1 PE=1 SV=2 | 5 | 3 | 3 | 58.4 | 6 | 529 | 6.29 | | | | | | | | | | | | |
| High | Q14494 | Endoplasmic reticulum membrane sensor NFE2L1 OS=Homo sapiens OX=9606 GN=NFE2L1 PE=1 SV=1 | 5 | 2 | 2 | 84.7 | 2 | 772 | 4.64 | | | | 1 | 1 | 2 | | | | 1 | 1 | 2 |
| High | Q96T23 | Remodeling and spacing factor 1 OS=Homo sapiens OX=9606 GN=RSF1 PE=1 SV=2 | 5 | 3 | 3 | 163.7 | 4 | 1441 | 5.01 | 1 | | 2 | | | | 1 | | 2 | | | |
| High | P29508 | Serpin B3 OS=Homo sapiens OX=9606 GN=SERPINB3 PE=1 SV=2 | 5 | 3 | 3 | 44.5 | 6 | 390 | 6.81 | | | | 2 | 2 | 1 | | | | 2 | 2 | 1 |
| High | Q96IR7 | 4-hydroxyphenylpyruvate dioxygenase-like protein OS=Homo sapiens OX=9606 GN=HPDL PE=1 SV=1 | 5 | 2 | 2 | 39.4 | 11 | 371 | 7.03 | 1 | 1 | 1 | | | | 1 | 1 | 1 | | | |
| High | O00429 | Dynamin-1-like protein OS=Homo sapiens OX=9606 GN=DNM1L PE=1 SV=2 | 5 | 4 | 4 | 81.8 | 10 | 736 | 6.81 | 1 | | 2 | | 1 | | 1 | | 2 | | 1 | |
| High | Q8ND56 | Protein LSM14 homolog A OS=Homo sapiens OX=9606 GN=LSM14A PE=1 SV=3 | 5 | 2 | 2 | 50.5 | 4 | 463 | 9.52 | | | 1 | 1 | 1 | | | | 1 | 1 | 1 | |
| High | O14734 | Acyl-coenzyme A thioesterase 8 OS=Homo sapiens OX=9606 GN=ACOT8 PE=1 SV=1 | 5 | 2 | 2 | 35.9 | 13 | 319 | 7.56 | 1 | 1 | 1 | | | | 1 | 1 | 1 | | | |
| High | Q15375 | Ephrin type-A receptor 7 OS=Homo sapiens OX=9606 GN=EPHA7 PE=1 SV=3 | 5 | 3 | 1 | 112 | 3 | 998 | 5.8 | | | | 2 | | 1 | | | | 2 | | 1 |
| High | P11279 | Lysosome-associated membrane glycoprotein 1 OS=Homo sapiens OX=9606 GN=LAMP1 PE=1 SV=3 | 5 | 2 | 2 | 44.9 | 4 | 417 | 8.75 | 2 | | 2 | | | 1 | 2 | | 2 | | | 1 |
| High | Q9Y6G3 | 39S ribosomal protein L42, mitochondrial OS=Homo sapiens OX=9606 GN=MRPL42 PE=1 SV=1 | 5 | 1 | 1 | 16.7 | 11 | 142 | 8.35 | | | | | 1 | 1 | | | | | 1 | 1 |
| High | Q9H0S4 | Probable ATP-dependent RNA helicase DDX47 OS=Homo sapiens OX=9606 GN=DDX47 PE=1 SV=1 | 5 | 3 | 3 | 50.6 | 10 | 455 | 9.1 | | | 1 | | 1 | | | | 1 | | 1 | |
| High | Q9Y4A5 | Transformation/transcription domain-associated protein OS=Homo sapiens OX=9606 GN=TRRAP PE=1 SV=3 | 5 | 3 | 3 | 437.3 | 1 | 3859 | 8.19 | 1 | | 1 | | 1 | | 1 | | 1 | | 1 | |
| High | Q15785 | Mitochondrial import receptor subunit TOM34 OS=Homo sapiens OX=9606 GN=TOMM34 PE=1 SV=2 | 5 | 3 | 3 | 34.5 | 9 | 309 | 8.98 | | 1 | | 1 | | 2 | | 1 | | 1 | | 2 |
| High | Q5T200 | Zinc finger CCCH domain-containing protein 13 OS=Homo sapiens OX=9606 GN=ZC3H13 PE=1 SV=1 | 5 | 3 | 3 | 196.5 | 3 | 1668 | 9.42 | | | | | 1 | | | | | | 1 | |
| High | Q13884 | Beta-1-syntrophin OS=Homo sapiens OX=9606 GN=SNTB1 PE=1 SV=3 | 5 | 3 | 3 | 58 | 9 | 538 | 8.63 | | | 1 | | | | | | 1 | | | |
| High | Q9H4L5 | Oxysterol-binding protein-related protein 3 OS=Homo sapiens OX=9606 GN=OSBPL3 PE=1 SV=1 | 5 | 1 | 1 | 101.2 | 2 | 887 | 6.87 | | 1 | 1 | | | | | 1 | 1 | | | |
| High | O95149 | Snurportin-1 OS=Homo sapiens OX=9606 GN=SNUPN PE=1 SV=1 | 5 | 2 | 2 | 41.1 | 5 | 360 | 6.64 | | | 1 | 1 | 1 | 2 | | | 1 | 1 | 1 | 2 |
| High | Q13443 | Disintegrin and metalloproteinase domain-containing protein 9 OS=Homo sapiens OX=9606 GN=ADAM9 PE=1 SV=1 | 5 | 2 | 2 | 90.5 | 3 | 819 | 7.52 | | | | | | | | | | | | |
| High | Q13618 | Cullin-3 OS=Homo sapiens OX=9606 GN=CUL3 PE=1 SV=2 | 5 | 2 | 2 | 88.9 | 3 | 768 | 8.48 | | | 1 | | | | | | 1 | | | |
| High | Q9Y2U8 | Inner nuclear membrane protein Man1 OS=Homo sapiens OX=9606 GN=LEMD3 PE=1 SV=2 | 5 | 2 | 2 | 99.9 | 5 | 911 | 7.55 | | | | | 1 | | | | | | 1 | |
| High | P16435 | NADPH--cytochrome P450 reductase OS=Homo sapiens OX=9606 GN=POR PE=1 SV=2 | 5 | 1 | 1 | 76.6 | 1 | 677 | 5.58 | 1 | 1 | 1 | | | | 1 | 1 | 1 | | | |
| High | Q9BXV9 | EKC/KEOPS complex subunit GON7 OS=Homo sapiens OX=9606 GN=GON7 PE=1 SV=2 | 5 | 2 | 2 | 10.9 | 41 | 100 | 4.27 | 1 | 1 | 1 | | | | 1 | 1 | 1 | | | |
| High | Q5T0W9 | Protein FAM83B OS=Homo sapiens OX=9606 GN=FAM83B PE=1 SV=1 | 5 | 3 | 3 | 114.7 | 4 | 1011 | 8.97 | 3 | 2 | | | | | 3 | 2 | | | | |
| High | P07196 | Neurofilament light polypeptide OS=Homo sapiens OX=9606 GN=NEFL PE=1 SV=3 | 5 | 2 | 1 | 61.5 | 3 | 543 | 4.65 | | | | 2 | 2 | 1 | | | | 2 | 2 | 1 |
| High | Q9UHA4 | Ragulator complex protein LAMTOR3 OS=Homo sapiens OX=9606 GN=LAMTOR3 PE=1 SV=1 | 5 | 2 | 2 | 13.6 | 27 | 124 | 7.34 | | 2 | 1 | | | | | 2 | 1 | | | |
| High | P14735 | Insulin-degrading enzyme OS=Homo sapiens OX=9606 GN=IDE PE=1 SV=4 | 5 | 2 | 2 | 117.9 | 2 | 1019 | 6.61 | | | 1 | | | 1 | | | 1 | | | 1 |
| High | O00750 | Phosphatidylinositol 4-phosphate 3-kinase C2 domain-containing subunit beta OS=Homo sapiens OX=9606 GN=PIK3C2B PE=1 SV= | 5 | 2 | 2 | 184.7 | 1 | 1634 | 7.3 | | | 1 | | 1 | | | | 1 | | 1 | |
| High | O15357 | Phosphatidylinositol 3,4,5-trisphosphate 5-phosphatase 2 OS=Homo sapiens OX=9606 GN=INPPL1 PE=1 SV=2 | 5 | 3 | 3 | 138.5 | 3 | 1258 | 6.54 | | 1 | 1 | 1 | | | | 1 | 1 | 1 | | |
| High | Q9Y3S1 | Serine/threonine-protein kinase WNK2 OS=Homo sapiens OX=9606 GN=WNK2 PE=1 SV=4 | 5 | 2 | 2 | 242.5 | 1 | 2297 | 6.13 | | | | 1 | | | | | | 1 | | |
| High | P08651 | Nuclear factor 1 C-type OS=Homo sapiens OX=9606 GN=NFIC PE=1 SV=2 | 5 | 2 | 1 | 55.6 | 6 | 508 | 8.38 | | | | 1 | 1 | 1 | | | | 1 | 1 | 1 |
| High | Q15311 | RalA-binding protein 1 OS=Homo sapiens OX=9606 GN=RALBP1 PE=1 SV=3 | 5 | 4 | 4 | 76 | 8 | 655 | 5.88 | 1 | | 1 | | | | 1 | | 1 | | | |
| High | Q15024 | Exosome complex component RRP42 OS=Homo sapiens OX=9606 GN=EXOSC7 PE=1 SV=3 | 5 | 2 | 2 | 31.8 | 11 | 291 | 5.19 | | | | | 2 | | | | | | 1 | |
| High | Q9Y3C1 | Nucleolar protein 16 OS=Homo sapiens OX=9606 GN=NOP16 PE=1 SV=2 | 5 | 2 | 2 | 21.2 | 11 | 178 | 9.94 | | | | | 1 | 1 | | | | | 1 | 1 |
| High | P54760 | Ephrin type-B receptor 4 OS=Homo sapiens OX=9606 GN=EPHB4 PE=1 SV=2 | 5 | 3 | 1 | 108.2 | 3 | 987 | 6.9 | | 1 | | 1 | | 1 | | 1 | | 1 | | 1 |
| High | P56134 | ATP synthase subunit f, mitochondrial OS=Homo sapiens OX=9606 GN=ATP5MF PE=1 SV=3 | 5 | 2 | 2 | 10.9 | 26 | 94 | 9.67 | 2 | 2 | 1 | | | | 2 | 2 | 1 | | | |
| High | Q9Y312 | Protein AAR2 homolog OS=Homo sapiens OX=9606 GN=AAR2 PE=1 SV=2 | 5 | 2 | 2 | 43.4 | 9 | 384 | 5.96 | 1 | 2 | 1 | | | | 1 | 2 | 1 | | | |
| High | P12273 | Prolactin-inducible protein OS=Homo sapiens OX=9606 GN=PIP PE=1 SV=1 | 5 | 2 | 2 | 16.6 | 13 | 146 | 8.05 | | | | 2 | 2 | 1 | | | | 2 | 2 | 1 |
| High | Q5D862 | SWISS-PROT:Q5D862 Tax_Id=9606 Gene_Symbol=FLG2 Filaggrin-2 | 5 | 4 | 4 | 247.9 | 2 | 2391 | 8.31 | | 1 | 3 | | 1 | | | 1 | 3 | | 1 | |
| High | Q9NX14 | NADH dehydrogenase [ubiquinone] 1 beta subcomplex subunit 11, mitochondrial OS=Homo sapiens OX=9606 GN=NDUFB11 PE=1 | 5 | 1 | 1 | 17.3 | 7 | 153 | 5.22 | 1 | 1 | | | | | 1 | 1 | | | | |
| High | Q15404 | Ras suppressor protein 1 OS=Homo sapiens OX=9606 GN=RSU1 PE=1 SV=3 | 5 | 2 | 2 | 31.5 | 6 | 277 | 8.65 | | | 1 | | | | | | 1 | | | |
| High | O60825 | 6-phosphofructo-2-kinase/fructose-2,6-bisphosphatase 2 OS=Homo sapiens OX=9606 GN=PFKFB2 PE=1 SV=2 | 5 | 3 | 1 | 58.4 | 6 | 505 | 8.38 | | | | | 1 | | | | | | 1 | |
| High | P50502 | Hsc70-interacting protein OS=Homo sapiens OX=9606 GN=ST13 PE=1 SV=2 | 5 | 2 | 2 | 41.3 | 4 | 369 | 5.27 | | | 2 | 1 | | 1 | | | 2 | 1 | | 1 |
| High | Q9P2D1 | Chromodomain-helicase-DNA-binding protein 7 OS=Homo sapiens OX=9606 GN=CHD7 PE=1 SV=3 | 5 | 3 | 2 | 335.7 | 1 | 2997 | 6.34 | 2 | 1 | 2 | | | | 2 | 1 | 2 | | | |
| High | O96005 | Cleft lip and palate transmembrane protein 1 OS=Homo sapiens OX=9606 GN=CLPTM1 PE=1 SV=1 | 5 | 3 | 3 | 76 | 3 | 669 | 6.3 | 2 | 1 | 1 | | | | 2 | 1 | 1 | | | |
| High | Q9H9T3 | Elongator complex protein 3 OS=Homo sapiens OX=9606 GN=ELP3 PE=1 SV=2 | 5 | 3 | 3 | 62.2 | 8 | 547 | 8.88 | 1 | 1 | | | | 1 | 1 | 1 | | | | 1 |
| High | Q8IWA0 | WD repeat-containing protein 75 OS=Homo sapiens OX=9606 GN=WDR75 PE=1 SV=1 | 5 | 1 | 1 | 94.4 | 1 | 830 | 5.96 | | 1 | 1 | | | | | 1 | 1 | | | |
| High | Q8TAA5 | GrpE protein homolog 2, mitochondrial OS=Homo sapiens OX=9606 GN=GRPEL2 PE=1 SV=1 | 5 | 3 | 3 | 25.4 | 12 | 225 | 7.72 | 1 | 1 | | | | | 1 | 1 | | | | |
| High | Q9BW91 | ADP-ribose pyrophosphatase, mitochondrial OS=Homo sapiens OX=9606 GN=NUDT9 PE=1 SV=1 | 5 | 1 | 1 | 39.1 | 2 | 350 | 8.22 | | | 1 | | 1 | | | | 1 | | 1 | |
| High | Q99442 | Translocation protein SEC62 OS=Homo sapiens OX=9606 GN=SEC62 PE=1 SV=1 | 5 | 3 | 3 | 45.8 | 8 | 399 | 7.12 | | | | | | | | | | | | |
| High | Q9BVT8 | Transmembrane and ubiquitin-like domain-containing protein 1 OS=Homo sapiens OX=9606 GN=TMUB1 PE=1 SV=1 | 5 | 2 | 2 | 26.2 | 9 | 246 | 5.72 | | | 1 | | | | | | 1 | | | |
| High | O75616 | GTPase Era, mitochondrial OS=Homo sapiens OX=9606 GN=ERAL1 PE=1 SV=2 | 5 | 1 | 1 | 48.3 | 3 | 437 | 8.84 | | | 1 | | 1 | | | | 1 | | 1 | |
| High | O60229 | Kalirin OS=Homo sapiens OX=9606 GN=KALRN PE=1 SV=3 | 5 | 3 | 3 | 340 | 1 | 2986 | 6.07 | | | 1 | | | | | | 1 | | | |
| High | O00411 | DNA-directed RNA polymerase, mitochondrial OS=Homo sapiens OX=9606 GN=POLRMT PE=1 SV=2 | 5 | 2 | 2 | 138.5 | 1 | 1230 | 8.98 | | | | 1 | | | | | | 1 | | |
| High | Q8WUW1 | Protein BRICK1 OS=Homo sapiens OX=9606 GN=BRK1 PE=1 SV=1 | 5 | 1 | 1 | 8.7 | 17 | 75 | 5.45 | | | 2 | | | | | | 1 | | | |
| High | Q9NX57 | Ras-related protein Rab-20 OS=Homo sapiens OX=9606 GN=RAB20 PE=1 SV=1 | 5 | 3 | 3 | 26.3 | 14 | 234 | 6.55 | 1 | | | | | | 1 | | | | | |
| High | Q9C0B9 | Zinc finger CCHC domain-containing protein 2 OS=Homo sapiens OX=9606 GN=ZCCHC2 PE=1 SV=6 | 5 | 4 | 4 | 125.9 | 5 | 1178 | 7.01 | | | | 2 | 1 | 2 | | | | 2 | 1 | 2 |
| High | Q9H2M9 | Rab3 GTPase-activating protein non-catalytic subunit OS=Homo sapiens OX=9606 GN=RAB3GAP2 PE=1 SV=1 | 5 | 2 | 2 | 155.9 | 2 | 1393 | 5.62 | | | 1 | | | | | | 1 | | | |
| High | P35754 | Glutaredoxin-1 OS=Homo sapiens OX=9606 GN=GLRX PE=1 SV=2 | 5 | 2 | 2 | 11.8 | 30 | 106 | 8.09 | | | 1 | | 1 | | | | 1 | | 1 | |
| High | Q9GZZ9 | Ubiquitin-like modifier-activating enzyme 5 OS=Homo sapiens OX=9606 GN=UBA5 PE=1 SV=1 | 5 | 2 | 2 | 44.8 | 8 | 404 | 4.84 | 1 | 1 | | | | | 1 | 1 | | | | |
| High | Q86WB0 | Nuclear-interacting partner of ALK OS=Homo sapiens OX=9606 GN=ZC3HC1 PE=1 SV=1 | 5 | 2 | 2 | 55.2 | 4 | 502 | 5.62 | | | 1 | 1 | | | | | 1 | 1 | | |
| High | Q9Y4C1 | Lysine-specific demethylase 3A OS=Homo sapiens OX=9606 GN=KDM3A PE=1 SV=4 | 5 | 4 | 4 | 147.2 | 5 | 1321 | 8.07 | | | 1 | | | | | | 1 | | | |
| High | Q9NUQ3 | Gamma-taxilin OS=Homo sapiens OX=9606 GN=TXLNG PE=1 SV=2 | 5 | 2 | 2 | 60.5 | 3 | 528 | 7.52 | | | 1 | 1 | 1 | 1 | | | 1 | 1 | 1 | 1 |
| High | P11274 | Breakpoint cluster region protein OS=Homo sapiens OX=9606 GN=BCR PE=1 SV=2 | 5 | 4 | 3 | 142.7 | 4 | 1271 | 7.03 | | | | | 1 | | | | | | 1 | |
| High | Q9BUL9 | Ribonuclease P protein subunit p25 OS=Homo sapiens OX=9606 GN=RPP25 PE=1 SV=1 | 5 | 2 | 2 | 20.6 | 11 | 199 | 9.61 | | | | 1 | 1 | | | | | 1 | 1 | |
| High | Q86WR7 | Proline and serine-rich protein 2 OS=Homo sapiens OX=9606 GN=PROSER2 PE=1 SV=2 | 5 | 2 | 2 | 45.8 | 6 | 435 | 7.2 | 1 | 2 | 1 | | | | 1 | 2 | 1 | | | |

| | | | | | | | | | | | | | | | | | | | | | |
|---|---|---|---|---|---|---|---|---|---|---|---|---|---|---|---|---|---|---|---|---|---|
| High | Q01415 | N-acetylgalactosamine kinase OS=Homo sapiens OX=9606 GN=GALK2 PE=1 SV=1 | 5 | 1 | 1 | 50.3 | 2 | 458 | 6.61 | | 1 | 1 | | | 1 | | 1 | 1 | | | 1 |
| High | Q9Y365 | START domain-containing protein 10 OS=Homo sapiens OX=9606 GN=STARD10 PE=1 SV=2 | 5 | 2 | 2 | 33 | 10 | 291 | 7.12 | | 1 | | | | | | 1 | | | | |
| High | O00203 | AP-3 complex subunit beta-1 OS=Homo sapiens OX=9606 GN=AP3B1 PE=1 SV=3 | 5 | 2 | 2 | 121.2 | 2 | 1094 | 6.04 | 1 | 2 | 1 | | | | 1 | 2 | 1 | | | |
| High | O94826 | Mitochondrial import receptor subunit TOM70 OS=Homo sapiens OX=9606 GN=TOMM70 PE=1 SV=1 | 5 | 2 | 2 | 67.4 | 5 | 608 | 7.12 | | 1 | | | | | | 1 | | | | |
| High | P52735 | Guanine nucleotide exchange factor VAV2 OS=Homo sapiens OX=9606 GN=VAV2 PE=1 SV=2 | 5 | 2 | 2 | 101.2 | 3 | 878 | 7.08 | 1 | 1 | 1 | | | | 1 | 1 | 1 | | | |
| High | Q96I99 | Succinate--CoA ligase [GDP-forming] subunit beta, mitochondrial OS=Homo sapiens OX=9606 GN=SUCLG2 PE=1 SV=2 | 5 | 2 | 2 | 46.5 | 5 | 432 | 6.39 | | 1 | 1 | 1 | | | | 1 | 1 | 1 | | |
| High | Q8WW01 | tRNA-splicing endonuclease subunit Sen15 OS=Homo sapiens OX=9606 GN=TSEN15 PE=1 SV=1 | 5 | 3 | 3 | 18.6 | 33 | 171 | 4.58 | | | | | | 1 | | | | | | 1 |
| High | A4D161 | Protein FAM221A OS=Homo sapiens OX=9606 GN=FAM221A PE=1 SV=1 | 5 | 2 | 2 | 33.1 | 8 | 298 | 6.74 | | | 1 | | | | | | 1 | | | |
| High | Q96L92 | Sorting nexin-27 OS=Homo sapiens OX=9606 GN=SNX27 PE=1 SV=2 | 5 | 1 | 1 | 61.2 | 2 | 541 | 6.49 | | | 1 | 1 | | | | | 1 | 1 | | |
| High | Q9NP92 | 39S ribosomal protein S30, mitochondrial OS=Homo sapiens OX=9606 GN=MRPS30 PE=1 SV=2 | 5 | 3 | 3 | 50.3 | 15 | 439 | 7.97 | 1 | 1 | | | 1 | | 1 | 1 | | | 1 | |
| High | Q9H4A6 | Golgi phosphoprotein 3 OS=Homo sapiens OX=9606 GN=GOLPH3 PE=1 SV=1 | 5 | 3 | 3 | 33.8 | 17 | 298 | 6.44 | 1 | | 1 | 1 | | | 1 | | 1 | 1 | | |
| High | Q9C0C9 | (E3-independent) E2 ubiquitin-conjugating enzyme OS=Homo sapiens OX=9606 GN=UBE2O PE=1 SV=3 | 5 | 2 | 2 | 141.2 | 1 | 1292 | 5.12 | 1 | | 1 | | | | 1 | | 1 | | | |
| High | Q6GMV2 | Histone-lysine N-trimethyltransferase SMYD5 OS=Homo sapiens OX=9606 GN=SMYD5 PE=1 SV=2 | 5 | 1 | 1 | 47.3 | 3 | 418 | 5.05 | | | 1 | | 1 | | | | 1 | | 1 | |
| High | Q6P1N0 | Coiled-coil and C2 domain-containing protein 1A OS=Homo sapiens OX=9606 GN=CC2D1A PE=1 SV=1 | 5 | 2 | 2 | 104 | 3 | 951 | 8.09 | | | | | | | | | | | | |
| High | P02751 | Fibronectin OS=Homo sapiens OX=9606 GN=FN1 PE=1 SV=5 | 5 | 2 | 2 | 272.2 | 1 | 2477 | 5.5 | | | | 1 | | 1 | | | | 1 | | 1 |
| High | Q15392 | Delta(24)-sterol reductase OS=Homo sapiens OX=9606 GN=DHCR24 PE=1 SV=2 | 5 | 2 | 2 | 60.1 | 5 | 516 | 8.16 | 1 | 1 | 1 | | | | 1 | 1 | 1 | | | |
| High | O75821 | Eukaryotic translation initiation factor 3 subunit G OS=Homo sapiens OX=9606 GN=EIF3G PE=1 SV=2 | 5 | 2 | 2 | 35.6 | 10 | 320 | 6.13 | 2 | 2 | 1 | | | | 2 | 2 | 1 | | | |
| High | Q9NP97 | Dynein light chain roadblock-type 1 OS=Homo sapiens OX=9606 GN=DYNLRB1 PE=1 SV=3 | 5 | 2 | 2 | 10.9 | 34 | 96 | 7.25 | 1 | 1 | 1 | | | | 1 | 1 | 1 | | | |
| High | Q9HAU5 | Regulator of nonsense transcripts 2 OS=Homo sapiens OX=9606 GN=UPF2 PE=1 SV=1 | 5 | 4 | 4 | 147.7 | 4 | 1272 | 5.69 | | | 1 | 2 | | | | | 1 | 2 | | |
| High | Q7L3T8 | Probable proline--tRNA ligase, mitochondrial OS=Homo sapiens OX=9606 GN=PARS2 PE=1 SV=1 | 5 | 3 | 3 | 53.2 | 8 | 475 | 8.1 | | | | | | | | | | | | |
| High | Q7Z2Z2 | Elongation factor-like GTPase 1 OS=Homo sapiens OX=9606 GN=EFL1 PE=1 SV=2 | 5 | 2 | 2 | 125.4 | 3 | 1120 | 5.91 | | | 1 | | 1 | | | | 1 | | 1 | |
| High | O15162 | Phospholipid scramblase 1 OS=Homo sapiens OX=9606 GN=PLSCR1 PE=1 SV=1 | 5 | 2 | 2 | 35 | 7 | 318 | 4.94 | | 1 | | | | | | 1 | | | | |
| High | Q969X5 | Endoplasmic reticulum-Golgi intermediate compartment protein 1 OS=Homo sapiens OX=9606 GN=ERGIC1 PE=1 SV=1 | 5 | 2 | 2 | 32.6 | 6 | 290 | 7.06 | | | | | 1 | | | | | | 1 | |
| High | P21953 | 2-oxoisovalerate dehydrogenase subunit beta, mitochondrial OS=Homo sapiens OX=9606 GN=BCKDHB PE=1 SV=2 | 5 | 4 | 4 | 43.1 | 14 | 392 | 6.29 | | | 1 | | | | | | 1 | | | |
| High | Q9BV86 | N-terminal Xaa-Pro-Lys N-methyltransferase 1 OS=Homo sapiens OX=9606 GN=NTMT1 PE=1 SV=3 | 5 | 1 | 1 | 25.4 | 7 | 223 | 5.52 | | | | | 1 | 1 | | | | | 1 | 1 |
| High | O60934 | Nibrin OS=Homo sapiens OX=9606 GN=NBN PE=1 SV=1 | 5 | 4 | 4 | 84.9 | 7 | 754 | 6.9 | | | 1 | | | | | | 1 | | | |
| High | Q9UGP4 | LIM domain-containing protein 1 OS=Homo sapiens OX=9606 GN=LIMD1 PE=1 SV=1 | 5 | 2 | 2 | 72.1 | 5 | 676 | 6.65 | | | | | | 1 | | | | | | 1 |
| High | Q8WXX5 | DnaJ homolog subfamily C member 9 OS=Homo sapiens OX=9606 GN=DNAJC9 PE=1 SV=1 | 5 | 3 | 3 | 29.9 | 13 | 260 | 5.73 | | | | | 1 | 1 | | | | | 1 | 1 |
| High | Q96H55 | Unconventional myosin-XIX OS=Homo sapiens OX=9606 GN=MYO19 PE=1 SV=2 | 5 | 2 | 2 | 109.1 | 2 | 970 | 7.71 | | | 2 | 1 | | | | | 2 | 1 | | |
| High | P83111 | Serine beta-lactamase-like protein LACTB, mitochondrial OS=Homo sapiens OX=9606 GN=LACTB PE=1 SV=2 | 5 | 2 | 2 | 60.7 | 3 | 547 | 8.53 | | | 1 | | | | | | 1 | | | |
| High | Q9Y287 | Integral membrane protein 2B OS=Homo sapiens OX=9606 GN=ITM2B PE=1 SV=1 | 5 | 2 | 2 | 30.3 | 14 | 266 | 5.14 | 1 | 1 | 1 | | | | 1 | 1 | 1 | | | |
| High | Q9Y3E5 | Peptidyl-tRNA hydrolase 2, mitochondrial OS=Homo sapiens OX=9606 GN=PTRH2 PE=1 SV=1 | 5 | 2 | 2 | 19.2 | 21 | 179 | 8.73 | 1 | | | | | 1 | 1 | | | | | 1 |
| High | Q9BVG4 | Protein PBDC1 OS=Homo sapiens OX=9606 GN=PBDC1 PE=1 SV=1 | 5 | 1 | 1 | 26 | 4 | 233 | 4.79 | | | 1 | | 1 | | | | 1 | | 1 | |
| High | Q16625 | Occludin OS=Homo sapiens OX=9606 GN=OCLN PE=1 SV=1 | 5 | 1 | 1 | 59.1 | 4 | 522 | 6.09 | 1 | | 1 | | | | 1 | | 1 | | | |
| High | Q15459 | Splicing factor 3A subunit 1 OS=Homo sapiens OX=9606 GN=SF3A1 PE=1 SV=1 | 5 | 3 | 3 | 88.8 | 6 | 793 | 5.22 | 1 | | | | | 2 | 1 | | | | | 2 |
| High | Q9GZT3 | SRA stem-loop-interacting RNA-binding protein, mitochondrial OS=Homo sapiens OX=9606 GN=SLIRP PE=1 SV=1 | 5 | 1 | 1 | 12.3 | 13 | 109 | 10.24 | | | 1 | | 1 | | | | 1 | | 1 | |
| High | C9J798 | Ras GTPase-activating protein 4B OS=Homo sapiens OX=9606 GN=RASA4B PE=3 SV=2 | 5 | 4 | 4 | 90.4 | 7 | 803 | 7.74 | | 1 | | | | | | 1 | | | | |
| High | Q5T2T1 | MAGUK p55 subfamily member 7 OS=Homo sapiens OX=9606 GN=MPP7 PE=1 SV=1 | 5 | 2 | 2 | 65.5 | 5 | 576 | 7.14 | 1 | 1 | 1 | | | | 1 | 1 | 1 | | | |
| High | Q96PZ0 | Pseudouridylate synthase 7 homolog OS=Homo sapiens OX=9606 GN=PUS7 PE=1 SV=2 | 5 | 4 | 4 | 75 | 12 | 661 | 6.37 | | | 1 | | | | | | 1 | | | |
| High | P22307 | Sterol carrier protein 2 OS=Homo sapiens OX=9606 GN=SCP2 PE=1 SV=2 | 5 | 1 | 1 | 59 | 1 | 547 | 6.89 | | | 1 | | 1 | | | | 1 | | 1 | |
| High | P08574 | Cytochrome c1, heme protein, mitochondrial OS=Homo sapiens OX=9606 GN=CYC1 PE=1 SV=3 | 5 | 2 | 2 | 35.4 | 10 | 325 | 9 | 2 | 1 | 1 | | | | 2 | 1 | 1 | | | |
| High | Q9UK45 | U6 snRNA-associated Sm-like protein LSm7 OS=Homo sapiens OX=9606 GN=LSM7 PE=1 SV=1 | 5 | 2 | 2 | 11.6 | 33 | 103 | 5.27 | | | | | 1 | | | | | | 1 | |
| High | O95340 | Bifunctional 3'-phosphoadenosine 5'-phosphosulfate synthase 2 OS=Homo sapiens OX=9606 GN=PAPSS2 PE=1 SV=2 | 5 | 2 | 2 | 69.5 | 5 | 614 | 8.03 | | | 1 | | 1 | | | | 1 | | 1 | |
| High | O95365 | Zinc finger and BTB domain-containing protein 7A OS=Homo sapiens OX=9606 GN=ZBTB7A PE=1 SV=1 | 5 | 2 | 2 | 61.4 | 6 | 584 | 5.19 | 1 | 1 | | | | | 1 | 1 | | | | |
| High | O15270 | Serine palmitoyltransferase 2 OS=Homo sapiens OX=9606 GN=SPTLC2 PE=1 SV=1 | 5 | 2 | 2 | 62.9 | 4 | 562 | 7.78 | | | | | 2 | | | | | | 2 | |
| High | Q8TEW0 | Partitioning defective 3 homolog OS=Homo sapiens OX=9606 GN=PARD3 PE=1 SV=2 | 5 | 4 | 4 | 151.3 | 3 | 1356 | 7.68 | 3 | | | | | | 3 | | | | | |
| High | P40937 | Replication factor C subunit 5 OS=Homo sapiens OX=9606 GN=RFC5 PE=1 SV=1 | 5 | 2 | 2 | 38.5 | 10 | 340 | 7.2 | | | 1 | | | | | | 1 | | | |
| High | Q9Y294 | Histone chaperone ASF1A OS=Homo sapiens OX=9606 GN=ASF1A PE=1 SV=1 | 5 | 2 | 2 | 23 | 9 | 204 | 4.41 | | 1 | | | 2 | 1 | | 1 | | | 2 | 1 |
| High | Q13601 | KRR1 small subunit processome component homolog OS=Homo sapiens OX=9606 GN=KRR1 PE=1 SV=4 | 5 | 3 | 3 | 43.6 | 7 | 381 | 9.77 | | | | | 2 | | | | | | 2 | |
| High | P49407 | Beta-arrestin-1 OS=Homo sapiens OX=9606 GN=ARRB1 PE=1 SV=2 | 5 | 3 | 3 | 47 | 9 | 418 | 6.2 | | | 1 | | | | | | 1 | | | |
| High | Q8WUP2 | Filamin-binding LIM protein 1 OS=Homo sapiens OX=9606 GN=FBLIM1 PE=1 SV=2 | 5 | 2 | 2 | 40.6 | 9 | 373 | 6.02 | | | | 1 | | 1 | | | | 1 | | 1 |
| High | Q9UJY4 | ADP-ribosylation factor-binding protein GGA2 OS=Homo sapiens OX=9606 GN=GGA2 PE=1 SV=3 | 5 | 3 | 3 | 67.1 | 8 | 613 | 6.55 | | 1 | | | | | | 1 | | | | |
| High | P15090 | Fatty acid-binding protein, adipocyte OS=Homo sapiens OX=9606 GN=FABP4 PE=1 SV=3 | 5 | 2 | 2 | 14.7 | 16 | 132 | 7.14 | | | | 2 | 1 | 2 | | | | 2 | 1 | 2 |
| High | O43847 | Nardilysin OS=Homo sapiens OX=9606 GN=NRDC PE=1 SV=3 | 5 | 3 | 3 | 131.6 | 3 | 1151 | 4.98 | | | 1 | | 2 | | | | 1 | | 2 | |
| High | O95453 | Poly(A)-specific ribonuclease PARN OS=Homo sapiens OX=9606 GN=PARN PE=1 SV=1 | 5 | 2 | 2 | 73.4 | 3 | 639 | 6.2 | | | | 1 | | 1 | | | | 1 | | 1 |
| High | O14733 | Dual specificity mitogen-activated protein kinase kinase 7 OS=Homo sapiens OX=9606 GN=MAP2K7 PE=1 SV=2 | 5 | 2 | 2 | 47.5 | 6 | 419 | 9.16 | 2 | 1 | | | 1 | | 2 | 1 | | | 1 | |
| High | Q6UWE0 | E3 ubiquitin-protein ligase LRSAM1 OS=Homo sapiens OX=9606 GN=LRSAM1 PE=1 SV=1 | 5 | 2 | 2 | 83.5 | 4 | 723 | 5.94 | | | 1 | | | | | | 1 | | | |
| High | O95858 | Tetraspanin-15 OS=Homo sapiens OX=9606 GN=TSPAN15 PE=1 SV=1 | 5 | 1 | 1 | 33.1 | 3 | 294 | 5.48 | | 1 | 1 | | | | | 1 | 1 | | | |
| High | Q9H299 | SH3 domain-binding glutamic acid-rich-like protein 3 OS=Homo sapiens OX=9606 GN=SH3BGRL3 PE=1 SV=1 | 5 | 1 | 1 | 10.4 | 11 | 93 | 4.93 | 1 | 1 | 1 | 1 | | 1 | 1 | 1 | 1 | 1 | | 1 |
| High | P17026 | Zinc finger protein 22 OS=Homo sapiens OX=9606 GN=ZNF22 PE=1 SV=3 | 5 | 2 | 2 | 25.9 | 8 | 224 | 10.04 | | | | | | | | | | | | |
| High | P04181 | Ornithine aminotransferase, mitochondrial OS=Homo sapiens OX=9606 GN=OAT PE=1 SV=1 | 5 | 1 | 1 | 48.5 | 2 | 439 | 7.03 | | 1 | 1 | | | 1 | | 1 | 1 | | | 1 |
| High | Q86XZ4 | Spermatogenesis-associated serine-rich protein 2 OS=Homo sapiens OX=9606 GN=SPATS2 PE=1 SV=1 | 5 | 3 | 3 | 59.5 | 10 | 545 | 8.9 | | | | | | 1 | | | | | | 1 |
| High | Q15555 | Microtubule-associated protein RP/EB family member 2 OS=Homo sapiens OX=9606 GN=MAPRE2 PE=1 SV=1 | 5 | 2 | 2 | 37 | 7 | 327 | 5.57 | 1 | 1 | 1 | | 1 | | 1 | 1 | 1 | | 1 | |
| High | P61960 | Ubiquitin-fold modifier 1 OS=Homo sapiens OX=9606 GN=UFM1 PE=1 SV=1 | 5 | 2 | 2 | 9.1 | 27 | 85 | 9.31 | | | | 1 | 2 | | | | | 1 | 2 | |
| High | Q9H6S0 | 3'-5' RNA helicase YTHDC2 OS=Homo sapiens OX=9606 GN=YTHDC2 PE=1 SV=2 | 5 | 3 | 3 | 160.1 | 2 | 1430 | 8.4 | | | | | | | | | | | | |
| High | Q93099 | Homogentisate 1,2-dioxygenase OS=Homo sapiens OX=9606 GN=HGD PE=1 SV=2 | 5 | 3 | 3 | 49.9 | 13 | 445 | 6.96 | | 1 | | | | | | 1 | | | | |
| High | Q9NPJ3 | Acyl-coenzyme A thioesterase 13 OS=Homo sapiens OX=9606 GN=ACOT13 PE=1 SV=1 | 5 | 1 | 1 | 15 | 7 | 140 | 9.14 | | | | | 1 | 1 | | | | | 1 | 1 |
| High | Q9H4A5 | Golgi phosphoprotein 3-like OS=Homo sapiens OX=9606 GN=GOLPH3L PE=1 SV=1 | 5 | 1 | 1 | 32.7 | 6 | 285 | 5.83 | 1 | 1 | 1 | | | | 1 | 1 | 1 | | | |
| High | Q9Y597 | BTB/POZ domain-containing protein KCTD3 OS=Homo sapiens OX=9606 GN=KCTD3 PE=1 SV=2 | 5 | 2 | 1 | 88.9 | 3 | 815 | 7.03 | | | | | 1 | | | | | | 1 | |
| High | Q15653 | NF-kappa-B inhibitor beta OS=Homo sapiens OX=9606 GN=NFKBIB PE=1 SV=2 | 5 | 1 | 1 | 37.7 | 4 | 356 | 4.84 | 1 | | 1 | | | | 1 | | 1 | | | |
| High | O75832 | 26S proteasome non-ATPase regulatory subunit 10 OS=Homo sapiens OX=9606 GN=PSMD10 PE=1 SV=1 | 5 | 3 | 3 | 24.4 | 16 | 226 | 6.1 | | | | 1 | 1 | 1 | | | | 1 | 1 | 1 |
| High | Q9Y5Q8 | General transcription factor 3C polypeptide 5 OS=Homo sapiens OX=9606 GN=GTF3C5 PE=1 SV=2 | 5 | 1 | 1 | 59.5 | 2 | 519 | 6.9 | | | | 1 | 1 | 1 | | | | 1 | 1 | 1 |
| High | Q96A35 | 39S ribosomal protein L24, mitochondrial OS=Homo sapiens OX=9606 GN=MRPL24 PE=1 SV=1 | 5 | 4 | 4 | 24.9 | 23 | 216 | 9.29 | | | | | | | | | | | | |
| High | P25685 | DnaJ homolog subfamily B member 1 OS=Homo sapiens OX=9606 GN=DNAJB1 PE=1 SV=4 | 5 | 5 | 4 | 38 | 17 | 340 | 8.63 | | 1 | 1 | | | | | 1 | 1 | | | |
| High | Q96GX9 | Methylthioribulose-1-phosphate dehydratase OS=Homo sapiens OX=9606 GN=APIP PE=1 SV=1 | 4 | 1 | 1 | 27.1 | 7 | 242 | 7.12 | 1 | 1 | 1 | | | | 1 | 1 | 1 | | | |
| High | Q7L5A8 | Fatty acid 2-hydroxylase OS=Homo sapiens OX=9606 GN=FA2H PE=1 SV=1 | 4 | 2 | 2 | 42.8 | 6 | 372 | 8.66 | 1 | 1 | 1 | | | | 1 | 1 | 1 | | | |
| High | P31751 | RAC-beta serine/threonine-protein kinase OS=Homo sapiens OX=9606 GN=AKT2 PE=1 SV=2 | 4 | 2 | 2 | 55.7 | 6 | 481 | 6.37 | | | 1 | | | | | | 1 | | | |
| High | Q12979 | Active breakpoint cluster region-related protein OS=Homo sapiens OX=9606 GN=ABR PE=1 SV=2 | 4 | 2 | 1 | 97.5 | 3 | 859 | 6.55 | | | 1 | | 1 | | | | 1 | | 1 | |
| High | Q4G0F5 | Vacuolar protein sorting-associated protein 26B OS=Homo sapiens OX=9606 GN=VPS26B PE=1 SV=2 | 4 | 2 | 1 | 39.1 | 8 | 336 | 7.36 | | 1 | | | 1 | | | 1 | | | 1 | |
| High | Q9BU68 | Proline-rich protein 15-like protein OS=Homo sapiens OX=9606 GN=PRR15L PE=3 SV=1 | 4 | 1 | 1 | 11.7 | 17 | 103 | 9.5 | 1 | 1 | 1 | | | | 1 | 1 | 1 | | | |
| High | Q9BVS4 | Serine/threonine-protein kinase RIO2 OS=Homo sapiens OX=9606 GN=RIOK2 PE=1 SV=2 | 4 | 1 | 1 | 63.2 | 4 | 552 | 5.94 | | | 1 | | | | | | 1 | | | |
| High | Q9BVC4 | Target of rapamycin complex subunit LST8 OS=Homo sapiens OX=9606 GN=MLST8 PE=1 SV=1 | 4 | 2 | 2 | 35.9 | 12 | 326 | 5.86 | | 1 | | | | | | 1 | | | | |
| High | O15530 | 3-phosphoinositide-dependent protein kinase 1 OS=Homo sapiens OX=9606 GN=PDPK1 PE=1 SV=1 | 4 | 2 | 2 | 63.1 | 5 | 556 | 7.36 | 1 | 1 | 1 | | | | 1 | 1 | 1 | | | |
| High | P48739 | Phosphatidylinositol transfer protein beta isoform OS=Homo sapiens OX=9606 GN=PITPNB PE=1 SV=2 | 4 | 3 | 3 | 31.5 | 13 | 271 | 6.87 | | | | 1 | | 1 | | | | 1 | | 1 |
| High | Q96IW7 | Vesicle-trafficking protein SEC22a OS=Homo sapiens OX=9606 GN=SEC22A PE=1 SV=1 | 4 | 1 | 1 | 34.9 | 2 | 307 | 8.24 | 1 | | 1 | | | | 1 | | 1 | | | |
| High | P49441 | Inositol polyphosphate 1-phosphatase OS=Homo sapiens OX=9606 GN=INPP1 PE=1 SV=1 | 4 | 2 | 2 | 44 | 9 | 399 | 5.26 | 1 | 1 | 1 | | | | 1 | 1 | 1 | | | |
| High | O00267 | Transcription elongation factor SPT5 OS=Homo sapiens OX=9606 GN=SUPT5H PE=1 SV=1 | 4 | 2 | 2 | 120.9 | 3 | 1087 | 5.06 | 1 | | | | 1 | | 1 | | | | 1 | |
| High | P48059 | LIM and senescent cell antigen-like-containing domain protein 1 OS=Homo sapiens OX=9606 GN=LIMS1 PE=1 SV=4 | 4 | 3 | 3 | 37.2 | 9 | 325 | 8.05 | | | | | | | | | | | | |
| High | Q9NQT5 | Exosome complex component RRP40 OS=Homo sapiens OX=9606 GN=EXOSC3 PE=1 SV=3 | 4 | 3 | 3 | 29.6 | 12 | 275 | 8.1 | | | | | | | | | | | | |
| High | Q9NV35 | Nucleotide triphosphate diphosphatase NUDT15 OS=Homo sapiens OX=9606 GN=NUDT15 PE=1 SV=1 | 4 | 1 | 1 | 18.6 | 7 | 164 | 6.14 | | | 1 | 1 | | 1 | | | 1 | 1 | | 1 |
| High | Q13445 | Transmembrane emp24 domain-containing protein 1 OS=Homo sapiens OX=9606 GN=TMED1 PE=1 SV=1 | 4 | 1 | 1 | 25.2 | 7 | 227 | 4.48 | 1 | 1 | 1 | | | | 1 | 1 | 1 | | | |
| High | O75964 | ATP synthase subunit g, mitochondrial OS=Homo sapiens OX=9606 GN=ATP5MG PE=1 SV=3 | 4 | 1 | 1 | 11.4 | 11 | 103 | 9.64 | | 1 | 1 | | | | | 1 | 1 | | | |
| High | P16220 | Cyclic AMP-responsive element-binding protein 1 OS=Homo sapiens OX=9606 GN=CREB1 PE=1 SV=3 | 4 | 2 | 2 | 35.1 | 12 | 327 | 5.27 | | | | 2 | 1 | 1 | | | | 2 | 1 | 1 |
| High | P01857 | Immunoglobulin heavy constant gamma 1 OS=Homo sapiens OX=9606 GN=IGHG1 PE=1 SV=1 | 4 | 2 | 1 | 36.1 | 7 | 330 | 8.19 | | | | 1 | 1 | 2 | | | | 1 | 1 | 2 |
| High | Q96Q11 | CCA tRNA nucleotidyltransferase 1, mitochondrial OS=Homo sapiens OX=9606 GN=TRNT1 PE=1 SV=2 | 4 | 2 | 2 | 50.1 | 6 | 434 | 8.1 | | | 1 | | | 1 | | | 1 | | | 1 |
| High | Q08188 | Protein-glutamine gamma-glutamyltransferase E OS=Homo sapiens OX=9606 GN=TGM3 PE=1 SV=4 | 4 | 3 | 3 | 76.6 | 3 | 693 | 5.86 | | | | 2 | 1 | 1 | | | | 2 | 1 | 1 |
| High | Q5JPH6 | Probable glutamate--tRNA ligase, mitochondrial OS=Homo sapiens OX=9606 GN=EARS2 PE=1 SV=2 | 4 | 2 | 2 | 58.7 | 5 | 523 | 8.76 | 1 | 1 | 1 | | | | 1 | 1 | 1 | | | |
| High | P61201 | COP9 signalosome complex subunit 2 OS=Homo sapiens OX=9606 GN=COPS2 PE=1 SV=1 | 4 | 1 | 1 | 51.6 | 6 | 443 | 5.53 | | | 1 | | | | | | 1 | | | |
| High | O75689 | Arf-GAP with dual PH domain-containing protein 1 OS=Homo sapiens OX=9606 GN=ADAP1 PE=1 SV=2 | 4 | 2 | 2 | 43.4 | 7 | 374 | 8.97 | | | 1 | 1 | | 1 | | | 1 | 1 | | 1 |
| High | Q16775 | Hydroxyacylglutathione hydrolase, mitochondrial OS=Homo sapiens OX=9606 GN=HAGH PE=1 SV=2 | 4 | 1 | 1 | 33.8 | 5 | 308 | 8.12 | | | 1 | | | | | | 1 | | | |
| High | O94819 | Kelch repeat and BTB domain-containing protein 11 OS=Homo sapiens OX=9606 GN=KBTBD11 PE=1 SV=1 | 4 | 2 | 2 | 65.7 | 4 | 623 | 6.07 | | | | | | | | | | | | |
| High | P13798 | Acylamino-acid-releasing enzyme OS=Homo sapiens OX=9606 GN=APEH PE=1 SV=4 | 4 | 2 | 2 | 81.2 | 4 | 732 | 5.48 | 1 | 1 | 1 | | 1 | | 1 | 1 | 1 | | 1 | |
| High | Q7L5N1 | COP9 signalosome complex subunit 6 OS=Homo sapiens OX=9606 GN=COPS6 PE=1 SV=1 | 4 | 3 | 3 | 36.1 | 12 | 327 | 5.73 | | | | | 1 | | | | | | 1 | |
| High | Q68DH5 | G-protein coupled receptor-associated protein LMBRD2 OS=Homo sapiens OX=9606 GN=LMBRD2 PE=1 SV=1 | 4 | 3 | 3 | 81.1 | 6 | 695 | 7.5 | 2 | 1 | 1 | | | | 2 | 1 | 1 | | | |
| High | Q5W0U4 | B box and SPRY domain-containing protein OS=Homo sapiens OX=9606 GN=BSPRY PE=2 SV=1 | 4 | 2 | 2 | 44.4 | 5 | 402 | 6.42 | | | | 1 | | 1 | | | | 1 | | 1 |
| High | Q9Y371 | Endophilin-B1 OS=Homo sapiens OX=9606 GN=SH3GLB1 PE=1 SV=1 | 4 | 2 | 2 | 40.8 | 6 | 365 | 6.04 | | 1 | | | | | | 1 | | | | |
| High | Q15629 | Translocating chain-associated membrane protein 1 OS=Homo sapiens OX=9606 GN=TRAM1 PE=1 SV=3 | 4 | 1 | 1 | 43 | 3 | 374 | 9.63 | 1 | 1 | 1 | | | | 1 | 1 | 1 | | | |
| High | O15243 | Leptin receptor gene-related protein OS=Homo sapiens OX=9606 GN=LEPROT PE=1 SV=1 | 4 | 1 | 1 | 14.2 | 10 | 131 | 6.51 | 1 | 1 | | | | | 1 | 1 | | | | |
| High | Q8N884 | Cyclic GMP-AMP synthase OS=Homo sapiens OX=9606 GN=CGAS PE=1 SV=2 | 4 | 2 | 2 | 58.8 | 4 | 522 | 9.48 | 1 | 1 | 2 | | | | 1 | 1 | 2 | | | |
| High | Q9UNQ0 | Broad substrate specificity ATP-binding cassette transporter ABCG2 OS=Homo sapiens OX=9606 GN=ABCG2 PE=1 SV=3 | 4 | 3 | 3 | 72.3 | 4 | 655 | 8.69 | | | 1 | | | | | | 1 | | | |
| High | Q9BSC4 | Nucleolar protein 10 OS=Homo sapiens OX=9606 GN=NOL10 PE=1 SV=1 | 4 | 1 | 1 | 80.3 | 1 | 688 | 8.46 | | | | | 1 | | | | | | 1 | |
| High | Q9UL15 | BAG family molecular chaperone regulator 5 OS=Homo sapiens OX=9606 GN=BAG5 PE=1 SV=1 | 4 | 2 | 2 | 51.2 | 5 | 447 | 6.05 | 1 | 2 | | | | 1 | 1 | 2 | | | | 1 |
| High | P05413 | Fatty acid-binding protein, heart OS=Homo sapiens OX=9606 GN=FABP3 PE=1 SV=4 | 4 | 2 | 2 | 14.8 | 12 | 133 | 6.8 | | | | 1 | 1 | 2 | | | | 1 | 1 | 2 |
| High | Q13217 | DnaJ homolog subfamily C member 3 OS=Homo sapiens OX=9606 GN=DNAJC3 PE=1 SV=1 | 4 | 1 | 1 | 57.5 | 3 | 504 | 6.15 | | | 1 | | | | | | 1 | | | |
| High | Q96DH6 | RNA-binding protein Musashi homolog 2 OS=Homo sapiens OX=9606 GN=MSI2 PE=1 SV=1 | 4 | 3 | 3 | 35.2 | 12 | 328 | 8.48 | | | | | | | | | | | | |
| High | Q05443 | SWISS-PROT:Q05443 (Bos taurus) Lumican precursor | 4 | 3 | 3 | 38.7 | 8 | 342 | 6.35 | | | | 1 | 1 | 2 | | | | 1 | 1 | 2 |
| High | P30043 | Flavin reductase (NADPH) OS=Homo sapiens OX=9606 GN=BLVRB PE=1 SV=3 | 4 | 1 | 1 | 22.1 | 5 | 206 | 7.65 | 1 | | | | | | 1 | | | | | |
| High | P63272 | Transcription elongation factor SPT4 OS=Homo sapiens OX=9606 GN=SUPT4H1 PE=1 SV=1 | 4 | 1 | 1 | 13.2 | 6 | 117 | 8.06 | | | | | 1 | 1 | | | | | 1 | 1 |
| High | P29375 | Lysine-specific demethylase 5A OS=Homo sapiens OX=9606 GN=KDM5A PE=1 SV=3 | 4 | 2 | 1 | 192 | 2 | 1690 | 6.49 | | | | | | | | | | | | |
| High | O75528 | Transcriptional adapter 3 OS=Homo sapiens OX=9606 GN=TADA3 PE=1 SV=1 | 4 | 1 | 1 | 48.9 | 5 | 432 | 6.27 | | | | | 1 | 1 | | | | | 1 | 1 |
| High | O15460 | Prolyl 4-hydroxylase subunit alpha-2 OS=Homo sapiens OX=9606 GN=P4HA2 PE=1 SV=1 | 4 | 4 | 4 | 60.9 | 10 | 535 | 5.71 | | | 2 | | | | | | 2 | | | |
| High | Q8IUD2 | ELKS/Rab6-interacting/CAST family member 1 OS=Homo sapiens OX=9606 GN=ERC1 PE=1 SV=1 | 4 | 3 | 3 | 128 | 3 | 1116 | 5.97 | | | 3 | | | | | | 3 | | | |
| High | Q8TC07 | TBC1 domain family member 15 OS=Homo sapiens OX=9606 GN=TBC1D15 PE=1 SV=2 | 4 | 3 | 3 | 79.4 | 4 | 691 | 5.67 | | 2 | | | | | | 2 | | | | |
| High | P55198 | Protein AF-17 OS=Homo sapiens OX=9606 GN=MLLT6 PE=1 SV=3 | 4 | 1 | 1 | 112 | 1 | 1093 | 8.62 | | | | 1 | | 1 | | | | 1 | | 1 |
| High | Q8IZX4 | Transcription initiation factor TFIID subunit 1-like OS=Homo sapiens OX=9606 GN=TAF1L PE=1 SV=1 | 4 | 2 | 2 | 207.2 | 1 | 1826 | 5.4 | | | | | | | | | | | | |
| High | P48147 | Prolyl endopeptidase OS=Homo sapiens OX=9606 GN=PREP PE=1 SV=2 | 4 | 3 | 3 | 80.6 | 6 | 710 | 5.86 | 1 | 1 | | | | | 1 | 1 | | | | |
| High | Q9Y4Z0 | U6 snRNA-associated Sm-like protein LSm4 OS=Homo sapiens OX=9606 GN=LSM4 PE=1 SV=1 | 4 | 1 | 1 | 15.3 | 7 | 139 | 9.99 | | | | | 1 | | | | | | 1 | |
| High | Q5VYS8 | Terminal uridylyltransferase 7 OS=Homo sapiens OX=9606 GN=TUT7 PE=1 SV=1 | 4 | 2 | 2 | 171.1 | 2 | 1495 | 6.83 | | | | | 1 | | | | | | 1 | |
| High | Q9H8N7 | Zinc finger protein 395 OS=Homo sapiens OX=9606 GN=ZNF395 PE=1 SV=2 | 4 | 1 | 1 | 54.9 | 4 | 513 | 7.43 | | | 1 | | | | | | 1 | | | |
| High | P20591 | Interferon-induced GTP-binding protein Mx1 OS=Homo sapiens OX=9606 GN=MX1 PE=1 SV=4 | 4 | 2 | 2 | 75.5 | 5 | 662 | 5.83 | | | | 1 | 1 | 2 | | | | 1 | 1 | 2 |
| High | Q96HP4 | Oxidoreductase NAD-binding domain-containing protein 1 OS=Homo sapiens OX=9606 GN=OXNAD1 PE=1 SV=1 | 4 | 2 | 2 | 34.8 | 6 | 312 | 8.37 | | 1 | | | | | | 1 | | | | |
| High | P49755 | Transmembrane emp24 domain-containing protein 10 OS=Homo sapiens OX=9606 GN=TMED10 PE=1 SV=2 | 4 | 3 | 3 | 25 | 19 | 219 | 7.44 | 1 | 1 | 1 | | 1 | | 1 | 1 | 1 | | 1 | |
| High | Q9BXM0 | Periaxin OS=Homo sapiens OX=9606 GN=PRX PE=1 SV=2 | 4 | 2 | 2 | 154.8 | 1 | 1461 | 7.5 | | | | 1 | 2 | 1 | | | | 1 | 2 | 1 |
| High | Q9H019 | Mitochondrial fission regulator 1-like OS=Homo sapiens OX=9606 GN=MTFR1L PE=1 SV=2 | 4 | 2 | 2 | 31.9 | 11 | 292 | 6.1 | 1 | 1 | 1 | | | | 1 | 1 | 1 | | | |
| High | Q92766 | Ras-responsive element-binding protein 1 OS=Homo sapiens OX=9606 GN=RREB1 PE=1 SV=3 | 4 | 2 | 2 | 181.3 | 2 | 1687 | 6.98 | | | 1 | | | | | | 1 | | | |
| High | Q9NVZ3 | Adaptin ear-binding coat-associated protein 2 OS=Homo sapiens OX=9606 GN=NECAP2 PE=1 SV=1 | 4 | 2 | 2 | 28.3 | 13 | 263 | 8.38 | 1 | | 1 | | 1 | | 1 | | 1 | | 1 | |
| High | Q96BN8 | Ubiquitin thioesterase otulin OS=Homo sapiens OX=9606 GN=OTULIN PE=1 SV=3 | 4 | 2 | 2 | 40.2 | 5 | 352 | 5.47 | | | 1 | | | | | | 1 | | | |
| High | Q9BRR6 | ADP-dependent glucokinase OS=Homo sapiens OX=9606 GN=ADPGK PE=1 SV=1 | 4 | 1 | 1 | 54.1 | 5 | 497 | 6.2 | 1 | 1 | | | | | 1 | 1 | | | | |
| High | P68402 | Platelet-activating factor acetylhydrolase IB subunit alpha2 OS=Homo sapiens OX=9606 GN=PAFAH1B2 PE=1 SV=1 | 4 | 1 | 1 | 25.6 | 8 | 229 | 5.92 | 1 | 2 | 1 | | | | 1 | 1 | 1 | | | |
| High | Q92688 | Acidic leucine-rich nuclear phosphoprotein 32 family member B OS=Homo sapiens OX=9606 GN=ANP32B PE=1 SV=1 | 4 | 3 | 3 | 28.8 | 14 | 251 | 4.06 | | 1 | | | 1 | | | 1 | | | 1 | |
| High | Q16637 | Survival motor neuron protein OS=Homo sapiens OX=9606 GN=SMN1 PE=1 SV=1 | 4 | 2 | 2 | 31.8 | 10 | 294 | 6.55 | | | | 1 | 1 | 1 | | | | 1 | 1 | 1 |
| High | Q9Y2P8 | RNA 3'-terminal phosphate cyclase-like protein OS=Homo sapiens OX=9606 GN=RCL1 PE=1 SV=3 | 4 | 2 | 2 | 40.8 | 7 | 373 | 9.26 | | | | | | | | | | | | |
| High | Q9NXJ5 | Pyroglutamyl-peptidase 1 OS=Homo sapiens OX=9606 GN=PGPEP1 PE=1 SV=1 | 4 | 2 | 2 | 23.1 | 13 | 209 | 5.9 | 1 | 1 | 1 | | | | 1 | 1 | 1 | | | |
| High | Q8TER5 | Rho guanine nucleotide exchange factor 40 OS=Homo sapiens OX=9606 GN=ARHGEF40 PE=1 SV=3 | 4 | 3 | 3 | 164.6 | 3 | 1519 | 6.15 | | | | | | | | | | | | |
| High | Q8NHH9 | Atlastin-2 OS=Homo sapiens OX=9606 GN=ATL2 PE=1 SV=2 | 4 | 2 | 2 | 66.2 | 7 | 583 | 5.48 | | 1 | 1 | | | | | 1 | 1 | | | |
| High | Q96NL6 | Sodium channel and clathrin linker 1 OS=Homo sapiens OX=9606 GN=SCLT1 PE=1 SV=2 | 4 | 2 | 2 | 80.9 | 2 | 688 | 6.07 | 1 | 1 | 2 | | | | 1 | 1 | 2 | | | |
| High | Q8IYT2 | Cap-specific mRNA (nucleoside-2'-O-)-methyltransferase 2 OS=Homo sapiens OX=9606 GN=CMTR2 PE=1 SV=2 | 4 | 1 | 1 | 88.1 | 1 | 770 | 7.01 | 1 | 1 | | | | | 1 | 1 | | | | |
| High | P61769 | Beta-2-microglobulin OS=Homo sapiens OX=9606 GN=B2M PE=1 SV=1 | 4 | 2 | 2 | 13.7 | 27 | 119 | 6.52 | 1 | 2 | 1 | | | | 1 | 2 | 1 | | | |
| High | Q6P1J9 | Parafibromin OS=Homo sapiens OX=9606 GN=CDC73 PE=1 SV=1 | 4 | 3 | 3 | 60.5 | 6 | 531 | 9.61 | | | | | 1 | | | | | | 1 | |
| High | Q96RN5 | Mediator of RNA polymerase II transcription subunit 15 OS=Homo sapiens OX=9606 GN=MED15 PE=1 SV=2 | 4 | 1 | 1 | 86.7 | 3 | 788 | 9.42 | | | 1 | | | | | | 1 | | | |
| High | P25686 | DnaJ homolog subfamily B member 2 OS=Homo sapiens OX=9606 GN=DNAJB2 PE=1 SV=3 | 4 | 2 | 1 | 35.6 | 8 | 324 | 5.95 | 1 | 1 | 2 | | | | 1 | 1 | 2 | | | |
| High | Q7Z589 | BRCA2-interacting transcriptional repressor EMSY OS=Homo sapiens OX=9606 GN=EMSY PE=1 SV=2 | 4 | 2 | 2 | 141.4 | 2 | 1322 | 9.33 | | | | | 1 | | | | | | 1 | |
| High | Q5JSZ5 | Protein PRRC2B OS=Homo sapiens OX=9606 GN=PRRC2B PE=1 SV=2 | 4 | 1 | 1 | 242.8 | 1 | 2229 | 8.34 | 1 | 1 | 1 | | | | 1 | 1 | 1 | | | |
| High | Q96JC1 | Vam6/Vps39-like protein OS=Homo sapiens OX=9606 GN=VPS39 PE=1 SV=2 | 4 | 1 | 1 | 101.7 | 2 | 886 | 6.99 | 1 | | 1 | | | | 1 | | 1 | | | |
| High | P12830 | Cadherin-1 OS=Homo sapiens OX=9606 GN=CDH1 PE=1 SV=3 | 4 | 1 | 1 | 97.4 | 1 | 882 | 4.73 | | | 1 | | 1 | | | | 1 | | 1 | |
| High | P49005 | DNA polymerase delta subunit 2 OS=Homo sapiens OX=9606 GN=POLD2 PE=1 SV=1 | 4 | 2 | 2 | 51.3 | 5 | 469 | 5.58 | | | | | | | | | | | | |
| High | P12955 | Xaa-Pro dipeptidase OS=Homo sapiens OX=9606 GN=PEPD PE=1 SV=3 | 4 | 2 | 2 | 54.5 | 3 | 493 | 6 | | | | 1 | 2 | 1 | | | | 1 | 2 | 1 |
| High | Q9NWZ8 | Gem-associated protein 8 OS=Homo sapiens OX=9606 GN=GEMIN8 PE=1 SV=1 | 4 | 2 | 2 | 28.6 | 12 | 242 | 6.8 | | | | | 2 | | | | | | 2 | |
| High | Q8N7H5 | RNA polymerase II-associated factor 1 homolog OS=Homo sapiens OX=9606 GN=PAF1 PE=1 SV=2 | 4 | 2 | 2 | 59.9 | 5 | 531 | 4.63 | | | | | | | | | | | | |
| High | O60293 | Zinc finger C3H1 domain-containing protein OS=Homo sapiens OX=9606 GN=ZFC3H1 PE=1 SV=3 | 4 | 3 | 3 | 226.2 | 2 | 1989 | 8.13 | | | | | | | | | | | | |
| High | Q15269 | Periodic tryptophan protein 2 homolog OS=Homo sapiens OX=9606 GN=PWP2 PE=1 SV=2 | 4 | 1 | 1 | 102.4 | 1 | 919 | 6.15 | | | | | 1 | | | | | | 1 | |
| High | Q8N5N7 | 39S ribosomal protein L50, mitochondrial OS=Homo sapiens OX=9606 GN=MRPL50 PE=1 SV=2 | 4 | 2 | 2 | 18.3 | 22 | 158 | 7.88 | 1 | | | | | | 1 | | | | | |
| High | Q9NQW7 | Xaa-Pro aminopeptidase 1 OS=Homo sapiens OX=9606 GN=XPNPEP1 PE=1 SV=3 | 4 | 1 | 1 | 69.9 | 1 | 623 | 5.67 | 1 | 1 | 1 | | | | 1 | 1 | 1 | | | |
| High | Q5T5U3 | Rho GTPase-activating protein 21 OS=Homo sapiens OX=9606 GN=ARHGAP21 PE=1 SV=2 | 4 | 2 | 2 | 217.3 | 1 | 1958 | 7.8 | | | 1 | | 1 | | | | 1 | | 1 | |
| High | Q9BXK1 | Krueppel-like factor 16 OS=Homo sapiens OX=9606 GN=KLF16 PE=1 SV=1 | 4 | 2 | 2 | 25.4 | 17 | 252 | 9.88 | | | | 1 | 2 | 1 | | | | 1 | 2 | 1 |
| High | Q96B26 | Exosome complex component RRP43 OS=Homo sapiens OX=9606 GN=EXOSC8 PE=1 SV=1 | 4 | 2 | 2 | 30 | 7 | 276 | 5.3 | | | | | | 1 | | | | | | 1 |
| High | O43172 | U4/U6 small nuclear ribonucleoprotein Prp4 OS=Homo sapiens OX=9606 GN=PRPF4 PE=1 SV=2 | 4 | 2 | 2 | 58.4 | 8 | 522 | 7.42 | | | | | | | | | | | | |
| High | P61923 | Coatomer subunit zeta-1 OS=Homo sapiens OX=9606 GN=COPZ1 PE=1 SV=1 | 4 | 2 | 2 | 20.2 | 12 | 177 | 4.81 | 2 | 1 | | | | | 2 | 1 | | | | |
| High | O43639 | Cytoplasmic protein NCK2 OS=Homo sapiens OX=9606 GN=NCK2 PE=1 SV=2 | 4 | 1 | 1 | 42.9 | 5 | 380 | 6.95 | 1 | 1 | 1 | | | | 1 | 1 | 1 | | | |

| | | | | | | | | | | | | | | | | | | | | | |
|---|---|---|---|---|---|---|---|---|---|---|---|---|---|---|---|---|---|---|---|---|---|
| High | Q15042 | Rab3 GTPase-activating protein catalytic subunit OS=Homo sapiens OX=9606 GN=RAB3GAP1 PE=1 SV=3 | 4 | 2 | 2 | 110.5 | 2 | 981 | 5.55 | 2 | 1 | 1 | | | | 2 | 1 | 1 | | | |
| High | Q16204 | Coiled-coil domain-containing protein 6 OS=Homo sapiens OX=9606 GN=CCDC6 PE=1 SV=2 | 4 | 4 | 4 | 53.3 | 9 | 474 | 7.34 | | | | 1 | | | | | | 1 | | |
| High | O15111 | Inhibitor of nuclear factor kappa-B kinase subunit alpha OS=Homo sapiens OX=9606 GN=CHUK PE=1 SV=2 | 4 | 1 | 1 | 84.6 | 2 | 745 | 6.73 | 1 | 1 | 1 | | | | 1 | 1 | 1 | | | |
| High | P24928 | DNA-directed RNA polymerase II subunit RPB1 OS=Homo sapiens OX=9606 GN=POLR2A PE=1 SV=2 | 4 | 2 | 2 | 217 | 2 | 1970 | 7.37 | | | | | | | | | | | | |
| High | Q9NRR3 | CDC42 small effector protein 2 OS=Homo sapiens OX=9606 GN=CDC42SE2 PE=1 SV=1 | 4 | 1 | 1 | 9.2 | 23 | 84 | 8.35 | | | | | | | | | | | | |
| High | Q8NE86 | Calcium uniporter protein, mitochondrial OS=Homo sapiens OX=9606 GN=MCU PE=1 SV=1 | 4 | 2 | 2 | 39.8 | 9 | 351 | 8.65 | 1 | 2 | 1 | | | | 1 | 2 | 1 | | | |
| High | Q13409 | Cytoplasmic dynein 1 intermediate chain 2 OS=Homo sapiens OX=9606 GN=DYNC1I2 PE=1 SV=3 | 4 | 1 | 1 | 71.4 | 2 | 638 | 5.2 | | | | | 1 | 1 | | | | | 1 | 1 |
| High | Q8NFH5 | Nucleoporin NUP35 OS=Homo sapiens OX=9606 GN=NUP35 PE=1 SV=1 | 4 | 1 | 1 | 34.8 | 6 | 326 | 9.09 | | | 1 | | | | | | 1 | | | |
| High | Q01804 | OTU domain-containing protein 4 OS=Homo sapiens OX=9606 GN=OTUD4 PE=1 SV=4 | 4 | 1 | 1 | 124 | 1 | 1114 | 6.71 | 1 | 1 | 1 | | | | 1 | 1 | 1 | | | |
| High | P11717 | Cation-independent mannose-6-phosphate receptor OS=Homo sapiens OX=9606 GN=IGF2R PE=1 SV=3 | 4 | 2 | 2 | 274.2 | 1 | 2491 | 5.94 | | | 1 | | | | | | 1 | | | |
| High | O95674 | Phosphatidate cytidylyltransferase 2 OS=Homo sapiens OX=9606 GN=CDS2 PE=1 SV=1 | 4 | 1 | 1 | 51.4 | 4 | 445 | 7.09 | 1 | 1 | 2 | | | | 1 | 1 | 1 | | | |
| High | O94818 | Nucleolar protein 4 OS=Homo sapiens OX=9606 GN=NOL4 PE=1 SV=2 | 4 | 1 | 1 | 71.3 | 1 | 638 | 5.62 | | | | 1 | 1 | | | | | 1 | 1 | |
| High | P27986 | Phosphatidylinositol 3-kinase regulatory subunit alpha OS=Homo sapiens OX=9606 GN=PIK3R1 PE=1 SV=2 | 4 | 3 | 3 | 83.5 | 7 | 724 | 6.16 | | | 1 | | 1 | | | | 1 | | 1 | |
| High | P42356 | Phosphatidylinositol 4-kinase alpha OS=Homo sapiens OX=9606 GN=PI4KA PE=1 SV=4 | 4 | 2 | 2 | 236.7 | 1 | 2102 | 7.06 | 1 | 2 | 1 | | | | 1 | 2 | 1 | | | |
| High | Q93062 | RNA-binding protein with multiple splicing OS=Homo sapiens OX=9606 GN=RBPMS PE=1 SV=1 | 4 | 2 | 2 | 21.8 | 20 | 196 | 8.07 | 1 | | 1 | | | | 1 | | 1 | | | |
| High | Q9NP61 | ADP-ribosylation factor GTPase-activating protein 3 OS=Homo sapiens OX=9606 GN=ARFGAP3 PE=1 SV=1 | 4 | 2 | 2 | 56.9 | 5 | 516 | 7.36 | | 1 | | | | | | 1 | | | | |
| High | Q6ZN55 | Zinc finger protein 574 OS=Homo sapiens OX=9606 GN=ZNF574 PE=1 SV=2 | 4 | 2 | 2 | 98.8 | 3 | 896 | 8.07 | | | | | 1 | | | | | | 1 | |
| High | Q8IWB7 | WD repeat and FYVE domain-containing protein 1 OS=Homo sapiens OX=9606 GN=WDFY1 PE=1 SV=1 | 4 | 2 | 2 | 46.3 | 8 | 410 | 7.33 | 1 | 1 | 2 | | | | 1 | 1 | 2 | | | |
| High | Q9H9A5 | CCR4-NOT transcription complex subunit 10 OS=Homo sapiens OX=9606 GN=CNOT10 PE=1 SV=1 | 4 | 2 | 2 | 82.3 | 3 | 744 | 7.78 | | | | | | | | | | | | |
| High | P42696 | RNA-binding protein 34 OS=Homo sapiens OX=9606 GN=RBM34 PE=1 SV=2 | 4 | 3 | 3 | 48.5 | 7 | 430 | 10.11 | | | | | 1 | 1 | | | | | 1 | 1 |
| High | P12429 | Annexin A3 OS=Homo sapiens OX=9606 GN=ANXA3 PE=1 SV=3 | 4 | 2 | 2 | 36.4 | 5 | 323 | 5.92 | | | | 2 | 1 | 1 | | | | 2 | 1 | 1 |
| High | Q9BUK6 | Protein misato homolog 1 OS=Homo sapiens OX=9606 GN=MSTO1 PE=1 SV=1 | 4 | 3 | 3 | 61.8 | 8 | 570 | 6.11 | | | 3 | 1 | | | | | 3 | 1 | | |
| High | P19784 | Casein kinase II subunit alpha' OS=Homo sapiens OX=9606 GN=CSNK2A2 PE=1 SV=1 | 4 | 3 | 3 | 41.2 | 13 | 350 | 8.56 | | | 1 | | | | | | 1 | | | |
| High | Q9BRJ2 | 39S ribosomal protein L45, mitochondrial OS=Homo sapiens OX=9606 GN=MRPL45 PE=1 SV=2 | 4 | 1 | 1 | 35.3 | 3 | 306 | 9.03 | | | | | 1 | | | | | | 1 | |
| High | P01040 | Cystatin-A OS=Homo sapiens OX=9606 GN=CSTA PE=1 SV=1 | 4 | 3 | 3 | 11 | 38 | 98 | 5.5 | | | | 1 | 3 | | | | | 1 | 3 | |
| High | P51572 | B-cell receptor-associated protein 31 OS=Homo sapiens OX=9606 GN=BCAP31 PE=1 SV=3 | 4 | 2 | 2 | 28 | 7 | 246 | 8.44 | | | 1 | | | | | | 1 | | | |
| High | P20226 | TATA-box-binding protein OS=Homo sapiens OX=9606 GN=TBP PE=1 SV=2 | 4 | 2 | 2 | 37.7 | 7 | 339 | 9.79 | 1 | 2 | 1 | | | | 1 | 2 | 1 | | | |
| High | Q9H814 | Phosphorylated adapter RNA export protein OS=Homo sapiens OX=9606 GN=PHAX PE=1 SV=1 | 4 | 2 | 2 | 44.4 | 8 | 394 | 5.4 | | | | 1 | 1 | 1 | | | | 1 | 1 | 1 |
| High | P50548 | ETS domain-containing transcription factor ERF OS=Homo sapiens OX=9606 GN=ERF PE=1 SV=2 | 4 | 1 | 1 | 58.7 | 4 | 548 | 7.3 | 1 | 1 | 1 | | | | 1 | 1 | 1 | | | |
| High | Q9HA92 | Radical S-adenosyl methionine domain-containing protein 1, mitochondrial OS=Homo sapiens OX=9606 GN=RSAD1 PE=1 SV=2 | 4 | 2 | 2 | 48.7 | 4 | 442 | 7.78 | | | | | | | | | | | | |
| High | Q86WJ1 | Chromodomain-helicase-DNA-binding protein 1-like OS=Homo sapiens OX=9606 GN=CHD1L PE=1 SV=3 | 4 | 3 | 3 | 100.9 | 4 | 897 | 6.9 | | 1 | 1 | | | | | 1 | 1 | | | |
| High | Q96MU7 | YTH domain-containing protein 1 OS=Homo sapiens OX=9606 GN=YTHDC1 PE=1 SV=3 | 4 | 1 | 1 | 84.6 | 2 | 727 | 6.23 | | | 1 | | | | | | 1 | | | |
| High | Q9GZP9 | Derlin-2 OS=Homo sapiens OX=9606 GN=DERL2 PE=1 SV=1 | 4 | 1 | 1 | 27.5 | 3 | 239 | 7.28 | | 1 | 1 | 1 | | 1 | | 1 | 1 | 1 | | 1 |
| High | Q8WUA7 | TBC1 domain family member 22A OS=Homo sapiens OX=9606 GN=TBC1D22A PE=1 SV=2 | 4 | 3 | 3 | 59.1 | 7 | 517 | 5.97 | 2 | 1 | 1 | | | | 2 | 1 | 1 | | | |
| High | Q8NHM5 | Lysine-specific demethylase 2B OS=Homo sapiens OX=9606 GN=KDM2B PE=1 SV=1 | 4 | 2 | 2 | 152.5 | 2 | 1336 | 8.56 | | | | 1 | 2 | 1 | | | | 1 | 2 | 1 |
| High | Q16594 | Transcription initiation factor TFIID subunit 9 OS=Homo sapiens OX=9606 GN=TAF9 PE=1 SV=1 | 4 | 3 | 3 | 29 | 16 | 264 | 8.66 | | | | | 1 | | | | | | 1 | |
| High | Q8N4J0 | Carnosine N-methyltransferase OS=Homo sapiens OX=9606 GN=CARNMT1 PE=1 SV=1 | 4 | 1 | 1 | 47.2 | 3 | 409 | 6.32 | | | | | 1 | | | | | | 1 | |
| High | P82664 | 28S ribosomal protein S10, mitochondrial OS=Homo sapiens OX=9606 GN=MRPS10 PE=1 SV=2 | 4 | 2 | 2 | 23 | 15 | 201 | 8 | | | | | | | | | | | | |
| High | P61225 | Ras-related protein Rap-2b OS=Homo sapiens OX=9606 GN=RAP2B PE=1 SV=1 | 4 | 2 | 2 | 20.5 | 14 | 183 | 4.81 | | | | | | | | | | | | |
| High | O43395 | U4/U6 small nuclear ribonucleoprotein Prp3 OS=Homo sapiens OX=9606 GN=PRPF3 PE=1 SV=2 | 4 | 2 | 2 | 77.5 | 6 | 683 | 9.5 | | | | 1 | | | | | | 1 | | |
| High | Q9NVA2 | Septin-11 OS=Homo sapiens OX=9606 GN=SEPTIN11 PE=1 SV=3 | 4 | 2 | 1 | 49.4 | 7 | 429 | 6.81 | 1 | | 1 | 1 | | | 1 | | 1 | 1 | | |
| High | O43747 | AP-1 complex subunit gamma-1 OS=Homo sapiens OX=9606 GN=AP1G1 PE=1 SV=5 | 4 | 2 | 2 | 91.3 | 2 | 822 | 6.8 | | 1 | 1 | | 1 | | | 1 | 1 | | 1 | |
| High | Q9H3P2 | Negative elongation factor A OS=Homo sapiens OX=9606 GN=NELFA PE=1 SV=3 | 4 | 1 | 1 | 57.2 | 4 | 528 | 9.03 | | | | 1 | | 1 | | | | 1 | | 1 |
| High | Q9HCK0 | Zinc finger and BTB domain-containing protein 26 OS=Homo sapiens OX=9606 GN=ZBTB26 PE=1 SV=2 | 4 | 1 | 1 | 49.9 | 4 | 441 | 7.08 | | | 1 | | | | | | 1 | | | |
| High | P49593 | Protein phosphatase 1F OS=Homo sapiens OX=9606 GN=PPM1F PE=1 SV=3 | 4 | 1 | 1 | 49.8 | 2 | 454 | 5.1 | | 1 | | | | | | 1 | | | | |
| High | Q8IWW6 | Rho GTPase-activating protein 12 OS=Homo sapiens OX=9606 GN=ARHGAP12 PE=1 SV=1 | 4 | 2 | 2 | 96.2 | 4 | 846 | 7.61 | | 1 | 1 | | | 1 | | 1 | 1 | | | 1 |
| High | A1X283 | SH3 and PX domain-containing protein 2B OS=Homo sapiens OX=9606 GN=SH3PXD2B PE=1 SV=3 | 4 | 2 | 2 | 101.5 | 4 | 911 | 8.69 | | | | 1 | 1 | 2 | | | | 1 | 1 | 2 |
| High | Q8IYU8 | Calcium uptake protein 2, mitochondrial OS=Homo sapiens OX=9606 GN=MICU2 PE=1 SV=2 | 4 | 2 | 2 | 49.6 | 9 | 434 | 9.09 | 1 | | 2 | | | | 1 | | 2 | | | |
| High | Q14142 | Tripartite motif-containing protein 14 OS=Homo sapiens OX=9606 GN=TRIM14 PE=1 SV=2 | 4 | 2 | 2 | 49.7 | 6 | 442 | 7.87 | | | | | | | | | | | | |
| High | Q99627 | COP9 signalosome complex subunit 8 OS=Homo sapiens OX=9606 GN=COPS8 PE=1 SV=1 | 4 | 1 | 1 | 23.2 | 10 | 209 | 5.38 | | | | | 1 | 1 | | | | | 1 | 1 |
| High | Q9NVU0 | DNA-directed RNA polymerase III subunit RPC5 OS=Homo sapiens OX=9606 GN=POLR3E PE=1 SV=1 | 4 | 2 | 2 | 79.8 | 4 | 708 | 6.47 | | | | | | | | | | | | |
| High | Q9NU22 | Midasin OS=Homo sapiens OX=9606 GN=MDN1 PE=1 SV=2 | 4 | 3 | 3 | 632.4 | 1 | 5596 | 5.68 | | 2 | | | | | | 1 | | | | |
| High | Q12768 | WASH complex subunit 5 OS=Homo sapiens OX=9606 GN=WASHC5 PE=1 SV=1 | 4 | 1 | 1 | 134.2 | 1 | 1159 | 6.98 | 1 | 1 | | | | | 1 | 1 | | | | |
| High | Q99549 | M-phase phosphoprotein 8 OS=Homo sapiens OX=9606 GN=MPHOSPH8 PE=1 SV=2 | 4 | 2 | 2 | 97.1 | 4 | 860 | 6.06 | | | | | | | | | | | | |
| High | P82912 | 28S ribosomal protein S11, mitochondrial OS=Homo sapiens OX=9606 GN=MRPS11 PE=1 SV=2 | 4 | 1 | 1 | 20.6 | 8 | 194 | 10.81 | | | | 1 | | | | | | 1 | | |
| High | Q8N5P1 | Zinc finger CCCH domain-containing protein 8 OS=Homo sapiens OX=9606 GN=ZC3H8 PE=1 SV=2 | 4 | 1 | 1 | 33.6 | 4 | 291 | 8.28 | | | | 1 | | | | | | 1 | | |
| High | P37837 | Transaldolase OS=Homo sapiens OX=9606 GN=TALDO1 PE=1 SV=2 | 4 | 2 | 2 | 37.5 | 6 | 337 | 6.81 | 1 | 1 | 2 | | | | 1 | 1 | 2 | | | |
| High | Q86X10 | Ral GTPase-activating protein subunit beta OS=Homo sapiens OX=9606 GN=RALGAPB PE=1 SV=1 | 4 | 2 | 2 | 166.7 | 2 | 1494 | 6.79 | 1 | 1 | | | | | 1 | 1 | | | | |
| High | Q9UKM7 | Endoplasmic reticulum mannosyl-oligosaccharide 1,2-alpha-mannosidase OS=Homo sapiens OX=9606 GN=MAN1B1 PE=1 SV=2 | 4 | 2 | 2 | 79.5 | 5 | 699 | 7.72 | 1 | 1 | | | | | 1 | 1 | | | | |
| High | O14646 | Chromodomain-helicase-DNA-binding protein 1 OS=Homo sapiens OX=9606 GN=CHD1 PE=1 SV=2 | 4 | 2 | 1 | 196.6 | 1 | 1710 | 7.14 | 1 | 1 | 1 | | | | 1 | 1 | 1 | | | |
| High | P25325 | 3-mercaptopyruvate sulfurtransferase OS=Homo sapiens OX=9606 GN=MPST PE=1 SV=3 | 4 | 2 | 2 | 33.2 | 13 | 297 | 6.6 | | | 1 | | | | | | 1 | | | |
| High | Q9H4L4 | Sentrin-specific protease 3 OS=Homo sapiens OX=9606 GN=SENP3 PE=1 SV=2 | 4 | 2 | 2 | 65 | 4 | 574 | 8.56 | 1 | 1 | | | | | 1 | 1 | | | | |
| High | Q14002 | Carcinoembryonic antigen-related cell adhesion molecule 7 OS=Homo sapiens OX=9606 GN=CEACAM7 PE=1 SV=2 | 4 | 3 | 3 | 29.3 | 14 | 265 | 5.54 | | | | | | | | | | | | |
| High | Q96IJ6 | Mannose-1-phosphate guanyltransferase alpha OS=Homo sapiens OX=9606 GN=GMPPA PE=1 SV=1 | 4 | 2 | 2 | 46.3 | 5 | 420 | 7.21 | 1 | 1 | | | | | 1 | 1 | | | | |
| High | P00505 | Aspartate aminotransferase, mitochondrial OS=Homo sapiens OX=9606 GN=GOT2 PE=1 SV=3 | 4 | 1 | 1 | 47.5 | 2 | 430 | 9.01 | | | 1 | 1 | 1 | 1 | | | 1 | 1 | 1 | 1 |
| High | O43933 | Peroxisome biogenesis factor 1 OS=Homo sapiens OX=9606 GN=PEX1 PE=1 SV=1 | 4 | 2 | 2 | 142.8 | 1 | 1283 | 6.33 | | | 1 | | | | | | 1 | | | |
| High | Q8WTW3 | Conserved oligomeric Golgi complex subunit 1 OS=Homo sapiens OX=9606 GN=COG1 PE=1 SV=1 | 4 | 2 | 2 | 108.9 | 2 | 980 | 7.31 | 1 | 2 | 1 | | | | 1 | 2 | 1 | | | |
| High | Q9UBB6 | Neurochondrin OS=Homo sapiens OX=9606 GN=NCDN PE=1 SV=1 | 4 | 2 | 2 | 78.8 | 4 | 729 | 5.48 | 1 | 1 | 1 | | | | 1 | 1 | 1 | | | |
| High | Q9Y666 | Solute carrier family 12 member 7 OS=Homo sapiens OX=9606 GN=SLC12A7 PE=1 SV=3 | 4 | 1 | 1 | 119 | 2 | 1083 | 6.71 | | | 1 | | | | | | 1 | | | |
| High | O15228 | Dihydroxyacetone phosphate acyltransferase OS=Homo sapiens OX=9606 GN=GNPAT PE=1 SV=1 | 4 | 3 | 3 | 77.1 | 4 | 680 | 6.57 | | | | | 1 | | | | | | 1 | |
| High | P50395 | Rab GDP dissociation inhibitor beta OS=Homo sapiens OX=9606 GN=GDI2 PE=1 SV=2 | 4 | 2 | 1 | 50.6 | 6 | 445 | 6.47 | 1 | 1 | 1 | | | | 1 | 1 | 1 | | | |
| High | O00204 | Sulfotransferase 2B1 OS=Homo sapiens OX=9606 GN=SULT2B1 PE=1 SV=2 | 4 | 2 | 2 | 41.3 | 11 | 365 | 5.4 | 1 | | 1 | | | | 1 | | 1 | | | |
| High | P07108 | Acyl-CoA-binding protein OS=Homo sapiens OX=9606 GN=DBI PE=1 SV=2 | 4 | 1 | 1 | 10 | 23 | 87 | 6.57 | 1 | 1 | 1 | | | | 1 | 1 | 1 | | | |
| High | Q9H5V8 | CUB domain-containing protein 1 OS=Homo sapiens OX=9606 GN=CDCP1 PE=1 SV=3 | 4 | 1 | 1 | 92.9 | 2 | 836 | 7.96 | 1 | 1 | | | | | 1 | 1 | | | | |
| High | P35226 | Polycomb complex protein BMI-1 OS=Homo sapiens OX=9606 GN=BMI1 PE=1 SV=2 | 4 | 2 | 2 | 36.9 | 6 | 326 | 8.63 | | | | 1 | 1 | | | | | 1 | 1 | |
| High | Q9Y3C8 | Ubiquitin-fold modifier-conjugating enzyme 1 OS=Homo sapiens OX=9606 GN=UFC1 PE=1 SV=3 | 4 | 2 | 2 | 19.4 | 11 | 167 | 7.4 | | | | | | 1 | | | | | | 1 |
| High | Q13613 | Myotubularin-related protein 1 OS=Homo sapiens OX=9606 GN=MTMR1 PE=1 SV=4 | 4 | 3 | 3 | 74.6 | 9 | 665 | 7.14 | 1 | 2 | | | | | 1 | 2 | | | | |
| High | Q9UNE7 | E3 ubiquitin-protein ligase CHIP OS=Homo sapiens OX=9606 GN=STUB1 PE=1 SV=2 | 4 | 1 | 1 | 34.8 | 4 | 303 | 5.87 | | | | | 1 | 1 | | | | | 1 | 1 |
| High | O95801 | Tetratricopeptide repeat protein 4 OS=Homo sapiens OX=9606 GN=TTC4 PE=1 SV=3 | 4 | 1 | 1 | 44.7 | 3 | 387 | 5.6 | 1 | | | 1 | | 1 | 1 | | | 1 | | 1 |
| High | Q5JVS0 | Intracellular hyaluronan-binding protein 4 OS=Homo sapiens OX=9606 GN=HABP4 PE=1 SV=1 | 4 | 1 | 1 | 45.8 | 6 | 413 | 7.46 | 1 | 1 | | | 1 | | 1 | 1 | | | 1 | |
| High | P14314 | Glucosidase 2 subunit beta OS=Homo sapiens OX=9606 GN=PRKCSH PE=1 SV=2 | 4 | 2 | 2 | 59.4 | 4 | 528 | 4.41 | | 2 | 1 | | | | | 2 | 1 | | | |
| High | Q96A49 | Synapse-associated protein 1 OS=Homo sapiens OX=9606 GN=SYAP1 PE=1 SV=1 | 4 | 2 | 2 | 39.9 | 6 | 352 | 4.53 | | | | | | | | | | | | |
| High | Q96J02 | E3 ubiquitin-protein ligase Itchy homolog OS=Homo sapiens OX=9606 GN=ITCH PE=1 SV=2 | 4 | 2 | 1 | 102.7 | 3 | 903 | 6.3 | 1 | 1 | | | | | 1 | 1 | | | | |
| High | Q5SNT2 | Transmembrane protein 201 OS=Homo sapiens OX=9606 GN=TMEM201 PE=1 SV=1 | 4 | 2 | 2 | 72.2 | 3 | 666 | 9.22 | 1 | 1 | 1 | | | | 1 | 1 | 1 | | | |
| High | Q9H3S7 | Tyrosine-protein phosphatase non-receptor type 23 OS=Homo sapiens OX=9606 GN=PTPN23 PE=1 SV=1 | 4 | 2 | 2 | 178.9 | 2 | 1636 | 6.92 | | | | | 1 | | | | | | 1 | |
| High | Q9H400 | Lck-interacting transmembrane adapter 1 OS=Homo sapiens OX=9606 GN=LIME1 PE=1 SV=1 | 4 | 2 | 2 | 31.3 | 11 | 295 | 9.58 | | | | | | | | | | | | |
| High | Q08554 | Desmocollin-1 OS=Homo sapiens OX=9606 GN=DSC1 PE=1 SV=2 | 4 | 1 | 1 | 99.9 | 1 | 894 | 5.43 | | | 1 | 1 | 1 | 1 | | | 1 | 1 | 1 | 1 |
| High | Q96HW7 | Integrator complex subunit 4 OS=Homo sapiens OX=9606 GN=INTS4 PE=1 SV=2 | 4 | 2 | 2 | 108.1 | 3 | 963 | 6.44 | 1 | 1 | 1 | | | | 1 | 1 | 1 | | | |
| High | Q9NR12 | PDZ and LIM domain protein 7 OS=Homo sapiens OX=9606 GN=PDLIM7 PE=1 SV=1 | 4 | 3 | 3 | 49.8 | 7 | 457 | 8.41 | 1 | | 2 | | | | 1 | | 1 | | | |
| High | P02668 | SWISS-PROT:P02668 Kappa-casein [Contains: Casoxin C; Casoxin 6; Casoxin A; Casoxin B; Casoplatelin] - Bos taurus (Bovine). | 4 | 2 | 2 | 19 | 12 | 169 | 6.3 | | | | | 1 | | | | | | 1 | |
| High | O75884 | Serine hydrolase RBBP9 OS=Homo sapiens OX=9606 GN=RBBP9 PE=1 SV=2 | 4 | 2 | 2 | 21 | 11 | 186 | 6.2 | | | 1 | | | | | | 1 | | | |
| High | P62894 | SWISS-PROT:P62894 Cytochrome c - Bos taurus (Bovine). | 4 | 2 | 1 | 11.7 | 17 | 105 | 9.5 | | | 1 | 1 | 1 | 1 | | | 1 | 1 | 1 | 1 |
| High | Q16527 | Cysteine and glycine-rich protein 2 OS=Homo sapiens OX=9606 GN=CSRP2 PE=1 SV=3 | 4 | 3 | 3 | 20.9 | 18 | 193 | 8.62 | | | | 2 | 1 | 1 | | | | 2 | 1 | 1 |
| High | Q9H9Q4 | Non-homologous end-joining factor 1 OS=Homo sapiens OX=9606 GN=NHEJ1 PE=1 SV=1 | 3 | 1 | 1 | 33.3 | 7 | 299 | 5.97 | | | 1 | | | | | | 1 | | | |
| High | Q96D71 | RalBP1-associated Eps domain-containing protein 1 OS=Homo sapiens OX=9606 GN=REPS1 PE=1 SV=3 | 3 | 1 | 1 | 86.6 | 1 | 796 | 5.69 | | | | | | | | | | | | |
| High | Q96S66 | Chloride channel CLIC-like protein 1 OS=Homo sapiens OX=9606 GN=CLCC1 PE=1 SV=1 | 3 | 1 | 1 | 62 | 5 | 551 | 5.55 | | | | | 1 | | | | | | 1 | |
| High | P53701 | Holocytochrome c-type synthase OS=Homo sapiens OX=9606 GN=HCCS PE=1 SV=1 | 3 | 3 | 3 | 30.6 | 16 | 268 | 6.68 | | | | | | | | | | | | |
| High | Q8N9N2 | Activating signal cointegrator 1 complex subunit 1 OS=Homo sapiens OX=9606 GN=ASCC1 PE=1 SV=1 | 3 | 1 | 1 | 45.5 | 2 | 400 | 5.54 | | | | 1 | 1 | 1 | | | | 1 | 1 | 1 |
| High | Q5F1R6 | DnaJ homolog subfamily C member 21 OS=Homo sapiens OX=9606 GN=DNAJC21 PE=1 SV=2 | 3 | 1 | 1 | 62 | 3 | 531 | 5.47 | | | | | | | | | | | | |
| High | Q9GZM7 | Tubulointerstitial nephritis antigen-like OS=Homo sapiens OX=9606 GN=TINAGL1 PE=1 SV=1 | 3 | 1 | 1 | 52.4 | 2 | 467 | 6.99 | | | | | | | | | | | | |
| High | P49848 | Transcription initiation factor TFIID subunit 6 OS=Homo sapiens OX=9606 GN=TAF6 PE=1 SV=1 | 3 | 2 | 2 | 72.6 | 3 | 677 | 8.6 | | | | | | 1 | | | | | | 1 |
| High | Q96C36 | Pyrroline-5-carboxylate reductase 2 OS=Homo sapiens OX=9606 GN=PYCR2 PE=1 SV=1 | 3 | 1 | 1 | 33.6 | 2 | 320 | 7.77 | | | | | | | | | | | | |
| High | O94988 | Protein FAM13A OS=Homo sapiens OX=9606 GN=FAM13A PE=1 SV=2 | 3 | 1 | 1 | 116.9 | 2 | 1023 | 6.48 | | | 1 | | | | | | 1 | | | |
| High | P09914 | Interferon-induced protein with tetratricopeptide repeats 1 OS=Homo sapiens OX=9606 GN=IFIT1 PE=1 SV=2 | 3 | 1 | 1 | 55.3 | 4 | 478 | 7.2 | 1 | | | | 1 | 1 | 1 | | | | 1 | 1 |
| High | Q9P2R6 | Arginine-glutamic acid dipeptide repeats protein OS=Homo sapiens OX=9606 GN=RERE PE=1 SV=2 | 3 | 1 | 1 | 172.3 | 1 | 1566 | 7.69 | | | 1 | | 1 | | | | 1 | | 1 | |
| High | Q13043 | Serine/threonine-protein kinase 4 OS=Homo sapiens OX=9606 GN=STK4 PE=1 SV=2 | 3 | 2 | 1 | 55.6 | 5 | 487 | 5.07 | 1 | | | | 1 | | 1 | | | | 1 | |
| High | Q9Y4H2 | Insulin receptor substrate 2 OS=Homo sapiens OX=9606 GN=IRS2 PE=1 SV=2 | 3 | 3 | 3 | 137.2 | 6 | 1338 | 8.65 | | | | | 1 | | | | | | 1 | |
| High | Q7Z333 | Probable helicase senataxin OS=Homo sapiens OX=9606 GN=SETX PE=1 SV=4 | 3 | 2 | 2 | 302.7 | 1 | 2677 | 7.17 | | | | | 1 | | | | | | 1 | |
| High | P63146 | Ubiquitin-conjugating enzyme E2 B OS=Homo sapiens OX=9606 GN=UBE2B PE=1 SV=1 | 3 | 1 | 1 | 17.3 | 11 | 152 | 5.01 | 1 | 1 | 1 | | | | 1 | 1 | 1 | | | |
| High | Q96TC7 | Regulator of microtubule dynamics protein 3 OS=Homo sapiens OX=9606 GN=RMDN3 PE=1 SV=2 | 3 | 1 | 1 | 52.1 | 4 | 470 | 5.1 | 1 | 1 | 1 | | | | 1 | 1 | 1 | | | |
| High | Q6UW68 | Transmembrane protein 205 OS=Homo sapiens OX=9606 GN=TMEM205 PE=1 SV=1 | 3 | 1 | 1 | 21.2 | 9 | 189 | 8.62 | | | 1 | | | | | | 1 | | | |
| High | P13674 | Prolyl 4-hydroxylase subunit alpha-1 OS=Homo sapiens OX=9606 GN=P4HA1 PE=1 SV=2 | 3 | 2 | 2 | 61 | 5 | 534 | 6.01 | | | 1 | | | | | | 1 | | | |
| High | O14562 | Ubiquitin domain-containing protein UBFD1 OS=Homo sapiens OX=9606 GN=UBFD1 PE=1 SV=2 | 3 | 2 | 2 | 33.4 | 7 | 309 | 5.77 | | | | | | | | | | | | |
| High | P05496 | ATP synthase F(0) complex subunit C1, mitochondrial OS=Homo sapiens OX=9606 GN=ATP5MC1 PE=1 SV=2 | 3 | 1 | 1 | 14.3 | 23 | 136 | 9.74 | | | | 1 | | 1 | | | | 1 | | 1 |
| High | Q9BUZ4 | TNF receptor-associated factor 4 OS=Homo sapiens OX=9606 GN=TRAF4 PE=1 SV=1 | 3 | 1 | 1 | 53.5 | 3 | 470 | 8.15 | | | | | | | | | | | | |
| High | P26583 | High mobility group protein B2 OS=Homo sapiens OX=9606 GN=HMGB2 PE=1 SV=2 | 3 | 1 | 1 | 24 | 6 | 209 | 7.81 | 1 | 1 | 1 | | | | 1 | 1 | 1 | | | |
| High | P33240 | Cleavage stimulation factor subunit 2 OS=Homo sapiens OX=9606 GN=CSTF2 PE=1 SV=1 | 3 | 1 | 1 | 60.9 | 2 | 577 | 6.83 | | | | | 1 | 1 | | | | | 1 | 1 |
| High | Q969T9 | WW domain-binding protein 2 OS=Homo sapiens OX=9606 GN=WBP2 PE=1 SV=1 | 3 | 1 | 1 | 28.1 | 4 | 261 | 5.91 | 1 | | 1 | | | | 1 | | 1 | | | |
| High | Q9HBY8 | Serine/threonine-protein kinase Sgk2 OS=Homo sapiens OX=9606 GN=SGK2 PE=1 SV=2 | 3 | 1 | 1 | 41.1 | 2 | 367 | 6.74 | 1 | 1 | | | | | 1 | 1 | | | | |
| High | Q8N3D4 | EH domain-binding protein 1-like protein 1 OS=Homo sapiens OX=9606 GN=EHBP1L1 PE=1 SV=2 | 3 | 1 | 1 | 161.8 | 1 | 1523 | 4.83 | | | 1 | | | | | | 1 | | | |
| High | Q8N163 | Cell cycle and apoptosis regulator protein 2 OS=Homo sapiens OX=9606 GN=CCAR2 PE=1 SV=2 | 3 | 1 | 1 | 102.8 | 1 | 923 | 5.22 | 1 | | | | | | 1 | | | | | |
| High | Q96GX2 | Ataxin-7-like protein 3B OS=Homo sapiens OX=9606 GN=ATXN7L3B PE=1 SV=2 | 3 | 2 | 2 | 10.8 | 31 | 97 | 4.36 | | | 1 | | 1 | | | | 1 | | 1 | |
| High | Q8N5Z5 | BTB/POZ domain-containing protein KCTD17 OS=Homo sapiens OX=9606 GN=KCTD17 PE=1 SV=3 | 3 | 2 | 2 | 35.6 | 8 | 321 | 4.86 | | | | | 2 | 1 | | | | | 2 | 1 |
| High | Q5TCZ1 | SH3 and PX domain-containing protein 2A OS=Homo sapiens OX=9606 GN=SH3PXD2A PE=1 SV=1 | 3 | 1 | 1 | 125.2 | 1 | 1133 | 8.9 | | | | | | | | | | | | |
| High | P51648 | Aldehyde dehydrogenase family 3 member A2 OS=Homo sapiens OX=9606 GN=ALDH3A2 PE=1 SV=1 | 3 | 1 | 1 | 54.8 | 3 | 485 | 7.88 | | | | | 1 | | | | | | 1 | |
| High | O60573 | Eukaryotic translation initiation factor 4E type 2 OS=Homo sapiens OX=9606 GN=EIF4E2 PE=1 SV=1 | 3 | 2 | 2 | 28.3 | 16 | 245 | 8.88 | | | 1 | | | | | | 1 | | | |
| High | Q13405 | 39S ribosomal protein L49, mitochondrial OS=Homo sapiens OX=9606 GN=MRPL49 PE=1 SV=1 | 3 | 2 | 2 | 19.2 | 16 | 166 | 9.45 | | | | | | | | | | | | |
| High | Q8IZQ5 | Selenoprotein H OS=Homo sapiens OX=9606 GN=SELENOH PE=1 SV=2 | 3 | 1 | 1 | 13.4 | 9 | 122 | 9.74 | | | | 1 | | | | | | 1 | | |
| High | Q9UGU0 | Transcription factor 20 OS=Homo sapiens OX=9606 GN=TCF20 PE=1 SV=3 | 3 | 3 | 3 | 211.6 | 2 | 1960 | 9.04 | | 1 | 2 | | | | | 1 | 2 | | | |
| High | Q9UBP0 | Spastin OS=Homo sapiens OX=9606 GN=SPAST PE=1 SV=1 | 3 | 1 | 1 | 67.2 | 3 | 616 | 9.64 | 1 | | 1 | | | | 1 | | 1 | | | |
| High | Q7L2H7 | Eukaryotic translation initiation factor 3 subunit M OS=Homo sapiens OX=9606 GN=EIF3M PE=1 SV=1 | 3 | 2 | 2 | 42.5 | 9 | 374 | 5.63 | 1 | 1 | | | | | 1 | 1 | | | | |
| High | Q9H078 | Caseinolytic peptidase B protein homolog OS=Homo sapiens OX=9606 GN=CLPB PE=1 SV=1 | 3 | 1 | 1 | 78.7 | 3 | 707 | 9.01 | | | | | 1 | | | | | | 1 | |
| High | Q92844 | TRAF family member-associated NF-kappa-B activator OS=Homo sapiens OX=9606 GN=TANK PE=1 SV=2 | 3 | 2 | 2 | 47.8 | 9 | 425 | 5.68 | | | | | | | | | | | | |
| High | P49903 | Selenide, water dikinase 1 OS=Homo sapiens OX=9606 GN=SEPHS1 PE=1 SV=2 | 3 | 1 | 1 | 42.9 | 4 | 392 | 5.97 | | | 1 | | | | | | 1 | | | |
| High | Q9UBT2 | SUMO-activating enzyme subunit 2 OS=Homo sapiens OX=9606 GN=UBA2 PE=1 SV=2 | 3 | 2 | 2 | 71.2 | 3 | 640 | 5.29 | | 1 | 1 | | | | | 1 | 1 | | | |
| High | O95352 | Ubiquitin-like modifier-activating enzyme ATG7 OS=Homo sapiens OX=9606 GN=ATG7 PE=1 SV=1 | 3 | 2 | 2 | 77.9 | 3 | 703 | 6.24 | | | | | | | | | | | | |
| High | Q9BX46 | RNA-binding protein 24 OS=Homo sapiens OX=9606 GN=RBM24 PE=1 SV=1 | 3 | 1 | 1 | 24.8 | 7 | 236 | 8.51 | | | 1 | | | | | | 1 | | | |
| High | O00178 | GTP-binding protein 1 OS=Homo sapiens OX=9606 GN=GTPBP1 PE=1 SV=3 | 3 | 1 | 1 | 72.4 | 2 | 669 | 8.34 | | | 1 | | | | | | 1 | | | |
| High | Q5VT52 | Regulation of nuclear pre-mRNA domain-containing protein 2 OS=Homo sapiens OX=9606 GN=RPRD2 PE=1 SV=1 | 3 | 2 | 2 | 155.9 | 3 | 1461 | 7.42 | | | | | | | | | | | | |
| High | O76062 | Delta(14)-sterol reductase TM7SF2 OS=Homo sapiens OX=9606 GN=TM7SF2 PE=1 SV=3 | 3 | 1 | 1 | 46.4 | 3 | 418 | 8.87 | 1 | 1 | 1 | | | | 1 | 1 | 1 | | | |
| High | Q9UJ83 | 2-hydroxyacyl-CoA lyase 1 OS=Homo sapiens OX=9606 GN=HACL1 PE=1 SV=2 | 3 | 2 | 2 | 63.7 | 6 | 578 | 7.36 | | | | | | | | | | | | |
| High | Q14802 | FXYD domain-containing ion transport regulator 3 OS=Homo sapiens OX=9606 GN=FXYD3 PE=1 SV=1 | 3 | 1 | 1 | 9.3 | 22 | 87 | 7.3 | | | | | | | | | | | | |
| High | Q92692 | Nectin-2 OS=Homo sapiens OX=9606 GN=NECTIN2 PE=1 SV=1 | 3 | 2 | 2 | 57.7 | 3 | 538 | 4.82 | | | | | | | | | | | | |
| High | P46937 | Transcriptional coactivator YAP1 OS=Homo sapiens OX=9606 GN=YAP1 PE=1 SV=2 | 3 | 2 | 2 | 54.4 | 7 | 504 | 5.17 | | | | | 1 | | | | | | 1 | |
| High | Q9H9B1 | Histone-lysine N-methyltransferase EHMT1 OS=Homo sapiens OX=9606 GN=EHMT1 PE=1 SV=4 | 3 | 1 | 1 | 141.4 | 1 | 1298 | 5.76 | | | | | | | | | | | | |
| High | Q9BTE1 | Dynactin subunit 5 OS=Homo sapiens OX=9606 GN=DCTN5 PE=1 SV=1 | 3 | 2 | 2 | 20.1 | 12 | 182 | 8.02 | | | | 1 | | 1 | | | | 1 | | 1 |
| High | Q6ZU15 | Septin-14 OS=Homo sapiens OX=9606 GN=SEPTIN14 PE=1 SV=2 | 3 | 2 | 1 | 50 | 6 | 432 | 6.24 | | | | 1 | | | | | | 1 | | |
| High | Q9Y5S9 | RNA-binding protein 8A OS=Homo sapiens OX=9606 GN=RBM8A PE=1 SV=1 | 3 | 2 | 2 | 19.9 | 16 | 174 | 5.72 | | | 1 | | 1 | | | | 1 | | 1 | |
| High | Q9BRK4 | Leucine zipper putative tumor suppressor 2 OS=Homo sapiens OX=9606 GN=LZTS2 PE=1 SV=2 | 3 | 1 | 1 | 72.7 | 2 | 669 | 6.51 | 1 | 1 | | | | | 1 | 1 | | | | |
| High | P06732 | Creatine kinase M-type OS=Homo sapiens OX=9606 GN=CKM PE=1 SV=2 | 3 | 2 | 2 | 43.1 | 6 | 381 | 7.25 | | | | 2 | | 1 | | | | 2 | | 1 |
| High | O95398 | Rap guanine nucleotide exchange factor 3 OS=Homo sapiens OX=9606 GN=RAPGEF3 PE=1 SV=6 | 3 | 2 | 2 | 103.7 | 3 | 923 | 7.56 | | | | | | | | | | | | |
| High | Q9NXE8 | Pre-mRNA-splicing factor CWC25 homolog OS=Homo sapiens OX=9606 GN=CWC25 PE=1 SV=1 | 3 | 2 | 2 | 49.6 | 5 | 425 | 10.18 | | | | | 2 | 1 | | | | | 2 | 1 |
| High | P60059 | Protein transport protein Sec61 subunit gamma OS=Homo sapiens OX=9606 GN=SEC61G PE=1 SV=1 | 3 | 1 | 1 | 7.7 | 18 | 68 | 9.99 | | | | | | | | | | | | |
| High | Q99538 | Legumain OS=Homo sapiens OX=9606 GN=LGMN PE=1 SV=1 | 3 | 2 | 2 | 49.4 | 7 | 433 | 6.55 | 1 | | 2 | | | | 1 | | 1 | | | |
| High | Q96HI0 | Sentrin-specific protease 5 OS=Homo sapiens OX=9606 GN=SENP5 PE=1 SV=3 | 3 | 3 | 3 | 86.6 | 5 | 755 | 9.17 | | | | | | | | | | | | |
| High | Q5SWX8 | Protein odr-4 homolog OS=Homo sapiens OX=9606 GN=ODR4 PE=1 SV=1 | 3 | 2 | 2 | 51.1 | 6 | 454 | 5.92 | 1 | 1 | | | | | 1 | 1 | | | | |
| High | Q12955 | Ankyrin-3 OS=Homo sapiens OX=9606 GN=ANK3 PE=1 SV=3 | 3 | 1 | 1 | 480.1 | 0 | 4377 | 6.49 | | | | | 1 | | | | | | 1 | |
| High | P30989 | Neurotensin receptor type 1 OS=Homo sapiens OX=9606 GN=NTSR1 PE=1 SV=2 | 3 | 1 | 1 | 46.2 | 4 | 418 | 9.35 | 1 | 1 | | | | | 1 | 1 | | | | |
| High | P09884 | DNA polymerase alpha catalytic subunit OS=Homo sapiens OX=9606 GN=POLA1 PE=1 SV=2 | 3 | 2 | 2 | 165.8 | 2 | 1462 | 5.85 | 1 | 1 | | | | | 1 | 1 | | | | |
| High | P61009 | Signal peptidase complex subunit 3 OS=Homo sapiens OX=9606 GN=SPCS3 PE=1 SV=1 | 3 | 1 | 1 | 20.3 | 5 | 180 | 8.62 | 1 | | | | | | 1 | | | | | |
| High | Q8IUR0 | Trafficking protein particle complex subunit 5 OS=Homo sapiens OX=9606 GN=TRAPPC5 PE=1 SV=1 | 3 | 1 | 1 | 20.8 | 4 | 188 | 9.66 | | | | | | | | | | | | |
| High | O60563 | Cyclin-T1 OS=Homo sapiens OX=9606 GN=CCNT1 PE=1 SV=1 | 3 | 1 | 1 | 80.6 | 2 | 726 | 8.78 | | | | | | | | | | | | |
| High | Q92917 | G-patch domain and KOW motifs-containing protein OS=Homo sapiens OX=9606 GN=GPKOW PE=1 SV=2 | 3 | 2 | 2 | 52.2 | 6 | 476 | 6.15 | | | | | | | | | | | | |
| High | Q8NF37 | Lysophosphatidylcholine acyltransferase 1 OS=Homo sapiens OX=9606 GN=LPCAT1 PE=1 SV=2 | 3 | 2 | 2 | 59.1 | 4 | 534 | 6.02 | | | | | 1 | | | | | | 1 | |
| High | Q9NPF5 | DNA methyltransferase 1-associated protein 1 OS=Homo sapiens OX=9606 GN=DMAP1 PE=1 SV=1 | 3 | 1 | 1 | 53 | 3 | 467 | 9.5 | | | | | 1 | | | | | | 1 | |
| High | Q5EBM0 | UMP-CMP kinase 2, mitochondrial OS=Homo sapiens OX=9606 GN=CMPK2 PE=1 SV=3 | 3 | 1 | 1 | 49.4 | 3 | 449 | 7.01 | | | | | 1 | 1 | | | | | 1 | 1 |
| High | Q9Y450 | HBS1-like protein OS=Homo sapiens OX=9606 GN=HBS1L PE=1 SV=1 | 3 | 2 | 2 | 75.4 | 4 | 684 | 6.61 | | | | | | | | | | | | |
| High | O15037 | Protein KHNYN OS=Homo sapiens OX=9606 GN=KHNYN PE=1 SV=3 | 3 | 1 | 1 | 74.5 | 3 | 678 | 7.02 | 1 | 1 | 1 | | | | 1 | 1 | 1 | | | |
| High | Q15800 | Methylsterol monooxygenase 1 OS=Homo sapiens OX=9606 GN=MSMO1 PE=1 SV=1 | 3 | 1 | 1 | 35.2 | 5 | 293 | 7.23 | 1 | 1 | 1 | | | | 1 | 1 | 1 | | | |

| | | | | | | | | | | | | | | | | | | | | | |
|---|---|---|---|---|---|---|---|---|---|---|---|---|---|---|---|---|---|---|---|---|---|
| High | Q9P0M9 | 39S ribosomal protein L27, mitochondrial OS=Homo sapiens OX=9606 GN=MRPL27 PE=1 SV=1 | 3 | 1 | 1 | 16.1 | 7 | 148 | 10.42 | | | | 1 | | 1 | | | | 1 | | 1 |
| High | O43237 | Cytoplasmic dynein 1 light intermediate chain 2 OS=Homo sapiens OX=9606 GN=DYNC1LI2 PE=1 SV=1 | 3 | 2 | 2 | 54.1 | 4 | 492 | 6.38 | | | | | | | | | | | | |
| High | P51948 | CDK-activating kinase assembly factor MAT1 OS=Homo sapiens OX=9606 GN=MNAT1 PE=1 SV=1 | 3 | 1 | 1 | 35.8 | 5 | 309 | 6.09 | | | 1 | | | | | | 1 | | | |
| High | P54709 | Sodium/potassium-transporting ATPase subunit beta-3 OS=Homo sapiens OX=9606 GN=ATP1B3 PE=1 SV=1 | 3 | 1 | 1 | 31.5 | 5 | 279 | 8.35 | | | | | 1 | | | | | | 1 | |
| High | Q9BVQ7 | Spermatogenesis-associated protein 5-like protein 1 OS=Homo sapiens OX=9606 GN=SPATA5L1 PE=1 SV=2 | 3 | 1 | 1 | 80.7 | 3 | 753 | 8.09 | | 1 | | | | | | 1 | | | | |
| High | Q7L7V1 | Putative pre-mRNA-splicing factor ATP-dependent RNA helicase DHX32 OS=Homo sapiens OX=9606 GN=DHX32 PE=1 SV=1 | 3 | 1 | 1 | 84.4 | 2 | 743 | 4.97 | | | 1 | | | | | | 1 | | | |
| High | O75027 | Iron-sulfur clusters transporter ABCB7, mitochondrial OS=Homo sapiens OX=9606 GN=ABCB7 PE=1 SV=2 | 3 | 1 | 1 | 82.6 | 2 | 752 | 9.33 | 1 | 1 | 1 | | | | 1 | 1 | 1 | | | |
| High | Q6ZN54 | Differentially expressed in FDCP 8 homolog OS=Homo sapiens OX=9606 GN=DEF8 PE=1 SV=2 | 3 | 1 | 1 | 58.7 | 3 | 512 | 6.52 | 1 | 1 | 1 | | | | 1 | 1 | 1 | | | |
| High | O60488 | Long-chain-fatty-acid--CoA ligase 4 OS=Homo sapiens OX=9606 GN=ACSL4 PE=1 SV=2 | 3 | 2 | 1 | 79.1 | 5 | 711 | 8.38 | | | | | | | | | | | | |
| High | Q00653 | Nuclear factor NF-kappa-B p100 subunit OS=Homo sapiens OX=9606 GN=NFKB2 PE=1 SV=4 | 3 | 2 | 2 | 96.7 | 3 | 900 | 6.25 | | 1 | | | 1 | | | 1 | | | 1 | |
| High | P49585 | Choline-phosphate cytidylyltransferase A OS=Homo sapiens OX=9606 GN=PCYT1A PE=1 SV=2 | 3 | 2 | 2 | 41.7 | 5 | 367 | 7.25 | | | | 1 | | 1 | | | | 1 | | 1 |
| High | Q14210 | Lymphocyte antigen 6D OS=Homo sapiens OX=9606 GN=LY6D PE=1 SV=1 | 3 | 1 | 1 | 13.3 | 7 | 128 | 8.21 | | | | 1 | 1 | 1 | | | | 1 | 1 | 1 |
| High | P00568 | Adenylate kinase isoenzyme 1 OS=Homo sapiens OX=9606 GN=AK1 PE=1 SV=3 | 3 | 1 | 1 | 21.6 | 7 | 194 | 8.63 | | | 1 | | 1 | | | | 1 | | 1 | |
| High | Q53HC5 | Kelch-like protein 26 OS=Homo sapiens OX=9606 GN=KLHL26 PE=1 SV=2 | 3 | 1 | 1 | 68.1 | 1 | 615 | 6.48 | | | | 1 | 1 | 1 | | | | 1 | 1 | 1 |
| High | Q9Y673 | Dolichyl-phosphate beta-glucosyltransferase OS=Homo sapiens OX=9606 GN=ALG5 PE=1 SV=1 | 3 | 2 | 2 | 36.9 | 10 | 324 | 9.28 | | | 1 | | | | | | 1 | | | |
| High | Q8N9N8 | Probable RNA-binding protein EIF1AD OS=Homo sapiens OX=9606 GN=EIF1AD PE=1 SV=1 | 3 | 1 | 1 | 19 | 10 | 165 | 5.21 | | | | | 1 | | | | | | 1 | |
| High | Q8NHQ9 | ATP-dependent RNA helicase DDX55 OS=Homo sapiens OX=9606 GN=DDX55 PE=1 SV=3 | 3 | 2 | 2 | 68.5 | 4 | 600 | 9.25 | | | | | 1 | | | | | | 1 | |
| High | Q14232 | Translation initiation factor eIF-2B subunit alpha OS=Homo sapiens OX=9606 GN=EIF2B1 PE=1 SV=1 | 3 | 1 | 1 | 33.7 | 3 | 305 | 7.33 | | | 1 | | | | | | 1 | | | |
| High | P78314 | SH3 domain-binding protein 2 OS=Homo sapiens OX=9606 GN=SH3BP2 PE=1 SV=2 | 3 | 2 | 2 | 62.2 | 4 | 561 | 7.78 | | 1 | | | | | | 1 | | | | |
| High | Q9HD20 | Endoplasmic reticulum transmembrane helix translocase OS=Homo sapiens OX=9606 GN=ATP13A1 PE=1 SV=2 | 3 | 1 | 1 | 132.9 | 1 | 1204 | 8.13 | | | 1 | | | | | | 1 | | | |
| High | Q92599 | Septin-8 OS=Homo sapiens OX=9606 GN=SEPTIN8 PE=1 SV=4 | 3 | 2 | 1 | 55.7 | 5 | 483 | 6.28 | | 1 | | 1 | | | | 1 | | 1 | | |
| High | Q96AQ6 | Pre-B-cell leukemia transcription factor-interacting protein 1 OS=Homo sapiens OX=9606 GN=PBXIP1 PE=1 SV=1 | 3 | 2 | 2 | 80.6 | 5 | 731 | 5.33 | | | | | | | | | | | | |
| High | Q8IZ69 | tRNA (uracil-5-)-methyltransferase homolog A OS=Homo sapiens OX=9606 GN=TRMT2A PE=1 SV=2 | 3 | 2 | 2 | 68.7 | 6 | 625 | 7.94 | 1 | 1 | | | | | 1 | 1 | | | | |
| High | Q8IY81 | pre-rRNA 2'-O-ribose RNA methyltransferase FTSJ3 OS=Homo sapiens OX=9606 GN=FTSJ3 PE=1 SV=2 | 3 | 2 | 2 | 96.5 | 4 | 847 | 8.4 | | | | 1 | | | | | | 1 | | |
| High | Q7Z4H7 | HAUS augmin-like complex subunit 6 OS=Homo sapiens OX=9606 GN=HAUS6 PE=1 SV=2 | 3 | 1 | 1 | 108.6 | 2 | 955 | 6.47 | | 1 | 1 | | | | | 1 | 1 | | | |
| High | Q9Y2H6 | Fibronectin type-III domain-containing protein 3A OS=Homo sapiens OX=9606 GN=FNDC3A PE=1 SV=4 | 3 | 2 | 2 | 131.8 | 3 | 1198 | 6.71 | | | | | | | | | | | | |
| High | P55957 | BH3-interacting domain death agonist OS=Homo sapiens OX=9606 GN=BID PE=1 SV=1 | 3 | 1 | 1 | 22 | 8 | 195 | 5.44 | 1 | 1 | 1 | | | | 1 | 1 | 1 | | | |
| High | Q13227 | G protein pathway suppressor 2 OS=Homo sapiens OX=9606 GN=GPS2 PE=1 SV=3 | 3 | 1 | 1 | 36.7 | 2 | 327 | 9.52 | | | | | | | | | | | | |
| High | Q9NY59 | Sphingomyelin phosphodiesterase 3 OS=Homo sapiens OX=9606 GN=SMPD3 PE=1 SV=1 | 3 | 1 | 1 | 71 | 2 | 655 | 5.78 | | | | | | | | | | | | |
| High | Q03169 | Tumor necrosis factor alpha-induced protein 2 OS=Homo sapiens OX=9606 GN=TNFAIP2 PE=1 SV=2 | 3 | 2 | 2 | 72.6 | 4 | 654 | 6.46 | | 1 | 2 | | | | | 1 | 2 | | | |
| High | O75306 | NADH dehydrogenase [ubiquinone] iron-sulfur protein 2, mitochondrial OS=Homo sapiens OX=9606 GN=NDUFS2 PE=1 SV=2 | 3 | 2 | 2 | 52.5 | 3 | 463 | 7.55 | 1 | | 1 | | | | 1 | | 1 | | | |
| High | Q6DCA0 | AMMECR1-like protein OS=Homo sapiens OX=9606 GN=AMMECR1L PE=1 SV=1 | 3 | 1 | 1 | 34.5 | 2 | 310 | 8.98 | | | | 1 | 1 | 1 | | | | 1 | 1 | 1 |
| High | P15497 | SWISS-PROT:P15497 (Bos taurus) Apolipoprotein A-I precursor | 3 | 1 | 1 | 30.3 | 4 | 265 | 5.97 | | | | 1 | 1 | 1 | | | | 1 | 1 | 1 |
| High | O00566 | U3 small nucleolar ribonucleoprotein protein MPP10 OS=Homo sapiens OX=9606 GN=MPHOSPH10 PE=1 SV=2 | 3 | 1 | 1 | 78.8 | 1 | 681 | 4.86 | | | | | | | | | | | | |
| High | Q9BXS5 | AP-1 complex subunit mu-1 OS=Homo sapiens OX=9606 GN=AP1M1 PE=1 SV=3 | 3 | 1 | 1 | 48.6 | 4 | 423 | 7.3 | | | 1 | | | | | | 1 | | | |
| High | Q8NCW5 | NAD(P)H-hydrate epimerase OS=Homo sapiens OX=9606 GN=NAXE PE=1 SV=2 | 3 | 2 | 2 | 31.7 | 10 | 288 | 7.66 | | | | | | | | | | | | |
| High | O43402 | ER membrane protein complex subunit 8 OS=Homo sapiens OX=9606 GN=EMC8 PE=1 SV=1 | 3 | 1 | 1 | 23.8 | 4 | 210 | 6.4 | | | | | | | | | | | | |
| High | Q9C0C7 | Activating molecule in BECN1-regulated autophagy protein 1 OS=Homo sapiens OX=9606 GN=AMBRA1 PE=1 SV=2 | 3 | 2 | 2 | 142.4 | 3 | 1298 | 7.18 | | | | 1 | 1 | | | | | 1 | 1 | |
| High | Q5UIP0 | Telomere-associated protein RIF1 OS=Homo sapiens OX=9606 GN=RIF1 PE=1 SV=2 | 3 | 2 | 2 | 274.3 | 1 | 2472 | 5.52 | | | | | | | | | | | | |
| High | O00244 | Copper transport protein ATOX1 OS=Homo sapiens OX=9606 GN=ATOX1 PE=1 SV=1 | 3 | 1 | 1 | 7.4 | 12 | 68 | 7.24 | | | 1 | | 1 | | | | 1 | | 1 | |
| High | Q9UBB4 | Ataxin-10 OS=Homo sapiens OX=9606 GN=ATXN10 PE=1 SV=1 | 3 | 1 | 1 | 53.5 | 2 | 475 | 5.25 | | 1 | 1 | | | 1 | | 1 | 1 | | | 1 |
| High | Q15004 | PCNA-associated factor OS=Homo sapiens OX=9606 GN=PCLAF PE=1 SV=1 | 3 | 2 | 2 | 12 | 28 | 111 | 9.83 | | | | 1 | 1 | 1 | | | | 1 | 1 | 1 |
| High | P61803 | Dolichyl-diphosphooligosaccharide--protein glycosyltransferase subunit DAD1 OS=Homo sapiens OX=9606 GN=DAD1 PE=1 SV=3 | 3 | 1 | 1 | 12.5 | 11 | 113 | 7.08 | 1 | 1 | 1 | | | | 1 | 1 | 1 | | | |
| High | P00519 | Tyrosine-protein kinase ABL1 OS=Homo sapiens OX=9606 GN=ABL1 PE=1 SV=4 | 3 | 1 | 1 | 122.8 | 1 | 1130 | 8.66 | | | 1 | | | | | | 1 | | | |
| High | Q92504 | Zinc transporter SLC39A7 OS=Homo sapiens OX=9606 GN=SLC39A7 PE=1 SV=2 | 3 | 2 | 2 | 50.1 | 3 | 469 | 6.87 | | 1 | | | | | | 1 | | | | |
| High | Q03426 | Mevalonate kinase OS=Homo sapiens OX=9606 GN=MVK PE=1 SV=1 | 3 | 2 | 2 | 42.4 | 12 | 396 | 6.46 | 1 | | 1 | | | | 1 | | 1 | | | |
| High | P28065 | Proteasome subunit beta type-9 OS=Homo sapiens OX=9606 GN=PSMB9 PE=1 SV=2 | 3 | 1 | 1 | 23.3 | 5 | 219 | 5.03 | | | | 1 | 1 | 1 | | | | 1 | 1 | 1 |
| High | O15120 | 1-acyl-sn-glycerol-3-phosphate acyltransferase beta OS=Homo sapiens OX=9606 GN=AGPAT2 PE=1 SV=1 | 3 | 1 | 1 | 30.9 | 3 | 278 | 9.01 | 1 | 1 | | | | | 1 | 1 | | | | |
| High | Q04864 | Proto-oncogene c-Rel OS=Homo sapiens OX=9606 GN=REL PE=1 SV=1 | 3 | 1 | 1 | 68.5 | 2 | 619 | 5.86 | | | | | | | | | | | | |
| High | Q68D10 | Protein SPT2 homolog OS=Homo sapiens OX=9606 GN=SPTY2D1 PE=1 SV=3 | 3 | 2 | 2 | 75.6 | 3 | 685 | 9.79 | 1 | | | | | | 1 | | | | | |
| High | P49773 | Adenosine 5'-monophosphoramidase HINT1 OS=Homo sapiens OX=9606 GN=HINT1 PE=1 SV=2 | 3 | 1 | 1 | 13.8 | 7 | 126 | 6.95 | 1 | | | | 1 | | 1 | | | | 1 | |
| High | Q96GQ5 | RUS family member 1 OS=Homo sapiens OX=9606 GN=RUSF1 PE=1 SV=2 | 3 | 2 | 2 | 51 | 8 | 468 | 6.93 | 1 | 1 | 1 | | | | 1 | 1 | 1 | | | |
| High | Q9UIV1 | CCR4-NOT transcription complex subunit 7 OS=Homo sapiens OX=9606 GN=CNOT7 PE=1 SV=3 | 3 | 2 | 2 | 32.7 | 9 | 285 | 4.84 | | | 1 | | | | | | 1 | | | |
| High | Q8NDH3 | Probable aminopeptidase NPEPL1 OS=Homo sapiens OX=9606 GN=NPEPL1 PE=1 SV=3 | 3 | 2 | 2 | 55.8 | 5 | 523 | 6.87 | | 1 | 1 | | | | | 1 | 1 | | | |
| High | P13716 | Delta-aminolevulinic acid dehydratase OS=Homo sapiens OX=9606 GN=ALAD PE=1 SV=1 | 3 | 1 | 1 | 36.3 | 3 | 330 | 6.79 | 1 | | | | 1 | | 1 | | | | 1 | |
| High | P55854 | Small ubiquitin-related modifier 3 OS=Homo sapiens OX=9606 GN=SUMO3 PE=1 SV=2 | 3 | 1 | 1 | 11.6 | 12 | 103 | 5.49 | | | | | | | | | | | | |
| High | Q86Y37 | CDK2-associated and cullin domain-containing protein 1 OS=Homo sapiens OX=9606 GN=CACUL1 PE=1 SV=1 | 3 | 2 | 2 | 41 | 9 | 369 | 5.19 | 1 | 1 | | | | | 1 | 1 | | | | |
| High | Q9BV38 | WD repeat-containing protein 18 OS=Homo sapiens OX=9606 GN=WDR18 PE=1 SV=2 | 3 | 2 | 2 | 47.4 | 6 | 432 | 6.7 | 1 | | 1 | 1 | | | 1 | | 1 | 1 | | |
| High | Q9UIA9 | Exportin-7 OS=Homo sapiens OX=9606 GN=XPO7 PE=1 SV=3 | 3 | 3 | 3 | 123.8 | 4 | 1087 | 6.32 | | 1 | 1 | | | | | 1 | 1 | | | |
| High | Q9NQ84 | G-protein coupled receptor family C group 5 member C OS=Homo sapiens OX=9606 GN=GPRC5C PE=1 SV=2 | 3 | 2 | 2 | 48.2 | 5 | 441 | 8.43 | | | | | | | | | | | | |
| High | Q9BZ29 | Dedicator of cytokinesis protein 9 OS=Homo sapiens OX=9606 GN=DOCK9 PE=1 SV=2 | 3 | 2 | 2 | 236.3 | 1 | 2069 | 7.49 | 1 | 1 | | | | | 1 | 1 | | | | |
| High | P07741 | Adenine phosphoribosyltransferase OS=Homo sapiens OX=9606 GN=APRT PE=1 SV=2 | 3 | 2 | 2 | 19.6 | 10 | 180 | 6.02 | 1 | | 1 | | | | 1 | | 1 | | | |
| High | Q9Y6I9 | Testis-expressed protein 264 OS=Homo sapiens OX=9606 GN=TEX264 PE=1 SV=1 | 3 | 1 | 1 | 34.2 | 4 | 313 | 4.86 | | | | | | | | | | | | |
| High | Q14534 | Squalene monooxygenase OS=Homo sapiens OX=9606 GN=SQLE PE=1 SV=3 | 3 | 2 | 2 | 63.9 | 5 | 574 | 8.63 | | | 2 | | | | | | 2 | | | |
| High | Q9NYI0 | PH and SEC7 domain-containing protein 3 OS=Homo sapiens OX=9606 GN=PSD3 PE=1 SV=2 | 3 | 1 | 1 | 116 | 2 | 1048 | 5.99 | | | 1 | | | | | | 1 | | | |
| High | Q9NZT1 | Calmodulin-like protein 5 OS=Homo sapiens OX=9606 GN=CALML5 PE=1 SV=2 | 3 | 1 | 1 | 15.9 | 5 | 146 | 4.44 | | | | 1 | 1 | 1 | | | | 1 | 1 | 1 |
| High | Q05469 | Hormone-sensitive lipase OS=Homo sapiens OX=9606 GN=LIPE PE=1 SV=4 | 3 | 2 | 2 | 116.5 | 1 | 1076 | 6.7 | | | | | | | | | | | | |
| High | Q9Y262 | Eukaryotic translation initiation factor 3 subunit L OS=Homo sapiens OX=9606 GN=EIF3L PE=1 SV=1 | 3 | 3 | 3 | 66.7 | 6 | 564 | 6.34 | | | | | 1 | | | | | | 1 | |
| High | Q96P48 | Arf-GAP with Rho-GAP domain, ANK repeat and PH domain-containing protein 1 OS=Homo sapiens OX=9606 GN=ARAP1 PE=1 SV= | 3 | 2 | 2 | 162.1 | 2 | 1450 | 6.23 | | | | | | | | | | | | |
| High | Q9UKK9 | ADP-sugar pyrophosphatase OS=Homo sapiens OX=9606 GN=NUDT5 PE=1 SV=1 | 3 | 1 | 1 | 24.3 | 4 | 219 | 4.94 | | | 1 | | 1 | | | | 1 | | 1 | |
| High | P53384 | Cytosolic Fe-S cluster assembly factor NUBP1 OS=Homo sapiens OX=9606 GN=NUBP1 PE=1 SV=2 | 3 | 1 | 1 | 34.5 | 6 | 320 | 5.33 | | | | | | | | | | | | |
| High | Q9H7B2 | Ribosome production factor 2 homolog OS=Homo sapiens OX=9606 GN=RPF2 PE=1 SV=2 | 3 | 1 | 1 | 35.6 | 6 | 306 | 9.99 | 1 | | 1 | | | | 1 | | 1 | | | |
| High | Q9UL33 | Trafficking protein particle complex subunit 2-like protein OS=Homo sapiens OX=9606 GN=TRAPPC2L PE=1 SV=1 | 3 | 1 | 1 | 16.1 | 14 | 140 | 6.77 | | | 1 | | | | | | 1 | | | |
| High | Q9BUN8 | Derlin-1 OS=Homo sapiens OX=9606 GN=DERL1 PE=1 SV=1 | 3 | 1 | 1 | 28.8 | 4 | 251 | 9.51 | 1 | 1 | 1 | | | | 1 | 1 | 1 | | | |
| High | Q15032 | R3H domain-containing protein 1 OS=Homo sapiens OX=9606 GN=R3HDM1 PE=1 SV=3 | 3 | 2 | 2 | 120.6 | 3 | 1099 | 8.75 | | 1 | 2 | | | | | 1 | 2 | | | |
| High | Q16540 | 39S ribosomal protein L23, mitochondrial OS=Homo sapiens OX=9606 GN=MRPL23 PE=1 SV=1 | 3 | 1 | 1 | 17.8 | 10 | 153 | 9.69 | | | | 1 | | | | | | 1 | | |
| High | O14497 | AT-rich interactive domain-containing protein 1A OS=Homo sapiens OX=9606 GN=ARID1A PE=1 SV=3 | 3 | 2 | 2 | 241.9 | 2 | 2285 | 6.7 | | | | | | | | | | | | |
| High | Q8N357 | Solute carrier family 35 member F6 OS=Homo sapiens OX=9606 GN=SLC35F6 PE=1 SV=1 | 3 | 1 | 1 | 40.2 | 3 | 371 | 6.93 | 1 | 1 | 1 | | | | 1 | 1 | 1 | | | |
| High | O43292 | Glycosylphosphatidylinositol anchor attachment 1 protein OS=Homo sapiens OX=9606 GN=GPAA1 PE=1 SV=3 | 3 | 1 | 1 | 67.6 | 1 | 621 | 8.06 | 1 | 1 | 1 | | | | 1 | 1 | 1 | | | |
| High | Q8IWI9 | MAX gene-associated protein OS=Homo sapiens OX=9606 GN=MGA PE=1 SV=4 | 3 | 1 | 1 | 336 | 0 | 3065 | 6.9 | | | | | | | | | | | | |
| High | Q9H974 | Queuine tRNA-ribosyltransferase accessory subunit 2 OS=Homo sapiens OX=9606 GN=QTRT2 PE=1 SV=1 | 3 | 1 | 1 | 46.7 | 5 | 415 | 6.81 | 1 | 1 | 1 | | | | 1 | 1 | 1 | | | |
| High | Q9UPP1 | Histone lysine demethylase PHF8 OS=Homo sapiens OX=9606 GN=PHF8 PE=1 SV=3 | 3 | 3 | 3 | 117.8 | 4 | 1060 | 8.72 | | | | | | | | | | | | |
| High | Q96JN0 | Ligand-dependent corepressor OS=Homo sapiens OX=9606 GN=LCOR PE=1 SV=2 | 3 | 1 | 1 | 47 | 3 | 433 | 9.38 | 1 | 1 | 1 | | | | 1 | 1 | 1 | | | |
| High | O43709 | Probable 18S rRNA (guanine-N(7))-methyltransferase OS=Homo sapiens OX=9606 GN=BUD23 PE=1 SV=2 | 3 | 2 | 2 | 31.9 | 7 | 281 | 8.73 | | | | | 1 | 1 | | | | | 1 | 1 |
| High | O14925 | Mitochondrial import inner membrane translocase subunit Tim23 OS=Homo sapiens OX=9606 GN=TIMM23 PE=1 SV=1 | 3 | 1 | 1 | 21.9 | 5 | 209 | 8.6 | 1 | 1 | 1 | | | | 1 | 1 | 1 | | | |
| High | Q8N9E0 | Protein FAM133A OS=Homo sapiens OX=9606 GN=FAM133A PE=1 SV=1 | 3 | 1 | 1 | 28.9 | 3 | 248 | 10.1 | | | | 1 | 1 | 1 | | | | 1 | 1 | 1 |
| High | Q96M96 | FYVE, RhoGEF and PH domain-containing protein 4 OS=Homo sapiens OX=9606 GN=FGD4 PE=1 SV=2 | 3 | 1 | 1 | 86.6 | 1 | 766 | 6.13 | | | | | | | | | | | | |
| High | Q9NRR4 | Ribonuclease 3 OS=Homo sapiens OX=9606 GN=DROSHA PE=1 SV=2 | 3 | 1 | 1 | 159.2 | 1 | 1374 | 7.87 | | 1 | 1 | | | | | 1 | 1 | | | |
| High | P62745 | Rho-related GTP-binding protein RhoB OS=Homo sapiens OX=9606 GN=RHOB PE=1 SV=1 | 3 | 2 | 1 | 22.1 | 17 | 196 | 5.24 | 1 | 1 | 1 | | | | 1 | 1 | 1 | | | |
| High | P82909 | Alpha-ketoglutarate dehydrogenase component 4 OS=Homo sapiens OX=9606 GN=MRPS36 PE=1 SV=2 | 3 | 2 | 2 | 11.5 | 27 | 103 | 9.99 | 1 | 1 | | | | | 1 | 1 | | | | |
| High | Q9NPR2 | Semaphorin-4B OS=Homo sapiens OX=9606 GN=SEMA4B PE=1 SV=4 | 3 | 2 | 2 | 92.7 | 3 | 837 | 7.05 | 1 | 1 | | | | | 1 | 1 | | | | |
| High | Q66K14 | TBC1 domain family member 9B OS=Homo sapiens OX=9606 GN=TBC1D9B PE=1 SV=3 | 3 | 1 | 1 | 140.4 | 2 | 1250 | 5.25 | | | 1 | | | | | | 1 | | | |
| High | Q15054 | DNA polymerase delta subunit 3 OS=Homo sapiens OX=9606 GN=POLD3 PE=1 SV=2 | 3 | 2 | 2 | 51.4 | 5 | 466 | 9.35 | | | 1 | | | | | | 1 | | | |
| High | O60504 | Vinexin OS=Homo sapiens OX=9606 GN=SORBS3 PE=1 SV=2 | 3 | 1 | 1 | 75.3 | 2 | 671 | 9.45 | | | | | | | | | | | | |
| High | Q8NEZ4 | Histone-lysine N-methyltransferase 2C OS=Homo sapiens OX=9606 GN=KMT2C PE=1 SV=3 | 3 | 2 | 2 | 541 | 0 | 4911 | 6.49 | | | | 1 | 1 | 1 | | | | 1 | 1 | 1 |
| High | Q9NR09 | Baculoviral IAP repeat-containing protein 6 OS=Homo sapiens OX=9606 GN=BIRC6 PE=1 SV=3 | 3 | 2 | 2 | 529.9 | 1 | 4857 | 6.05 | | | | | | | | | | | | |
| High | O95154 | Aflatoxin B1 aldehyde reductase member 3 OS=Homo sapiens OX=9606 GN=AKR7A3 PE=1 SV=2 | 3 | 3 | 2 | 37.2 | 13 | 331 | 7.15 | | 1 | | | | | | 1 | | | | |
| High | Q00403 | Transcription initiation factor IIB OS=Homo sapiens OX=9606 GN=GTF2B PE=1 SV=1 | 3 | 1 | 1 | 34.8 | 5 | 316 | 8.35 | | 1 | | | 1 | | | 1 | | | 1 | |
| High | Q9C0D5 | Protein TANC1 OS=Homo sapiens OX=9606 GN=TANC1 PE=1 SV=3 | 3 | 2 | 2 | 202.1 | 1 | 1861 | 8.32 | | | | | | | | | | | | |
| High | Q4G0X4 | BTB/POZ domain-containing protein KCTD21 OS=Homo sapiens OX=9606 GN=KCTD21 PE=1 SV=1 | 3 | 1 | 1 | 29.6 | 7 | 260 | 6.54 | 1 | 1 | 1 | | | | 1 | 1 | 1 | | | |
| High | Q969S9 | Ribosome-releasing factor 2, mitochondrial OS=Homo sapiens OX=9606 GN=GFM2 PE=1 SV=1 | 3 | 2 | 2 | 86.5 | 2 | 779 | 6.51 | 1 | 1 | | | | | 1 | 1 | | | | |
| High | Q8IXM3 | 39S ribosomal protein L41, mitochondrial OS=Homo sapiens OX=9606 GN=MRPL41 PE=1 SV=1 | 3 | 2 | 2 | 15.4 | 18 | 137 | 9.57 | | | | | | 1 | | | | | | 1 |
| High | Q9H300 | Presenilins-associated rhomboid-like protein, mitochondrial OS=Homo sapiens OX=9606 GN=PARL PE=1 SV=2 | 3 | 1 | 1 | 42.2 | 2 | 379 | 9.79 | 1 | 1 | 1 | | | | 1 | 1 | 1 | | | |
| High | P02663 | SWISS-PROT:P02663 Alpha-S2-casein [Contains: Casocidin-1 - Bos taurus (Bovine). | 3 | 2 | 2 | 24.3 | 9 | 207 | 8.27 | | | | | | | | | | | | |
| High | P49840 | Glycogen synthase kinase-3 alpha OS=Homo sapiens OX=9606 GN=GSK3A PE=1 SV=2 | 3 | 1 | 1 | 50.9 | 3 | 483 | 8.75 | 1 | 1 | 1 | | | | 1 | 1 | 1 | | | |
| High | Q6ZU65 | Ubinuclein-2 OS=Homo sapiens OX=9606 GN=UBN2 PE=1 SV=2 | 3 | 2 | 2 | 146 | 3 | 1347 | 9.19 | | | 1 | | | | | | 1 | | | |
| High | Q7Z4V5 | Hepatoma-derived growth factor-related protein 2 OS=Homo sapiens OX=9606 GN=HDGFL2 PE=1 SV=1 | 3 | 2 | 2 | 74.3 | 4 | 671 | 7.49 | 1 | | | | | | 1 | | | | | |
| High | Q5T750 | Skin-specific protein 32 OS=Homo sapiens OX=9606 GN=XP32 PE=1 SV=1 | 3 | 1 | 1 | 26.2 | 3 | 250 | 7.97 | | | 1 | | 1 | | | | 1 | | 1 | |
| High | P28062 | Proteasome subunit beta type-8 OS=Homo sapiens OX=9606 GN=PSMB8 PE=1 SV=3 | 3 | 2 | 2 | 30.3 | 9 | 276 | 7.43 | | | | 1 | 1 | 1 | | | | 1 | 1 | 1 |
| High | O43159 | Ribosomal RNA-processing protein 8 OS=Homo sapiens OX=9606 GN=RRP8 PE=1 SV=2 | 3 | 2 | 2 | 50.7 | 6 | 456 | 9.42 | | | | | | | | | | | | |
| High | O95619 | YEATS domain-containing protein 4 OS=Homo sapiens OX=9606 GN=YEATS4 PE=1 SV=1 | 3 | 1 | 1 | 26.5 | 4 | 227 | 8.41 | 1 | 1 | 1 | | | | 1 | 1 | 1 | | | |
| High | O60936 | Nucleolar protein 3 OS=Homo sapiens OX=9606 GN=NOL3 PE=1 SV=2 | 3 | 1 | 1 | 22.6 | 9 | 208 | 4.18 | | | | | | | | | | | | |
| High | Q13610 | Periodic tryptophan protein 1 homolog OS=Homo sapiens OX=9606 GN=PWP1 PE=1 SV=1 | 3 | 2 | 2 | 55.8 | 5 | 501 | 4.77 | | | | | | | | | | | | |
| High | P46100 | Transcriptional regulator ATRX OS=Homo sapiens OX=9606 GN=ATRX PE=1 SV=6 | 3 | 2 | 2 | 282.4 | 1 | 2492 | 6.55 | | | 2 | | | | | | 2 | | | |
| High | Q13572 | Inositol-tetrakisphosphate 1-kinase OS=Homo sapiens OX=9606 GN=ITPK1 PE=1 SV=2 | 3 | 2 | 2 | 45.6 | 7 | 414 | 6.16 | 1 | 1 | | | | | 1 | 1 | | | | |
| High | Q9NXC5 | GATOR complex protein MIOS OS=Homo sapiens OX=9606 GN=MIOS PE=1 SV=2 | 3 | 1 | 1 | 98.5 | 1 | 875 | 6.73 | | | 1 | | | | | | 1 | | | |
| High | P00846 | ATP synthase subunit a OS=Homo sapiens OX=9606 GN=MT-ATP6 PE=1 SV=1 | 3 | 1 | 1 | 24.8 | 4 | 226 | 10.1 | 1 | 1 | 1 | | | | 1 | 1 | 1 | | | |
| High | Q9NVM9 | Integrator complex subunit 13 OS=Homo sapiens OX=9606 GN=INTS13 PE=1 SV=2 | 3 | 2 | 2 | 80.2 | 3 | 706 | 6.7 | | | | | 1 | | | | | | 1 | |
| High | Q13459 | Unconventional myosin-IXb OS=Homo sapiens OX=9606 GN=MYO9B PE=1 SV=3 | 3 | 2 | 2 | 243.2 | 1 | 2157 | 8.75 | | 1 | 1 | | | | | 1 | 1 | | | |
| High | O60870 | DNA/RNA-binding protein KIN17 OS=Homo sapiens OX=9606 GN=KIN PE=1 SV=2 | 3 | 1 | 1 | 45.3 | 2 | 393 | 8.95 | | | 1 | | 1 | 1 | | | 1 | | 1 | 1 |
| High | Q969E8 | Pre-rRNA-processing protein TSR2 homolog OS=Homo sapiens OX=9606 GN=TSR2 PE=1 SV=1 | 3 | 1 | 1 | 20.9 | 5 | 191 | 4.39 | | | | 1 | | | | | | 1 | | |
| High | P27707 | Deoxycytidine kinase OS=Homo sapiens OX=9606 GN=DCK PE=1 SV=1 | 3 | 2 | 2 | 30.5 | 7 | 260 | 5.21 | | | | 1 | 1 | | | | | 1 | 1 | |
| High | Q13564 | NEDD8-activating enzyme E1 regulatory subunit OS=Homo sapiens OX=9606 GN=NAE1 PE=1 SV=1 | 3 | 3 | 3 | 60.2 | 9 | 534 | 5.4 | | | | 1 | | | | | | 1 | | |
| High | Q15649 | Zinc finger HIT domain-containing protein 3 OS=Homo sapiens OX=9606 GN=ZNHIT3 PE=1 SV=2 | 3 | 2 | 2 | 17.6 | 12 | 155 | 5.67 | | | | | | 1 | | | | | | 1 |
| High | Q96JY6 | PDZ and LIM domain protein 2 OS=Homo sapiens OX=9606 GN=PDLIM2 PE=1 SV=1 | 3 | 1 | 1 | 37.4 | 6 | 352 | 8.72 | | | | | | | | | | | | |
| High | Q13868 | Exosome complex component RRP4 OS=Homo sapiens OX=9606 GN=EXOSC2 PE=1 SV=2 | 3 | 3 | 3 | 32.8 | 12 | 293 | 7.5 | | | | | | | | | | | | |
| High | Q92796 | Disks large homolog 3 OS=Homo sapiens OX=9606 GN=DLG3 PE=1 SV=2 | 3 | 2 | 2 | 90.3 | 4 | 817 | 7.03 | | | | | | | | | | | | |
| High | O94979 | Protein transport protein Sec31A OS=Homo sapiens OX=9606 GN=SEC31A PE=1 SV=3 | 3 | 1 | 1 | 132.9 | 1 | 1220 | 6.89 | | | | | | | | | | | | |
| High | P20585 | DNA mismatch repair protein Msh3 OS=Homo sapiens OX=9606 GN=MSH3 PE=1 SV=4 | 3 | 1 | 1 | 127.3 | 1 | 1137 | 8.02 | | | | 1 | | 1 | | | | 1 | | 1 |
| High | P29218 | Inositol monophosphatase 1 OS=Homo sapiens OX=9606 GN=IMPA1 PE=1 SV=1 | 3 | 2 | 2 | 30.2 | 5 | 277 | 5.26 | | | | | | | | | | | | |
| High | Q8NB37 | Glutamine amidotransferase-like class 1 domain-containing protein 1 OS=Homo sapiens OX=9606 GN=GATD1 PE=1 SV=1 | 3 | 2 | 2 | 23.3 | 14 | 220 | 6.61 | | | | | | | | | | | | |
| High | P20645 | Cation-dependent mannose-6-phosphate receptor OS=Homo sapiens OX=9606 GN=M6PR PE=1 SV=1 | 3 | 1 | 1 | 31 | 5 | 277 | 5.83 | | 1 | 1 | | | | | 1 | 1 | | | |
| High | Q9H2W6 | 39S ribosomal protein L46, mitochondrial OS=Homo sapiens OX=9606 GN=MRPL46 PE=1 SV=1 | 3 | 2 | 2 | 31.7 | 12 | 279 | 7.05 | 1 | 1 | | | | | 1 | 1 | | | | |
| High | P09132 | Signal recognition particle 19 kDa protein OS=Homo sapiens OX=9606 GN=SRP19 PE=1 SV=3 | 3 | 1 | 1 | 16.1 | 10 | 144 | 9.85 | | | 1 | 1 | | 1 | | | 1 | 1 | | 1 |
| High | Q8NDA8 | Maestro heat-like repeat-containing protein family member 1 OS=Homo sapiens OX=9606 GN=MROH1 PE=2 SV=3 | 3 | 2 | 2 | 181.1 | 1 | 1641 | 6.89 | 1 | | 1 | | | | 1 | | 1 | | | |
| High | Q5VYX0 | Renalase OS=Homo sapiens OX=9606 GN=RNLS PE=1 SV=1 | 3 | 2 | 2 | 37.8 | 5 | 342 | 6.52 | | | | | 1 | | | | | | 1 | |
| High | Q2NKX8 | DNA excision repair protein ERCC-6-like OS=Homo sapiens OX=9606 GN=ERCC6L PE=1 SV=1 | 3 | 1 | 1 | 141 | 1 | 1250 | 5.31 | 1 | | | | | | 1 | | | | | |
| High | P55001 | Microfibrillar-associated protein 2 OS=Homo sapiens OX=9606 GN=MFAP2 PE=1 SV=1 | 3 | 1 | 1 | 20.8 | 5 | 183 | 4.97 | | | | 1 | 1 | 1 | | | | 1 | 1 | 1 |
| High | O60879 | Protein diaphanous homolog 2 OS=Homo sapiens OX=9606 GN=DIAPH2 PE=1 SV=1 | 3 | 1 | 1 | 125.5 | 1 | 1101 | 6.58 | | | | | | | | | | | | |
| High | Q8NCA5 | Protein FAM98A OS=Homo sapiens OX=9606 GN=FAM98A PE=1 SV=2 | 3 | 3 | 3 | 55.2 | 10 | 518 | 8.95 | | | 1 | | 1 | | | | 1 | | 1 | |
| High | P51797 | H(+)/Cl(-) exchange transporter 6 OS=Homo sapiens OX=9606 GN=CLCN6 PE=1 SV=2 | 3 | 1 | 1 | 97.2 | 1 | 869 | 6.81 | | | 1 | | | | | | 1 | | | |
| High | P15036 | Protein C-ets-2 OS=Homo sapiens OX=9606 GN=ETS2 PE=1 SV=1 | 3 | 1 | 1 | 53 | 2 | 469 | 5.03 | | | | | | | | | | | | |
| High | P04156 | Major prion protein OS=Homo sapiens OX=9606 GN=PRNP PE=1 SV=1 | 3 | 1 | 1 | 27.6 | 5 | 253 | 9 | | | | | | | | | | | | |
| High | Q5XG87 | Terminal nucleotidyltransferase 4A OS=Homo sapiens OX=9606 GN=TENT4A PE=1 SV=3 | 2 | 1 | 1 | 82.3 | 1 | 772 | 9.52 | | | | 1 | | 1 | | | | 1 | | 1 |
| High | P0DPB6 | DNA-directed RNA polymerases I and III subunit RPAC2 OS=Homo sapiens OX=9606 GN=POLR1D PE=1 SV=1 | 2 | 2 | 2 | 15.2 | 8 | 133 | 5.8 | | | | | 1 | | | | | | 1 | |
| High | Q71RC2 | La-related protein 4 OS=Homo sapiens OX=9606 GN=LARP4 PE=1 SV=3 | 2 | 1 | 1 | 80.5 | 3 | 724 | 6.61 | | | | | | | | | | | | |
| High | Q5T280 | Putative methyltransferase C9orf114 OS=Homo sapiens OX=9606 GN=SPOUT1 PE=1 SV=3 | 2 | 1 | 1 | 42 | 4 | 376 | 7.43 | | | 1 | | 1 | | | | 1 | | 1 | |
| High | P10644 | cAMP-dependent protein kinase type I-alpha regulatory subunit OS=Homo sapiens OX=9606 GN=PRKAR1A PE=1 SV=1 | 2 | 1 | 1 | 43 | 3 | 381 | 5.35 | 1 | 1 | | | | | 1 | 1 | | | | |
| High | Q9P1F3 | Costars family protein ABRACL OS=Homo sapiens OX=9606 GN=ABRACL PE=1 SV=1 | 2 | 1 | 1 | 9.1 | 20 | 81 | 6.29 | 1 | 1 | | | | | 1 | 1 | | | | |
| High | Q9NR28 | Diablo IAP-binding mitochondrial protein OS=Homo sapiens OX=9606 GN=DIABLO PE=1 SV=1 | 2 | 1 | 1 | 27.1 | 4 | 239 | 5.9 | | | | | | | | | | | | |
| High | Q99590 | Protein SCAF11 OS=Homo sapiens OX=9606 GN=SCAF11 PE=1 SV=2 | 2 | 1 | 1 | 164.6 | 1 | 1463 | 8.41 | | | | | | | | | | | | |
| High | Q9NZC9 | SWI/SNF-related matrix-associated actin-dependent regulator of chromatin subfamily A-like protein 1 OS=Homo sapiens OX=9606 | 2 | 2 | 2 | 105.9 | 2 | 954 | 9.06 | 1 | | 1 | | | | 1 | | 1 | | | |
| High | Q7Z2W9 | 39S ribosomal protein L21, mitochondrial OS=Homo sapiens OX=9606 GN=MRPL21 PE=1 SV=2 | 2 | 1 | 1 | 22.8 | 5 | 205 | 9.89 | | | | | | | | | | | | |
| High | Q8WW12 | PEST proteolytic signal-containing nuclear protein OS=Homo sapiens OX=9606 GN=PCNP PE=1 SV=2 | 2 | 1 | 1 | 18.9 | 9 | 178 | 7.49 | | | 1 | | | | | | 1 | | | |
| High | Q9UHW5 | GPN-loop GTPase 3 OS=Homo sapiens OX=9606 GN=GPN3 PE=1 SV=2 | 2 | 1 | 1 | 32.7 | 6 | 284 | 4.5 | | 1 | 1 | | | | | 1 | 1 | | | |
| High | Q8IZV5 | Retinol dehydrogenase 10 OS=Homo sapiens OX=9606 GN=RDH10 PE=1 SV=1 | 2 | 1 | 1 | 38.1 | 5 | 341 | 7.4 | | | | | | | | | | | | |
| High | O15550 | Lysine-specific demethylase 6A OS=Homo sapiens OX=9606 GN=KDM6A PE=1 SV=2 | 2 | 2 | 2 | 154.1 | 2 | 1401 | 7.44 | | | | | 1 | | | | | | 1 | |
| High | Q8N1P7 | Beta/gamma crystallin domain-containing protein 2 OS=Homo sapiens OX=9606 GN=CRYBG2 PE=2 SV=2 | 2 | 1 | 1 | 177.8 | 1 | 1661 | 6.05 | | 1 | | | | | | 1 | | | | |
| High | Q8NF64 | Zinc finger MIZ domain-containing protein 2 OS=Homo sapiens OX=9606 GN=ZMIZ2 PE=1 SV=2 | 2 | 2 | 2 | 96.5 | 2 | 920 | 7.12 | | | | | | | | | | | | |
| High | Q96EL3 | 39S ribosomal protein L53, mitochondrial OS=Homo sapiens OX=9606 GN=MRPL53 PE=1 SV=1 | 2 | 1 | 1 | 12.1 | 7 | 112 | 8.76 | | | | | | | | | | | | |
| High | O95707 | Ribonuclease P protein subunit p29 OS=Homo sapiens OX=9606 GN=POP4 PE=1 SV=2 | 2 | 1 | 1 | 25.4 | 8 | 220 | 10.07 | | | | | | | | | | | | |
| High | Q13347 | Eukaryotic translation initiation factor 3 subunit I OS=Homo sapiens OX=9606 GN=EIF3I PE=1 SV=1 | 2 | 2 | 2 | 36.5 | 6 | 325 | 5.64 | | | | | | | | | | | | |
| High | Q2M389 | WASH complex subunit 4 OS=Homo sapiens OX=9606 GN=WASHC4 PE=1 SV=2 | 2 | 1 | 1 | 136.3 | 1 | 1173 | 7.44 | 1 | 1 | | | | | 1 | 1 | | | | |
| High | Q96DI7 | U5 small nuclear ribonucleoprotein 40 kDa protein OS=Homo sapiens OX=9606 GN=SNRNP40 PE=1 SV=1 | 2 | 1 | 1 | 39.3 | 4 | 357 | 8.1 | | | | | | | | | | | | |
| High | O60658 | High affinity cAMP-specific and IBMX-insensitive 3',5'-cyclic phosphodiesterase 8A OS=Homo sapiens OX=9606 GN=PDE8A PE=1 S | 2 | 1 | 1 | 93.2 | 2 | 829 | 6.11 | | | 1 | | | | | | 1 | | | |
| High | P05166 | Propionyl-CoA carboxylase beta chain, mitochondrial OS=Homo sapiens OX=9606 GN=PCCB PE=1 SV=3 | 2 | 2 | 2 | 58.2 | 5 | 539 | 7.64 | 1 | | | | | | 1 | | | | | |
| High | Q96Q05 | Trafficking protein particle complex subunit 9 OS=Homo sapiens OX=9606 GN=TRAPPC9 PE=1 SV=2 | 2 | 2 | 2 | 128.4 | 2 | 1148 | 6.62 | | | | | | | | | | | | |
| High | Q9UJW0 | Dynactin subunit 4 OS=Homo sapiens OX=9606 GN=DCTN4 PE=1 SV=1 | 2 | 1 | 1 | 52.3 | 5 | 460 | 7.34 | | | 1 | | | | | | 1 | | | |
| High | O75110 | Probable phospholipid-transporting ATPase IIA OS=Homo sapiens OX=9606 GN=ATP9A PE=1 SV=3 | 2 | 2 | 2 | 118.5 | 3 | 1047 | 7.77 | | | 1 | | | | | | 1 | | | |

| | | | | | | | | | | | | | | | | | | | | | |
|---|---|---|---|---|---|---|---|---|---|---|---|---|---|---|---|---|---|---|---|---|---|
| High | Q9Y4P8 | WD repeat domain phosphoinositide-interacting protein 2 OS=Homo sapiens OX=9606 GN=WIPI2 PE=1 SV=1 | 2 | 1 | 1 | 49.4 | 4 | 454 | 6.46 | | | 1 | | | | | | 1 | | | |
| High | P36402 | Transcription factor 7 OS=Homo sapiens OX=9606 GN=TCF7 PE=1 SV=3 | 2 | 1 | 1 | 41.6 | 3 | 384 | 6.81 | 1 | 1 | | | | | 1 | 1 | | | | |
| High | Q5VTL8 | Pre-mRNA-splicing factor 38B OS=Homo sapiens OX=9606 GN=PRPF38B PE=1 SV=1 | 2 | 1 | 1 | 64.4 | 2 | 546 | 10.54 | | | | 1 | | 1 | | | | 1 | | 1 |
| High | P07585 | Decorin OS=Homo sapiens OX=9606 GN=DCN PE=1 SV=1 | 2 | 1 | 1 | 39.7 | 3 | 359 | 8.54 | | | | 1 | | 1 | | | | 1 | | 1 |
| High | Q96SB3 | Neurabin-2 OS=Homo sapiens OX=9606 GN=PPP1R9B PE=1 SV=3 | 2 | 2 | 2 | 89.3 | 3 | 817 | 4.97 | | | | | | | | | | | | |
| High | Q13617 | Cullin-2 OS=Homo sapiens OX=9606 GN=CUL2 PE=1 SV=2 | 2 | 1 | 1 | 86.9 | 1 | 745 | 6.92 | | | | | | | | | | | | |
| High | P10253 | Lysosomal alpha-glucosidase OS=Homo sapiens OX=9606 GN=GAA PE=1 SV=4 | 2 | 1 | 1 | 105.3 | 2 | 952 | 6 | 1 | 1 | | | | | 1 | 1 | | | | |
| High | Q53SF7 | Cordon-bleu protein-like 1 OS=Homo sapiens OX=9606 GN=COBLL1 PE=1 SV=3 | 2 | 1 | 1 | 123.8 | 2 | 1128 | 6.58 | | | 1 | | | | | | 1 | | | |
| High | Q9NV96 | Cell cycle control protein 50A OS=Homo sapiens OX=9606 GN=TMEM30A PE=1 SV=1 | 2 | 1 | 1 | 40.7 | 2 | 361 | 8.59 | | | | | | | | | | | | |
| High | Q9P2S5 | WD repeat-containing protein WRAP73 OS=Homo sapiens OX=9606 GN=WRAP73 PE=1 SV=1 | 2 | 1 | 1 | 51.6 | 4 | 460 | 6.81 | | | | | | | | | | | | |
| High | Q15118 | [Pyruvate dehydrogenase (acetyl-transferring)] kinase isozyme 1, mitochondrial OS=Homo sapiens OX=9606 GN=PDK1 PE=1 SV=1 | 2 | 2 | 2 | 49.2 | 6 | 436 | 8.81 | | | 1 | | | | | | 1 | | | |
| High | Q2NL82 | Pre-rRNA-processing protein TSR1 homolog OS=Homo sapiens OX=9606 GN=TSR1 PE=1 SV=1 | 2 | 2 | 2 | 91.8 | 3 | 804 | 7.42 | | | | | | | | | | | | |
| High | P07225 | Vitamin K-dependent protein S OS=Homo sapiens OX=9606 GN=PROS1 PE=1 SV=1 | 2 | 1 | 1 | 75.1 | 3 | 676 | 5.67 | | 1 | | | | | | 1 | | | | |
| High | Q14137 | Ribosome biogenesis protein BOP1 OS=Homo sapiens OX=9606 GN=BOP1 PE=1 SV=2 | 2 | 2 | 2 | 83.6 | 6 | 746 | 6.19 | | | | | | | | | | | | |
| High | Q92989 | Polyribonucleotide 5'-hydroxyl-kinase Clp1 OS=Homo sapiens OX=9606 GN=CLP1 PE=1 SV=1 | 2 | 2 | 2 | 47.6 | 4 | 425 | 6.62 | | | | | | | | | | | | |
| High | Q9P246 | Stromal interaction molecule 2 OS=Homo sapiens OX=9606 GN=STIM2 PE=1 SV=2 | 2 | 2 | 2 | 83.9 | 4 | 746 | 6.76 | | | | | | | | | | | | |
| High | Q8NI60 | Atypical kinase COQ8A, mitochondrial OS=Homo sapiens OX=9606 GN=COQ8A PE=1 SV=1 | 2 | 2 | 2 | 71.9 | 4 | 647 | 6.99 | | | 2 | | | | | | 2 | | | |
| High | Q86YZ3 | SWISS-PROT:Q86YZ3 Tax_Id=9606 Gene_Symbol=HRNR Hornerin | 2 | 1 | 1 | 282.2 | 2 | 2850 | 10.04 | | | | | 1 | | | | | | 1 | |
| High | Q9NPD8 | Ubiquitin-conjugating enzyme E2 T OS=Homo sapiens OX=9606 GN=UBE2T PE=1 SV=1 | 2 | 2 | 2 | 22.5 | 13 | 197 | 7.99 | | | | | 1 | 1 | | | | | 1 | 1 |
| High | Q9UHI6 | Probable ATP-dependent RNA helicase DDX20 OS=Homo sapiens OX=9606 GN=DDX20 PE=1 SV=2 | 2 | 2 | 2 | 92.2 | 4 | 824 | 6.95 | 1 | | | | | | 1 | | | | | |
| High | P46783 | 40S ribosomal protein S10 OS=Homo sapiens OX=9606 GN=RPS10 PE=1 SV=1 | 2 | 1 | 1 | 18.9 | 5 | 165 | 10.15 | | 1 | | | | | | 1 | | | | |
| High | Q6IQ49 | Replication stress response regulator SDE2 OS=Homo sapiens OX=9606 GN=SDE2 PE=1 SV=1 | 2 | 1 | 1 | 49.7 | 5 | 451 | 6.05 | | | | | | | | | | | | |
| High | Q5T3J3 | Ligand-dependent nuclear receptor-interacting factor 1 OS=Homo sapiens OX=9606 GN=LRIF1 PE=1 SV=1 | 2 | 2 | 2 | 84.5 | 5 | 769 | 9.72 | | | | | | | | | | | | |
| High | Q9BZZ5 | Apoptosis inhibitor 5 OS=Homo sapiens OX=9606 GN=API5 PE=1 SV=3 | 2 | 1 | 1 | 59 | 2 | 524 | 7.34 | | | | | | 1 | | | | | | 1 |
| High | Q7Z6L1 | Tectonin beta-propeller repeat-containing protein 1 OS=Homo sapiens OX=9606 GN=TECPR1 PE=1 SV=1 | 2 | 1 | 1 | 129.6 | 1 | 1165 | 6.24 | | | | | | | | | | | | |
| High | Q05932 | Folylpolyglutamate synthase, mitochondrial OS=Homo sapiens OX=9606 GN=FPGS PE=1 SV=3 | 2 | 1 | 1 | 64.6 | 3 | 587 | 7.94 | | | 1 | | | | | | 1 | | | |
| High | Q5EBL8 | PDZ domain-containing protein 11 OS=Homo sapiens OX=9606 GN=PDZD11 PE=1 SV=2 | 2 | 1 | 1 | 16.1 | 19 | 140 | 7.21 | 1 | 1 | | | | | 1 | 1 | | | | |
| High | Q9UHP3 | Ubiquitin carboxyl-terminal hydrolase 25 OS=Homo sapiens OX=9606 GN=USP25 PE=1 SV=4 | 2 | 1 | 1 | 122.1 | 1 | 1055 | 5.34 | | | | | | | | | | | | |
| High | Q86U44 | N6-adenosine-methyltransferase catalytic subunit OS=Homo sapiens OX=9606 GN=METTL3 PE=1 SV=2 | 2 | 1 | 1 | 64.4 | 2 | 580 | 6.42 | 1 | 1 | | | | | 1 | 1 | | | | |
| High | Q9H981 | Actin-related protein 8 OS=Homo sapiens OX=9606 GN=ACTR8 PE=1 SV=2 | 2 | 1 | 1 | 70.4 | 1 | 624 | 7.78 | | | | | 1 | | | | | | 1 | |
| High | Q12765 | Secernin-1 OS=Homo sapiens OX=9606 GN=SCRN1 PE=1 SV=2 | 2 | 1 | 1 | 46.4 | 4 | 414 | 4.75 | | | | | | | | | | | | |
| High | O75822 | Eukaryotic translation initiation factor 3 subunit J OS=Homo sapiens OX=9606 GN=EIF3J PE=1 SV=2 | 2 | 2 | 2 | 29 | 9 | 258 | 4.83 | | | 1 | | | | | | 1 | | | |
| High | Q9NX00 | Transmembrane protein 160 OS=Homo sapiens OX=9606 GN=TMEM160 PE=1 SV=1 | 2 | 1 | 1 | 19.6 | 4 | 188 | 8.03 | | | | | 1 | 1 | | | | | 1 | 1 |
| High | Q7Z7H5 | Transmembrane emp24 domain-containing protein 4 OS=Homo sapiens OX=9606 GN=TMED4 PE=1 SV=1 | 2 | 1 | 1 | 25.9 | 4 | 227 | 8.28 | 1 | 1 | | | | | 1 | 1 | | | | |
| High | Q96RF0 | Sorting nexin-18 OS=Homo sapiens OX=9606 GN=SNX18 PE=1 SV=2 | 2 | 1 | 1 | 68.9 | 3 | 628 | 5.68 | | | | | | | | | | | | |
| High | P11766 | Alcohol dehydrogenase class-3 OS=Homo sapiens OX=9606 GN=ADH5 PE=1 SV=4 | 2 | 1 | 1 | 39.7 | 4 | 374 | 7.49 | | | 1 | | | | | | 1 | | | |
| High | Q969F9 | Hermansky-Pudlak syndrome 3 protein OS=Homo sapiens OX=9606 GN=HPS3 PE=1 SV=1 | 2 | 1 | 1 | 113.7 | 2 | 1004 | 6.43 | 1 | 1 | | | | | 1 | 1 | | | | |
| High | P51452 | Dual specificity protein phosphatase 3 OS=Homo sapiens OX=9606 GN=DUSP3 PE=1 SV=1 | 2 | 2 | 2 | 20.5 | 12 | 185 | 7.8 | | | 1 | | | 1 | | | 1 | | | 1 |
| High | Q15904 | V-type proton ATPase subunit S1 OS=Homo sapiens OX=9606 GN=ATP6AP1 PE=1 SV=2 | 2 | 1 | 1 | 52 | 2 | 470 | 6.14 | | | | | | | | | | | | |
| High | P05109 | Protein S100-A8 OS=Homo sapiens OX=9606 GN=S100A8 PE=1 SV=1 | 2 | 1 | 1 | 10.8 | 12 | 93 | 7.03 | | | | 1 | 1 | | | | | 1 | 1 | |
| High | O75044 | SLIT-ROBO Rho GTPase-activating protein 2 OS=Homo sapiens OX=9606 GN=SRGAP2 PE=1 SV=3 | 2 | 2 | 2 | 120.8 | 3 | 1071 | 6.7 | | 1 | | | | | | 1 | | | | |
| High | P31271 | Homeobox protein Hox-A13 OS=Homo sapiens OX=9606 GN=HOXA13 PE=1 SV=3 | 2 | 2 | 2 | 39.7 | 7 | 388 | 9.13 | 1 | | | | | | 1 | | | | | |
| High | P03886 | NADH-ubiquinone oxidoreductase chain 1 OS=Homo sapiens OX=9606 GN=MT-ND1 PE=1 SV=1 | 2 | 1 | 1 | 35.6 | 3 | 318 | 6.55 | 1 | 1 | | | | | 1 | 1 | | | | |
| High | Q9HB40 | Retinoid-inducible serine carboxypeptidase OS=Homo sapiens OX=9606 GN=SCPEP1 PE=1 SV=1 | 2 | 1 | 1 | 50.8 | 2 | 452 | 5.81 | 1 | | 1 | | | | 1 | | 1 | | | |
| High | Q66PJ3 | ADP-ribosylation factor-like protein 6-interacting protein 4 OS=Homo sapiens OX=9606 GN=ARL6IP4 PE=1 SV=3 | 2 | 1 | 1 | 26.4 | 8 | 237 | 10.86 | | | | | | | | | | | | |
| High | Q9C002 | Normal mucosa of esophagus-specific gene 1 protein OS=Homo sapiens OX=9606 GN=NMES1 PE=1 SV=1 | 2 | 2 | 2 | 9.6 | 28 | 83 | 9.47 | | | | | | | | | | | | |
| High | Q9NW82 | WD repeat-containing protein 70 OS=Homo sapiens OX=9606 GN=WDR70 PE=1 SV=1 | 2 | 2 | 2 | 73.2 | 4 | 654 | 6.33 | | | | | | | | | | | | |
| High | Q99418 | Cytohesin-2 OS=Homo sapiens OX=9606 GN=CYTH2 PE=1 SV=2 | 2 | 2 | 2 | 46.5 | 7 | 400 | 5.54 | | | 1 | | | 1 | | | 1 | | | 1 |
| High | Q9BZL1 | Ubiquitin-like protein 5 OS=Homo sapiens OX=9606 GN=UBL5 PE=1 SV=1 | 2 | 1 | 1 | 8.5 | 12 | 73 | 8.44 | 1 | 1 | | | | | 1 | 1 | | | | |
| High | Q13608 | Peroxisome assembly factor 2 OS=Homo sapiens OX=9606 GN=PEX6 PE=1 SV=2 | 2 | 2 | 2 | 104 | 1 | 980 | 6.34 | | | | | | | | | | | | |
| High | Q9P031 | Thyroid transcription factor 1-associated protein 26 OS=Homo sapiens OX=9606 GN=CCDC59 PE=1 SV=2 | 2 | 1 | 1 | 28.7 | 5 | 241 | 9.88 | | | | | | | | | | | | |
| High | Q9UM00 | Calcium load-activated calcium channel OS=Homo sapiens OX=9606 GN=TMCO1 PE=1 SV=2 | 2 | 1 | 1 | 27.1 | 6 | 239 | 10.26 | 1 | 1 | | | | | 1 | 1 | | | | |
| High | P49459 | Ubiquitin-conjugating enzyme E2 A OS=Homo sapiens OX=9606 GN=UBE2A PE=1 SV=2 | 2 | 1 | 1 | 17.3 | 7 | 152 | 5.15 | | | | | | | | | | | | |
| High | Q96QZ7 | Membrane-associated guanylate kinase, WW and PDZ domain-containing protein 1 OS=Homo sapiens OX=9606 GN=MAGI1 PE=1 S | 2 | 1 | 1 | 164.5 | 1 | 1491 | 7.58 | | | | | | | | | | | | |
| High | Q9P2W9 | Syntaxin-18 OS=Homo sapiens OX=9606 GN=STX18 PE=1 SV=1 | 2 | 1 | 1 | 38.7 | 5 | 335 | 5.49 | | 1 | 1 | | | | | 1 | 1 | | | |
| High | O15254 | Peroxisomal acyl-coenzyme A oxidase 3 OS=Homo sapiens OX=9606 GN=ACOX3 PE=1 SV=2 | 2 | 1 | 1 | 77.6 | 3 | 700 | 7.25 | | | | | | | | | | | | |
| High | P42765 | 3-ketoacyl-CoA thiolase, mitochondrial OS=Homo sapiens OX=9606 GN=ACAA2 PE=1 SV=2 | 2 | 1 | 1 | 41.9 | 2 | 397 | 8.09 | | 1 | 1 | | | | | 1 | 1 | | | |
| High | Q9UQR1 | Zinc finger protein 148 OS=Homo sapiens OX=9606 GN=ZNF148 PE=1 SV=2 | 2 | 2 | 2 | 88.9 | 2 | 794 | 6.48 | 1 | | 1 | | | | 1 | | 1 | | | |
| High | Q9Y6W3 | Calpain-7 OS=Homo sapiens OX=9606 GN=CAPN7 PE=1 SV=1 | 2 | 1 | 1 | 92.6 | 1 | 813 | 7.65 | 1 | 1 | | | | | 1 | 1 | | | | |
| High | P10606 | Cytochrome c oxidase subunit 5B, mitochondrial OS=Homo sapiens OX=9606 GN=COX5B PE=1 SV=2 | 2 | 1 | 1 | 13.7 | 6 | 129 | 8.81 | | | | | | | | | | | | |
| High | Q13751 | Laminin subunit beta-3 OS=Homo sapiens OX=9606 GN=LAMB3 PE=1 SV=1 | 2 | 2 | 2 | 129.5 | 2 | 1172 | 7.21 | | | | | | | | | | | | |
| High | O60306 | RNA helicase aquarius OS=Homo sapiens OX=9606 GN=AQR PE=1 SV=4 | 2 | 2 | 2 | 171.2 | 2 | 1485 | 6.37 | | | | | 1 | | | | | | 1 | |
| High | Q8N0Z8 | tRNA pseudouridine synthase-like 1 OS=Homo sapiens OX=9606 GN=PUSL1 PE=1 SV=1 | 2 | 1 | 1 | 33.2 | 3 | 303 | 9.85 | | 1 | | | | | | 1 | | | | |
| High | O15321 | Transmembrane 9 superfamily member 1 OS=Homo sapiens OX=9606 GN=TM9SF1 PE=2 SV=2 | 2 | 1 | 1 | 68.8 | 1 | 606 | 7.17 | 1 | 1 | | | | | 1 | 1 | | | | |
| High | P19532 | Transcription factor E3 OS=Homo sapiens OX=9606 GN=TFE3 PE=1 SV=4 | 2 | 2 | 2 | 61.5 | 3 | 575 | 5.58 | | 1 | 1 | | | | | 1 | 1 | | | |
| High | Q13642 | Four and a half LIM domains protein 1 OS=Homo sapiens OX=9606 GN=FHL1 PE=1 SV=4 | 2 | 1 | 1 | 36.2 | 4 | 323 | 8.97 | | | | 1 | | 1 | | | | 1 | | 1 |
| High | Q9BV68 | E3 ubiquitin-protein ligase RNF126 OS=Homo sapiens OX=9606 GN=RNF126 PE=1 SV=2 | 2 | 1 | 1 | 33.8 | 8 | 311 | 5.47 | | | | | | | | | | | | |
| High | Q969X1 | Protein lifeguard 3 OS=Homo sapiens OX=9606 GN=TMBIM1 PE=1 SV=2 | 2 | 1 | 1 | 34.6 | 5 | 311 | 7.72 | 1 | | 1 | | | | 1 | | 1 | | | |
| High | Q9H2H8 | Peptidyl-prolyl cis-trans isomerase-like 3 OS=Homo sapiens OX=9606 GN=PPIL3 PE=1 SV=1 | 2 | 1 | 1 | 18.1 | 12 | 161 | 6.79 | 1 | | | | | | 1 | | | | | |
| High | O75691 | Small subunit processome component 20 homolog OS=Homo sapiens OX=9606 GN=UTP20 PE=1 SV=3 | 2 | 2 | 2 | 318.2 | 1 | 2785 | 7.39 | | | | | | | | | | | | |
| High | O43504 | Ragulator complex protein LAMTOR5 OS=Homo sapiens OX=9606 GN=LAMTOR5 PE=1 SV=1 | 2 | 1 | 1 | 9.6 | 26 | 91 | 4.87 | | | | | | | | | | | | |
| High | Q9P227 | Rho GTPase-activating protein 23 OS=Homo sapiens OX=9606 GN=ARHGAP23 PE=1 SV=2 | 2 | 1 | 1 | 162.1 | 1 | 1491 | 9.19 | | | | | | | | | | | | |
| High | P11047 | Laminin subunit gamma-1 OS=Homo sapiens OX=9606 GN=LAMC1 PE=1 SV=3 | 2 | 1 | 1 | 177.5 | 1 | 1609 | 5.12 | | | | | 1 | 1 | | | | | 1 | 1 |
| High | P54098 | DNA polymerase subunit gamma-1 OS=Homo sapiens OX=9606 GN=POLG PE=1 SV=1 | 2 | 1 | 1 | 139.5 | 1 | 1239 | 6.9 | | | 1 | | | | | | 1 | | | |
| High | P46976 | Glycogenin-1 OS=Homo sapiens OX=9606 GN=GYG1 PE=1 SV=4 | 2 | 2 | 2 | 39.4 | 7 | 350 | 5.53 | | | | | | | | | | | | |
| High | P61457 | Pterin-4-alpha-carbinolamine dehydratase OS=Homo sapiens OX=9606 GN=PCBD1 PE=1 SV=2 | 2 | 1 | 1 | 12 | 10 | 104 | 6.8 | 1 | | | | | | 1 | | | | | |
| High | P07477 | SWISS-PROT:P07477 Tax_Id=9606 Gene_Symbol=PRSS1 Trypsin-1 precursor | 2 | 1 | 1 | 26.5 | 8 | 247 | 6.51 | 1 | | | | | | 1 | | | | | |
| High | Q8N594 | MPN domain-containing protein OS=Homo sapiens OX=9606 GN=MPND PE=1 SV=1 | 2 | 2 | 2 | 50.6 | 6 | 471 | 5.72 | | 1 | 1 | | | | | 1 | 1 | | | |
| High | P51159 | Ras-related protein Rab-27A OS=Homo sapiens OX=9606 GN=RAB27A PE=1 SV=3 | 2 | 2 | 2 | 24.9 | 8 | 221 | 5.22 | | | | | | | | | | | | |
| High | Q8WU90 | Zinc finger CCCH domain-containing protein 15 OS=Homo sapiens OX=9606 GN=ZC3H15 PE=1 SV=1 | 2 | 2 | 2 | 48.6 | 6 | 426 | 5.31 | | | | | 1 | | | | | | 1 | |
| High | O75717 | WD repeat and HMG-box DNA-binding protein 1 OS=Homo sapiens OX=9606 GN=WDHD1 PE=1 SV=1 | 2 | 1 | 1 | 125.9 | 1 | 1129 | 5.62 | | | | | | | | | | | | |
| High | O43929 | Origin recognition complex subunit 4 OS=Homo sapiens OX=9606 GN=ORC4 PE=1 SV=2 | 2 | 2 | 2 | 50.3 | 6 | 436 | 8 | 1 | 1 | | | | | 1 | 1 | | | | |
| High | O60664 | Perilipin-3 OS=Homo sapiens OX=9606 GN=PLIN3 PE=1 SV=3 | 2 | 1 | 1 | 47 | 4 | 434 | 5.44 | 1 | 1 | | | | | 1 | 1 | | | | |
| High | Q15828 | Cystatin-M OS=Homo sapiens OX=9606 GN=CST6 PE=1 SV=1 | 2 | 1 | 1 | 16.5 | 7 | 149 | 8.09 | | | | 1 | | 1 | | | | 1 | | 1 |
| High | Q9Y572 | Receptor-interacting serine/threonine-protein kinase 3 OS=Homo sapiens OX=9606 GN=RIPK3 PE=1 SV=2 | 2 | 1 | 1 | 56.9 | 4 | 518 | 6.47 | | | 1 | | | | | | 1 | | | |
| High | Q04446 | 1,4-alpha-glucan-branching enzyme OS=Homo sapiens OX=9606 GN=GBE1 PE=1 SV=3 | 2 | 1 | 1 | 80.4 | 1 | 702 | 6.32 | | | | | | | | | | | | |
| High | Q9BVG9 | Phosphatidylserine synthase 2 OS=Homo sapiens OX=9606 GN=PTDSS2 PE=1 SV=1 | 2 | 1 | 1 | 56.2 | 5 | 487 | 6.25 | 1 | | 1 | | | | 1 | | 1 | | | |
| High | Q99797 | Mitochondrial intermediate peptidase OS=Homo sapiens OX=9606 GN=MIPEP PE=1 SV=2 | 2 | 1 | 1 | 80.6 | 1 | 713 | 7.05 | | | | | | | | | | | | |
| High | Q99943 | 1-acyl-sn-glycerol-3-phosphate acyltransferase alpha OS=Homo sapiens OX=9606 GN=AGPAT1 PE=1 SV=2 | 2 | 1 | 1 | 31.7 | 7 | 283 | 9.38 | | | | | | | | | | | | |
| High | P54792 | Putative segment polarity protein dishevelled homolog DVL1P1 OS=Homo sapiens OX=9606 GN=DVL1P1 PE=5 SV=1 | 2 | 1 | 1 | 73.2 | 1 | 670 | 6.46 | | | | | 1 | 1 | | | | | 1 | 1 |
| High | P03928 | ATP synthase protein 8 OS=Homo sapiens OX=9606 GN=MT-ATP8 PE=1 SV=1 | 2 | 1 | 1 | 8 | 13 | 68 | 9.91 | | | 2 | | | | | | 1 | | | |
| High | Q9HB58 | Sp110 nuclear body protein OS=Homo sapiens OX=9606 GN=SP110 PE=1 SV=5 | 2 | 2 | 2 | 78.3 | 3 | 689 | 8.87 | | | | | | 1 | | | | | | 1 |
| High | Q8IY18 | Structural maintenance of chromosomes protein 5 OS=Homo sapiens OX=9606 GN=SMC5 PE=1 SV=2 | 2 | 1 | 1 | 128.7 | 1 | 1101 | 8.38 | | | | | | | | | | | | |
| High | P17174 | Aspartate aminotransferase, cytoplasmic OS=Homo sapiens OX=9606 GN=GOT1 PE=1 SV=3 | 2 | 1 | 1 | 46.2 | 2 | 413 | 7.01 | | | 2 | | | | | | 1 | | | |
| High | P53990 | IST1 homolog OS=Homo sapiens OX=9606 GN=IST1 PE=1 SV=1 | 2 | 2 | 2 | 39.7 | 8 | 364 | 5.35 | | | | | | | | | | | | |
| High | P61073 | C-X-C chemokine receptor type 4 OS=Homo sapiens OX=9606 GN=CXCR4 PE=1 SV=1 | 2 | 1 | 1 | 39.7 | 3 | 352 | 8.21 | 1 | | 1 | | | | 1 | | 1 | | | |
| High | P51809 | Vesicle-associated membrane protein 7 OS=Homo sapiens OX=9606 GN=VAMP7 PE=1 SV=3 | 2 | 1 | 1 | 24.9 | 5 | 220 | 8.6 | 1 | 1 | | | | | 1 | 1 | | | | |
| High | Q9Y3X0 | Coiled-coil domain-containing protein 9 OS=Homo sapiens OX=9606 GN=CCDC9 PE=1 SV=1 | 2 | 2 | 2 | 59.7 | 4 | 531 | 5.4 | | 1 | | 1 | | | | 1 | | 1 | | |
| High | Q8NEW0 | Zinc transporter 7 OS=Homo sapiens OX=9606 GN=SLC30A7 PE=2 SV=1 | 2 | 1 | 1 | 41.6 | 4 | 376 | 6.95 | 1 | 1 | | | | | 1 | 1 | | | | |
| High | Q92908 | Transcription factor GATA-6 OS=Homo sapiens OX=9606 GN=GATA6 PE=1 SV=2 | 2 | 1 | 1 | 60 | 2 | 595 | 8.35 | | | | | | | | | | | | |
| High | Q96GC9 | Vacuole membrane protein 1 OS=Homo sapiens OX=9606 GN=VMP1 PE=1 SV=1 | 2 | 1 | 1 | 46.2 | 2 | 406 | 6.95 | | | | | | | | | | | | |
| High | Q9Y2Y1 | DNA-directed RNA polymerase III subunit RPC10 OS=Homo sapiens OX=9606 GN=POLR3K PE=1 SV=3 | 2 | 1 | 1 | 12.3 | 13 | 108 | 7.83 | | | | | | | | | | | | |
| High | Q14644 | Ras GTPase-activating protein 3 OS=Homo sapiens OX=9606 GN=RASA3 PE=1 SV=3 | 2 | 2 | 2 | 95.6 | 2 | 834 | 7.15 | | | | | 1 | | | | | | 1 | |
| High | Q9BQE5 | Apolipoprotein L2 OS=Homo sapiens OX=9606 GN=APOL2 PE=1 SV=1 | 2 | 1 | 1 | 37.1 | 3 | 337 | 6.74 | | | | | | | | | | | | |
| High | Q9Y679 | Lipid droplet-regulating VLDL assembly factor AUP1 OS=Homo sapiens OX=9606 GN=AUP1 PE=1 SV=2 | 2 | 2 | 2 | 45.8 | 6 | 410 | 8.65 | 1 | | | | | | 1 | | | | | |
| High | Q7Z404 | Transmembrane channel-like protein 4 OS=Homo sapiens OX=9606 GN=TMC4 PE=1 SV=3 | 2 | 1 | 1 | 79.2 | 2 | 712 | 8.94 | | | 1 | | | | | | 1 | | | |
| High | Q8NBT2 | Kinetochore protein Spc24 OS=Homo sapiens OX=9606 GN=SPC24 PE=1 SV=3 | 2 | 1 | 1 | 22.4 | 7 | 197 | 4.7 | 1 | 1 | | | | | 1 | 1 | | | | |
| High | Q9Y4P1 | Cysteine protease ATG4B OS=Homo sapiens OX=9606 GN=ATG4B PE=1 SV=2 | 2 | 2 | 2 | 44.3 | 8 | 393 | 5.07 | | | | | | | | | | | | |
| High | Q5T0N5 | Formin-binding protein 1-like OS=Homo sapiens OX=9606 GN=FNBP1L PE=1 SV=3 | 2 | 1 | 1 | 70 | 1 | 605 | 6.64 | | | 1 | | | | | | 1 | | | |
| High | O95786 | Antiviral innate immune response receptor RIG-I OS=Homo sapiens OX=9606 GN=DDX58 PE=1 SV=2 | 2 | 1 | 1 | 106.5 | 1 | 925 | 6.4 | | | | | 1 | | | | | | 1 | |
| High | Q07654 | Trefoil factor 3 OS=Homo sapiens OX=9606 GN=TFF3 PE=1 SV=3 | 2 | 1 | 1 | 8.6 | 14 | 80 | 5.92 | | | | | | | | | | | | |
| High | Q9Y2Q9 | 28S ribosomal protein S28, mitochondrial OS=Homo sapiens OX=9606 GN=MRPS28 PE=1 SV=1 | 2 | 2 | 2 | 20.8 | 11 | 187 | 9.1 | | | | | | 1 | | | | | | 1 |
| High | P53611 | Geranylgeranyl transferase type-2 subunit beta OS=Homo sapiens OX=9606 GN=RABGGTB PE=1 SV=2 | 2 | 1 | 1 | 36.9 | 3 | 331 | 5.03 | | | | 1 | | 1 | | | | 1 | | 1 |
| High | Q4KMP7 | TBC1 domain family member 10B OS=Homo sapiens OX=9606 GN=TBC1D10B PE=1 SV=3 | 2 | 1 | 1 | 87.1 | 3 | 808 | 9.19 | | | | | | | | | | | | |
| High | P55210 | Caspase-7 OS=Homo sapiens OX=9606 GN=CASP7 PE=1 SV=1 | 2 | 1 | 1 | 34.3 | 8 | 303 | 6.07 | | | | 1 | | | | | | 1 | | |
| High | P06746 | DNA polymerase beta OS=Homo sapiens OX=9606 GN=POLB PE=1 SV=3 | 2 | 1 | 1 | 38.2 | 3 | 335 | 8.95 | | | | 1 | | 1 | | | | 1 | | 1 |
| High | O14732 | Inositol monophosphatase 2 OS=Homo sapiens OX=9606 GN=IMPA2 PE=1 SV=1 | 2 | 2 | 2 | 31.3 | 8 | 288 | 6.61 | | | | 1 | | | | | | 1 | | |
| High | Q9NPI1 | Bromodomain-containing protein 7 OS=Homo sapiens OX=9606 GN=BRD7 PE=1 SV=1 | 2 | 2 | 2 | 74.1 | 6 | 651 | 6.39 | | | | | | | | | | | | |
| High | Q13469 | Nuclear factor of activated T-cells, cytoplasmic 2 OS=Homo sapiens OX=9606 GN=NFATC2 PE=1 SV=2 | 2 | 1 | 1 | 100.1 | 2 | 925 | 7.28 | 1 | 1 | | | | | 1 | 1 | | | | |
| High | Q9NZM3 | Intersectin-2 OS=Homo sapiens OX=9606 GN=ITSN2 PE=1 SV=3 | 2 | 1 | 1 | 193.3 | 1 | 1697 | 8.12 | | | | | | | | | | | | |
| High | Q8WWY3 | U4/U6 small nuclear ribonucleoprotein Prp31 OS=Homo sapiens OX=9606 GN=PRPF31 PE=1 SV=2 | 2 | 2 | 2 | 55.4 | 7 | 499 | 5.78 | | | | | | | | | | | | |
| High | Q6Q0C0 | E3 ubiquitin-protein ligase TRAF7 OS=Homo sapiens OX=9606 GN=TRAF7 PE=1 SV=1 | 2 | 1 | 1 | 74.6 | 1 | 670 | 7.15 | | | | 1 | | 1 | | | | 1 | | 1 |
| High | P19075 | Tetraspanin-8 OS=Homo sapiens OX=9606 GN=TSPAN8 PE=1 SV=1 | 2 | 1 | 1 | 26 | 4 | 237 | 5.6 | | | | | | | | | | | | |
| High | Q14978 | Nucleolar and coiled-body phosphoprotein 1 OS=Homo sapiens OX=9606 GN=NOLC1 PE=1 SV=2 | 2 | 1 | 1 | 73.6 | 1 | 699 | 9.47 | | | | | 1 | 1 | | | | | 1 | 1 |
| High | Q2PZI1 | Probable C-mannosyltransferase DPY19L1 OS=Homo sapiens OX=9606 GN=DPY19L1 PE=2 SV=1 | 2 | 1 | 1 | 77.3 | 3 | 675 | 8.95 | | | 1 | | | | | | 1 | | | |
| High | Q9BPX6 | Calcium uptake protein 1, mitochondrial OS=Homo sapiens OX=9606 GN=MICU1 PE=1 SV=1 | 2 | 1 | 1 | 54.3 | 3 | 476 | 8.41 | | | 1 | | | | | | 1 | | | |
| High | O14966 | Ras-related protein Rab-7L1 OS=Homo sapiens OX=9606 GN=RAB29 PE=1 SV=1 | 2 | 1 | 1 | 23.1 | 5 | 203 | 7.18 | | | | | | | | | | | | |
| High | P51790 | H(+)/Cl(-) exchange transporter 3 OS=Homo sapiens OX=9606 GN=CLCN3 PE=1 SV=2 | 2 | 2 | 2 | 90.9 | 2 | 818 | 6.28 | | 1 | 1 | | | | | 1 | 1 | | | |
| High | Q9BZK3 | Putative nascent polypeptide-associated complex subunit alpha-like protein OS=Homo sapiens OX=9606 GN=NACA4P PE=5 SV=1 | 2 | 1 | 1 | 23.3 | 7 | 213 | 4.59 | | | | | | | | | | | | |
| High | Q9BVM2 | Protein DPCD OS=Homo sapiens OX=9606 GN=DPCD PE=1 SV=2 | 2 | 1 | 1 | 23.2 | 4 | 203 | 9.03 | | | | | | | | | | | | |
| High | Q5C9Z4 | Nucleolar MIF4G domain-containing protein 1 OS=Homo sapiens OX=9606 GN=NOM1 PE=1 SV=1 | 2 | 1 | 1 | 96.2 | 1 | 860 | 8.1 | | | | | | | | | | | | |
| High | P53567 | CCAAT/enhancer-binding protein gamma OS=Homo sapiens OX=9606 GN=CEBPG PE=1 SV=1 | 2 | 1 | 1 | 16.4 | 21 | 150 | 9.77 | 1 | 1 | | | | | 1 | 1 | | | | |
| High | O15169 | Axin-1 OS=Homo sapiens OX=9606 GN=AXIN1 PE=1 SV=2 | 2 | 1 | 1 | 95.6 | 2 | 862 | 6.96 | | | | | | | | | | | | |
| High | O94913 | Pre-mRNA cleavage complex 2 protein Pcf11 OS=Homo sapiens OX=9606 GN=PCF11 PE=1 SV=3 | 2 | 1 | 1 | 172.9 | 1 | 1555 | 8.48 | | | | | | | | | | | | |
| High | Q8N3U4 | Cohesin subunit SA-2 OS=Homo sapiens OX=9606 GN=STAG2 PE=1 SV=3 | 2 | 2 | 2 | 141.2 | 2 | 1231 | 5.43 | 1 | 1 | | | | | 1 | 1 | | | | |
| High | Q15005 | Signal peptidase complex subunit 2 OS=Homo sapiens OX=9606 GN=SPCS2 PE=1 SV=3 | 2 | 2 | 2 | 25 | 8 | 226 | 8.47 | 1 | | | | | | 1 | | | | | |
| High | Q6IQ23 | Pleckstrin homology domain-containing family A member 7 OS=Homo sapiens OX=9606 GN=PLEKHA7 PE=1 SV=2 | 2 | 2 | 2 | 127.1 | 2 | 1121 | 9.35 | | | 1 | | | | | | 1 | | | |
| High | Q02809 | Procollagen-lysine,2-oxoglutarate 5-dioxygenase 1 OS=Homo sapiens OX=9606 GN=PLOD1 PE=1 SV=2 | 2 | 1 | 1 | 83.5 | 2 | 727 | 6.95 | | | 1 | | | | | | 1 | | | |
| High | Q9BQP7 | Mitochondrial genome maintenance exonuclease 1 OS=Homo sapiens OX=9606 GN=MGME1 PE=1 SV=1 | 2 | 1 | 1 | 39.4 | 6 | 344 | 7.68 | | | | | 1 | | | | | | 1 | |
| High | O95155 | Ubiquitin conjugation factor E4 B OS=Homo sapiens OX=9606 GN=UBE4B PE=1 SV=1 | 2 | 1 | 1 | 146.1 | 1 | 1302 | 6.55 | 1 | | 1 | | | | 1 | | 1 | | | |
| High | Q9BQC6 | Ribosomal protein 63, mitochondrial OS=Homo sapiens OX=9606 GN=MRPL57 PE=1 SV=1 | 2 | 1 | 1 | 12.3 | 13 | 102 | 11.44 | | | 1 | | | | | | 1 | | | |
| High | Q9NZM5 | Ribosome biogenesis protein NOP53 OS=Homo sapiens OX=9606 GN=NOP53 PE=1 SV=2 | 2 | 1 | 1 | 54.4 | 4 | 478 | 10.32 | | | | | | | | | | | | |
| High | Q92541 | RNA polymerase-associated protein RTF1 homolog OS=Homo sapiens OX=9606 GN=RTF1 PE=1 SV=4 | 2 | 2 | 2 | 80.3 | 4 | 710 | 8.15 | | | | | | | | | | | | |
| High | P55809 | Succinyl-CoA:3-ketoacid coenzyme A transferase 1, mitochondrial OS=Homo sapiens OX=9606 GN=OXCT1 PE=1 SV=1 | 2 | 1 | 1 | 56.1 | 6 | 520 | 7.46 | | | | | 2 | | | | | | 1 | |
| High | Q12788 | Transducin beta-like protein 3 OS=Homo sapiens OX=9606 GN=TBL3 PE=1 SV=2 | 2 | 1 | 1 | 89 | 2 | 808 | 6.9 | | | | | 1 | | | | | | 1 | |
| High | Q96CM8 | Medium-chain acyl-CoA ligase ACSF2, mitochondrial OS=Homo sapiens OX=9606 GN=ACSF2 PE=1 SV=2 | 2 | 1 | 1 | 68.1 | 3 | 615 | 7.55 | | | 1 | | | | | | 1 | | | |
| High | Q5T0D9 | Tumor protein p63-regulated gene 1-like protein OS=Homo sapiens OX=9606 GN=TPRG1L PE=1 SV=1 | 2 | 1 | 1 | 30.2 | 7 | 272 | 7.34 | 1 | 1 | | | | | 1 | 1 | | | | |
| High | Q6ZNJ1 | Neurobeachin-like protein 2 OS=Homo sapiens OX=9606 GN=NBEAL2 PE=1 SV=2 | 2 | 1 | 1 | 302.3 | 1 | 2754 | 6.38 | | | 1 | | | | | | 1 | | | |
| High | P32121 | Beta-arrestin-2 OS=Homo sapiens OX=9606 GN=ARRB2 PE=1 SV=2 | 2 | 1 | 1 | 46.1 | 4 | 409 | 7.69 | | | | | 1 | | | | | | 1 | |
| High | Q02040 | A-kinase anchor protein 17A OS=Homo sapiens OX=9606 GN=AKAP17A PE=1 SV=2 | 2 | 2 | 2 | 80.7 | 3 | 695 | 9.73 | | 1 | | | | | | 1 | | | | |
| High | Q9BQ13 | BTB/POZ domain-containing protein KCTD14 OS=Homo sapiens OX=9606 GN=KCTD14 PE=1 SV=2 | 2 | 1 | 1 | 29.6 | 5 | 255 | 8.59 | | | | | | | | | | | | |
| High | P13569 | Cystic fibrosis transmembrane conductance regulator OS=Homo sapiens OX=9606 GN=CFTR PE=1 SV=3 | 2 | 1 | 1 | 168 | 1 | 1480 | 8.73 | | 1 | 1 | | | | | 1 | 1 | | | |
| High | Q13144 | Translation initiation factor eIF-2B subunit epsilon OS=Homo sapiens OX=9606 GN=EIF2B5 PE=1 SV=3 | 2 | 2 | 2 | 80.3 | 4 | 721 | 5.08 | 1 | | | | | | 1 | | | | | |
| High | P08582 | Melanotransferrin OS=Homo sapiens OX=9606 GN=MELTF PE=1 SV=2 | 2 | 1 | 1 | 80.2 | 3 | 738 | 5.94 | | | | | | | | | | | | |
| High | P07451 | Carbonic anhydrase 3 OS=Homo sapiens OX=9606 GN=CA3 PE=1 SV=3 | 2 | 1 | 1 | 29.5 | 3 | 260 | 7.34 | | | | 1 | | 1 | | | | 1 | | 1 |
| High | Q9NWW5 | Ceroid-lipofuscinosis neuronal protein 6 OS=Homo sapiens OX=9606 GN=CLN6 PE=1 SV=1 | 2 | 1 | 1 | 35.9 | 7 | 311 | 8.87 | | | | | | | | | | | | |
| High | Q96FS4 | Signal-induced proliferation-associated protein 1 OS=Homo sapiens OX=9606 GN=SIPA1 PE=1 SV=1 | 2 | 1 | 1 | 112.1 | 1 | 1042 | 6.6 | | | 1 | 1 | | | | | 1 | 1 | | |
| High | Q92625 | Ankyrin repeat and SAM domain-containing protein 1A OS=Homo sapiens OX=9606 GN=ANKS1A PE=1 SV=4 | 2 | 1 | 1 | 123 | 2 | 1134 | 6.38 | 1 | 1 | | | | | 1 | 1 | | | | |
| High | Q9BW71 | HIRA-interacting protein 3 OS=Homo sapiens OX=9606 GN=HIRIP3 PE=1 SV=3 | 2 | 1 | 1 | 61.9 | 2 | 556 | 8.54 | | | | | | | | | | | | |
| High | Q9H1I8 | Activating signal cointegrator 1 complex subunit 2 OS=Homo sapiens OX=9606 GN=ASCC2 PE=1 SV=3 | 2 | 1 | 1 | 86.3 | 2 | 757 | 5.16 | 1 | | | | | | 1 | | | | | |
| High | O60281 | Zinc finger protein 292 OS=Homo sapiens OX=9606 GN=ZNF292 PE=1 SV=3 | 2 | 2 | 2 | 304.6 | 1 | 2723 | 7.39 | | | | | | | | | | | | |
| High | P63098 | Calcineurin subunit B type 1 OS=Homo sapiens OX=9606 GN=PPP3R1 PE=1 SV=2 | 2 | 1 | 1 | 19.3 | 5 | 170 | 4.81 | | | 1 | | | | | | 1 | | | |
| High | Q9NTZ6 | RNA-binding protein 12 OS=Homo sapiens OX=9606 GN=RBM12 PE=1 SV=1 | 2 | 1 | 1 | 97.3 | 1 | 932 | 8.63 | | | | | 1 | | | | | | 1 | |
| High | Q6UVK1 | Chondroitin sulfate proteoglycan 4 OS=Homo sapiens OX=9606 GN=CSPG4 PE=1 SV=2 | 2 | 1 | 1 | 250.4 | 1 | 2322 | 5.47 | 1 | | 1 | | | | 1 | | 1 | | | |
| High | Q6ZVM7 | TOM1-like protein 2 OS=Homo sapiens OX=9606 GN=TOM1L2 PE=1 SV=1 | 2 | 2 | 2 | 55.5 | 6 | 507 | 4.79 | | | 1 | | | | | | 1 | | | |
| High | P02794 | Ferritin heavy chain OS=Homo sapiens OX=9606 GN=FTH1 PE=1 SV=2 | 2 | 1 | 1 | 21.2 | 4 | 183 | 5.55 | | | | 1 | | 1 | | | | 1 | | 1 |
| High | Q13867 | Bleomycin hydrolase OS=Homo sapiens OX=9606 GN=BLMH PE=1 SV=1 | 2 | 2 | 2 | 52.5 | 5 | 455 | 6.27 | | | 2 | | | | | | 2 | | | |
| High | Q8TEX9 | Importin-4 OS=Homo sapiens OX=9606 GN=IPO4 PE=1 SV=2 | 2 | 1 | 1 | 118.6 | 2 | 1081 | 4.96 | | | | | | | | | | | | |
| High | Q9BTD8 | RNA-binding protein 42 OS=Homo sapiens OX=9606 GN=RBM42 PE=1 SV=1 | 2 | 1 | 1 | 50.4 | 4 | 480 | 9.63 | | | | | | | | | | | | |

| | | | | | | | | | | | | | | | | | | | | | |
|---|---|---|---|---|---|---|---|---|---|---|---|---|---|---|---|---|---|---|---|---|---|
| High | Q5VZ89 | DENN domain-containing protein 4C OS=Homo sapiens OX=9606 GN=DENND4C PE=1 SV=3 | 2 | 1 | 1 | 212.6 | 0 | 1909 | 6.83 | | | | | | | | | | | | |
| High | P55081 | Microfibrillar-associated protein 1 OS=Homo sapiens OX=9606 GN=MFAP1 PE=1 SV=2 | 2 | 1 | 1 | 51.9 | 2 | 439 | 4.98 | | | | 1 | | 1 | | | | 1 | | 1 |
| High | Q9UDY4 | DnaJ homolog subfamily B member 4 OS=Homo sapiens OX=9606 GN=DNAJB4 PE=1 SV=1 | 2 | 2 | 1 | 37.8 | 8 | 337 | 8.5 | | | | | | | | | | | | |
| High | P51617 | Interleukin-1 receptor-associated kinase 1 OS=Homo sapiens OX=9606 GN=IRAK1 PE=1 SV=2 | 2 | 1 | 1 | 76.5 | 2 | 712 | 6.62 | | 1 | 1 | | | | | 1 | 1 | | | |
| High | Q969Z0 | FAST kinase domain-containing protein 4 OS=Homo sapiens OX=9606 GN=TBRG4 PE=1 SV=1 | 2 | 2 | 2 | 70.7 | 4 | 631 | 7.42 | 1 | 1 | | | | | 1 | 1 | | | | |
| High | O75962 | Triple functional domain protein OS=Homo sapiens OX=9606 GN=TRIO PE=1 SV=2 | 2 | 1 | 1 | 346.7 | 0 | 3097 | 6.37 | | | | | 1 | | | | | | 1 | |
| High | O95602 | DNA-directed RNA polymerase I subunit RPA1 OS=Homo sapiens OX=9606 GN=POLR1A PE=1 SV=2 | 2 | 1 | 1 | 194.7 | 1 | 1720 | 7.03 | | | | | | | | | | | | |
| High | Q2TAY7 | WD40 repeat-containing protein SMU1 OS=Homo sapiens OX=9606 GN=SMU1 PE=1 SV=2 | 2 | 2 | 2 | 57.5 | 4 | 513 | 7.18 | | | | | 1 | | | | | | 1 | |
| High | P21817 | Ryanodine receptor 1 OS=Homo sapiens OX=9606 GN=RYR1 PE=1 SV=3 | 2 | 2 | 2 | 564.8 | 1 | 5038 | 5.3 | | 1 | 1 | | | | | 1 | 1 | | | |
| High | Q9BRG2 | SH2 domain-containing protein 3A OS=Homo sapiens OX=9606 GN=SH2D3A PE=1 SV=1 | 2 | 2 | 2 | 63.1 | 6 | 576 | 7.42 | | | 1 | | | | | | 1 | | | |
| High | Q03405 | Urokinase plasminogen activator surface receptor OS=Homo sapiens OX=9606 GN=PLAUR PE=1 SV=1 | 2 | 2 | 2 | 37 | 10 | 335 | 6.65 | | 1 | | | | | | 1 | | | | |
| High | Q9P0U3 | Sentrin-specific protease 1 OS=Homo sapiens OX=9606 GN=SENP1 PE=1 SV=2 | 2 | 1 | 1 | 73.4 | 1 | 644 | 8.47 | | | | | | | | | | | | |
| High | Q6UX65 | DNA damage-regulated autophagy modulator protein 2 OS=Homo sapiens OX=9606 GN=DRAM2 PE=1 SV=1 | 2 | 1 | 1 | 29.7 | 8 | 266 | 8.03 | 1 | 1 | | | | | 1 | 1 | | | | |
| High | Q16651 | Prostasin OS=Homo sapiens OX=9606 GN=PRSS8 PE=1 SV=1 | 2 | 1 | 1 | 36.4 | 4 | 343 | 5.85 | | | | | | | | | | | | |
| High | Q5BJD5 | Transmembrane protein 41B OS=Homo sapiens OX=9606 GN=TMEM41B PE=1 SV=1 | 2 | 1 | 1 | 32.5 | 7 | 291 | 9.58 | 2 | | | | | | 1 | | | | | |
| High | Q9Y291 | 28S ribosomal protein S33, mitochondrial OS=Homo sapiens OX=9606 GN=MRPS33 PE=1 SV=1 | 2 | 1 | 1 | 12.6 | 8 | 106 | 10.11 | | | | | | | | | | | | |
| High | C9JLW8 | Mapk-regulated corepressor-interacting protein 1 OS=Homo sapiens OX=9606 GN=MCRIP1 PE=1 SV=1 | 2 | 1 | 1 | 10.9 | 9 | 97 | 9.41 | | | | 1 | | 1 | | | | 1 | | 1 |
| High | Q9UH99 | SUN domain-containing protein 2 OS=Homo sapiens OX=9606 GN=SUN2 PE=1 SV=3 | 2 | 1 | 1 | 80.3 | 1 | 717 | 6.73 | | | 1 | | | | | | 1 | | | |
| High | Q15633 | RISC-loading complex subunit TARBP2 OS=Homo sapiens OX=9606 GN=TARBP2 PE=1 SV=3 | 2 | 1 | 1 | 39 | 4 | 366 | 6.54 | | | | | | | | | | | | |
| High | Q69YN4 | Protein virilizer homolog OS=Homo sapiens OX=9606 GN=VIRMA PE=1 SV=2 | 2 | 1 | 1 | 201.9 | 0 | 1812 | 5.01 | | | | | | | | | | | | |
| High | Q9UKJ3 | G patch domain-containing protein 8 OS=Homo sapiens OX=9606 GN=GPATCH8 PE=1 SV=2 | 2 | 1 | 1 | 164.1 | 1 | 1502 | 8.66 | | | | | | | | | | | | |
| High | P37287 | Phosphatidylinositol N-acetylglucosaminyltransferase subunit A OS=Homo sapiens OX=9606 GN=PIGA PE=1 SV=1 | 2 | 1 | 1 | 54.1 | 2 | 484 | 8.34 | | | | | | | | | | | | |
| High | Q99504 | Eyes absent homolog 3 OS=Homo sapiens OX=9606 GN=EYA3 PE=1 SV=3 | 2 | 1 | 1 | 62.6 | 2 | 573 | 5.21 | | | 1 | | | | | | 1 | | | |
| High | Q96JB5 | CDK5 regulatory subunit-associated protein 3 OS=Homo sapiens OX=9606 GN=CDK5RAP3 PE=1 SV=2 | 2 | 1 | 1 | 56.9 | 3 | 506 | 4.75 | | | | | | | | | | | | |
| High | Q14CX7 | N-alpha-acetyltransferase 25, NatB auxiliary subunit OS=Homo sapiens OX=9606 GN=NAA25 PE=1 SV=1 | 2 | 1 | 1 | 112.2 | 1 | 972 | 6.64 | | | 1 | | | | | | 1 | | | |
| High | Q15596 | Nuclear receptor coactivator 2 OS=Homo sapiens OX=9606 GN=NCOA2 PE=1 SV=2 | 2 | 2 | 2 | 159.1 | 3 | 1464 | 6.64 | | | | | | | | | | | | |
| High | Q9BUB7 | Transmembrane protein 70, mitochondrial OS=Homo sapiens OX=9606 GN=TMEM70 PE=1 SV=2 | 2 | 1 | 1 | 29 | 4 | 260 | 8.91 | 1 | 1 | | | | | 1 | 1 | | | | |
| High | Streptavid | (S.avidinii) | 2 | 1 | 1 | 16.6 | 12 | 160 | 6.55 | 1 | | | | 1 | | 1 | | | | 1 | |
| High | O94763 | Unconventional prefoldin RPB5 interactor 1 OS=Homo sapiens OX=9606 GN=URI1 PE=1 SV=3 | 2 | 2 | 2 | 59.8 | 8 | 535 | 5.05 | | | | | | | | | | | | |
| High | Q01469 | Fatty acid-binding protein 5 OS=Homo sapiens OX=9606 GN=FABP5 PE=1 SV=3 | 2 | 1 | 1 | 15.2 | 7 | 135 | 7.01 | | | | 1 | | 1 | | | | 1 | | 1 |
| High | Q86TB9 | Protein PAT1 homolog 1 OS=Homo sapiens OX=9606 GN=PATL1 PE=1 SV=2 | 2 | 2 | 2 | 86.8 | 2 | 770 | 6.67 | | | | | 1 | | | | | | 1 | |
| High | P09960 | Leukotriene A-4 hydrolase OS=Homo sapiens OX=9606 GN=LTA4H PE=1 SV=2 | 2 | 1 | 1 | 69.2 | 2 | 611 | 6.18 | | | | | | | | | | | | |
| High | Q9GZL7 | Ribosome biogenesis protein WDR12 OS=Homo sapiens OX=9606 GN=WDR12 PE=1 SV=2 | 2 | 1 | 1 | 47.7 | 2 | 423 | 5.9 | | | | | | | | | | | | |
| High | Q9H9P8 | L-2-hydroxyglutarate dehydrogenase, mitochondrial OS=Homo sapiens OX=9606 GN=L2HGDH PE=1 SV=3 | 2 | 1 | 1 | 50.3 | 2 | 463 | 8.15 | | | | | | | | | | | | |
| High | Q96C57 | Protein CUSTOS OS=Homo sapiens OX=9606 GN=CUSTOS PE=1 SV=2 | 2 | 1 | 1 | 28.2 | 7 | 262 | 9.42 | 1 | 1 | | | | | 1 | 1 | | | | |
| High | Q9H840 | Gem-associated protein 7 OS=Homo sapiens OX=9606 GN=GEMIN7 PE=1 SV=1 | 2 | 1 | 1 | 14.5 | 10 | 131 | 7.4 | | | | | | | | | | | | |
| High | O60232 | Protein ZNRD2 OS=Homo sapiens OX=9606 GN=ZNRD2 PE=1 SV=1 | 2 | 1 | 1 | 21.5 | 4 | 199 | 5.24 | | | | | | | | | | | | |
| High | Q53EZ4 | Centrosomal protein of 55 kDa OS=Homo sapiens OX=9606 GN=CEP55 PE=1 SV=3 | 2 | 2 | 2 | 54.1 | 5 | 464 | 7.01 | | | 1 | | | | | | 1 | | | |
| High | Q9HD47 | Ran guanine nucleotide release factor OS=Homo sapiens OX=9606 GN=RANGRF PE=1 SV=1 | 2 | 1 | 1 | 20.4 | 7 | 186 | 4.94 | | | | | | | | | | | | |
| High | Q9NVD7 | Alpha-parvin OS=Homo sapiens OX=9606 GN=PARVA PE=1 SV=1 | 2 | 1 | 1 | 42.2 | 4 | 372 | 5.95 | | | | | | | | | | | | |
| High | Q07065 | Cytoskeleton-associated protein 4 OS=Homo sapiens OX=9606 GN=CKAP4 PE=1 SV=2 | 2 | 1 | 1 | 66 | 3 | 602 | 5.92 | | | | | | | | | | | | |
| High | Q96NT0 | Coiled-coil domain-containing protein 115 OS=Homo sapiens OX=9606 GN=CCDC115 PE=1 SV=1 | 2 | 1 | 1 | 19.7 | 8 | 180 | 6.95 | | 1 | | | | | | 1 | | | | |
| High | Q8NFA0 | Ubiquitin carboxyl-terminal hydrolase 32 OS=Homo sapiens OX=9606 GN=USP32 PE=1 SV=1 | 2 | 1 | 1 | 181.5 | 1 | 1604 | 6.44 | | | 1 | | | | | | 1 | | | |
| High | A1KXE4 | Myelin-associated neurite-outgrowth inhibitor OS=Homo sapiens OX=9606 GN=FAM168B PE=1 SV=1 | 2 | 1 | 1 | 20.3 | 9 | 195 | 9.23 | | | | 1 | | 1 | | | | 1 | | 1 |
| High | P17275 | Transcription factor JunB OS=Homo sapiens OX=9606 GN=JUNB PE=1 SV=1 | 2 | 1 | 1 | 35.9 | 2 | 347 | 9.22 | | | | | | 1 | | | | | | 1 |
| High | P49354 | Protein farnesyltransferase/geranylgeranyltransferase type-1 subunit alpha OS=Homo sapiens OX=9606 GN=FNTA PE=1 SV=1 | 2 | 1 | 1 | 44.4 | 3 | 379 | 5.08 | | 1 | | | | | | 1 | | | | |
| High | P67812 | Signal peptidase complex catalytic subunit SEC11A OS=Homo sapiens OX=9606 GN=SEC11A PE=1 SV=1 | 2 | 1 | 1 | 20.6 | 7 | 179 | 9.48 | 1 | 1 | | | | | 1 | 1 | | | | |
| High | Q9H490 | Phosphatidylinositol glycan anchor biosynthesis class U protein OS=Homo sapiens OX=9606 GN=PIGU PE=1 SV=3 | 2 | 1 | 1 | 50 | 3 | 435 | 7.72 | | | | | | | | | | | | |
| High | P00711 | SWISS-PROT:P00711 Alpha-lactalbumin - Bos taurus (Bovine). | 2 | 1 | 1 | 14.2 | 14 | 123 | 5 | | | | | | 1 | | | | | | 1 |
| High | Q9BRQ6 | MICOS complex subunit MIC25 OS=Homo sapiens OX=9606 GN=CHCHD6 PE=1 SV=1 | 2 | 1 | 1 | 26.4 | 5 | 235 | 8.85 | 1 | | 1 | | | | 1 | | 1 | | | |
| High | Q9H8Y5 | Ankyrin repeat and zinc finger domain-containing protein 1 OS=Homo sapiens OX=9606 GN=ANKZF1 PE=1 SV=1 | 2 | 2 | 2 | 80.9 | 3 | 726 | 8.41 | | | | | | | | | | | | |
| High | Q9Y388 | RNA-binding motif protein, X-linked 2 OS=Homo sapiens OX=9606 GN=RBMX2 PE=1 SV=2 | 2 | 2 | 2 | 37.3 | 7 | 322 | 9.83 | | | | | 2 | | | | | | 2 | |
| High | P60604 | Ubiquitin-conjugating enzyme E2 G2 OS=Homo sapiens OX=9606 GN=UBE2G2 PE=1 SV=1 | 2 | 1 | 1 | 18.6 | 13 | 165 | 4.7 | 1 | 1 | | | | | 1 | 1 | | | | |
| High | Q7Z624 | Calmodulin-lysine N-methyltransferase OS=Homo sapiens OX=9606 GN=CAMKMT PE=1 SV=2 | 1 | 1 | 1 | 36.1 | 4 | 323 | 6.83 | | | | | | | | | | | | |
| High | P12757 | Ski-like protein OS=Homo sapiens OX=9606 GN=SKIL PE=1 SV=2 | 1 | 1 | 1 | 76.9 | 2 | 684 | 7.11 | | | | | | | | | | | | |
| High | Q86U28 | Iron-sulfur cluster assembly 2 homolog, mitochondrial OS=Homo sapiens OX=9606 GN=ISCA2 PE=1 SV=2 | 1 | 1 | 1 | 16.5 | 8 | 154 | 5.25 | | | | | | | | | | | | |
| High | Q9ULM3 | YEATS domain-containing protein 2 OS=Homo sapiens OX=9606 GN=YEATS2 PE=1 SV=2 | 1 | 1 | 1 | 150.7 | 1 | 1422 | 8.98 | | | | | | | | | | | | |
| High | O43166 | Signal-induced proliferation-associated 1-like protein 1 OS=Homo sapiens OX=9606 GN=SIPA1L1 PE=1 SV=4 | 1 | 1 | 1 | 199.9 | 0 | 1804 | 8.19 | | | | | | | | | | | | |
| High | Q86YV5 | Inactive tyrosine-protein kinase PRAG1 OS=Homo sapiens OX=9606 GN=PRAG1 PE=1 SV=4 | 1 | 1 | 1 | 149.5 | 1 | 1406 | 7.2 | | | | | | | | | | | | |
| High | P27695 | DNA-(apurinic or apyrimidinic site) endonuclease OS=Homo sapiens OX=9606 GN=APEX1 PE=1 SV=2 | 1 | 1 | 1 | 35.5 | 7 | 318 | 8.12 | | | | | 1 | | | | | | 1 | |
| High | Q7L5Y9 | E3 ubiquitin-protein transferase MAEA OS=Homo sapiens OX=9606 GN=MAEA PE=1 SV=1 | 1 | 1 | 1 | 45.3 | 2 | 396 | 8.69 | | | | | 1 | | | | | | 1 | |
| High | P15428 | 15-hydroxyprostaglandin dehydrogenase [NAD(+)] OS=Homo sapiens OX=9606 GN=HPGD PE=1 SV=1 | 1 | 1 | 1 | 29 | 3 | 266 | 5.86 | | | | | | | | | | | | |
| High | Q13561 | Dynactin subunit 2 OS=Homo sapiens OX=9606 GN=DCTN2 PE=1 SV=4 | 1 | 1 | 1 | 44.2 | 2 | 401 | 5.21 | | | | | | | | | | | | |
| High | Q9NZM1 | Myoferlin OS=Homo sapiens OX=9606 GN=MYOF PE=1 SV=1 | 1 | 1 | 1 | 234.6 | 1 | 2061 | 6.18 | | | | | | | | | | | | |
| High | Q9Y6R9 | Centrosomal protein CCDC61 OS=Homo sapiens OX=9606 GN=CCDC61 PE=1 SV=3 | 1 | 1 | 1 | 57.3 | 2 | 512 | 10.27 | | | | | | | | | | | | |
| High | Q9ULH1 | Arf-GAP with SH3 domain, ANK repeat and PH domain-containing protein 1 OS=Homo sapiens OX=9606 GN=ASAP1 PE=1 SV=4 | 1 | 1 | 1 | 125.4 | 2 | 1129 | 7.31 | | | | | | | | | | | | |
| High | Q9P0J1 | [Pyruvate dehydrogenase [acetyl-transferring]]-phosphatase 1, mitochondrial OS=Homo sapiens OX=9606 GN=PDP1 PE=1 SV=3 | 1 | 1 | 1 | 61 | 3 | 537 | 6.67 | | | 1 | | | | | | 1 | | | |
| High | Q9UKG1 | DCC-interacting protein 13-alpha OS=Homo sapiens OX=9606 GN=APPL1 PE=1 SV=1 | 1 | 1 | 1 | 79.6 | 2 | 709 | 5.41 | | 1 | | | | | | 1 | | | | |
| High | Q9NWV4 | CXXC motif containing zinc binding protein OS=Homo sapiens OX=9606 GN=CZIB PE=1 SV=1 | 1 | 1 | 1 | 18 | 11 | 160 | 5.01 | | | 1 | | | | | | 1 | | | |
| High | Q01196 | Runt-related transcription factor 1 OS=Homo sapiens OX=9606 GN=RUNX1 PE=1 SV=3 | 1 | 1 | 1 | 48.7 | 2 | 453 | 9.35 | | | 1 | | | | | | 1 | | | |
| High | P19387 | DNA-directed RNA polymerase II subunit RPB3 OS=Homo sapiens OX=9606 GN=POLR2C PE=1 SV=2 | 1 | 1 | 1 | 31.4 | 4 | 275 | 4.92 | | | | | | | | | | | | |
| High | Q5VWZ2 | Lysophospholipase-like protein 1 OS=Homo sapiens OX=9606 GN=LYPLAL1 PE=1 SV=3 | 1 | 1 | 1 | 26.3 | 4 | 237 | 7.84 | | | | | | | | | | | | |
| High | Q6WKZ4 | Rab11 family-interacting protein 1 OS=Homo sapiens OX=9606 GN=RAB11FIP1 PE=1 SV=3 | 1 | 1 | 1 | 137.1 | 2 | 1283 | 5.43 | | | | | | | | | | | | |
| High | Q3LXA3 | Triokinase/FMN cyclase OS=Homo sapiens OX=9606 GN=TKFC PE=1 SV=2 | 1 | 1 | 1 | 58.9 | 2 | 575 | 7.49 | | | | | | | | | | | | |
| High | Q5TH69 | Brefeldin A-inhibited guanine nucleotide-exchange protein 3 OS=Homo sapiens OX=9606 GN=ARFGEF3 PE=1 SV=3 | 1 | 1 | 1 | 240.5 | 0 | 2177 | 5.82 | | | 1 | | | | | | 1 | | | |
| High | Q03701 | CCAAT/enhancer-binding protein zeta OS=Homo sapiens OX=9606 GN=CEBPZ PE=1 SV=3 | 1 | 1 | 1 | 120.9 | 1 | 1054 | 5.94 | | | | | | | | | | | | |
| High | Q96L73 | Histone-lysine N-methyltransferase, H3 lysine-36 specific OS=Homo sapiens OX=9606 GN=NSD1 PE=1 SV=1 | 1 | 1 | 1 | 296.5 | 1 | 2696 | 8.03 | 1 | | | | | | 1 | | | | | |
| High | Q9H3N1 | Thioredoxin-related transmembrane protein 1 OS=Homo sapiens OX=9606 GN=TMX1 PE=1 SV=1 | 1 | 1 | 1 | 31.8 | 4 | 280 | 4.98 | | | | | | | | | | | | |
| High | Q460N5 | Protein mono-ADP-ribosyltransferase PARP14 OS=Homo sapiens OX=9606 GN=PARP14 PE=1 SV=3 | 1 | 1 | 1 | 202.7 | 1 | 1801 | 7.18 | 1 | | | | | | 1 | | | | | |
| High | P52565 | Rho GDP-dissociation inhibitor 1 OS=Homo sapiens OX=9606 GN=ARHGDIA PE=1 SV=3 | 1 | 1 | 1 | 23.2 | 8 | 204 | 5.11 | | | 1 | | | | | | 1 | | | |
| High | Q9H4L7 | SWI/SNF-related matrix-associated actin-dependent regulator of chromatin subfamily A containing DEAD/H box 1 OS=Homo sapie | 1 | 1 | 1 | 117.3 | 1 | 1026 | 5.55 | | | 1 | | | | | | 1 | | | |
| High | Q9H7X7 | Intraflagellar transport protein 22 homolog OS=Homo sapiens OX=9606 GN=IFT22 PE=1 SV=1 | 1 | 1 | 1 | 20.8 | 12 | 185 | 5.15 | | | | | | | | | | | | |
| High | P39019 | 40S ribosomal protein S19 OS=Homo sapiens OX=9606 GN=RPS19 PE=1 SV=2 | 1 | 1 | 1 | 16.1 | 9 | 145 | 10.32 | 1 | | | | | | 1 | | | | | |
| High | Q9H329 | Band 4.1-like protein 4B OS=Homo sapiens OX=9606 GN=EPB41L4B PE=2 SV=2 | 1 | 1 | 1 | 99.7 | 1 | 900 | 9.04 | | | | | | | | | | | | |
| High | Q86Y39 | NADH dehydrogenase [ubiquinone] 1 alpha subcomplex subunit 11 OS=Homo sapiens OX=9606 GN=NDUFA11 PE=1 SV=3 | 1 | 1 | 1 | 14.8 | 16 | 141 | 8.72 | | | 1 | | | | | | 1 | | | |
| High | Q9NR31 | GTP-binding protein SAR1a OS=Homo sapiens OX=9606 GN=SAR1A PE=1 SV=1 | 1 | 1 | 1 | 22.4 | 6 | 198 | 6.68 | | | | | | | | | | | | |
| High | Q8NCE0 | tRNA-splicing endonuclease subunit Sen2 OS=Homo sapiens OX=9606 GN=TSEN2 PE=1 SV=2 | 1 | 1 | 1 | 53.2 | 5 | 465 | 7.85 | | | | | | | | | | | | |
| High | Q96HU1 | Small G protein signaling modulator 3 OS=Homo sapiens OX=9606 GN=SGSM3 PE=1 SV=1 | 1 | 1 | 1 | 85.3 | 1 | 749 | 6 | | | | | | | | | | | | |
| High | P35030 | Trypsin-3 OS=Homo sapiens OX=9606 GN=PRSS3 PE=1 SV=2 | 1 | 1 | 1 | 32.5 | 4 | 304 | 7.49 | | | | | 1 | | | | | | 1 | |
| High | Q9H269 | Vacuolar protein sorting-associated protein 16 homolog OS=Homo sapiens OX=9606 GN=VPS16 PE=1 SV=2 | 1 | 1 | 1 | 94.6 | 1 | 839 | 6.77 | | | | | | | | | | | | |
| High | Q99973 | Telomerase protein component 1 OS=Homo sapiens OX=9606 GN=TEP1 PE=1 SV=2 | 1 | 1 | 1 | 290.3 | 1 | 2627 | 7.97 | | | | | | | | | | | | |
| High | Q15363 | Transmembrane emp24 domain-containing protein 2 OS=Homo sapiens OX=9606 GN=TMED2 PE=1 SV=1 | 1 | 1 | 1 | 22.7 | 8 | 201 | 5.17 | | | 1 | | | | | | 1 | | | |
| High | P48443 | Retinoic acid receptor RXR-gamma OS=Homo sapiens OX=9606 GN=RXRG PE=1 SV=1 | 1 | 1 | 1 | 50.8 | 2 | 463 | 7.62 | | | 1 | | | | | | 1 | | | |
| High | P18615 | Negative elongation factor E OS=Homo sapiens OX=9606 GN=NELFE PE=1 SV=3 | 1 | 1 | 1 | 43.2 | 6 | 380 | 9.33 | | | 1 | | | | | | 1 | | | |
| High | Q9BSF4 | Mitochondrial import inner membrane translocase subunit Tim29 OS=Homo sapiens OX=9606 GN=TIMM29 PE=1 SV=2 | 1 | 1 | 1 | 29.2 | 5 | 260 | 8.09 | | | 1 | | | | | | 1 | | | |
| High | Q14790 | Caspase-8 OS=Homo sapiens OX=9606 GN=CASP8 PE=1 SV=1 | 1 | 1 | 1 | 55.4 | 2 | 479 | 5.1 | 1 | | | | | | 1 | | | | | |
| High | O75052 | Carboxyl-terminal PDZ ligand of neuronal nitric oxide synthase protein OS=Homo sapiens OX=9606 GN=NOS1AP PE=1 SV=3 | 1 | 1 | 1 | 56.1 | 2 | 506 | 6.3 | | | | | | | | | | | | |
| High | Q13111 | Chromatin assembly factor 1 subunit A OS=Homo sapiens OX=9606 GN=CHAF1A PE=1 SV=3 | 1 | 1 | 1 | 106.8 | 2 | 956 | 5.94 | | | | | | | | | | | | |
| High | E9PRG8 | Uncharacterized protein C11orf98 OS=Homo sapiens OX=9606 GN=C11orf98 PE=4 SV=2 | 1 | 1 | 1 | 14.2 | 11 | 123 | 11.53 | | | | | | | | | | | | |
| High | P61086 | Ubiquitin-conjugating enzyme E2 K OS=Homo sapiens OX=9606 GN=UBE2K PE=1 SV=3 | 1 | 1 | 1 | 22.4 | 5 | 200 | 5.44 | | | | | 1 | | | | | | 1 | |
| High | Q15582 | Transforming growth factor-beta-induced protein ig-h3 OS=Homo sapiens OX=9606 GN=TGFBI PE=1 SV=1 | 1 | 1 | 1 | 74.6 | 3 | 683 | 7.71 | | | | | 1 | | | | | | 1 | |
| High | A0AVI4 | E3 ubiquitin-protein ligase TM129 OS=Homo sapiens OX=9606 GN=TMEM129 PE=1 SV=1 | 1 | 1 | 1 | 40.4 | 5 | 362 | 7.66 | | | 1 | | | | | | 1 | | | |
| High | Q9H270 | Vacuolar protein sorting-associated protein 11 homolog OS=Homo sapiens OX=9606 GN=VPS11 PE=1 SV=1 | 1 | 1 | 1 | 107.8 | 3 | 941 | 7.05 | 1 | | | | | | 1 | | | | | |
| High | P34955 | SWISS-PROT:P34955 (Bos taurus) Alpha-1-antiproteinase precursor | 1 | 1 | 1 | 46.1 | 2 | 416 | 6.52 | | | | | 1 | | | | | | 1 | |
| High | Q86W56 | Poly(ADP-ribose) glycohydrolase OS=Homo sapiens OX=9606 GN=PARG PE=1 SV=1 | 1 | 1 | 1 | 111 | 1 | 976 | 6.43 | | | | | | | | | | | | |
| High | P57772 | Selenocysteine-specific elongation factor OS=Homo sapiens OX=9606 GN=EEFSEC PE=1 SV=4 | 1 | 1 | 1 | 65.3 | 2 | 596 | 8.35 | | | | | | | | | | | | |
| High | Q9NZD8 | Maspardin OS=Homo sapiens OX=9606 GN=SPG21 PE=1 SV=1 | 1 | 1 | 1 | 34.9 | 6 | 308 | 6.28 | | | | | | | | | | | | |
| High | Q96JC9 | ELL-associated factor 1 OS=Homo sapiens OX=9606 GN=EAF1 PE=1 SV=1 | 1 | 1 | 1 | 29 | 7 | 268 | 5.4 | | | | | | | | | | | | |
| High | Q5TA45 | Integrator complex subunit 11 OS=Homo sapiens OX=9606 GN=INTS11 PE=1 SV=2 | 1 | 1 | 1 | 67.6 | 2 | 600 | 8.06 | | | 1 | | | | | | 1 | | | |
| High | Q5TCQ9 | Membrane-associated guanylate kinase, WW and PDZ domain-containing protein 3 OS=Homo sapiens OX=9606 GN=MAGI3 PE=1 | 1 | 1 | 1 | 162.8 | 1 | 1481 | 8.09 | | | | | | | | | | | | |
| High | Q9NVR5 | Protein kintoun OS=Homo sapiens OX=9606 GN=DNAAF2 PE=1 SV=2 | 1 | 1 | 1 | 91.1 | 2 | 837 | 5.2 | | | | | | | | | | | | |
| High | P17813 | Endoglin OS=Homo sapiens OX=9606 GN=ENG PE=1 SV=2 | 1 | 1 | 1 | 70.5 | 3 | 658 | 6.61 | | | | | | | | | | | | |
| High | O75330 | Hyaluronan mediated motility receptor OS=Homo sapiens OX=9606 GN=HMMR PE=1 SV=2 | 1 | 1 | 1 | 84 | 2 | 724 | 5.83 | | | | | | | | | | | | |
| High | P07199 | Major centromere autoantigen B OS=Homo sapiens OX=9606 GN=CENPB PE=1 SV=2 | 1 | 1 | 1 | 65.1 | 2 | 599 | 4.55 | | | 1 | | | | | | 1 | | | |
| High | Q9BZI7 | Regulator of nonsense transcripts 3B OS=Homo sapiens OX=9606 GN=UPF3B PE=1 SV=1 | 1 | 1 | 1 | 57.7 | 2 | 483 | 9.48 | | | 1 | | | | | | 1 | | | |
| High | A7KAX9 | Rho GTPase-activating protein 32 OS=Homo sapiens OX=9606 GN=ARHGAP32 PE=1 SV=1 | 1 | 1 | 1 | 230.4 | 0 | 2087 | 6.74 | | | | | | | | | | | | |
| High | Q9BVI4 | Nucleolar complex protein 4 homolog OS=Homo sapiens OX=9606 GN=NOC4L PE=1 SV=1 | 1 | 1 | 1 | 58.4 | 2 | 516 | 7.49 | | | | | | | | | | | | |
| High | Q86SK9 | Stearoyl-CoA desaturase 5 OS=Homo sapiens OX=9606 GN=SCD5 PE=1 SV=2 | 1 | 1 | 1 | 37.6 | 5 | 330 | 9.61 | 1 | | | | | | 1 | | | | | |
| High | P08047 | Transcription factor Sp1 OS=Homo sapiens OX=9606 GN=SP1 PE=1 SV=3 | 1 | 1 | 1 | 80.6 | 4 | 785 | 7.34 | | | | | | | | | | | | |
| High | O15541 | E3 ubiquitin-protein ligase RNF113A OS=Homo sapiens OX=9606 GN=RNF113A PE=1 SV=1 | 1 | 1 | 1 | 38.8 | 3 | 343 | 5.69 | | | | | | | | | | | | |
| High | Q8N511 | Transmembrane protein 199 OS=Homo sapiens OX=9606 GN=TMEM199 PE=1 SV=1 | 1 | 1 | 1 | 23.1 | 4 | 208 | 9.01 | 1 | | | | | | 1 | | | | | |
| High | O95834 | Echinoderm microtubule-associated protein-like 2 OS=Homo sapiens OX=9606 GN=EML2 PE=1 SV=1 | 1 | 1 | 1 | 70.6 | 2 | 649 | 6.32 | | | | | | | | | | | | |
| High | P63316 | Troponin C, slow skeletal and cardiac muscles OS=Homo sapiens OX=9606 GN=TNNC1 PE=1 SV=1 | 1 | 1 | 1 | 18.4 | 11 | 161 | 4.18 | | | | | 1 | | | | | | 1 | |
| High | Q9ULR3 | Protein phosphatase 1H OS=Homo sapiens OX=9606 GN=PPM1H PE=1 SV=2 | 1 | 1 | 1 | 56.4 | 3 | 514 | 6.6 | | | | | | | | | | | | |
| High | P40855 | Peroxisomal biogenesis factor 19 OS=Homo sapiens OX=9606 GN=PEX19 PE=1 SV=1 | 1 | 1 | 1 | 32.8 | 5 | 299 | 4.34 | | | | | | | | | | | | |
| High | Q9Y4B6 | DDB1- and CUL4-associated factor 1 OS=Homo sapiens OX=9606 GN=DCAF1 PE=1 SV=3 | 1 | 1 | 1 | 168.9 | 1 | 1507 | 5.06 | | | | | | | | | | | | |
| High | Q9BZA7 | Protocadherin-11 X-linked OS=Homo sapiens OX=9606 GN=PCDH11X PE=2 SV=1 | 1 | 1 | 1 | 147.5 | 1 | 1347 | 5.21 | | | | | | | | | | | | |
| High | P15104 | Glutamine synthetase OS=Homo sapiens OX=9606 GN=GLUL PE=1 SV=4 | 1 | 1 | 1 | 42 | 5 | 373 | 6.89 | | | 1 | | | | | | 1 | | | |
| High | Q9H7E2 | Tudor domain-containing protein 3 OS=Homo sapiens OX=9606 GN=TDRD3 PE=1 SV=1 | 1 | 1 | 1 | 73.1 | 2 | 651 | 9.23 | | | | | | | | | | | | |
| High | P49821 | NADH dehydrogenase [ubiquinone] flavoprotein 1, mitochondrial OS=Homo sapiens OX=9606 GN=NDUFV1 PE=1 SV=4 | 1 | 1 | 1 | 50.8 | 2 | 464 | 8.21 | | | | | | | | | | | | |
| High | P29372 | DNA-3-methyladenine glycosylase OS=Homo sapiens OX=9606 GN=MPG PE=1 SV=3 | 1 | 1 | 1 | 32.8 | 4 | 298 | 9.57 | 1 | | | | | | 1 | | | | | |
| High | Q12962 | Transcription initiation factor TFIID subunit 10 OS=Homo sapiens OX=9606 GN=TAF10 PE=1 SV=1 | 1 | 1 | 1 | 21.7 | 4 | 218 | 6.57 | | | | | | | | | | | | |
| High | Q14197 | Peptidyl-tRNA hydrolase ICT1, mitochondrial OS=Homo sapiens OX=9606 GN=MRPL58 PE=1 SV=1 | 1 | 1 | 1 | 23.6 | 5 | 206 | 10.07 | | | | | | | | | | | | |
| High | O43572 | A-kinase anchor protein 10, mitochondrial OS=Homo sapiens OX=9606 GN=AKAP10 PE=1 SV=2 | 1 | 1 | 1 | 73.8 | 3 | 662 | 6.39 | | | 1 | | | | | | 1 | | | |
| High | Q6AI12 | Ankyrin repeat domain-containing protein 40 OS=Homo sapiens OX=9606 GN=ANKRD40 PE=1 SV=2 | 1 | 1 | 1 | 41.1 | 5 | 368 | 4.97 | | | | | | | | | | | | |
| High | Q9UGK8 | Secretion-regulating guanine nucleotide exchange factor OS=Homo sapiens OX=9606 GN=SERGEF PE=1 SV=2 | 1 | 1 | 1 | 49 | 2 | 458 | 6.46 | | | | | | | | | | | | |
| High | Q9H8H0 | Nucleolar protein 11 OS=Homo sapiens OX=9606 GN=NOL11 PE=1 SV=1 | 1 | 1 | 1 | 81.1 | 1 | 719 | 6.07 | | | | | | | | | | | | |
| High | P42336 | Phosphatidylinositol 4,5-bisphosphate 3-kinase catalytic subunit alpha isoform OS=Homo sapiens OX=9606 GN=PIK3CA PE=1 SV=2 | 1 | 1 | 1 | 124.2 | 2 | 1068 | 7.23 | | | | | | | | | | | | |
| High | Q8TAD8 | Smad nuclear-interacting protein 1 OS=Homo sapiens OX=9606 GN=SNIP1 PE=1 SV=1 | 1 | 1 | 1 | 45.8 | 4 | 396 | 9.99 | | | | | | | | | | | | |
| High | Q15061 | WD repeat-containing protein 43 OS=Homo sapiens OX=9606 GN=WDR43 PE=1 SV=3 | 1 | 1 | 1 | 74.8 | 4 | 677 | 5.57 | | | | | | | | | | | | |
| High | Q684P5 | Rap1 GTPase-activating protein 2 OS=Homo sapiens OX=9606 GN=RAP1GAP2 PE=1 SV=2 | 1 | 1 | 1 | 80 | 1 | 730 | 6.67 | | | 1 | | | | | | 1 | | | |
| High | O95359 | Transforming acidic coiled-coil-containing protein 2 OS=Homo sapiens OX=9606 GN=TACC2 PE=1 SV=3 | 1 | 1 | 1 | 309.2 | 0 | 2948 | 4.79 | | | | | | | | | | | | |
| High | Q5QJE6 | Deoxynucleotidyltransferase terminal-interacting protein 2 OS=Homo sapiens OX=9606 GN=DNTTIP2 PE=1 SV=2 | 1 | 1 | 1 | 84.4 | 2 | 756 | 6.16 | | | 1 | | | | | | 1 | | | |
| High | O00189 | AP-4 complex subunit mu-1 OS=Homo sapiens OX=9606 GN=AP4M1 PE=1 SV=2 | 1 | 1 | 1 | 49.9 | 3 | 453 | 7.2 | | | 1 | | | | | | 1 | | | |
| High | Q8TBE9 | N-acylneuraminate-9-phosphatase OS=Homo sapiens OX=9606 GN=NANP PE=1 SV=1 | 1 | 1 | 1 | 27.8 | 5 | 248 | 6.4 | | | | | | | | | | | | |
| High | Q9H0N5 | Pterin-4-alpha-carbinolamine dehydratase 2 OS=Homo sapiens OX=9606 GN=PCBD2 PE=1 SV=4 | 1 | 1 | 1 | 14.4 | 12 | 130 | 9.11 | 1 | | | | | | 1 | | | | | |
| High | Q9NWU1 | 3-oxoacyl-[acyl-carrier-protein] synthase, mitochondrial OS=Homo sapiens OX=9606 GN=OXSM PE=1 SV=1 | 1 | 1 | 1 | 48.8 | 3 | 459 | 7.66 | | | | | | | | | | | | |
| High | O43678 | NADH dehydrogenase [ubiquinone] 1 alpha subcomplex subunit 2 OS=Homo sapiens OX=9606 GN=NDUFA2 PE=1 SV=3 | 1 | 1 | 1 | 10.9 | 21 | 99 | 9.57 | | | | | | | | | | | | |
| High | O60783 | 28S ribosomal protein S14, mitochondrial OS=Homo sapiens OX=9606 GN=MRPS14 PE=1 SV=1 | 1 | 1 | 1 | 15.1 | 12 | 128 | 11.41 | | | | | | | | | | | | |
| High | Q9BW30 | Tubulin polymerization-promoting protein family member 3 OS=Homo sapiens OX=9606 GN=TPPP3 PE=1 SV=1 | 1 | 1 | 1 | 19 | 5 | 176 | 9.13 | | | | | 1 | | | | | | 1 | |
| High | Q8TD19 | Serine/threonine-protein kinase Nek9 OS=Homo sapiens OX=9606 GN=NEK9 PE=1 SV=2 | 1 | 1 | 1 | 107.1 | 1 | 979 | 5.74 | | | | | | | | | | | | |
| High | Q5VV41 | Rho guanine nucleotide exchange factor 16 OS=Homo sapiens OX=9606 GN=ARHGEF16 PE=1 SV=1 | 1 | 1 | 1 | 80.1 | 1 | 709 | 7.36 | | | 1 | | | | | | 1 | | | |
| High | P10619 | Lysosomal protective protein OS=Homo sapiens OX=9606 GN=CTSA PE=1 SV=2 | 1 | 1 | 1 | 54.4 | 3 | 480 | 6.61 | | | | | 1 | | | | | | 1 | |
| High | Q9BT25 | HAUS augmin-like complex subunit 8 OS=Homo sapiens OX=9606 GN=HAUS8 PE=1 SV=3 | 1 | 1 | 1 | 44.8 | 2 | 410 | 7.06 | | | 1 | | | | | | 1 | | | |
| High | Q969K7 | Transmembrane protein 54 OS=Homo sapiens OX=9606 GN=TMEM54 PE=1 SV=1 | 1 | 1 | 1 | 23.8 | 4 | 222 | 7.83 | | | | | | | | | | | | |
| High | Q9ULQ0 | Striatin-interacting protein 2 OS=Homo sapiens OX=9606 GN=STRIP2 PE=1 SV=2 | 1 | 1 | 1 | 95.3 | 1 | 834 | 5.91 | | | | | | | | | | | | |
| High | Q8IY22 | C-Maf-inducing protein OS=Homo sapiens OX=9606 GN=CMIP PE=1 SV=3 | 1 | 1 | 1 | 86.3 | 3 | 773 | 6.7 | | | | | | | | | | | | |
| High | P35869 | Aryl hydrocarbon receptor OS=Homo sapiens OX=9606 GN=AHR PE=1 SV=2 | 1 | 1 | 1 | 96.1 | 1 | 848 | 6.38 | | | | | | | | | | | | |
| High | Q8IXH7 | Negative elongation factor C/D OS=Homo sapiens OX=9606 GN=NELFCD PE=1 SV=2 | 1 | 1 | 1 | 66.2 | 2 | 590 | 5.1 | | | | | | | | | | | | |
| High | Q8IYL3 | UPF0688 protein C1orf174 OS=Homo sapiens OX=9606 GN=C1orf174 PE=1 SV=2 | 1 | 1 | 1 | 26 | 6 | 243 | 6.9 | | | | | | | | | | | | |
| High | O95396 | Adenylyltransferase and sulfurtransferase MOCS3 OS=Homo sapiens OX=9606 GN=MOCS3 PE=1 SV=1 | 1 | 1 | 1 | 49.6 | 4 | 460 | 6.21 | | 1 | | | | | | 1 | | | | |
| High | Q9UPN4 | Centrosomal protein of 131 kDa OS=Homo sapiens OX=9606 GN=CEP131 PE=1 SV=3 | 1 | 1 | 1 | 122.1 | 1 | 1083 | 8.69 | | | | | | | | | | | | |
| High | P09913 | Interferon-induced protein with tetratricopeptide repeats 2 OS=Homo sapiens OX=9606 GN=IFIT2 PE=1 SV=1 | 1 | 1 | 1 | 54.6 | 2 | 472 | 6.76 | | | | | 1 | | | | | | 1 | |
| High | Q9Y2H0 | Disks large-associated protein 4 OS=Homo sapiens OX=9606 GN=DLGAP4 PE=1 SV=3 | 1 | 1 | 1 | 107.9 | 1 | 992 | 7.08 | | | | | | | | | | | | |
| High | Q8NFU3 | Thiosulfate:glutathione sulfurtransferase OS=Homo sapiens OX=9606 GN=TSTD1 PE=1 SV=3 | 1 | 1 | 1 | 12.5 | 12 | 115 | 6.07 | | | | | | | | | | | | |
| High | Q96LB3 | Intraflagellar transport protein 74 homolog OS=Homo sapiens OX=9606 GN=IFT74 PE=1 SV=1 | 1 | 1 | 1 | 69.2 | 3 | 600 | 6 | | 1 | | | | | | 1 | | | | |
| High | Q9NW08 | DNA-directed RNA polymerase III subunit RPC2 OS=Homo sapiens OX=9606 GN=POLR3B PE=1 SV=2 | 1 | 1 | 1 | 127.7 | 1 | 1133 | 8.5 | | | | | 1 | | | | | | 1 | |
| High | O95478 | Ribosome biogenesis protein NSA2 homolog OS=Homo sapiens OX=9606 GN=NSA2 PE=1 SV=1 | 1 | 1 | 1 | 30 | 4 | 260 | 10.27 | | | | | | | | | | | | |
| High | Q6PJT7 | Zinc finger CCCH domain-containing protein 14 OS=Homo sapiens OX=9606 GN=ZC3H14 PE=1 SV=1 | 1 | 1 | 1 | 82.8 | 1 | 736 | 7.31 | | | | | | | | | | | | |

| | | | | | | | | | | | | | | | | | | | | | |
|---|---|---|---|---|---|---|---|---|---|---|---|---|---|---|---|---|---|---|---|---|---|
| High | O00165 | HCLS1-associated protein X-1 OS=Homo sapiens OX=9606 GN=HAX1 PE=1 SV=2 | 1 | 1 | 1 | 31.6 | 4 | 279 | 4.92 | | | 1 | | | | | | 1 | | | |
| High | Q9HCN4 | GPN-loop GTPase 1 OS=Homo sapiens OX=9606 GN=GPN1 PE=1 SV=1 | 1 | 1 | 1 | 41.7 | 4 | 374 | 4.92 | | | 1 | | | | | | 1 | | | |
| High | Q7Z2K6 | Endoplasmic reticulum metallopeptidase 1 OS=Homo sapiens OX=9606 GN=ERMP1 PE=1 SV=2 | 1 | 1 | 1 | 100.2 | 1 | 904 | 7.52 | | | 1 | | | | | | 1 | | | |
| High | Q9H3R2 | Mucin-13 OS=Homo sapiens OX=9606 GN=MUC13 PE=1 SV=3 | 1 | 1 | 1 | 54.6 | 4 | 512 | 5.07 | | | | | | | | | | | | |
| High | Q12874 | Splicing factor 3A subunit 3 OS=Homo sapiens OX=9606 GN=SF3A3 PE=1 SV=1 | 1 | 1 | 1 | 58.8 | 2 | 501 | 5.38 | | | | | | | | | | | | |
| High | O14880 | Microsomal glutathione S-transferase 3 OS=Homo sapiens OX=9606 GN=MGST3 PE=1 SV=1 | 1 | 1 | 1 | 16.5 | 14 | 152 | 9.38 | | | | | | | | | | | | |
| High | Q92835 | Phosphatidylinositol 3,4,5-trisphosphate 5-phosphatase 1 OS=Homo sapiens OX=9606 GN=INPP5D PE=1 SV=2 | 1 | 1 | 1 | 133.2 | 1 | 1189 | 7.59 | | | | | | | | | | | | |
| High | Q96MC6 | Hippocampus abundant transcript 1 protein OS=Homo sapiens OX=9606 GN=MFSD14A PE=1 SV=2 | 1 | 1 | 1 | 53 | 4 | 490 | 8.4 | 1 | | | | | | 1 | | | | | |
| High | O00400 | Acetyl-coenzyme A transporter 1 OS=Homo sapiens OX=9606 GN=SLC33A1 PE=1 SV=1 | 1 | 1 | 1 | 60.9 | 2 | 549 | 7.33 | | | | | | | | | | | | |
| High | Q96AZ6 | Interferon-stimulated gene 20 kDa protein OS=Homo sapiens OX=9606 GN=ISG20 PE=1 SV=2 | 1 | 1 | 1 | 20.4 | 9 | 181 | 8.92 | | | | | | | | | | | | |
| High | Q9NUQ2 | 1-acyl-sn-glycerol-3-phosphate acyltransferase epsilon OS=Homo sapiens OX=9606 GN=AGPAT5 PE=1 SV=3 | 1 | 1 | 1 | 42 | 6 | 364 | 9.1 | | | | | | | | | | | | |
| High | O60238 | BCL2/adenovirus E1B 19 kDa protein-interacting protein 3-like OS=Homo sapiens OX=9606 GN=BNIP3L PE=1 SV=1 | 1 | 1 | 1 | 23.9 | 7 | 219 | 5.85 | | | 1 | | | | | | 1 | | | |
| High | O75431 | Metaxin-2 OS=Homo sapiens OX=9606 GN=MTX2 PE=1 SV=1 | 1 | 1 | 1 | 29.7 | 4 | 263 | 6.29 | | | | | | 1 | | | | | | 1 |
| High | Q8NFV4 | Protein ABHD11 OS=Homo sapiens OX=9606 GN=ABHD11 PE=1 SV=1 | 1 | 1 | 1 | 34.7 | 5 | 315 | 9.48 | | | 1 | | | | | | 1 | | | |
| High | P31749 | RAC-alpha serine/threonine-protein kinase OS=Homo sapiens OX=9606 GN=AKT1 PE=1 SV=2 | 1 | 1 | 1 | 55.7 | 3 | 480 | 6.07 | | | | | | | | | | | | |
| High | O14949 | Cytochrome b-c1 complex subunit 8 OS=Homo sapiens OX=9606 GN=UQCRQ PE=1 SV=4 | 1 | 1 | 1 | 9.9 | 16 | 82 | 10.08 | | 1 | | | | | | 1 | | | | |
| High | Q6PJ69 | Tripartite motif-containing protein 65 OS=Homo sapiens OX=9606 GN=TRIM65 PE=1 SV=3 | 1 | 1 | 1 | 57.3 | 3 | 517 | 6.7 | | | 1 | | | | | | 1 | | | |
| High | O75817 | Ribonuclease P protein subunit p20 OS=Homo sapiens OX=9606 GN=POP7 PE=1 SV=2 | 1 | 1 | 1 | 15.6 | 11 | 140 | 8.94 | | | | | | | | | | | | |
| High | Q9NUD5 | Zinc finger CCHC domain-containing protein 3 OS=Homo sapiens OX=9606 GN=ZCCHC3 PE=1 SV=2 | 1 | 1 | 1 | 43.5 | 4 | 403 | 8.53 | | | | | | | | | | | | |
| High | Q96EH3 | Mitochondrial assembly of ribosomal large subunit protein 1 OS=Homo sapiens OX=9606 GN=MALSU1 PE=1 SV=1 | 1 | 1 | 1 | 26.2 | 9 | 234 | 5.49 | | | | | | | | | | | | |
| High | Q9BTL3 | RNA guanine-N7 methyltransferase activating subunit OS=Homo sapiens OX=9606 GN=RAMAC PE=1 SV=1 | 1 | 1 | 1 | 14.4 | 11 | 118 | 8.94 | | | 1 | | | | | | 1 | | | |
| High | Q9BSR8 | Protein YIPF4 OS=Homo sapiens OX=9606 GN=YIPF4 PE=1 SV=1 | 1 | 1 | 1 | 27.1 | 4 | 244 | 4.65 | | | 1 | | | | | | 1 | | | |
| High | Q14139 | Ubiquitin conjugation factor E4 A OS=Homo sapiens OX=9606 GN=UBE4A PE=1 SV=2 | 1 | 1 | 1 | 122.5 | 1 | 1066 | 5.24 | | | | | | | | | | | | |
| High | Q8TBF4 | Zinc finger CCHC-type and RNA-binding motif-containing protein 1 OS=Homo sapiens OX=9606 GN=ZCRB1 PE=1 SV=2 | 1 | 1 | 1 | 24.6 | 7 | 217 | 8.53 | | | | | | | | | | | | |
| High | Q96QT4 | Transient receptor potential cation channel subfamily M member 7 OS=Homo sapiens OX=9606 GN=TRPM7 PE=1 SV=1 | 1 | 1 | 1 | 212.6 | 1 | 1865 | 7.88 | | | | | | | | | | | | |
| High | Q9NRY4 | Rho GTPase-activating protein 35 OS=Homo sapiens OX=9606 GN=ARHGAP35 PE=1 SV=3 | 1 | 1 | 1 | 170.4 | 1 | 1499 | 6.64 | | | | | | | | | | | | |
| High | P08243 | Asparagine synthetase [glutamine-hydrolyzing] OS=Homo sapiens OX=9606 GN=ASNS PE=1 SV=4 | 1 | 1 | 1 | 64.3 | 2 | 561 | 6.86 | | | 1 | | | | | | 1 | | | |
| High | Q9NV70 | Exocyst complex component 1 OS=Homo sapiens OX=9606 GN=EXOC1 PE=1 SV=4 | 1 | 1 | 1 | 101.9 | 1 | 894 | 6.61 | | | | | | | | | | | | |
| High | Q96EK5 | KIF-binding protein OS=Homo sapiens OX=9606 GN=KIFBP PE=1 SV=1 | 1 | 1 | 1 | 71.8 | 2 | 621 | 5.49 | 1 | | | | | | 1 | | | | | |
| High | P31937 | 3-hydroxyisobutyrate dehydrogenase, mitochondrial OS=Homo sapiens OX=9606 GN=HIBADH PE=1 SV=2 | 1 | 1 | 1 | 35.3 | 4 | 336 | 8.13 | | | 1 | | | | | | 1 | | | |
| High | Q99102 | Mucin-4 OS=Homo sapiens OX=9606 GN=MUC4 PE=1 SV=5 | 1 | 1 | 1 | 542 | 1 | 5412 | 5.3 | | | | | | | | | | | | |
| High | Q99470 | Stromal cell-derived factor 2 OS=Homo sapiens OX=9606 GN=SDF2 PE=1 SV=2 | 1 | 1 | 1 | 23 | 7 | 211 | 7.33 | | 1 | | | | | | 1 | | | | |
| High | Q9H0H0 | Integrator complex subunit 2 OS=Homo sapiens OX=9606 GN=INTS2 PE=1 SV=2 | 1 | 1 | 1 | 134.2 | 1 | 1204 | 6.05 | | | 1 | | | | | | 1 | | | |
| High | Q9BXS4 | Transmembrane protein 59 OS=Homo sapiens OX=9606 GN=TMEM59 PE=1 SV=1 | 1 | 1 | 1 | 36.2 | 4 | 323 | 5.1 | | | | | | | | | | | | |
| High | Q5T8D3 | Acyl-CoA-binding domain-containing protein 5 OS=Homo sapiens OX=9606 GN=ACBD5 PE=1 SV=1 | 1 | 1 | 1 | 60.1 | 3 | 534 | 5.33 | | | | | | | | | | | | |
| High | Q9BSM1 | Polycomb group RING finger protein 1 OS=Homo sapiens OX=9606 GN=PCGF1 PE=1 SV=2 | 1 | 1 | 1 | 30.3 | 4 | 259 | 8.91 | | | | | 1 | | | | | | 1 | |
| High | Q12974 | Protein tyrosine phosphatase type IVA 2 OS=Homo sapiens OX=9606 GN=PTP4A2 PE=1 SV=1 | 1 | 1 | 1 | 19.1 | 8 | 167 | 8.37 | | 1 | | | | | | 1 | | | | |
| High | Q9NTJ5 | Phosphatidylinositol-3-phosphatase SAC1 OS=Homo sapiens OX=9606 GN=SACM1L PE=1 SV=2 | 1 | 1 | 1 | 66.9 | 2 | 587 | 7.12 | | | 1 | | | | | | 1 | | | |
| High | Q6UWP8 | Suprabasin OS=Homo sapiens OX=9606 GN=SBSN PE=1 SV=2 | 1 | 1 | 1 | 60.5 | 3 | 590 | 7.01 | | | | | 1 | | | | | | 1 | |
| High | Q6ZRP7 | Sulfhydryl oxidase 2 OS=Homo sapiens OX=9606 GN=QSOX2 PE=1 SV=3 | 1 | 1 | 1 | 77.5 | 2 | 698 | 7.72 | | | | | | | | | | | | |
| High | Q15833 | Syntaxin-binding protein 2 OS=Homo sapiens OX=9606 GN=STXBP2 PE=1 SV=2 | 1 | 1 | 1 | 66.4 | 3 | 593 | 6.55 | | 1 | | | | | | 1 | | | | |
| High | Q9NSP4 | Centromere protein M OS=Homo sapiens OX=9606 GN=CENPM PE=1 SV=1 | 1 | 1 | 1 | 19.7 | 6 | 180 | 7.2 | | | | | | 1 | | | | | | 1 |
| High | Q9Y3T9 | Nucleolar complex protein 2 homolog OS=Homo sapiens OX=9606 GN=NOC2L PE=1 SV=4 | 1 | 1 | 1 | 84.9 | 1 | 749 | 5.62 | | | | | | | | | | | | |
| High | Q8TAP8 | Protein phosphatase 1 regulatory subunit 35 OS=Homo sapiens OX=9606 GN=PPP1R35 PE=1 SV=1 | 1 | 1 | 1 | 27.9 | 9 | 253 | 8.68 | | | | | | | | | | | | |
| High | P17612 | cAMP-dependent protein kinase catalytic subunit alpha OS=Homo sapiens OX=9606 GN=PRKACA PE=1 SV=2 | 1 | 1 | 1 | 40.6 | 5 | 351 | 8.79 | | | | | | | | | | | | |
| High | P06756 | Integrin alpha-V OS=Homo sapiens OX=9606 GN=ITGAV PE=1 SV=2 | 1 | 1 | 1 | 116 | 2 | 1048 | 5.68 | | | | | | | | | | | | |
| High | Q9H1K1 | Iron-sulfur cluster assembly enzyme ISCU, mitochondrial OS=Homo sapiens OX=9606 GN=ISCU PE=1 SV=2 | 1 | 1 | 1 | 18 | 10 | 167 | 9.48 | | | | | | | | | | | | |
| High | P13498 | Cytochrome b-245 light chain OS=Homo sapiens OX=9606 GN=CYBA PE=1 SV=3 | 1 | 1 | 1 | 21 | 4 | 195 | 9.54 | | | | | | | | | | | | |
| High | Q9Y6A4 | Cilia- and flagella-associated protein 20 OS=Homo sapiens OX=9606 GN=CFAP20 PE=1 SV=1 | 1 | 1 | 1 | 22.8 | 4 | 193 | 9.76 | | | | | | | | | | | | |
| High | Q6I9Y2 | THO complex subunit 7 homolog OS=Homo sapiens OX=9606 GN=THOC7 PE=1 SV=3 | 1 | 1 | 1 | 23.7 | 7 | 204 | 5.67 | | | | | | | | | | | | |
| High | Q9HC07 | Transmembrane protein 165 OS=Homo sapiens OX=9606 GN=TMEM165 PE=1 SV=1 | 1 | 1 | 1 | 34.9 | 8 | 324 | 7.02 | | | 1 | | | | | | 1 | | | |
| High | Q14689 | Disco-interacting protein 2 homolog A OS=Homo sapiens OX=9606 GN=DIP2A PE=1 SV=2 | 1 | 1 | 1 | 170.3 | 1 | 1571 | 8.03 | | | | | | | | | | | | |
| High | O60921 | Checkpoint protein HUS1 OS=Homo sapiens OX=9606 GN=HUS1 PE=1 SV=1 | 1 | 1 | 1 | 31.7 | 4 | 280 | 6.84 | | | 1 | | | | | | 1 | | | |
| High | Q9Y6H1 | Coiled-coil-helix-coiled-coil-helix domain-containing protein 2 OS=Homo sapiens OX=9606 GN=CHCHD2 PE=1 SV=1 | 1 | 1 | 1 | 15.5 | 7 | 151 | 9.22 | | 1 | | | | | | 1 | | | | |
| High | Q8N6N3 | UPF0690 protein C1orf52 OS=Homo sapiens OX=9606 GN=C1orf52 PE=1 SV=1 | 1 | 1 | 1 | 20.6 | 8 | 182 | 5.21 | | | | | | | | | | | | |
| High | Q16719 | Kynureninase OS=Homo sapiens OX=9606 GN=KYNU PE=1 SV=1 | 1 | 1 | 1 | 52.3 | 6 | 465 | 7.03 | | | | | 1 | | | | | | 1 | |
| High | Q9BYD1 | 39S ribosomal protein L13, mitochondrial OS=Homo sapiens OX=9606 GN=MRPL13 PE=1 SV=1 | 1 | 1 | 1 | 20.7 | 4 | 178 | 9.16 | | | | | 1 | | | | | | 1 | |
| High | Q5K651 | Sterile alpha motif domain-containing protein 9 OS=Homo sapiens OX=9606 GN=SAMD9 PE=1 SV=1 | 1 | 1 | 1 | 184.2 | 0 | 1589 | 7.83 | | | | | 1 | | | | | | 1 | |
| High | O75907 | Diacylglycerol O-acyltransferase 1 OS=Homo sapiens OX=9606 GN=DGAT1 PE=1 SV=2 | 1 | 1 | 1 | 55.2 | 4 | 488 | 9.32 | | | | | | | | | | | | |
| High | P07357 | Complement component C8 alpha chain OS=Homo sapiens OX=9606 GN=C8A PE=1 SV=2 | 1 | 1 | 1 | 65.1 | 5 | 584 | 6.47 | | | | | | | | | | | | |
| High | Q86UV5 | Ubiquitin carboxyl-terminal hydrolase 48 OS=Homo sapiens OX=9606 GN=USP48 PE=1 SV=1 | 1 | 1 | 1 | 119 | 1 | 1035 | 6.05 | | | | | | | | | | | | |
| High | Q9BWH6 | RNA polymerase II-associated protein 1 OS=Homo sapiens OX=9606 GN=RPAP1 PE=1 SV=3 | 1 | 1 | 1 | 152.7 | 1 | 1393 | 6.38 | | | 1 | | | | | | 1 | | | |
| High | Q8IY17 | Patatin-like phospholipase domain-containing protein 6 OS=Homo sapiens OX=9606 GN=PNPLA6 PE=1 SV=3 | 1 | 1 | 1 | 150.9 | 1 | 1375 | 7.74 | 1 | | | | | | 1 | | | | | |
| High | Q8N5V2 | Ephexin-1 OS=Homo sapiens OX=9606 GN=NGEF PE=1 SV=2 | 1 | 1 | 1 | 82.4 | 2 | 710 | 5.57 | | 1 | | | | | | 1 | | | | |
| High | Q96EY9 | Probable inactive tRNA-specific adenosine deaminase-like protein 3 OS=Homo sapiens OX=9606 GN=ADAT3 PE=1 SV=1 | 1 | 1 | 1 | 38 | 7 | 351 | 7.88 | | | | | | | | | | | | |
| High | Q15654 | Thyroid receptor-interacting protein 6 OS=Homo sapiens OX=9606 GN=TRIP6 PE=1 SV=3 | 1 | 1 | 1 | 50.3 | 4 | 476 | 7.37 | | | | | | | | | | | | |
| High | O95985 | DNA topoisomerase 3-beta-1 OS=Homo sapiens OX=9606 GN=TOP3B PE=1 SV=1 | 1 | 1 | 1 | 96.6 | 2 | 862 | 8.12 | | 1 | | | | | | 1 | | | | |
| High | Q15075 | Early endosome antigen 1 OS=Homo sapiens OX=9606 GN=EEA1 PE=1 SV=2 | 1 | 1 | 1 | 162.4 | 1 | 1411 | 5.68 | | | 1 | | | | | | 1 | | | |
| High | Q8IYB7 | DIS3-like exonuclease 2 OS=Homo sapiens OX=9606 GN=DIS3L2 PE=1 SV=4 | 1 | 1 | 1 | 99.2 | 2 | 885 | 6.1 | | | | | | | | | | | | |
| High | O75251 | NADH dehydrogenase [ubiquinone] iron-sulfur protein 7, mitochondrial OS=Homo sapiens OX=9606 GN=NDUFS7 PE=1 SV=3 | 1 | 1 | 1 | 23.5 | 4 | 213 | 9.99 | | 1 | | | | | | 1 | | | | |
| High | Q9BT40 | Inositol polyphosphate 5-phosphatase K OS=Homo sapiens OX=9606 GN=INPP5K PE=1 SV=3 | 1 | 1 | 1 | 51.1 | 2 | 448 | 6.54 | | 1 | | | | | | 1 | | | | |
| High | Q9H7Z7 | Prostaglandin E synthase 2 OS=Homo sapiens OX=9606 GN=PTGES2 PE=1 SV=1 | 1 | 1 | 1 | 41.9 | 3 | 377 | 9.16 | 1 | | | | | | 1 | | | | | |
| High | Q8WX93 | Palladin OS=Homo sapiens OX=9606 GN=PALLD PE=1 SV=3 | 1 | 1 | 1 | 150.5 | 1 | 1383 | 7.09 | | | | 1 | | | | | | 1 | | |
| High | O15525 | Transcription factor MafG OS=Homo sapiens OX=9606 GN=MAFG PE=1 SV=1 | 1 | 1 | 1 | 17.8 | 6 | 162 | 10.04 | | | | | | 1 | | | | | | 1 |
| High | Q96A32 | Myosin regulatory light chain 2, skeletal muscle isoform OS=Homo sapiens OX=9606 GN=MYLPF PE=1 SV=1 | 1 | 1 | 1 | 19 | 8 | 169 | 5.01 | | | | 1 | | | | | | 1 | | |
| High | Q15125 | 3-beta-hydroxysteroid-Delta(8),Delta(7)-isomerase OS=Homo sapiens OX=9606 GN=EBP PE=1 SV=3 | 1 | 1 | 1 | 26.3 | 4 | 230 | 7.9 | | | 1 | | | | | | 1 | | | |
| High | Q9BX40 | Protein LSM14 homolog B OS=Homo sapiens OX=9606 GN=LSM14B PE=1 SV=1 | 1 | 1 | 1 | 42 | 3 | 385 | 9.69 | | | | | | | | | | | | |
| High | Q92544 | Transmembrane 9 superfamily member 4 OS=Homo sapiens OX=9606 GN=TM9SF4 PE=1 SV=2 | 1 | 1 | 1 | 74.5 | 3 | 642 | 6.54 | | 1 | | | | | | 1 | | | | |
| High | Q6ZRI6 | Uncharacterized protein C15orf39 OS=Homo sapiens OX=9606 GN=C15orf39 PE=1 SV=3 | 1 | 1 | 1 | 110.6 | 2 | 1047 | 7.64 | | | | | | | | | | | | |
| High | Q9UM13 | Anaphase-promoting complex subunit 10 OS=Homo sapiens OX=9606 GN=ANAPC10 PE=1 SV=1 | 1 | 1 | 1 | 21.2 | 6 | 185 | 9.06 | 1 | | | | | | 1 | | | | | |
| High | Q9Y2B0 | Protein canopy homolog 2 OS=Homo sapiens OX=9606 GN=CNPY2 PE=1 SV=1 | 1 | 1 | 1 | 20.6 | 8 | 182 | 4.92 | 1 | | | | | | 1 | | | | | |
| High | Q96PE2 | Rho guanine nucleotide exchange factor 17 OS=Homo sapiens OX=9606 GN=ARHGEF17 PE=1 SV=1 | 1 | 1 | 1 | 221.5 | 0 | 2063 | 6.29 | | | | | | | | | | | | |
| High | Q8NC56 | LEM domain-containing protein 2 OS=Homo sapiens OX=9606 GN=LEMD2 PE=1 SV=1 | 1 | 1 | 1 | 56.9 | 3 | 503 | 9 | | 1 | | | | | | 1 | | | | |
| High | Q5XXA6 | Anoctamin-1 OS=Homo sapiens OX=9606 GN=ANO1 PE=1 SV=1 | 1 | 1 | 1 | 114 | 2 | 986 | 8.54 | | | | | | | | | | | | |
| High | P35568 | Insulin receptor substrate 1 OS=Homo sapiens OX=9606 GN=IRS1 PE=1 SV=1 | 1 | 1 | 1 | 131.5 | 1 | 1242 | 8.54 | | | | | | | | | | | | |
| High | Q5JVF3 | PCI domain-containing protein 2 OS=Homo sapiens OX=9606 GN=PCID2 PE=1 SV=2 | 1 | 1 | 1 | 46 | 3 | 399 | 8.53 | | | | | | | | | | | | |
| High | P50135 | Histamine N-methyltransferase OS=Homo sapiens OX=9606 GN=HNMT PE=1 SV=1 | 1 | 1 | 1 | 33.3 | 8 | 292 | 5.34 | | | | | | | | | | | | |
| High | O43681 | ATPase GET3 OS=Homo sapiens OX=9606 GN=GET3 PE=1 SV=2 | 1 | 1 | 1 | 38.8 | 3 | 348 | 4.91 | | | | | | | | | | | | |
| High | Q9BW85 | Splicing factor YJU2 OS=Homo sapiens OX=9606 GN=YJU2 PE=1 SV=1 | 1 | 1 | 1 | 37.1 | 5 | 323 | 5.92 | | | | | | | | | | | | |
| High | Q96QA5 | Gasdermin-A OS=Homo sapiens OX=9606 GN=GSDMA PE=1 SV=4 | 1 | 1 | 1 | 49.3 | 6 | 445 | 5.29 | | | 1 | | | | | | 1 | | | |
| High | Q8N392 | Rho GTPase-activating protein 18 OS=Homo sapiens OX=9606 GN=ARHGAP18 PE=1 SV=3 | 1 | 1 | 1 | 74.9 | 2 | 663 | 6.44 | | | 1 | | | | | | 1 | | | |
| High | Q8TB03 | Uncharacterized protein CXorf38 OS=Homo sapiens OX=9606 GN=CXorf38 PE=1 SV=1 | 1 | 1 | 1 | 36.6 | 6 | 319 | 6.3 | | | 1 | | | | | | 1 | | | |
| High | Q96FX7 | tRNA (adenine(58)-N(1))-methyltransferase catalytic subunit TRMT61A OS=Homo sapiens OX=9606 GN=TRMT61A PE=1 SV=1 | 1 | 1 | 1 | 31.4 | 9 | 289 | 7.36 | | | | | | | | | | | | |
| High | Q6NZI2 | Caveolae-associated protein 1 OS=Homo sapiens OX=9606 GN=CAVIN1 PE=1 SV=1 | 1 | 1 | 1 | 43.5 | 5 | 390 | 5.6 | | | | 1 | | | | | | 1 | | |
| High | O95376 | E3 ubiquitin-protein ligase ARIH2 OS=Homo sapiens OX=9606 GN=ARIH2 PE=1 SV=1 | 1 | 1 | 1 | 57.8 | 3 | 493 | 5.63 | | | | | | | | | | | | |
| High | A5D8V6 | Vacuolar protein sorting-associated protein 37C OS=Homo sapiens OX=9606 GN=VPS37C PE=1 SV=2 | 1 | 1 | 1 | 38.6 | 3 | 355 | 5.21 | | | | | | | | | | | | |
| High | Q96SZ6 | Mitochondrial tRNA methylthiotransferase CDK5RAP1 OS=Homo sapiens OX=9606 GN=CDK5RAP1 PE=1 SV=2 | 1 | 1 | 1 | 67.6 | 3 | 601 | 8.21 | | | | | | | | | | | | |
| High | O94804 | Serine/threonine-protein kinase 10 OS=Homo sapiens OX=9606 GN=STK10 PE=1 SV=1 | 1 | 1 | 1 | 112.1 | 2 | 968 | 6.95 | | | | | | | | | | | | |
| High | P19404 | NADH dehydrogenase [ubiquinone] flavoprotein 2, mitochondrial OS=Homo sapiens OX=9606 GN=NDUFV2 PE=1 SV=2 | 1 | 1 | 1 | 27.4 | 4 | 249 | 8.06 | | | | | | | | | | | | |
| High | Q9Y4W2 | Ribosomal biogenesis protein LAS1L OS=Homo sapiens OX=9606 GN=LAS1L PE=1 SV=2 | 1 | 1 | 1 | 83 | 2 | 734 | 4.73 | | | 1 | | | | | | 1 | | | |
| High | Q9NVH2 | Integrator complex subunit 7 OS=Homo sapiens OX=9606 GN=INTS7 PE=1 SV=1 | 1 | 1 | 1 | 106.8 | 1 | 962 | 8.02 | | | 1 | | | | | | 1 | | | |
| High | Q8IUH5 | Palmitoyltransferase ZDHHC17 OS=Homo sapiens OX=9606 GN=ZDHHC17 PE=1 SV=2 | 1 | 1 | 1 | 72.6 | 3 | 632 | 7.5 | | | 1 | | | | | | 1 | | | |
| High | Q13503 | Mediator of RNA polymerase II transcription subunit 21 OS=Homo sapiens OX=9606 GN=MED21 PE=1 SV=1 | 1 | 1 | 1 | 15.6 | 13 | 144 | 4.45 | | | | | | | | | | | | |
| High | Q13586 | Stromal interaction molecule 1 OS=Homo sapiens OX=9606 GN=STIM1 PE=1 SV=3 | 1 | 1 | 1 | 77.4 | 2 | 685 | 6.67 | | | | | | | | | | | | |
| High | Q9BTE7 | DCN1-like protein 5 OS=Homo sapiens OX=9606 GN=DCUN1D5 PE=1 SV=1 | 1 | 1 | 1 | 27.5 | 5 | 237 | 5.58 | | | | | | | | | | | | |
| High | Q9UPT8 | Zinc finger CCCH domain-containing protein 4 OS=Homo sapiens OX=9606 GN=ZC3H4 PE=1 SV=3 | 1 | 1 | 1 | 140.2 | 2 | 1303 | 6.27 | | | | | | | | | | | | |
| High | Q8N0Z3 | Spindle and centriole-associated protein 1 OS=Homo sapiens OX=9606 GN=SPICE1 PE=1 SV=1 | 1 | 1 | 1 | 96.2 | 2 | 855 | 7.44 | | | | | | | | | | | | |
| High | Q15056 | Eukaryotic translation initiation factor 4H OS=Homo sapiens OX=9606 GN=EIF4H PE=1 SV=5 | 1 | 1 | 1 | 27.4 | 5 | 248 | 7.23 | | | 1 | | | | | | 1 | | | |
| High | Q13277 | Syntaxin-3 OS=Homo sapiens OX=9606 GN=STX3 PE=1 SV=3 | 1 | 1 | 1 | 33.1 | 3 | 289 | 5.44 | | | | | | | | | | | | |
| High | O96008 | Mitochondrial import receptor subunit TOM40 homolog OS=Homo sapiens OX=9606 GN=TOMM40 PE=1 SV=1 | 1 | 1 | 1 | 37.9 | 3 | 361 | 7.25 | | | | | | | | | | | | |
| High | Q14680 | Maternal embryonic leucine zipper kinase OS=Homo sapiens OX=9606 GN=MELK PE=1 SV=3 | 1 | 1 | 1 | 74.6 | 1 | 651 | 8.72 | | | | | 1 | | | | | | 1 | |
| High | Q13952 | Nuclear transcription factor Y subunit gamma OS=Homo sapiens OX=9606 GN=NFYC PE=1 SV=3 | 1 | 1 | 1 | 50.3 | 2 | 458 | 6.1 | | | 1 | | | | | | 1 | | | |
